# A bridge-like lipid transfer protein is critical for generation of invasive stages in malaria parasites

**DOI:** 10.1101/2025.07.25.666630

**Authors:** Andrés Guillén-Samander, Nika Perepelkina, Vendula Horáčková, Hannah M. Behrens, Joëlle Paolo Mesén-Ramirez, Ana Ribeiro-Holbein, Per Haberkant, Frank Stein, Tobias Spielmann

## Abstract

Malaria blood stages build and maintain an intricate system of membranes during their cycle of rapid growth and schizogony (daughter-cell formation), requiring precise mechanisms of lipid synthesis and trafficking. Lipid transfer proteins (LTPs) at ER membrane contact sites (MCSs) have emerged as key for lipid distribution processes but remain largely unexplored in protozoans. Here we use the ER adaptor VAP to identify essential mechanisms of lipid transfer at ER-MCSs in *P. falciparum*. One PfVAP-interacting LTP was the bridge-like PfVPS13L1, which allows bulk flow of lipids between two apposed membranes. PfVPS13L1 bridges the ER with the nascent inner membrane complex (IMC), a *de novo*-generated organelle required for schizogony. Its loss-of-function reduced IMC growth and led to smaller anucleated progeny, impairing schizogony. Our data supports a model in which VPS13L1 is critical for the formation of apicomplexan invasive stages by mediating bulk transfer of lipids from the ER to the growing IMC.

## Introduction

*Plasmodium falciparum* parasites invade, grow and replicate in human red blood cells (RBCs), leading to exponential proliferation of the parasites in the blood and the symptoms of malaria. During their development in RBCs, the parasites establish and expand an intricate system of intra and extra-cellular membranes, which requires precise mechanisms of lipid synthesis and distribution. In eukaryotic cells, lipids are mostly synthesized in the endoplasmic reticulum (ER) and are transported via vesicular trafficking or via lipid transfer proteins (LTPs)^1^. The latter happens primarily in regions of close apposition between organelles, so called membrane contact sites (MCSs), where LTPs are capable of extracting lipids from a bilayer into their hydrophobic cavity and deliver them to the apposed bilayer, thus enabling non-vesicle mediated transfer of lipids between organelles^2,3^. Eukaryotic LTPs can be classified as shuttle-like – proteins with a small hydrophobic cavity that can transfer 1-2 lipid molecules at a time usually engaging in counter transport reactions – or bridge-like – proteins that span the entire distance between donor and acceptor membranes and have a large hydrophobic groove which accommodates many lipids, allowing for bulk transport of lipids to support membrane growth^1,2^.

In recent years, cell biological studies in opisthokonts have revealed the importance of LTPs for various cellular processes involving membrane remodeling, homeostasis, biogenesis and repair^3,4^. Although similarity searches identified many LTPs in *P. falciparum* parasites^5^, only two have been investigated: PfSTART1 and PfNCR1, both with sterol-transferring modules, and essential for the intraerythrocytic stage^6,7^. However, as both are transported beyond the plasma membrane (PM) of the parasite, the relevance of LTPs for intracellular membrane dynamics remains unknown. Importantly, PfSTART1 and PfNCR1 are also the target of novel antimalarials^8,9^, highlighting the importance of studying LTPs for drug discovery. Additionally, MCSs have been observed between several organelles in *Plasmodium* and other apicomplexans^10-12^, but very little is known about potential lipid transferring processes, their relevance for the parasite and the proteins involved.

The ER, as a central hub of cellular lipid distribution, contacts and engages in lipid transferring processes with all other organelles in eukaryotic cells^3,13-16^. *P. falciparum* parasites are not expected to be an exception as MCSs between the ER and other organelles have been observed by different imaging techniques^12,17-19^. In plants and opisthokonts, the ER protein VAP plays an essential role in the formation of ER-MCS by acting as an ER adaptor for proteins that contain a FFAT [two phenylalanines (FF) in an acidic tract (AT)] motif, many of which are LTPs^20-22^. In this study, we used proximity biotinylation to unbiasedly identify LTP partners of the *P. falciparum* ortholog of VAP (PfVAP). We identified a homolog of the bridge-like LTP (BLTP) VPS13, here named PfVPS13L1, that binds PfVAP via its N-terminal end and the inner membrane complex (IMC) through its C-terminal end, indicating localization to ER-IMC MCSs. The IMC is an organelle generated *de novo* during parasite schizogony, the process where up to 36 daughter cells (merozoites) are formed from a multinucleated cell. IMC biogenesis requires large amounts of lipids to wrap around the nuclei and delimit the newly formed merozoites, the stage invading new RBCs. Conditional inactivation of PfVPS13L1 impaired IMC growth and led to the generation of mini merozoite-like structures while a large part of the cytosol of the parent was left behind. Our work supports a model in which direct transfer of lipids in bulk from the ER to the growing IMC by the PfVPS13L1 bridge is essential to support IMC membrane extension and hence formation of functional progeny.

## Results

### PfVAP is an essential ER protein that binds FFAT motifs

Similar to opisthokont VAP, PfVAP (PF3D7_1439800) is a small protein with an N-terminal Major Sperm Protein (MSP) domain (responsible for FFAT motif binding in other organisms), followed by a coiled coil region and a C-terminal transmembrane domain expected to anchor the protein to the ER (Fig. 1a). Using the selection linked integration (SLI) system for genome modification, two parasite lines expressing a version of the protein fused to either Halo or a multipurpose GFP-tag (GFP-2xFKBP) on its N-terminus were generated (Halo-PfVAP^endo^ and N-GFP-2xFKBP-PfVAP^endo^ parasites, Extended Data Fig. 1a-b). Both cell lines showed a bright PfVAP signal in all stages of the asexual intraerythrocytic cycle of the parasite and a clear localization to the ER, as shown by colocalization with the fluorescently tagged TM domain of Sec61β (mSc-TM^Sec61β^) (Fig. 1b and Extended Data Fig. 1c), a protein commonly used as an ER marker in other organisms^18,23^. The ER marker surrounded the nuclei with additional features specific to different stages: in rings and trophozoites tubular extensions were observed as previously described^24^, whereas schizonts contained several ER accumulations that disappeared after segmentation into daughter cells (Fig. 1b). By confocal microscopy, Halo-PfVAP largely colocalized with mSc-TM^Sec61β^, but was enriched in some puncta along the ER (Extended Data Fig. 1d), a feature commonly reported for proteins that are engaged at MCSs^25-27^. More prominent hotspots were observed with GFP-2xFKBP-PfVAP (Extended Data Fig. 1c&e), resembling a phenotype observed in mammalian cells when ER proteins are tagged with GFP due to this tag’s dimerization properties^28^. Despite these artifactual accumulations being seemingly innocuous for the parasites, we opted for using parasites with non-GFP tagged PfVAP for further analysis of localization and interactions.

**Figure 1.**
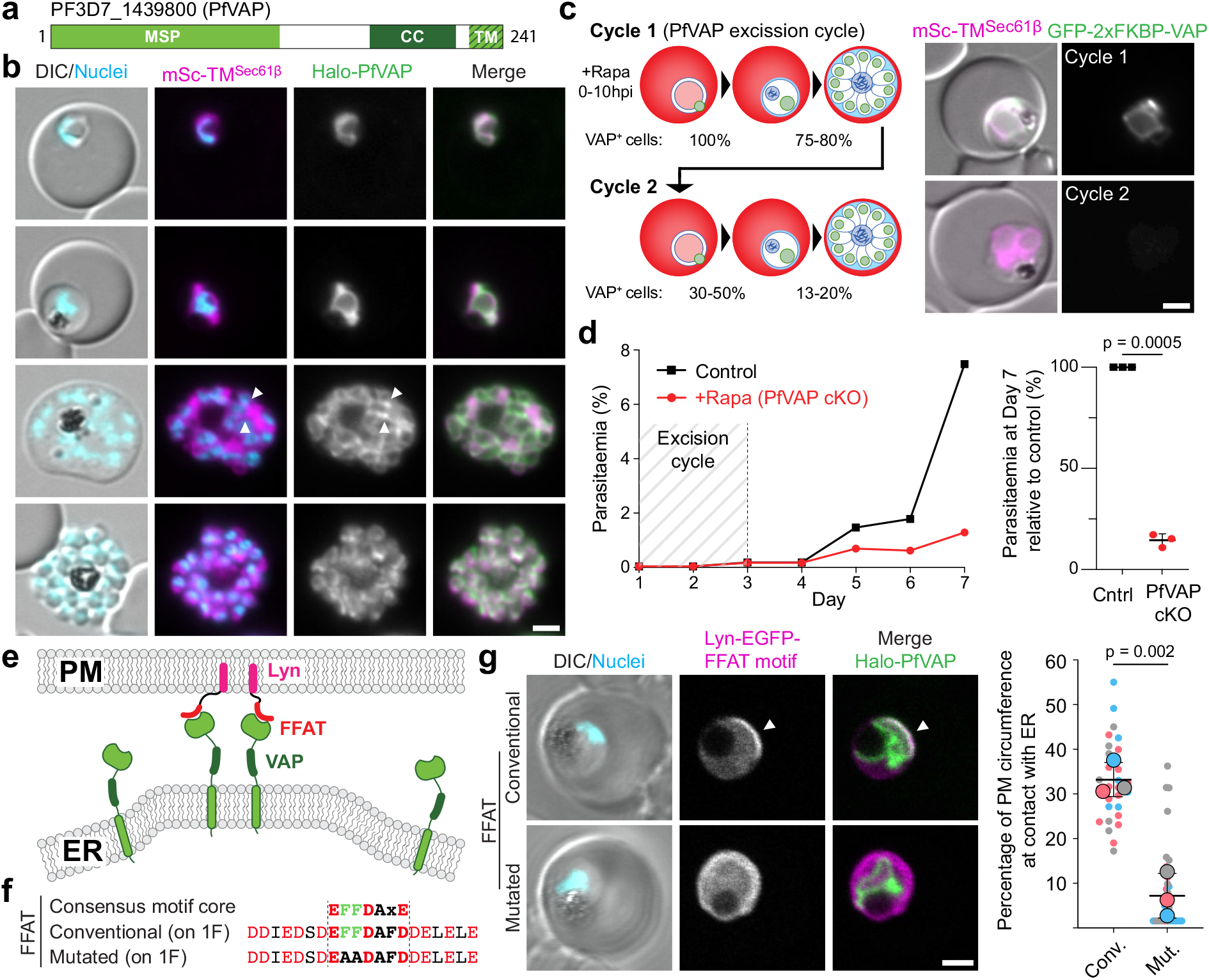
PfVAP is an essential ER protein and binds FFAT motifs. **a**, Domain cartoon of PfVAP. MSP, Major Sperm Protein domain; CC, coiled coil; TM, transmembrane. **b**, Fluorescence microscopy images of Halo-PfVAP^endo^ parasites (Extended Data Fig. 1a-b) episomally co-expressing the TM domain of Sec61β fused to mScarlet (mSc-TM^Sec61β^). Arrowheads, PfVAP enriched in hotspots at ER (also in Extended Data Fig. 1d). **c**, Schematic and quantification of PfVAP loss in GFP-2xFKBP-PfVAP^endo^ parasites episomally co-expressing mSc-TM^Sec61β^ after diCre-mediated gene excision induced using rapalog (+Rapa, PfVAP cKO) [left panel, 3-4 experimental replicates, with 36, 17, 62 and 38 parasites scored for the first timepoint (cycle 1, day 2); 53, 20, 68 parasites for the second timepoint (cycle 2, day 3); 61, 33, 41, 88 parasites for the third timepoint (cycle 2, day 4)] and representative fluorescence microscopy images from both cycles parasites (right panel). **d**, Growth of GFP-2xFKBP-PfVAP^endo^ parasites with rapalog (PfVAPcKO) or without (control), measured by flow cytometry over 7 days (left panel) and growth relative to control (Cntrl) on day 7 in 3 experimental replicates (right panel; bars, average and SD; p-value, two-tailed paired t test). **e**, Schematic of the experimental design to test PfVAP and FFAT motif binding. The motif is anchored to the PM via Lyn, promoting ER-PM MCS formation if bound to PfVAP. **f**, Conventional and mutated FFAT sequences used, based on the opisthokont consensus. **g**, Representative confocal microscopy images of Halo-PfVAP^endo^ parasites episomally expressing the indicated FFAT motif (left panel) and quantification of ER-PM MCSs distance (percentage of total PM circumference, right panel), shown as superplot^102^, from n=3 independent experiments (right panel) with a total of 28 and 29 parasites (1- or 2-nuclei trophozoite stage) quantified for the conventional (Conv.) and mutated (Mut.) motif, respectively; colors indicate independent experiments (small dots, individual parasites; large dots, average of each experiment; black lines, mean and SD; p-value, two-tailed unpaired t test of the means). DIC, differential interference contrast; Nuclei, Hoechst 33342; scale bars, 2 μm.

During genome editing, loxP sites were added at both ends of the *Pfvap* gene (Extended Data Fig. 1a-b) to allow for gene excision with a rapalog-dimerization inducible Cre^29-32^. Growing synchronous GFP-2xFKBP-PfVAP^endo^ parasites with rapalog for a full cycle induced excision of the gene and led to loss of detectable PfVAP in ∼80% of the parasites in the following cycle (Fig. 1c and Extended Data Fig. 1f). This led to a ∼90% reduction in growth over two cycles when compared to control (Fig. 1d), demonstrating the importance of PfVAP for intraerythrocytic cycle progression. The remaining growth may be due to the failure of full PfVAP removal in a percentage of parasites.

As in other organisms VAP acts as an ER receptor for FFAT-motif containing proteins^20^, we tested whether FFAT binding is a conserved function of PfVAP. For this, we designed an assay in which the expression of a FFAT motif anchored to the PM (via a Lyn targeting sequence) would mediate the formation of ER-PM MCSs if it bound PfVAP (Fig. 1e). Indeed, when a conventional FFAT motif^20^ was used in the assay, Halo-PfVAP and the ER in general were recruited to the PM in a section where the Lyn-EGFP-FFAT construct was enriched, but these features were not observed when a FFAT motif with the core phenylalanines mutated to alanines was used (Fig. 1f-g). These results indicate that the FFAT-motif binding capacity of the MSP domain of PfVAP is conserved and, more importantly, PfVAP was confirmed as a good candidate to identify FFAT-motif containing, potential lipid transfer proteins, in *P. falciparum*.

### Identification of VAP-binding proteins

In order to identify VAP interactors in the parasite, we carried out DiQ-BioID^33,34^. For this we generated a cell line endogenously expressing PfVAP fused to two copies of FKBP (Extended Data Fig. 1a-b) and co-expressed the promiscuous biotinilysing enzyme miniTurbo^35^, fused to mCherry and FRB for rapalog-mediated conditional dimerization to recruit it to PfVAP (Fig. 2a). This enables proximity labelling in an induction-dependent manner to subtract background labelling.

**Figure 2.**
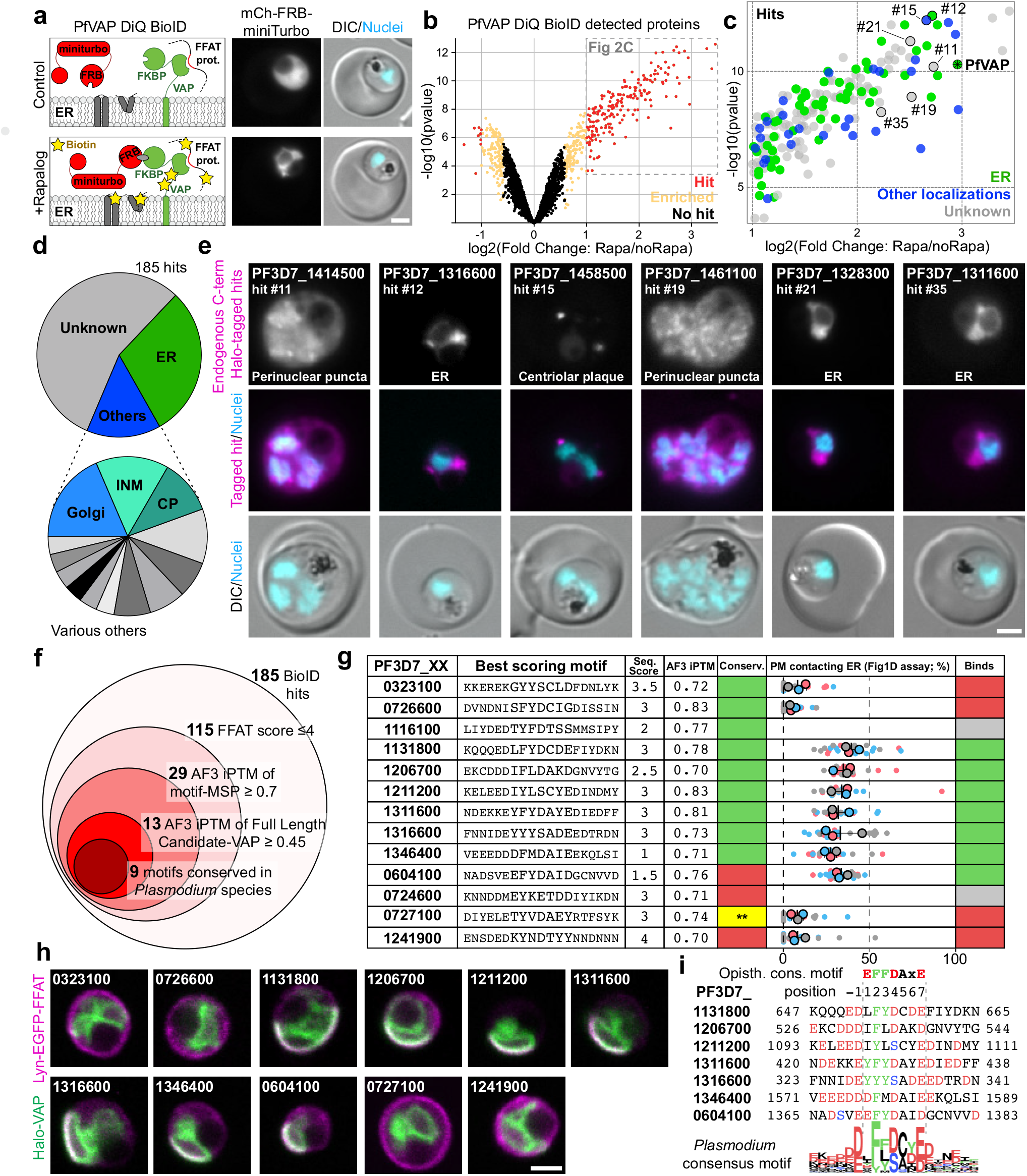
PfVAP DiQ-BioID identifies FFAT-motif containing proteins. **a**, Schematic (left) and representative fluorescence microscopy images (right) showing the rapalog-mediated recruitment of the episomally expressed miniTurbo biotinilyzer to PfVAP for the identification of proximal proteins in 2xFKBP-PfVAP^endo^ parasites. **b**, Volcano plot of the PfVAP DiQ-BioID outlined in (a) showing enrichment of proteins in plus rapalog (miniTurbo recruited to PfVAP) over control (miniTurbo free-floating in cytosol) color coded as indicated. Hits, absolute fold change >2, and a false discovery rate (FDR) <0.05 (185 proteins); enriched, absolute fold change >1.5, FDR <0.2 (252 proteins). Averages from n = 3 independent replicates analysed side by side in the same run (data in Supplementary File 1). Moderated t-test was applied as implemented in the limma package. **c**, Enlargement from (b) showing proteins considered hits color coded according to localization. Numbers indicate proteins analysed in (e). **d**, Pie charts showing proportion of the hits in (c) with the indicated localization. Annotation of localization according to Supplementary File 1B. INM: Inner Nuclear Membrane; CP: Centriolar Plaque. **e**, Representative fluorescence microscopy images of parasites in which the indicated protein (each corresponding to a hit numbered in (c) were endogenously tagged with Halo-SW in their C-terminus. Classification of localization indicated. **f**, Schematic of hit filtering based on FFAT motifs starting with the 185 hits of the PfVAP DiQ-BioID (a-c) as indicated in the materials and methods and Supplementary File 3. Scoring cut-offs are indicated; iPTM, interface predicted template modelling. **g**, Table summarizing the 13 highest scoring FFAT motifs from (f) (Seq. Score, FFAT score based on Slee, J.A. & Levine, T.P.^40^; AF3 iPTM, MSP domain PVAP AlphaFold3 iPTM value^41^) and experimental data shown in (h) quantified and displayed as a superplot^102^ as in Fig. 1g [3 independent replicates and a total of at least 20 parasites (1- or 2-nuclei trophozoite stage); binding was considered when there was no significant difference (p-value >0.05) with the positive control (Fig. 1g) and indicated in green (grey, not tested; red, no binding), two-tailed unpaired t test of the means (p-values: PF3D7_0321300, 0.0026; PF3D7_0726600, 0.0003; PF3D7_1131800, 0.0917; PF3D7_1206700, 0.5591; PF3D7_1211200, 0.727; PF3D7_1311600, 0.8825; PF3D7_1316600, 0.9431; PF3D7_1346400, 0.1544; PF3D7_0604100, 0.7238; PF3D7_0727100, 0.0012; PF3D7_1241900, 0.0015)]. Conservation of motifs (Conserv.) is indicated in green (no conservation in red); ** (yellow), motif that is conserved but part of a folded region, hence unlikely to bind PfVAP. (Fig. 2 continued) **h**, Representative confocal microscopy example images of Halo-PfVAP^endo^ parasites episomally expressing the FFAT motifs indicated and used to quantify the resulting ER-PM MCSs distance as in Fig. 1g and summarized in (g). **i**, Alignment of the 7 binding motifs identified, displayed respective to the opisthokont consensus and a sequence logo of the Plasmodium consensus built from their conserved homologous sequences in 10 different Plasmodium species. Amino acids are colored according to their properties: green, aromatic; red, acidic; blue, phosphorylatable. Note difference to consensus opisthokont motif in positions 1 and -1. DIC, differential interference contrast; Nuclei, Hoechst 33342; scale bars, 2 μm.

Quantitative MS analysis of biotinylated proteins on target (+rapalog) over control revealed a total of 185 significant hits in proximity to PfVAP (Fig. 2b-c, Supplementary File 1). Given the high expression of this protein and its localization throughout the ER in all stages of the parasite, the detection of many hits is expected as this system identifies all proteins in proximity and not only direct interactors of PfVAP. Upon classification based on previous literature and protein function, the majority of proteins with a known or expected localization turned out to be ER proteins (Fig. 2c and Extended Data Fig. 2a-b), followed by other ER proximal compartments, such as the inner nuclear membrane (INM; the same compartment as the ER but facing the nucleoplasm, where VAP is also found in other eukaryotes^36^), the nuclear envelope embedded centriolar plaque (CP)^37^, or the Golgi apparatus. Additionally, there was an overrepresentation of transmembrane proteins, expected due to the recruitment of the biotinyliser to the ER membrane (Fig. 2d and Extended Data Fig. 2c-f).

We modified the endogenous loci of 6 hits by adding a C-terminal Halo-SW tag (Halo-sandwich, corresponding to Halo flanked on each side by 2xFKBP) (Extended Data Fig. 2g), 4 of which were proteins of unknown localization and 2 with expected localizations^38,39^. Three of them showed a typical ER pattern, and three showed puncta in the surroundings of the nuclei (Fig. 2e), one of which was at the CP as shown by being adjacent to the inner CP marked by tubulin (Extended Data Fig. 2h), in agreement with a previous study in *P. berghei* ^39^. All the localizations were compatible with a proximity to the ER and, hence, to PfVAP.

Additionally, we observed an overrepresentation of high-scoring FFAT motifs in the DiQ-BioID hits, likely indicating the identification of direct PfVAP binders (Extended Data Fig. 2i-j, Supplementary File 2&3). We filtered the list of 185 hits based on their highest scoring FFAT motif sequences^40^ and predicted AlphaFold3^41^ interaction to identify 13 proteins containing motifs with a high likelihood of PfVAP binding, 9 of which were conserved in *Plasmodium spp* (Fig. 2f-g, Supplementary File 3).

To validate this strategy to find PfVAP binders, we used the above described PfVAP-binding assay (Fig. 1e-f) with the selected FFAT motifs. Out of 11 tested motifs, 7 resulted in the artificial formation of ER-PM MCSs (Fig. 2g-h), indicative of binding to PfVAP and a direct interaction of the corresponding full-length candidates with PfVAP. Alignment of the interacting motifs and their orthologous sequences in 10 different *Plasmodium* species revealed some differences with the opisthokont consensus motif (Fig. 2i): a lack of importance of the amino acid in position 1 of the *Plasmodium* motif, which in opisthokonts is usually a negatively charged residue, and instead the presence of a negatively charged residue in position -1 of the *Plasmodium* motif. Taking this into account, we propose a *Plasmodium* specific FFAT scoring system, which we used to predict motifs across the *Plasmodium* genome and which showed an even higher enrichment of FFAT motifs in the PfVAP DiQ-BioID hit list, lending support to the *Plasmodium* specific consensus and further validating the hit list (Extended Data Fig. 2k-l and Supplementary File 2). Of the 7 proteins here confirmed to have a PfVAP-interacting motif, 2 contained domains characteristic of LTPs (PF3D7_1131800 and PF3D7_1346400), prompting us to study them in more depth.

### Lipid transfer proteins identified in proximity to the ER

Besides the 2 FFAT-motif containing LTPs, 5 other proteins with domains typical for LTPs, resulting in 7 ER proximal LTPs, were identified in our PfVAP DiQ-BioID (Fig. 3a). Of these, 3 were shuttle-like and 4 bridge-like LTPs (Fig. 3b-d). The shuttle-like proteins all had lipid transfer domains belonging to different families: a VAD1 Analog of a StART (VASt) domain, a Phosphatidylinositol Transfer Protein (PITP) domain and an OSBP-Related Domain (ORD)^1,42^ (Extended Data Fig. 3a-b). We named them, accordingly, PfGRAMD1 (PF3D7_0803600) based on its homology to the human GRAMD1 proteins^43^, PfPITP (PF3D7_1351000), which also has a second lipid transfer domain in its C-terminus with remote similarity to an ORD, and PfOSBP (PF3D7_1131800, containing a FFAT motif in Fig. 2g-i).

**Figure 3.**
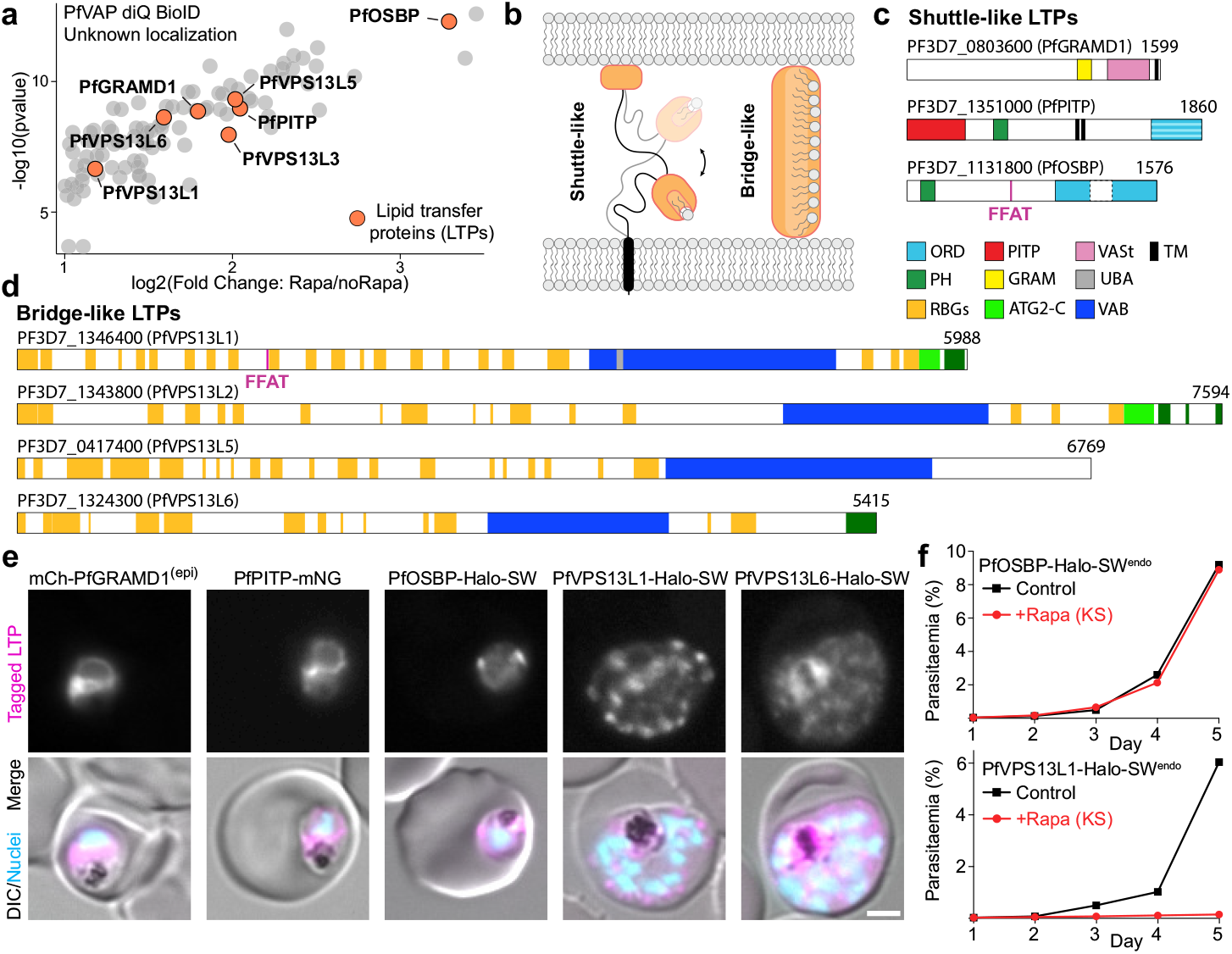
Lipid transfer proteins (LTPs) in proximity to PfVAP. **a**, 7 LTPs identified in the PfVAP diQ-BioID hits (plot as in Fig. 1c). **b**, General sketch of shuttle- or bridge-like LTPs. **c-d**, To scale representation of the 3 shuttle-like (c) and 4 bridge-kike LTPs (d; all remote homologs of opisthokont VPS13) highlighted in (a) showing the domains identified in the AF3 predicted structures. TM, transmembrane domain. Numbers, total amino acid sequence length. In d, β-strands presumably forming the lipid transfer groove are in orange, even if a full RBG domain was not observed (more detailed RBG domains annotation of PfVPS13L1 in Fig. 4a and Extended Data Fig 5a). **e**, Representative fluorescence microscopy images of parasites episomally (epi) (mCh-PfGRAMD1) or endogenously expressing the indicated tagged LTPs. **f**, Growth of parasites after knock-sideways (KS, rapalog) of the indicated PfVAP binding LTPs compared to control, measured by flow cytometry (n = 3 (PfVPS13L1) or 4 (PfOSBP) independent experiments; Comparison of day 5 parasitemia for all replicates in Extended Data 4b). DIC, differential interference contrast; Nuclei, Hoechst 33342; scale bars, 2 μm.

We targeted the genes encoding the shuttle-like LTPs for endogenous tagging and obtained genome edited lines for PfPITP and PfOSBP, but not for PfGRAMD1, which we expressed episomally (Extended Data Fig. 3c). mCh-PfGRAMD1 and PfPITP-mNG localized to the ER, likely via their TM domains (Fig. 3e). Interestingly, both had other domains that can serve as lipid-binding adapters (PH and GRAM, Fig. 3c and Extended Data Fig. 3b), which could target the proteins to a specific ER-MCS in a regulated manner. PfOSBP-Halo-SW was mostly found in puncta proximal to the nucleus in trophozoites, colocalizing with the ER and some-times overlapping with or adjacent to mNG-Rab6, a marker of the Golgi apparatus (Fig. 3e and Extended Data Fig. 3d-g). Interestingly, this localization was dynamic, as in some cells PfOSBP-Halo-SW was also found at the PM, while in other cells it was exclusively at the ER (Extended Data Fig. 3d-f) and this distribution changed throughout the parasite cycle (Extended Data Fig. 3e). Localization to the ER is likely mediated by its FFAT motif binding to PfVAP, and its additional PH domain could direct the protein to a specific ER-MCSs in a regulated manner (Extended Data Fig. 3b). Accordingly, episomal expression of a tandem construct of its PH domain showed an enrichment at the Golgi, as seen by colocalization with a tagged adaptor protein complex subunit AP-1μ(Extended Data Fig. 3h). In humans, the PH domain of OSBP binds Phosphatidylinositol 4-phosphate (PI4P) through a surface that is conserved in the PH domain of PfOSBP^44,45^ (Extended Data Fig. 3i). Similarly to other eukaryotes, in *P. falciparum* parasites PI4P is enriched at the Golgi, and to a lesser extent at the PM^46^. It is therefore likely that PfOSBP is recruited dynamically to ER-Golgi MCSs or ER-PM MCSs depending on PI4P levels in these membranes.

The four bridge-like LTPs (BLTPs) found were homologous to the opisthokont VPS13 proteins (Fig. 3d). Two of them had sequence homology in the characteristic N-terminal chorein domain, but the other two had no identifiable sequence similarity and were found by using AlphaFold3^41^ to predict the structures of all the large (above 2000 aa) and unknown proteins in the list of VAP DiQ-BioID hits. Further homology searches showed that there is a total of six VPS13-like proteins in *P. falciparum*, which we considered for naming them PfVPS13L1-6 (Extended Data Fig. 4a). We here focused on the VPS13-like proteins in the VAP hit list (PfVPS13L1, 2, 5 and 6). Unfortunately, we could not gather data for PfVPS13L5 as the SLI-mediated tagging did not yield any parasites. PfVPS13L3-Halo-SW signal was undetectable and PfVPS13L6-Halo-SW showed a very low fluorescence signal suggesting a cytosolic localization with accumulations (Fig. 3e and Extended Data Fig. 4b). Parasites where PfVPS13L1 was tagged with a Halo-SW (from here on referred to as PfVPS13L1-Halo-SW^endo^, and PfVPS13L1-GFP-SW^endo^ for the GFP version) also gave a low signal but in early schizont stages, when its expression peaked, a punctate localization proximal to the nuclei was clearly discernable (Fig. 3e and Extended Data Fig. 4b). Since PfVPS13L1 has a functional FFAT motif (Fig. 2g-i), it is expected to be bound to PfVAP, and the punctate pattern could reflect its enrichment at an ER-MCSs with another organelle.

We then determined the essentiality of the two FFAT-motif containing LTPs, PfOSBP and PfVPS13L1, that localized in puncta that may represent ER-MCSs. For this we episomally expressed a Lyn-fused FRB construct in the respective lines. As the SW constructs have 4 copies of FKBP, addition of rapalog promotes the recruitment of the LTPs to the PM, allowing us to study their loss-of-function by mislocalization (Knock-sideways, KS^32,47,48^). Mislocalization of the target protein to the PM was efficient in both cell lines (Extended Data Fig. 4d-e). This had no effect on parasite growth in the case of PfOSBP (Fig. 3f and Extended Data Fig. 4c-d), in accordance with a previous study where a targeted gene disruption was achieved^49^. In contrast, removal of PfVPS13L1 was lethal for the parasites (Fig. 3f and Extended Data Fig. 4c&e), which prompted us to focus our further investigation on this BLTP’s function. Knock-sideways was also done for the non-FFAT containing PfVPS13L6, but this protein turned out to be dispensable (Extended Data Fig. 4c&f).

### PfVPS13L1 bridges the ER and the IMC

Similar to other VPS13 proteins^4^, PfVPS13L1 is a very large protein (5899 residues) predicted by AlphaFold3^41^ to fold as a rod formed by a twisted β-sheet which features three folded domains on its C-terminal end: 1) a VPS13 adaptor binding (VAB) domain that sticks out of the rod and is formed by six β-sheet repeats, 2) an ATG2-C, formed by amphipathic helices, and 3) a PH domain; the latter two domains arranged in tandem at the C-terminal end of the rod. In addition, PfVPS13L1 is predicted to have a UBA domain on the N-terminal end of the VAB domain, similar to metazoan VPS13D proteins^50,51^ (Fig. 4a-c, Extended Data Fig. 5a-e and Movie S1). The ∼22 nm rod is formed by a series of 12 repeating β-groove (RBG) domains, a repeating unit of 5 antiparallel β-strands and 1 α-helix (except for the first and last RBG domains which have fewer β-strands) (Extended Data Fig. 5a-b), which gives the name to the superfamily VPS13 belongs to^52,53^. Although the size of its predicted 3D structure is similar to that of yeast Vps13 or human VPS13A (both also with 12 RBG domains), PfVPS13L1 comprises around twice the number of amino acids^4^, largely due to flexible stretches protruding from the folded core (compare Fig. 4b and c). This, in addition to the further divergence of the other VPS13L protein sequences, hampered the proper prediction of their full structures.

**Figure 4.**
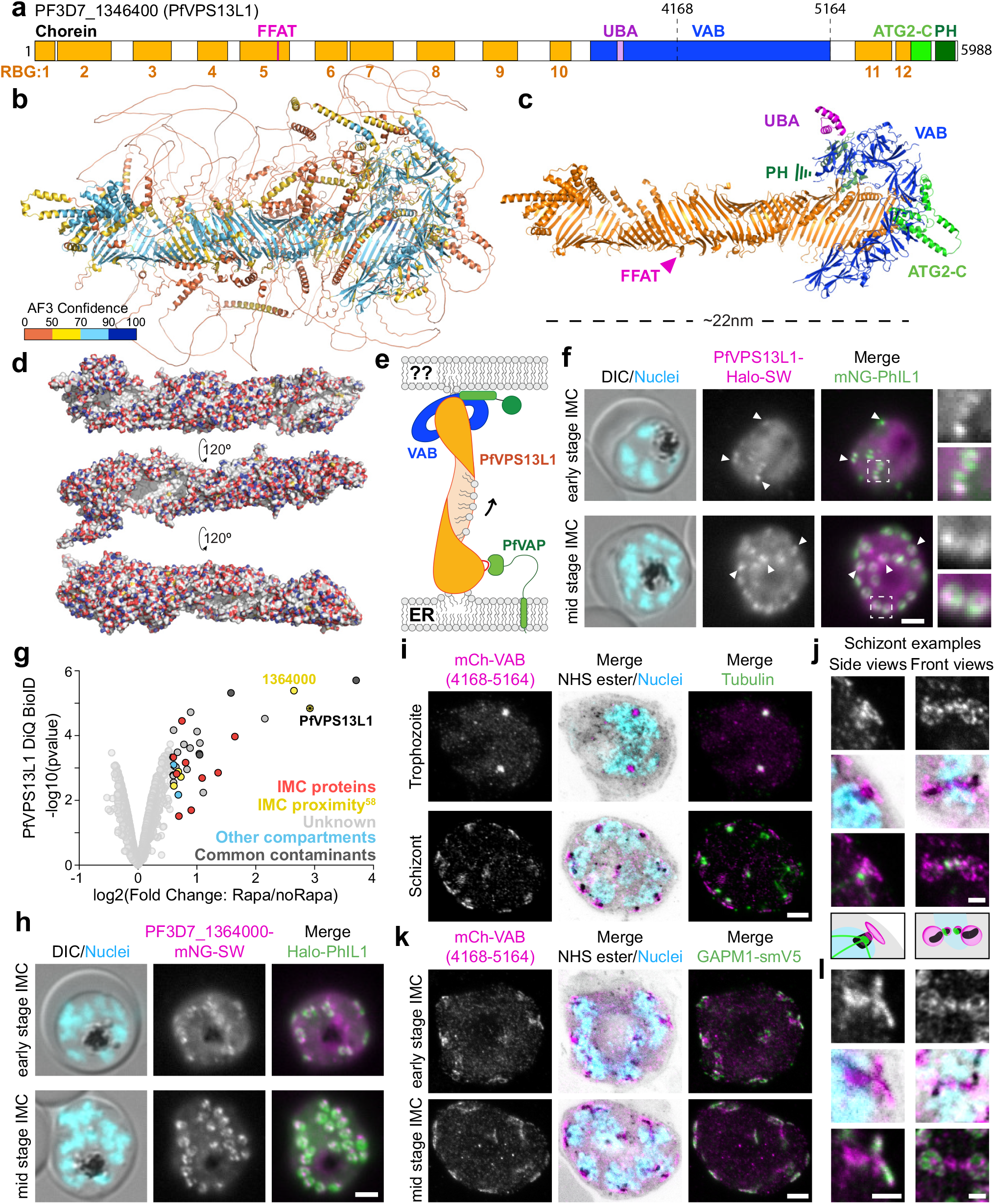
PfVPS13L1 bridges the ER to the nascent IMC. **a**, Domain cartoon of PfVPS13L1, indicating the RBG domains that compose the lipid transferring bridge, the PfVAP binding FFAT motif and the C-terminal adaptor and membrane binding domains. Black numbers, amino acids; orange numbers, RBG domains. **b**, Ribbon representation of AlphaFold3 predicted structure of PfVPS13L1 colored by prediction confidence. **c**, Structure from (b) with flexible regions not part of annotated folded domains removed for clarity, and colored by domain annotation. **d**, Surface representation of the rod composed by the 12 RBG domains from 3 angles colored by element (oxygen, red; nitrogen, blue; carbon, white), highlighting the internal white groove, indicating its hydrophobicity. **e**, Schematic of PfVPS13L1 expected recruitment to an ER-MCSs based on its N-terminal FFAT motif binding to the ER via PfVAP. **f**, Representative fluorescence microscopy images of parasites expressing PfVPS13L1-Halo-SW from the endogenous locus (Extended Data Fig. 4b) and episomally expressing the IMC marker PhIL1 fused to mNeonGreen (mNG-PhIL1). Arrowheads, overlap in early- and mid-stage of IMC formation. Boxed areas are enlarged 2.5x. **g**, Volcano plot of the DiQ-BioID of the C-terminus of PfVPS13L1 showing enrichment (3 independent samples per condition; second experimental repeat and more detail in Extended Data Fig. 7c) of proteins in plus rapalog (miniTurbo recruited to PfVPS13L1 C-terminus) over control (miniTurbo free-floating in cytosol) as outlined in Extended Data Fig. 7a. Enriched proteins (absolute fold change >1.5, p-value <0.05) are highlighted by a black stroke and color coded as indicated (data in Supplementary File 4). Moderated t-test was applied as implemented in the limma package. IMC proximity based on previous Phil1 BioID^58^; common contaminants based on previous DiQ-BioIDs^33^. **h**, Representative fluorescence microscopy images of parasites with endogenously mNG-SW-tagged PF3D7_1364000 and episomally expressing Halo-PhIL1 in early- and mid-stages of IMC formation. (Fig. 4 continued) **i-l**, Ultra Expansion Microscopy (U-ExM) images of parasites expressing a construct containing 5 of the 6 repeats of the C-terminal VAB domain of PfVPS13L1 (see Extended Data Fig. 5c) fused to mCherry (mCh-VAB) co-immunostained for α-tubulin (i, j) or episomally co-expressing GAPM1-smV5 (k-l). mCh-VAB colocalized with the centriolar plaque (marked by tubulin and NHS density) in trophozoites but formed a ring-like structure adjacent to the outer centriolar plaque in schizonts (i, examples enlarged in j) that colocalized with GAPM1-smV5 (k, enlarged in l). The enlargements show examples where the ring was captured from side and front perspectives, with a schematic below (j), color coded as (j), simplifying the observation. U-ExM images are maximum intensity projections of Z-slices, with 10 (i, top), 8 (i, bottom), 12 (k, top), 6 (k, bottom) and 3 (j&l) slices. DIC, differential interference contrast; Nuclei, Hoechst 33342; scale bars, 2 μm in (f, h, j & l); 5 μm in (i & k).

The RBG domains have a hydrophobic surface facing the internal side of the twisted β-sheet that forms the rod (Fig. 4d and Extended Data Fig. 5b), which in BLTPs of other organisms has been shown to be able to accommodate the hydrophobic tails of several lipids^54,56^. When the rod is located between two organellar membranes, lipids would be able to flow in bulk from one organelle to the other. The FFAT motif of PfVPS13L1 is found in a flexible stretch on the fifth RBG domain, localized closer to the N-terminal end of the rod (Fig. 4c and Extended Data Fig. 5f), suggesting that this end is bound to the ER (Fig. 4e). Interestingly, 3 out of the 4 human VPS13 proteins have an FFAT motif found in a similar region (right before the 5th RBG domain), which also leads to the attachment of their N-terminal end to the ER, considered the lipid donor membrane^4^. On the C-terminal end of the protein, the specific interactions of the adaptor domains (VAB, PH domains) with protein or lipid partners usually dictate the identity of the receiving membrane (Fig. 4e). The amphipathic helices of the ATG2-C C-terminal domain are thought to partially insert themselves into the bilayer, potentially facilitating the lipid delivery process^57^.

VPS13 proteins are expected to mediate bulk flow of lipids and are essential for the *de novo* organelle biogenesis in other organisms^4^. In *P. falciparum*, the inner membrane complex (IMC) is generated *de novo* around the nuclei of schizonts, the stage where we observed the peak of PfVPS13L1 expression (Extended Data Fig. 6a). A previous BioID detecting IMC proteins in *P. falciparum* contained PfVPS13L1 as a top candidate^58^, prompting us to assess its localization in relation to the IMC marker mNG-PhIL1^59^. PfVPS13L1-Halo-SW showed a colocalization at early and mid stages of IMC formation (Fig. 4f and Extended Data Fig. 6a-c). Specifically, PfVPS13L1-Halo-SW was enriched in hotspots in a subsection of the IMC, which by confocal microscopy sometimes appeared to be on the basal side of the nascent IMC (Extended Data Fig. 6b) and were also directly colocalizing or adjacent to the ER (Extended Data Fig. 6d-e), compatible with the idea that PfVPS13L1 was present at the MCS between both organelles. To confirm that this connection was mediated through its C-terminal end, we performed DiQ-BioID to identify proteins in proximity to this end of PfVPS13L1 (Extended Data Fig. 7a-b, Supplementary File 4). Indeed, our DiQ-BioID results showed an enrichment of bona fide IMC proteins and proteins expected to be in the IMC (based on previously published proximity biotinylation experiments^58^) (Fig. 4g and Extended Data Fig. 7c-d). One protein, PF3D7_1364000, was consistently the highest enriched hit, suggesting a potential interaction. Genome modified parasites expressing PF3D7_-1364000 with a mNG-SW tag revealed that this protein localizes to the IMC, most prominently in a subsection (Fig 4h and Extended Data Fig. 7e-f), similar, albeit brighter, to what was observed for PfVPS13L1. Taken together, these results confirm that the C-terminal end of the PfVPS13L1 bridge is proximal to the IMC membrane.

As the C-terminal adaptor domains of VPS13 proteins are expected to determine the organellar specificity, we sought to confirm the IMC interaction by episomally expressing constructs containing these domains of PfVPS13L1 (Extended Data Fig. 8a). The construct encoding two copies of the PH domain showed a cytosolic signal (Extended Data Fig. 8b), but the one containing the tandem of ATG2-C and PH domains showed an enrichment in small-rounded compartments also observable by DIC in late RBC stages, likely corresponding to lipid droplets (LDs) (Extended Data Fig. 8c). This is not surprising, as the amphipathic helices in the ATG2-C domain target LDs in other VPS13 proteins such as in human VPS13A and VPS13C^26^, but this is not expected to be the main site of action for these proteins as their role in LD homeostasis is minimal^60^. The construct containing the VAB domain displayed a clear punctate localization adjacent to the inner CP indicated by a tubulin marker in trophozoite and schizont stages, and dispersed at the end of parasite segmentation (Extended Data Fig. 8d), similar to what is known to occur with the outer CP^61^. Ultra-expansion microscopy (U-ExM) revealed that while this signal colocalized with the CP in trophozoites, it acquired a ring-like morphology in schizonts that was adjacent to the CP, on its apical side (Fig. 4i-j), where the nascent IMC generates from^62^. Accordingly, this signal appeared to localize between the CP and the Golgi, partially colocalizing with an IMC marker in confocal microscopy (Extended Data Fig. 8e), and by U-ExM the VAB ring was observed colocalizing with the early stages of the IMC, as marked by the IMC protein GAPM1 (Fig. 4k-l). A recent publication reported the existence of a ring-like IMC initiation scaffold near the apical side of developing merozoites, adjacent to the CP, that disperses in late schizont stages^63^, which would be in accordance with what we observe for the VAB domain of PfVPS13L1. Taken together, our localization data show that the conserved FFAT domain of PfVPS13L1 binds the ER (Fig. 2g-h) while its VAB domain on the other end of the protein binds the IMC, likely at the initiation scaffold of this organelle (Fig. 4f-l). This indicates that PfVPS13L1 forms a lipid transferring bridge at MCSs between the ER around the nucleus and the growing IMC.

### PfVPS13L1 is essential for IMC formation and proper segmentation

To understand the function of PfVPS3L1, we first assessed in what stage of the intraerythrocytic cycle it was essential by inducing its mislocalization with rapalog in different stages. Despite a small percentage of parasites dying in the trophozoite stage after mislocalization of PfVPS13L1, the largest defect was observed in later stages which failed to produce new progeny, evident from an absence of new rings in the next cycle (Fig. 5a and Extended Data Fig. 9a). To confirm this observation, we induced the loss of PfVPS13L1 function around 36hpi (i.e. right before the schizont stage and hence IMC formation) and evaluated how many new rings the schizonts were able to produce in the next cycle. Almost no new parasites were observed after mislocalization of PfVPS13L1, indicating the essentiality of PfVPS13L1 for late intraerythrocytic stages, which could include segmentation, egress or invasion of new RBC (Fig. 5b-c).

**Figure 5.**
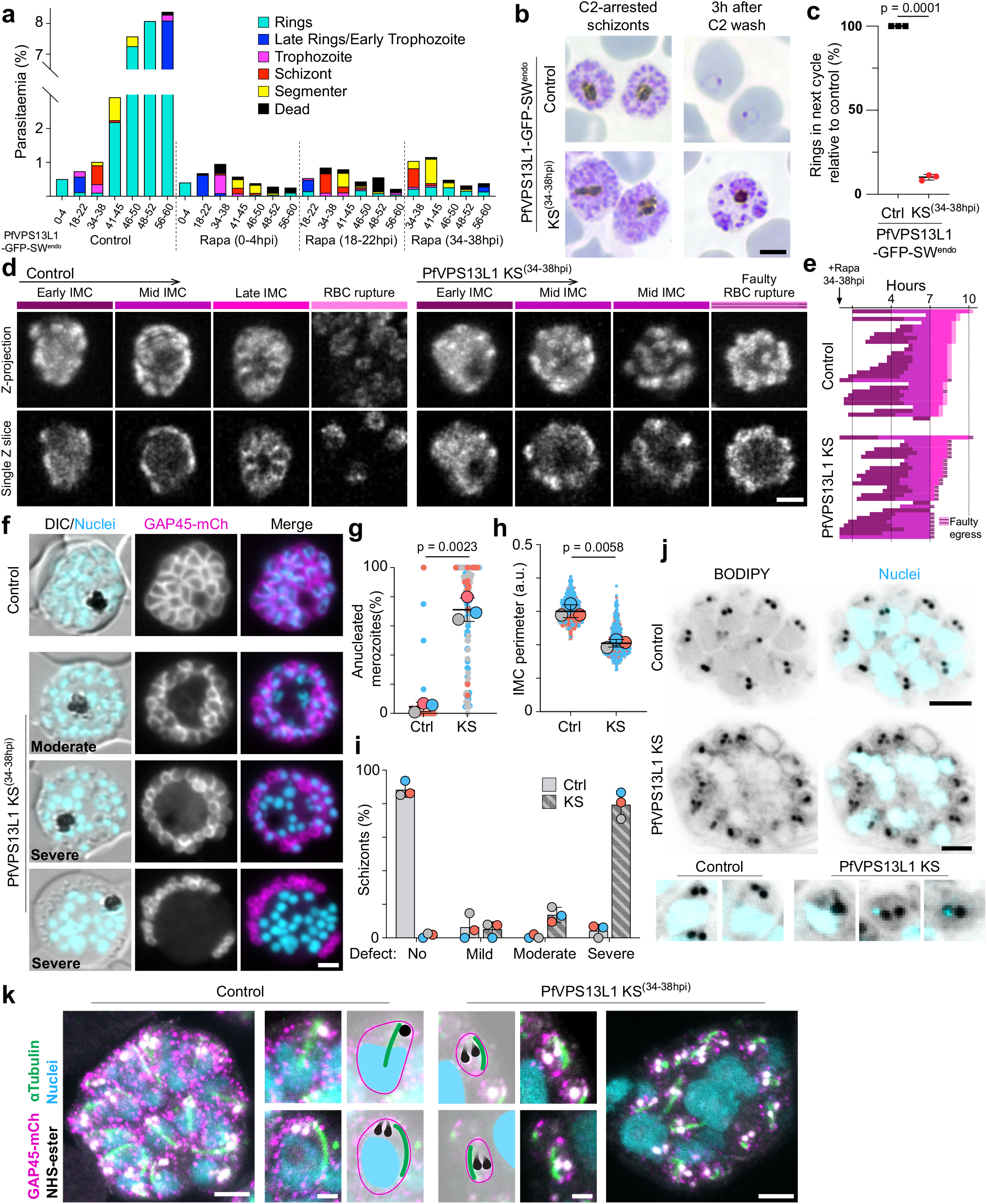
PfVPS13L1 is essential for proper IMC formation and schizont segmentation. **a**, Parasite growth and stage progression assay in synchronous parasites (4 h stage window) upon PfVPS13L1 KS (using the PfVPS13L1-GFP-SW^endo^ parasites (Extended data Fig. 4b) episomally expressing the Lyn mislocaliser) induced at the indicated time points based on Giemsa smears (representative images are shown in Extended Data Fig. 9a). **b-c**, Representative example Giemsa smears (b) and quantification (c) of rings produced after PfVPS13L1 KS (rapalog added at 34-38hpi) in comparison to no rapalog controls. Compound 2 was used to arrest segmenters in both conditions and rings assessed after wash out (n = 3 independent experiments; black lines, mean with SD; p-value, paired t test). **d**, Representative maximum intensity projections and single Z-slices of selected time points of 3D-timelapse imaging of PfVPS13L1-GFP-SW^endo^ parasites episomally expressing Halo-PhIL1 (IMC) in control and PfVPS13L1 KS (+rapa) schizonts. PfVPS13L1 mislocalization triggered at 34-38hpi schizonts, imaging (20 min interval) started at 36-40hpi schizonts. **e**, Quantification of time lapse experiment in (d). IMC development stages [indicated in (d)] were classified in each imaged schizont (n=29 and 27 cells, in control and KS, respectively, from 2 independent experiments) and progression represented graphically. Faulty egress: RBC rupture with merozoites not egressing or remaining attached to a large residual body. **f**, Fluorescence microscopy images of control and late (34-38hpi)-induced PfVPS13L1 KS schizonts arrested before egress with compound 2. **g**, Superplots^102^ showing percentage of anucleated merozoites (defined as compartments delimited by the IMC marker without a nucleus) per schizont after KS of PfVPS13L1 compared to control (Ctrl). Three independent replicates with a total of 87 and 103 cells for control and KS, respectively; colors, independent experiments; individual parasites, small dots; average per experiment, large dot; black lines, mean and SD; p-values, two-tailed paired t test of the means). (Fig. 5 continued) **h**, Superplot as in (g) but measuring IMC perimeter. a.u., arbitrary units; p-value, two-tailed paired t test of the means. All visible IMC compartments were measured in 15, 16, 16 control cells and 16, 19, 16 KS cells across three independent experiments. **i**, Quantification of the phenotypes observed in (f) classified as mild (1-2 nuclei with no associated IMC), moderate (up to 6 nuclei with no associated IMC) or severe (at least half of the nuclei without associated IMC). Bar graphs, percentage of each phenotype scored, (3 independent experiments, each represented by one color of dot); p-values (two-tailed paired t test of the means): 0.0016, 0.7134, 0.0055 and 0.0014 for no, mild, moderate and severe phenotype, respectively. **j-k**, U-ExM images of control and late-induced KS schizonts stained with Bodipy (j) or NHS-ester, anti-mCh (for GAP45-mCh) and anti-αTubulin (k) showing accumulated nuclei in the residual body and anucleated merozoites upon PfVPS13L1 mislocalization (PfVPS13L1 KS). Some of the smaller merozoites carried a small amount of DNA (I, bottom panel). Enlargements show individual merozoites taken from different slices, including images with artificial graphical highlight of key features (rhoptries, black; IMC, magenta; Tubulin, green; nuclei, cyan). U-ExM images are maximum intensity projections of Z-slices, with 6 (j; Control and KS), 4 (j; enlargements), 13 (k; Control), 11 (k; KS) and 6 (k; enlargements) slices. DIC, differential interference contrast; Nuclei, Hoechst 33342; scale bars, 2 μm in [d, f & k (enlargements)] and 5 μm in [b, j & k (whole cell)].

As we had determined the localization of PfVPS13L1 to be at ER-IMC MCSs, the most parsimonious explanation would be that this BLTP is transferring lipids in bulk to support the *de novo* IMC formation. We evaluated this hypothesis by monitoring IMC growth upon loss of function induction in late stages. Live-cell confocal microscopy showed that, although an initial IMC seed was observed in all cases, the further growth of this membrane was severely reduced when PfVPS13L1 was not functional, leading to a faulty egress upon RBC rupture that left behind a large residual body (Fig. 5d-e and Movie S2).

To better understand the morphological defects in the schizonts where PfVPS13L1 was mislocalized, we arrested the parasites shortly before egress by using Compound 2^64^. While in the control cells the IMC membranes formed and distributed all around the daughter parasites, always surrounding a nucleus, severe defects were observed upon PfVPS13L1-Halo-SW mislocalization: the IMC appeared smaller and disconnected from the nuclei, with abundant DNA material and parasite cytoplasm being leftover in an atypically large residual body (Fig. 5f-g and Extended Data Fig. 9b). Segmentation still occurred but resulted in the formation of often anucleated small merozoite-like structures (Fig. 5f-h and Extended Data Fig. 9b-e). These defects ranged in severity, probably depending on the exact stage each individual schizont was in when PfVPS13L1 mislocalization was induced (Fig. 5f-i). U-ExM confirmed these phenotypes (Fig. 5j-k). The U-ExM also showed that the rhoptries (another organelle formed in late stages) were still properly formed in the small defective merozoites and that tubulin appeared as extended structures characteristic of fully developed merozoites. These findings argue against a general development arrest and indicate a more exclusive effect of PfVPS13L1 loss of function in IMC formation (Fig. 5j-k and Extended Data Fig. 9f). Overall, this indicated a failure to sufficiently expand the IMC when PfVPS13L1 function was missing which in turn led to a failure to accommodate enough cytoplasm and a full nucleus into daughter cells dur-ing schizogony, leading to the generation of mini merozoite-like structures and leaving behind a large residual body (Fig. 6).

**Figure 6.**
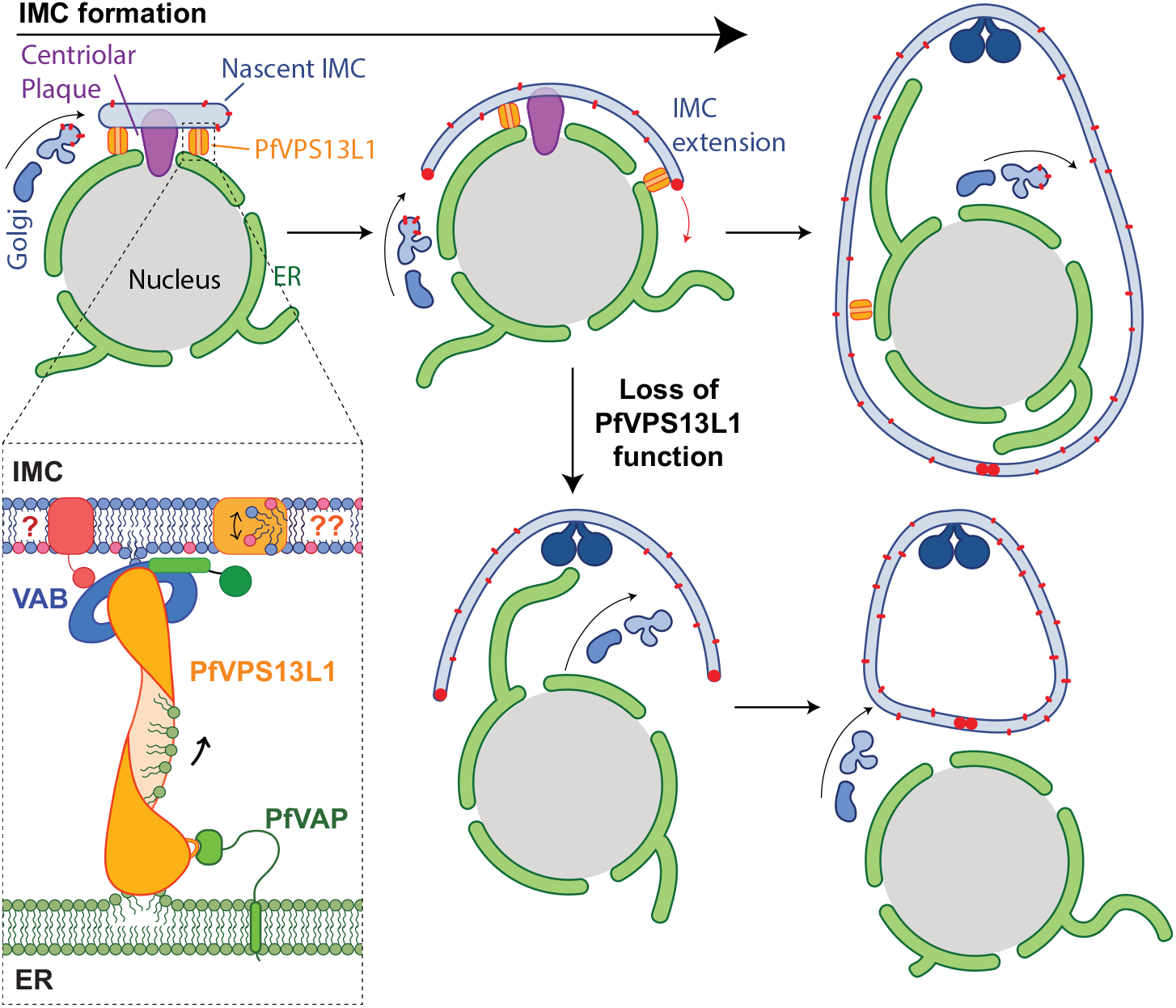
Lipid transfer by PfVPS13L1 at ER-IMC membrane contact sites promotes IMC biogenesis. Schematic summarizing findings of this study. PfVPS13L1 bridges the ER to the nascent IMC via binding of its N-terminal FFAT motif to PfVAP in the ER and of its VAB domain to an unknown partner in the IMC generating an MCS and the direct transport of lipids to support IMC extension.

## Discussion

Protein-mediated direct lipid transfer between organelles is emerging as a key cell biological function in eukaryotes but has not been studied in malaria parasites – a protozoan of high medical importance with a complex life cycle involving rapid organelle expansion and extensive membrane rearrangements. Here we show that a VPS13-like BLTP is needed for the expansion of the IMC, a key structure underlying the PM of alveolates that in malaria parasite schizogony is generated *de novo* and is needed as a scaffold for progeny formation and for host cell invasion^65^. Parasites without the function of this protein were incapable of generating functional progeny, suggesting lipid transfer fulfils key roles in the biology of this parasite and likely also in other apicomplexans. Our results support a model by which PfVPS13L1 enables the rapid and massive membrane extension of the IMC by supplying it with a bulk flow of lipids from the ER (Fig. 6). The N-terminal end of the bridge binds the ER via an interaction of its FFAT motif to PfVAP and the C-terminal VAB domain binds the IMC (Fig. 6). In accordance with this model, MCSs between the ER and the IMC have been reported to occur in *P. falciparum* gametocytes, a stage where the IMC is largely expanded^12^. The disassociation between nuclei and IMC observed in the PfVPS13L1 loss of function phenotype would indicate that its lipid transfer function is coupled with a tethering role to maintain this MCS, as similar, albeit more extreme, disconnections have been observed upon the loss of proteins that hold the inner and outer centriolar plaques together^66^.

Lipid transfer by VPS13 and other BLTPs is usually coordinated with lipid scrambling at the interacting membranes that ensure accurate distribution of lipids in the bilayers during their extraction and delivery^2,67,68^. While it remains to be determined which IMC TM protein could fulfil such a function, we identified proteins with multiple TM domains in the PfVPS13L1 C-terminal proxiome (Supplementary File 4). In addition, the most enriched protein in the PfVPS13L1 C-terminal proxiome was PF3D7_1364000, a protein containing a pair of pore-forming hemolysin domains^69,70^. A potential direct interaction could suggest a functional coordination between these membrane embedding domains and the bulk lipid delivery of PfVPS13L1. Interestingly, a human VPS13 homolog is recruited to the membranes of lysosomes upon rupture^71^, indicating a tendency of these proteins to be recruited to porous membranes.

IMC and segmentation phenotypes have been observed upon the loss of function of well-established IMC proteins, such as the alveolins IMC1g and IMC1c, or the initial IMC scaffolding protein FBXO1, which upon removal lead to the formation of anucleated merozoites and large residual bodies^63,72^. Addi-tionally, the localization of FBXO1 resembles that observed for the VAB domain of PfVPS13L1, which supports a lipid transfer function of PfVPS13L1 to expand the IMC from its early stages of formation. All of these studies point to a general role of the IMC in ensuring proper segmentation into progeny. Interestingly, impairing vesicular transfer disrupts the IMC in a different way, with IMC targeted proteins remaining cytosolic or localized to a different membrane, indicating the lack of an IMC membrane^73,74^. These observations do not exclude the role of the non-vesicular transfer model of growth proposed here, because an initial IMC “seed”, likely Golgi-derived^65^, would be required for PfVPS13L1 to deliver lipids to (Fig. 6). In fact, membrane seeds are observed in similar organelle biogenesis processes. Namely, the growth of the autophagosome and of the prospore membrane in yeast, in which the BLTPs ATG2 and Vps13, respectively, transfer lipids from the ER to a Golgi-derived membrane seed^2,75-77^. The latter process is highly homologous to IMC formation, as the prospore membrane grows adjacently to the nucleus and the nuclear embedded CP, and in parallel to nuclear division during yeast meiosis^76^. These similarities, and this study as the first evidence for a BLTP function in organelle biogenesis in a protozoan, highlight this as a conserved mechanism for membrane expansion across all eukaryotes.

In addition to identifying PfVPS13L1, we also provide the proteome of the cytosolic face of the ER from living intraerythrocytic stages of *P. falciparum* parasites, obtained by proximity biotinylation using PfVAP as bait. These results showed conservation of FFAT-binding of VAP in the parasite and were used to refine the PfVAP-binding FFAT motif to present a genus-specific consensus. Seven proteins with an experimentally confirmed interacting motif were identified. Two of these proteins (PF3D7_1311600 and PF3D7_1316600) localized to the ER, even though they lack a TM region, and hence are expected to be peripherally recruited to the ER, congruent with a recruitment by PfVAP. Studies of other proteins with FFAT motifs could reveal previously unknown functions of the ER in intraerythrocytic stages. For example, one of the motif-containing proteins identified is PfSIP2 (PF3D7_0604100), an AP2 transcription factor binding heterochromatin known to be enriched at the inner nuclear membrane (INM)^78^, where VAP has known interactors in other organisms^36^. Our study raises the possibility that PfVAP serves as a way to anchoring PfSIP2 and potentially chromatin to the INM.

Besides localizing PfVPS13L1 at the ER-IMC MCSs, our study also identified the LTP PfOSBP at the ER-Golgi MCSs, a function conserved for this protein in opisthokonts, and up to 5 other LTPs in proximity to the ER. This study thus largely expands the repertoire of studied *P. falciparum* LTPs and MCSs. As shown by previous studies^8,9^, LTPs represent a potential target for antimalarial drugs that could act by blocking their lipid transfer capabilities.

## Methods

### Parasite culture, synchronization and transfections

*P. falciparum* 3D7 parasites were cultured in RPMI1640 medium containing 0.5% Albumax II (Gibco #11021045), 10 mM Glucose, 12 mM sodium bicarbonate and 20 μg/ml Gentamycin (Ratiopharm), with a 5% haematocrit of human O+ erythrocytes at 37 °C and in a controlled gas environment of 5% O2, 5% CO2, 90% N2, following standard culture protocol. Transfusion human O+ blood was commercially purchased from Universitätsklinikum Hamburg-Eppendorf (Approval number 10569a/96-1). Age, sex and identity of blood donors was not known. For synchronization of ring stage parasites, cultures were centrifuged at 2300 g for 3 minutes and the erythrocyte pellet was resuspended in 10 pellet volumes of 5% sorbitol, incubated for 10minutes at 37 °C, followed by one wash with medium. For enrichment of late schizont stage parasites, 10 ml of cultures were slowly overlaid on 3 ml of 60% Percoll, and centrifuged at 2300 g for 3 minutes without brake. The layer of schizonts above the Percoll was collected and washed with medium once. For transfections, Percoll-enriched late schizonts (originating from 400 μl of erythrocytes at 5% parasitaemia per transfection) were resuspended in 90 μl of a buffer containing 90 mM NaPO4, 5 mM KCl, 0.15 mM CaCl2 and 50 mM HEPES pH 7.6, and electroporated with 50 μg of plasmid in 10μl of TE buffer, using the Amaxa system (Lonza Nucleofector II AAD-1001N, program U-033) as previously described^79^. Electroporated parasites were incubated shaking (800 rpm) at 37 °C with 200 μl of fresh erythrocytes and 500 μl of medium for 30-60 minutes and were then transferred to standard culture conditions. A day after transfection, parasites containing the plasmid were selected with the corresponding drug (4 nM WR99210 (Jacobus Pharmaceuticals), 0.9 μM DSM1 (Merck #5.33304.0001), 400 μg/mL G418 (Merck #A1720), or 2 μg/μl blasticidin S (Invitrogen #R21001) for plasmids encoding the hDHFR, yDHODH, Neo-R, or BSD resistance genes, respectively). Medium and drugs were changed daily for 4 days, and parasites were periodically monitored.

### Plasmids

Plasmids generated for this study were cloned by Gibson ligation reaction (a mixture of T5 exonuclease (Epicenter #T5E4111K), Phusion DNA polymerase (NEB #M0530S), Taq DNA ligase (NEB #M0208L), dNTPs (Roth #K039.1), NAD (Sigma-Aldrich #n6522), DTT, MgCl2 and Tris-HCl pH 7.5), linearizing a backbone with restriction enzymes and ligating in the insert that was PCR amplified with primers containing at least 20 nucleotides of overlap with the target backbone. In cases where the insert was smaller than 80 nucleotides, these were ordered to be synthesized as oligonucleotides including the overlapping regions. All oligonucleotides used were ordered from Sigma-Aldrich and all restriction enzymes used were ordered from NEB. In cases where a re-codonized sequence was required, synthesized DNA fragments were ordered from BioCat GmbH (Heidelberg, DE). A list of all the plasmids generated in this study and their respective cloning reactions is found in Supplementary File 5, and the expected full plasmid sequences were deposited in a public database^80^. Other plasmids used for transfections in this work, mCh-FRB-miniTurbo, GRASP1-mCh and pLyn-FRB-mCh-nmd3-BSD (Addgene #85796) were generated by previous studies^32-73^.

Used as templates or backbones for further cloning reactions, pSLI-N-GFP-2xFKBP-loxP (Addgene #85792), pSLI-C-GFP-Sandwich (also pSLI-sandwich, Addgene #85790), pSLI-C-mNG-Sandwich (same as addgene #85792 with mNG replacing GFP), pSkipFlox (Addgene #85797), mSc-Rab6_-_NLS-FRB, Sf3A2 mCh-Kelch13, pLyn-FRB-mCh-nmd3-BSD (Addgene #85796), were previously generated in our laboratory^32-33^, and GAPM1-GFP and GAP45-mCh^62^ were gifts from T. Gilberger (CSSB, Hamburg, DE).

### Selection linked integration (SLI)

The SLI approach was also used for endogenously editing all genes analyzed in this study, with slightly different plasmids depending on whether the modification was N- or C-terminal, as the N-terminal modifications require the insertion of a second recodonized version of the gene (as shown in Extended Data Fig. 1a) but the C-terminal ones do not. Methodologically, SLI was carried out as described^32^. Briefly, parasites were transfected with a SLI plasmid and selected with WR99210. When the parasites had resurfaced and were growing at usual rates, the drug in the medium was replaced by a different drug to which only parasites that had properly integrated the plasmid in their genome will be resistant. These drugs were G418 (in the case of C-terminal gene modifications) or DSM1 (in the case of N-terminal gene modifications). Medium and drug were changed daily for 5 days and cultures were periodically monitored, thereafter every second day until parasites resurfaced. Once integrated parasites were growing at usual rates, correct integration was evaluated by a diagnostic PCR from genomic DNA (collected using Monarch #T3010L kit). Diagnostic PCR involved reactions spanning the two integration junctions and the absence of the unmodified gene locus and were performed using Firepol enzyme (Solis biodyne #01-01-02000). Genomic sequences of the original and modified loci, including the primers and amplicon sizes for diagnostic PCR, are provided in Supplementary File 6.

### Labelling of parasites and live-cell microscopy

For direct staining, parasites were incubated with 50 ng/ml Hoechst 33342 (Biomol #ABD-17533), 1 μM Tubulin Tracker Deep Red (Invitrogen #T34076) or 50 nM Halo-JF585 or - JFX650 ligands^81^ (kind gift from L. Lavis, Janelia Farm, Ashburn VA) for nuclear, tubulin or Halo-tag visualization, respectively. All staining reactions were carried out for 20 minutes at 37 °C in RPMI-1640 medium after which the parasites were washed once with medium, with an additional 10 minute incubation and wash in the case of Halo dyes.

For wide-field fluorescence microscopy, parasites in RPMI medium were placed between a glass slide and a cover slip and imaged in a Zeiss AxioImager microscope equipped with a Hamamatsu Orca C4742-95 camera and a plan apochromat objective (63x, 1.4 NA, Oil DIC). Images were acquired using the AxioVision software (version 4.7) and processed using FIJI^82^ to adjust for brightness and contrast and in a few cases apply a gaussian filter.

For laser scanning confocal microscopy of live cells was performed at 37 °C using an Olympus FluoView FV3000 system equipped with a universal plan apochromat objective (60×, 1.5 NA, oil) and a cell Vivo incubation system. Time lapse imaging of IMC formation was done as described^83^. Briefly, infected RBCs (34-38 hpi) were plated in a glass-bottomed dish (Ibidi #80427) previously coated with 0.5 mg/ml concanavalin A (Sigma-Aldrich #C0412) and allowed to attach for 10 minutes. Unbound cells were washed off with pre-warmed DPBS, the dish filled up completely with fresh medium (previously pre-adsorbed with uninfected RBCs), sealed with parafilm and placed into the microscope. Z-stacks were acquired every 20 minutes. Resulting images were processed with FIJI^82^ or Imaris (Oxford Instruments), to adjust for brightness and contrast and apply a gaussian filter.

### Ultra-expansion microscopy (U-ExM)

U-ExM was performed as previously described^61^. Parasites were placed on 3 mm PDL-coated coverslips and allowed to attach for 20 minutes at 37°C. Samples were then washed with PBS, fixed with 4% formaldehyde in PBS for 20 minutes, and left overnight in post-fix solution [1.4% formaldehyde and 2% acrylamide (Merck #A4058)], all at 37 °C. The coverslips were then washed twice with PBS and placed cells facing down on 35 μl of monomer solution [19% sodium acrylate (Sigma-Aldrich #408220)], 10% acrylamide and 0.1% N,N’-methylenebisacrylamide (Merck #M1533)] with freshly added TEMED (EMD Millipore #1.10732) and APS (Sigma-Aldrich #A3678) to a final concentration of 0.5% each. Gels were allowed to polymerize at 37 °C for 1 hour and were then incubated shaking at RT on denaturation buffer (200 mM SDS, 200 mM NaCl, 50 mM Tris-HCl pH 9). After the gels had separated from the coverslip, they were further incubated in denaturation buffer for 90 minutes at 95°C, and were either froze at -20 °C or placed in water for three rounds of expansion of 30 minutes each. Expanded gels were shrunk by placing them in PBS for two 15-minute rounds, and a slice of gel was cut for antibody staining. This slice was blocked with 3% BSA (Biomol #9048-46-8) in PBS for 30 minutes at RT, and left incubating with primary antibodies [anti-V5 (1/250, BioRad #MCA1360), anti-αTubulin (1/500, Thermo Fischer #32-3500), anti-mCh (1/1000, abcam #ab167453)] shaking overnight at RT. Gels were then washed three times with 0.5% Tween in PBS, shaking for 10 minutes at RT, and incubated with secondary antibodies (anti-mouse IgG Alexa Fluor™ 594 (Thermo Fischer #A11032), anti-mouse IgG Alexa Fluor™ 633 (Thermo Fischer #A21053), anti-rabbit IgG Alexa Fluor™ 546 (Thermo Fischer #A10040) and anti-rabbit IgG Alexa Fluor™ 647 (Thermo Fischer #A21244), all used 1/500) and/or NHS-ester (1/250, Thermo Fischer #46403), Bodipy TR Ceramide (1/500, Invitrogen #B34400) and Hoechst 33342 for 3h at RT. Gels were washed three times with 0.5% Tween in PBS and were then placed on a glass-bottom dish (Ibidi #80427) previously coated with PDL for imaging in an Olympus FluoView FV3000 laser scanning confocal microscope equipped with a universal plan super apochromat objective (60x, 1.3 NA, silicone). Images were processed using FIJI^82^ to adjust brightness and contrast, and obtain Z-stack max intensity projections of the number of Z-slices specified in each image.

### DiQ-BioID

#### Preparation of streptavidin sepharose beads for protease resistance

Streptavidin sepharose beads were treated to be protease resistant for reducing the presence of streptavidin in the eluted peptides after on-bead trypsin digestion. This treatment was carried out as previously described by others84. Briefly, sepharose beads (Cytiva #17511301) were treated with cyclohexanedione (Merck #C101400) for 4 h at RT, washed with PBS-0.1% Tween (PBS-T), treated with a solution of 4% formaldehyde and 0.1 M of sodium cyanoborohydride (Sigma-Aldrich #8.18053) for 2 h at RT and washed with 0.1 M Tris-HCl pH 7.5 and twice with PBS-T.

#### DiQ-BioID sample collection

DiQ-BioID experiments were perfomed as previously outlined^33^. Briefly, cultures of genetically edited parasite episomally expressing the miniTurbo biotinyliser were grown to 200 mL in biotin-free RPMI-1640 medium (Biozol #USB-R9002-01). Asynchronous cultures were used for PfVAP and cultures synchronized to 34-38 hpi (by subsequent Percoll-Sorbitol synchronization to obtain 0-4 hpi rings) were used for PfVPS13L1. The culture was split into two, and in one half the dimerization of the biotinilyser with the POI (PfVPS13L1 or PfVAP) was induced by addition of 250 nM rapalog. Immediately after, 50 μM of biotin (Sigma-Aldrich #B4639) was added to both cultures for 30 minutes of labelling at standard growth conditions. Thereafter, RBCs were harvested, washed with 1xPBS and parasites were isolated by lysis with 0.03% saponin (in 1xPBS) on ice for 10 minutes, washed five times with 1xPBS and lysed in 2 ml of lysis buffer (50mM Tris-HCL pH 7.5, 500 mM NaCl, 1% Triton-X-100, 0.4% SDS) supplemented with 1 mM DTT, 1 mM PMSF and 1 x protease inhibitor cocktail (Roche #11836170001). The lysate was frozen at -80 °C until affinity purification was carried out.

#### Affinity purification of biotin-labelled proteins

Lysates were thawed and frozen twice, then centrifuged at 16,000xg for 10 to 60 minutes to clear the lysate. The supernatant was diluted 3-fold in 50 mM Tris-HCl pH 7.5 and incubated with 50 μl of pre-treated (for protease-resistance) streptavidin sepharose beads. The cleared lysate was incubated with the sepharose beads rotating overnight at 4°C. The beads were washed twice with lysis buffer, once with dH20, twice with Tris-HCl pH 7.5 and three times in 100 mM TEAB (Sigma-Aldrich #T7408) pH 8.5, before adding 50 μl of elution buffer (2 M urea and 10 mM DTT in 100 mM Tris-HCl pH 7.5) and incubation for 20 minutes with shaking at 1400rpm at RT. On-bead digestion was then performed, first by adding 5 μl iodoacetamide to a final concentration of 50 mM and incubation for 10 minutes shaking in the dark for protein alkylation, and then by adding 500 ng of Trypsin (Sigma-Aldrich #T0303) and shaking for 2 hours, all at RT. The supernatant was collected and the beads were resuspended in 50 μl of elution buffer and incubated for 15 min. The supernatant was again collected and combined with the previous one and 200 ng of additional trypsin was added and left shaking at RT overnight. Tryptic peptides were frozen at -80°C and shipped to the EMBL proteomics core facility (Heidelberg, DE), where MS analysis was carried out.

#### Preparation of peptides for MS analysis

Peptides were dried and reconstituted in 10 μl of 100 mM Heps/NaOH, pH 8.5 and reacted for 1 h at room temperature with 80 μg of TMT6plex (Thermo Fischer #90066) dissolved in 4 μl of acetonitrile. Excess TMT reagent was quenched by the addition of 4 μl of an aqueous 5% hydroxylamine solution (Sigma-Aldrich #438227). Peptides were reconstituted in 0.1% formic acid, mixed and purified by a reverse phase clean-up step (OASIS HLB 96-well μElution Plate, Waters #186001828BA). Peptides were subjected to an offline fractionation under high pH conditions^85^. The resulting 6 fractions were analyzed on a Lumos system (Thermo Fischer).

#### LC-MS/MS analysis

Peptides were separated using an Ultimate 3000 nano RSLC system (Dionex) equipped with a trapping cartridge (Precolumn C18 PepMap100, 5 mm, 300 μm i.d., 5 μm, 100 Å) and an analytical column (Acclaim PepMap 100. 75 × 50 cm C18, 3 mm, 100 Å) connected to a nanospray-Flex ion source. The peptides were loaded onto the trap column at 30 μl per min using solvent A (0.1% formic acid) and eluted using a gradient from 2 to 38% Solvent B (0.1% formic acid in acetonitrile) over 52 min and then to 80% at 0.3 μl per min (all solvents were of LC-MS grade). The Orbitrap Fusion Lumos was operated in positive ion mode with a spray voltage of 2.4 kV and capillary temperature of 275 °C. Full scan MS spectra with a mass range of 375–1500 m/z were acquired in profile mode using a resolution of 60,000 with a maximum injection time of 50 ms, AGC operated in standard mode and a RF lens setting of 30%.

Fragmentation was triggered for 3 s cycle time for peptide-like features with charge states of 2–7 on the MS scan (data-dependent acquisition). Precursors were isolated using the quadrupole with a window of 0.7 m/z and fragmented with a normalized collision energy of 36%. Fragment mass spectra were acquired in profile mode and a resolution of 30,000 in profile mode. Maximum injection time was set to 94 ms or an AGC target of 200%. The dynamic exclu-sion was set to 60 s. Acquired data were analyzed using FragPipe^86^ and a Uniprot *Plasmodium falciparum* isolate 3D7 fasta database (UP000001450, ID 36329, 5.372 entries, date: 06.03.2023, downloaded: 17.05.2023) including common contaminants. The following modifications were considered: Carbamidomethyl (C, fixed), TMT6plex (K, fixed), Acetyl (N-term, variable), Oxidation (M, variable) and TMT6plex (N-term, variable). The mass error tolerance for full scan MS spectra was set to 10 ppm and for MS/MS spectra to 0.02 Da. A maximum of 2 missed cleavages were allowed. A minimum of 2 unique peptides with a peptide length of at least seven amino acids and a false discovery rate below 0.01 were required on the peptide and protein level^87^.

The raw output files of FragPipe were processed using the R programming language (ISBN 3-900051-07-0). Contaminants and reverse proteins were filtered out and only proteins that were quantified with at least 2 razor peptides were considered for the analysis. Log2 transformed raw TMT reporter ion intensities were first cleaned for batch effects using the ‘removeBatchEffect’ function of the limma package^88^ and further normalized using the ‘normalizeVSN’ function of the limma package (VSN - variance stabilization normalization^89^). Proteins were tested for differential expression using a moderated t-test by applying the limma package (‘lmFit’ and ‘eBayes’ functions). The replicate information was added as a factor in the design matrix given as an argument to the ‘lmFit’ function of limma.

### Loss-of-function induction by knock sideways or conditional gene excision

Knock-sideways (KS) was induced by addition of 250 nM Rapalog to a cell line expressing the POI endogenously fused to 4 FKBP domains and an episomally expressed FRB protein anchored to the PM via a Lyn-targeting region. Mislocalization to the PM was assessed by microscopy after 5 hours of induction. In the case of PfVPS13L1, as the IMC and the PM are in close proximity, the mislocalization was assessed by comparison with the IMC marker PhIL1 (no overlap: full mis-localization; overlap only of 1-2 PfVPS13L1 puncta: partial mislocalization).

Gene excision was induced by addition of 250 nM Rapalog to a cell line expressing PfVAP flanked by two loxP sites at its endogenous locus (Extended Data Fig. 1a-b) and an episomally expressed diCre protein using pSkipFlox as done previously^32^. Excision was assessed by PCR one cycle after induction, using primers annealing to sites outside the loxP flanked region which would result in a reduced amplicon size upon correct gene excision (Supplementary File 6).

### Parasite growth assays

Importance of proteins for parasite survival was tested by monitoring the growth of asynchronous control and diCre-excised or KS parasite cultures by flow cytometry-based measurements of parasitemia every day over 7 (for diCre excision) or 5 (for KS) days. Samples were stained with Hoechst 33342 and Dihydroethidium (Cayman #12013-5) for 20 minutes at room temperature, and the parasites were inactivated and the staining stopped with 400 μl of 0.003% glutaraldehyde (Roth #4157) containing medium. Parasitemia was measured as the percentage of positively stained cells over 100000 events counted with an LSRII flow cytometer (BD Biosciences) with a FACSDiva software (BD Biosciences) as described^32,90^.

For stage specific growth assays, parasites were first synchronized. For GFP-2xFKBP-PfVAP^endo^, ring stage para-sites were synchronized to 10-18hpi by two 10-minute incubation with 5% sorbitol 10 hours apart. For PfVPS13L1-GFP-SW^endo^, ring stage parasites were synchronized to 0-4hpi by an initial Percoll enrichment of schizonts and sorbitol four hours after adding the parasites to fresh erythrocytes^83^. KS or diCre excision was induced with 250 nM Rapalog (Takara Bio), and Giemsa (Merck) smears were taken at the indicated time points.

### Egress and invasion assays

PfVPS13L1-GFP-SW knocked-in parasites were synchronized by Percoll followed by sorbitol after four hours. Parasites were grown for another 34 hours (34-38hpi), at which point they were separated in two cultures, one which was treated with Rapalog and one kept as a control. Six hours after (40-44hpi), 2 μM Compound 2^64,91,92^(imidazopyridine referred to as Compound 2 (4-[7-[(dimethylamino)methyl]-2-(4-fluorophenyl)imidazo [1,2-α]pyridine-3-yl]pyrimidin-2-amine), a kind gift from Michael J. Blackman [MRC-NIMR, London, UK]) was added to arrest parasites at the segmenter stage. Giemsa smears were prepared at 44-48hpi (pre-egress), the Compound 2 was washed off, parasites were allowed to egress and invade new erythrocytes for four hours, and a second set of Giemsa smears was prepared at 48-52hpi (post-egress). The number of rings and schizonts in both sets of smears was counted and used to calculate the number of new rings in the post-egress sample per schizont in the pre-egress sample.

### FFAT prediction and AlphaFold3 mediated filtering

Filtering of PfVAP DiQ-BioID hits using FFAT motif identification was done by sequence scoring and AlphaFold3^41^ predicted binding of the motif to PfVAP. First, the lowest-scoring motif (i.e. most likely to be binding partner) was identified for each of the protein hits. Prediction of FFAT motifs within a protein sequence was carried out using the scoring matrix previously proposed based on motifs identified in yeast and human proteins^40^. A python code written in collaboration with ChatGPT (OpenAI, 2023) was used and is deposited in a public repository^80^. Briefly, the code scores any sequence of 7 amino acid residues following the scoring matrix, giving a value to each position of the motif based on the consensus sequence. The negative charge of the 6 residues upstream of the potential motif is also considered. Motifs scoring 4 or below are considered potential FFAT binders and were considered for AlphaFold3-mediated filtering. For this, structural predictions of the MSP domain and the selected FFAT motif interactions were obtained using the AlphaFold3 server, and motifs with an interface predicted template modelling (ipTM) of 0.7 were selected for further analysis. Full-length PfVAP and the full-length protein candidates containing the selected motifs were then submitted to the AlphaFold3 server and the structural predictions with an ipTM value of 0.45 or above (corresponding to the higher half of all the ipTM values) were selected as likely PfVAP binders. The 13 motifs remaining after this filtering were selected for episomal expression in Halo-PfVAP^endo^ parasites to assess binding. After determining binding, the FFAT motif scoring matrix was redefined to increase accuracy of identification in *Plasmodium*.

### Sequence and structural analysis

Protein sequences were retrieved from PlasmoDB (release 67). For conservation analysis across *Plasmodium* species (FFAT-containing proteins in Fig 2i and PfVPS13L proteins in Extended Data. Fig. 4a), sequences from reference strains of *P. falciparum, P. berghei, P. knowlesi, P. ovale, P. malariae, P. vivax-like, P. vivax, P. gallinaceaum, P. chabaudi, P. yoelii* and *P. coatneyi* were considered. Maximum-likelihood phylogenetic tree of PfVPS13L proteins was generated using MEGA11^93^ with 100 bootstrapping repeats.

LTPs were identified by sequence and structure homology, using HHPred^94^ and Foldseek^95^. Structures were downloaded from the AlphaFold public database^96,97^, for the shuttle-like LTPs, or predicted using AlphaFold3^41^, for the BLTPs, and analysed using PyMOL^98^. For the predictions, the BLTP sequences were split in 2 or 3 overlapping segments, which were then aligned in PyMOL to get the full-length structure of PfVPS13L1. The structures obtained were deposited in a public repository^80^. Shuttle-like LTP structures were aligned with indicated structures in Extended Data Fig. 3a using TM-align^99^.

## Data availability

The *Plasmodium falciparum* protein database used in this study can be accessed at PlasmoDB^100^, release 67. The mass spectrometry proteomics data have been deposited to the ProteomeXchange Consortium via the PRIDE^101^ partner repository with the dataset identifier PXD066159. AlphaFold3^41^ predicted structures and code for FFAT analysis have been deposited in the Zenodo public repository^80^. All other data are included in the article and supplementary files. Source data are provided.

## Acknowledgements

We thank J. S. Wichers-Misterek for essential discussion before the beginning of this project. We thank J. Cubillán-Marín and I. Henshall, from the malaria cell biology laboratory (BNITM, Hamburg) and T. Gilberger (CSSB, Hamburg) for discussion. The manuscript file uploaded to biorXiv was generated using the LaTeX template adapted by S. Royle available at https://github.com/quantixed/manuscript-templates. A. Guillén-Samander was supported by an EMBO long-term postdoctoral fellowship (ALTF 166-2022).

## Author contributions

A. Guillén-Samander and T. Spielmann conceptualized the project. A. Guillén-Samander designed and performed most experiments and analysed the data. N. Perepelkina performed the FFAT filtering; V. Horáčková, H. Behrens and J. Mesén-Ramirez each generated one parasite line used in this study; A. Ribeiro-Holbein contributed to data analysis; P. Haberkant and F. Stein performed the LC-MS/MS analysis. A. Guillén-Samander prepared the figures. The original draft was written by A. Guillén-Samander, revised and edited by T. Spielmann, and reviewed by all the authors.

## Supplementary files provided

**Supplementary File 1:** PfVAP DiQ-BioID results.

**Supplementary File 2:** Highest scoring FFAT motifs in *P. falciparum* proteome and DiQ-BioID results.

**Supplementary File 3:** Filtering of PfVAP DiQ-BioID results to identify potential direct interactors.

**Supplementary File 4:** PfVPS13L1 C-terminus DiQ-BioID results.

**Supplementary File 5:** Oligos and DNA fragments synthesized for this study.

**Supplementary File 6:** Genomic sequences of loci edited in this study.

## Movie legends

**Movie S1. PfVPS13L1 structure**. Rotating AlphaFold3 predicted structure (N-terminus to the left). Structure representations transition from 1) ribbon representation colored by AlphaFold3 prediction confidence (as in Fig. 4b), 2) ribbon representation colored by domain annotation (as in Fig. 4c), and 3) surface representation where the lipid transfer groove was colored by element (Oxygen, red; Nitrogen, blue; Carbon, white; Sulfur, yellow).

**Movie S2. PfVPS13L1 mislocalization leads to reduced IMC growth and faulty egress**. Timelapse confocal microscopy of PfVPS13L1-GFP-SWendo parasites episomally expressing the IMC marker Halo-PhIL1 and the PM mislocalization construct Lyn-FRB-mCh without (control) or with rapalog added at 34-38hpi (knock-sideways). Z-stacks were acquired every 20min, starting at 36-40hpi. 4 cells for each condition are included. Scale bar, 2 μm.

**Extended Data Figure 1.**
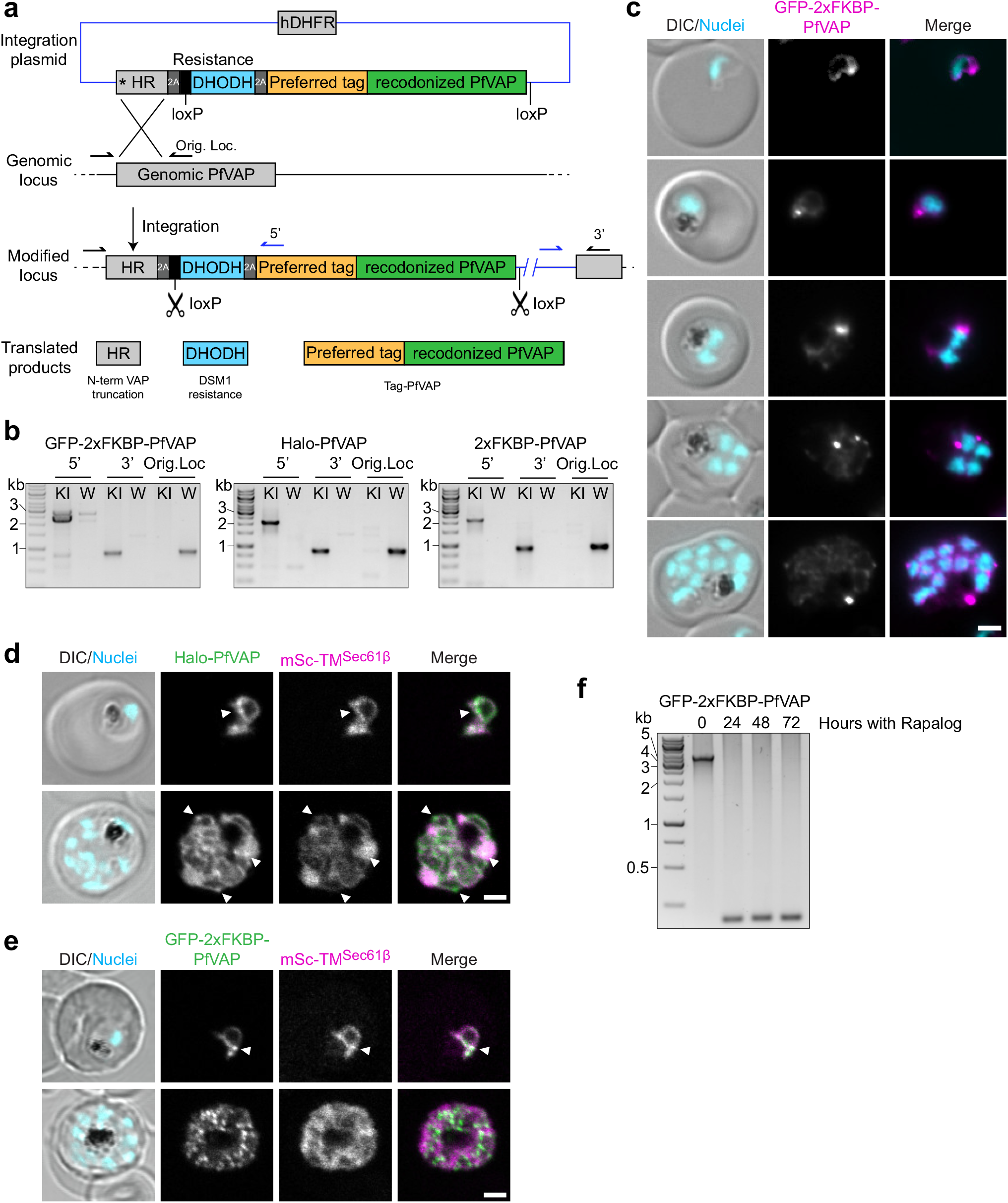
PfVAP is an essential ER protein. **a** Schematic depicting selection-linked integration (SLI) strategy to edit the genomic locus of PfVAP to express N-terminally tagged versions of the protein under its original promoter. The expression of the drug resistance gene yDHODH fused to PfVAP with a skip peptide (T2A) was used to select edited parasites. The asterisk indicates a stop codon. **b**, Agarose gels showing PCR products amplified from genomic DNA of the indicated cell lines confirming correct integration of genome-modified parasites: 5’ and 3’ are amplicons generated across the integration junctions and the original locus amplicon (Orig. Loc.) shows presence or absence of the unmodified locus, comparing parental (W) and knock-in (KI) lines. **c**, Fluorescence microscopy images of GFP-2xFKBP-PfVAP^endo^, showing the artifactual formation of PfVAP accumulations in the ER across all parasite stages, potentially mediated by GFP dimerization. **d-e**, Representative confocal images of parasites with endogenously edited PfVAP and episomally co-expressing mSc-TM^Sec61β^, reflecting an enrichment at hotspots in the ER (arrowheads). This is seen with Halo-tagged PfVAP (d), and artifactually exacerbated with GFP-tagged PfVAP (e). **f**, Agarose gels with PCR products amplified from genomic DNA confirming the excision of PfVAP upon rapalog-mediated diCre activation. DIC, differential interference contrast; Nuclei, Hoechst 33342; scale bars, 2 μm.

**Extended Data Figure 2.**
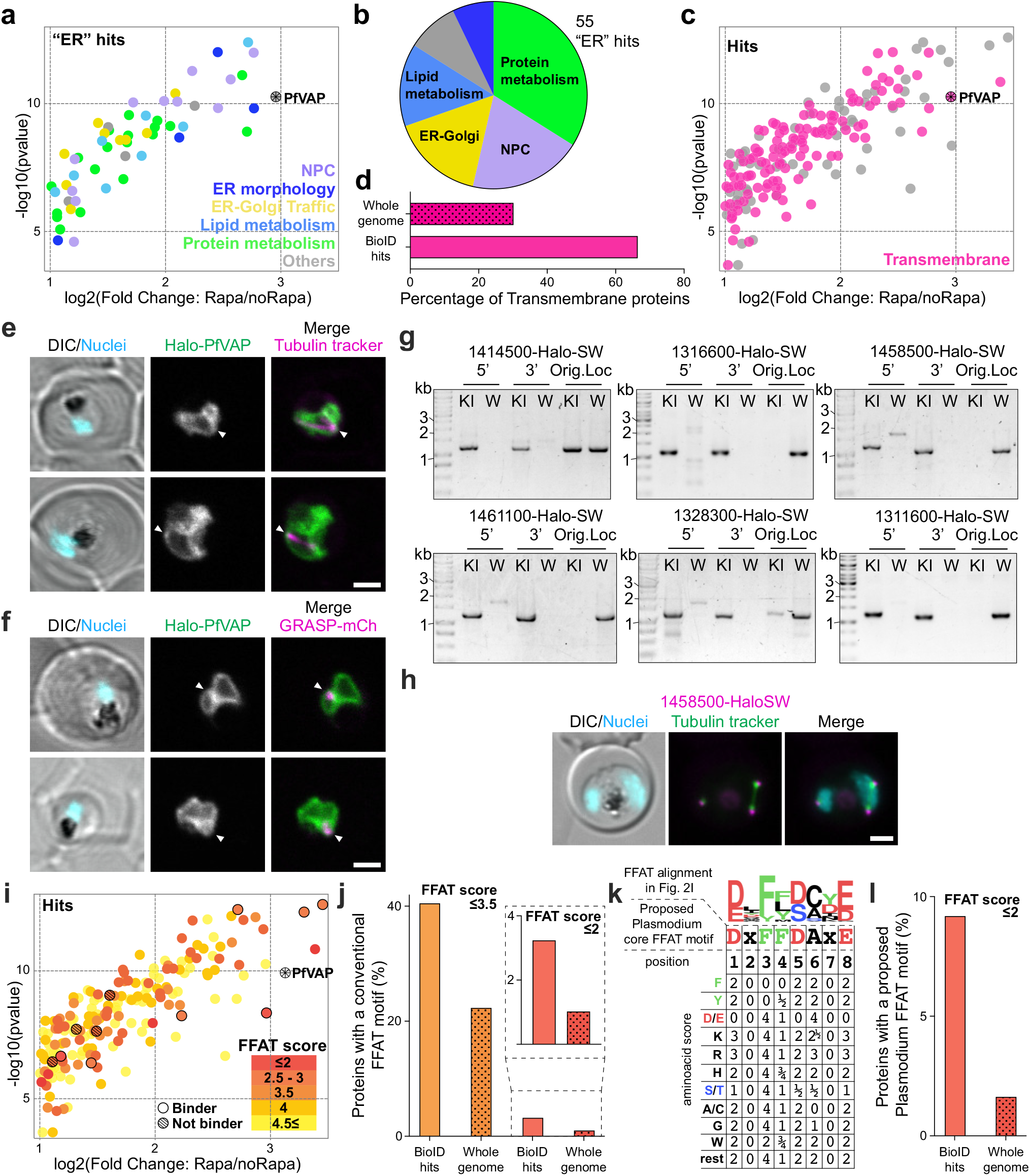
PfVAP DiQ-BioID hits are enriched in ER and FFAT-motif containing proteins. **a**, PfVAPDiQ-BioID hits annotated as “ER” proteins in Fig. 2c classified by function. NPC: Nuclear Pore Complex. **b**, Pie chart showing proportion of proteins from (a) with the indicated functional classification. **c**, PfVAP DiQ-BioID hits from Fig. 2c annotated by the presence of a transmembrane domain. **d**, Graph comparing the percentage of transmembrane proteins in the hits in (c) with the proportion of all proteins annotated from the *P. falciparum* genome. **e-f**, Representative confocal images showing Halo-PfVAP and the Centriolar Plaque (e) or the Golgi (f), as labelled by tubulin tracker and episomally expressed GRASP-mCherry (GRASP-mCh), respectively. Arrowheads show areas with overlap. Both compartments are annotated as “Others” in Fig. 2c and are in proximity to PfVAP. **g**, Agarose gels showing PCR products amplified from genomic DNA of the indicated cell lines confirming correct integration of genome-modified parasites. Features as in Extended Data Fig. 1b. In the case of the PF3D7_1414500-Halo-SW line, there is still a significant population of unedited parasites, with no fluorescence signal, but this does not affect localization experiments which was the sole purpose of this line for this study. **h**, Representative fluorescence microscopy images showing PF3D7_1458500 (SAS4) adjacent to the tubulin foci that labels the inner centriolar plaque, confirming the localization of this candidate to the outer centriolar plaque. **i**, PfVAP DiQ-BioID hits from Fig. 2c colored by the score of their highest scoring FFAT-motif, calculated based on the opisthokont FFAT consensus motif^40^. The proteins containing the motifs analyzed in Fig. 2g-h are highlighted with a black stroke, of which those determined as non-binders are filled with diagonal lines. **j**, Bar chart showing percentages of proteins containing high-scoring FFAT motifs of the 185 PfVAP DiQ-BioID hits (BioID hits) and in all proteins encoded in the *P. falciparum* genome (whole genome) as calculated in Slee, J.A. & Levine, T.P.^40^) using a cut-off of 3.5 or lower, or 2 or lower (inlay). **k**, Table of FFAT-motif scoring system proposed based on our findings. The scoring modifications were made to the core FFAT motif, which was extended from 7 to 8 amino acids to consider the presence of an acidic amino acid in position 1 (formerly position -1 in opisthokont motif). The scoring of the upstream acidic region was maintained. l, Bar chart showing percentages of proteins containing high-scoring FFAT motifs as in (j) but calculated based on the new proposed scoring system, which results in larger overrepresentation of motifs within the PfVAP DiQ-BioID hits. DIC, differential interference contrast; Nuclei, Hoechst 33342; scale bars, 2 μm.

**Extended Data Figure 3.**
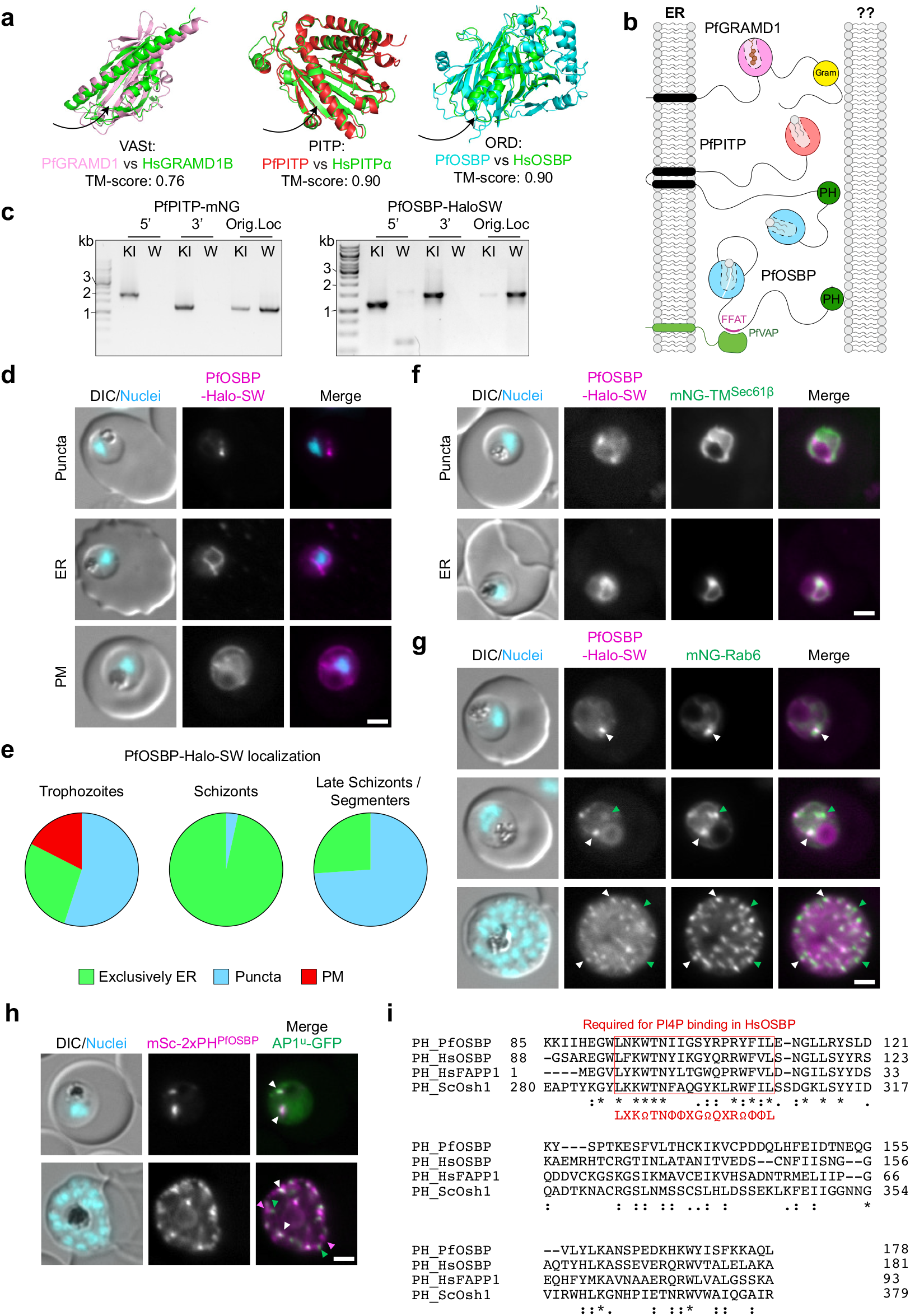
The Shuttle-like LTP PfOSBP mediates ER-Golgi MCSs. **a**, Structural alignment using TM-align^99^ between the AlphaFold predicted structures of domains found in P. falciparum LTPs and structures of human LTPs: ORD of HsOSBP (PDB 7V62^103^), HsPITPα (PDB 1UW5^104^) and VASt of HsGRAMD1B (AF-Q3KR37-F1-v4^96,97^). **b**, Schematics indicating potential arrangement of the shuttles found in this study at ER-MCSs. **c**, Agarose gels showing PCR products amplified from genomic DNA of the indicated cell lines confirming correct integration of genome-modified parasites. Features as in Extended data Fig. 1b. In the case of the PfPITP-mNG line, there are some unedited parasites, with no fluorescence signal, but this does not affect localization experiments which was the sole purpose of this line for this study. **d**, Representative fluorescence microscopy images of PfOSBP-Halo-SW dynamic localization to the ER (expected due to the interaction of its FFAT with PfVAP), either exclusively to the ER (ER) or with puncta or with the PM (PM). **e**, Proportion of the phenotypes from (d) from a total of 40 trophozoites, 29 schizonts and 23 late schizonts/segmenters from 4 independent experiments. **f-g**, Representative fluorescence microscopy images showing PfOSBP-Halo-SW edited parasites with episomal expression of the ER marker mNG-TM^Sec61β^ (f) or the Golgi marker mNG-Rab6 (g) in trophozoites. The PfOSBP-Halo-SW puncta colocalize with both markers, indicating that these are hotspots where the two organelles overlap. Arrowheads in (g) indicate colocalization (white) or only Rab6 positive (green) puncta. **h**, The PH domain of PfOSBP is recruited to the Golgi in trophozoites and partially in schizonts, as seen by colocalization of an episomal construct encoding a mScarlet tagged tandem of two copies of this PH domain (mSc-2xPH^PfOSBP^) with the μsubunit of the AP-1 complex expressed from the endogenous locus [AP-1μ-GFP (endo)]. Arrowheads in schizonts indicate colocalization (white), only mSc-2xPH^PfOSBP^ (magenta), or only AP-1μ(green) puncta. **i**, Sequence alignment between PH domain of PfOSBP and human and yeast homologues expected to bind PI4P. The red box indicates the PI4P-binding wedge conserved in PfOSBP. Ω, *ϕ* and X, indicate aromatic, hydrophobic and any residue, respectively. In the alignment,., : and *, indicate similar, highly similar, and identical residues, respectively. DIC, differential interference contrast; Nuclei, Hoechst 33342; scale bars, 2 μm.

**Extended Data Figure 4.**
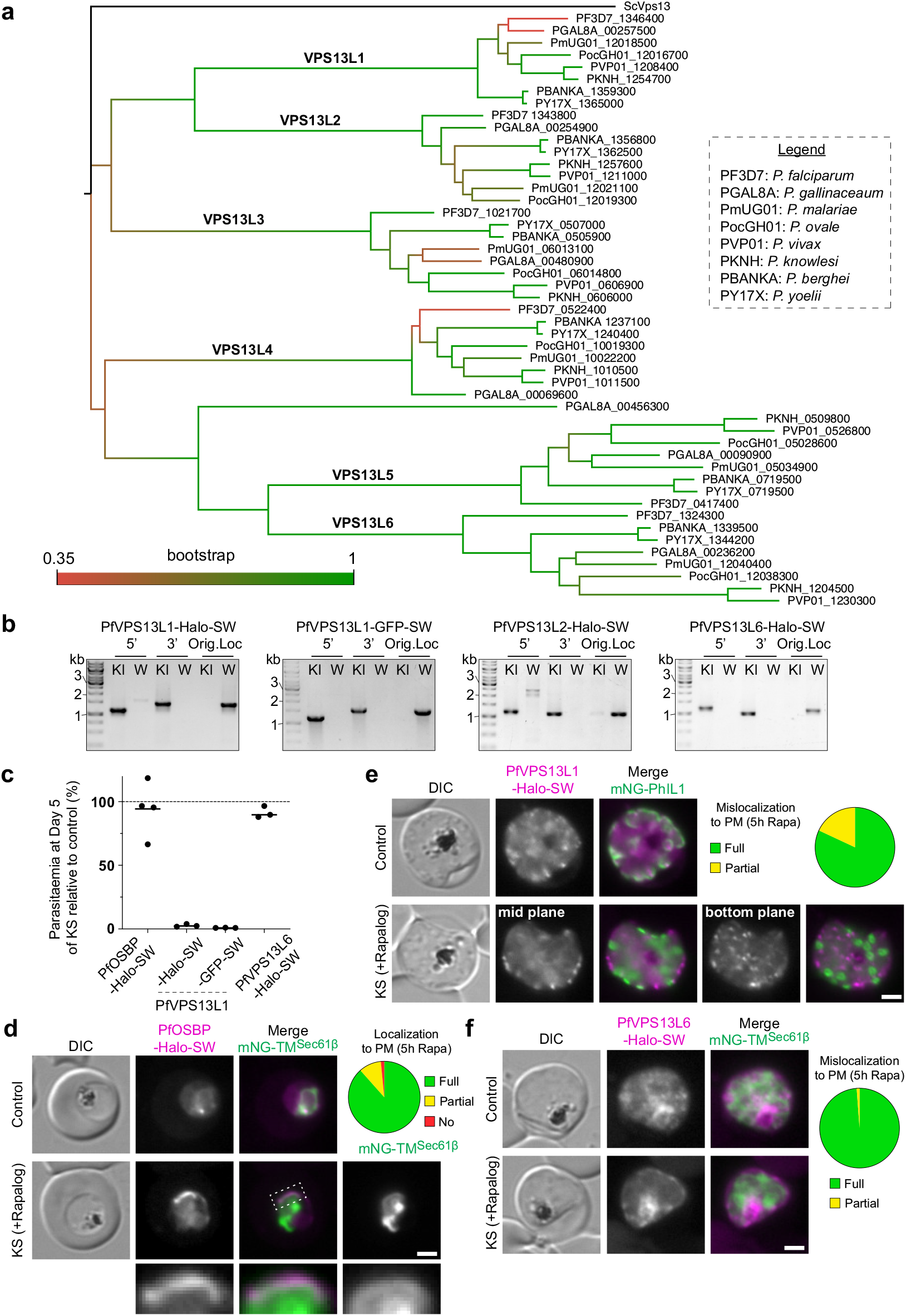
PfVPS13L1 is an essential BLTP. **a**, Phylogenetic tree of the six VPS13-like proteins identified in *P. falciparum* and their homologues in other *Plasmodium* species. **b**, Agarose gels showing PCR products amplified from genomic DNA of the indicated cell lines confirming correct integration of genome-modified parasites. Features as in Extended data Fig. 1b. **c**, Graph summarizing growth defects observed in independent replicates (each indicated by a dot; bar, mean) after 5 days upon knock-sideways (KS) of the indicated protein using the parasite lines generated in (Extended Data Fig. 3c and 4b) episomally expressing the Lyn-FRB mislocalizer. P-values, paired t test between the KS and control: PfOSBP-Halo-SW^endo^, 0.2971; PfVPS13L1-Halo-SW^endo^, <0.0001; PfVPS13L1-GFP-SW^endo^, <.0001; PfVPS13L6-Halo-SW^endo^, 0.0832. **d-f**, Representative fluorescence microscopy images and quantification of the KS of the indicated target proteins. Mislocalization of PfOSBP to the PM (d) also resulted in the formation of ER-PM MCSs, as seen by the ER marker (also shown in the 3.5x enlargement of the boxed area), confirming its attachment to the ER via PfVAP. Similarly, mislocalization of PfVPS13L1-Halo-SW resulted in puncta at the PM (e). Pie charts show localization after rapalog addition, (total of 71 (d), 48 (e) and 56 (f) parasites across 2 experiments for each parasite line). DIC, differential interference contrast; Nuclei, Hoechst 33342; scale bars, 2 μm.

**Extended Data Figure 5.**
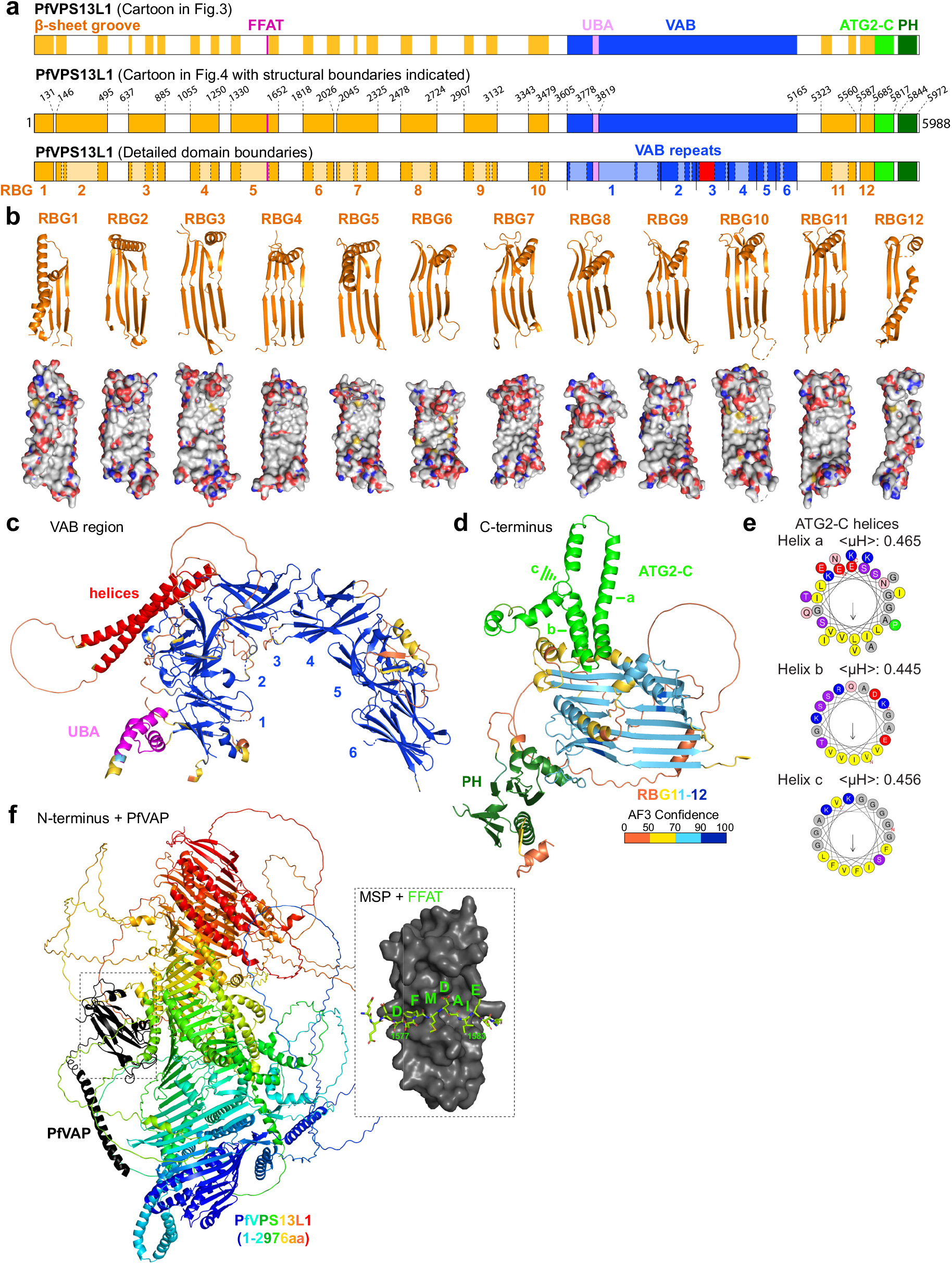
Structural details of PfVPS13L1. **a**, PfVPS13L1 domain cartoons with structural details annotated. Top, annotation of all the pieces that form the lipid transferring β-sheet rod, similarly to the cartoons in Fig. 3d. In the mid cartoon, these pieces are grouped to indicate the multiple repeating units (RBG domains) forming the rod, and the boundaries of each annotated region are indicated. Bottom, indicates the disordered regions that are part of each domain in a lighter shade of color. **b-d**, Details of AlphaFold3 predicted PfVPS13L1 structure. PfVPS13L1 is composed of: 12 RBG repeats (b), which have a hydrophobic face to accommodate the hydrophobic tails of lipids; a VPS13 Adapter Binding (VAB) domain formed by 6 β-strand repeats (c) that sticks out of the RBG-formed rod between RBG10 and RBG11; a UBA domain that is attached to the first repeat in the VAB domain (c); and a tandem of ATG2-C and PH domains found at the C-terminal end of the rod (d). A pair of helices stick out of the third repeat of the VAB domain, a feature not observed in any other eukaryotic VPS13 (c). The surface representation in (b) is colored by element (Oxygens are red, Nitrogens are blue and Carbons white), the ribbon representations in (c) and (d) are colored by AlphaFold3 confidence with the indicated domains colored according to the annotated cartoon in (a). **e**, Heliquest^105^ predictions of the hydrophobic moment of three helices of the ATG2-C domain confirming their amphipathicity, which allows the C-terminal end of the rod to closely interact with lipid bilayers. **f**, AlphaFold3 prediction of the interaction between the N-terminal region (residues 1-2976) of PfVPS13L1 (colored by N-to C-terminus) and PfVAP. The MSP domain of PfVAP is predicted to bind the FFAT motif tested in Fig. 2g-h (inset), which would anchor the N-terminal end of the rod to the ER.

**Extended Data Figure 6.**
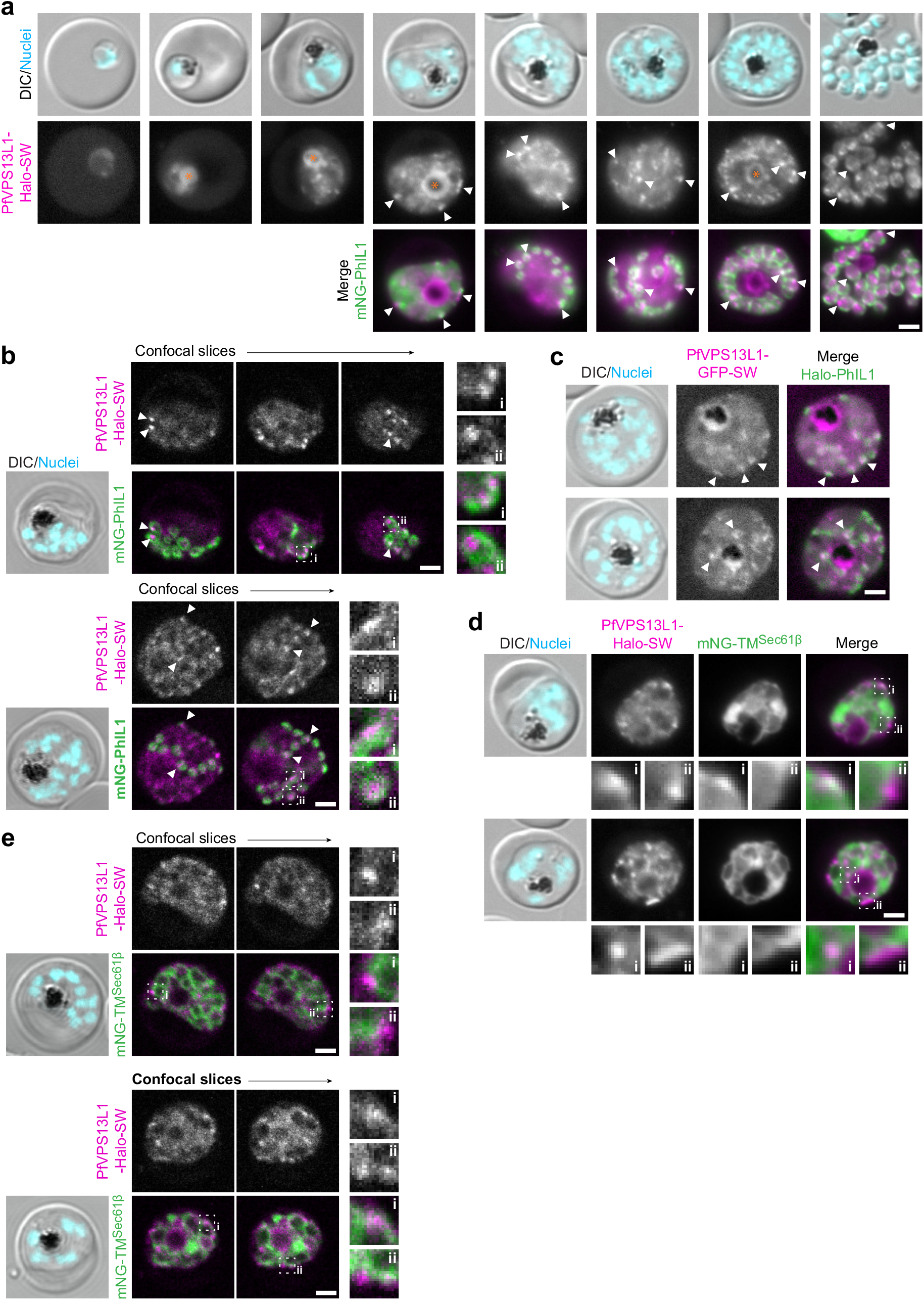
PfVPS13L1 colocalizes with the IMC during its early stages of formation. **a**, Representative fluorescence microscopy images of parasites expressing PfVPS13L1-Halo-SW from the endogenous locus (Extended Data Fig. 4b) and episomally expressing mNeon tagged PhIL1 (mNG-PhIL1) as an IMC marker. Little to no expression is observed in rings and trophozoites (orange asterisk indicates food vacuole background typical for high exposure images), whereas foci that colocalize with the IMC marker are observed in schizont stages (arrowheads). The protein remains in foci even after completion of IMC formation. **b**, Representative confocal microscopy images [using the cell line in (a)] showing PfVPS13L1 foci localize to a subsection of the IMC or directly adjacent to it. **c**, Representative fluorescence microscopy images of parasites expressing PfVPS13L1-GFP-SW from the endogenous locus (Extended Data Fig. 4b), which is in foci that colocalize with the episomally expressed IMC marker Halo-PhIL1 in early stages of IMC formation (arrowheads). **d-e**, Endogenous PfVPS13L1-Halo-SW hotspots are also colocalizing with or directly adjacent to the ER marker mNG-TM^Sec61β^, as seen by fluorescence (d) or confocal (e) microscopy. 3x enlargements of two boxed regions (i and ii in the merged slices) are shown to the right or at the bottom of each panel in (b-e). DIC, differential interference contrast; Nuclei, Hoechst 33342; scale bars, 2 μm.

**Extended Data Figure 7.**
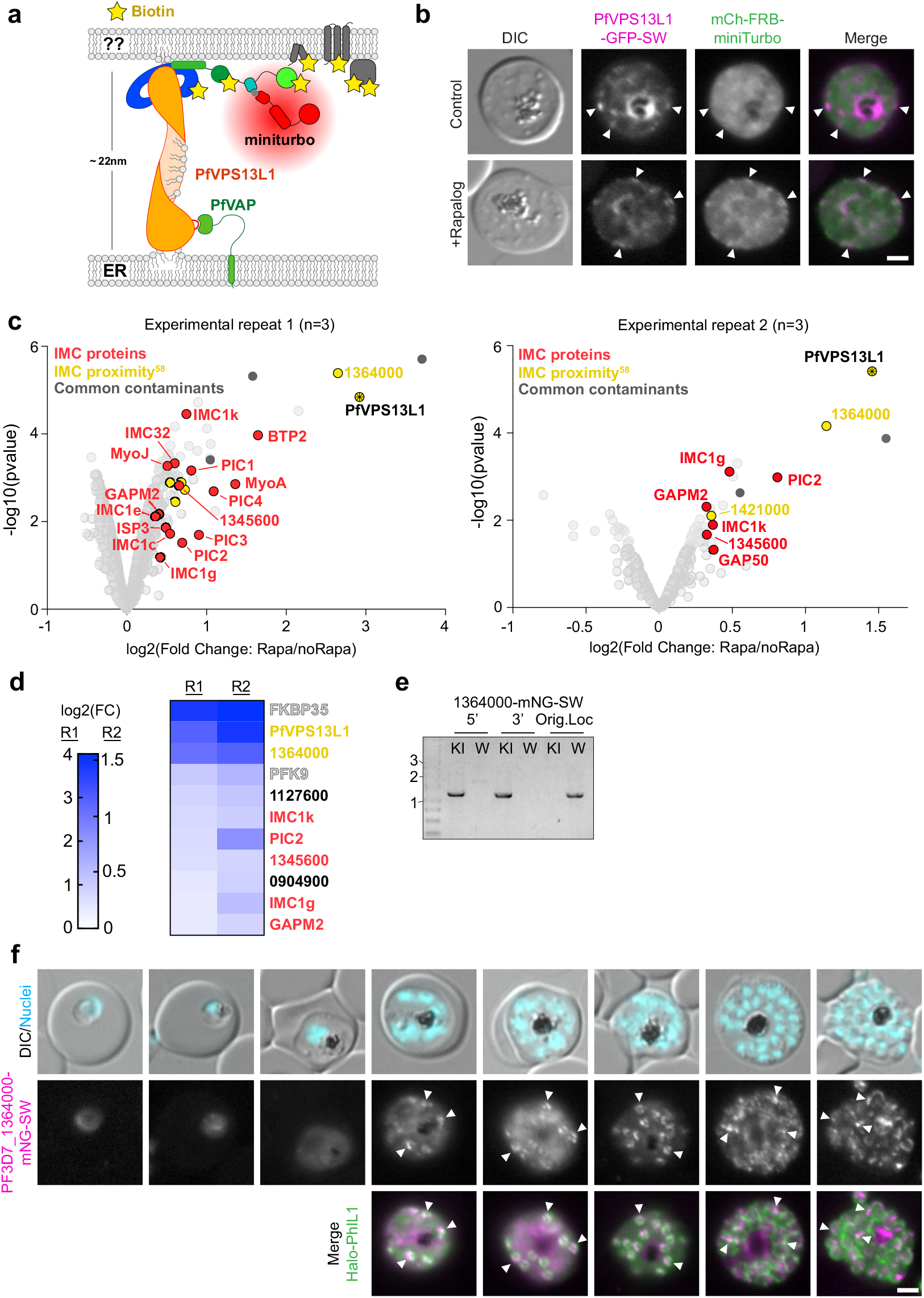
The C-terminal end of the PfVPS13L1 bridge interacts with the IMC. **a**, Schematic of PfVPS13L1 DiQ-BioID experiment. The labeling radius of the miniTurbo biotinylizer is smaller than the length of the PfVPS13L1 bridge (∼10 nm^106^), which is expected to allow for selective labelling of proteins in proximity to the C-terminal end of PfVPS13L1. **b**, Representative fluorescence microscopy images of parasites expressing PfVPS13L1-GFP-SW from the endogenous locus (Extended Data Fig. 4b) and episomally expressing the mCh-FRB-miniTurbo biotinilyzer, showing the rapalog-mediated recruitment of the biotinilyzer to the C-terminus of PfVPS13L1 as outlined in (a). Arrowheads indicate PfVPS13L1-GFP-SW foci. **c**, Individual LC-MS/MS experiments (each with 3 independent replicates) of C-terminal PfVPS13L1 DiQ-BioID. The left panel is the same experiment as shown in Fig. 4g. Proteins colored as indicated when meeting the cut offs (enrichment of at least 25% with a p value of at least 0.07). Moderated t-test was applied as implemented in the limma package. **d**, Heatmap with the 11 proteins found to pass the thresholds in (c) in both experimental repeats (R1 and R2) from (c). IMC proteins color coded as in c. FKBP35 and PFK9 are common BioID contaminants. **e**, Agarose gels showing PCR products amplified from genomic DNA of the indicated cell lines confirming correct integration of genome-modified parasites. Features as in Extended data Fig. 1b. **f**, Representative fluorescence microscopy images of the parasite line with endogenously tagged PF3D7_1364000-mNG-SW (e) and episomally expressing the IMC marker Halo-PhIL1 along the intraerythrocytic cycle. Colocalization with a subsection of the IMC is observed since early stages of its formation, and the protein remains in foci even after completion of IMC formation, similar to what was observed for PfVPS13L1 (Extended data Fig. 6a). Arrowheads indicate PF3D7_1364000 foci. DIC, differential interference contrast; Nuclei, Hoechst 33342; scale bars, 2 μm.

**Extended Data Figure 8.**
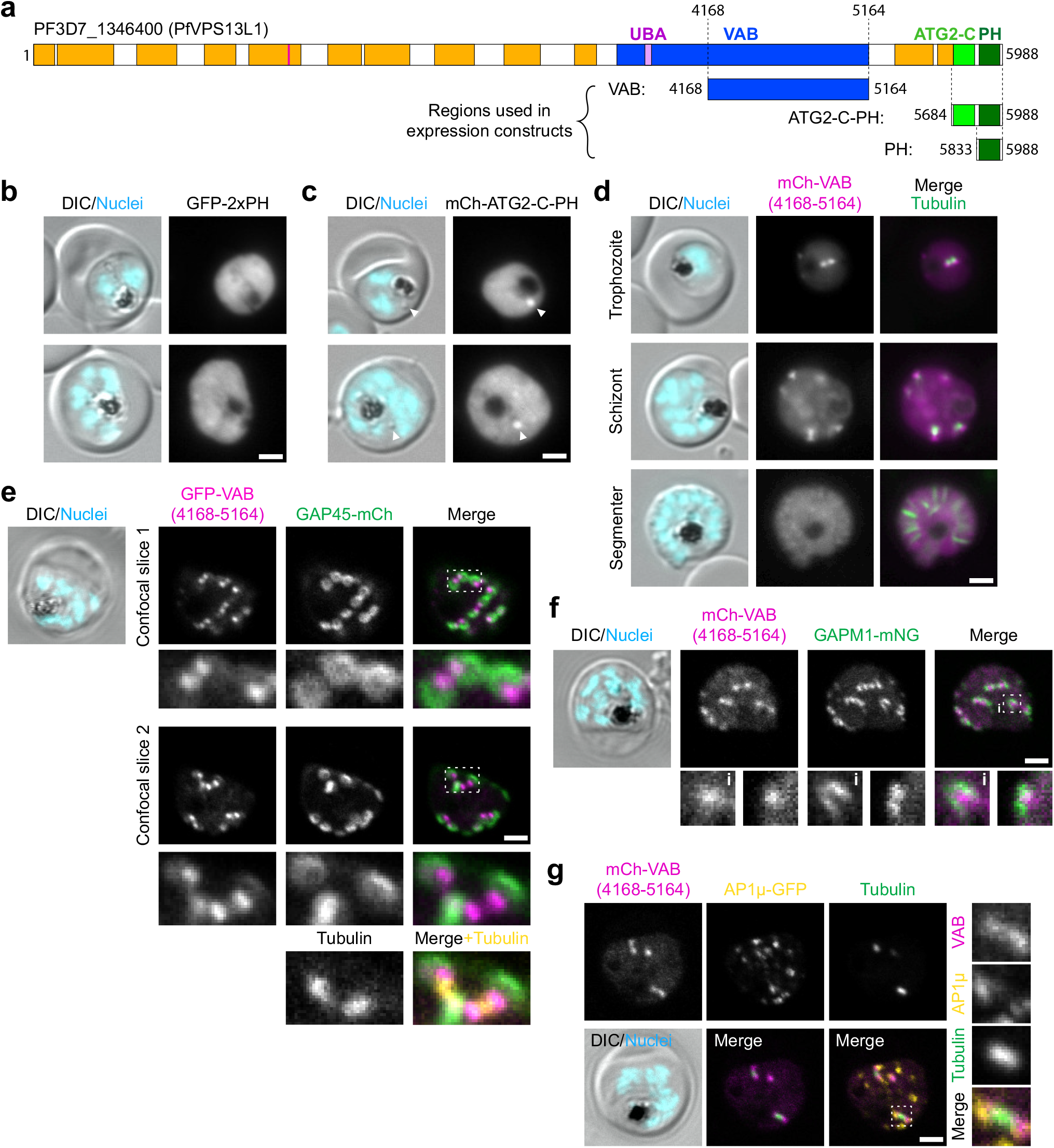
The C-terminal VAB domain of PfVPS13L1 regulates its localization to the IMC. **a**, PfVPS13L1, full length and truncations, domain cartoons. **b-d**, Representative fluorescence microscopy images showing the localization of episomally expressed C-terminal domains of PfVPS13L1 indicated in (a). The parasites expressing the VAB construct were stained with tubulin tracker (d). **e-f**, Confocal images of episomally expressed GFP-(e) or mCh-(f) tagged VAB showing partial colocalization with or a localization adjacent to the IMC markers GAP45-mCh (e) or GAPM1-mNG (f). The VAB hotspot is also adjacent to the tubulin tracker-stained inner centriolar plaque. **g**, Confocal images of tubulin tracker-stained parasites episomally expressing mCh-VAB and the Golgi apparatus as seen by AP-1μ-GFP (endo). Images in (g) are a maximum intensity Z-projection of two confocal slices. 3x enlargements of boxed regions in the merged slices are shown at the bottom or to the right of each panel in (e-g). The second enlargement in (f) is taken from a different confocal slice. DIC, differential interference contrast; Nuclei, Hoechst 33342; scale bars, 2 μm.

**Extended Data Figure 9.**
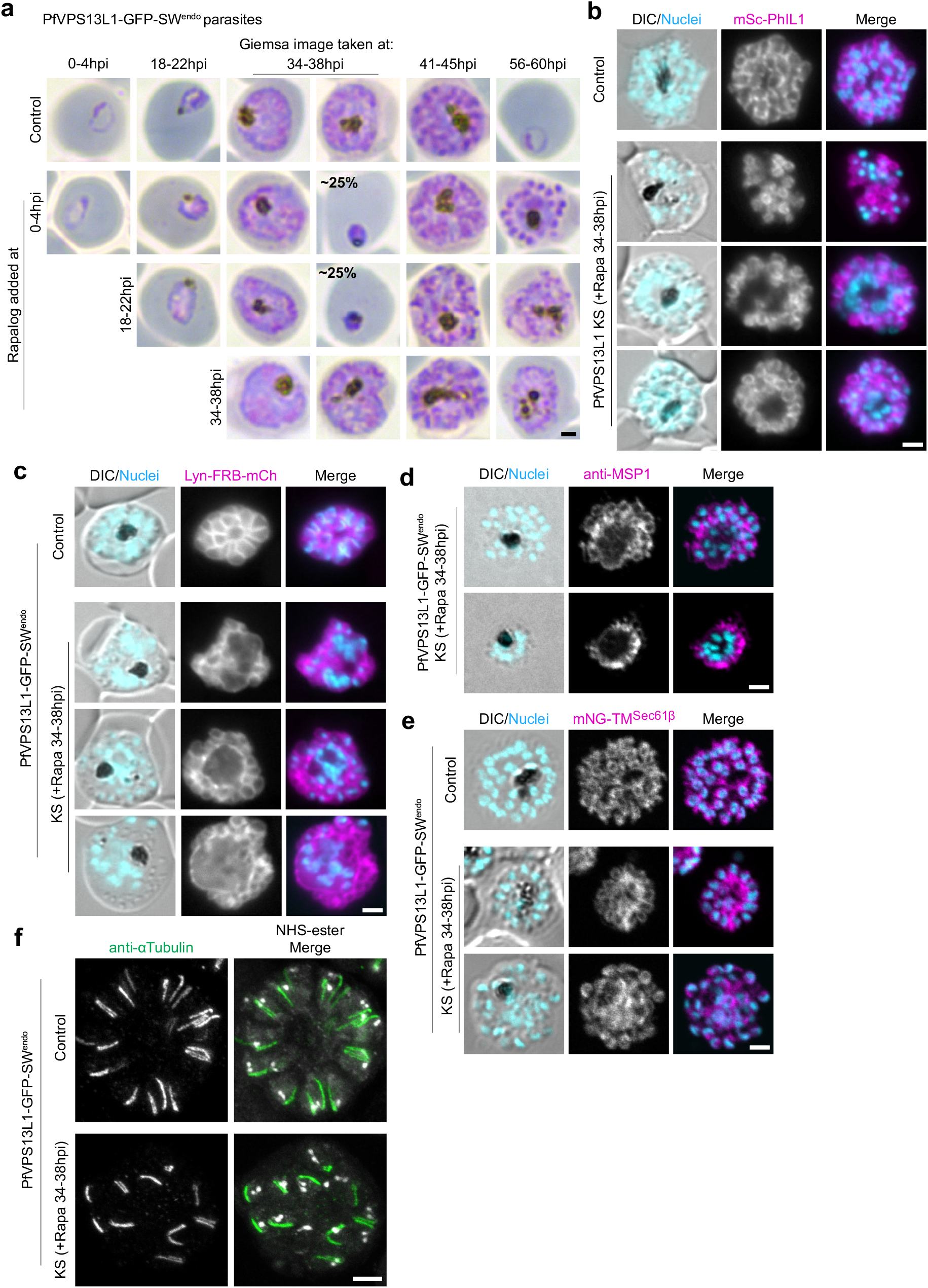
PfVPS13L1 loss-of-function leads to severe segmentation defects. **a**, Representative example microscopy images of the Giemsa smears of PfVPS13L1-GFP-SW^endo^ control and KS parasites (induced at different time points through addition of rapalog) from the stage quantification experiment in Fig. 5a. **b-c**, Fluorescence microscopy images of control (same parasites without rapalog) and late (34-38hpi) induced PfVPS13L1 KS schizonts arrested before egress with compound 2 and episomally expressing the IMC marker PhIL1 (b) or the PM marker Lyn-FRB-mCh (C). **d**, Representative confocal microscopy images of late (34-38hpi)-induced PfVPS13L1 KS compound 2-arrested schizonts immunostained with anti-MSP1, an antigen in the PM. **e**, Representative confocal microscopy images of PfVPS13L1-Halo-SW^endo^ control and late (34-38hpi) induced KS compound 2-arrested schizonts episomally expressing the ER marker mNG-TM^Sec61β^. **f**, U-ExM images of control and late-induced PfVPS13L1 KS compound 2-arrested schizonts immunostained with anti-αTubulin showing no noticeable differences in tubulin appearing as extended structures in segmented schizonts. U-ExM images are maximum intensity projections of Z-slices, with 4 (control) and 33 (KS) slices. DIC, differential interference contrast; Nuclei, Hoechst 33342; scale bars, 2 μm in (a-e) and 5 μm in (f).

