## Supplementary File 6 for "A bridge-like lipid transfer protein is critical for generation of invasive stages in malaria parasites"

**Original and modified genomic sequences**

Shown for each gene:

- The unedited genomic sequence.
- The modified genomic sequence after integration.
- The ATG is highlighted in green.
- Introns are in italics.
- The exons are shown in upper case.
- The homology region is shown in bold letters.
- The gene specific primers used for amplification of the original locus are highlighted in yellow.
- The common primers used for amplification of 5’ and 3’ junctions are highlighted in cyan.
- Other gene specific color labels are indicated for each sequence.
- Predicted size of PCR products are at the end of each genomic sequence.

For the case of PfVAP, the additional primers used to confirm correct excision are highlighted in red.

**Index:**

- **PF3D7_1439800 (PfVAP)	2-15**
- **PF3D7_1414500	16-26**
- **PF3D7_1316600	27-32**
- **PF3D7_1458500	33-39**
- **PF3D7_1461100	40-45**
- **PF3D7_1328300	46-49**
- **PF3D7_1311600	50-53**
- **PF3D7_1351000 (PfPITP)	54-60**
- **PF3D7_1131800 (PfOSBP)	61-66**
- **PF3D7_1346400 (PfVPS13L1)	67-84**
- **PF3D7_1343800 (PfVPS13L2)	85-98**
- **PF3D7_1324300 (PfVPS13L6)	99-110**
- **PF3D7_1364000	111-114**

### **> PF3D7_1439800 (PfVAP)**

tgaaggaaaaaaagaaaaaaaaaaggaaaaaaaaaaaaaaaaaaaaattaataataatatataatattaataatattttatatatttatatatttttcttacatatataatatatatatatatatataataataataataataataataatatatatgataaacttttattatttagtgaattttatataatttttactttatgttatataaataaatataaataaaaaataagtatatatatattttatatgtatataatatataaacaatgtgtttaatgtttttttaataaaaagatagtatttttatgaggcatatattattatatcttatattttattatatatatatatatatatttttttttttttttttttaatatcattgatcaacattgaattattttaaaaagaaaataaaaagggaaaaaaaaaaaaagaaaaaaaaaaagaataaaaaaagaagaaaaaaaagaataaaaaaagaaaaaaaaaaaagaataaaaaaagaaaaaaaagataaaactttcctaaatttttctcaaaaaaagaagaaaaaggattaaataaataattaaggagaatagtaagttaaatataagttatatataaacatataaggattaccaaaagggaaaatatacaactataaatattataatatcaaaaggataattatttaattaaataaaaaaaaaaaaaaaaaatagaatatatatatatttatatatataaaaaaattataaaaATG**AAACTTTTAAGAGTAACACCTGAAAAAAATATAGAATTTCCTCTTGTTCATTTTCAAGCAGTAACTCAAGTTGTTAAATTAGAAAATGTAAGTGATAAAAAAGTAGCTTTTAAAATAAAAACAACCGCTCCTAATAATTATTTAGTAAGACCATCATTTGGTTTAATAAGTGTAAGAGAAACAATAGAAATACAAATTATATTACAACCCTTGTCAGACAAAGATAATATATCAAACGATAAATTTCAGGTACAATGTTTAAATGTTGATGATAATACTACAGTAGATAAACAATTTTGGATAACAGTTAATAAAAATGAAATACAAGATCATAAACTTATTGTAGTTCTAAACGATGAAAATAATAGTAAATTAAATCATTCTTACATACCCTCAAATAATGTACCTCTCTCAGAAATGAATAACAAAAATATACACAATATGGGATACGTCGATAATAATAATATAAATCAAGATGACCCAAATTTAGCAGATGgtaaaaaatattacaaagaaatttatatctaatagtatatattaatattgtaatatatgtgtaaatcttatatgatgaattgtcacatatatcgtgttcttatatatataatgtgtgtgtgtactctacaagtgttcaagacaaaatgttatatatatatatatatgtatgtgtgtgtggatatatatatttatttattttatatatatatatatatatatattttttttttttttttttttttgttaagGTTTAAAAGGAGGTCTACCAGGTAT**GCAAAGGAAATATCATGAACTTTTAAATTATTGCGTTTTTGTTGATAAACAAAAAGCAGCCCTAGAAAAAGAAAACGAAAGTTTAAAAAATCAGTTAAAAGCATATAACAGTAATTCTAATAAATTCTTAATAGATAATAAATTAATTCCTATTATAATTGTAATGTTAGCTATAATAACAAAATATATGGGTTACTGGTAAtttcaaagttatacaaagaaaatatattaatatatatgtttacatttttgtgtaaggaaaatatcaagaatcaataaattataaaatagtaaattcgataagttataaaatgaaatatatgataaattataaaaatgattaataaaaaaaaaacccaaaaaaaaaaaaaaaaaaaaaaaaaagagagaagaaaacacatattaatatatatatatatatatatatttctatgtatacaaaggaatattaacattacaccagtattattaaataaatattaatatccaacaatatattattaagtacaattatacctattgaacagtattaaaagcaaaaacaaggaaatttatatatatatatatttatttatttatatgtatataaatacattctttatctatattgtatttagtttttaataattttttttatttttctcttccttactttttaatatgtttaaataattcataaacaagtagtaaaaaatttataatatatatatgaaatatgttatcaatatttaaaattgaagaaaat

**Original Locus (OL) amplicon size: 974bp**

**> GFP-2xFKBP-PfVAP yDHODH GFP recodonPfVAP loxP**

tatgtatataatatataaacaatgtgtttaatgtttttttaataaaaagatagtatttttatgaggcatatattattatatcttatattttattatatatatatatatatatttttttttttttttttttaatatcattgatcaacattgaattattttaaaaagaaaataaaaagggaaaaaaaaaaaaagaaaaaaaaaaagaataaaaaaagaagaaaaaaaagaataaaaaaagaaaaaaaaaaaagaataaaaaaagaaaaaaaagataaaactttcctaaatttttctcaaaaa

aagaagaaaaaggattaaataaataattaaggagaatagtaagttaaatataagttatatataaacatataaggattaccaaaagggaaaatatacaactataaatattataatatcaaaaggataattatttaattaaataaaaaaaaaaaaaaaaaatagaatatatatatatttatatatataaaaaaattataaaaATG**AAACTTTTAAGAGTAACACCTGAAAAAAATATAGAATTTCCTCTTGTTCATTTTCAAGCAGTAACTCAAGTTGTTAAATTAGAAAATGTAAGTGATAAAAAAGTAGCTTTTAAAATAAAAACAACCGCTCCTAATAATTATTTAGTAAGACCATCATTTGGTTTAATAAGTGTAAGAGAAACAATAGAAATACAAATTATATTACAACCCTTGTCAGACAAAGATAATATATCAAACGATAAATTTCAGGTACAATGTTTAAATGTTGATGATAATACTACAGTAGATAAACAATTTTGGATAACAGTTAATAAAAATGAAATACAAGATCATAAACTTATTGTAGTTCTAAACGATGAAAATAATAGTAAATTAAATCATTCTTACATACCCTCAAATAATGTACCTCTCTCAGAAATGAATAACAAAAATATACACAATATGGGATACGTCGATAATAATAATATAAATCAAGATGACCCAAATTTAGCAGATG*gtaaaaaatattacaaagaaatttatatctaatagtatatattaatattgtaatatatgtgtaaatcttatatgatgaattgtcacatatatcgtgttcttatatatataatgtgtgtgtgtactctacaagtgttcaagacaaaatgttatatatatatatatatgtatgtgtgtgtggatatatatatttatttattttatatatatatatatatatatattttttttttttttttttttttgttaag*GTTTAAAAGGAGGTCTACCAGGTAT**GCGTTTAAACGAGCAGAAGTTAATATCAGAAGAGGATTTGGGTGAACAAAAACTCATAAGCGAAGAAGATTTAATAACTTCGTATAGCATACATTATACGAAGTTATCCGGAGAAGGAAGAGGAAGTTTATTAACATGTGGAGATGTAGAAGAAAATCCAGGACCAATGACAGCCAGTTTAACTACCAAGTTCTTGAACAATACCTATGAAAACCCATTTATGAATGCATCCGGTGTTCATTGCATGACTACACAAGAATTAGATGAATTAGCAAACTCTAAAGCTGGCGCATTCATTACAAAGAGTGCTACAACCTTAGAAAGAGAAGGTAACCCTGAACCACGTTACATTTCTGTCCCTCTAGGCAGTATCAACTCCATGGGTTTACCAAACGAAGGTATCGACTACTATTTGTCCTATGTATTAAACCGTCAAAAGAATTATCCTGATGCACCTGCTATTTTCTTCTCAGTTGCTGGTATGAGCATTGATGAAAATTTAAATTTGTTGAGGAAAATCCAAGATAGCGAATTCAACGGTATTACCGAGTTAAACTTGTCTTGTCCTAATGTGCCTGGGAAACCACAAGTTGCTTATGACTTTGACTTGACAAAGGAAACCTTGGAAAAGGTTTTTGCCTTTTTCAAAAAACCTCTTGGTGTCAAGTTGCCTCCTTATTTTGATTTTGCCCATTTTGATATCATGGCAAAAATATTGAACGAGTTCCCATTAGCTTATGTCAACTCTATCAATAGTATAGGAAATGGTCTTTTCATTGATGTGGAGAAGGAGAGTGTAGTAGTGAAGCCAAAGAATGGTTTCGGGGGTATTGGAGGTGAATATGTTAAGCCAACCGCGCTCGCCAATGTTCGTGCATTTTACACTCGTTTGAGACCTGAAATCAAAGTTATCGGTACAGGTGGAATTAAGTCCGGTAAGGATGCATTTGAACATCTTCTATGTGGTGCCTCTATGCTACAGATTGGTACAGAATTACAAAAAGAGGGCGTCAAGATTTTTGAACGTATCGAAAAAGAATTAAAAGACATAATGGAAGCTAAGGGTTATACATCCATAGATCAGTTCCGTGGGAAGTTGAACAGCATTGGTGAAGGTAGAGGTTCTTTGTTGACTTGTGGTGATGTTGAAGAAAATCCAGGTCCAGCTAGCATGAGTAAAGGAGAAGAACTTTTCACTGGAGTTGTCCCAATTCTTGTTGAATTAGATGGTGATGTTAATGGGCACAAATTTTCTGTCAGTGGAGAGGGTGAAGGTGATGCAACATACGGAAAACTTACCCTTAAATTTATTTGCACTACTGGAAAACTACCAGTTCCATGGCCAACACTTGTCACTACTTTCGCGTATGGTCTTCAATGCTTTGCGAGATACCCAGATCATATGAAACAGCATGACTTTTTCAAGAGTGCCATGCCCGAAGGTTATGTACAGGAAAGAACTATATTTTTCAAAGATGACGGGAACTACAAGACACGTGCTGAAGTCAAGTTTGAAGGTGATACCCTTGTTAATAGAATCGAGTTAAAAGGTATTGATTTTAAAGAAGATGGAAACATTCTTGGACACAAATTGGAATACAACTATAACTCACACAATGTATACATCATGGCAGACAAACAAAAGAATGGAATCAAAGTTAACTTCAAAATTAGACACAACATTGAAGATGGAAGCGTTCAACTAGCAGACCATTATCAACAAAATACTCCAATTGGCGATGGCCCTGTCCTTTTACCAGACAACCATTACCTGTCCACACAATCTGCCCTTTCGAAAGATCCCAACGAAAAGAGAGACCACATGGTCCTTCTTGAGTTTGTAACAGCTGCTGGGATTACACATGGCATGGATGAGCTCTACAAACCTAGCTCAGGATTGAGATCAAGATCTGCTGCTGCTGGTGCTGGTGGTGCTGCTAGAGCTGCTCTGCAGAGAGGAGTACAAGTTGAAACAATATCACCAGGAGATGGTCGTACATTTCCAAAAAGAGGTCAAACTTGTGTTGTACATTATACTGGAATGCTTGAAGATGGAAAGAAATTTGATTCATCTCGTGATAGAAATAAACCATTTAAATTTATGCTAGGTAAACAAGAAGTAATACGAGGTTGGGAAGAAGGAGTTGCTCAAATGAGTGTAGGTCAAAGAGCAAAACTTACTATATCTCCAGATTATGCTTATGGTGCAACTGGACATCCAGGTATAATTCCACCTCATGCAACTCTTGTATTTGATGTGGAGCTTCTAAAACTAGAAACTAGAGGTGTTCAGGTTGAAACAATTTCACCTGGAGATGGCAGAACCTTTCCTAAAAGAGGACAGACTTGCGTAGTTCATTATACAGGCATGCTAGAGGATGGTAAGAAATTTGATTCTAGTCGAGATAGAAATAAGCCATTCAAGTTTATGCTAGGTAAACAGGAAGTAATAAGAGGTTGGGAAGAGGGTGTAGCACAGATGTCAGTTGGACAAAGAGCAAAGTTAACAATATCACCAGATTATGCATACGGTGCAACAGGCCATCCTGGCATCATCCCTCCACATGCAACTTTAGTATTCGACGTTGAATTGTTAAAGTTAGAGACAACGCGTGCTAGAGGTGCTGCTGCTGGTGCTGGAGGTGCAGGTAGACCTAGGATGAAGTTACTTCGTGTTACTCCAGAGAAGAACATTGAGTTCCCATTAGTACACTTCCAGGCTGTTACACAGGTAGTAAAGCTTGAGAACGTTTCAGACAAGAAGGTTGCATTCAAGATTAAGACTACAGCACCAAACAACTACCTTGTTCGTCCTAGTTTCGGACTTATTTCAGTTCGTGAGACTATTGAGATTCAGATAATTCTTCAGCCACTTAGTGATAAGGACAACATTAGTAATGACAAGTTCCAAGTTCAGTGCCTTAACGTAGACGACAACACAACTGTTGACAAGCAGTTCTGGATTACTGTAAACAAGAACGAGATTCAGGACCACAAGTTAATAGTTGTATTGAATGACGAGAACAACTCAAAGCTTAACCACAGCTATATTCCAAGTAACAACGTTCCATTAAGTGAGATGAACAATAAGAACATTCATAACATGGGTTATGTTGACAACAACAACATTAACCAGGACGATCCTAACCTTGCTGACGGACTTAAGGGTGGATTGCCTGGAATGCAGCGTAAGTACCACGAGTTACTTAACTACTGTGTATTCGTAGACAAGCAGAAGGCTGCATTGGAGAAGGAGAATGAGTCACTTAAGAACCAACTTAAGGCTTACAATTCAAACAGCAACAAGTTTCTTATTGACAACAAGCTTATACCAATAATTATAGTTATGCTTGCAATTATTACTAAGTACATGGGATATTGGTAGaggcctataacttcgtatagcatacattatacgaagttattatgactcgagggatatggcagcttaatgttcgtttttcttatttatatatttataccaattgattgtatttataactgtaaaaatgtgtatgttgtgtgcatatttttttttgtgcatgcacatgcatgtaaatagctaaaattatgaacattttattttttgttcagaaaaaaaaaactttacacacataaaatggctagtatgaatagccatattttatataaattaaatcctatgaatttatgaccatattaaaaatttagatatttatggaacataatatgtttgaaacaataagacaaaattattattattattattatttttactgttataattatgtgtctccttcaatgattcataaatagttggacttgatttttaaaatgtttataatatgattagcatagttaaataaaaaaagttgaaaaattaaaaaaaaacatataaacacaaatgatggtttttccttcaatttcgatatcaatttatagaaacaaaatatatacttgtataattttatttttttatataaatcattacatatataattatacaatattttttctaagagataattatatattaatatatataaaaaaaggtgttttttttttttttttttatttttatttttattttatggtaatattttattttccttattttataaattatattagtttatatgtgattaattttatatattatcaatttatatatttttaaatgcttacttaattatctttttttttttttttttttttttttcccctctttttatattaatttatttttgaaaaaattgatatatatatatatatataatatatatatatacatgtagtagtattaaacaatgtataatatatataaataatatatttatatatttcatttcaattttaattttttttggttttttttttttttctttttgtcatatttaaaaaaaattatattcatataagttatgcattttttataaacattattcaatatatgtataatataatatatatatatatattaatgtattattccaatgtgcatgataaaagaaaaaaataatatttataaaaaaaaagaaaaataaaacaaaaaaagaaaaaaaaaaaaaaaaaaaaaaaaatacaaaaataaataatataatttataattatatattcttgtcacaataaaaatatatatatatatatatatatttataatatgtatattttaaactagaaaaggaataactaatattttatttattatcattcaagatttatattttataataataaatacctaatagaaatatatcaggatccatgcatggttcgctaaactgcatcgtcgctgtgtcccagaacatgggcatcggcaagaacggggactacccctggccaccgctcaggaacgaatttagatatttccagagaatgaccacaacctcttcagtagaaggtaaacagaatctggtgattatgggtaagaagacctggttctccattcctgagaagaatcgacctttaaagggtagaattaatttagttctcagcagagaactcaaggaacctccacaaggagctcattttctttccagaagtctagatgatgccttaaaacttactgaacaaccagaattagcaaataaagtagacatggtctggatagttggtggcagttctgtttataaggaagccatgaatcacccaggccatcttaaactatttgtgacaaggatcatgcaagactttgaaagtgacacgttttttccagaaattgatttggagaaatataaacttctgccagaatacccaggtgttctctctgatgtccaggaggagaaaggcattaagtacaaatttgaagtatatgagaagaatgattaagcttatttaataatagattaaaaatattataaaaataaaaacataaacacagaaattacaaaaaaaatacatatgaattttttttttgtaatcttccttataaatatagaataatgaatcatataaaacatatcattattcatttatttacatttaaaattattgtttcagtatctttaatttattatgtatatataaaaataacttacaattttattaataaacaatatatgtttattaattcatgttttgtaatttatgggatagcgattttttttactgtctgtatttttcttttttaattatgttttaattgtattttatttttattattgttctttttatagtattattttaaaacaaaatgtattttctaagaacttataataataataaatataaattttaataaaaattatatttatcttttacaatatgaacataaagtacaacattaatatatagcttttaatatttttattcctaatcatgtaaatcttaaatttttctttttaaacatatgttaaatatttatttctcattatatataagaacatatttattaaatctagaattctatagtgagtcgtattacaattcactggccgtcgttttacaacgtcgtgactgggaaaaccctggcgttacccaacttaatcgccttgcagcacatccccctttcgccagctggcgtaatagcgaagaggcccgcaccgatcgcccttcccaacagttgcgcagcctgaatggcgaatggcgcctgatgcggtattttctccttacgcatctgtgcggtatttcacaccgcatatggtgcactctcagtacaatctgctctgatgccgcatagttaagccagccccgacacccgccaacacccgctgacgcgccctgacgggcttgtctgctcccggcatccgcttacagacaagctgtgaccgtctccgggagctgcatgtgtcagaggttttcaccgtcatcaccgaaacgcgcgagacgaaagggcctcgtgatacgcctatttttataggttaatgtcatgataataatggtttcttagacgtcaggtggcacttttcggggaaatgtgcgcggaacccctatttgtttatttttctaaatacattcaaatatgtatccgctcatgagacaataaccctgataaatgcttcaataatattgaaaaaggaagagtatgagtattcaacatttccgtgtcgcccttattcccttttttgcggcattttgccttcctgtttttgctcacccagaaacgctggtgaaagtaaaagatgctgaagatcagttgggtgcacgagtgggttacatcgaactggatctcaacagcggtaagatccttgagagttttcgccccgaagaacgttttccaatgatgagcacttttaaagttctgctatgtggcgcggtattatcccgtattgacgccgggcaagagcaactcggtcgccgcatacactattctcagaatgacttggttgagtactcaccagtcacagaaaagcatcttacggatggcatgacagtaagagaattatgcagtgctgccataaccatgagtgataacactgcggccaacttacttctgacaacgatcggaggaccgaaggagctaaccgcttttttgcacaacatgggggatcatgtaactcgccttgatcgttgggaaccggagctgaatgaagccataccaaacgacgagcgtgacaccacgatgcctgtagcaatgccaacaacgttgcgcaaactattaactggcgaactacttactctagcttcccggcaacaattaatagactggatggaggcggataaagttgcaggaccacttctgcgctcggcccttccggctggctggtttattgctgataaatctggagccggtgagcgtgggtctcgcggtatcattgcagcactggggccagatggtaagccctcccgtatcgtagttatctacacgacggggagtcaggcaactatggatgaacgaaatagacagatcgctgagataggtgcctcactgattaagcattggtaactgtcagaccaagtttactcatatatactttagattgatttaaaacttcatttttaatttaaaaggatctaggtgaagatcctttttgataatctcatgaccaaaatcccttaacgtgagttttcgttccactgagcgtcagaccccgtagaaaagatcaaaggatcttcttgagatcctttttttctgcgcgtaatctgctgcttgcaaacaaaaaaaccaccgctaccagcggtggtttgtttgccggatcaagagctaccaactctttttccgaaggtaactggcttcagcagagcgcagataccaaatactgtccttctagtgtagccgtagttaggccaccacttcaagaactctgtagcaccgcctacatacctcgctctgctaatcctgttaccagtggctgctgccagtggcgataagtcgtgtcttaccgggttggactcaagacgatagttaccggataaggcgcagcggtcgggctgaacggggggttcgtgcacacagcccagcttggagcgaacgacctacaccgaactgagatacctacagcgtgagctatgagaaagcgccacgcttcccgaagggagaaaggcggacaggtatccggtaagcggcagggtcggaacaggagagcgcacgagggagcttccagggggaaacgcctggtatctttatagtcctgtcgggtttcgccacctctgacttgagcgtcgatttttgtgatgctcgtcaggggggcggagcctatcgaaaaacgccagcaacgcggcctttttacggttcctggccttttgctggccttttgctcacatgttctttcctgcgttatcccctgattctgtggataaccgtattaccgcctttgagtgagctgataccgctcgccgcagccgaacgaccgagcgcagcgagtcagtgagcgaggaagcggaagagcgcccaatacgcaaaccgcctctccccgcgcgttggccgattcattaatgcagctggcacgacaggtttcccgactggaaagcgggcagtgagcgcaacgcaattaatgtgagttagctcactcattaggcaccccaggctttacactttatgcttccggctcgtatgttgtgtggaattgtgagcggataacaatttcacacaggaaacagctatgaccatgattacgccaagctatttaggtgacactatagaatactcgcggccgcTAGaaacttttaagagtaacacctgaaaaaaatatagaatttcctcttgttcattttcaagcagtaactcaagttgttaaattagaaaatgtaagtgataaaaaagtagcttttaaaataaaaacaaccgctcctaataattatttagtaagaccatcatttggtttaataagtgtaagagaaacaatagaaatacaaattatattacaacccttgtcagacaaagataatatatcaaacgataaatttcaggtacaatgtttaaatgttgatgataatactacagtagataaacaattttggataacagttaataaaaatgaaatacaagatcataaacttattgtagttctaaacgatgaaaataatagtaaattaaatcattcttacataccctcaaataatgtacctctctcagaaatgaataacaaaaatatacacaatatgggatacgtcgataataataatataaatcaagatgacccaaatttagcagatggtaaaaaatattacaaagaaatttatatctaatagtatatattaatattgtaatatatgtgtaaatcttatatgatgaattgtcacatatatcgtgttcttatatatataatgtgtgtgtgtactctacaagtgttcaagacaaaatgttatatatatatatatatgtatgtgtgtgtggatatatatatttatttattttatatatatatatatatatatattttttttttttttttttttttgttaaggtttaaaaggaggtctaccaggtatgcaaaggaaatatcatgaacttttaaattattgcgtttttgttgataaacaaaaagcagccctagaaaaagaaaacgaaagtttaaaaaatcagttaaaagcatataacagtaattctaataaattcttaatagataataaattaattcctattataattgtaatgttagctataataacaaaatatatgggttactggtaatttcaaagttatacaaagaaaatatattaatatatatgtttacatttttgtgtaaggaaaatatcaagaatcaataaattataaaatagtaaattcgataagttataaaatgaaatatatgataaattataaaaatgattaataaaaaaaaaacccaaaaaaaaaaaaaaaaaaaaaaaaaagagagaagaaaacacatattaatatatatatatatatatatatttctatgtatacaaaggaatattaacattacaccagtattattaaataaatattaatatccaacaatatattattaagtacaattatacctattgaacagtattaaaagcaaaaacaaggaaatttatatatatatatatttatttatttatatgtatataaatacattctttatctatattgtatttagtttttaataattttttttatttttctcttccttactttttaatatgtttaaataattcataaacaagtagtaaaaaatttataat

**5’ amplicon size: 2352bps**

**3’ amplicon size: 934bps**

**> GFP-2xFKBP-PfVAP (after excision) yDHODH recodonPfVAP loxP**

tatgtatataatatataaacaatgtgtttaatgtttttttaataaaaagatagtatttttatgaggcatatattattatatcttatattttattatatatatatatatatatttttttttttttttttttaatatcattgatcaacattgaattattttaaaaagaaaataaaaagggaaaaaaaaaaaaagaaaaaaaaaaagaataaaaaaagaagaaaaaaaagaataaaaaaagaaaaaaaaaaaagaataaaaaaagaaaaaaaagataaaactttcctaaatttttctcaaaaa

aagaagaaaaaggattaaataaataattaaggagaatagtaagttaaatataagttatatataaacatataaggattaccaaaagggaaaatatacaactataaatattataatatcaaaaggataattatttaattaaataaaaaaaaaaaaaaaaaatagaatatatatatatttatatatataaaaaaattataaaaATG**AAACTTTTAAGAGTAACACCTGAAAAAAATATAGAATTTCCTCTTGTTCATTTTCAAGCAGTAACTCAAGTTGTTAAATTAGAAAATGTAAGTGATA**

**AAAAAGTAGCTTTTAAAATAAAAACAACCGCTCCTAATAATTATTTAGTAAGACCATCATTTGGTTTAATAAGTGTAAGAGAAACAATAGAAATACAAATTATATTACAACCCTTGTCAGACAAAGATAATATATCAAACGATAAATTTCAGGTACAATGTTTAAATGTTGATGATAATACTACAGTAGATAAACAATTTTGGATAACAGTTAATAAAAATGAAATACAAGATCATAAACTTATTGTAGTTCTAAACGATGAAAATAATAGTAAATTAAATCATTCTTACATACCCTCAA**

**ATAATGTACCTCTCTCAGAAATGAATAACAAAAATATACACAATATGGGATACGTCGATAATAATAATATAAATCAAGATGACCCAAATTTAGCAGATGgtaaaaaatattacaaagaaatttatatctaatagtatatattaatattgtaatatatgtgtaaatcttatatgatgaattgtcacatatatcgtgttcttatatatataatgtgtgtgtgtactctacaagtgttcaagacaaaatgttatatatatatatatatgtatgtgtgtgtggatatatatatttatttatttt**

**atatatatatatatatatatattttttttttttttttttttttgttaagGTTTAAAAGGAGGTCTACCAGGTAT**GCGTTTAAACGAGCAGAAGTTAATATCAGAAGAGGATTTGGGTGAACAAAAACTCATAAGCGAAGAAGATTTAATAACTTCGTATAGCATACATTATACGAAGTTATtatgactcgagggatatggcagcttaatgttcgtttttcttatttatatatttataccaattgattgtatttataactgtaaaaatgtgtatgttgtgtgcatatttttttttgtgcatgcacatgcatgtaaatagctaaaattatgaacattttattttttgttcagaaaaaaaaaactttacacacataaaatggctagtatgaatagccatattttatataaattaaatcctatgaatttatgaccatattaaaaatttagatatttatggaacataatatgtttgaaacaataagacaaaattattattattattattatttttactgttataattatgtgtctccttcaatgattcataaatagttggacttgatttttaaaatgtttataatatgattagcatagttaaataaaaaaagttgaaaaattaaaaaaaaacatataaacacaaatgatggtttttccttcaatttcgatatcaatttatagaaacaaaatatatacttgtataattttatttttttatataaatcattacatatataattatacaatattttttctaagagataattatatattaatatatataaaaaaaggtgttttttttttttttttttatttttatttttattttatggtaatattttattttccttattttataaattatattagtttatatgtgattaattttatatattatcaatttatatatttttaaatgcttacttaattatctttttttttttttttttttttttttcccctctttttatattaatttatttttgaaaaaattgatatatatatatatatataatatatatatatacatgtagtagtattaaacaatgtataatatatataaataatatatttatatatttcatttcaattttaattttttttggttttttttttttttctttttgtcatatttaaaaaaaattatattcatataagttatgcattttttataaacattattcaatatatgtataatataatatatatatatatattaatgtattattccaatgtgcatgataaaagaaaaaaataatatttataaaaaaaaagaaaaataaaacaaaaaaagaaaaaaaaaaaaaaaaaaaaaaaaatacaaaaataaataatataatttataattatatattcttgtcacaataaaaatatatatatatatatatatatttataatatgtatattttaaactagaaaaggaataactaatattttatttattatcattcaagatttatattttataataataaatacctaatagaaatatatcaggatccatgcatggttcgctaaactgcatcgtcgctgtgtcccagaacatgggcatcggcaagaacggggactacccctggccaccgctcaggaacgaatttagatatttccagagaatgaccacaacctcttcagtagaaggtaaacagaatctggtgattatgggtaagaagacctggttctccattcctgagaagaatcgacctttaaagggtagaattaatttagttctcagcagagaactcaaggaacctccacaaggagctcattttctttccagaagtctagatgatgccttaaaacttactgaacaaccagaattagcaaataaagtagacatggtctggatagttggtggcagttctgtttataaggaagccatgaatcacccaggccatcttaaactatttgtgacaaggatcatgcaagactttgaaagtgacacgttttttccagaaattgatttggagaaatataaacttctgccagaatacccaggtgttctctctgatgtccaggaggagaaaggcattaagtacaaatttgaagtatatgagaagaatgattaagcttatttaataatagattaaaaatattataaaaataaaaacataaacacagaaattacaaaaaaaatacatatgaattttttttttgtaatcttccttataaatatagaataatgaatcatataaaacatatcattattcatttatttacatttaaaattattgtttcagtatctttaatttattatgtatatataaaaataacttacaattttattaataaacaatatatgtttattaattcatgttttgtaatttatgggatagcgattttttttactgtctgtatttttcttttttaattatgttttaattgtattttatttttattattgttctttttatagtattattttaaaacaaaatgtattttctaagaacttataataataataaatataaattttaataaaaattatatttatcttttacaatatgaacataaagtacaacattaatatatagcttttaatatttttattcctaatcatgtaaatcttaaatttttctttttaaacatatgttaaatatttatttctcattatatataagaacatatttattaaatctagaattctatagtgagtcgtattacaattcactggccgtcgttttacaacgtcgtgactgggaaaaccctggcgttacccaacttaatcgccttgcagcacatccccctttcgccagctggcgtaatagcgaagaggcccgcaccgatcgcccttcccaacagttgcgcagcctgaatggcgaatggcgcctgatgcggtattttctccttacgcatctgtgcggtatttcacaccgcatatggtgcactctcagtacaatctgctctgatgccgcatagttaagccagccccgacacccgccaacacccgctgacgcgccctgacgggcttgtctgctcccggcatccgcttacagacaagctgtgaccgtctccgggagctgcatgtgtcagaggttttcaccgtcatcaccgaaacgcgcgagacgaaagggcctcgtgatacgcctatttttataggttaatgtcatgataataatggtttcttagacgtcaggtggcacttttcggggaaatgtgcgcggaacccctatttgtttatttttctaaatacattcaaatatgtatccgctcatgagacaataaccctgataaatgcttcaataatattgaaaaaggaagagtatgagtattcaacatttccgtgtcgcccttattcccttttttgcggcattttgccttcctgtttttgctcacccagaaacgctggtgaaagtaaaagatgctgaagatcagttgggtgcacgagtgggttacatcgaactggatctcaacagcggtaagatccttgagagttttcgccccgaagaacgttttccaatgatgagcacttttaaagttctgctatgtggcgcggtattatcccgtattgacgccgggcaagagcaactcggtcgccgcatacactattctcagaatgacttggttgagtactcaccagtcacagaaaagcatcttacggatggcatgacagtaagagaattatgcagtgctgccataaccatgagtgataacactgcggccaacttacttctgacaacgatcggaggaccgaaggagctaaccgcttttttgcacaacatgggggatcatgtaactcgccttgatcgttgggaaccggagctgaatgaagccataccaaacgacgagcgtgacaccacgatgcctgtagcaatgccaacaacgttgcgcaaactattaactggcgaactacttactctagcttcccggcaacaattaatagactggatggaggcggataaagttgcaggaccacttctgcgctcggcccttccggctggctggtttattgctgataaatctggagccggtgagcgtgggtctcgcggtatcattgcagcactggggccagatggtaagccctcccgtatcgtagttatctacacgacggggagtcaggcaactatggatgaacgaaatagacagatcgctgagataggtgcctcactgattaagcattggtaactgtcagaccaagtttactcatatatactttagattgatttaaaacttcatttttaatttaaaaggatctaggtgaagatcctttttgataatctcatgaccaaaatcccttaacgtgagttttcgttccactgagcgtcagaccccgtagaaaagatcaaaggatcttcttgagatcctttttttctgcgcgtaatctgctgcttgcaaacaaaaaaaccaccgctaccagcggtggtttgtttgccggatcaagagctaccaactctttttccgaaggtaactggcttcagcagagcgcagataccaaatactgtccttctagtgtagccgtagttaggccaccacttcaagaactctgtagcaccgcctacatacctcgctctgctaatcctgttaccagtggctgctgccagtggcgataagtcgtgtcttaccgggttggactcaagacgatagttaccggataaggcgcagcggtcgggctgaacggggggttcgtgcacacagcccagcttggagcgaacgacctacaccgaactgagatacctacagcgtgagctatgagaaagcgccacgcttcccgaagggagaaaggcggacaggtatccggtaagcggcagggtcggaacaggagagcgcacgagggagcttccagggggaaacgcctggtatctttatagtcctgtcgggtttcgccacctctgacttgagcgtcgatttttgtgatgctcgtcaggggggcggagcctatcgaaaaacgccagcaacgcggcctttttacggttcctggccttttgctggccttttgctcacatgttctttcctgcgttatcccctgattctgtggataaccgtattaccgcctttgagtgagctgataccgctcgccgcagccgaacgaccgagcgcagcgagtcagtgagcgaggaagcggaagagcgcccaatacgcaaaccgcctctccccgcgcgttggccgattcattaatgcagctggcacgacaggtttcccgactggaaagcgggcagtgagcgcaacgcaattaatgtgagttagctcactcattaggcaccccaggctttacactttatgcttccggctcgtatgttgtgtggaattgtgagcggataacaatttcacacaggaaacagctatgaccatgattacgccaagctatttaggtgacactatagaatactcgcggccgctagaaacttttaagagtaacacctgaaaaaaatatagaatttcctcttgttcattttcaagcagtaactcaagttgttaaattagaaaatgtaagtgataaaaaagtagcttttaaaataaaaacaaccgctcctaataattatttagtaagaccatcatttggtttaataagtgtaagagaaacaatagaaatacaaattatattacaacccttgtcagacaaagataatatatcaaacgataaatttcaggtacaatgtttaaatgttgatgataatactacagtagataaacaattttggataacagttaataaaaatgaaatacaagatcataaacttattgtagttctaaacgatgaaaataatagtaaattaaatcattcttacataccctcaaataatgtacctctctcagaaatgaataacaaaaatatacacaatatgggatacgtcgataataataatataaatcaagatgacccaaatttagcagatggtaaaaaatattacaaagaaatttatatctaatagtatatattaatattgtaatatatgtgtaaatcttatatgatgaattgtcacatatatcgtgttcttatatatataatgtgtgtgtgtactctacaagtgttcaagacaaaatgttatatatatatatatatgtatgtgtgtgtggatatatatatttatttattttatatatatatatatatatatattttttttttttttttttttttgttaaggtttaaaaggaggtctaccaggtatgcaaaggaaatatcatgaacttttaaattattgcgtttttgttgataaacaaaaagcagccctagaaaaagaaaacgaaagtttaaaaaatcagttaaaagcatataacagtaattctaataaattcttaatagataataaattaattcctattataattgtaatgttagctataataacaaaatatatgggttactggtaatttcaaagttatacaaagaaaatatattaatatatatgtttacatttttgtgtaaggaaaatatcaagaatcaataaattataaaatagtaaattcgataagttataaaatgaaatatatgataaattataaaaatgattaataaaaaaaaaacccaaaaaaaaaaaaaaaaaaaaaaaaaagagagaagaaaacacatattaatatatatatatatatatatatttctatgtatacaaaggaatattaacattacaccagtattattaaataaatattaatatccaacaatatattattaagtacaattatacctattgaacagtattaaaagcaaaaacaaggaaatttatatatatatatatttatttatttatatgtatataaatacattctttatctatattgtatttagtttttaataattttttttatttttctcttccttactttttaatatgtttaaataattcataaacaagtagtaaaaaatttataat

**Unexcised amplicon size: 3494bps**

**Excised amplicon size: 176bps**

**> Halo-PfVAP yDHODH Halo recodonPfVAP**

tgaaggaaaaaaagaaaaaaaaaaggaaaaaaaaaaaaaaaaaaaaattaataataatatataatattaataatattttatatatttatatatttttcttacatatataatatatatatatatatataataataataataataataataatatatatgataaacttttattatttagtgaattttatataatttttactttatgttatataaataaatataaataaaaaataagtatatatatattttatatgtatataatatataaacaatgtgtttaatgtttttttaataaaaagatagtatttttatgaggcatatattattatatcttatattttattatatatatatatatatatttttttttttttttttttaatatcattgatcaacattgaattattttaaaaagaaaataaaaagggaaaaaaaaaaaaagaaaaaaaaaaagaataaaaaaagaagaaaaaaaagaataaaaaaagaaaaaaaaaaaagaataaaaaaagaaaaaaaagataaaactttcctaaatttttctcaaaaaaagaagaaaaaggattaaataaataattaaggagaatagtaagttaaatataagttatatataaacatataaggattaccaaaagggaaaatatacaactataaatattataatatcaaaaggataattatttaattaaataaaaaaaaaaaaaaaaaatagaatatatatatatttatatatataaaaaaattataaaaATG**AAACTTTTAAGAGTAACACCTGAAAAAAATATAGAATTTCCTCTTGTTCATTTTCAAGCAGTAACTCAAGTTGTTAAATTAGAAAATGTAAGTGATAAAAAAGTAGCTTTTAAAATAAAAACAACCGCTCCTAATAATTATTTAGTAAGACCATCATTTGGTTTAATAAGTGTAAGAGAAACAATAGAAATACAAATTATATTACAACCCTTGTCAGACAAAGATAATATATCAAACGATAAATTTCAGGTACAATGTTTAAATGTTGATGATAATACTACAGTAGATAAACAATTTTGGATAACAGTTAATAAAAATGAAATACAAGATCATAAACTTATTGTAGTTCTAAACGATGAAAATAATAGTAAATTAAATCATTCTTACATACCCTCAAATAATGTACCTCTCTCAGAAATGAATAACAAAAATATACACAATATGGGATACGTCGATAATAATAATATAAATCAAGATGACCCAAATTTAGCAGATG*gtaaaaaatattacaaagaaatttatatctaatagtatatattaatattgtaatatatgtgtaaatcttatatgatgaattgtcacatatatcgtgttcttatatatataatgtgtgtgtgtactctacaagtgttcaagacaaaatgttatatatatatatatatgtatgtgtgtgtggatatatatatttatttattttatatatatatatatatatatattttttttttttttttttttttgttaag*GTTTAAAAGGAGGTCTACCAGGTAT**GCGTTTAAACGAGCAGAAGTTAATATCAGAAGAGGATTTGGGTGAACAAAAACTCATAAGCGAAGAAGATTTAATAACTTCGTATAGCATACATTATACGAAGTTATCCGGAGAAGGAAGAGGAAGTTTATTAACATGTGGAGATGTAGAAGAAAATCCAGGACCAATGACAGCCAGTTTAACTACCAAGTTCTTGAACAATACCTATGAAAACCCATTTATGAATGCATCCGGTGTTCATTGCATGACTACACAAGAATTAGATGAATTAGCAAACTCTAAAGCTGGCGCATTCATTACAAAGAGTGCTACAACCTTAGAAAGAGAAGGTAACCCTGAACCACGTTACATTTCTGTCCCTCTAGGCAGTATCAACTCCATGGGTTTACCAAACGAAGGTATCGACTACTATTTGTCCTATGTATTAAACCGTCAAAAGAATTATCCTGATGCACCTGCTATTTTCTTCTCAGTTGCTGGTATGAGCATTGATGAAAATTTAAATTTGTTGAGGAAAATCCAAGATAGCGAATTCAACGGTATTACCGAGTTAAACTTGTCTTGTCCTAATGTGCCTGGGAAACCACAAGTTGCTTATGACTTTGACTTGACAAAGGAAACCTTGGAAAAGGTTTTTGCCTTTTTCAAAAAACCTCTTGGTGTCAAGTTGCCTCCTTATTTTGATTTTGCCCATTTTGATATCATGGCAAAAATATTGAACGAGTTCCCATTAGCTTATGTCAACTCTATCAATAGTATAGGAAATGGTCTTTTCATTGATGTGGAGAAGGAGAGTGTAGTAGTGAAGCCAAAGAATGGTTTCGGGGGTATTGGAGGTGAATATGTTAAGCCAACCGCGCTCGCCAATGTTCGTGCATTTTACACTCGTTTGAGACCTGAAATCAAAGTTATCGGTACAGGTGGAATTAAGTCCGGTAAGGATGCATTTGAACATCTTCTATGTGGTGCCTCTATGCTACAGATTGGTACAGAATTACAAAAAGAGGGCGTCAAGATTTTTGAACGTATCGAAAAAGAATTAAAAGACATAATGGAAGCTAAGGGTTATACATCCATAGATCAGTTCCGTGGGAAGTTGAACAGCATTGGTGAAGGTAGAGGTTCTTTGTTGACTTGTGGTGATGTTGAAGAAAATCCAGGTCCAGCTAGCATGGCAGAAATTGGTACGGGTTTTCCATTTGATCCTCATTATGTGGAGGTGCTGGGGGAAAGGATGCATTATGTTGACGTAGGACCAAGAGATGGTACTCCAGTGTTATTTTTGCATGGAAACCCAACCTCGAGTTATGTATGGAGAAATATAATTCCACATGTAGCACCAACACATAGATGTATAGCTCCTGATTTAATTGGTATGGGAAAAAGTGATAAACCTGACTTAGGATATTTTTTTGATGATCATGTCCGTTTTATGGATGCTTTCATTGAAGCTCTTGGCCTTGAAGAAGTAGTATTAGTTATACATGATTGGGGATCCGCCTTAGGATTTCATTGGGCCAAGAGGAATCCTGAAAGAGTAAAAGGAATAGCATTCATGGAATTCATACGACCAATCCCCACATGGGATGAATGGCCAGAATTTGCACGCGAAACATTTCAAGCTTTTAGAACTACAGATGTTGGTAGAAAATTAATAATAGATCAAAATGTATTTATAGAAGGAACTTTACCTATGGGTGTTGTAAGGCCGTTAACAGAAGTTGAAATGGACCACTACCGTGAACCTTTTTTAAATCCAGTAGATAGAGAGCCCTTATGGAGATTTCCTAATGAATTACCTATTGCAGGTGAACCCGCGAATATTGTTGCTTTAGTAGAAGAATATATGGATTGGTTACATCAGTCTCCTGTTCCTAAACTTCTATTTTGGGGTACACCTGGAGTTCTAATACCACCAGCTGAAGCAGCAAGATTAGCAAAATCATTACCAAATTGTAAAGCTGTTGATATAGGTCCTGGGTTGAATTTATTACAAGAAGATAATCCAGATTTGATTGGATCTGAGATAGCTAGATGGCTAAGTACATTAGAAATTTCAGGTACGCGTGCTAGAGGTGCTGCTGCTGGTGCTGGAGGTGCAGGTAGACCTAGGATGAAGTTACTTCGTGTTACTCCAGAGAAGAACATTGAGTTCCCATTAGTACACTTCCAGGCTGTTACACAGGTAGTAAAGCTTGAGAACGTTTCAGACAAGAAGGTTGCATTCAAGATTAAGACTACAGCACCAAACAACTACCTTGTTCGTCCTAGTTTCGGACTTATTTCAGTTCGTGAGACTATTGAGATTCAGATAATTCTTCAGCCACTTAGTGATAAGGACAACATTAGTAATGACAAGTTCCAAGTTCAGTGCCTTAACGTAGACGACAACACAACTGTTGACAAGCAGTTCTGGATTACTGTAAACAAGAACGAGATTCAGGACCACAAGTTAATAGTTGTATTGAATGACGAGAACAACTCAAAGCTTAACCACAGCTATATTCCAAGTAACAACGTTCCATTAAGTGAGATGAACAATAAGAACATTCATAACATGGGTTATGTTGACAACAACAACATTAACCAGGACGATCCTAACCTTGCTGACGGACTTAAGGGTGGATTGCCTGGAATGCAGCGTAAGTACCACGAGTTACTTAACTACTGTGTATTCGTAGACAAGCAGAAGGCTGCATTGGAGAAGGAGAATGAGTCACTTAAGAACCAACTTAAGGCTTACAATTCAAACAGCAACAAGTTTCTTATTGACAACAAGCTTATACCAATAATTATAGTTATGCTTGCAATTATTACTAAGTACATGGGATATTGGTAGaggcctataacttcgtatagcatacattatacgaagttattatgactcgagggatatggcagcttaatgttcgtttttcttatttatatatttataccaattgattgtatttataactgtaaaaatgtgtatgttgtgtgcatatttttttttgtgcatgcacatgcatgtaaatagctaaaattatgaacattttattttttgttcagaaaaaaaaaactttacacacataaaatggctagtatgaatagccatattttatataaattaaatcctatgaatttatgaccatattaaaaatttagatatttatggaacataatatgtttgaaacaataagacaaaattattattattattattatttttactgttataattatgtgtctccttcaatgattcataaatagttggacttgatttttaaaatgtttataatatgattagcatagttaaataaaaaaagttgaaaaattaaaaaaaaacatataaacacaaatgatggtttttccttcaatttcgatatcaatttatagaaacaaaatatatacttgtataattttatttttttatataaatcattacatatataattatacaatattttttctaagagataattatatattaatatatataaaaaaaggtgttttttttttttttttttatttttatttttattttatggtaatattttattttccttattttataaattatattagtttatatgtgattaattttatatattatcaatttatatatttttaaatgcttacttaattatctttttttttttttttttttttttttcccctctttttatattaatttatttttgaaaaaattgatatatatatatatatataatatatatatatacatgtagtagtattaaacaatgtataatatatataaataatatatttatatatttcatttcaattttaattttttttggttttttttttttttctttttgtcatatttaaaaaaaattatattcatataagttatgcattttttataaacattattcaatatatgtataatataatatatatatatatattaatgtattattccaatgtgcatgataaaagaaaaaaataatatttataaaaaaaaagaaaaataaaacaaaaaaagaaaaaaaaaaaaaaaaaaaaaaaaatacaaaaataaataatataatttataattatatattcttgtcacaataaaaatatatatatatatatatatatttataatatgtatattttaaactagaaaaggaataactaatattttatttattatcattcaagatttatattttataataataaatacctaatagaaatatatcaggatccatgcatggttcgctaaactgcatcgtcgctgtgtcccagaacatgggcatcggcaagaacggggactacccctggccaccgctcaggaacgaatttagatatttccagagaatgaccacaacctcttcagtagaaggtaaacagaatctggtgattatgggtaagaagacctggttctccattcctgagaagaatcgacctttaaagggtagaattaatttagttctcagcagagaactcaaggaacctccacaaggagctcattttctttccagaagtctagatgatgccttaaaacttactgaacaaccagaattagcaaataaagtagacatggtctggatagttggtggcagttctgtttataaggaagccatgaatcacccaggccatcttaaactatttgtgacaaggatcatgcaagactttgaaagtgacacgttttttccagaaattgatttggagaaatataaacttctgccagaatacccaggtgttctctctgatgtccaggaggagaaaggcattaagtacaaatttgaagtatatgagaagaatgattaagcttatttaataatagattaaaaatattataaaaataaaaacataaacacagaaattacaaaaaaaatacatatgaattttttttttgtaatcttccttataaatatagaataatgaatcatataaaacatatcattattcatttatttacatttaaaattattgtttcagtatctttaatttattatgtatatataaaaataacttacaattttattaataaacaatatatgtttattaattcatgttttgtaatttatgggatagcgattttttttactgtctgtatttttcttttttaattatgttttaattgtattttatttttattattgttctttttatagtattattttaaaacaaaatgtattttctaagaacttataataataataaatataaattttaataaaaattatatttatcttttacaatatgaacataaagtacaacattaatatatagcttttaatatttttattcctaatcatgtaaatcttaaatttttctttttaaacatatgttaaatatttatttctcattatatataagaacatatttattaaatctagaattctatagtgagtcgtattacaattcactggccgtcgttttacaacgtcgtgactgggaaaaccctggcgttacccaacttaatcgccttgcagcacatccccctttcgccagctggcgtaatagcgaagaggcccgcaccgatcgcccttcccaacagttgcgcagcctgaatggcgaatggcgcctgatgcggtattttctccttacgcatctgtgcggtatttcacaccgcatatggtgcactctcagtacaatctgctctgatgccgcatagttaagccagccccgacacccgccaacacccgctgacgcgccctgacgggcttgtctgctcccggcatccgcttacagacaagctgtgaccgtctccgggagctgcatgtgtcagaggttttcaccgtcatcaccgaaacgcgcgagacgaaagggcctcgtgatacgcctatttttataggttaatgtcatgataataatggtttcttagacgtcaggtggcacttttcggggaaatgtgcgcggaacccctatttgtttatttttctaaatacattcaaatatgtatccgctcatgagacaataaccctgataaatgcttcaataatattgaaaaaggaagagtatgagtattcaacatttccgtgtcgcccttattcccttttttgcggcattttgccttcctgtttttgctcacccagaaacgctggtgaaagtaaaagatgctgaagatcagttgggtgcacgagtgggttacatcgaactggatctcaacagcggtaagatccttgagagttttcgccccgaagaacgttttccaatgatgagcacttttaaagttctgctatgtggcgcggtattatcccgtattgacgccgggcaagagcaactcggtcgccgcatacactattctcagaatgacttggttgagtactcaccagtcacagaaaagcatcttacggatggcatgacagtaagagaattatgcagtgctgccataaccatgagtgataacactgcggccaacttacttctgacaacgatcggaggaccgaaggagctaaccgcttttttgcacaacatgggggatcatgtaactcgccttgatcgttgggaaccggagctgaatgaagccataccaaacgacgagcgtgacaccacgatgcctgtagcaatgccaacaacgttgcgcaaactattaactggcgaactacttactctagcttcccggcaacaattaatagactggatggaggcggataaagttgcaggaccacttctgcgctcggcccttccggctggctggtttattgctgataaatctggagccggtgagcgtgggtctcgcggtatcattgcagcactggggccagatggtaagccctcccgtatcgtagttatctacacgacggggagtcaggcaactatggatgaacgaaatagacagatcgctgagataggtgcctcactgattaagcattggtaactgtcagaccaagtttactcatatatactttagattgatttaaaacttcatttttaatttaaaaggatctaggtgaagatcctttttgataatctcatgaccaaaatcccttaacgtgagttttcgttccactgagcgtcagaccccgtagaaaagatcaaaggatcttcttgagatcctttttttctgcgcgtaatctgctgcttgcaaacaaaaaaaccaccgctaccagcggtggtttgtttgccggatcaagagctaccaactctttttccgaaggtaactggcttcagcagagcgcagataccaaatactgtccttctagtgtagccgtagttaggccaccacttcaagaactctgtagcaccgcctacatacctcgctctgctaatcctgttaccagtggctgctgccagtggcgataagtcgtgtcttaccgggttggactcaagacgatagttaccggataaggcgcagcggtcgggctgaacggggggttcgtgcacacagcccagcttggagcgaacgacctacaccgaactgagatacctacagcgtgagctatgagaaagcgccacgcttcccgaagggagaaaggcggacaggtatccggtaagcggcagggtcggaacaggagagcgcacgagggagcttccagggggaaacgcctggtatctttatagtcctgtcgggtttcgccacctctgacttgagcgtcgatttttgtgatgctcgtcaggggggcggagcctatcgaaaaacgccagcaacgcggcctttttacggttcctggccttttgctggccttttgctcacatgttctttcctgcgttatcccctgattctgtggataaccgtattaccgcctttgagtgagctgataccgctcgccgcagccgaacgaccgagcgcagcgagtcagtgagcgaggaagcggaagagcgcccaatacgcaaaccgcctctccccgcgcgttggccgattcattaatgcagctggcacgacaggtttcccgactggaaagcgggcagtgagcgcaacgcaattaatgtgagttagctcactcattaggcaccccaggctttacactttatgcttccggctcgtatgttgtgtggaattgtgagcggataacaatttcacacaggaaacagctatgaccatgattacgccaagctatttaggtgacactatagaatactcgcggccgctagaaacttttaagagtaacacctgaaaaaaatatagaatttcctcttgttcattttcaagcagtaactcaagttgttaaattagaaaatgtaagtgataaaaaagtagcttttaaaataaaaacaaccgctcctaataattatttagtaagaccatcatttggtttaataagtgtaagagaaacaatagaaatacaaattatattacaacccttgtcagacaaagataatatatcaaacgataaatttcaggtacaatgtttaaatgttgatgataatactacagtagataaacaattttggataacagttaataaaaatgaaatacaagatcataaacttattgtagttctaaacgatgaaaataatagtaaattaaatcattcttacataccctcaaataatgtacctctctcagaaatgaataacaaaaatatacacaatatgggatacgtcgataataataatataaatcaagatgacccaaatttagcagatggtaaaaaatattacaaagaaatttatatctaatagtatatattaatattgtaatatatgtgtaaatcttatatgatgaattgtcacatatatcgtgttcttatatatataatgtgtgtgtgtactctacaagtgttcaagacaaaatgttatatatatatatatatgtatgtgtgtgtggatatatatatttatttattttatatatatatatatatatatattttttttttttttttttttttgttaaggtttaaaaggaggtctaccaggtatgcaaaggaaatatcatgaacttttaaattattgcgtttttgttgataaacaaaaagcagccctagaaaaagaaaacgaaagtttaaaaaatcagttaaaagcatataacagtaattctaataaattcttaatagataataaattaattcctattataattgtaatgttagctataataacaaaatatatgggttactggtaatttcaaagttatacaaagaaaatatattaatatatatgtttacatttttgtgtaaggaaaatatcaagaatcaataaattataaaatagtaaattcgataagttataaaatgaaatatatgataaattataaaaatgattaataaaaaaaaaacccaaaaaaaaaaaaaaaaaaaaaaaaaagagagaagaaaacacatattaatatatatatatatatatatatttctatgtatacaaaggaatattaacattacaccagtattattaaataaatattaatatccaacaatatattattaagtacaattatacctattgaacagtattaaaagcaaaaacaaggaaatttatatatatatatatttatttatttatatgtatataaatacattctttatctatattgtatttagtttttaataattttttttatttttctcttccttactttttaatatgtttaaataattcataaacaagtagtaaaaaatttataatatatatatgaaatatgttatcaatatttaaaattgaagaaaat

**5’ amplicon size: 2185bps**

**3’ amplicon size: 934bps**

**> 2xFKBP-PfVAP yDHODH recodonPfVAP**

tgaaggaaaaaaagaaaaaaaaaaggaaaaaaaaaaaaaaaaaaaaattaataataatatataatattaataatattttatatatttatatatttttcttacatatataatatatatatatatatataataataataataataataataatatatatgataaacttttattatttagtgaattttatataatttttactttatgttatataaataaatataaataaaaaataagtatatatatattttatatgtatataatatataaacaatgtgtttaatgtttttttaataaaaagatagtatttttatgaggcatatattattatatcttatattttattatatatatatatatatatttttttttttttttttttaatatcattgatcaacattgaattattttaaaaagaaaataaaaagggaaaaaaaaaaaaagaaaaaaaaaaagaataaaaaaagaagaaaaaaaagaataaaaaaagaaaaaaaaaaaagaataaaaaaagaaaaaaaagataaaactttcctaaatttttctcaaaaaaagaagaaaaaggattaaataaataattaaggagaatagtaagttaaatataagttatatataaacatataaggattaccaaaagggaaaatatacaactataaatattataatatcaaaaggataattatttaattaaataaaaaaaaaaaaaaaaaatagaatatatatatatttatatatataaaaaaattataaaaATG**AAACTTTTAAGAGTAACACCTGAAAAAAATATAGAATTTCCTCTTGTTCATTTTCAAGCAGTAACTCAAGTTGTTAAATTAGAAAATGTAAGTGATAAAAAAGTAGCTTTTAAAATAAAAACAACCGCTCCTAATAATTATTTAGTAAGACCATCATTTGGTTTAATAAGTGTAAGAGAAACAATAGAAATACAAATTATATTACAACCCTTGTCAGACAAAGATAATATATCAAACGATAAATTTCAGGTACAATGTTTAAATGTTGATGATAATACTACAGTAGATAAACAATTTTGGATAACAGTTAATAAAAATGAAATACAAGATCATAAACTTATTGTAGTTCTAAACGATGAAAATAATAGTAAATTAAATCATTCTTACATACCCTCAAATAATGTACCTCTCTCAGAAATGAATAACAAAAATATACACAATATGGGATACGTCGATAATAATAATATAAATCAAGATGACCCAAATTTAGCAGATG*gtaaaaaatattacaaagaaatttatatctaatagtatatattaatattgtaatatatgtgtaaatcttatatgatgaattgtcacatatatcgtgttcttatatatataatgtgtgtgtgtactctacaagtgttcaagacaaaatgttatatatatatatatatgtatgtgtgtgtggatatatatatttatttattttatatatatatatatatatatattttttttttttttttttttttgttaag*GTTTAAAAGGAGGTCTACCAGGTAT**GCGTTTAAACGAGCAGAAGTTAATATCAGAAGAGGATTTGGGTGAACAAAAACTCATAAGCGAAGAAGATTTAATAACTTCGTATAGCATACATTATACGAAGTTATCCGGAGAAGGAAGAGGAAGTTTATTAACATGTGGAGATGTAGAAGAAAATCCAGGACCAATGACAGCCAGTTTAACTACCAAGTTCTTGAACAATACCTATGAAAACCCATTTATGAATGCATCCGGTGTTCATTGCATGACTACACAAGAATTAGATGAATTAGCAAACTCTAAAGCTGGCGCATTCATTACAAAGAGTGCTACAACCTTAGAAAGAGAAGGTAACCCTGAACCACGTTACATTTCTGTCCCTCTAGGCAGTATCAACTCCATGGGTTTACCAAACGAAGGTATCGACTACTATTTGTCCTATGTATTAAACCGTCAAAAGAATTATCCTGATGCACCTGCTATTTTCTTCTCAGTTGCTGGTATGAGCATTGATGAAAATTTAAATTTGTTGAGGAAAATCCAAGATAGCGAATTCAACGGTATTACCGAGTTAAACTTGTCTTGTCCTAATGTGCCTGGGAAACCACAAGTTGCTTATGACTTTGACTTGACAAAGGAAACCTTGGAAAAGGTTTTTGCCTTTTTCAAAAAACCTCTTGGTGTCAAGTTGCCTCCTTATTTTGATTTTGCCCATTTTGATATCATGGCAAAAATATTGAACGAGTTCCCATTAGCTTATGTCAACTCTATCAATAGTATAGGAAATGGTCTTTTCATTGATGTGGAGAAGGAGAGTGTAGTAGTGAAGCCAAAGAATGGTTTCGGGGGTATTGGAGGTGAATATGTTAAGCCAACCGCGCTCGCCAATGTTCGTGCATTTTACACTCGTTTGAGACCTGAAATCAAAGTTATCGGTACAGGTGGAATTAAGTCCGGTAAGGATGCATTTGAACATCTTCTATGTGGTGCCTCTATGCTACAGATTGGTACAGAATTACAAAAAGAGGGCGTCAAGATTTTTGAACGTATCGAAAAAGAATTAAAAGACATAATGGAAGCTAAGGGTTATACATCCATAGATCAGTTCCGTGGGAAGTTGAACAGCATTGGTGAAGGTAGAGGTTCTTTGTTGACTTGTGGTGATGTTGAAGAAAATCCAGGTCCAGCTAGCATGACCATGTACCCATACGATGTTCCAGATTATGCTACGATGTACCCGTACGACGTGCCGGACTACGCGAGATCTGCTGCTGCTGGTGCTGGTGGTGCTGCTAGAGCTGCTCTGCAGAGAGGAGTACAAGTTGAAACAATATCACCAGGAGATGGTCGTACATTTCCAAAAAGAGGTCAAACTTGTGTTGTACATTATACTGGAATGCTTGAAGATGGAAAGAAATTTGATTCATCTCGTGATAGAAATAAACCATTTAAATTTATGCTAGGTAAACAAGAAGTAATACGAGGTTGGGAAGAAGGAGTTGCTCAAATGAGTGTAGGTCAAAGAGCAAAACTTACTATATCTCCAGATTATGCTTATGGTGCAACTGGACATCCAGGTATAATTCCACCTCATGCAACTCTTGTATTTGATGTGGAGCTTCTAAAACTAGAAACTAGAGGTGTTCAGGTTGAAACAATTTCACCTGGAGATGGCAGAACCTTTCCTAAAAGAGGACAGACTTGCGTAGTTCATTATACAGGCATGCTAGAGGATGGTAAGAAATTTGATTCTAGTCGAGATAGAAATAAGCCATTCAAGTTTATGCTAGGTAAACAGGAAGTAATAAGAGGTTGGGAAGAGGGTGTAGCACAGATGTCAGTTGGACAAAGAGCAAAGTTAACAATATCACCAGATTATGCATACGGTGCAACAGGCCATCCTGGCATCATCCCTCCACATGCAACTTTAGTATTCGACGTTGAATTGTTAAAGTTAGAGACAACGCGTGCTAGAGGTGCTGCTGCTGGTGCTGGAGGTGCAGGTAGACCTAGGATGAAGTTACTTCGTGTTACTCCAGAGAAGAACATTGAGTTCCCATTAGTACACTTCCAGGCTGTTACACAGGTAGTAAAGCTTGAGAACGTTTCAGACAAGAAGGTTGCATTCAAGATTAAGACTACAGCACCAAACAACTACCTTGTTCGTCCTAGTTTCGGACTTATTTCAGTTCGTGAGACTATTGAGATTCAGATAATTCTTCAGCCACTTAGTGATAAGGACAACATTAGTAATGACAAGTTCCAAGTTCAGTGCCTTAACGTAGACGACAACACAACTGTTGACAAGCAGTTCTGGATTACTGTAAACAAGAACGAGATTCAGGACCACAAGTTAATAGTTGTATTGAATGACGAGAACAACTCAAAGCTTAACCACAGCTATATTCCAAGTAACAACGTTCCATTAAGTGAGATGAACAATAAGAACATTCATAACATGGGTTATGTTGACAACAACAACATTAACCAGGACGATCCTAACCTTGCTGACGGACTTAAGGGTGGATTGCCTGGAATGCAGCGTAAGTACCACGAGTTACTTAACTACTGTGTATTCGTAGACAAGCAGAAGGCTGCATTGGAGAAGGAGAATGAGTCACTTAAGAACCAACTTAAGGCTTACAATTCAAACAGCAACAAGTTTCTTATTGACAACAAGCTTATACCAATAATTATAGTTATGCTTGCAATTATTACTAAGTACATGGGATATTGGTAGaggcctataacttcgtatagcatacattatacgaagttattatgactcgagggatatggcagcttaatgttcgtttttcttatttatatatttataccaattgattgtatttataactgtaaaaatgtgtatgttgtgtgcatatttttttttgtgcatgcacatgcatgtaaatagctaaaattatgaacattttattttttgttcagaaaaaaaaaactttacacacataaaatggctagtatgaatagccatattttatataaattaaatcctatgaatttatgaccatattaaaaatttagatatttatggaacataatatgtttgaaacaataagacaaaattattattattattattatttttactgttataattatgtgtctccttcaatgattcataaatagttggacttgatttttaaaatgtttataatatgattagcatagttaaataaaaaaagttgaaaaattaaaaaaaaacatataaacacaaatgatggtttttccttcaatttcgatatcaatttatagaaacaaaatatatacttgtataattttatttttttatataaatcattacatatataattatacaatattttttctaagagataattatatattaatatatataaaaaaaggtgttttttttttttttttttatttttatttttattttatggtaatattttattttccttattttataaattatattagtttatatgtgattaattttatatattatcaatttatatatttttaaatgcttacttaattatctttttttttttttttttttttttttcccctctttttatattaatttatttttgaaaaaattgatatatatatatatatataatatatatatatacatgtagtagtattaaacaatgtataatatatataaataatatatttatatatttcatttcaattttaattttttttggttttttttttttttctttttgtcatatttaaaaaaaattatattcatataagttatgcattttttataaacattattcaatatatgtataatataatatatatatatatattaatgtattattccaatgtgcatgataaaagaaaaaaataatatttataaaaaaaaagaaaaataaaacaaaaaaagaaaaaaaaaaaaaaaaaaaaaaaaatacaaaaataaataatataatttataattatatattcttgtcacaataaaaatatatatatatatatatatatttataatatgtatattttaaactagaaaaggaataactaatattttatttattatcattcaagatttatattttataataataaatacctaatagaaatatatcaggatccatgcatggttcgctaaactgcatcgtcgctgtgtcccagaacatgggcatcggcaagaacggggactacccctggccaccgctcaggaacgaatttagatatttccagagaatgaccacaacctcttcagtagaaggtaaacagaatctggtgattatgggtaagaagacctggttctccattcctgagaagaatcgacctttaaagggtagaattaatttagttctcagcagagaactcaaggaacctccacaaggagctcattttctttccagaagtctagatgatgccttaaaacttactgaacaaccagaattagcaaataaagtagacatggtctggatagttggtggcagttctgtttataaggaagccatgaatcacccaggccatcttaaactatttgtgacaaggatcatgcaagactttgaaagtgacacgttttttccagaaattgatttggagaaatataaacttctgccagaatacccaggtgttctctctgatgtccaggaggagaaaggcattaagtacaaatttgaagtatatgagaagaatgattaagcttatttaataatagattaaaaatattataaaaataaaaacataaacacagaaattacaaaaaaaatacatatgaattttttttttgtaatcttccttataaatatagaataatgaatcatataaaacatatcattattcatttatttacatttaaaattattgtttcagtatctttaatttattatgtatatataaaaataacttacaattttattaataaacaatatatgtttattaattcatgttttgtaatttatgggatagcgattttttttactgtctgtatttttcttttttaattatgttttaattgtattttatttttattattgttctttttatagtattattttaaaacaaaatgtattttctaagaacttataataataataaatataaattttaataaaaattatatttatcttttacaatatgaacataaagtacaacattaatatatagcttttaatatttttattcctaatcatgtaaatcttaaatttttctttttaaacatatgttaaatatttatttctcattatatataagaacatatttattaaatctagaattctatagtgagtcgtattacaattcactggccgtcgttttacaacgtcgtgactgggaaaaccctggcgttacccaacttaatcgccttgcagcacatccccctttcgccagctggcgtaatagcgaagaggcccgcaccgatcgcccttcccaacagttgcgcagcctgaatggcgaatggcgcctgatgcggtattttctccttacgcatctgtgcggtatttcacaccgcatatggtgcactctcagtacaatctgctctgatgccgcatagttaagccagccccgacacccgccaacacccgctgacgcgccctgacgggcttgtctgctcccggcatccgcttacagacaagctgtgaccgtctccgggagctgcatgtgtcagaggttttcaccgtcatcaccgaaacgcgcgagacgaaagggcctcgtgatacgcctatttttataggttaatgtcatgataataatggtttcttagacgtcaggtggcacttttcggggaaatgtgcgcggaacccctatttgtttatttttctaaatacattcaaatatgtatccgctcatgagacaataaccctgataaatgcttcaataatattgaaaaaggaagagtatgagtattcaacatttccgtgtcgcccttattcccttttttgcggcattttgccttcctgtttttgctcacccagaaacgctggtgaaagtaaaagatgctgaagatcagttgggtgcacgagtgggttacatcgaactggatctcaacagcggtaagatccttgagagttttcgccccgaagaacgttttccaatgatgagcacttttaaagttctgctatgtggcgcggtattatcccgtattgacgccgggcaagagcaactcggtcgccgcatacactattctcagaatgacttggttgagtactcaccagtcacagaaaagcatcttacggatggcatgacagtaagagaattatgcagtgctgccataaccatgagtgataacactgcggccaacttacttctgacaacgatcggaggaccgaaggagctaaccgcttttttgcacaacatgggggatcatgtaactcgccttgatcgttgggaaccggagctgaatgaagccataccaaacgacgagcgtgacaccacgatgcctgtagcaatgccaacaacgttgcgcaaactattaactggcgaactacttactctagcttcccggcaacaattaatagactggatggaggcggataaagttgcaggaccacttctgcgctcggcccttccggctggctggtttattgctgataaatctggagccggtgagcgtgggtctcgcggtatcattgcagcactggggccagatggtaagccctcccgtatcgtagttatctacacgacggggagtcaggcaactatggatgaacgaaatagacagatcgctgagataggtgcctcactgattaagcattggtaactgtcagaccaagtttactcatatatactttagattgatttaaaacttcatttttaatttaaaaggatctaggtgaagatcctttttgataatctcatgaccaaaatcccttaacgtgagttttcgttccactgagcgtcagaccccgtagaaaagatcaaaggatcttcttgagatcctttttttctgcgcgtaatctgctgcttgcaaacaaaaaaaccaccgctaccagcggtggtttgtttgccggatcaagagctaccaactctttttccgaaggtaactggcttcagcagagcgcagataccaaatactgtccttctagtgtagccgtagttaggccaccacttcaagaactctgtagcaccgcctacatacctcgctctgctaatcctgttaccagtggctgctgccagtggcgataagtcgtgtcttaccgggttggactcaagacgatagttaccggataaggcgcagcggtcgggctgaacggggggttcgtgcacacagcccagcttggagcgaacgacctacaccgaactgagatacctacagcgtgagctatgagaaagcgccacgcttcccgaagggagaaaggcggacaggtatccggtaagcggcagggtcggaacaggagagcgcacgagggagcttccagggggaaacgcctggtatctttatagtcctgtcgggtttcgccacctctgacttgagcgtcgatttttgtgatgctcgtcaggggggcggagcctatcgaaaaacgccagcaacgcggcctttttacggttcctggccttttgctggccttttgctcacatgttctttcctgcgttatcccctgattctgtggataaccgtattaccgcctttgagtgagctgataccgctcgccgcagccgaacgaccgagcgcagcgagtcagtgagcgaggaagcggaagagcgcccaatacgcaaaccgcctctccccgcgcgttggccgattcattaatgcagctggcacgacaggtttcccgactggaaagcgggcagtgagcgcaacgcaattaatgtgagttagctcactcattaggcaccccaggctttacactttatgcttccggctcgtatgttgtgtggaattgtgagcggataacaatttcacacaggaaacagctatgaccatgattacgccaagctatttaggtgacactatagaatactcgcggccgctagaaacttttaagagtaacacctgaaaaaaatatagaatttcctcttgttcattttcaagcagtaactcaagttgttaaattagaaaatgtaagtgataaaaaagtagcttttaaaataaaaacaaccgctcctaataattatttagtaagaccatcatttggtttaataagtgtaagagaaacaatagaaatacaaattatattacaacccttgtcagacaaagataatatatcaaacgataaatttcaggtacaatgtttaaatgttgatgataatactacagtagataaacaattttggataacagttaataaaaatgaaatacaagatcataaacttattgtagttctaaacgatgaaaataatagtaaattaaatcattcttacataccctcaaataatgtacctctctcagaaatgaataacaaaaatatacacaatatgggatacgtcgataataataatataaatcaagatgacccaaatttagcagatggtaaaaaatattacaaagaaatttatatctaatagtatatattaatattgtaatatatgtgtaaatcttatatgatgaattgtcacatatatcgtgttcttatatatataatgtgtgtgtgtactctacaagtgttcaagacaaaatgttatatatatatatatatgtatgtgtgtgtggatatatatatttatttattttatatatatatatatatatatattttttttttttttttttttttgttaaggtttaaaaggaggtctaccaggtatgcaaaggaaatatcatgaacttttaaattattgcgtttttgttgataaacaaaaagcagccctagaaaaagaaaacgaaagtttaaaaaatcagttaaaagcatataacagtaattctaataaattcttaatagataataaattaattcctattataattgtaatgttagctataataacaaaatatatgggttactggtaatttcaaagttatacaaagaaaatatattaatatatatgtttacatttttgtgtaaggaaaatatcaagaatcaataaattataaaatagtaaattcgataagttataaaatgaaatatatgataaattataaaaatgattaataaaaaaaaaacccaaaaaaaaaaaaaaaaaaaaaaaaaagagagaagaaaacacatattaatatatatatatatatatatatttctatgtatacaaaggaatattaacattacaccagtattattaaataaatattaatatccaacaatatattattaagtacaattatacctattgaacagtattaaaagcaaaaacaaggaaatttatatatatatatatttatttatttatatgtatataaatacattctttatctatattgtatttagtttttaataattttttttatttttctcttccttactttttaatatgtttaaataattcataaacaagtagtaaaaaatttataatatatatatgaaatatgttatcaatatttaaaattgaagaaaat

**5’ amplicon size: 2260bps**

**3’ amplicon size: 934bps**

## **> PF3D7_1414500**

aatattattatatctttattacaaaataataatagaaaaaatagaaataatattgtggatattttttaatacatacatacatatataatattatatttatatataaaatatatacatagataatacatatatatatttatttatctatatatgctggattaatattaatatatatatatatattatatatatatatatatatatatatgtaataaatatatatgttcgtcatataaaataactaaaaaaaataaaaaaaaaaattaaaataaaataaaaataattaatatttaattgaaatgtttatacatttttcagaacatcaactaaatatatgaatatatatgtataatatatataatacatatgtatattatatatatatatatatataaatataattatgtatttttgtatgtttaatattatttttttcgtttattctatatatgaatatattattatataacatattaatgttttaattttcatgtttgatatagaattcaatcggtattttaattcttgtattttaattgaatgaatatatatatatatatatatatatataatatatatatatttatatttattaatatgtttgtaaatatgataatatggatcaatcttttcatatataaaaaaaaaaaaaaaatttaccattatataaatattcttactttatacattatgtagttaaaagacttaaaatatatgcattttggtattctttttttttttttttttctatttagttttgttttacatgtaaataaacttaaatttgattatttttttattttaatttttattatacttttttataaatgtgtgtagttattactttatttttttacatacatatatatatatatatatatatatatatgtatatatttttacaatctgaatgccttaagtttatttgatacatatatttccatttttttttggaaaataaaaaagataaaaatataaaaatttttgataacgtataaattgtttttttttttttttttaatatatattaaaggatgcggctttgattatatatatatatatataatataatgtgaaaaagtctaataaaagaaaaaaaaaaaaaaaaaaaaaaaaagaagtaaaataatatatatatatttatgaaaaaaatcttcagatataaatatgttttatcttattttattaatttatagtgtacatattaataatatattatataatatatgattatgtagaaattaagaaaaaacaaaaaaaaataaaacaaatactttttttatgtgattttattttattctatgttaaagaatgatatacaaatttacattatacttctgctgaatttgtaaaaatttataattaatgtaatgatttctttgatttattatttctttcaaatttttatattttactgtgcatatttataatattataatgtaaatgcaataaatatattttatatatatatatatatatatatatatatatatatatatacatttatatttttttatattcctgttacacataatatataaataaatattttcacatttgttatattttttattttattttacttccatttatttatttatattctttcttttttttcttttttttgtttttacatattttaaaaataatataccgttatgtaaaaaatgtgttcattcatataaacggtttctcattatatatataataaatatatataaatttttataaatttatatatttatcatataaatgaatatacacatatatatatatatatatatatatatatatatatatatatatttttatttatatatgattaaaaaaaaaaagaaaaaattgtatgatatggtataattacaatactgaagtaattttttttgtaaatttttatatcatttattgttttaatttatttatttatttatttttttttttttgaaaaaaaataataatatgttataaaagtagacataatataaattagttgttaatttttatatataatattacatcatattgtttatatttgttttgttggttttattaccttttttttttttctttttttttttctttttttttctatataaaaatttataacaatatatttgtttacttaaaaataatatatatatatatatatatatatatatgtatatatttttgaagagggaaaaatatggtgttcttacattcttttcatttttatcctttttaaaaatttattaagtaaaccaattgataaatgtgatcacatattttggtgtgtttcactttttttttttatttctaaaaaaaagaaaagaaaaaaagaactttttatatttaattggtacttgttcttttaatagaataagttgtgtgcaaatcttatagaaaaATGAATCAATATGATCAATGGAATCGTAGAATTCGG*gtaggaaaaaaaaaaaaaaataaagaaaaataaaattaaatatatatactttgttgaattaattatattatattgttttattttattttaattatttattttatttttttttttttttttccttttag*ACAATATGGTACTGTAGTAGTCTGTATGTAGAATATAAGAACGCTTTGAGGAAATCCAAAAAAATGCCGGTGGAGAAAAAAAATGAATACTGGGAAAAGAAGCATGAAGAATTTGCTACAAAGATGTTAAATAATATATATGAATTAAGAG*gtaaggaatattattataaaagttaaaaaaaaaaaaaaaaaaaaaaaaaataatatatataaatataaatatataatacatgaatgcgttgtctatttttcatattttataaaaaaataccactacacgtttatattatctttatttatttatttgtttgtttgtttattataaatag*GTTGGTGGGTTAAAGTAGGACAATTTTTAAGTACCCAGGAAAATATCATGCCTGTTGCTTATATAGAAAAATTTACGAAGCTGCAAGATATGATGCCTACGTCTTCATTTGATAAAATAGAAAATATTTTAAAAAAGGAATTAG*gtaatttaaaaaaaaaaataaaaaataaatgtgtctatgtatgcataaaagtatgagtatatgtatgtataaatttatgtacaaatgtatgcgtatatgtatgtataaatttatgtacaaatgtattcgtatatgtatgtataaatttatgtacaaaagtatgcgtatatgtatatataaatatatgtattttttttttttttttgtttgattcctag*GTAACATTTATGAAATGTTTGAATACATCGATAAAGAACCACTAGCTAGCGCTTCCATTGGACAAGTACATAAAGCGAAATTAAAAAAATATGATAAACATATATGCGATATAGACAGTAGAAAGCATAAGGATTATAATGTAATAATAAAAATTCAACATGAAGGAATAGATCAGTTTTTGTCATCAGATATAAGTACTTTAAAAAAAGTGTCTTGGGCATTTGGATTAATAGATAAGAATTTTTATTTTACAGATTTTATAGATGAATGGCAAGATTCTGCTTCTCGAGAATTAAATTATAAATATGAATTATATCATCAATTGTTAGCTTATAACAGTTTTAAAAAATCAGGTATCCCTGTAAAAATACCTAAAATTTATTGTGCTCATACAACTTCTAAGGTATTGGTTATGGAATATATTAAAGGCTTTAAAATAACTGATACTGAATTACTTAAAAAATATAATGTAGATACATATAAAATGGTCTATAAAATTATTGACTATTTTGCTTATCAAATACATAATGATGGTTTTTTTCATGGGGACCCACATCCGGGAAATATATTAGTCATGATGGAAGAAAAGAAAAAAAGAAAAAGGAAAAAAAAATATAATAAAGATAAAATAAATAACTCGAAAATGAAAATGAAAAATGAAAATATATATATGAACAAGAGAAAAAAAAATCATAAAAAAAGTAATAGGAATTCTAAGAAAAAGTTAAGAAGTTGTAAAAAAAAAAAACTAGAAATAAATAAAGAAAGAGAAAAAGGAGAAGAAAAGAAAAAAAATATGTATAATAATAATGATATAAATTATTGTGAAAATAAAAATGAAGATATTATAACAAACAATCCCAAATTACAAAATAATAGTAGTGATATGAAAAATATAATAAAATTTACAAGTTATTCCAAAAGTGTATCTTCATCTAATGCTTCATTGATGGATCATTTTGTGGATTCAAGATATAATATTGATAAAATGAGAGCTCCTAATAATTTTTCGAATCCTTTTCATAAATCTTTTAGTTTTGTTGATTATTCACTTAATCAAGATGTAGATTCCGATAAAATGAATCAAGAAGATTATGAATATATTTTGAAGGAAAATATGAGTGATGGTTTAGATAATATGAAAAATTGTCTCTCAATATCCCATGAATATATTTCATGCGATCAATCAAGAAGTGGAGTAACATCTATTGCGACAAGTAGGAAAACACAAAATAGGGATTCTAACAAAGCTAGTGAAAATAGTATTATGGAATCTCGAGAAACAACGATGTTTTCTTTTGTTAACAATATGGAGGAGTTAGTCAAGATGAAATCCGAGACTAAAACGGATCTACAAATTAAAGGTAATGATAAAAAAGTGGTTAAAGGTAGTAAGTGTGACAATAAAAATGATCAAAATGTTAATAATGATAATATTAATAATGATAATATTAATAATGATAAAATTAATAATGATAATATTAATAATGATAATATTAATAATAATAATATTGTTAATAATAATAACAATATTTCTAATGGTGAGAAAGAGAAAAACAAAAGTGGTCCGATATACAAATCCAAATTCAAACTATCAGATTTAAAAAAAAGAAAAAATGCAAAATATAATTTTGTACCTGTTATTATTGATTGGGGTTTAATTAAACAATTAGATAGCGTAATGAAATTCGCCTTTTGTAAATTAGTTTATAATATTAGTTGTATGAATGTATTAAATATTATAGAAGCCTTTGAAGATATGGGTTTCTGCTTTAAAGAAGATTTTACATACGATCCTGAAATATATATTGAGAATTTGAAAACATACTTTTTAAGCAAATTAGAAGAATCTAAAAGTAAAGGTAATGAAGGAGAAAAGGAATTTGGTAGTGCTAATACAATTGATGAAGCAAATAAGAGTAATAAGAATATGGAAATATTAAAAAATATTGAAAAAAAAGATGTTATGGATAAAAATCCGATAAGTGATGTACCTAAAGATATTATATTTTTTATGAGAGTAGCATCATTATTACATGGATTATGTACACAATTAAATGTAAAAATTAATTATTTAAATATTTTTTCCAGAAGAGCTAAAGAAGCTTTAGAAAAAATATATATACCAATAAATAATAGTATACACACAATACCAATAGATAAATGTCCAAATACATTTGTAGAAAAAAGAATTCATCAATTTATAAAAACATTATATGAAAAAAAAAAAATATTAGGATGTCAAGTAGCCATTATACATAAAAAAAAGCTAGTTGTTAATACATGTGTAGGAATTACAAGTACAACAGATAAAAGGCCTATAACAAAACATTCTTTATTTAATGGATATTCATTAAACAAAATTATTTTAAATATAGCTCTATATCATCTTATATGTAACTCAGTAAATGAACAAACGTCAGGTGAAAGTATTCTGCTCGGTATGGATAAGAAAAAGGATAAGCTAGGGGAAAAGATAAGTCAAAAAATTCAAGAAAAAATAAATGAAAAGATAAAAATTAAATCAAGTGCGAAGGACAAAAATCAATCCGATGAAAAAATTAAAATGAAGATAAACGAAAAGGATAAGGATAAGGATAAGGGTAAAGGTAAGGAAAAGAATAAAAATAATAATAAGCAAAACAATGATAGGGATAAAAATAATAATGAGGAAAACAATAATAATGAGGACAATAATAAGGATAACAATAATAATAATAATAATAATAATAATAATAATAATAATAATAATAATAATAATAAATTATTAAACGAAATGGATCAAAGTAAACACAATGCAAATGAAAAAACATCCAATATGAAAAATAACAATAAACTTTATGAAGAAGAAGATGTAGAGGATGAAGTTGCTGATGATAATAAAACAGATAATGAAAATAATTATATTACCGAAAAATTAAATGAATTTAATAAAATAGAAAACGATTATTTTTCAGCAAATGTTAATGAAAATATAATTAAATGTGAAGAGTTTGGTGAGAGCAAGGATATTATAAGTCCTTTATCTGTCAATAGTTTCATAAGTTTTGATAGTTATAATAGTTATAATAATAATGTTAGTTGGATCAGTAGTATGAGTAGGAATAGTTACAAAAGTATGATTGAAAGGATAGAAGATGATGTGGAATATAAAGAAATTAATGATGATCTACCTAATGATAGTAAGGTTAGTTGTATGAGCGAATACAAAAAGGATGATAAATATAAAGGGAGAAATTATATACATGATCACAATAATGATGATGTACATCATCATAATAATCATAATAATCATAATAATCACAATCATCACCATAATCATCACCATAATCATCACCATAATCATCACCATAATCATCACCATAATCATCACCATAATCATCACCATAATCATCACCACCATCACCATCATGATAATAATATAAATAAAAGTAATAAAGATACAGAGTCCAATCCAAATAAATCCATAAATATATATCCAAATATATCATATAACAAAATGGATACCTTAATACAAAAAAAACAAAGTTATATTGATGAGAAAAACTTTATAAAAAGTATAGAAAGCAAACAAATAAGAAATTTCAAAAATATTATTAACGATCATATTTGTAATTACTGGGATGGATTTATATGTAATAATAAAAAAAATATAACGATAAAAGATATATTAACTTTAAAGTGTTTTATTAAGAAACCTTTTCATGATAAAATAACGTTAAGCAAATTTATAGATTACGATCAAATGATAAATATGATTGAGAACTCTAAGCATTATAAGAGTTCAGATAAGACGTCGAAATATGGAGAGTATTTATATTTAATCGATACGTATATTATATCTGAGTTAATACATAATATATCAGGATTACATTATTATGAGTATATATATAAATATATAGTGAAACCATTAAATTTGACTGAAGAAATGTTTGTACCTATACCTTCAGAATTATTAAAAAATCAAAAATACGAAGATTCAAAAAAGAAAAATGTCAACAATCAAAATAATGTAAATACCACTGGTGGTCTTAGTAATAATAATTTAGATAAAACAAAAAAATATATAAAGGATAAAAAGACAAATGTTATAGATAAGAAATCTATTGTTATGGATAAATGGCCTAACATGATGAAGGATTATACAACCGGTGCTTTAAAAAATAATGAATACAAGTCAACTTCATTTATTGGTGGTAAGAAATTTAAAAAGTCTAGTACTATTTCCAACTTTATAGACGATATAAATGATGATATGAATTTTTATATTATGAACGCGGAACTTTTAAAGTTTGAAAATATTAATTTAAAGAATACCATATCTAACAAAAAAAAATTTGATAGTAATCAAAAAGTTTTATCCAAAAAAAGAAGTATAAGTGTAGATGTGTATTATTCTAAGAAAAAAATTAAAG*gtaacacaataaatagaataaaataagaacaagtgaatagataaaaaaaaaaaaataaaaaaaaaaataaacacatcaaaattgatatataaagaaaaaatatattcataaagatgtaatatgtaatatgtaatatgtaatatatatatatatatatatatatatatatatatttatttatttatttattttttgtttatttatag*ATGGTAGTAAGAAGGAGAAAAAAAAAAATAAGGATAAGAAAGATCAAAAGGGAAGTAAAACCAAAATGGATTTTGAAAAAAGAGACATTGCAGAAATGAAGAAAGTAGATAGGAATGAACAGAAAGACCAAACGAATCAACAGAAAGATCAAACGAATGAACAGAAAAACCAACCGAGTCAACAAAACGAACAAATGTATGATGACATAGTTGAAACGTATGAAGATATAATTGAAACATATGAAGACAAAATAGAACTAAACCATGATAATGATGATAGTAACATAAAAACGAATGATGTTTATCATGACATTATAGAAGAGAAAGAAGACAATGAATATTATAATAATATTAAGAATGAATATGAGAATAAAAGGAATAATAATAAAATTGAAATATCAAAGGATTCTTTAAATAAGAATCCATCTAATATAGAATGTAAAATGAATATTTTAATAAATAACGAACTTATAGATAATGAAAATGCAAGAAAATTAGATTCAAGTAATCTTCTTGAAAGTACCTTAAGGAATAAAGAATCCTTTAGTTTTAAAAATAAATTTCTGAATCATGTAGATAATTTAAAATTAAAAACGAAGAAAATAAATGATTCTATAAAAAATATGTATGAAAATAATGAGAATATTTTTTTGAAAAATTTCTTTCTATCATCTAAGAAATATGATAATATAGAAAATAACTTAGATCACAGAAATGTAGAGGCTAATTGTGTAGATAATAATATGTGTGAAAGTAATAGTCTGAATAATAGTTGTTATAATAATAGTTTTTATAATAGTTTTTATAATAGTCATAGTAATAGTAAAAATAGTAGTAAAATAAGAAACTACAATAATATGGATAAGGAAAATAATAATGTTAATAGTGATAATATAAATTTCAATCCCAATGATAATAAAAAGAAAAATAGTTATAATAATAATAATAATAATAATAATAACAATATTATGGTGAGTTCTAGCCCTTATATTTATAATTACAACAATAAATATATATTAAAAGAAAATTATGATAAAAAATTTAATATGAACAAATTTAATAAAGTAATGTTATCTAATCAAAAAGAACCAGTTCATAAAAATAATAAAAAAGATAATTTATTTTTATTTGAAGAAAGAGGTAAGAGATCACATAGTTATAGTGACTGTAGAAATGAATCACACCATGTAATGAAAAATTCTTATATGGAGAATATTAACAATATAAATAATTATAGTAATAGTAGTATGAATTACTTAAACTATTATATTAGTAATAAGAAAAGTGTAAATTATATGAATGCTAATGGGGTCGTCATAAATTATGATGATAATAATAATAATGTAAATAATGTTAATAATGTAAATAATATGAATAATATAAATAACAATGAAACTAATAATACTAACAAAATGAGCTTCTTTACAAACACTATCTTGAATACTTTTCAAGATAAAAATAATAAGACAAAAAAGAAAAAAGAAGTGAAATGTTTAAGTAATGAAGATAAATATTTTTATAATTTAGAAGAAAGAAGAAAATATTATAAATATAATTTTGATAAAAATATCAGAAAGTTATTTAACAATGTGGATACAAAAACGAGTGGTGAAGAATCAGCAAATAGTCTATTGAAAGAAAAATATAATAAATATCATAAGAGAAATAAAAAATATAAAAAATTGTTCGAATATATCAAACAATTAAAAGATCATATAAAAAAAAAAAGCGATCATATTAGAGAGGCAAATGAAGCATTTATAAGAAGTATAAAACAAAAGCAAGAGGACGAAAATAAAAAAAAAGAAGAGAATAAAAAAAAAGAAGAGAATAAAACAACTGATGGGGGTAAAACAACTGATGGGGGTAAAACAACTGATGAGGGTAAAATAATTGAAGAGGGTAAAATAATTGAAGAGGGTAAAACAACTGATGAGGGTAAAACAACTGATGAGGGTAAAACAATTGATGAGTGTAAAACAACTGATGAGTGTAAAACAACTGATGAGTGTAAAACAACTGATGAGTGTAAAACAACTGATGAAGGTAAAACAATTGAAGAGTGCAAAATCATTGAAGATAATAAAAAGGGTAGTGAAGATTACAATAAAAATGAAGAGAATAAATCGAAAGAGGAAAAAACACCAAACGATGACAAATACGAATTTGTTCAAGATACTGAAGAGCTCATAAAAAAGGTTACGAAAAAAGATCAAGAGTTGTTGAAAAAATTAATAGAATATAAATACCAAATATTAAAAAAACAAAATAATAACCTTATGAAAAAAAAAAATAAAGAAAATATATATATGAACGATTATTCAGATAATATAGAGATGTTTAGTGATACTGAAACGGATTCTACTGATTATTATGATGAAGATGATGCAGATTATGATGAACATATGGTTGAATATAATAATTATCATGATGAAAGTAATCAGAATACAAAAGATTATATATTTTATGATAAGAAAAAAACGAGAAGAGAACTAAAAGAATTACGACGTTTTAATGAAAGGAAGAAAAGGTTAGAAACATATTTAATTATGAATAAGTTATATAATCAAAATAAAAAAGGAGTGATATCAAAAGAATATCATACAATTAATAAAAAAACGGATGTATATTGGCGACTAGCATTTGCAAGAAGAAATATCAATGTATTAGAAAACGTAACA**CAAGAGGATGTGAATAATCAATTAGAAATGTTAAAGTCAAATGGGAAGGTACAAAAAAAGGGAGCAGCACATAATAAGGAAAATGAAAATGAGGAAAAAAATAAAAATAATAAAAATAATTATATTCATAATAGTAGTAATAATATAAAGAATGATCAAGAGAAATCATCACTAAATGATAATAAAAATTTTGATAACAAACTTAAAATGTATGTAGATAATTTTAATATTAAACAATATAATAAATATATATCATTATTCCAATTAATGCAATCGAAACCATATATTTTAGATCCATTAATTTATGATTCAAAAAAAATTTTGGATAAATTTATACCAATCAATGGTAGATTTACAGCAAAAGCATTATGTAAATTATTTGCATTTGTGAATAACGAATTTTTTTTTCCATCTTATATTATTAATAAAATAAGAAAGAAATATACTATTGATAAAAGTGTAGAGGCATTGATTTTAACAGGAGCTATGAGTAGAACATGGGGTATGGGTTTTCAACTTTTTGAATGTGAATATAATGCTAATATTAATGATGAATTTAATTTATATAAAAAAAAAAATAAATATAAAAAAAAAAATGTAGAATCTAGAAAAAAAAAAAAAATAAATAAAAAAAAAAAAAAAATTGTTGGATATGGACAATCCGATTTTTCAGGATGTTTGGCTATATCATTTCCAGAAATTGATTTATCTATCACAATACTTTTATCTGACGTTTTTAAAGGAGCAGAC*gtaatagaaaatgaaaataattcaacaattatgtgttatatatgatatatgttaagtctcaatatttatttaatttatttaatttatttaatttatttaatttattcattttttttttttttttttttttctttcttttctatcgttttcttag*GTGTGTCATCTCTTATTAGAATTTATATTAAAACTATACGGATTGAAACCTCAATGGAAAGTGCCTGTCAAAATGTCTGAACTTGTCAAAGTTTTT**TAAaattaaaaaaaatgacagaaaaaagaaagcatcttttttgtcgtcctacaattcgtaatacacacacatatatatatatattatatatatgtgtttatttcttcctaatttttttagttgtacttttttttttttttattattatatatatttttttttacccagtagacacaggaatatactcttaataattcccctttattttttaatttgtcaaattatacaaatttatttttaacatagaattttataaaattattatactagaaagagatttattttattttattttatttttttatttatatttatttatttattttttatttccgacaaaaagaaatatatggctaaattaaatataaatagcgtaaatgttatgttgcatatataaatcattaagataataatttatttgattcatatactatatatatgtgttatttaatatgaaacagttttttttttttaattttaatatttttcgttgttttattttatttgtttttatttcctattttctgtttttttttttttttttttttattccttgcgtttaaaatgtttttataataaatacataaataaaattcatattaataatttaaataataaatatatatatatatatatttatatatatatgtattttttaaaacatataaaacattatatatattatatatatatatatatatattgatatatatatatatgtatatattatatatatttaactaaattttaaacatatacatgtctattttatataaaactatataaaaatatgtataatatatatatatatatatatatataatggaaaaaaaaaaaaaaaaaaatagaaaaaagaacagataagaaaagaaaaatactacttttatagataatatttgttttgttcaaaaaaaaaaaaaaatatatatatatatatataaataaataaataaataaaaaaaaaaaaaatatatatatatatatataaataaataaataaataaaaaaaaaaaaaatatatatatatatatttgagtaaacaattatgaaaaaatatatacattaatatatatatatatatatatatatatatattatgtatatacataaacatgcaaaaacaatatcttataacaaatataaattaaaaaaaaaaaaaaaaga

**Original Locus (OL) amplicon size: 1114bp**

**> PF3D7_1414500-HALO-SW Halo Neo-R**

aatattattatatctttattacaaaataataatagaaaaaatagaaataatattgtggatattttttaatacatacatacatatataatattatatttatatataaaatatatacatagataatacatatatatatttatttatctatatatgctggattaatattaatatatatatatatattatatatatatatatatatatatatgtaataaatatatatgttcgtcatataaaataactaaaaaaaataaaaaaaaaaattaaaataaaataaaaataattaatatttaattgaaatgtttatacatttttcagaacatcaactaaatatatgaatatatatgtataatatatataatacatatgtatattatatatatatatatatataaatataattatgtatttttgtatgtttaatattatttttttcgtttattctatatatgaatatattattatataacatattaatgttttaattttcatgtttgatatagaattcaatcggtattttaattcttgtattttaattgaatgaatatatatatatatatatatatatataatatatatatatttatatttattaatatgtttgtaaatatgataatatggatcaatcttttcatatataaaaaaaaaaaaaaaatttaccattatataaatattcttactttatacattatgtagttaaaagacttaaaatatatgcattttggtattctttttttttttttttttctatttagttttgttttacatgtaaataaacttaaatttgattatttttttattttaatttttattatacttttttataaatgtgtgtagttattactttatttttttacatacatatatatatatatatatatatatatatgtatatatttttacaatctgaatgccttaagtttatttgatacatatatttccatttttttttggaaaataaaaaagataaaaatataaaaatttttgataacgtataaattgtttttttttttttttttaatatatattaaaggatgcggctttgattatatatatatatatataatataatgtgaaaaagtctaataaaagaaaaaaaaaaaaaaaaaaaaaaaaagaagtaaaataatatatatatatttatgaaaaaaatcttcagatataaatatgttttatcttattttattaatttatagtgtacatattaataatatattatataatatatgattatgtagaaattaagaaaaaacaaaaaaaaataaaacaaatactttttttatgtgattttattttattctatgttaaagaatgatatacaaatttacattatacttctgctgaatttgtaaaaatttataattaatgtaatgatttctttgatttattatttctttcaaatttttatattttactgtgcatatttataatattataatgtaaatgcaataaatatattttatatatatatatatatatatatatatatatatatatatacatttatatttttttatattcctgttacacataatatataaataaatattttcacatttgttatattttttattttattttacttccatttatttatttatattctttcttttttttcttttttttgtttttacatattttaaaaataatataccgttatgtaaaaaatgtgttcattcatataaacggtttctcattatatatataataaatatatataaatttttataaatttatatatttatcatataaatgaatatacacatatatatatatatatatatatatatatatatatatatatatttttatttatatatgattaaaaaaaaaaagaaaaaattgtatgatatggtataattacaatactgaagtaattttttttgtaaatttttatatcatttattgttttaatttatttatttatttatttttttttttttgaaaaaaaataataatatgttataaaagtagacataatataaattagttgttaatttttatatataatattacatcatattgtttatatttgttttgttggttttattaccttttttttttttctttttttttttctttttttttctatataaaaatttataacaatatatttgtttacttaaaaataatatatatatatatatatatatatatatgtatatatttttgaagagggaaaaatatggtgttcttacattcttttcatttttatcctttttaaaaatttattaagtaaaccaattgataaatgtgatcacatattttggtgtgtttcactttttttttttatttctaaaaaaaagaaaagaaaaaaagaactttttatatttaattggtacttgttcttttaatagaataagttgtgtgcaaatcttatagaaaaATGAATCAATATGATCAATGGAATCGTAGAATTCGG*gtaggaaaaaaaaaaaaaaataaagaaaaataaaattaaatatatatactttgttgaattaattatattatattgttttattttattttaattatttattttatttttttttttttttttccttttag*ACAATATGGTACTGTAGTAGTCTGTATGTAGAATATAAGAACGCTTTGAGGAAATCCAAAAAAATGCCGGTGGAGAAAAAAAATGAATACTGGGAAAAGAAGCATGAAGAATTTGCTACAAAGATGTTAAATAATATATATGAATTAAGAG*gtaaggaatattattataaaagttaaaaaaaaaaaaaaaaaaaaaaaaaataatatatataaatataaatatataatacatgaatgcgttgtctatttttcatattttataaaaaaataccactacacgtttatattatctttatttatttatttgtttgtttgtttattataaatag*GTTGGTGGGTTAAAGTAGGACAATTTTTAAGTACCCAGGAAAATATCATGCCTGTTGCTTATATAGAAAAATTTACGAAGCTGCAAGATATGATGCCTACGTCTTCATTTGATAAAATAGAAAATATTTTAAAAAAGGAATTAG*gtaatttaaaaaaaaaaataaaaaataaatgtgtctatgtatgcataaaagtatgagtatatgtatgtataaatttatgtacaaatgtatgcgtatatgtatgtataaatttatgtacaaatgtattcgtatatgtatgtataaatttatgtacaaaagtatgcgtatatgtatatataaatatatgtattttttttttttttttgtttgattcctag*GTAACATTTATGAAATGTTTGAATACATCGATAAAGAACCACTAGCTAGCGCTTCCATTGGACAAGTACATAAAGCGAAATTAAAAAAATATGATAAACATATATGCGATATAGACAGTAGAAAGCATAAGGATTATAATGTAATAATAAAAATTCAACATGAAGGAATAGATCAGTTTTTGTCATCAGATATAAGTACTTTAAAAAAAGTGTCTTGGGCATTTGGATTAATAGATAAGAATTTTTATTTTACAGATTTTATAGATGAATGGCAAGATTCTGCTTCTCGAGAATTAAATTATAAATATGAATTATATCATCAATTGTTAGCTTATAACAGTTTTAAAAAATCAGGTATCCCTGTAAAAATACCTAAAATTTATTGTGCTCATACAACTTCTAAGGTATTGGTTATGGAATATATTAAAGGCTTTAAAATAACTGATACTGAATTACTTAAAAAATATAATGTAGATACATATAAAATGGTCTATAAAATTATTGACTATTTTGCTTATCAAATACATAATGATGGTTTTTTTCATGGGGACCCACATCCGGGAAATATATTAGTCATGATGGAAGAAAAGAAAAAAAGAAAAAGGAAAAAAAAATATAATAAAGATAAAATAAATAACTCGAAAATGAAAATGAAAAATGAAAATATATATATGAACAAGAGAAAAAAAAATCATAAAAAAAGTAATAGGAATTCTAAGAAAAAGTTAAGAAGTTGTAAAAAAAAAAAACTAGAAATAAATAAAGAAAGAGAAAAAGGAGAAGAAAAGAAAAAAAATATGTATAATAATAATGATATAAATTATTGTGAAAATAAAAATGAAGATATTATAACAAACAATCCCAAATTACAAAATAATAGTAGTGATATGAAAAATATAATAAAATTTACAAGTTATTCCAAAAGTGTATCTTCATCTAATGCTTCATTGATGGATCATTTTGTGGATTCAAGATATAATATTGATAAAATGAGAGCTCCTAATAATTTTTCGAATCCTTTTCATAAATCTTTTAGTTTTGTTGATTATTCACTTAATCAAGATGTAGATTCCGATAAAATGAATCAAGAAGATTATGAATATATTTTGAAGGAAAATATGAGTGATGGTTTAGATAATATGAAAAATTGTCTCTCAATATCCCATGAATATATTTCATGCGATCAATCAAGAAGTGGAGTAACATCTATTGCGACAAGTAGGAAAACACAAAATAGGGATTCTAACAAAGCTAGTGAAAATAGTATTATGGAATCTCGAGAAACAACGATGTTTTCTTTTGTTAACAATATGGAGGAGTTAGTCAAGATGAAATCCGAGACTAAAACGGATCTACAAATTAAAGGTAATGATAAAAAAGTGGTTAAAGGTAGTAAGTGTGACAATAAAAATGATCAAAATGTTAATAATGATAATATTAATAATGATAATATTAATAATGATAAAATTAATAATGATAATATTAATAATGATAATATTAATAATAATAATATTGTTAATAATAATAACAATATTTCTAATGGTGAGAAAGAGAAAAACAAAAGTGGTCCGATATACAAATCCAAATTCAAACTATCAGATTTAAAAAAAAGAAAAAATGCAAAATATAATTTTGTACCTGTTATTATTGATTGGGGTTTAATTAAACAATTAGATAGCGTAATGAAATTCGCCTTTTGTAAATTAGTTTATAATATTAGTTGTATGAATGTATTAAATATTATAGAAGCCTTTGAAGATATGGGTTTCTGCTTTAAAGAAGATTTTACATACGATCCTGAAATATATATTGAGAATTTGAAAACATACTTTTTAAGCAAATTAGAAGAATCTAAAAGTAAAGGTAATGAAGGAGAAAAGGAATTTGGTAGTGCTAATACAATTGATGAAGCAAATAAGAGTAATAAGAATATGGAAATATTAAAAAATATTGAAAAAAAAGATGTTATGGATAAAAATCCGATAAGTGATGTACCTAAAGATATTATATTTTTTATGAGAGTAGCATCATTATTACATGGATTATGTACACAATTAAATGTAAAAATTAATTATTTAAATATTTTTTCCAGAAGAGCTAAAGAAGCTTTAGAAAAAATATATATACCAATAAATAATAGTATACACACAATACCAATAGATAAATGTCCAAATACATTTGTAGAAAAAAGAATTCATCAATTTATAAAAACATTATATGAAAAAAAAAAAATATTAGGATGTCAAGTAGCCATTATACATAAAAAAAAGCTAGTTGTTAATACATGTGTAGGAATTACAAGTACAACAGATAAAAGGCCTATAACAAAACATTCTTTATTTAATGGATATTCATTAAACAAAATTATTTTAAATATAGCTCTATATCATCTTATATGTAACTCAGTAAATGAACAAACGTCAGGTGAAAGTATTCTGCTCGGTATGGATAAGAAAAAGGATAAGCTAGGGGAAAAGATAAGTCAAAAAATTCAAGAAAAAATAAATGAAAAGATAAAAATTAAATCAAGTGCGAAGGACAAAAATCAATCCGATGAAAAAATTAAAATGAAGATAAACGAAAAGGATAAGGATAAGGATAAGGGTAAAGGTAAGGAAAAGAATAAAAATAATAATAAGCAAAACAATGATAGGGATAAAAATAATAATGAGGAAAACAATAATAATGAGGACAATAATAAGGATAACAATAATAATAATAATAATAATAATAATAATAATAATAATAATAATAATAATAATAAATTATTAAACGAAATGGATCAAAGTAAACACAATGCAAATGAAAAAACATCCAATATGAAAAATAACAATAAACTTTATGAAGAAGAAGATGTAGAGGATGAAGTTGCTGATGATAATAAAACAGATAATGAAAATAATTATATTACCGAAAAATTAAATGAATTTAATAAAATAGAAAACGATTATTTTTCAGCAAATGTTAATGAAAATATAATTAAATGTGAAGAGTTTGGTGAGAGCAAGGATATTATAAGTCCTTTATCTGTCAATAGTTTCATAAGTTTTGATAGTTATAATAGTTATAATAATAATGTTAGTTGGATCAGTAGTATGAGTAGGAATAGTTACAAAAGTATGATTGAAAGGATAGAAGATGATGTGGAATATAAAGAAATTAATGATGATCTACCTAATGATAGTAAGGTTAGTTGTATGAGCGAATACAAAAAGGATGATAAATATAAAGGGAGAAATTATATACATGATCACAATAATGATGATGTACATCATCATAATAATCATAATAATCATAATAATCACAATCATCACCATAATCATCACCATAATCATCACCATAATCATCACCATAATCATCACCATAATCATCACCATAATCATCACCATAATCATCACCACCATCACCATCATGATAATAATATAAATAAAAGTAATAAAGATACAGAGTCCAATCCAAATAAATCCATAAATATATATCCAAATATATCATATAACAAAATGGATACCTTAATACAAAAAAAACAAAGTTATATTGATGAGAAAAACTTTATAAAAAGTATAGAAAGCAAACAAATAAGAAATTTCAAAAATATTATTAACGATCATATTTGTAATTACTGGGATGGATTTATATGTAATAATAAAAAAAATATAACGATAAAAGATATATTAACTTTAAAGTGTTTTATTAAGAAACCTTTTCATGATAAAATAACGTTAAGCAAATTTATAGATTACGATCAAATGATAAATATGATTGAGAACTCTAAGCATTATAAGAGTTCAGATAAGACGTCGAAATATGGAGAGTATTTATATTTAATCGATACGTATATTATATCTGAGTTAATACATAATATATCAGGATTACATTATTATGAGTATATATATAAATATATAGTGAAACCATTAAATTTGACTGAAGAAATGTTTGTACCTATACCTTCAGAATTATTAAAAAATCAAAAATACGAAGATTCAAAAAAGAAAAATGTCAACAATCAAAATAATGTAAATACCACTGGTGGTCTTAGTAATAATAATTTAGATAAAACAAAAAAATATATAAAGGATAAAAAGACAAATGTTATAGATAAGAAATCTATTGTTATGGATAAATGGCCTAACATGATGAAGGATTATACAACCGGTGCTTTAAAAAATAATGAATACAAGTCAACTTCATTTATTGGTGGTAAGAAATTTAAAAAGTCTAGTACTATTTCCAACTTTATAGACGATATAAATGATGATATGAATTTTTATATTATGAACGCGGAACTTTTAAAGTTTGAAAATATTAATTTAAAGAATACCATATCTAACAAAAAAAAATTTGATAGTAATCAAAAAGTTTTATCCAAAAAAAGAAGTATAAGTGTAGATGTGTATTATTCTAAGAAAAAAATTAAAGgtaacacaataaatagaataaaataagaacaagtgaatagataaaaaaaaaaaaataaaaaaaaaaataaacacatcaaaattgatatataaagaaaaaatatattcataaagatgtaatatgtaatatgtaatatgtaatatatatatatatatatatatatatatatatatttatttatttatttattttttgtttatttatagATGGTAGTAAGAAGGAGAAAAAAAAAAATAAGGATAAGAAAGATCAAAAGGGAAGTAAAACCAAAATGGATTTTGAAAAAAGAGACATTGCAGAAATGAAGAAAGTAGATAGGAATGAACAGAAAGACCAAACGAATCAACAGAAAGATCAAACGAATGAACAGAAAAACCAACCGAGTCAACAAAACGAACAAATGTATGATGACATAGTTGAAACGTATGAAGATATAATTGAAACATATGAAGACAAAATAGAACTAAACCATGATAATGATGATAGTAACATAAAAACGAATGATGTTTATCATGACATTATAGAAGAGAAAGAAGACAATGAATATTATAATAATATTAAGAATGAATATGAGAATAAAAGGAATAATAATAAAATTGAAATATCAAAGGATTCTTTAAATAAGAATCCATCTAATATAGAATGTAAAATGAATATTTTAATAAATAACGAACTTATAGATAATGAAAATGCAAGAAAATTAGATTCAAGTAATCTTCTTGAAAGTACCTTAAGGAATAAAGAATCCTTTAGTTTTAAAAATAAATTTCTGAATCATGTAGATAATTTAAAATTAAAAACGAAGAAAATAAATGATTCTATAAAAAATATGTATGAAAATAATGAGAATATTTTTTTGAAAAATTTCTTTCTATCATCTAAGAAATATGATAATATAGAAAATAACTTAGATCACAGAAATGTAGAGGCTAATTGTGTAGATAATAATATGTGTGAAAGTAATAGTCTGAATAATAGTTGTTATAATAATAGTTTTTATAATAGTTTTTATAATAGTCATAGTAATAGTAAAAATAGTAGTAAAATAAGAAACTACAATAATATGGATAAGGAAAATAATAATGTTAATAGTGATAATATAAATTTCAATCCCAATGATAATAAAAAGAAAAATAGTTATAATAATAATAATAATAATAATAATAACAATATTATGGTGAGTTCTAGCCCTTATATTTATAATTACAACAATAAATATATATTAAAAGAAAATTATGATAAAAAATTTAATATGAACAAATTTAATAAAGTAATGTTATCTAATCAAAAAGAACCAGTTCATAAAAATAATAAAAAAGATAATTTATTTTTATTTGAAGAAAGAGGTAAGAGATCACATAGTTATAGTGACTGTAGAAATGAATCACACCATGTAATGAAAAATTCTTATATGGAGAATATTAACAATATAAATAATTATAGTAATAGTAGTATGAATTACTTAAACTATTATATTAGTAATAAGAAAAGTGTAAATTATATGAATGCTAATGGGGTCGTCATAAATTATGATGATAATAATAATAATGTAAATAATGTTAATAATGTAAATAATATGAATAATATAAATAACAATGAAACTAATAATACTAACAAAATGAGCTTCTTTACAAACACTATCTTGAATACTTTTCAAGATAAAAATAATAAGACAAAAAAGAAAAAAGAAGTGAAATGTTTAAGTAATGAAGATAAATATTTTTATAATTTAGAAGAAAGAAGAAAATATTATAAATATAATTTTGATAAAAATATCAGAAAGTTATTTAACAATGTGGATACAAAAACGAGTGGTGAAGAATCAGCAAATAGTCTATTGAAAGAAAAATATAATAAATATCATAAGAGAAATAAAAAATATAAAAAATTGTTCGAATATATCAAACAATTAAAAGATCATATAAAAAAAAAAAGCGATCATATTAGAGAGGCAAATGAAGCATTTATAAGAAGTATAAAACAAAAGCAAGAGGACGAAAATAAAAAAAAAGAAGAGAATAAAAAAAAAGAAGAGAATAAAACAACTGATGGGGGTAAAACAACTGATGGGGGTAAAACAACTGATGAGGGTAAAATAATTGAAGAGGGTAAAATAATTGAAGAGGGTAAAACAACTGATGAGGGTAAAACAACTGATGAGGGTAAAACAATTGATGAGTGTAAAACAACTGATGAGTGTAAAACAACTGATGAGTGTAAAACAACTGATGAGTGTAAAACAACTGATGAAGGTAAAACAATTGAAGAGTGCAAAATCATTGAAGATAATAAAAAGGGTAGTGAAGATTACAATAAAAATGAAGAGAATAAATCGAAAGAGGAAAAAACACCAAACGATGACAAATACGAATTTGTTCAAGATACTGAAGAGCTCATAAAAAAGGTTACGAAAAAAGATCAAGAGTTGTTGAAAAAATTAATAGAATATAAATACCAAATATTAAAAAAACAAAATAATAACCTTATGAAAAAAAAAAATAAAGAAAATATATATATGAACGATTATTCAGATAATATAGAGATGTTTAGTGATACTGAAACGGATTCTACTGATTATTATGATGAAGATGATGCAGATTATGATGAACATATGGTTGAATATAATAATTATCATGATGAAAGTAATCAGAATACAAAAGATTATATATTTTATGATAAGAAAAAAACGAGAAGAGAACTAAAAGAATTACGACGTTTTAATGAAAGGAAGAAAAGGTTAGAAACATATTTAATTATGAATAAGTTATATAATCAAAATAAAAAAGGAGTGATATCAAAAGAATATCATACAATTAATAAAAAAACGGATGTATATTGGCGACTAGCATTTGCAAGAAGAAATATCAATGTATTAGAAAACGTAACA**CAAGAGGATGTGAATAATCAATTAGAAATGTTAAAGTCAAATGGGAAGGTACAAAAAAAGGGAGCAGCACATAATAAGGAAAATGAAAATGAGGAAAAAAATAAAAATAATAAAAATAATTATATTCATAATAGTAGTAATAATATAAAGAATGATCAAGAGAAATCATCACTAAATGATAATAAAAATTTTGATAACAAACTTAAAATGTATGTAGATAATTTTAATATTAAACAATATAATAAATATATATCATTATTCCAATTAATGCAATCGAAACCATATATTTTAGATCCATTAATTTATGATTCAAAAAAAATTTTGGATAAATTTATACCAATCAATGGTAGATTTACAGCAAAAGCATTATGTAAATTATTTGCATTTGTGAATAACGAATTTTTTTTTCCATCTTATATTATTAATAAAATAAGAAAGAAATATACTATTGATAAAAGTGTAGAGGCATTGATTTTAACAGGAGCTATGAGTAGAACATGGGGTATGGGTTTTCAACTTTTTGAATGTGAATATAATGCTAATATTAATGATGAATTTAATTTATATAAAAAAAAAAATAAATATAAAAAAAAAAATGTAGAATCTAGAAAAAAAAAAAAAATAAATAAAAAAAAAAAAAAAATTGTTGGATATGGACAATCCGATTTTTCAGGATGTTTGGCTATATCATTTCCAGAAATTGATTTATCTATCACAATACTTTTATCTGACGTTTTTAAAGGAGCAGAC*gtaatagaaaatgaaaataattcaacaattatgtgttatatatgatatatgttaagtctcaatatttatttaatttatttaatttatttaatttatttaatttattcattttttttttttttttttttttctttcttttctatcgttttcttag*GTGTGTCATCTCTTATTAGAATTTATATTAAAACTATACGGATTGAAACCTCAATGGAAAGTGCCTGTCAAAATGTCTGAACTTGTCAAAGTTTTT**CCTAGGTCAGGATTGAGATCAAGATCTGCTGCTGCTGGTGCTGGTGGTGCTGCTAGAGCTGCTCTGCAGAGAGGAGTACAAGTTGAAACAATATCACCAGGAGATGGTCGTACATTTCCAAAAAGAGGTCAAACTTGTGTTGTACATTATACTGGAATGCTTGAAGATGGAAAGAAATTTGATTCATCTCGTGATAGAAATAAACCATTTAAATTTATGCTAGGTAAACAAGAAGTAATACGAGGTTGGGAAGAAGGAGTTGCTCAAATGAGTGTAGGTCAAAGAGCAAAACTTACTATATCTCCAGATTATGCTTATGGTGCAACTGGACATCCAGGTATAATTCCACCTCATGCAACTCTTGTATTTGATGTGGAGCTTCTAAAACTAGAAACTAGAGGTGTTCAGGTTGAAACAATTTCACCTGGAGATGGCAGAACCTTTCCTAAAAGAGGACAGACTTGCGTAGTTCATTATACAGGCATGCTAGAGGATGGTAAGAAATTTGATTCTAGTCGAGATAGAAATAAGCCATTCAAGTTTATGCTAGGTAAACAGGAAGTAATAAGAGGTTGGGAAGAGGGTGTAGCACAGATGTCAGTTGGACAAAGAGCAAAGTTAACAATATCACCAGATTATGCATACGGTGCAACAGGCCATCCTGGCATCATCCCTCCACATGCAACTTTAGTATTCGACGTTGAATTGTTAAAGTTAGAGACAACGCGTGCTAGAGGTGCTGCTGCTGGTGCTGGAGGTGCAGGTAGACGTACGATGGCAGAAATTGGTACGGGTTTTCCATTTGATCCTCATTATGTGGAGGTGCTGGGGGAAAGGATGCATTATGTTGACGTAGGACCAAGAGATGGTACTCCAGTGTTATTTTTGCATGGAAACCCAACCTCGAGTTATGTATGGAGAAATATAATTCCACATGTAGCACCAACACATAGATGTATAGCTCCTGATTTAATTGGTATGGGAAAAAGTGATAAACCTGACTTAGGATATTTTTTTGATGATCATGTCCGTTTTATGGATGCTTTCATTGAAGCTCTTGGCCTTGAAGAAGTAGTATTAGTTATACATGATTGGGGATCCGCCTTAGGATTTCATTGGGCCAAGAGGAATCCTGAAAGAGTAAAAGGAATAGCATTCATGGAATTCATACGACCAATCCCCACATGGGATGAATGGCCAGAATTTGCACGCGAAACATTTCAAGCTTTTAGAACTACAGATGTTGGTAGAAAATTAATAATAGATCAAAATGTATTTATAGAAGGAACTTTACCTATGGGTGTTGTAAGGCCGTTAACAGAAGTTGAAATGGACCACTACCGTGAACCTTTTTTAAATCCAGTAGATAGAGAGCCCTTATGGAGATTTCCTAATGAATTACCTATTGCAGGTGAACCCGCGAATATTGTTGCTTTAGTAGAAGAATATATGGATTGGTTACATCAGTCTCCTGTTCCTAAACTTCTATTTTGGGGTACACCTGGAGTTCTAATACCACCAGCTGAAGCAGCAAGATTAGCAAAATCATTACCAAATTGTAAAGCTGTTGATATAGGTCCTGGGTTGAATTTATTACAAGAAGATAATCCAGATTTGATTGGATCTGAGATAGCTAGATGGCTAAGTACATTAGAAATTTCAGGTACCGGTGCCAGGGGAGCAGCCGCAGGAGCAGGGGGGGCAGGAAGGCGTGGTGTTCAGGTCGAGACTATTAGCCCTGGAGATGGACGCACGTTTCCTAAGCGTGGACAGACATGCGTAGTTCACTACACAGGTATGTTGGAGGACGGTAAAAAGTTCGACAGCTCACGCGACCGCAATAAACCTTTCAAGTTTATGCTTGGCAAGCAGGAGGTTATTCGTGGATGGGAGGAGGGTGTAGCACAGATGTCTGTTGGACAGCGTGCTAAGTTGACAATTTCACCTGACTATGCTTATGGCGCTACGGGCCATCCCGGGATCATTCCGCCACATGCGACTCTGGTATTCGACGTTGAATTATTAAAGTTAGAGACAGCTAGAGGGGCCGCTGCAGGTGCTGGTGGAGCTGGAAGACGTGGAGTACAAGTAGAGACTATCTCTCCAGGTGACGGTCGCACTTTCCCAAAGCGTGGCCAAACCTGTGTTGTACATTACACTGGTATGCTGGAGGATGGGAAAAAGTTCGATTCCAGTCGCGACCGTAACAAACCGTTCAAATTCATGTTGGGAAAGCAGGAAGTGATCCGCGGGTGGGAGGAAGGCGTGGCGCAAATGAGCGTCGGTCAGCGGGCTAAATTGACCATTTCCCCTGACTACGCGTATGGGGCTACTGGGCACCCAGGGATTATTCCGCCTCACGCTACACTTGTGTTTGATGTCGAACTTTTGAAACTGGAAACTGTCGACGGAGAAGGAAGAGGAAGTTTATTAACATGTGGAGATGTAGAAGAAAATCCAGGACCAATGATTGAACAAGATGGATTGCACGCAGGTTCTCCGGCCGCTTGGGTGGAGAGGCTATTCGGCTATGACTGGGCACAACAGACAATCGGCTGCTCTGATGCCGCCGTGTTCCGGCTGTCAGCGCAGGGGCGCCCGGTTCTTTTTGTCAAGACCGACCTGTCCGGTGCCCTGAATGAACTGCAGGACGAGGCAGCGCGGCTATCGTGGCTGGCCACGACGGGCGTTCCTTGCGCAGCTGTGCTCGACGTTGTCACTGAAGCGGGAAGGGACTGGCTGCTATTGGGCGAAGTGCCGGGGCAGGATCTCCTGTCATCTCACCTTGCTCCTGCCGAGAAAGTATCCATCATGGCTGATGCAATGCGGCGGCTGCATACGCTTGATCCGGCTACCTGCCCATTCGACCACCAAGCGAAACATCGCATCGAGCGAGCACGTACTCGGATGGAAGCCGGTCTTGTCGATCAGGATGATCTGGACGAAGAGCATCAGGGGCTCGCGCCAGCCGAACTGTTCGCCAGGCTCAAGGCGCGCATGCCCGACGGCGAGGATCTCGTCGTGACCCATGGCGATGCCTGCTTGCCGAATATCATGGTGGAAAATGGCCGCTTTTCTGGATTCATCGACTGTGGCCGGCTGGGTGTGGCGGACCGCTATCAGGACATAGCGTTGGCTACCCGTGATATTGCTGAAGAGCTTGGCGGCGAATGGGCTGACCGCTTCCTCGTGCTTTACGGTATCGCCGCTCCCGATTCGCAGCGCATCGCCTTCTATCGCCTTCTTGACGAGTTCTTCTAActcgagggatatggcagcttaatgttcgtttttcttatttatatatttataccaattgattgtatttataactgtaaaaatgtgtatgttgtgtgcatatttttttttgtgcatgcacatgcatgtaaatagctaaaattatgaacattttattttttgttcagaaaaaaaaaactttacacacataaaatggctagtatgaatagccatattttatataaattaaatcctatgaatttatgaccatattaaaaatttagatatttatggaacataatatgtttgaaacaataagacaaaattattattattattattatttttactgttataattatgtgtctccttcaatgattcataaatagttggacttgatttttaaaatgtttataatatgattagcatagttaaataaaaaaagttgaaaaattaaaaaaaaacatataaacacaaatgatggtttttccttcaatttcgatatcaatttatagaaacaaaatatatacttgtataattttatttttttatataaatcattacatatataattatacaatattttttctaagagataattatatattaatatatataaaaaaaggtgttttttttttttttttttatttttatttttattttatggtaatattttattttccttattttataaattatattagtttatatgtgattaattttatatattatcaatttatatatttttaaatgcttacttaattatctttttttttttttttttttttttttcccctctttttatattaatttatttttgaaaaaattgatatatatatatatatataatatatatatatacatgtagtagtattaaacaatgtataatatatataaataatatatttatatatttcatttcaattttaattttttttggttttttttttttttctttttgtcatatttaaaaaaaattatattcatataagttatgcattttttataaacattattcaatatatgtataatataatatatatatatatattaatgtattattccaatgtgcatgataaaagaaaaaaataatatttataaaaaaaaagaaaaataaaacaaaaaaagaaaaaaaaaaaaaaaaaaaaaaaaatacaaaaataaataatataatttataattatatattcttgtcacaataaaaatatatatatatatatatatatttataatatgtatattttaaactagaaaaggaataactaatattttatttattatcattcaagatttatattttataataataaatacctaatagaaatatatcaggatccatgcatggttcgctaaactgcatcgtcgctgtgtcccagaacatgggcatcggcaagaacggggactacccctggccaccgctcaggaacgaatttagatatttccagagaatgaccacaacctcttcagtagaaggtaaacagaatctggtgattatgggtaagaagacctggttctccattcctgagaagaatcgacctttaaagggtagaattaatttagttctcagcagagaactcaaggaacctccacaaggagctcattttctttccagaagtctagatgatgccttaaaacttactgaacaaccagaattagcaaataaagtagacatggtctggatagttggtggcagttctgtttataaggaagccatgaatcacccaggccatcttaaactatttgtgacaaggatcatgcaagactttgaaagtgacacgttttttccagaaattgatttggagaaatataaacttctgccagaatacccaggtgttctctctgatgtccaggaggagaaaggcattaagtacaaatttgaagtatatgagaagaatgattaagcttatttaataatagattaaaaatattataaaaataaaaacataaacacagaaattacaaaaaaaatacatatgaattttttttttgtaatcttccttataaatatagaataatgaatcatataaaacatatcattattcatttatttacatttaaaattattgtttcagtatctttaatttattatgtatatataaaaataacttacaattttattaataaacaatatatgtttattaattcatgttttgtaatttatgggatagcgattttttttactgtctgtatttttcttttttaattatgttttaattgtattttatttttattattgttctttttatagtattattttaaaacaaaatgtattttctaagaacttataataataataaatataaattttaataaaaattatatttatcttttacaatatgaacataaagtacaacattaatatatagcttttaatatttttattcctaatcatgtaaatcttaaatttttctttttaaacatatgttaaatatttatttctcattatatataagaacatatttattaaatctagaattctatagtgagtcgtattacaattcactggccgtcgttttacaacgtcgtgactgggaaaaccctggcgttacccaacttaatcgccttgcagcacatccccctttcgccagctggcgtaatagcgaagaggcccgcaccgatcgcccttcccaacagttgcgcagcctgaatggcgaatggcgcctgatgcggtattttctccttacgcatctgtgcggtatttcacaccgcatatggtgcactctcagtacaatctgctctgatgccgcatagttaagccagccccgacacccgccaacacccgctgacgcgccctgacgggcttgtctgctcccggcatccgcttacagacaagctgtgaccgtctccgggagctgcatgtgtcagaggttttcaccgtcatcaccgaaacgcgcgagacgaaagggcctcgtgatacgcctatttttataggttaatgtcatgataataatggtttcttagacgtcaggtggcacttttcggggaaatgtgcgcggaacccctatttgtttatttttctaaatacattcaaatatgtatccgctcatgagacaataaccctgataaatgcttcaataatattgaaaaaggaagagtatgagtattcaacatttccgtgtcgcccttattcccttttttgcggcattttgccttcctgtttttgctcacccagaaacgctggtgaaagtaaaagatgctgaagatcagttgggtgcacgagtgggttacatcgaactggatctcaacagcggtaagatccttgagagttttcgccccgaagaacgttttccaatgatgagcacttttaaagttctgctatgtggcgcggtattatcccgtattgacgccgggcaagagcaactcggtcgccgcatacactattctcagaatgacttggttgagtactcaccagtcacagaaaagcatcttacggatggcatgacagtaagagaattatgcagtgctgccataaccatgagtgataacactgcggccaacttacttctgacaacgatcggaggaccgaaggagctaaccgcttttttgcacaacatgggggatcatgtaactcgccttgatcgttgggaaccggagctgaatgaagccataccaaacgacgagcgtgacaccacgatgcctgtagcaatgccaacaacgttgcgcaaactattaactggcgaactacttactctagcttcccggcaacaattaatagactggatggaggcggataaagttgcaggaccacttctgcgctcggcccttccggctggctggtttattgctgataaatctggagccggtgagcgtgggtctcgcggtatcattgcagcactggggccagatggtaagccctcccgtatcgtagttatctacacgacggggagtcaggcaactatggatgaacgaaatagacagatcgctgagataggtgcctcactgattaagcattggtaactgtcagaccaagtttactcatatatactttagattgatttaaaacttcatttttaatttaaaaggatctaggtgaagatcctttttgataatctcatgaccaaaatcccttaacgtgagttttcgttccactgagcgtcagaccccgtagaaaagatcaaaggatcttcttgagatcctttttttctgcgcgtaatctgctgcttgcaaacaaaaaaaccaccgctaccagcggtggtttgtttgccggatcaagagctaccaactctttttccgaaggtaactggcttcagcagagcgcagataccaaatactgtccttctagtgtagccgtagttaggccaccacttcaagaactctgtagcaccgcctacatacctcgctctgctaatcctgttaccagtggctgctgccagtggcgataagtcgtgtcttaccgggttggactcaagacgatagttaccggataaggcgcagcggtcgggctgaacggggggttcgtgcacacagcccagcttggagcgaacgacctacaccgaactgagatacctacagcgtgagctatgagaaagcgccacgcttcccgaagggagaaaggcggacaggtatccggtaagcggcagggtcggaacaggagagcgcacgagggagcttccagggggaaacgcctggtatctttatagtcctgtcgggtttcgccacctctgacttgagcgtcgatttttgtgatgctcgtcaggggggcggagcctatcgaaaaacgccagcaacgcggcctttttacggttcctggccttttgctggccttttgctcacatgttctttcctgcgttatcccctgattctgtggataaccgtattaccgcctttgagtgagctgataccgctcgccgcagccgaacgaccgagcgcagcgagtcagtgagcgaggaagcggaagagcgcccaatacgcaaaccgcctctccccgcgcgttggccgattcattaatgcagctggcacgacaggtttcccgactggaaagcgggcagtgagcgcaacgcaattaatgtgagttagctcactcattaggcaccccaggctttacactttatgcttccggctcgtatgttgtgtggaattgtgagcggataacaatttcacacaggaaacagctatgaccatgattacgccaagctatttaggtgacactatagaagcggccgcTAgCAAGAGGATGTGAATAATCAATTAGAAATGTTAAAGTCAAATGGGAAGGTACAAAAAAAGGGAGCAGCACATAATAAGGAAAATGAAAATGAGGAAAAAAATAAAAATAATAAAAATAATTATATTCATAATAGTAGTAATAATATAAAGAATGATCAAGAGAAATCATCACTAAATGATAATAAAAATTTTGATAACAAACTTAAAATGTATGTAGATAATTTTAATATTAAACAATATAATAAATATATATCATTATTCCAATTAATGCAATCGAAACCATATATTTTAGATCCATTAATTTATGATTCAAAAAAAATTTTGGATAAATTTATACCAATCAATGGTAGATTTACAGCAAAAGCATTATGTAAATTATTTGCATTTGTGAATAACGAATTTTTTTTTCCATCTTATATTATTAATAAAATAAGAAAGAAATATACTATTGATAAAAGTGTAGAGGCATTGATTTTAACAGGAGCTATGAGTAGAACATGGGGTATGGGTTTTCAACTTTTTGAATGTGAATATAATGCTAATATTAATGATGAATTTAATTTATATAAAAAAAAAAATAAATATAAAAAAAAAAATGTAGAATCTAGAAAAAAAAAAAAAATAAATAAAAAAAAAAAAAAAATTGTTGGATATGGACAATCCGATTTTTCAGGATGTTTGGCTATATCATTTCCAGAAATTGATTTATCTATCACAATACTTTTATCTGACGTTTTTAAAGGAGCAGACgtaatagaaaatgaaaataattcaacaattatgtgttatatatgatatatgttaagtctcaatatttatttaatttatttaatttatttaatttatttaatttattcattttttttttttttttttttttctttcttttctatcgttttcttagGTGTGTCATCTCTTATTAGAATTTATATTAAAACTATACGGATTGAAACCTCAATGGAAAGTGCCTGTCAAAATGTCTGAACTTGTCAAAGTTTTTTAAaattaaaaaaaatgacagaaaaaagaaagcatcttttttgtcgtcctacaattcgtaatacacacacatatatatatatattatatatatgtgtttatttcttcctaatttttttagttgtacttttttttttttttattattatatatatttttttttacccagtagacacaggaatatactcttaataattcccctttattttttaatttgtcaaattatacaaatttatttttaacatagaattttataaaattattatactagaaagagatttattttattttattttatttttttatttatatttatttatttattttttatttccgacaaaaagaaatatatggctaaattaaatataaatagcgtaaatgttatgttgcatatataaatcattaagataataatttatttgattcatatactatatatatgtgttatttaatatgaaacagttttttttttttaattttaatatttttcgttgttttattttatttgtttttatttcctattttctgtttttttttttttttttttttattccttgcgtttaaaatgtttttataataaatacataaataaaattcatattaataatttaaataataaatatatatatatatatatttatatatatatgtattttttaaaacatataaaacattatatatattatatatatatatatatatattgatatatatatatatgtatatattatatatatttaactaaattttaaacatatacatgtctattttatataaaactatataaaaatatgtataatatatatatatatatatatatataatggaaaaaaaaaaaaaaaaaaatagaaaaaagaacagataagaaaagaaaaatactacttttatagataatatttgttttgttcaaaaaaaaaaaaaaatatatatatatatatataaataaataaataaataaaaaaaaaaaaaatatatatatatatatataaataaataaataaataaaaaaaaaaaaaatatatatatatatatttgagtaaacaattatgaaaaaatatatacattaatatatatatatatatatatatatatatattatgtatatacataaacatgcaaaaacaatatcttataacaaatataaattaaaaaaaaaaaaaaaaga

**5’ amplicon size: 1194bps**

**3’ amplicon size: 1145bps**

## **> PF3D7_1316600**

ttttttttattttgttttgactaaaatatgcactaaaattttttttgaataatattattaattgtagtaaatattaattataattttttgtacattttgtgttttataacacattagtcataaaatagattatatatatatatatatatatatatatatataataccatattgattatatataatattatatatatatgtatatatctaattattaactaattatttaaatatatatattttatattaacttcatattataaatgtataataaagcgtttcccttttttttaattattaaaatcatgagataatataaaATGATAATTAAGGTTAATTCGGTTCAAACACAGAAG*gtaaaaattaaaattattaagataaatattaatactggaggttatacatatatatatatatatatatataatatatatataatattaaggagaattatattgtgcgaatacgtaaaatgaagttgaactattataaattacttagaaaaaaaggggtgcaatattgtattgaataaaaaatatataaagacccctaatgtgaatacagctatatagtaaattacacatatatattatatatatatatatatatatatatatatatatatatttatttatttatttatttgttgtacttttgaacattttaattttag*AGCCAAGATGGTAACAAAGTTCATAATAGTCAAGATGATATGGATAATGAGGAAAAAAGCATAAGGATATATGCAGATG*gttaaaaaatataaagaatatatgaaacatgacatatatatatatgtgtatatattttcttatttttttcttaattttttttgttggtttttccttctttttag*GTGTTTATGACCTGCTACATTTGGGTCATATGAAACAGCTTGAGCAAGCCAAACATGTAGATAAGAATGTTACCCTAATTGTGGGTGTTACAGGAGATAATGAGACTAGAAAGTTTAAAGGTCAAATTGTTCAAACTCTAGAAGAAAGAACAGAGACTTTAAAACATATACGATGGGTAGATGAAATAATTTCTCCATGCCCATGGGTGATTACTCCTGAATTTATGGAGGAACATAAAATAGATTATGTTGCACATGATGATATACCATATGCAAATAATCAGAAAAAGAAGAAAAAAAAAAAGTCTAAAGGGAAATCATTTAGTTTCGATGAAGAAAATGAAGATATATATGCTTGGTTAAAAAGAGCAGGGAAATTTAAAGCAACACAAAGAACAGAAGGTGTATCAACTACAGATTTAATTGTAAGAATATTAAAAAATTATGAGGATTATATTGAAAGGTCATTACAACGTGGTATACATCCTAATGAATTAAATATTGGTGTAACCAAAGCTCAGTCAATAAAAATGAAAAAAAATTTAATACGTTGGGGAGAAAAAGTAACAGATGAATTAACGAAGGTAACTTTAACAGACAAGCCATTAGGTACTGATTTTGATCAGGGTATTGATATAATAAGAGATAAAGTACATGATTTATTTAAATTATGGAGATATCATTCAAAAAAACTTTTAAAAGATTTTGCAAAATCATTTGATCCAATGTTTATTATAATTAGAAAAAGATATAGGAAAGATAATTTATCAGCTATGTACTTATCTGATTCAAATTATTTTTCTAGTTCTATGAAAGATGAATTAAAAAAAAAGAAAAGTTTAAGTAGTACATTTAATAATATTGATGAATATTATTATTCAGCTGATGAAGAAGATACACGGGATAATAGTTTTTATCGAGTTATAAATAAAATTGCTAATCATACTTATTATAGTGAAAAAGAAAATTATGAAACCTGTTTTGAATTAGAACATTCTTTAAGAGATAAAAAAAAAAGTTTAATATTTAAAGATAAAAATATGAAATTATCACCTGATGATTTATTACATTTTACTAATCCTATGAATTCAAAGTATGATCATAATTATTATTATTATAATAATACATATGATAAATTAAGTAACGTAACTTTTAATTTAAACGGAATAAAAAAAAAAAGTTATGATAATGAATCTTCCTATATCAAAACAGCTTATCATGATATAATTGATGAAGATAAAAATTTATTAAATAAAGAAAAAAGTTTTAATAAAGATGAAAAAAAAAAAAAAAAAAGTATTCTTGAGACAGATCAATATAAAAATTGTGAGGATCAACAATTTGATAATAAATATAATCATATAATAATTGGAAAGAAAAAAAGTGATAGCACTTATGATAATAGTTTGGATAAAGAATCATCAAATGAAAAAAAATTATCAATTGATAAAGGACAATATTCTAATATGGATAGTTCAAATTATTTTCATGATTGTAAAACCATGCTAAGTGAACATAATGAATCTATTGAATCTAGTAATAATGATATAAATGGAAAGCAAAAGGAACACATTAAAAAA**GGAAATTCTGAGAATCAAGATGTAGATCCAGATACAAACCCAGATGCAGTTCCAGATGATGATGATGATGATGATAATAGTAATGATGAAAGTGAATATGAAAGTAGTCAAATGGATAGTGAAAAAAATAAGGGATCAATAAAAAATTCAAAAAATGTTGTTATATATGCTGACGGAGTATATGATATGCTTCATTTAGGGCATATGAAACAATTGGAGCAAGCCAAAAAACTTTTTGAAAATACTACTTTAATAGTAGGTGTAACTAGCGATAATGAAACCAAATTATTTAAAGGTCAAGTTGTTCAAACTCTTGAAGAAAGAACAGAGACTTTAAAACATATACGATGGGTAGATGAAATAATTTCTCCATGCCCATGGGTTGTAACTCCAGAATTTTTGGAAAAATATAAAATAGATTATGTTGCACATGATGATATACCATATGCAAATAATCAGAAAAAGAAAAAGAAAAAAAAATCTAAAGGGAAGTCATTTAGTTTTGATGAAGAAAATGAAGATATATATGCTTGGTTAAAAAGAGCAGGCAAATTTAAAGCAACACAAAGAACAGAAGGTGTATCAACTACAGATTTAATAGTAAGAATATTAAAAAATTATGAGGATTATATTGAAAGGTCATTACAACGTGGTATACATCCTAATGAATTAAATATTGGTGTAACCAAAGCTCAGTCAATAAAAATGAAAAAAAATTTAATAAGATGGGGAGAAAAAGTGACAGATGAATTAACTAAAGTTACTTTAACAGACAAGCCATTAGGTACGGATTTTGATCAAGGAGTTGAAAATCTTCAAGTCAAATTTAAAGAACTTTTTAAAATATGGAAAAATGCATCGAATAAATTGATAACTGATTTTACAAGAAAACTTGAAGCAACATCTTATTTAACATCTATTCAAAATATCATAGATTATGAAATAGAAAATGATGATTATGCTAGTAGTAATTTTGATGATGAAACCAGTAGT**TAAaatgtaatttgtcttttatttatatttgacatgatataataaaataaaaaataatataaaatataataagaagaaaaataaaaagaattgtacatataaatcatatacagattatatacatataatgtgtataaaatattttttttatcttgtcatagtttttaatatatgaatttatataaaattttttttttttgtttttattatatctgcctaaatttctcaatttatcaatttttatgaagcattctctcatctttatcttttccccttctttttatttcatttctctttttccttttatttcttcttttttatatatagtttttatgtttatgtaaataattaaataatcttcttttttaaaggtaattcttttttttctttttttttttttatatataatattttataaatttttatgtacatttttgtattttgtattttgtattttgaattttccatttttcattcttttttttttttttttttttttttgtgtgtatattatgaaaaaaagataaaatagttaattattaaatttgtactcatatatattatatatataatatatatgtatatatgtgtaatctatatatattgttttcaccttcttactattgatcaaattataaaaaaaaaaaggaaagaaaatgaagataaggtattcaaatgtacaaaatgttttatataactatatatgatatatatattatgatttatttatgtatgtattttattatgatttattaatttttgttttaatacatgccttgtaaattattaataatttatcacaatgtgcgtaatttttaaaatatactttcttaattatccaaaatatttattggatccataatgtgtaaatatatggacatattattataatatgcacaataataaaaaaatatgcagcctacattgatatatatatatatatatatatatatatacatatacatatatttgaaatttagaatattatatatttttgtctatcatgaattttatattgtttcattttattttattttttttgtttttgtgtttttatataaatcttatataacttattaaaaatttctgaaattttcttgttgaacattatagaaaagtacttagtttttaaaaagataaattatttttgttgttttgtttaaagt

**Original Locus (OL) amplicon size: 1065bp**

**> PF3D7_1316600-HALO-SW Halo Neo-R**

ttttttttattttgttttgactaaaatatgcactaaaattttttttgaataatattattaattgtagtaaatattaattataattttttgtacattttgtgttttataacacattagtcataaaatagattatatatatatatatatatatatatatatataataccatattgattatatataatattatatatatatgtatatatctaattattaactaattatttaaatatatatattttatattaacttcatattataaatgtataataaagcgtttcccttttttttaattattaaaatcatgagataatataaaATGATAATTAAGGTTAATTCGGTTCAAACACAGAAG*gtaaaaattaaaattattaagataaatattaatactggaggttatacatatatatatatatatatatataatatatatataatattaaggagaattatattgtgcgaatacgtaaaatgaagttgaactattataaattacttagaaaaaaaggggtgcaatattgtattgaataaaaaatatataaagacccctaatgtgaatacagctatatagtaaattacacatatatattatatatatatatatatatatatatatatatatatatttatttatttatttatttgttgtacttttgaacattttaattttag*AGCCAAGATGGTAACAAAGTTCATAATAGTCAAGATGATATGGATAATGAGGAAAAAAGCATAAGGATATATGCAGATG*gttaaaaaatataaagaatatatgaaacatgacatatatatatatgtgtatatattttcttatttttttcttaattttttttgttggtttttccttctttttag*GTGTTTATGACCTGCTACATTTGGGTCATATGAAACAGCTTGAGCAAGCCAAACATGTAGATAAGAATGTTACCCTAATTGTGGGTGTTACAGGAGATAATGAGACTAGAAAGTTTAAAGGTCAAATTGTTCAAACTCTAGAAGAAAGAACAGAGACTTTAAAACATATACGATGGGTAGATGAAATAATTTCTCCATGCCCATGGGTGATTACTCCTGAATTTATGGAGGAACATAAAATAGATTATGTTGCACATGATGATATACCATATGCAAATAATCAGAAAAAGAAGAAAAAAAAAAAGTCTAAAGGGAAATCATTTAGTTTCGATGAAGAAAATGAAGATATATATGCTTGGTTAAAAAGAGCAGGGAAATTTAAAGCAACACAAAGAACAGAAGGTGTATCAACTACAGATTTAATTGTAAGAATATTAAAAAATTATGAGGATTATATTGAAAGGTCATTACAACGTGGTATACATCCTAATGAATTAAATATTGGTGTAACCAAAGCTCAGTCAATAAAAATGAAAAAAAATTTAATACGTTGGGGAGAAAAAGTAACAGATGAATTAACGAAGGTAACTTTAACAGACAAGCCATTAGGTACTGATTTTGATCAGGGTATTGATATAATAAGAGATAAAGTACATGATTTATTTAAATTATGGAGATATCATTCAAAAAAACTTTTAAAAGATTTTGCAAAATCATTTGATCCAATGTTTATTATAATTAGAAAAAGATATAGGAAAGATAATTTATCAGCTATGTACTTATCTGATTCAAATTATTTTTCTAGTTCTATGAAAGATGAATTAAAAAAAAAGAAAAGTTTAAGTAGTACATTTAATAATATTGATGAATATTATTATTCAGCTGATGAAGAAGATACACGGGATAATAGTTTTTATCGAGTTATAAATAAAATTGCTAATCATACTTATTATAGTGAAAAAGAAAATTATGAAACCTGTTTTGAATTAGAACATTCTTTAAGAGATAAAAAAAAAAGTTTAATATTTAAAGATAAAAATATGAAATTATCACCTGATGATTTATTACATTTTACTAATCCTATGAATTCAAAGTATGATCATAATTATTATTATTATAATAATACATATGATAAATTAAGTAACGTAACTTTTAATTTAAACGGAATAAAAAAAAAAAGTTATGATAATGAATCTTCCTATATCAAAACAGCTTATCATGATATAATTGATGAAGATAAAAATTTATTAAATAAAGAAAAAAGTTTTAATAAAGATGAAAAAAAAAAAAAAAAAAGTATTCTTGAGACAGATCAATATAAAAATTGTGAGGATCAACAATTTGATAATAAATATAATCATATAATAATTGGAAAGAAAAAAAGTGATAGCACTTATGATAATAGTTTGGATAAAGAATCATCAAATGAAAAAAAATTATCAATTGATAAAGGACAATATTCTAATATGGATAGTTCAAATTATTTTCATGATTGTAAAACCATGCTAAGTGAACATAATGAATCTATTGAATCTAGTAATAATGATATAAATGGAAAGCAAAAGGAACACATTAAAAAA**GGAAATTCTGAGAATCAAGATGTAGATCCAGATACAAACCCAGATGCAGTTCCAGATGATGATGATGATGATGATAATAGTAATGATGAAAGTGAATATGAAAGTAGTCAAATGGATAGTGAAAAAAATAAGGGATCAATAAAAAATTCAAAAAATGTTGTTATATATGCTGACGGAGTATATGATATGCTTCATTTAGGGCATATGAAACAATTGGAGCAAGCCAAAAAACTTTTTGAAAATACTACTTTAATAGTAGGTGTAACTAGCGATAATGAAACCAAATTATTTAAAGGTCAAGTTGTTCAAACTCTTGAAGAAAGAACAGAGACTTTAAAACATATACGATGGGTAGATGAAATAATTTCTCCATGCCCATGGGTTGTAACTCCAGAATTTTTGGAAAAATATAAAATAGATTATGTTGCACATGATGATATACCATATGCAAATAATCAGAAAAAGAAAAAGAAAAAAAAATCTAAAGGGAAGTCATTTAGTTTTGATGAAGAAAATGAAGATATATATGCTTGGTTAAAAAGAGCAGGCAAATTTAAAGCAACACAAAGAACAGAAGGTGTATCAACTACAGATTTAATAGTAAGAATATTAAAAAATTATGAGGATTATATTGAAAGGTCATTACAACGTGGTATACATCCTAATGAATTAAATATTGGTGTAACCAAAGCTCAGTCAATAAAAATGAAAAAAAATTTAATAAGATGGGGAGAAAAAGTGACAGATGAATTAACTAAAGTTACTTTAACAGACAAGCCATTAGGTACGGATTTTGATCAAGGAGTTGAAAATCTTCAAGTCAAATTTAAAGAACTTTTTAAAATATGGAAAAATGCATCGAATAAATTGATAACTGATTTTACAAGAAAACTTGAAGCAACATCTTATTTAACATCTATTCAAAATATCATAGATTATGAAATAGAAAATGATGATTATGCTAGTAGTAATTTTGATGATGAAACCAGTAGT**CCTAGGTCAGGATTGAGATCAAGATCTGCTGCTGCTGGTGCTGGTGGTGCTGCTAGAGCTGCTctgcagAGAGGAGTACAAGTTGAAACAATATCACCAGGAGATGGTCGTACATTTCCAAAAAGAGGTCAAACTTGTGTTGTACATTATACTGGAATGCTTGAAGATGGAAAGAAATTTGATTCATCTCGTGATAGAAATAAACCATTTAAATTTATGCTAGGTAAACAAGAAGTAATACGAGGTTGGGAAGAAGGAGTTGCTCAAATGAGTGTAGGTCAAAGAGCAAAACTTACTATATCTCCAGATTATGCTTATGGTGCAACTGGACATCCAGGTATAATTCCACCTCATGCAACTCTTGTATTTGATGTGGAGCTTCTAAAACTAGAAACTAGAGGTGTTCAGGTTGAAACAATTTCACCTGGAGATGGCAGAACCTTTCCTAAAAGAGGACAGACTTGCGTAGTTCATTATACAGGCATGCTAGAGGATGGTAAGAAATTTGATTCTAGTCGAGATAGAAATAAGCCATTCAAGTTTATGCTAGGTAAACAGGAAGTAATAAGAGGTTGGGAAGAGGGTGTAGCACAGATGTCAGTTGGACAAAGAGCAAAGTTAACAATATCACCAGATTATGCATACGGTGCAACAGGCCATCCTGGCATCATCCCTCCACATGCAACTTTAGTATTCGACGTTGAATTGTTAAAGTTAGAGACAacgcgtGCTAGAGGTGCTGCTGCTGGTGCTGGAGGTGCAGGTAGAcgtacgATGGCAGAAATTGGTACGGGTTTTCCATTTGATCCTCATTATGTGGAGGTGCTGGGGGAAAGGATGCATTATGTTGACGTAGGACCAAGAGATGGTACTCCAGTGTTATTTTTGCATGGAAACCCAACCTCGAGTTATGTATGGAGAAATATAATTCCACATGTAGCACCAACACATAGATGTATAGCTCCTGATTTAATTGGTATGGGAAAAAGTGATAAACCTGACTTAGGATATTTTTTTGATGATCATGTCCGTTTTATGGATGCTTTCATTGAAGCTCTTGGCCTTGAAGAAGTAGTATTAGTTATACATGATTGGGGATCCGCCTTAGGATTTCATTGGGCCAAGAGGAATCCTGAAAGAGTAAAAGGAATAGCATTCATGGAATTCATACGACCAATCCCCACATGGGATGAATGGCCAGAATTTGCACGCGAAACATTTCAAGCTTTTAGAACTACAGATGTTGGTAGAAAATTAATAATAGATCAAAATGTATTTATAGAAGGAACTTTACCTATGGGTGTTGTAAGGCCGTTAACAGAAGTTGAAATGGACCACTACCGTGAACCTTTTTTAAATCCAGTAGATAGAGAGCCCTTATGGAGATTTCCTAATGAATTACCTATTGCAGGTGAACCCGCGAATATTGTTGCTTTAGTAGAAGAATATATGGATTGGTTACATCAGTCTCCTGTTCCTAAACTTCTATTTTGGGGTACACCTGGAGTTCTAATACCACCAGCTGAAGCAGCAAGATTAGCAAAATCATTACCAAATTGTAAAGCTGTTGATATAGGTCCTGGGTTGAATTTATTACAAGAAGATAATCCAGATTTGATTGGATCTGAGATAGCTAGATGGCTAAGTACATTAGAAATTTCAGGTACCGGTGCCAGGGGAGCAGCCGCAGGAGCAGGGGGGGCAGGAAGGCGTGGTGTTCAGGTCGAGACTATTAGCCCTGGAGATGGACGCACGTTTCCTAAGCGTGGACAGACATGCGTAGTTCACTACACAGGTATGTTGGAGGACGGTAAAAAGTTCGACAGCTCACGCGACCGCAATAAACCTTTCAAGTTTATGCTTGGCAAGCAGGAGGTTATTCGTGGATGGGAGGAGGGTGTAGCACAGATGTCTGTTGGACAGCGTGCTAAGTTGACAATTTCACCTGACTATGCTTATGGCGCTACGGGCCATCCCGGGATCATTCCGCCACATGCGACTCTGGTATTCGACGTTGAATTATTAAAGTTAGAGACAGCTAGAGGGGCCGCTGCAGGTGCTGGTGGAGCTGGAAGACGTGGAGTACAAGTAGAGACTATCTCTCCAGGTGACGGTCGCACTTTCCCAAAGCGTGGCCAAACCTGTGTTGTACATTACACTGGTATGCTGGAGGATGGGAAAAAGTTCGATTCCAGTCGCGACCGTAACAAACCGTTCAAATTCATGTTGGGAAAGCAGGAAGTGATCCGCGGGTGGGAGGAAGGCGTGGCGCAAATGAGCGTCGGTCAGCGGGCTAAATTGACCATTTCCCCTGACTACGCGTATGGGGCTACTGGGCACCCAGGGATTATTCCGCCTCACGCTACACTTGTGTTTGATGTCGAACTTTTGAAACTGGAAACTGTCGACGGAGAAGGAAGAGGAAGTTTATTAACATGTGGAGATGTAGAAGAAAATCCAGGACCAATGATTGAACAAGATGGATTGCACGCAGGTTCTCCGGCCGCTTGGGTGGAGAGGCTATTCGGCTATGACTGGGCACAACAGACAATCGGCTGCTCTGATGCCGCCGTGTTCCGGCTGTCAGCGCAGGGGCGCCCGGTTCTTTTTGTCAAGACCGACCTGTCCGGTGCCCTGAATGAACTGCAGGACGAGGCAGCGCGGCTATCGTGGCTGGCCACGACGGGCGTTCCTTGCGCAGCTGTGCTCGACGTTGTCACTGAAGCGGGAAGGGACTGGCTGCTATTGGGCGAAGTGCCGGGGCAGGATCTCCTGTCATCTCACCTTGCTCCTGCCGAGAAAGTATCCATCATGGCTGATGCAATGCGGCGGCTGCATACGCTTGATCCGGCTACCTGCCCATTCGACCACCAAGCGAAACATCGCATCGAGCGAGCACGTACTCGGATGGAAGCCGGTCTTGTCGATCAGGATGATCTGGACGAAGAGCATCAGGGGCTCGCGCCAGCCGAACTGTTCGCCAGGCTCAAGGCGCGCATGCCCGACGGCGAGGATCTCGTCGTGACCCATGGCGATGCCTGCTTGCCGAATATCATGGTGGAAAATGGCCGCTTTTCTGGATTCATCGACTGTGGCCGGCTGGGTGTGGCGGACCGCTATCAGGACATAGCGTTGGCTACCCGTGATATTGCTGAAGAGCTTGGCGGCGAATGGGCTGACCGCTTCCTCGTGCTTTACGGTATCGCCGCTCCCGATTCGCAGCGCATCGCCTTCTATCGCCTTCTTGACGAGTTCTTCTAActcgagggatatggcagcttaatgttcgtttttcttatttatatatttataccaattgattgtatttataactgtaaaaatgtgtatgttgtgtgcatatttttttttgtgcatgcacatgcatgtaaatagctaaaattatgaacattttattttttgttcagaaaaaaaaaactttacacacataaaatggctagtatgaatagccatattttatataaattaaatcctatgaatttatgaccatattaaaaatttagatatttatggaacataatatgtttgaaacaataagacaaaattattattattattattatttttactgttataattatgtgtctccttcaatgattcataaatagttggacttgatttttaaaatgtttataatatgattagcatagttaaataaaaaaagttgaaaaattaaaaaaaaacatataaacacaaatgatggtttttccttcaatttcgatatcaatttatagaaacaaaatatatacttgtataattttatttttttatataaatcattacatatataattatacaatattttttctaagagataattatatattaatatatataaaaaaaggtgttttttttttttttttttatttttatttttattttatggtaatattttattttccttattttataaattatattagtttatatgtgattaattttatatattatcaatttatatatttttaaatgcttacttaattatctttttttttttttttttttttttttcccctctttttatattaatttatttttgaaaaaattgatatatatatatatatataatatatatatatacatgtagtagtattaaacaatgtataatatatataaataatatatttatatatttcatttcaattttaattttttttggttttttttttttttctttttgtcatatttaaaaaaaattatattcatataagttatgcattttttataaacattattcaatatatgtataatataatatatatatatatattaatgtattattccaatgtgcatgataaaagaaaaaaataatatttataaaaaaaaagaaaaataaaacaaaaaaagaaaaaaaaaaaaaaaaaaaaaaaaatacaaaaataaataatataatttataattatatattcttgtcacaataaaaatatatatatatatatatatatttataatatgtatattttaaactagaaaaggaataactaatattttatttattatcattcaagatttatattttataataataaatacctaatagaaatatatcaggatccatgcatggttcgctaaactgcatcgtcgctgtgtcccagaacatgggcatcggcaagaacggggactacccctggccaccgctcaggaacgaatttagatatttccagagaatgaccacaacctcttcagtagaaggtaaacagaatctggtgattatgggtaagaagacctggttctccattcctgagaagaatcgacctttaaagggtagaattaatttagttctcagcagagaactcaaggaacctccacaaggagctcattttctttccagaagtctagatgatgccttaaaacttactgaacaaccagaattagcaaataaagtagacatggtctggatagttggtggcagttctgtttataaggaagccatgaatcacccaggccatcttaaactatttgtgacaaggatcatgcaagactttgaaagtgacacgttttttccagaaattgatttggagaaatataaacttctgccagaatacccaggtgttctctctgatgtccaggaggagaaaggcattaagtacaaatttgaagtatatgagaagaatgattaagcttatttaataatagattaaaaatattataaaaataaaaacataaacacagaaattacaaaaaaaatacatatgaattttttttttgtaatcttccttataaatatagaataatgaatcatataaaacatatcattattcatttatttacatttaaaattattgtttcagtatctttaatttattatgtatatataaaaataacttacaattttattaataaacaatatatgtttattaattcatgttttgtaatttatgggatagcgattttttttactgtctgtatttttcttttttaattatgttttaattgtattttatttttattattgttctttttatagtattattttaaaacaaaatgtattttctaagaacttataataataataaatataaattttaataaaaattatatttatcttttacaatatgaacataaagtacaacattaatatatagcttttaatatttttattcctaatcatgtaaatcttaaatttttctttttaaacatatgttaaatatttatttctcattatatataagaacatatttattaaatctagaattctatagtgagtcgtattacaattcactggccgtcgttttacaacgtcgtgactgggaaaaccctggcgttacccaacttaatcgccttgcagcacatccccctttcgccagctggcgtaatagcgaagaggcccgcaccgatcgcccttcccaacagttgcgcagcctgaatggcgaatggcgcctgatgcggtattttctccttacgcatctgtgcggtatttcacaccgcatatggtgcactctcagtacaatctgctctgatgccgcatagttaagccagccccgacacccgccaacacccgctgacgcgccctgacgggcttgtctgctcccggcatccgcttacagacaagctgtgaccgtctccgggagctgcatgtgtcagaggttttcaccgtcatcaccgaaacgcgcgagacgaaagggcctcgtgatacgcctatttttataggttaatgtcatgataataatggtttcttagacgtcaggtggcacttttcggggaaatgtgcgcggaacccctatttgtttatttttctaaatacattcaaatatgtatccgctcatgagacaataaccctgataaatgcttcaataatattgaaaaaggaagagtatgagtattcaacatttccgtgtcgcccttattcccttttttgcggcattttgccttcctgtttttgctcacccagaaacgctggtgaaagtaaaagatgctgaagatcagttgggtgcacgagtgggttacatcgaactggatctcaacagcggtaagatccttgagagttttcgccccgaagaacgttttccaatgatgagcacttttaaagttctgctatgtggcgcggtattatcccgtattgacgccgggcaagagcaactcggtcgccgcatacactattctcagaatgacttggttgagtactcaccagtcacagaaaagcatcttacggatggcatgacagtaagagaattatgcagtgctgccataaccatgagtgataacactgcggccaacttacttctgacaacgatcggaggaccgaaggagctaaccgcttttttgcacaacatgggggatcatgtaactcgccttgatcgttgggaaccggagctgaatgaagccataccaaacgacgagcgtgacaccacgatgcctgtagcaatgccaacaacgttgcgcaaactattaactggcgaactacttactctagcttcccggcaacaattaatagactggatggaggcggataaagttgcaggaccacttctgcgctcggcccttccggctggctggtttattgctgataaatctggagccggtgagcgtgggtctcgcggtatcattgcagcactggggccagatggtaagccctcccgtatcgtagttatctacacgacggggagtcaggcaactatggatgaacgaaatagacagatcgctgagataggtgcctcactgattaagcattggtaactgtcagaccaagtttactcatatatactttagattgatttaaaacttcatttttaatttaaaaggatctaggtgaagatcctttttgataatctcatgaccaaaatcccttaacgtgagttttcgttccactgagcgtcagaccccgtagaaaagatcaaaggatcttcttgagatcctttttttctgcgcgtaatctgctgcttgcaaacaaaaaaaccaccgctaccagcggtggtttgtttgccggatcaagagctaccaactctttttccgaaggtaactggcttcagcagagcgcagataccaaatactgtccttctagtgtagccgtagttaggccaccacttcaagaactctgtagcaccgcctacatacctcgctctgctaatcctgttaccagtggctgctgccagtggcgataagtcgtgtcttaccgggttggactcaagacgatagttaccggataaggcgcagcggtcgggctgaacggggggttcgtgcacacagcccagcttggagcgaacgacctacaccgaactgagatacctacagcgtgagctatgagaaagcgccacgcttcccgaagggagaaaggcggacaggtatccggtaagcggcagggtcggaacaggagagcgcacgagggagcttccagggggaaacgcctggtatctttatagtcctgtcgggtttcgccacctctgacttgagcgtcgatttttgtgatgctcgtcaggggggcggagcctatcgaaaaacgccagcaacgcggcctttttacggttcctggccttttgctggccttttgctcacatgttctttcctgcgttatcccctgattctgtggataaccgtattaccgcctttgagtgagctgataccgctcgccgcagccgaacgaccgagcgcagcgagtcagtgagcgaggaagcggaagagcgcccaatacgcaaaccgcctctccccgcgcgttggccgattcattaatgcagctggcacgacaggtttcccgactggaaagcgggcagtgagcgcaacgcaattaatgtgagttagctcactcattaggcaccccaggctttacactttatgcttccggctcgtatgttgtgtggaattgtgagcggataacaatttcacacaggaaacagctatgaccatgattacgccaagctatttaggtgacactatagaagcggccgcTAgGGAAATTCTGAGAATCAAGATGTAGATCCAGATACAAACCCAGATGCAGTTCCAGATGATGATGATGATGATGATAATAGTAATGATGAAAGTGAATATGAAAGTAGTCAAATGGATAGTGAAAAAAATAAGGGATCAATAAAAAATTCAAAAAATGTTGTTATATATGCTGACGGAGTATATGATATGCTTCATTTAGGGCATATGAAACAATTGGAGCAAGCCAAAAAACTTTTTGAAAATACTACTTTAATAGTAGGTGTAACTAGCGATAATGAAACCAAATTATTTAAAGGTCAAGTTGTTCAAACTCTTGAAGAAAGAACAGAGACTTTAAAACATATACGATGGGTAGATGAAATAATTTCTCCATGCCCATGGGTTGTAACTCCAGAATTTTTGGAAAAATATAAAATAGATTATGTTGCACATGATGATATACCATATGCAAATAATCAGAAAAAGAAAAAGAAAAAAAAATCTAAAGGGAAGTCATTTAGTTTTGATGAAGAAAATGAAGATATATATGCTTGGTTAAAAAGAGCAGGCAAATTTAAAGCAACACAAAGAACAGAAGGTGTATCAACTACAGATTTAATAGTAAGAATATTAAAAAATTATGAGGATTATATTGAAAGGTCATTACAACGTGGTATACATCCTAATGAATTAAATATTGGTGTAACCAAAGCTCAGTCAATAAAAATGAAAAAAAATTTAATAAGATGGGGAGAAAAAGTGACAGATGAATTAACTAAAGTTACTTTAACAGACAAGCCATTAGGTACGGATTTTGATCAAGGAGTTGAAAATCTTCAAGTCAAATTTAAAGAACTTTTTAAAATATGGAAAAATGCATCGAATAAATTGATAACTGATTTTACAAGAAAACTTGAAGCAACATCTTATTTAACATCTATTCAAAATATCATAGATTATGAAATAGAAAATGATGATTATGCTAGTAGTAATTTTGATGATGAAACCAGTAGTTAAaatgtaatttgtcttttatttatatttgacatgatataataaaataaaaaataatataaaatataataagaagaaaaataaaaagaattgtacatataaatcatatacagattatatacatataatgtgtataaaatattttttttatcttgtcatagtttttaatatatgaatttatataaaattttttttttttgtttttattatatctgcctaaatttctcaatttatcaatttttatgaagcattctctcatctttatcttttccccttctttttatttcatttctctttttccttttatttcttcttttttatatatagtttttatgtttatgtaaataattaaataatcttcttttttaaaggtaattcttttttttctttttttttttttatatataatattttataaatttttatgtacatttttgtattttgtattttgtattttgaattttccatttttcattcttttttttttttttttttttttttgtgtgtatattatgaaaaaaagataaaatagttaattattaaatttgtactcatatatattatatatataatatatatgtatatatgtgtaatctatatatattgttttcaccttcttactattgatcaaattataaaaaaaaaaaggaaagaaaatgaagataaggtattcaaatgtacaaaatgttttatataactatatatgatatatatattatgatttatttatgtatgtattttattatgatttattaatttttgttttaatacatgccttgtaaattattaataatttatcacaatgtgcgtaatttttaaaatatactttcttaattatccaaaatatttattggatccataatgtgtaaatatatggacatattattataatatgcacaataataaaaaaatatgcagcctacattgatatatatatatatatatatatatatatacatatacatatatttgaaatttagaatattatatatttttgtctatcatgaattttatattgtttcattttattttattttttttgtttttgtgtttttatataaatcttatataacttattaaaaatttctgaaattttcttgttgaacattatagaaaagtacttagtttttaaaaagataaattatttttgttgttttgtttaaagt

**5’ amplicon size: 1161bps**

**3’ amplicon size: 1122bps**

## **> PF3D7_1458500**

ttactattagagaacaattaaaattatttattttattttatttttatttttttcataaaagggaataaaaaaatatcatattcttattttatatatacgtaaatttctaaatgtgtatttttttttaaaagttatttcttaaatatttttaataatatatgttttagtttacatttattataattttttaatttaaattttttggtaacatattactaagataaagtattgcttttataaagtatttcatcaaaggttgatataaaatagaaaaagacaaaaaatattatttttattaaaacaataacatacttaaatttcatattgaaatattattttgtattattaattgttagaaataactgatattgatttattttcaatttcaaactttatatcattagcaaaaataataaataaatgaataaataaataaataaataaatatattgtacatatatatatatatatatatatatatatacttattgattatttatatataaaaatataatatttcttatacacaaagaaataatattaataataattgaattaatatatgttgatacatgaagctgtacataatatttgtcatttaaattttatatttttttaatttttaataaatggtttgatataataATGAAAAAAATTAAAGACGAATTTGATGATTATTATGATGCGTATGAAGAAAATTGGTCCCCCGATACATTTGCCGAATTGGAATCAAGCCAAAGTTTAGAAAAAACAATAAAGAAAAAAATGTACAAAGAAGAATTTGAGCAAAACAATTTTTTTTGTGCGGCAAAGGAAATATATTCAAACAACTCCGAAAATGTGCAAGGGGATAATAATGATATAATAAAAAAAGAAATAACTAATGATGATAATAATGAAATGAAAAAGGAAAATATAGGAAATAATGATAATAATAATAATAATAATAATAATAGTCATTTGGTATTAGATATCGACGGAAATATCGATTTTACTGAGGAAAACGTTGAAGGTTCTAATAAAAATGAATATGAAAATGATTTTGATGAAAGTAAATGGACTCATCATTTTAATGAAGAAAATAAAAAGGAAGATACAAACACGTTATATGAAGAAAGGAAAGATGAAGTAGATGAAAACAGAAAAAGTCATGAAGAAATTAATTTATTTAAAGATATATATGAAGAATTAGAAAATTCCCAAAATGAAAAGGAAAAAAGCTATGTGGATTATTCATATAATGAAAATTATTCTAATAAAAAAAATGAAAATTCTGATTCTTTTTTCAATAATTTAAATATATATAGTAATAATATAGAAAGAAATCATGATGAGAAGGATGATTCGTTAAGTGTCTTTAATAGGAACGAATCGGATTTAAATTTATTAGATGATATAAATGATAGCAATACTTATAACAATAGGAGCAAGGATAAAAATGATTTCAGTCCATGTAGTAATAATAATAATAATAATGATAATAATAATTATAATTATAATAATAATAATTATAATAATAATAATGATAATAATAATATGAATGATATTTTCGAGAAGTGTCATATATATGAAAAAAACGAAGAATATGAGAATTATTCAAACAATTCATTTTCTCATATTTTACAGAAAGAAGAAAAAAATTATGATAATACAATGGAAAATAAAACCGATATGATGAATGTAAACAATTTCGAAAGATATAATTTAAAAGATGAAAACGTAAGTTTACAAAATGATGATATCATGAGCCTTATCAGTGAAAGGTTACAAGAAGATAAAAATAAAAATTTTATGGATTATAGTTTTTATAATAATACTTGTACAAGTTATAATGATAAGGATATAAAAAATATCGTATCAAATTATGATATGTCAAATGATATATCGAATAATTTATCTAATATTATTTCAAATATGATATCGAATAATCTATCTAATATCATATCAAATAATATATCAAATTATAATATATCCGATAGTATTAATTCAAATCATATAATTTCAAATAAAAATAATTTAAAATATAATAATAGTATTAATAATAATGACAGCACTATAGATAATGGAACGAATAGTTCCAATCATATAAATTATAGTAATAATAATATGAAGGATAATACATCTTGTATATATGATAATTATGATCGTAGATTGCATAATAGTTCATCATCCTATAGAGATATGTGCCATGAAAGGAATGATAAAGACACCAATATTAAAAGTGATATTATTACTACATGTTTAGATAATGATTATTCTAATATGAATTATAGTTCTGATAATTCTTATGTAGATAAGGAGATCAAGCGTATGTTTGATGAAGATAATTTACAAGATGGTTCAATAGAATATAACAATAATTATAAACAATTTGATAAATCGATAAACAAAGAACATAATAGTGAAAAGACAGACACACGAAACGAGATAAATAATACTAAAAATAATGATACAAAGATAAATATAAATAGGTATAATGAAGAATGTTACCCTAGAGAACATTTGGAAAAACTTGAAAAGGAATTAATAATGAATGAAAAAAATAAAATAAATAATTATACTGAAAGTGATATATATAATACTAGCGAAAAAAAAACAAAATGTGCATTAGAAAAAGAAGTAAATGATGAAGAGATTAAAACAAGGGAAAATAGTATTGATATTTGTGTAGATCTAAATGATGAAGAACCTTGGGACGAAAATATAAAAAGAGAAAACAACAATGACAAGAAAAGATTAACTTATAAGAATGAAAAATATAATAAATACGCAATTTTGAAAGATAAGGGTTCAAAAATGAACAACAACAACAACAACAACAACAACAATAATAATAATAATAATAATAATAATATTGAAAGGAACAACGTGAATGTTATAGGGGAGAAAAAGAAGAAACAAACTAAAAAGGAAACATCACAAAATAATAATAATCATAATATTAATAAGAAAGAAAAGATTATATCTACAAAATTAATAAACCAAACGAACCAAAGTGATAACCTAAAGGAACTAGGAAATGAATTAAATAATCAGATAAATAAACTCGAAAAAGAACAAGATAAAGTTAAAAAGCTAGAATATGAATTAATAGCTAAAAGTGCAGAAATGGAATTAGAAAGAGAAGAAATGAAAAATAACATGGAACAAGAAAAAAAAAAAATGATGAAAACTATTGAAGAAGAAAAAAAAAAATGGAATAAAGAAAAAAAAAGAATCGAAGGAGAAGTAGAGAAACAAAGAAATATTATAATGCATAAAAGAAAATTAACAAACGAAATAGCAATGTTCAAAAATAGAATAAAAGAATTAGAAGAAAATTTAGAAAAAGAAAAAAAAGAACATAGAATTCTTGTAGAGAAATTAAAAAAAAAGATAGACAATTTAAAAATTGAAAATGAAAAAATTAAAACTGAATTAAAAATATCAAATGAATATAGAAATAAATTAGAAGTATATCAACAAAATACTATTATGAAATTAGCTACAACTGTTAATAAAAAAAAAAACCAAAAGTATAAAAGTAAAAATTTTCTAAATACATCATCTGAATCGTCCATTCAAGAAGAAATATATAATAATAGTAATAATAATAATAATAATAATAATAATAATAATAATAATAGTATAGACAATAGCGATATTATAGATAATCACCATTATGCTAAAAATAAATCACATGATACAAAAAAAATTAATAAATTTGATATAGACAAAAGCTTGTTGGACTCTTCAGATAATTCTGATGAAATAAAAAAAATTATTAATCATAAAAAGAATAATATACATAAAGATTTAGAAATTATATATAATAGTTCTACTAACACAGATGATGGAATTTATAACAACAAAATAAAAAACAATATAAAAATAAAAAGTAATAATAATAATAATAATAATATGAACTTGAAAAATAACAAAAATAAATTTACATTGGATAAATTATTTATTCAAAACAAAAAAGATAAAGATAGTTATAAAAATTCAGATAATAATAAAATTGATGAAGAATATATAAATAAAAATTTAAAGAGAATGAGATCAATAATTAAAAATAAAAAAAAATACCTAGAAGATAAAATGTCTAATCATGAAAATGATGATACTTGTAGTTTACTAACTACTACAAACGAAATTAATTTGAATGACAAAGAAGTATTTTTTTATGAAAACTGTAAAAAGAATAAGGATAAAATTTCATTTAATGATGATATAAATTATGCAAATAATAAATCAAATATAAAATATGATAAAAGAACATCAAATAAAGAGGGCGGTGATATTATATATGAAAATATGAATAAAAAGAAAAACAAAGGTAGTACATTAAATGATAACAATAATAATAATAAAATAATGAAACCGTCTTTGGTTAATAATACAAATGAAATATATATGAGTGATCTTAATACGAATGAGCATAATGAAACTTCGTACAAGCCACATAGAAATTATTCTACTAGTAATCAAAAAGGAAAAGATGATATTATAAATAATAAAATATTATCTAAAAAAAAAACAAATAGTACGGATAGTTATAATAAAAATAATAATATA**GGTAATGATAATATTTTAAAACCTTTTGGTAAGGACTGGGATTTTGTAATAAACTTTGATTTCGACGAATTATTCAATCAATGTGAAAATGTTATAGAATCTATTTTTTCCTCATCCAAAAAAATAAAATATAGACAAGCATTTATTGATGGAAAGGTGGAAACTTTATTTGATGATGGGTTAAAATTAATTGAGAAAAATAGAAATAAAAAAATTATGCATCCAAGTAATATAACAATATATCTGTATCCTACCAAAGATTACAAGGCACATTTTCCCAATTCGTACATG*gtaaaatttaagaaaaaataaaaaaaaaaaaatttaaatgagcacatatatatatttatgtatatatatatgaccatattggtcttgctatatatcatacatttttataataatattatgtctgctttttag*TTATTTCGATTTGTTAATAAAGGAATATATCAGGTGAACATACCAAATAAATGTCAACTAAACAA*gtaaataagaaaaaaaaaaaaaaaaataataataataataaaacatcgttagagattattttttccattacatatatgtacatatatatatatatatatatatatatatacatttttttttatatgtatgtatttttctttttatttttattttcttcag*ATTTCCAAGTGGACAAGTGGATTGTAAATATACTGATGGTCATATACAAATACTATTTTGTGATGGAAAAAGAAAAGAAATACTACCCAACAGGGAAGAGTATGTTATATTACGAAACG*gtaaaatattaaaattaatataaaaatacacttatataaaatattaagaataaaacatatgtatattctttttttcttatttaaacatttttttattttaatacatttataactacaacgtcaatattataaataaatatgataaaaatatatttttttttttttttttttttttttttttccatttccctacag*GAACTATAAAAAAGTTAAAT**TAAtatgatgtgttcacttatatgaatatatgttggatatatgattaactcatgaaatttaatgttctattcataattgaggatataacaatataaacatttaaaatgtcaatgaaaaataaaggaacaaataaattttatattttat

**Original Locus (OL) amplicon size: 1126bp**s

**> PF3D7_1458500-HALO-SW Halo Neo-R**

ttactattagagaacaattaaaattatttattttattttatttttatttttttcataaaagggaataaaaaaatatcatattcttattttatatatacgtaaatttctaaatgtgtatttttttttaaaagttatttcttaaatatttttaataatatatgttttagtttacatttattataattttttaatttaaattttttggtaacatattactaagataaagtattgcttttataaagtatttcatcaaaggttgatataaaatagaaaaagacaaaaaatattatttttattaaaacaataacatacttaaatttcatattgaaatattattttgtattattaattgttagaaataactgatattgatttattttcaatttcaaactttatatcattagcaaaaataataaataaatgaataaataaataaataaataaatatattgtacatatatatatatatatatatatatatatacttattgattatttatatataaaaatataatatttcttatacacaaagaaataatattaataataattgaattaatatatgttgatacatgaagctgtacataatatttgtcatttaaattttatatttttttaatttttaataaatggtttgatataataATGAAAAAAATTAAAGACGAATTTGATGATTATTATGATGCGTATGAAGAAAATTGGTCCCCCGATACATTTGCCGAATTGGAATCAAGCCAAAGTTTAGAAAAAACAATAAAGAAAAAAATGTACAAAGAAGAATTTGAGCAAAACAATTTTTTTTGTGCGGCAAAGGAAATATATTCAAACAACTCCGAAAATGTGCAAGGGGATAATAATGATATAATAAAAAAAGAAATAACTAATGATGATAATAATGAAATGAAAAAGGAAAATATAGGAAATAATGATAATAATAATAATAATAATAATAATAGTCATTTGGTATTAGATATCGACGGAAATATCGATTTTACTGAGGAAAACGTTGAAGGTTCTAATAAAAATGAATATGAAAATGATTTTGATGAAAGTAAATGGACTCATCATTTTAATGAAGAAAATAAAAAGGAAGATACAAACACGTTATATGAAGAAAGGAAAGATGAAGTAGATGAAAACAGAAAAAGTCATGAAGAAATTAATTTATTTAAAGATATATATGAAGAATTAGAAAATTCCCAAAATGAAAAGGAAAAAAGCTATGTGGATTATTCATATAATGAAAATTATTCTAATAAAAAAAATGAAAATTCTGATTCTTTTTTCAATAATTTAAATATATATAGTAATAATATAGAAAGAAATCATGATGAGAAGGATGATTCGTTAAGTGTCTTTAATAGGAACGAATCGGATTTAAATTTATTAGATGATATAAATGATAGCAATACTTATAACAATAGGAGCAAGGATAAAAATGATTTCAGTCCATGTAGTAATAATAATAATAATAATGATAATAATAATTATAATTATAATAATAATAATTATAATAATAATAATGATAATAATAATATGAATGATATTTTCGAGAAGTGTCATATATATGAAAAAAACGAAGAATATGAGAATTATTCAAACAATTCATTTTCTCATATTTTACAGAAAGAAGAAAAAAATTATGATAATACAATGGAAAATAAAACCGATATGATGAATGTAAACAATTTCGAAAGATATAATTTAAAAGATGAAAACGTAAGTTTACAAAATGATGATATCATGAGCCTTATCAGTGAAAGGTTACAAGAAGATAAAAATAAAAATTTTATGGATTATAGTTTTTATAATAATACTTGTACAAGTTATAATGATAAGGATATAAAAAATATCGTATCAAATTATGATATGTCAAATGATATATCGAATAATTTATCTAATATTATTTCAAATATGATATCGAATAATCTATCTAATATCATATCAAATAATATATCAAATTATAATATATCCGATAGTATTAATTCAAATCATATAATTTCAAATAAAAATAATTTAAAATATAATAATAGTATTAATAATAATGACAGCACTATAGATAATGGAACGAATAGTTCCAATCATATAAATTATAGTAATAATAATATGAAGGATAATACATCTTGTATATATGATAATTATGATCGTAGATTGCATAATAGTTCATCATCCTATAGAGATATGTGCCATGAAAGGAATGATAAAGACACCAATATTAAAAGTGATATTATTACTACATGTTTAGATAATGATTATTCTAATATGAATTATAGTTCTGATAATTCTTATGTAGATAAGGAGATCAAGCGTATGTTTGATGAAGATAATTTACAAGATGGTTCAATAGAATATAACAATAATTATAAACAATTTGATAAATCGATAAACAAAGAACATAATAGTGAAAAGACAGACACACGAAACGAGATAAATAATACTAAAAATAATGATACAAAGATAAATATAAATAGGTATAATGAAGAATGTTACCCTAGAGAACATTTGGAAAAACTTGAAAAGGAATTAATAATGAATGAAAAAAATAAAATAAATAATTATACTGAAAGTGATATATATAATACTAGCGAAAAAAAAACAAAATGTGCATTAGAAAAAGAAGTAAATGATGAAGAGATTAAAACAAGGGAAAATAGTATTGATATTTGTGTAGATCTAAATGATGAAGAACCTTGGGACGAAAATATAAAAAGAGAAAACAACAATGACAAGAAAAGATTAACTTATAAGAATGAAAAATATAATAAATACGCAATTTTGAAAGATAAGGGTTCAAAAATGAACAACAACAACAACAACAACAACAACAATAATAATAATAATAATAATAATAATATTGAAAGGAACAACGTGAATGTTATAGGGGAGAAAAAGAAGAAACAAACTAAAAAGGAAACATCACAAAATAATAATAATCATAATATTAATAAGAAAGAAAAGATTATATCTACAAAATTAATAAACCAAACGAACCAAAGTGATAACCTAAAGGAACTAGGAAATGAATTAAATAATCAGATAAATAAACTCGAAAAAGAACAAGATAAAGTTAAAAAGCTAGAATATGAATTAATAGCTAAAAGTGCAGAAATGGAATTAGAAAGAGAAGAAATGAAAAATAACATGGAACAAGAAAAAAAAAAAATGATGAAAACTATTGAAGAAGAAAAAAAAAAATGGAATAAAGAAAAAAAAAGAATCGAAGGAGAAGTAGAGAAACAAAGAAATATTATAATGCATAAAAGAAAATTAACAAACGAAATAGCAATGTTCAAAAATAGAATAAAAGAATTAGAAGAAAATTTAGAAAAAGAAAAAAAAGAACATAGAATTCTTGTAGAGAAATTAAAAAAAAAGATAGACAATTTAAAAATTGAAAATGAAAAAATTAAAACTGAATTAAAAATATCAAATGAATATAGAAATAAATTAGAAGTATATCAACAAAATACTATTATGAAATTAGCTACAACTGTTAATAAAAAAAAAAACCAAAAGTATAAAAGTAAAAATTTTCTAAATACATCATCTGAATCGTCCATTCAAGAAGAAATATATAATAATAGTAATAATAATAATAATAATAATAATAATAATAATAATAATAGTATAGACAATAGCGATATTATAGATAATCACCATTATGCTAAAAATAAATCACATGATACAAAAAAAATTAATAAATTTGATATAGACAAAAGCTTGTTGGACTCTTCAGATAATTCTGATGAAATAAAAAAAATTATTAATCATAAAAAGAATAATATACATAAAGATTTAGAAATTATATATAATAGTTCTACTAACACAGATGATGGAATTTATAACAACAAAATAAAAAACAATATAAAAATAAAAAGTAATAATAATAATAATAATAATATGAACTTGAAAAATAACAAAAATAAATTTACATTGGATAAATTATTTATTCAAAACAAAAAAGATAAAGATAGTTATAAAAATTCAGATAATAATAAAATTGATGAAGAATATATAAATAAAAATTTAAAGAGAATGAGATCAATAATTAAAAATAAAAAAAAATACCTAGAAGATAAAATGTCTAATCATGAAAATGATGATACTTGTAGTTTACTAACTACTACAAACGAAATTAATTTGAATGACAAAGAAGTATTTTTTTATGAAAACTGTAAAAAGAATAAGGATAAAATTTCATTTAATGATGATATAAATTATGCAAATAATAAATCAAATATAAAATATGATAAAAGAACATCAAATAAAGAGGGCGGTGATATTATATATGAAAATATGAATAAAAAGAAAAACAAAGGTAGTACATTAAATGATAACAATAATAATAATAAAATAATGAAACCGTCTTTGGTTAATAATACAAATGAAATATATATGAGTGATCTTAATACGAATGAGCATAATGAAACTTCGTACAAGCCACATAGAAATTATTCTACTAGTAATCAAAAAGGAAAAGATGATATTATAAATAATAAAATATTATCTAAAAAAAAAACAAATAGTACGGATAGTTATAATAAAAATAATAATATA**GGTAATGATAATATTTTAAAACCTTTTGGTAAGGACTGGGATTTTGTAATAAACTTTGATTTCGACGAATTATTCAATCAATGTGAAAATGTTATAGAATCTATTTTTTCCTCATCCAAAAAAATAAAATATAGACAAGCATTTATTGATGGAAAGGTGGAAACTTTATTTGATGATGGGTTAAAATTAATTGAGAAAAATAGAAATAAAAAAATTATGCATCCAAGTAATATAACAATATATCTGTATCCTACCAAAGATTACAAGGCACATTTTCCCAATTCGTACATG*gtaaaatttaagaaaaaataaaaaaaaaaaaatttaaatgagcacatatatatatttatgtatatatatatgaccatattggtcttgctatatatcatacatttttataataatattatgtctgctttttag*TTATTTCGATTTGTTAATAAAGGAATATATCAGGTGAACATACCAAATAAATGTCAACTAAACAA*gtaaataagaaaaaaaaaaaaaaaaataataataataataaaacatcgttagagattattttttccattacatatatgtacatatatatatatatatatatatatatatacatttttttttatatgtatgtatttttctttttatttttattttcttcag*ATTTCCAAGTGGACAAGTGGATTGTAAATATACTGATGGTCATATACAAATACTATTTTGTGATGGAAAAAGAAAAGAAATACTACCCAACAGGGAAGAGTATGTTATATTACGAAACG*gtaaaatattaaaattaatataaaaatacacttatataaaatattaagaataaaacatatgtatattctttttttcttatttaaacatttttttattttaatacatttataactacaacgtcaatattataaataaatatgataaaaatatatttttttttttttttttttttttttttttccatttccctacag*GAACTATAAAAAAGTTAAAT**CCTAGGTCAGGATTGAGATCAAGATCTGCTGCTGCTGGTGCTGGTGGTGCTGCTAGAGCTGCTCTGCAGAGAGGAGTACAAGTTGAAACAATATCACCAGGAGATGGTCGTACATTTCCAAAAAGAGGTCAAACTTGTGTTGTACATTATACTGGAATGCTTGAAGATGGAAAGAAATTTGATTCATCTCGTGATAGAAATAAACCATTTAAATTTATGCTAGGTAAACAAGAAGTAATACGAGGTTGGGAAGAAGGAGTTGCTCAAATGAGTGTAGGTCAAAGAGCAAAACTTACTATATCTCCAGATTATGCTTATGGTGCAACTGGACATCCAGGTATAATTCCACCTCATGCAACTCTTGTATTTGATGTGGAGCTTCTAAAACTAGAAACTAGAGGTGTTCAGGTTGAAACAATTTCACCTGGAGATGGCAGAACCTTTCCTAAAAGAGGACAGACTTGCGTAGTTCATTATACAGGCATGCTAGAGGATGGTAAGAAATTTGATTCTAGTCGAGATAGAAATAAGCCATTCAAGTTTATGCTAGGTAAACAGGAAGTAATAAGAGGTTGGGAAGAGGGTGTAGCACAGATGTCAGTTGGACAAAGAGCAAAGTTAACAATATCACCAGATTATGCATACGGTGCAACAGGCCATCCTGGCATCATCCCTCCACATGCAACTTTAGTATTCGACGTTGAATTGTTAAAGTTAGAGACAacgcgtGCTAGAGGTGCTGCTGCTGGTGCTGGAGGTGCAGGTAGAcgtacgATGGCAGAAATTGGTACGGGTTTTCCATTTGATCCTCATTATGTGGAGGTGCTGGGGGAAAGGATGCATTATGTTGACGTAGGACCAAGAGATGGTACTCCAGTGTTATTTTTGCATGGAAACCCAACCTCGAGTTATGTATGGAGAAATATAATTCCACATGTAGCACCAACACATAGATGTATAGCTCCTGATTTAATTGGTATGGGAAAAAGTGATAAACCTGACTTAGGATATTTTTTTGATGATCATGTCCGTTTTATGGATGCTTTCATTGAAGCTCTTGGCCTTGAAGAAGTAGTATTAGTTATACATGATTGGGGATCCGCCTTAGGATTTCATTGGGCCAAGAGGAATCCTGAAAGAGTAAAAGGAATAGCATTCATGGAATTCATACGACCAATCCCCACATGGGATGAATGGCCAGAATTTGCACGCGAAACATTTCAAGCTTTTAGAACTACAGATGTTGGTAGAAAATTAATAATAGATCAAAATGTATTTATAGAAGGAACTTTACCTATGGGTGTTGTAAGGCCGTTAACAGAAGTTGAAATGGACCACTACCGTGAACCTTTTTTAAATCCAGTAGATAGAGAGCCCTTATGGAGATTTCCTAATGAATTACCTATTGCAGGTGAACCCGCGAATATTGTTGCTTTAGTAGAAGAATATATGGATTGGTTACATCAGTCTCCTGTTCCTAAACTTCTATTTTGGGGTACACCTGGAGTTCTAATACCACCAGCTGAAGCAGCAAGATTAGCAAAATCATTACCAAATTGTAAAGCTGTTGATATAGGTCCTGGGTTGAATTTATTACAAGAAGATAATCCAGATTTGATTGGATCTGAGATAGCTAGATGGCTAAGTACATTAGAAATTTCAGGTACCGGTGCCAGGGGAGCAGCCGCAGGAGCAGGGGGGGCAGGAAGGCGTGGTGTTCAGGTCGAGACTATTAGCCCTGGAGATGGACGCACGTTTCCTAAGCGTGGACAGACATGCGTAGTTCACTACACAGGTATGTTGGAGGACGGTAAAAAGTTCGACAGCTCACGCGACCGCAATAAACCTTTCAAGTTTATGCTTGGCAAGCAGGAGGTTATTCGTGGATGGGAGGAGGGTGTAGCACAGATGTCTGTTGGACAGCGTGCTAAGTTGACAATTTCACCTGACTATGCTTATGGCGCTACGGGCCATCCCGGGATCATTCCGCCACATGCGACTCTGGTATTCGACGTTGAATTATTAAAGTTAGAGACAGCTAGAGGGGCCGCTGCAGGTGCTGGTGGAGCTGGAAGACGTGGAGTACAAGTAGAGACTATCTCTCCAGGTGACGGTCGCACTTTCCCAAAGCGTGGCCAAACCTGTGTTGTACATTACACTGGTATGCTGGAGGATGGGAAAAAGTTCGATTCCAGTCGCGACCGTAACAAACCGTTCAAATTCATGTTGGGAAAGCAGGAAGTGATCCGCGGGTGGGAGGAAGGCGTGGCGCAAATGAGCGTCGGTCAGCGGGCTAAATTGACCATTTCCCCTGACTACGCGTATGGGGCTACTGGGCACCCAGGGATTATTCCGCCTCACGCTACACTTGTGTTTGATGTCGAACTTTTGAAACTGGAAACTGTCGACGGAGAAGGAAGAGGAAGTTTATTAACATGTGGAGATGTAGAAGAAAATCCAGGACCAATGATTGAACAAGATGGATTGCACGCAGGTTCTCCGGCCGCTTGGGTGGAGAGGCTATTCGGCTATGACTGGGCACAACAGACAATCGGCTGCTCTGATGCCGCCGTGTTCCGGCTGTCAGCGCAGGGGCGCCCGGTTCTTTTTGTCAAGACCGACCTGTCCGGTGCCCTGAATGAACTGCAGGACGAGGCAGCGCGGCTATCGTGGCTGGCCACGACGGGCGTTCCTTGCGCAGCTGTGCTCGACGTTGTCACTGAAGCGGGAAGGGACTGGCTGCTATTGGGCGAAGTGCCGGGGCAGGATCTCCTGTCATCTCACCTTGCTCCTGCCGAGAAAGTATCCATCATGGCTGATGCAATGCGGCGGCTGCATACGCTTGATCCGGCTACCTGCCCATTCGACCACCAAGCGAAACATCGCATCGAGCGAGCACGTACTCGGATGGAAGCCGGTCTTGTCGATCAGGATGATCTGGACGAAGAGCATCAGGGGCTCGCGCCAGCCGAACTGTTCGCCAGGCTCAAGGCGCGCATGCCCGACGGCGAGGATCTCGTCGTGACCCATGGCGATGCCTGCTTGCCGAATATCATGGTGGAAAATGGCCGCTTTTCTGGATTCATCGACTGTGGCCGGCTGGGTGTGGCGGACCGCTATCAGGACATAGCGTTGGCTACCCGTGATATTGCTGAAGAGCTTGGCGGCGAATGGGCTGACCGCTTCCTCGTGCTTTACGGTATCGCCGCTCCCGATTCGCAGCGCATCGCCTTCTATCGCCTTCTTGACGAGTTCTTCTAActcgagggatatggcagcttaatgttcgtttttcttatttatatatttataccaattgattgtatttataactgtaaaaatgtgtatgttgtgtgcatatttttttttgtgcatgcacatgcatgtaaatagctaaaattatgaacattttattttttgttcagaaaaaaaaaactttacacacataaaatggctagtatgaatagccatattttatataaattaaatcctatgaatttatgaccatattaaaaatttagatatttatggaacataatatgtttgaaacaataagacaaaattattattattattattatttttactgttataattatgtgtctccttcaatgattcataaatagttggacttgatttttaaaatgtttataatatgattagcatagttaaataaaaaaagttgaaaaattaaaaaaaaacatataaacacaaatgatggtttttccttcaatttcgatatcaatttatagaaacaaaatatatacttgtataattttatttttttatataaatcattacatatataattatacaatattttttctaagagataattatatattaatatatataaaaaaaggtgttttttttttttttttttatttttatttttattttatggtaatattttattttccttattttataaattatattagtttatatgtgattaattttatatattatcaatttatatatttttaaatgcttacttaattatctttttttttttttttttttttttttcccctctttttatattaatttatttttgaaaaaattgatatatatatatatatataatatatatatatacatgtagtagtattaaacaatgtataatatatataaataatatatttatatatttcatttcaattttaattttttttggttttttttttttttctttttgtcatatttaaaaaaaattatattcatataagttatgcattttttataaacattattcaatatatgtataatataatatatatatatatattaatgtattattccaatgtgcatgataaaagaaaaaaataatatttataaaaaaaaagaaaaataaaacaaaaaaagaaaaaaaaaaaaaaaaaaaaaaaaatacaaaaataaataatataatttataattatatattcttgtcacaataaaaatatatatatatatatatatatttataatatgtatattttaaactagaaaaggaataactaatattttatttattatcattcaagatttatattttataataataaatacctaatagaaatatatcaggatccatgcatggttcgctaaactgcatcgtcgctgtgtcccagaacatgggcatcggcaagaacggggactacccctggccaccgctcaggaacgaatttagatatttccagagaatgaccacaacctcttcagtagaaggtaaacagaatctggtgattatgggtaagaagacctggttctccattcctgagaagaatcgacctttaaagggtagaattaatttagttctcagcagagaactcaaggaacctccacaaggagctcattttctttccagaagtctagatgatgccttaaaacttactgaacaaccagaattagcaaataaagtagacatggtctggatagttggtggcagttctgtttataaggaagccatgaatcacccaggccatcttaaactatttgtgacaaggatcatgcaagactttgaaagtgacacgttttttccagaaattgatttggagaaatataaacttctgccagaatacccaggtgttctctctgatgtccaggaggagaaaggcattaagtacaaatttgaagtatatgagaagaatgattaagcttatttaataatagattaaaaatattataaaaataaaaacataaacacagaaattacaaaaaaaatacatatgaattttttttttgtaatcttccttataaatatagaataatgaatcatataaaacatatcattattcatttatttacatttaaaattattgtttcagtatctttaatttattatgtatatataaaaataacttacaattttattaataaacaatatatgtttattaattcatgttttgtaatttatgggatagcgattttttttactgtctgtatttttcttttttaattatgttttaattgtattttatttttattattgttctttttatagtattattttaaaacaaaatgtattttctaagaacttataataataataaatataaattttaataaaaattatatttatcttttacaatatgaacataaagtacaacattaatatatagcttttaatatttttattcctaatcatgtaaatcttaaatttttctttttaaacatatgttaaatatttatttctcattatatataagaacatatttattaaatctagaattctatagtgagtcgtattacaattcactggccgtcgttttacaacgtcgtgactgggaaaaccctggcgttacccaacttaatcgccttgcagcacatccccctttcgccagctggcgtaatagcgaagaggcccgcaccgatcgcccttcccaacagttgcgcagcctgaatggcgaatggcgcctgatgcggtattttctccttacgcatctgtgcggtatttcacaccgcatatggtgcactctcagtacaatctgctctgatgccgcatagttaagccagccccgacacccgccaacacccgctgacgcgccctgacgggcttgtctgctcccggcatccgcttacagacaagctgtgaccgtctccgggagctgcatgtgtcagaggttttcaccgtcatcaccgaaacgcgcgagacgaaagggcctcgtgatacgcctatttttataggttaatgtcatgataataatggtttcttagacgtcaggtggcacttttcggggaaatgtgcgcggaacccctatttgtttatttttctaaatacattcaaatatgtatccgctcatgagacaataaccctgataaatgcttcaataatattgaaaaaggaagagtatgagtattcaacatttccgtgtcgcccttattcccttttttgcggcattttgccttcctgtttttgctcacccagaaacgctggtgaaagtaaaagatgctgaagatcagttgggtgcacgagtgggttacatcgaactggatctcaacagcggtaagatccttgagagttttcgccccgaagaacgttttccaatgatgagcacttttaaagttctgctatgtggcgcggtattatcccgtattgacgccgggcaagagcaactcggtcgccgcatacactattctcagaatgacttggttgagtactcaccagtcacagaaaagcatcttacggatggcatgacagtaagagaattatgcagtgctgccataaccatgagtgataacactgcggccaacttacttctgacaacgatcggaggaccgaaggagctaaccgcttttttgcacaacatgggggatcatgtaactcgccttgatcgttgggaaccggagctgaatgaagccataccaaacgacgagcgtgacaccacgatgcctgtagcaatgccaacaacgttgcgcaaactattaactggcgaactacttactctagcttcccggcaacaattaatagactggatggaggcggataaagttgcaggaccacttctgcgctcggcccttccggctggctggtttattgctgataaatctggagccggtgagcgtgggtctcgcggtatcattgcagcactggggccagatggtaagccctcccgtatcgtagttatctacacgacggggagtcaggcaactatggatgaacgaaatagacagatcgctgagataggtgcctcactgattaagcattggtaactgtcagaccaagtttactcatatatactttagattgatttaaaacttcatttttaatttaaaaggatctaggtgaagatcctttttgataatctcatgaccaaaatcccttaacgtgagttttcgttccactgagcgtcagaccccgtagaaaagatcaaaggatcttcttgagatcctttttttctgcgcgtaatctgctgcttgcaaacaaaaaaaccaccgctaccagcggtggtttgtttgccggatcaagagctaccaactctttttccgaaggtaactggcttcagcagagcgcagataccaaatactgtccttctagtgtagccgtagttaggccaccacttcaagaactctgtagcaccgcctacatacctcgctctgctaatcctgttaccagtggctgctgccagtggcgataagtcgtgtcttaccgggttggactcaagacgatagttaccggataaggcgcagcggtcgggctgaacggggggttcgtgcacacagcccagcttggagcgaacgacctacaccgaactgagatacctacagcgtgagctatgagaaagcgccacgcttcccgaagggagaaaggcggacaggtatccggtaagcggcagggtcggaacaggagagcgcacgagggagcttccagggggaaacgcctggtatctttatagtcctgtcgggtttcgccacctctgacttgagcgtcgatttttgtgatgctcgtcaggggggcggagcctatcgaaaaacgccagcaacgcggcctttttacggttcctggccttttgctggccttttgctcacatgttctttcctgcgttatcccctgattctgtggataaccgtattaccgcctttgagtgagctgataccgctcgccgcagccgaacgaccgagcgcagcgagtcagtgagcgaggaagcggaagagcgcccaatacgcaaaccgcctctccccgcgcgttggccgattcattaatgcagctggcacgacaggtttcccgactggaaagcgggcagtgagcgcaacgcaattaatgtgagttagctcactcattaggcaccccaggctttacactttatgcttccggctcgtatgttgtgtggaattgtgagcggataacaatttcacacaggaaacagctatgaccatgattacgccaagctatttaggtgacactatagaagcggccgcTAgGGTAATGATAATATTTTAAAACCTTTTGGTAAGGACTGGGATTTTGTAATAAACTTTGATTTCGACGAATTATTCAATCAATGTGAAAATGTTATAGAATCTATTTTTTCCTCATCCAAAAAAATAAAATATAGACAAGCATTTATTGATGGAAAGGTGGAAACTTTATTTGATGATGGGTTAAAATTAATTGAGAAAAATAGAAATAAAAAAATTATGCATCCAAGTAATATAACAATATATCTGTATCCTACCAAAGATTACAAGGCACATTTTCCCAATTCGTACATGgtaaaatttaagaaaaaataaaaaaaaaaaaatttaaatgagcacatatatatatttatgtatatatatatgaccatattggtcttgctatatatcatacatttttataataatattatgtctgctttttagTTATTTCGATTTGTTAATAAAGGAATATATCAGGTGAACATACCAAATAAATGTCAACTAAACAAgtaaataagaaaaaaaaaaaaaaaaataataataataataaaacatcgttagagattattttttccattacatatatgtacatatatatatatatatatatatatatatacatttttttttatatgtatgtatttttctttttatttttattttcttcagATTTCCAAGTGGACAAGTGGATTGTAAATATACTGATGGTCATATACAAATACTATTTTGTGATGGAAAAAGAAAAGAAATACTACCCAACAGGGAAGAGTATGTTATATTACGAAACGgtaaaatattaaaattaatataaaaatacacttatataaaatattaagaataaaacatatgtatattctttttttcttatttaaacatttttttattttaatacatttataactacaacgtcaatattataaataaatatgataaaaatatatttttttttttttttttttttttttttttccatttccctacagGAACTATAAAAAAGTTAAATTAAtatgatgtgttcacttatatgaatatatgttggatatatgattaactcatgaaatttaatgttctattcataattgaggatataacaatataaacatttaaaatgtcaatgaaaaataaaggaacaaataaattttatattttat

**5’ amplicon size: 1232bps**

**3’ amplicon size: 1100bps**

## **> PF3D7_1461100**

tttttttctttatatttctttttgggtttgttctaataaatatgttaaattaaaaaaataaacataaaataaaatagaggaaaagaaaagaATGAATGAAAATTTGAATAATGTATCAGAGGAATTTAAAAAGCAGCTTTTATCTG*gtatgttaatctatatatgatgagtaaatttatgggtgttataagtatatatatatatatattatttatttatttattttattactattttgtttgttttttttttttttttttttcatgtag*CTACTATAAAAAGGGAAGTAAAAGAAAATATGGAAATAAATGTTAAACAGAAACCTAATGATAATAAAAATAATAATGATTTTATAATTTATATGGAATTACACCTGGAAAATTTGGTGATTAGGAGATATAATTTTGTCCTAGG*gtaagagtataaaaaaaaaaaaataaaataaaatattaagatacaatatgtattgttgaatagtatttatattgtatatatataattaaatatatatatatatattgatgtatacatatatatatatatattttttttttttttttttttttttag*TAACGAATTGCAGTCTCCCATTCGGAATAGTGTTCAGATAACTAGGTTCCTAATTATGAAGAGTCAAAATCATATAATGGGTACATGTGTTTCTGACAAGATAATATTATGTTATAGCAATGATAATGATACAAAAATGATATTAAATAGGTTAATTGATGAATTAGATAAGAATTATTTTATTACCTTAATAATGGAAGGTGATACAAACAGTGATGATGAAAATGATTATGATGAAGATAATAATCACAATAATAATCACAATAATAATCATAATAATAATCACAATAATTATCACAATAATAATCACAATAATAATCACAATTATAATCACAATAATAATCACAATAATTATCACAATAATAATCAGAACAAATATGATAATCGTCATATTGAAAAGTGCTTGTTATCTTACATAATACAATTTACAATAGTTTGTAAATACCTAGAGTGCGGGAAATGGTTTCATATATGGGATAATATGAATACAATTATTGAATCCAAATTTTGTGGGAATAATAACTGTAAATATTTTAATTGTCTTGGATTTAAATTTAATATGTTAAATAGTGATTATAATATATTGAATGAAGAGGAAAAGGGATCTCATATATATATTATGAAATTACATTTAAGGGTAACAGTAAATATTTATAAGATATTACCCTTATGTGATATTAATGATAATATTAATGTATATTGTTTACCAAGGTGTTCTATGAAAGCAACAATATTGAATATGATTTCATCATCATCAGCAAATGATCCGAATATTAAAACATTTGATTATAAAAATTATTGGTTAGATATACATGGATATTTATTAAATGATATGAGTATAGATAAAATTGTACAAGTTAGATTATATAAAGGTATATTCAATTACCCACAAGGGGTATTATTAAGAGATAATATATATAAATTAAATATACCTATTAGAAAGGAGCATACATTATATATATGTAGTTATCTGAAGCAATTTCAATTACTAACAAAATCGAAAATGAATATATTGAAACATTCAGACATTTTACATTTTCAAAGTATATGTAATGAGTTAATGGAAAATAATATGAACATTAAGAAAAAGGAAGATGTCGAATTTTACAAAAAGATTGATCATCTATGTAATCATGTCATAGAAGATGATAAGGGAAATAACAACGAAAAAAACAACAATAATAATTATAGTAATAATGATAGTAATAATGATAGTAATAATAATAGTAATAATGATAGTAATAATAATAGTAATAGTTGTTATAATAATAGTTAT**AATAATGCCTATCGTACATATGGAAACACATTATTCAGTTCTATGAAATCCATAAGTTTAGATTTCATTAAAAAGGAAAAAACACTAGACGGACAAAAAGATAATCAGAAAAATAACAAGGAAGAACTTAGCAACAAAATTGATTTGTATTTAAAAGAGTCAGCTATAAATTACAGAATGATGAACAATGAAAAAAATACAAATATTTCAAATAATAAAACTTCATCTAACCATTCAAATATATCAAATATATCAAATAAATCAAACAAATCTAATATATCAAACAAATCTAATATATCAAAAATATCTAACGAATCGTTTAATTCAATTAATTCAAATCAGTCTGATGAATCTTTCAATTCTATTAAGTCCAATAATTCAAATAATACAAATGAACAAAATATTCATATAAAAAATGATACATTTTATAATAATGATAATCAGAATAAATCATACCATCAAAGTAGTAGTTTAAGGCAAAATGAAGAAAACCAACAGAACCATAAAAATAATATTCCCTATAATCATAATGAATTATACCAAAAAAAAAATAACGTTGGTATTGAAACAGAATATAAAAAGAACAGTACGGAAAATTTAAATAAATATCTAAATGAAGTGTCACAGAATTATAAACATATAAATAATACCAAGAATAAAAATAATAATTATAATATTACAGAAGATGATGATAATTATCTAATGCATGATTTAGACTCATTTGAAAGAGTTAATGCTCATGTAATTCTTTCTAAAGAATTTCAAAACTTTCTTGATAATCTAGATGAAGTTGAATTATATACAAATGAAAAATTAAAAATTATTGATTTTAATGAACGTGAAAAAAAAAGACCACATTTATATATTAAAAATTCAAATGAAAATACTCTTGAAAAGATCGATGAAAATGATAATAGCTTATCTAACGTACATATAGATACAACATTACCTATGGAGGATGTACCTGCATATACTGAACAAAATAAAAAAATTAAAAAG**TGAcatgaatatgagaattattaataatatgaggtttatttaaaacaaggagaaatacatggattttaaaatgaaaattgtgatcactaataaggagaaattatataaaaataataaatatatatatatatatatatatatatatataattaaatatatgttatttaaaattaaaattaatcttttttttgaaaatctttaatatgtataaaaatatatacattctttaatatataataatatatacatttataaattatttaaaaaatatatatttattttttttaaattttatttttttataagtattgattttaataagtaaataaataaataaataaataaatatatatatatatatatatatatatattataatatatttatcaagaataaaatattaattattttttttgttttcatatatatattattatttttttttatattttttttttttttgtatttaattaacttaatatatatattcaactctttttccgggtatgtaaaatatataaaaatatatatttatatataatatgaaataatataat

**Original Locus (OL) amplicon size: 1184bp**s

**> PF3D7_1461100-HALO-SW Halo Neo-R**

tttttttctttatatttctttttgggtttgttctaataaatatgttaaattaaaaaaataaacataaaataaaatagaggaaaagaaaagaATGAATGAAAATTTGAATAATGTATCAGAGGAATTTAAAAAGCAGCTTTTATCTG*gtatgttaatctatatatgatgagtaaatttatgggtgttataagtatatatatatatatattatttatttatttattttattactattttgtttgttttttttttttttttttttcatgtag*CTACTATAAAAAGGGAAGTAAAAGAAAATATGGAAATAAATGTTAAACAGAAACCTAATGATAATAAAAATAATAATGATTTTATAATTTATATGGAATTACACCTGGAAAATTTGGTGATTAGGAGATATAATTTTGTCCTAGG*gtaagagtataaaaaaaaaaaaataaaataaaatattaagatacaatatgtattgttgaatagtatttatattgtatatatataattaaatatatatatatatattgatgtatacatatatatatatatattttttttttttttttttttttttag*TAACGAATTGCAGTCTCCCATTCGGAATAGTGTTCAGATAACTAGGTTCCTAATTATGAAGAGTCAAAATCATATAATGGGTACATGTGTTTCTGACAAGATAATATTATGTTATAGCAATGATAATGATACAAAAATGATATTAAATAGGTTAATTGATGAATTAGATAAGAATTATTTTATTACCTTAATAATGGAAGGTGATACAAACAGTGATGATGAAAATGATTATGATGAAGATAATAATCACAATAATAATCACAATAATAATCATAATAATAATCACAATAATTATCACAATAATAATCACAATAATAATCACAATTATAATCACAATAATAATCACAATAATTATCACAATAATAATCAGAACAAATATGATAATCGTCATATTGAAAAGTGCTTGTTATCTTACATAATACAATTTACAATAGTTTGTAAATACCTAGAGTGCGGGAAATGGTTTCATATATGGGATAATATGAATACAATTATTGAATCCAAATTTTGTGGGAATAATAACTGTAAATATTTTAATTGTCTTGGATTTAAATTTAATATGTTAAATAGTGATTATAATATATTGAATGAAGAGGAAAAGGGATCTCATATATATATTATGAAATTACATTTAAGGGTAACAGTAAATATTTATAAGATATTACCCTTATGTGATATTAATGATAATATTAATGTATATTGTTTACCAAGGTGTTCTATGAAAGCAACAATATTGAATATGATTTCATCATCATCAGCAAATGATCCGAATATTAAAACATTTGATTATAAAAATTATTGGTTAGATATACATGGATATTTATTAAATGATATGAGTATAGATAAAATTGTACAAGTTAGATTATATAAAGGTATATTCAATTACCCACAAGGGGTATTATTAAGAGATAATATATATAAATTAAATATACCTATTAGAAAGGAGCATACATTATATATATGTAGTTATCTGAAGCAATTTCAATTACTAACAAAATCGAAAATGAATATATTGAAACATTCAGACATTTTACATTTTCAAAGTATATGTAATGAGTTAATGGAAAATAATATGAACATTAAGAAAAAGGAAGATGTCGAATTTTACAAAAAGATTGATCATCTATGTAATCATGTCATAGAAGATGATAAGGGAAATAACAACGAAAAAAACAACAATAATAATTATAGTAATAATGATAGTAATAATGATAGTAATAATAATAGTAATAATGATAGTAATAATAATAGTAATAGTTGTTATAATAATAGTTAT**AATAATGCCTATCGTACATATGGAAACACATTATTCAGTTCTATGAAATCCATAAGTTTAGATTTCATTAAAAAGGAAAAAACACTAGACGGACAAAAAGATAATCAGAAAAATAACAAGGAAGAACTTAGCAACAAAATTGATTTGTATTTAAAAGAGTCAGCTATAAATTACAGAATGATGAACAATGAAAAAAATACAAATATTTCAAATAATAAAACTTCATCTAACCATTCAAATATATCAAATATATCAAATAAATCAAACAAATCTAATATATCAAACAAATCTAATATATCAAAAATATCTAACGAATCGTTTAATTCAATTAATTCAAATCAGTCTGATGAATCTTTCAATTCTATTAAGTCCAATAATTCAAATAATACAAATGAACAAAATATTCATATAAAAAATGATACATTTTATAATAATGATAATCAGAATAAATCATACCATCAAAGTAGTAGTTTAAGGCAAAATGAAGAAAACCAACAGAACCATAAAAATAATATTCCCTATAATCATAATGAATTATACCAAAAAAAAAATAACGTTGGTATTGAAACAGAATATAAAAAGAACAGTACGGAAAATTTAAATAAATATCTAAATGAAGTGTCACAGAATTATAAACATATAAATAATACCAAGAATAAAAATAATAATTATAATATTACAGAAGATGATGATAATTATCTAATGCATGATTTAGACTCATTTGAAAGAGTTAATGCTCATGTAATTCTTTCTAAAGAATTTCAAAACTTTCTTGATAATCTAGATGAAGTTGAATTATATACAAATGAAAAATTAAAAATTATTGATTTTAATGAACGTGAAAAAAAAAGACCACATTTATATATTAAAAATTCAAATGAAAATACTCTTGAAAAGATCGATGAAAATGATAATAGCTTATCTAACGTACATATAGATACAACATTACCTATGGAGGATGTACCTGCATATACTGAACAAAATAAAAAAATTAAAAAG**CCTAGGTCAGGATTGAGATCAAGATCTGCTGCTGCTGGTGCTGGTGGTGCTGCTAGAGCTGCTCTGCAGAGAGGAGTACAAGTTGAAACAATATCACCAGGAGATGGTCGTACATTTCCAAAAAGAGGTCAAACTTGTGTTGTACATTATACTGGAATGCTTGAAGATGGAAAGAAATTTGATTCATCTCGTGATAGAAATAAACCATTTAAATTTATGCTAGGTAAACAAGAAGTAATACGAGGTTGGGAAGAAGGAGTTGCTCAAATGAGTGTAGGTCAAAGAGCAAAACTTACTATATCTCCAGATTATGCTTATGGTGCAACTGGACATCCAGGTATAATTCCACCTCATGCAACTCTTGTATTTGATGTGGAGCTTCTAAAACTAGAAACTAGAGGTGTTCAGGTTGAAACAATTTCACCTGGAGATGGCAGAACCTTTCCTAAAAGAGGACAGACTTGCGTAGTTCATTATACAGGCATGCTAGAGGATGGTAAGAAATTTGATTCTAGTCGAGATAGAAATAAGCCATTCAAGTTTATGCTAGGTAAACAGGAAGTAATAAGAGGTTGGGAAGAGGGTGTAGCACAGATGTCAGTTGGACAAAGAGCAAAGTTAACAATATCACCAGATTATGCATACGGTGCAACAGGCCATCCTGGCATCATCCCTCCACATGCAACTTTAGTATTCGACGTTGAATTGTTAAAGTTAGAGACAacgcgtGCTAGAGGTGCTGCTGCTGGTGCTGGAGGTGCAGGTAGAcgtacgATGGCAGAAATTGGTACGGGTTTTCCATTTGATCCTCATTATGTGGAGGTGCTGGGGGAAAGGATGCATTATGTTGACGTAGGACCAAGAGATGGTACTCCAGTGTTATTTTTGCATGGAAACCCAACCTCGAGTTATGTATGGAGAAATATAATTCCACATGTAGCACCAACACATAGATGTATAGCTCCTGATTTAATTGGTATGGGAAAAAGTGATAAACCTGACTTAGGATATTTTTTTGATGATCATGTCCGTTTTATGGATGCTTTCATTGAAGCTCTTGGCCTTGAAGAAGTAGTATTAGTTATACATGATTGGGGATCCGCCTTAGGATTTCATTGGGCCAAGAGGAATCCTGAAAGAGTAAAAGGAATAGCATTCATGGAATTCATACGACCAATCCCCACATGGGATGAATGGCCAGAATTTGCACGCGAAACATTTCAAGCTTTTAGAACTACAGATGTTGGTAGAAAATTAATAATAGATCAAAATGTATTTATAGAAGGAACTTTACCTATGGGTGTTGTAAGGCCGTTAACAGAAGTTGAAATGGACCACTACCGTGAACCTTTTTTAAATCCAGTAGATAGAGAGCCCTTATGGAGATTTCCTAATGAATTACCTATTGCAGGTGAACCCGCGAATATTGTTGCTTTAGTAGAAGAATATATGGATTGGTTACATCAGTCTCCTGTTCCTAAACTTCTATTTTGGGGTACACCTGGAGTTCTAATACCACCAGCTGAAGCAGCAAGATTAGCAAAATCATTACCAAATTGTAAAGCTGTTGATATAGGTCCTGGGTTGAATTTATTACAAGAAGATAATCCAGATTTGATTGGATCTGAGATAGCTAGATGGCTAAGTACATTAGAAATTTCAGGTACCGGTGCCAGGGGAGCAGCCGCAGGAGCAGGGGGGGCAGGAAGGCGTGGTGTTCAGGTCGAGACTATTAGCCCTGGAGATGGACGCACGTTTCCTAAGCGTGGACAGACATGCGTAGTTCACTACACAGGTATGTTGGAGGACGGTAAAAAGTTCGACAGCTCACGCGACCGCAATAAACCTTTCAAGTTTATGCTTGGCAAGCAGGAGGTTATTCGTGGATGGGAGGAGGGTGTAGCACAGATGTCTGTTGGACAGCGTGCTAAGTTGACAATTTCACCTGACTATGCTTATGGCGCTACGGGCCATCCCGGGATCATTCCGCCACATGCGACTCTGGTATTCGACGTTGAATTATTAAAGTTAGAGACAGCTAGAGGGGCCGCTGCAGGTGCTGGTGGAGCTGGAAGACGTGGAGTACAAGTAGAGACTATCTCTCCAGGTGACGGTCGCACTTTCCCAAAGCGTGGCCAAACCTGTGTTGTACATTACACTGGTATGCTGGAGGATGGGAAAAAGTTCGATTCCAGTCGCGACCGTAACAAACCGTTCAAATTCATGTTGGGAAAGCAGGAAGTGATCCGCGGGTGGGAGGAAGGCGTGGCGCAAATGAGCGTCGGTCAGCGGGCTAAATTGACCATTTCCCCTGACTACGCGTATGGGGCTACTGGGCACCCAGGGATTATTCCGCCTCACGCTACACTTGTGTTTGATGTCGAACTTTTGAAACTGGAAACTGTCGACGGAGAAGGAAGAGGAAGTTTATTAACATGTGGAGATGTAGAAGAAAATCCAGGACCAATGATTGAACAAGATGGATTGCACGCAGGTTCTCCGGCCGCTTGGGTGGAGAGGCTATTCGGCTATGACTGGGCACAACAGACAATCGGCTGCTCTGATGCCGCCGTGTTCCGGCTGTCAGCGCAGGGGCGCCCGGTTCTTTTTGTCAAGACCGACCTGTCCGGTGCCCTGAATGAACTGCAGGACGAGGCAGCGCGGCTATCGTGGCTGGCCACGACGGGCGTTCCTTGCGCAGCTGTGCTCGACGTTGTCACTGAAGCGGGAAGGGACTGGCTGCTATTGGGCGAAGTGCCGGGGCAGGATCTCCTGTCATCTCACCTTGCTCCTGCCGAGAAAGTATCCATCATGGCTGATGCAATGCGGCGGCTGCATACGCTTGATCCGGCTACCTGCCCATTCGACCACCAAGCGAAACATCGCATCGAGCGAGCACGTACTCGGATGGAAGCCGGTCTTGTCGATCAGGATGATCTGGACGAAGAGCATCAGGGGCTCGCGCCAGCCGAACTGTTCGCCAGGCTCAAGGCGCGCATGCCCGACGGCGAGGATCTCGTCGTGACCCATGGCGATGCCTGCTTGCCGAATATCATGGTGGAAAATGGCCGCTTTTCTGGATTCATCGACTGTGGCCGGCTGGGTGTGGCGGACCGCTATCAGGACATAGCGTTGGCTACCCGTGATATTGCTGAAGAGCTTGGCGGCGAATGGGCTGACCGCTTCCTCGTGCTTTACGGTATCGCCGCTCCCGATTCGCAGCGCATCGCCTTCTATCGCCTTCTTGACGAGTTCTTCTAActcgagggatatggcagcttaatgttcgtttttcttatttatatatttataccaattgattgtatttataactgtaaaaatgtgtatgttgtgtgcatatttttttttgtgcatgcacatgcatgtaaatagctaaaattatgaacattttattttttgttcagaaaaaaaaaactttacacacataaaatggctagtatgaatagccatattttatataaattaaatcctatgaatttatgaccatattaaaaatttagatatttatggaacataatatgtttgaaacaataagacaaaattattattattattattatttttactgttataattatgtgtctccttcaatgattcataaatagttggacttgatttttaaaatgtttataatatgattagcatagttaaataaaaaaagttgaaaaattaaaaaaaaacatataaacacaaatgatggtttttccttcaatttcgatatcaatttatagaaacaaaatatatacttgtataattttatttttttatataaatcattacatatataattatacaatattttttctaagagataattatatattaatatatataaaaaaaggtgttttttttttttttttttatttttatttttattttatggtaatattttattttccttattttataaattatattagtttatatgtgattaattttatatattatcaatttatatatttttaaatgcttacttaattatctttttttttttttttttttttttttcccctctttttatattaatttatttttgaaaaaattgatatatatatatatatataatatatatatatacatgtagtagtattaaacaatgtataatatatataaataatatatttatatatttcatttcaattttaattttttttggttttttttttttttctttttgtcatatttaaaaaaaattatattcatataagttatgcattttttataaacattattcaatatatgtataatataatatatatatatatattaatgtattattccaatgtgcatgataaaagaaaaaaataatatttataaaaaaaaagaaaaataaaacaaaaaaagaaaaaaaaaaaaaaaaaaaaaaaaatacaaaaataaataatataatttataattatatattcttgtcacaataaaaatatatatatatatatatatatttataatatgtatattttaaactagaaaaggaataactaatattttatttattatcattcaagatttatattttataataataaatacctaatagaaatatatcaggatccatgcatggttcgctaaactgcatcgtcgctgtgtcccagaacatgggcatcggcaagaacggggactacccctggccaccgctcaggaacgaatttagatatttccagagaatgaccacaacctcttcagtagaaggtaaacagaatctggtgattatgggtaagaagacctggttctccattcctgagaagaatcgacctttaaagggtagaattaatttagttctcagcagagaactcaaggaacctccacaaggagctcattttctttccagaagtctagatgatgccttaaaacttactgaacaaccagaattagcaaataaagtagacatggtctggatagttggtggcagttctgtttataaggaagccatgaatcacccaggccatcttaaactatttgtgacaaggatcatgcaagactttgaaagtgacacgttttttccagaaattgatttggagaaatataaacttctgccagaatacccaggtgttctctctgatgtccaggaggagaaaggcattaagtacaaatttgaagtatatgagaagaatgattaagcttatttaataatagattaaaaatattataaaaataaaaacataaacacagaaattacaaaaaaaatacatatgaattttttttttgtaatcttccttataaatatagaataatgaatcatataaaacatatcattattcatttatttacatttaaaattattgtttcagtatctttaatttattatgtatatataaaaataacttacaattttattaataaacaatatatgtttattaattcatgttttgtaatttatgggatagcgattttttttactgtctgtatttttcttttttaattatgttttaattgtattttatttttattattgttctttttatagtattattttaaaacaaaatgtattttctaagaacttataataataataaatataaattttaataaaaattatatttatcttttacaatatgaacataaagtacaacattaatatatagcttttaatatttttattcctaatcatgtaaatcttaaatttttctttttaaacatatgttaaatatttatttctcattatatataagaacatatttattaaatctagaattctatagtgagtcgtattacaattcactggccgtcgttttacaacgtcgtgactgggaaaaccctggcgttacccaacttaatcgccttgcagcacatccccctttcgccagctggcgtaatagcgaagaggcccgcaccgatcgcccttcccaacagttgcgcagcctgaatggcgaatggcgcctgatgcggtattttctccttacgcatctgtgcggtatttcacaccgcatatggtgcactctcagtacaatctgctctgatgccgcatagttaagccagccccgacacccgccaacacccgctgacgcgccctgacgggcttgtctgctcccggcatccgcttacagacaagctgtgaccgtctccgggagctgcatgtgtcagaggttttcaccgtcatcaccgaaacgcgcgagacgaaagggcctcgtgatacgcctatttttataggttaatgtcatgataataatggtttcttagacgtcaggtggcacttttcggggaaatgtgcgcggaacccctatttgtttatttttctaaatacattcaaatatgtatccgctcatgagacaataaccctgataaatgcttcaataatattgaaaaaggaagagtatgagtattcaacatttccgtgtcgcccttattcccttttttgcggcattttgccttcctgtttttgctcacccagaaacgctggtgaaagtaaaagatgctgaagatcagttgggtgcacgagtgggttacatcgaactggatctcaacagcggtaagatccttgagagttttcgccccgaagaacgttttccaatgatgagcacttttaaagttctgctatgtggcgcggtattatcccgtattgacgccgggcaagagcaactcggtcgccgcatacactattctcagaatgacttggttgagtactcaccagtcacagaaaagcatcttacggatggcatgacagtaagagaattatgcagtgctgccataaccatgagtgataacactgcggccaacttacttctgacaacgatcggaggaccgaaggagctaaccgcttttttgcacaacatgggggatcatgtaactcgccttgatcgttgggaaccggagctgaatgaagccataccaaacgacgagcgtgacaccacgatgcctgtagcaatgccaacaacgttgcgcaaactattaactggcgaactacttactctagcttcccggcaacaattaatagactggatggaggcggataaagttgcaggaccacttctgcgctcggcccttccggctggctggtttattgctgataaatctggagccggtgagcgtgggtctcgcggtatcattgcagcactggggccagatggtaagccctcccgtatcgtagttatctacacgacggggagtcaggcaactatggatgaacgaaatagacagatcgctgagataggtgcctcactgattaagcattggtaactgtcagaccaagtttactcatatatactttagattgatttaaaacttcatttttaatttaaaaggatctaggtgaagatcctttttgataatctcatgaccaaaatcccttaacgtgagttttcgttccactgagcgtcagaccccgtagaaaagatcaaaggatcttcttgagatcctttttttctgcgcgtaatctgctgcttgcaaacaaaaaaaccaccgctaccagcggtggtttgtttgccggatcaagagctaccaactctttttccgaaggtaactggcttcagcagagcgcagataccaaatactgtccttctagtgtagccgtagttaggccaccacttcaagaactctgtagcaccgcctacatacctcgctctgctaatcctgttaccagtggctgctgccagtggcgataagtcgtgtcttaccgggttggactcaagacgatagttaccggataaggcgcagcggtcgggctgaacggggggttcgtgcacacagcccagcttggagcgaacgacctacaccgaactgagatacctacagcgtgagctatgagaaagcgccacgcttcccgaagggagaaaggcggacaggtatccggtaagcggcagggtcggaacaggagagcgcacgagggagcttccagggggaaacgcctggtatctttatagtcctgtcgggtttcgccacctctgacttgagcgtcgatttttgtgatgctcgtcaggggggcggagcctatcgaaaaacgccagcaacgcggcctttttacggttcctggccttttgctggccttttgctcacatgttctttcctgcgttatcccctgattctgtggataaccgtattaccgcctttgagtgagctgataccgctcgccgcagccgaacgaccgagcgcagcgagtcagtgagcgaggaagcggaagagcgcccaatacgcaaaccgcctctccccgcgcgttggccgattcattaatgcagctggcacgacaggtttcccgactggaaagcgggcagtgagcgcaacgcaattaatgtgagttagctcactcattaggcaccccaggctttacactttatgcttccggctcgtatgttgtgtggaattgtgagcggataacaatttcacacaggaaacagctatgaccatgattacgccaagctatttaggtgacactatagaagcggccgcTAgAATAATGCCTATCGTACATATGGAAACACATTATTCAGTTCTATGAAATCCATAAGTTTAGATTTCATTAAAAAGGAAAAAACACTAGACGGACAAAAAGATAATCAGAAAAATAACAAGGAAGAACTTAGCAACAAAATTGATTTGTATTTAAAAGAGTCAGCTATAAATTACAGAATGATGAACAATGAAAAAAATACAAATATTTCAAATAATAAAACTTCATCTAACCATTCAAATATATCAAATATATCAAATAAATCAAACAAATCTAATATATCAAACAAATCTAATATATCAAAAATATCTAACGAATCGTTTAATTCAATTAATTCAAATCAGTCTGATGAATCTTTCAATTCTATTAAGTCCAATAATTCAAATAATACAAATGAACAAAATATTCATATAAAAAATGATACATTTTATAATAATGATAATCAGAATAAATCATACCATCAAAGTAGTAGTTTAAGGCAAAATGAAGAAAACCAACAGAACCATAAAAATAATATTCCCTATAATCATAATGAATTATACCAAAAAAAAAATAACGTTGGTATTGAAACAGAATATAAAAAGAACAGTACGGAAAATTTAAATAAATATCTAAATGAAGTGTCACAGAATTATAAACATATAAATAATACCAAGAATAAAAATAATAATTATAATATTACAGAAGATGATGATAATTATCTAATGCATGATTTAGACTCATTTGAAAGAGTTAATGCTCATGTAATTCTTTCTAAAGAATTTCAAAACTTTCTTGATAATCTAGATGAAGTTGAATTATATACAAATGAAAAATTAAAAATTATTGATTTTAATGAACGTGAAAAAAAAAGACCACATTTATATATTAAAAATTCAAATGAAAATACTCTTGAAAAGATCGATGAAAATGATAATAGCTTATCTAACGTACATATAGATACAACATTACCTATGGAGGATGTACCTGCATATACTGAACAAAATAAAAAAATTAAAAAGTGAcatgaatatgagaattattaataatatgaggtttatttaaaacaaggagaaatacatggattttaaaatgaaaattgtgatcactaataaggagaaattatataaaaataataaatatatatatatatatatatatatatatataattaaatatatgttatttaaaattaaaattaatcttttttttgaaaatctttaatatgtataaaaatatatacattctttaatatataataatatatacatttataaattatttaaaaaatatatatttattttttttaaattttatttttttataagtattgattttaataagtaaataaataaataaataaataaatatatatatatatatatatatatatattataatatatttatcaagaataaaatattaattattttttttgttttcatatatatattattatttttttttatattttttttttttttgtatttaattaacttaatatatatattcaactctttttccgggtatgtaaaatatataaaaatatatatttatatataatatgaaataatataat

**5’ amplicon size: 1234bps**

**3’ amplicon size: 1174bps**

## **> PF3D7_1328300**

tgtacgtatatcacatatatatatatatatatatatatatatatatatatatattacaactatcatttcatacaataaataaaaataacgaagaaaaaaataaataatacatttttttttattatgaatatatttttttacatatatatttttatatttataaatataaaagattaaattatttatatttccatattacacgtttcttcaacattaatacataaaacaaataattacattatatatttatttatacgcatacatgttatatattcctcgttattattgtaaaataaaaaaagaagaaaaagaaaaaagaaaaagagaaagagaaagagaaagagaaaaaaaaaaaaaaaaaaaaaaaaaaaagtaaaaacggcataatatattaatatatattaataaagtacaggaaatatttttacccatatatataacaatatatatatatatatatatatatatatatataataaaagattacagttacatatttttaaacatattatatgtatgcaaaagcgtgatattttatggtttttgtaattttcatatacatttaagaataaaaaaaaatatatatatatggaaaaaaaaaaaaaaaaaaaaaaaaaaaaaaaagcacaaaataatatcaatattacataacaacaaaacatgaaagaaagaaaaaaaatggaaaatcaataaaaattatattaaatgttataaaaaaatttttatgtatttatataaatattatatataataaagtgtaatattaaaaatgtaattatagagtatataaatattgagaaaaagaagggaaattttataattattttatgtataactatatatatatatatatatatatatatatatatatatatatttatttatttttttatttattttttacgatacattaataatatttgacaagtaaaataaataaataaataaatgtatacctatataatatatatatatatatatataagttatttattttgattcagcatattttataattttatgtatagaaatttaataaggccttattaaattccttgttaaagtttttcacctttattttatatttgttaataatcaaaaatatatatataaaataattatatattagtttattataacgtgtaccattaattagtaatatatatatatttatatattttatttatttatttacacagtatagggacatttcccttataatttgacatatccttttatcccattttatcttatattttgtattttttttttttttttttttgaaaaagaaaaaaATGTCGCTAAAAAACAAAGTGCAAGTGAATGATTTAAATAAGACAATTTTTAAAGAAAATTACAAAATGAAAAGGAGTACCAAAGAAGTAGAACTAATGGAAACTATAGAAAATTTGAGTGATGATATAGGATCCAATAATTTTTCTTATGATAGTAATGACAGCTCAAAAATACATAATAGGAAAAATAATGATAAAAGTATTGATATGAACAAAAAAGATAATATAAAAAATAATAATAAACAAAAAAATGCTCCTAATTATAATAAAAACAAACAAGAGTATTATTCCAATGTACCTCAAATATAT**GAAATATATGATACTACACCAAAAGATGAAGGTGAATTAAGGACCTCGGATAACAATGGGGGGGAAGAAAATAGTTTTCTTGCTAATACTATAGCTGTTGTGTATATTATTATAGCAATAG*gtaacataaaatataaggactcatatatatatatatatatatatatatatatatatttatatttatttatataatcttttcatatatatatcatataatttttttttttttttttttttttgcttattccccattgcag*TTATTATCTTGCACTTGTATTTAAACGAAGAGCATATCGTAATAAAAACCATTACTTTATTAAAATTAAAATTTGTTGATTTAATAAAGAGCATACCAGTATTCAAAAATGTTATGCGAACTGTGTCTATAACA*gtaaggatttgaaataaataaataaataaataaataaatatatatatatatatatatatatatatatatataaatatatatgtatatgcgtgaattttattttattttattttattatttttttttttttaccccccccgcag*AATAGCAAAATTAACGGTTTCCTAGATATGACGATGACTCATTCAAAAATTTTGAAACCCTATATGGAAGCTTTAAAAGAACTTAATACCGAATTTTCCATATTATTTATCATATTAATATTAATGTTTTTAACGGCGACATGCATACTTCATGTAGTTTATGGTATCCTTACGATGAAGAATGAAAAAGACATGTACGAAAAGAATAGTGTAGTTTATGTAACGAATAAAATGACAGTAGACGAATATGAAGATAGATCTTTTACTTACAGTGAACTTGCTAAATTACATGATGATAAAGATTATATTTGTTTAAAAAATAAAAGAGCAGGTGAAGGTATGGAATCATGGAATTGGCAAATAAGAAAAAATAAATATAATACATATGATAAAGATATTTGCAGTGATATTGAACTTACAGATATAGATAAT**TAAtaaataatataaacaaatatttccttataaaaataaaataaaatatatatatatatatatatatatttttattaattattatataattatcgaggatgtgttatataattatccatcattatttttttttttattgtttctttcttttttttctaaatctatatatatatataaaatattttacacattacacaaatgtagttatttgttaaatattataaattaagatttaatatttagtacatattttataatctctatttttttttttttaaatattgttctttttttaaaattaacaaaatgcattatttttgtatgtattaagaaagaagtaaatcatttatataatcattttttttaatatatatttataaatgcatactgtgtaatcatttttaaattacaacatggaaataaaatattatatatatatatatatatatatttacttaatgtatttgtgtatattttttttcctatttactaaatatatatataatatatatatattatataataatacctttgatagcattgtgaatataaaattatacaaatttttattttttgcataatttaatttatgtcatattatattttatatttaatcatttatatatatatatttttttttttttatgttttttttataaactattaattcttgagattaccaaatgtgaagttttataacatatatatgtttatgtagtactatatatcc

**Original Locus (OL) amplicon size: 1244bp**s

**> PF3D7_1328300-HALO-SW Halo Neo-R**

tgtacgtatatcacatatatatatatatatatatatatatatatatatatatattacaactatcatttcatacaataaataaaaataacgaagaaaaaaataaataatacatttttttttattatgaatatatttttttacatatatatttttatatttataaatataaaagattaaattatttatatttccatattacacgtttcttcaacattaatacataaaacaaataattacattatatatttatttatacgcatacatgttatatattcctcgttattattgtaaaataaaaaaagaagaaaaagaaaaaagaaaaagagaaagagaaagagaaagagaaaaaaaaaaaaaaaaaaaaaaaaaaaagtaaaaacggcataatatattaatatatattaataaagtacaggaaatatttttacccatatatataacaatatatatatatatatatatatatatatatataataaaagattacagttacatatttttaaacatattatatgtatgcaaaagcgtgatattttatggtttttgtaattttcatatacatttaagaataaaaaaaaatatatatatatggaaaaaaaaaaaaaaaaaaaaaaaaaaaaaaaagcacaaaataatatcaatattacataacaacaaaacatgaaagaaagaaaaaaaatggaaaatcaataaaaattatattaaatgttataaaaaaatttttatgtatttatataaatattatatataataaagtgtaatattaaaaatgtaattatagagtatataaatattgagaaaaagaagggaaattttataattattttatgtataactatatatatatatatatatatatatatatatatatatatatttatttatttttttatttattttttacgatacattaataatatttgacaagtaaaataaataaataaataaatgtatacctatataatatatatatatatatatataagttatttattttgattcagcatattttataattttatgtatagaaatttaataaggccttattaaattccttgttaaagtttttcacctttattttatatttgttaataatcaaaaatatatatataaaataattatatattagtttattataacgtgtaccattaattagtaatatatatatatttatatattttatttatttatttacacagtatagggacatttcccttataatttgacatatccttttatcccattttatcttatattttgtattttttttttttttttttttgaaaaagaaaaaaATGTCGCTAAAAAACAAAGTGCAAGTGAATGATTTAAATAAGACAATTTTTAAAGAAAATTACAAAATGAAAAGGAGTACCAAAGAAGTAGAACTAATGGAAACTATAGAAAATTTGAGTGATGATATAGGATCCAATAATTTTTCTTATGATAGTAATGACAGCTCAAAAATACATAATAGGAAAAATAATGATAAAAGTATTGATATGAACAAAAAAGATAATATAAAAAATAATAATAAACAAAAAAATGCTCCTAATTATAATAAAAACAAACAAGAGTATTATTCCAATGTACCTCAAATATAT**GAAATATATGATACTACACCAAAAGATGAAGGTGAATTAAGGACCTCGGATAACAATGGGGGGGAAGAAAATAGTTTTCTTGCTAATACTATAGCTGTTGTGTATATTATTATAGCAATAG*gtaacataaaatataaggactcatatatatatatatatatatatatatatatatatttatatttatttatataatcttttcatatatatatcatataatttttttttttttttttttttttgcttattccccattgcag*TTATTATCTTGCACTTGTATTTAAACGAAGAGCATATCGTAATAAAAACCATTACTTTATTAAAATTAAAATTTGTTGATTTAATAAAGAGCATACCAGTATTCAAAAATGTTATGCGAACTGTGTCTATAACA*gtaaggatttgaaataaataaataaataaataaataaatatatatatatatatatatatatatatatatataaatatatatgtatatgcgtgaattttattttattttattttattatttttttttttttaccccccccgcag*AATAGCAAAATTAACGGTTTCCTAGATATGACGATGACTCATTCAAAAATTTTGAAACCCTATATGGAAGCTTTAAAAGAACTTAATACCGAATTTTCCATATTATTTATCATATTAATATTAATGTTTTTAACGGCGACATGCATACTTCATGTAGTTTATGGTATCCTTACGATGAAGAATGAAAAAGACATGTACGAAAAGAATAGTGTAGTTTATGTAACGAATAAAATGACAGTAGACGAATATGAAGATAGATCTTTTACTTACAGTGAACTTGCTAAATTACATGATGATAAAGATTATATTTGTTTAAAAAATAAAAGAGCAGGTGAAGGTATGGAATCATGGAATTGGCAAATAAGAAAAAATAAATATAATACATATGATAAAGATATTTGCAGTGATATTGAACTTACAGATATAGATAAT**CCTAGGTCAGGATTGAGATCAAGATCTGCTGCTGCTGGTGCTGGTGGTGCTGCTAGAGCTGCTCTGCAGAGAGGAGTACAAGTTGAAACAATATCACCAGGAGATGGTCGTACATTTCCAAAAAGAGGTCAAACTTGTGTTGTACATTATACTGGAATGCTTGAAGATGGAAAGAAATTTGATTCATCTCGTGATAGAAATAAACCATTTAAATTTATGCTAGGTAAACAAGAAGTAATACGAGGTTGGGAAGAAGGAGTTGCTCAAATGAGTGTAGGTCAAAGAGCAAAACTTACTATATCTCCAGATTATGCTTATGGTGCAACTGGACATCCAGGTATAATTCCACCTCATGCAACTCTTGTATTTGATGTGGAGCTTCTAAAACTAGAAACTAGAGGTGTTCAGGTTGAAACAATTTCACCTGGAGATGGCAGAACCTTTCCTAAAAGAGGACAGACTTGCGTAGTTCATTATACAGGCATGCTAGAGGATGGTAAGAAATTTGATTCTAGTCGAGATAGAAATAAGCCATTCAAGTTTATGCTAGGTAAACAGGAAGTAATAAGAGGTTGGGAAGAGGGTGTAGCACAGATGTCAGTTGGACAAAGAGCAAAGTTAACAATATCACCAGATTATGCATACGGTGCAACAGGCCATCCTGGCATCATCCCTCCACATGCAACTTTAGTATTCGACGTTGAATTGTTAAAGTTAGAGACAACGCGTGCTAGAGGTGCTGCTGCTGGTGCTGGAGGTGCAGGTAGACGTACGATGGCAGAAATTGGTACGGGTTTTCCATTTGATCCTCATTATGTGGAGGTGCTGGGGGAAAGGATGCATTATGTTGACGTAGGACCAAGAGATGGTACTCCAGTGTTATTTTTGCATGGAAACCCAACCTCGAGTTATGTATGGAGAAATATAATTCCACATGTAGCACCAACACATAGATGTATAGCTCCTGATTTAATTGGTATGGGAAAAAGTGATAAACCTGACTTAGGATATTTTTTTGATGATCATGTCCGTTTTATGGATGCTTTCATTGAAGCTCTTGGCCTTGAAGAAGTAGTATTAGTTATACATGATTGGGGATCCGCCTTAGGATTTCATTGGGCCAAGAGGAATCCTGAAAGAGTAAAAGGAATAGCATTCATGGAATTCATACGACCAATCCCCACATGGGATGAATGGCCAGAATTTGCACGCGAAACATTTCAAGCTTTTAGAACTACAGATGTTGGTAGAAAATTAATAATAGATCAAAATGTATTTATAGAAGGAACTTTACCTATGGGTGTTGTAAGGCCGTTAACAGAAGTTGAAATGGACCACTACCGTGAACCTTTTTTAAATCCAGTAGATAGAGAGCCCTTATGGAGATTTCCTAATGAATTACCTATTGCAGGTGAACCCGCGAATATTGTTGCTTTAGTAGAAGAATATATGGATTGGTTACATCAGTCTCCTGTTCCTAAACTTCTATTTTGGGGTACACCTGGAGTTCTAATACCACCAGCTGAAGCAGCAAGATTAGCAAAATCATTACCAAATTGTAAAGCTGTTGATATAGGTCCTGGGTTGAATTTATTACAAGAAGATAATCCAGATTTGATTGGATCTGAGATAGCTAGATGGCTAAGTACATTAGAAATTTCAGGTACCGGTGCCAGGGGAGCAGCCGCAGGAGCAGGGGGGGCAGGAAGGCGTGGTGTTCAGGTCGAGACTATTAGCCCTGGAGATGGACGCACGTTTCCTAAGCGTGGACAGACATGCGTAGTTCACTACACAGGTATGTTGGAGGACGGTAAAAAGTTCGACAGCTCACGCGACCGCAATAAACCTTTCAAGTTTATGCTTGGCAAGCAGGAGGTTATTCGTGGATGGGAGGAGGGTGTAGCACAGATGTCTGTTGGACAGCGTGCTAAGTTGACAATTTCACCTGACTATGCTTATGGCGCTACGGGCCATCCCGGGATCATTCCGCCACATGCGACTCTGGTATTCGACGTTGAATTATTAAAGTTAGAGACAGCTAGAGGGGCCGCTGCAGGTGCTGGTGGAGCTGGAAGACGTGGAGTACAAGTAGAGACTATCTCTCCAGGTGACGGTCGCACTTTCCCAAAGCGTGGCCAAACCTGTGTTGTACATTACACTGGTATGCTGGAGGATGGGAAAAAGTTCGATTCCAGTCGCGACCGTAACAAACCGTTCAAATTCATGTTGGGAAAGCAGGAAGTGATCCGCGGGTGGGAGGAAGGCGTGGCGCAAATGAGCGTCGGTCAGCGGGCTAAATTGACCATTTCCCCTGACTACGCGTATGGGGCTACTGGGCACCCAGGGATTATTCCGCCTCACGCTACACTTGTGTTTGATGTCGAACTTTTGAAACTGGAAACTGTCGACGGAGAAGGAAGAGGAAGTTTATTAACATGTGGAGATGTAGAAGAAAATCCAGGACCAATGATTGAACAAGATGGATTGCACGCAGGTTCTCCGGCCGCTTGGGTGGAGAGGCTATTCGGCTATGACTGGGCACAACAGACAATCGGCTGCTCTGATGCCGCCGTGTTCCGGCTGTCAGCGCAGGGGCGCCCGGTTCTTTTTGTCAAGACCGACCTGTCCGGTGCCCTGAATGAACTGCAGGACGAGGCAGCGCGGCTATCGTGGCTGGCCACGACGGGCGTTCCTTGCGCAGCTGTGCTCGACGTTGTCACTGAAGCGGGAAGGGACTGGCTGCTATTGGGCGAAGTGCCGGGGCAGGATCTCCTGTCATCTCACCTTGCTCCTGCCGAGAAAGTATCCATCATGGCTGATGCAATGCGGCGGCTGCATACGCTTGATCCGGCTACCTGCCCATTCGACCACCAAGCGAAACATCGCATCGAGCGAGCACGTACTCGGATGGAAGCCGGTCTTGTCGATCAGGATGATCTGGACGAAGAGCATCAGGGGCTCGCGCCAGCCGAACTGTTCGCCAGGCTCAAGGCGCGCATGCCCGACGGCGAGGATCTCGTCGTGACCCATGGCGATGCCTGCTTGCCGAATATCATGGTGGAAAATGGCCGCTTTTCTGGATTCATCGACTGTGGCCGGCTGGGTGTGGCGGACCGCTATCAGGACATAGCGTTGGCTACCCGTGATATTGCTGAAGAGCTTGGCGGCGAATGGGCTGACCGCTTCCTCGTGCTTTACGGTATCGCCGCTCCCGATTCGCAGCGCATCGCCTTCTATCGCCTTCTTGACGAGTTCTTCTAActcgagggatatggcagcttaatgttcgtttttcttatttatatatttataccaattgattgtatttataactgtaaaaatgtgtatgttgtgtgcatatttttttttgtgcatgcacatgcatgtaaatagctaaaattatgaacattttattttttgttcagaaaaaaaaaactttacacacataaaatggctagtatgaatagccatattttatataaattaaatcctatgaatttatgaccatattaaaaatttagatatttatggaacataatatgtttgaaacaataagacaaaattattattattattattatttttactgttataattatgtgtctccttcaatgattcataaatagttggacttgatttttaaaatgtttataatatgattagcatagttaaataaaaaaagttgaaaaattaaaaaaaaacatataaacacaaatgatggtttttccttcaatttcgatatcaatttatagaaacaaaatatatacttgtataattttatttttttatataaatcattacatatataattatacaatattttttctaagagataattatatattaatatatataaaaaaaggtgttttttttttttttttttatttttatttttattttatggtaatattttattttccttattttataaattatattagtttatatgtgattaattttatatattatcaatttatatatttttaaatgcttacttaattatctttttttttttttttttttttttttcccctctttttatattaatttatttttgaaaaaattgatatatatatatatatataatatatatatatacatgtagtagtattaaacaatgtataatatatataaataatatatttatatatttcatttcaattttaattttttttggttttttttttttttctttttgtcatatttaaaaaaaattatattcatataagttatgcattttttataaacattattcaatatatgtataatataatatatatatatatattaatgtattattccaatgtgcatgataaaagaaaaaaataatatttataaaaaaaaagaaaaataaaacaaaaaaagaaaaaaaaaaaaaaaaaaaaaaaaatacaaaaataaataatataatttataattatatattcttgtcacaataaaaatatatatatatatatatatatttataatatgtatattttaaactagaaaaggaataactaatattttatttattatcattcaagatttatattttataataataaatacctaatagaaatatatcaggatccatgcatggttcgctaaactgcatcgtcgctgtgtcccagaacatgggcatcggcaagaacggggactacccctggccaccgctcaggaacgaatttagatatttccagagaatgaccacaacctcttcagtagaaggtaaacagaatctggtgattatgggtaagaagacctggttctccattcctgagaagaatcgacctttaaagggtagaattaatttagttctcagcagagaactcaaggaacctccacaaggagctcattttctttccagaagtctagatgatgccttaaaacttactgaacaaccagaattagcaaataaagtagacatggtctggatagttggtggcagttctgtttataaggaagccatgaatcacccaggccatcttaaactatttgtgacaaggatcatgcaagactttgaaagtgacacgttttttccagaaattgatttggagaaatataaacttctgccagaatacccaggtgttctctctgatgtccaggaggagaaaggcattaagtacaaatttgaagtatatgagaagaatgattaagcttatttaataatagattaaaaatattataaaaataaaaacataaacacagaaattacaaaaaaaatacatatgaattttttttttgtaatcttccttataaatatagaataatgaatcatataaaacatatcattattcatttatttacatttaaaattattgtttcagtatctttaatttattatgtatatataaaaataacttacaattttattaataaacaatatatgtttattaattcatgttttgtaatttatgggatagcgattttttttactgtctgtatttttcttttttaattatgttttaattgtattttatttttattattgttctttttatagtattattttaaaacaaaatgtattttctaagaacttataataataataaatataaattttaataaaaattatatttatcttttacaatatgaacataaagtacaacattaatatatagcttttaatatttttattcctaatcatgtaaatcttaaatttttctttttaaacatatgttaaatatttatttctcattatatataagaacatatttattaaatctagaattctatagtgagtcgtattacaattcactggccgtcgttttacaacgtcgtgactgggaaaaccctggcgttacccaacttaatcgccttgcagcacatccccctttcgccagctggcgtaatagcgaagaggcccgcaccgatcgcccttcccaacagttgcgcagcctgaatggcgaatggcgcctgatgcggtattttctccttacgcatctgtgcggtatttcacaccgcatatggtgcactctcagtacaatctgctctgatgccgcatagttaagccagccccgacacccgccaacacccgctgacgcgccctgacgggcttgtctgctcccggcatccgcttacagacaagctgtgaccgtctccgggagctgcatgtgtcagaggttttcaccgtcatcaccgaaacgcgcgagacgaaagggcctcgtgatacgcctatttttataggttaatgtcatgataataatggtttcttagacgtcaggtggcacttttcggggaaatgtgcgcggaacccctatttgtttatttttctaaatacattcaaatatgtatccgctcatgagacaataaccctgataaatgcttcaataatattgaaaaaggaagagtatgagtattcaacatttccgtgtcgcccttattcccttttttgcggcattttgccttcctgtttttgctcacccagaaacgctggtgaaagtaaaagatgctgaagatcagttgggtgcacgagtgggttacatcgaactggatctcaacagcggtaagatccttgagagttttcgccccgaagaacgttttccaatgatgagcacttttaaagttctgctatgtggcgcggtattatcccgtattgacgccgggcaagagcaactcggtcgccgcatacactattctcagaatgacttggttgagtactcaccagtcacagaaaagcatcttacggatggcatgacagtaagagaattatgcagtgctgccataaccatgagtgataacactgcggccaacttacttctgacaacgatcggaggaccgaaggagctaaccgcttttttgcacaacatgggggatcatgtaactcgccttgatcgttgggaaccggagctgaatgaagccataccaaacgacgagcgtgacaccacgatgcctgtagcaatgccaacaacgttgcgcaaactattaactggcgaactacttactctagcttcccggcaacaattaatagactggatggaggcggataaagttgcaggaccacttctgcgctcggcccttccggctggctggtttattgctgataaatctggagccggtgagcgtgggtctcgcggtatcattgcagcactggggccagatggtaagccctcccgtatcgtagttatctacacgacggggagtcaggcaactatggatgaacgaaatagacagatcgctgagataggtgcctcactgattaagcattggtaactgtcagaccaagtttactcatatatactttagattgatttaaaacttcatttttaatttaaaaggatctaggtgaagatcctttttgataatctcatgaccaaaatcccttaacgtgagttttcgttccactgagcgtcagaccccgtagaaaagatcaaaggatcttcttgagatcctttttttctgcgcgtaatctgctgcttgcaaacaaaaaaaccaccgctaccagcggtggtttgtttgccggatcaagagctaccaactctttttccgaaggtaactggcttcagcagagcgcagataccaaatactgtccttctagtgtagccgtagttaggccaccacttcaagaactctgtagcaccgcctacatacctcgctctgctaatcctgttaccagtggctgctgccagtggcgataagtcgtgtcttaccgggttggactcaagacgatagttaccggataaggcgcagcggtcgggctgaacggggggttcgtgcacacagcccagcttggagcgaacgacctacaccgaactgagatacctacagcgtgagctatgagaaagcgccacgcttcccgaagggagaaaggcggacaggtatccggtaagcggcagggtcggaacaggagagcgcacgagggagcttccagggggaaacgcctggtatctttatagtcctgtcgggtttcgccacctctgacttgagcgtcgatttttgtgatgctcgtcaggggggcggagcctatcgaaaaacgccagcaacgcggcctttttacggttcctggccttttgctggccttttgctcacatgttctttcctgcgttatcccctgattctgtggataaccgtattaccgcctttgagtgagctgataccgctcgccgcagccgaacgaccgagcgcagcgagtcagtgagcgaggaagcggaagagcgcccaatacgcaaaccgcctctccccgcgcgttggccgattcattaatgcagctggcacgacaggtttcccgactggaaagcgggcagtgagcgcaacgcaattaatgtgagttagctcactcattaggcaccccaggctttacactttatgcttccggctcgtatgttgtgtggaattgtgagcggataacaatttcacacaggaaacagctatgaccatgattacgccaagctatttaggtgacactatagaagcggccgcTAgGAAATATATGATACTACACCAAAAGATGAAGGTGAATTAAGGACCTCGGATAACAATGGGGGGGAAGAAAATAGTTTTCTTGCTAATACTATAGCTGTTGTGTATATTATTATAGCAATAGgtaacataaaatataaggactcatatatatatatatatatatatatatatatatatttatatttatttatataatcttttcatatatatatcatataatttttttttttttttttttttttgcttattccccattgcagTTATTATCTTGCACTTGTATTTAAACGAAGAGCATATCGTAATAAAAACCATTACTTTATTAAAATTAAAATTTGTTGATTTAATAAAGAGCATACCAGTATTCAAAAATGTTATGCGAACTGTGTCTATAACAgtaaggatttgaaataaataaataaataaataaataaatatatatatatatatatatatatatatatatataaatatatatgtatatgcgtgaattttattttattttattttattatttttttttttttaccccccccgcagAATAGCAAAATTAACGGTTTCCTAGATATGACGATGACTCATTCAAAAATTTTGAAACCCTATATGGAAGCTTTAAAAGAACTTAATACCGAATTTTCCATATTATTTATCATATTAATATTAATGTTTTTAACGGCGACATGCATACTTCATGTAGTTTATGGTATCCTTACGATGAAGAATGAAAAAGACATGTACGAAAAGAATAGTGTAGTTTATGTAACGAATAAAATGACAGTAGACGAATATGAAGATAGATCTTTTACTTACAGTGAACTTGCTAAATTACATGATGATAAAGATTATATTTGTTTAAAAAATAAAAGAGCAGGTGAAGGTATGGAATCATGGAATTGGCAAATAAGAAAAAATAAATATAATACATATGATAAAGATATTTGCAGTGATATTGAACTTACAGATATAGATAATTAAtaaataatataaacaaatatttccttataaaaataaaataaaatatatatatatatatatatatatttttattaattattatataattatcgaggatgtgttatataattatccatcattatttttttttttattgtttctttcttttttttctaaatctatatatatatataaaatattttacacattacacaaatgtagttatttgttaaatattataaattaagatttaatatttagtacatattttataatctctatttttttttttttaaatattgttctttttttaaaattaacaaaatgcattatttttgtatgtattaagaaagaagtaaatcatttatataatcattttttttaatatatatttataaatgcatactgtgtaatcatttttaaattacaacatggaaataaaatattatatatatatatatatatatatttacttaatgtatttgtgtatattttttttcctatttactaaatatatatataatatatatatattatataataatacctttgatagcattgtgaatataaaattatacaaatttttattttttgcataatttaatttatgtcatattatattttatatttaatcatttatatatatatatttttttttttttatgttttttttataaactattaattcttgagattaccaaatgtgaagttttataacatatatatgtttatgtagtactatatatcc

**5’ amplicon size: 1260bps**

**3’ amplicon size: 1178bps**

## **> PF3D7_1311600**

tatatatatatatatatatataatattatatatgttatttataaattaataataaataaataaataaatatatatatatatatatatatatatttatattataataggaattttttttttttcaatgtaattttaaaacttttaacttttttttttcttttatatatatatatatattttttttttttgatttttcaattttcattattcctatccgtatatttaaaaaaatatatatattttttacatatttataatttcttcattttttcctataaacaagaaaaagaacgaaaaaacaaaacacaacatatttatatatatatatatatgtttaatattaatgtccttataaacatttaaatttatttttgtaattgttaaatgttataatttaaatgtataaaaagacatatacatatatatatatatttataacataataatttaatcattattttaaaattattttgattataattaaaatgctatatcataatattaattctatgtataccaaagaaaataaataaacatacatacataaataaataaatataaataaatatatatataaataaatatatatatatatatatatatatatatatatatatatatatatatatgtatatatttagataagtttataagtacgATGACAGGAAGGATTAATAATCATGAGGAGTATATAAAGAATGAGAAAGAAATATATAACGTAAAAGAATCGGCAAATGAAAAGGAGAGTGATAATATGAAGAATGATGAATTTGTAATTTATAATGACAATTTGAAAGATGTAGAAGATATAAATAAGGAAAATGTAAATTATATAAAAAGCATAAATAAAATAGAATGTGATACAAATAATCACAATAATAATAATAATAATTATTATATTAATAATGATAATAAATTTTGTAATGATCATAAAAGTTTCGCTTCTTCATATAATTCTCAAAATTCAGACGTTGTAGGTACTTGTGAAGAAGGAAGCAATAAGTGTTATGAACCATCTAATAACATACATAATAATTTTATATTAAATGAAGATTATAAGAATAACACAAATGAAGAATATGATAAAATCGAAAGTGCACAAAATAGTGTATATTGCGAACAGGAAATTAATAACAATATAAATGAATCATATAATAATATAAATAATGAATGTACACAGAATTATATAAATAACACCGAAGAAATAACTAATCATATAATACTACAGGAAGAAAAAGAAATGGGAAATATAGAAAAAGAAGAACAAGATAATGATATGTATAAAGAACAAAAAATAAAAAGAGAAGATACAGAAAATAAAAATGAAATGGATTATATAAAGGATGAATCAAATAATATTTTTCCAAGCGATCAGATTAATTCTGTAACAAGTGACCAAGAAATTTATATTACATCAAACGATAAAAAGAATAATTTAACAAATGAAAAAAAAGTTTATATTTCAACAAACGATAAAAATAATGCTGTAAGAAAT**GATAAAACATTTTACCTCTCATCAAATGATAACCAAAGTGATTCCTTACCAAAGAAATCGATACGAAATAATGTTGAAGAATTTATAAGAAAACTTATGGAAAATGAAGATATGGCAAATAAATTTAATGTGTTTCATATAGGTATAAATAAAATCGAAGCGCTTAAAGCTTTATATAGAGAATGTAAGAATAATTTTAATAAAAGTTTTAGAGAAAAAATTAATAAATTACATGAATTAAGAAAATATTATAATGATAGTATATTTGAAAGGAATATAAGTGTTGATTATATTGCCAATACAGAAAAATTGAATGCTATAAAAAAAGAAAGATTGTCAAGATTATTTTTATATATTGAAGATATACATAAGTTAAAACAAGCCCTAAAAGTAAAACAACATAATGATATAGAATTAGAAAATGATGAAAAAAAAGAATATTTTTATGATGCTTATGAAGATATAGAAGATTTCTTTATATTCGATCATGCACTCTCATATCTAAATAATGATGTTGAAACAATTTTTACAGAATGTGAAAATTCTTCTTTTTTTAGAGAAGATTCAAGTATTGATAATTTCTTAGATGAAGTAAGTAGTAAAGCCGATGAAGAATATAATAGATTAAATATATTTGCAAGATTTTCTTTAAAAAATTATTATTTATTCGATATAAATAAAAACTTATGTTCTATTTATGCATTCACACCTATGATTATTAAAAAATTTGCAGAAAAAATTAATTCCTATAATTTAACAAAAAATGAAAAAGACATAAATAATGATAATGATGCTGAACATGAAAAAAATGATATAGATTTTACTATAGATTCAACCAATAATGAAAAACGTAATATATATGCTAGCTGTGGACTTGAAAGTAATCATAATTTAGATGCATACGTACAATTTCATTGTGATAAGTGGCAAGAAAATATGTATGATGTTGTTGAGGTTAATAGTCTCGAATTAGATTTTAATAGATTAAAATGTAATATTATG**TAAttcgatttattttatatacttacctgtcttaatatcacatattataaattaagatatattattataatataataaaatataatatatctaaagttaattattatttatttatttatttattaatttacttatttatttatttatttattaatttacttatttatttatttatttatttatttatttttattgttatcttttttttttttgtgcttaatttttaaataatataaagaatttaaaatattcacatttttgaatgtatcaatttatattttgtgttttcttgtttttgctctttaagaaaaaatatatatataaattaaaaggaaaatatgtataatatatatatatttatatatacaatacaaaaaaattatatatttgtatacatatcatttttacaaaagatgtgtgcatatatattcatccgtgttacctattttgtttgtttttttttttttttaatggttcattttatatttgaatatataaattatataatatacatatataatgtattcatttatttttcataaaaaatgtgcataaactcattttattgttatgtattattactgttgtgattataataataattattattattatggttattctaattaattattttgtattttgttcttttattctttatatttttcatattataatgacatttataaagacttttaaatatccaaatataaaataaaaaaacaatacttttttgttcaatagaataataaaattttcataaaatggaatgtgacaactatatttgttctacataacacataaaattaaaaacaaacttttttttaacaatatagtaattgtttaattacaaattagtaggttaaatatgtgtgtacaaatattaattagagctccttattaataaaaggggatcaatgaataaaagaaaaaaaaagaaaagtaaaaaaaaaaaaaaaaaagaaacaaaaataaatgtatatatataataattaacaaaataaaaaacgtatttttttaattttcatttaaataaaaaaataaaagaaaaatattataattatacgaaatgtgtataatatatatagcgaaaacattttcaaaatttgcaaaaccattatatga

**Original Locus (OL) amplicon size: 1169bp**s

**> PF3D7_1311600-HALO-SW Halo Neo-R**

atatatatatatatatatttatattataataggaattttttttttttcaatgtaattttaaaacttttaacttttttttttcttttatatatatatatatattttttttttttgatttttcaattttcattattcctatccgtatatttaaaaaaatatatatattttttacatatttataatttcttcattttttcctataaacaagaaaaagaacgaaaaaacaaaacacaacatatttatatatatatatatatgtttaatattaatgtccttataaacatttaaatttatttttgtaattgttaaatgttataatttaaatgtataaaaagacatatacatatatatatatatttataacataataatttaatcattattttaaaattattttgattataattaaaatgctatatcataatattaattctatgtataccaaagaaaataaataaacatacatacataaataaataaatataaataaatatatatataaataaatatatatatatatatatatatatatatatatatatatatatatatatgtatatatttagataagtttataagtacgATGACAGGAAGGATTAATAATCATGAGGAGTATATAAAGAATGAGAAAGAAATATATAACGTAAAAGAATCGGCAAATGAAAAGGAGAGTGATAATATGAAGAATGATGAATTTGTAATTTATAATGACAATTTGAAAGATGTAGAAGATATAAATAAGGAAAATGTAAATTATATAAAAAGCATAAATAAAATAGAATGTGATACAAATAATCACAATAATAATAATAATAATTATTATATTAATAATGATAATAAATTTTGTAATGATCATAAAAGTTTCGCTTCTTCATATAATTCTCAAAATTCAGACGTTGTAGGTACTTGTGAAGAAGGAAGCAATAAGTGTTATGAACCATCTAATAACATACATAATAATTTTATATTAAATGAAGATTATAAGAATAACACAAATGAAGAATATGATAAAATCGAAAGTGCACAAAATAGTGTATATTGCGAACAGGAAATTAATAACAATATAAATGAATCATATAATAATATAAATAATGAATGTACACAGAATTATATAAATAACACCGAAGAAATAACTAATCATATAATACTACAGGAAGAAAAAGAAATGGGAAATATAGAAAAAGAAGAACAAGATAATGATATGTATAAAGAACAAAAAATAAAAAGAGAAGATACAGAAAATAAAAATGAAATGGATTATATAAAGGATGAATCAAATAATATTTTTCCAAGCGATCAGATTAATTCTGTAACAAGTGACCAAGAAATTTATATTACATCAAACGATAAAAAGAATAATTTAACAAATGAAAAAAAAGTTTATATTTCAACAAACGATAAAAATAATGCTGTAAGAAAT**GATAAAACATTTTACCTCTCATCAAATGATAACCAAAGTGATTCCTTACCAAAGAAATCGATACGAAATAATGTTGAAGAATTTATAAGAAAACTTATGGAAAATGAAGATATGGCAAATAAATTTAATGTGTTTCATATAGGTATAAATAAAATCGAAGCGCTTAAAGCTTTATATAGAGAATGTAAGAATAATTTTAATAAAAGTTTTAGAGAAAAAATTAATAAATTACATGAATTAAGAAAATATTATAATGATAGTATATTTGAAAGGAATATAAGTGTTGATTATATTGCCAATACAGAAAAATTGAATGCTATAAAAAAAGAAAGATTGTCAAGATTATTTTTATATATTGAAGATATACATAAGTTAAAACAAGCCCTAAAAGTAAAACAACATAATGATATAGAATTAGAAAATGATGAAAAAAAAGAATATTTTTATGATGCTTATGAAGATATAGAAGATTTCTTTATATTCGATCATGCACTCTCATATCTAAATAATGATGTTGAAACAATTTTTACAGAATGTGAAAATTCTTCTTTTTTTAGAGAAGATTCAAGTATTGATAATTTCTTAGATGAAGTAAGTAGTAAAGCCGATGAAGAATATAATAGATTAAATATATTTGCAAGATTTTCTTTAAAAAATTATTATTTATTCGATATAAATAAAAACTTATGTTCTATTTATGCATTCACACCTATGATTATTAAAAAATTTGCAGAAAAAATTAATTCCTATAATTTAACAAAAAATGAAAAAGACATAAATAATGATAATGATGCTGAACATGAAAAAAATGATATAGATTTTACTATAGATTCAACCAATAATGAAAAACGTAATATATATGCTAGCTGTGGACTTGAAAGTAATCATAATTTAGATGCATACGTACAATTTCATTGTGATAAGTGGCAAGAAAATATGTATGATGTTGTTGAGGTTAATAGTCTCGAATTAGATTTTAATAGATTAAAATGTAATATTATG**CCTAGGTCAGGATTGAGATCAAGATCTGCTGCTGCTGGTGCTGGTGGTGCTGCTAGAGCTGCTCTGCAGAGAGGAGTACAAGTTGAAACAATATCACCAGGAGATGGTCGTACATTTCCAAAAAGAGGTCAAACTTGTGTTGTACATTATACTGGAATGCTTGAAGATGGAAAGAAATTTGATTCATCTCGTGATAGAAATAAACCATTTAAATTTATGCTAGGTAAACAAGAAGTAATACGAGGTTGGGAAGAAGGAGTTGCTCAAATGAGTGTAGGTCAAAGAGCAAAACTTACTATATCTCCAGATTATGCTTATGGTGCAACTGGACATCCAGGTATAATTCCACCTCATGCAACTCTTGTATTTGATGTGGAGCTTCTAAAACTAGAAACTAGAGGTGTTCAGGTTGAAACAATTTCACCTGGAGATGGCAGAACCTTTCCTAAAAGAGGACAGACTTGCGTAGTTCATTATACAGGCATGCTAGAGGATGGTAAGAAATTTGATTCTAGTCGAGATAGAAATAAGCCATTCAAGTTTATGCTAGGTAAACAGGAAGTAATAAGAGGTTGGGAAGAGGGTGTAGCACAGATGTCAGTTGGACAAAGAGCAAAGTTAACAATATCACCAGATTATGCATACGGTGCAACAGGCCATCCTGGCATCATCCCTCCACATGCAACTTTAGTATTCGACGTTGAATTGTTAAAGTTAGAGACAACGCGTGCTAGAGGTGCTGCTGCTGGTGCTGGAGGTGCAGGTAGACGTACGATGGCAGAAATTGGTACGGGTTTTCCATTTGATCCTCATTATGTGGAGGTGCTGGGGGAAAGGATGCATTATGTTGACGTAGGACCAAGAGATGGTACTCCAGTGTTATTTTTGCATGGAAACCCAACCTCGAGTTATGTATGGAGAAATATAATTCCACATGTAGCACCAACACATAGATGTATAGCTCCTGATTTAATTGGTATGGGAAAAAGTGATAAACCTGACTTAGGATATTTTTTTGATGATCATGTCCGTTTTATGGATGCTTTCATTGAAGCTCTTGGCCTTGAAGAAGTAGTATTAGTTATACATGATTGGGGATCCGCCTTAGGATTTCATTGGGCCAAGAGGAATCCTGAAAGAGTAAAAGGAATAGCATTCATGGAATTCATACGACCAATCCCCACATGGGATGAATGGCCAGAATTTGCACGCGAAACATTTCAAGCTTTTAGAACTACAGATGTTGGTAGAAAATTAATAATAGATCAAAATGTATTTATAGAAGGAACTTTACCTATGGGTGTTGTAAGGCCGTTAACAGAAGTTGAAATGGACCACTACCGTGAACCTTTTTTAAATCCAGTAGATAGAGAGCCCTTATGGAGATTTCCTAATGAATTACCTATTGCAGGTGAACCCGCGAATATTGTTGCTTTAGTAGAAGAATATATGGATTGGTTACATCAGTCTCCTGTTCCTAAACTTCTATTTTGGGGTACACCTGGAGTTCTAATACCACCAGCTGAAGCAGCAAGATTAGCAAAATCATTACCAAATTGTAAAGCTGTTGATATAGGTCCTGGGTTGAATTTATTACAAGAAGATAATCCAGATTTGATTGGATCTGAGATAGCTAGATGGCTAAGTACATTAGAAATTTCAGGTACCGGTGCCAGGGGAGCAGCCGCAGGAGCAGGGGGGGCAGGAAGGCGTGGTGTTCAGGTCGAGACTATTAGCCCTGGAGATGGACGCACGTTTCCTAAGCGTGGACAGACATGCGTAGTTCACTACACAGGTATGTTGGAGGACGGTAAAAAGTTCGACAGCTCACGCGACCGCAATAAACCTTTCAAGTTTATGCTTGGCAAGCAGGAGGTTATTCGTGGATGGGAGGAGGGTGTAGCACAGATGTCTGTTGGACAGCGTGCTAAGTTGACAATTTCACCTGACTATGCTTATGGCGCTACGGGCCATCCCGGGATCATTCCGCCACATGCGACTCTGGTATTCGACGTTGAATTATTAAAGTTAGAGACAGCTAGAGGGGCCGCTGCAGGTGCTGGTGGAGCTGGAAGACGTGGAGTACAAGTAGAGACTATCTCTCCAGGTGACGGTCGCACTTTCCCAAAGCGTGGCCAAACCTGTGTTGTACATTACACTGGTATGCTGGAGGATGGGAAAAAGTTCGATTCCAGTCGCGACCGTAACAAACCGTTCAAATTCATGTTGGGAAAGCAGGAAGTGATCCGCGGGTGGGAGGAAGGCGTGGCGCAAATGAGCGTCGGTCAGCGGGCTAAATTGACCATTTCCCCTGACTACGCGTATGGGGCTACTGGGCACCCAGGGATTATTCCGCCTCACGCTACACTTGTGTTTGATGTCGAACTTTTGAAACTGGAAACTGTCGACGGAGAAGGAAGAGGAAGTTTATTAACATGTGGAGATGTAGAAGAAAATCCAGGACCAATGATTGAACAAGATGGATTGCACGCAGGTTCTCCGGCCGCTTGGGTGGAGAGGCTATTCGGCTATGACTGGGCACAACAGACAATCGGCTGCTCTGATGCCGCCGTGTTCCGGCTGTCAGCGCAGGGGCGCCCGGTTCTTTTTGTCAAGACCGACCTGTCCGGTGCCCTGAATGAACTGCAGGACGAGGCAGCGCGGCTATCGTGGCTGGCCACGACGGGCGTTCCTTGCGCAGCTGTGCTCGACGTTGTCACTGAAGCGGGAAGGGACTGGCTGCTATTGGGCGAAGTGCCGGGGCAGGATCTCCTGTCATCTCACCTTGCTCCTGCCGAGAAAGTATCCATCATGGCTGATGCAATGCGGCGGCTGCATACGCTTGATCCGGCTACCTGCCCATTCGACCACCAAGCGAAACATCGCATCGAGCGAGCACGTACTCGGATGGAAGCCGGTCTTGTCGATCAGGATGATCTGGACGAAGAGCATCAGGGGCTCGCGCCAGCCGAACTGTTCGCCAGGCTCAAGGCGCGCATGCCCGACGGCGAGGATCTCGTCGTGACCCATGGCGATGCCTGCTTGCCGAATATCATGGTGGAAAATGGCCGCTTTTCTGGATTCATCGACTGTGGCCGGCTGGGTGTGGCGGACCGCTATCAGGACATAGCGTTGGCTACCCGTGATATTGCTGAAGAGCTTGGCGGCGAATGGGCTGACCGCTTCCTCGTGCTTTACGGTATCGCCGCTCCCGATTCGCAGCGCATCGCCTTCTATCGCCTTCTTGACGAGTTCTTCTAActcgagggatatggcagcttaatgttcgtttttcttatttatatatttataccaattgattgtatttataactgtaaaaatgtgtatgttgtgtgcatatttttttttgtgcatgcacatgcatgtaaatagctaaaattatgaacattttattttttgttcagaaaaaaaaaactttacacacataaaatggctagtatgaatagccatattttatataaattaaatcctatgaatttatgaccatattaaaaatttagatatttatggaacataatatgtttgaaacaataagacaaaattattattattattattatttttactgttataattatgtgtctccttcaatgattcataaatagttggacttgatttttaaaatgtttataatatgattagcatagttaaataaaaaaagttgaaaaattaaaaaaaaacatataaacacaaatgatggtttttccttcaatttcgatatcaatttatagaaacaaaatatatacttgtataattttatttttttatataaatcattacatatataattatacaatattttttctaagagataattatatattaatatatataaaaaaaggtgttttttttttttttttttatttttatttttattttatggtaatattttattttccttattttataaattatattagtttatatgtgattaattttatatattatcaatttatatatttttaaatgcttacttaattatctttttttttttttttttttttttttcccctctttttatattaatttatttttgaaaaaattgatatatatatatatatataatatatatatatacatgtagtagtattaaacaatgtataatatatataaataatatatttatatatttcatttcaattttaattttttttggttttttttttttttctttttgtcatatttaaaaaaaattatattcatataagttatgcattttttataaacattattcaatatatgtataatataatatatatatatatattaatgtattattccaatgtgcatgataaaagaaaaaaataatatttataaaaaaaaagaaaaataaaacaaaaaaagaaaaaaaaaaaaaaaaaaaaaaaaatacaaaaataaataatataatttataattatatattcttgtcacaataaaaatatatatatatatatatatatttataatatgtatattttaaactagaaaaggaataactaatattttatttattatcattcaagatttatattttataataataaatacctaatagaaatatatcaggatccatgcatggttcgctaaactgcatcgtcgctgtgtcccagaacatgggcatcggcaagaacggggactacccctggccaccgctcaggaacgaatttagatatttccagagaatgaccacaacctcttcagtagaaggtaaacagaatctggtgattatgggtaagaagacctggttctccattcctgagaagaatcgacctttaaagggtagaattaatttagttctcagcagagaactcaaggaacctccacaaggagctcattttctttccagaagtctagatgatgccttaaaacttactgaacaaccagaattagcaaataaagtagacatggtctggatagttggtggcagttctgtttataaggaagccatgaatcacccaggccatcttaaactatttgtgacaaggatcatgcaagactttgaaagtgacacgttttttccagaaattgatttggagaaatataaacttctgccagaatacccaggtgttctctctgatgtccaggaggagaaaggcattaagtacaaatttgaagtatatgagaagaatgattaagcttatttaataatagattaaaaatattataaaaataaaaacataaacacagaaattacaaaaaaaatacatatgaattttttttttgtaatcttccttataaatatagaataatgaatcatataaaacatatcattattcatttatttacatttaaaattattgtttcagtatctttaatttattatgtatatataaaaataacttacaattttattaataaacaatatatgtttattaattcatgttttgtaatttatgggatagcgattttttttactgtctgtatttttcttttttaattatgttttaattgtattttatttttattattgttctttttatagtattattttaaaacaaaatgtattttctaagaacttataataataataaatataaattttaataaaaattatatttatcttttacaatatgaacataaagtacaacattaatatatagcttttaatatttttattcctaatcatgtaaatcttaaatttttctttttaaacatatgttaaatatttatttctcattatatataagaacatatttattaaatctagaattctatagtgagtcgtattacaattcactggccgtcgttttacaacgtcgtgactgggaaaaccctggcgttacccaacttaatcgccttgcagcacatccccctttcgccagctggcgtaatagcgaagaggcccgcaccgatcgcccttcccaacagttgcgcagcctgaatggcgaatggcgcctgatgcggtattttctccttacgcatctgtgcggtatttcacaccgcatatggtgcactctcagtacaatctgctctgatgccgcatagttaagccagccccgacacccgccaacacccgctgacgcgccctgacgggcttgtctgctcccggcatccgcttacagacaagctgtgaccgtctccgggagctgcatgtgtcagaggttttcaccgtcatcaccgaaacgcgcgagacgaaagggcctcgtgatacgcctatttttataggttaatgtcatgataataatggtttcttagacgtcaggtggcacttttcggggaaatgtgcgcggaacccctatttgtttatttttctaaatacattcaaatatgtatccgctcatgagacaataaccctgataaatgcttcaataatattgaaaaaggaagagtatgagtattcaacatttccgtgtcgcccttattcccttttttgcggcattttgccttcctgtttttgctcacccagaaacgctggtgaaagtaaaagatgctgaagatcagttgggtgcacgagtgggttacatcgaactggatctcaacagcggtaagatccttgagagttttcgccccgaagaacgttttccaatgatgagcacttttaaagttctgctatgtggcgcggtattatcccgtattgacgccgggcaagagcaactcggtcgccgcatacactattctcagaatgacttggttgagtactcaccagtcacagaaaagcatcttacggatggcatgacagtaagagaattatgcagtgctgccataaccatgagtgataacactgcggccaacttacttctgacaacgatcggaggaccgaaggagctaaccgcttttttgcacaacatgggggatcatgtaactcgccttgatcgttgggaaccggagctgaatgaagccataccaaacgacgagcgtgacaccacgatgcctgtagcaatgccaacaacgttgcgcaaactattaactggcgaactacttactctagcttcccggcaacaattaatagactggatggaggcggataaagttgcaggaccacttctgcgctcggcccttccggctggctggtttattgctgataaatctggagccggtgagcgtgggtctcgcggtatcattgcagcactggggccagatggtaagccctcccgtatcgtagttatctacacgacggggagtcaggcaactatggatgaacgaaatagacagatcgctgagataggtgcctcactgattaagcattggtaactgtcagaccaagtttactcatatatactttagattgatttaaaacttcatttttaatttaaaaggatctaggtgaagatcctttttgataatctcatgaccaaaatcccttaacgtgagttttcgttccactgagcgtcagaccccgtagaaaagatcaaaggatcttcttgagatcctttttttctgcgcgtaatctgctgcttgcaaacaaaaaaaccaccgctaccagcggtggtttgtttgccggatcaagagctaccaactctttttccgaaggtaactggcttcagcagagcgcagataccaaatactgtccttctagtgtagccgtagttaggccaccacttcaagaactctgtagcaccgcctacatacctcgctctgctaatcctgttaccagtggctgctgccagtggcgataagtcgtgtcttaccgggttggactcaagacgatagttaccggataaggcgcagcggtcgggctgaacggggggttcgtgcacacagcccagcttggagcgaacgacctacaccgaactgagatacctacagcgtgagctatgagaaagcgccacgcttcccgaagggagaaaggcggacaggtatccggtaagcggcagggtcggaacaggagagcgcacgagggagcttccagggggaaacgcctggtatctttatagtcctgtcgggtttcgccacctctgacttgagcgtcgatttttgtgatgctcgtcaggggggcggagcctatcgaaaaacgccagcaacgcggcctttttacggttcctggccttttgctggccttttgctcacatgttctttcctgcgttatcccctgattctgtggataaccgtattaccgcctttgagtgagctgataccgctcgccgcagccgaacgaccgagcgcagcgagtcagtgagcgaggaagcggaagagcgcccaatacgcaaaccgcctctccccgcgcgttggccgattcattaatgcagctggcacgacaggtttcccgactggaaagcgggcagtgagcgcaacgcaattaatgtgagttagctcactcattaggcaccccaggctttacactttatgcttccggctcgtatgttgtgtggaattgtgagcggataacaatttcacacaggaaacagctatgaccatgattacgccaagctatttaggtgacactatagaagcggccgcTAgGATAAAACATTTTACCTCTCATCAAATGATAACCAAAGTGATTCCTTACCAAAGAAATCGATACGAAATAATGTTGAAGAATTTATAAGAAAACTTATGGAAAATGAAGATATGGCAAATAAATTTAATGTGTTTCATATAGGTATAAATAAAATCGAAGCGCTTAAAGCTTTATATAGAGAATGTAAGAATAATTTTAATAAAAGTTTTAGAGAAAAAATTAATAAATTACATGAATTAAGAAAATATTATAATGATAGTATATTTGAAAGGAATATAAGTGTTGATTATATTGCCAATACAGAAAAATTGAATGCTATAAAAAAAGAAAGATTGTCAAGATTATTTTTATATATTGAAGATATACATAAGTTAAAACAAGCCCTAAAAGTAAAACAACATAATGATATAGAATTAGAAAATGATGAAAAAAAAGAATATTTTTATGATGCTTATGAAGATATAGAAGATTTCTTTATATTCGATCATGCACTCTCATATCTAAATAATGATGTTGAAACAATTTTTACAGAATGTGAAAATTCTTCTTTTTTTAGAGAAGATTCAAGTATTGATAATTTCTTAGATGAAGTAAGTAGTAAAGCCGATGAAGAATATAATAGATTAAATATATTTGCAAGATTTTCTTTAAAAAATTATTATTTATTCGATATAAATAAAAACTTATGTTCTATTTATGCATTCACACCTATGATTATTAAAAAATTTGCAGAAAAAATTAATTCCTATAATTTAACAAAAAATGAAAAAGACATAAATAATGATAATGATGCTGAACATGAAAAAAATGATATAGATTTTACTATAGATTCAACCAATAATGAAAAACGTAATATATATGCTAGCTGTGGACTTGAAAGTAATCATAATTTAGATGCATACGTACAATTTCATTGTGATAAGTGGCAAGAAAATATGTATGATGTTGTTGAGGTTAATAGTCTCGAATTAGATTTTAATAGATTAAAATGTAATATTATGTAAttcgatttattttatatacttacctgtcttaatatcacatattataaattaagatatattattataatataataaaatataatatatctaaagttaattattatttatttatttatttattaatttacttatttatttatttatttattaatttacttatttatttatttatttatttatttatttttattgttatcttttttttttttgtgcttaatttttaaataatataaagaatttaaaatattcacatttttgaatgtatcaatttatattttgtgttttcttgtttttgctctttaagaaaaaatatatatataaattaaaaggaaaatatgtataatatatatatatttatatatacaatacaaaaaaattatatatttgtatacatatcatttttacaaaagatgtgtgcatatatattcatccgtgttacctattttgtttgtttttttttttttttaatggttcattttatatttgaatatataaattatataatatacatatataatgtattcatttatttttcataaaaaatgtgcataaactcattttattgttatgtattattactgttgtgattataataataattattattattatggttattctaattaattattttgtattttgttcttttattctttatatttttcatattataatgacatttataaagacttttaaatatccaaatataaaataaaaaaacaatacttttttgttcaatagaataataaaattttcataaaatggaatgtgacaactatatttgttctacataacacataaaattaaaaacaaacttttttttaacaatatagtaattgtttaat

**5’ amplicon size: 1257bps**

**3’ amplicon size: 1139bps**

### **> PF3D7_1351000 (PfPITP)**

ataagaatatgaaaaatatgcaagtgtttacattttatttgtaaaaaaaataacaatacc

ttacaaaagtgtgaataaataaataaataaataaataaatatatatatatatatatatat

atatatatatatgtaatgatcagtatgtgtaaggataaatatatttaagtatatttcttc

tttctatgttttttttaaattatacatagaataaatataaggaataatcatatatatata

tatatatatatatatgttatcatttcattcatatatttattttatgaattatatattttt

aataatatatagctctattttcttctttttgataattaaaagtatttataatttataaag

tttatttgtgtgcatattattacgtatataaataaatatatacatatatatatacatata

tatatatatctatatattgtacatatgtttgtatgtgtatcaattcatttagatatgttt

acatacacgcttaatataaaATG**AAAATTGTCGAATTTCGGCTTGCTATGCCTTTAACCA**

**TAGAAGAATATAAAGTATGTCAATTATATTTTGTAGCTAAAGCTTCATTAGAAGACGCAG**

**AAAATAATATAAATAGTAATTGTGAAAAAGGTGATGAGAATAATGATTCAAAAAAAGGAA**

**TAGTAATATTAAAAAATGAAAGTTATATAAATGAGGATGGTACATGTGGTCAGTATACAT**

**ATAAGAGAATAAATTTAATAAATAAGTTACCAAAGTGGCTATTAAATTTTATAGATCCTA**

**AATATTGTATTATAGATGAAAAGTCATGGAATGCATATCCATATTTAAAAACAGTTTATG**

**AGTCTAGTGGTTTTCCAAAAGCGAAGATTCAAGTTGAATCCGCACATTTTAATGGTTATG**

**ACACTGAAGAGAATGCTTTGAATTTATCAGAAGAAGCGCTCTCTTTAAGAAAAGTAATAT**

**ATGTTGATATAGTAAATGATAAAATATCTTATAAAGATTATAATGAAAGTGAAGATCCTT**

**CGTTATTTTATAGTGATAAAGCTAAAAGAGGTAAACTCGAAAAAAACTGGAAAGAAAATC**

**ATTCTATTATTATGACATGTTATAAGGTATTTACTATTAATATACCTTATTTTGGTATAT**

**TCTGTTCAAAATTAGAAAACTGGATTATATCTGCGCTCAGGGATAATATATTAAAATATC**

**ATAGGAAAGCATTCTGTTGGATTGATGAATGGATAGATTTGACAATTGATGATATAAGAA**

**ATTTAGAAAGAGATGTTCAAAAGAAATTGAATAAATTTTGGAACGACTCACAAGACGAAA**

**ATGTTACAGATGATATTTTGGATGAAGAAAATAACAATAAAAACGAAAATGTAAAAGATC**

**AATATGTTATACATCATAACAATAATGAAGAAAATAATGATACACTAAGTAATGATATAA**

**TAGATTCGTTAACTTTTAAAAATAACCAAAAAGATGATATACAAAGAAATATGTCTAATG**

**GT**AATAATATGATGAATAATGATAAAATATGTAATAATAATAATAATAATAATATTTATT

ATAATGGAAACATTCATAATAACGAATGTACTAATAAACAAAATTTATCTCAAGGTAATG

ATCAAGGAAGGAAAGAAATATCAAAAGACCTGCCCAATGGAACTTATATTACATCATCAC

TAAATATGGATAATGATAATTTAGATTTTATATGTACCAAATATAATGAGGCTTATGATG

ATAATGATAAAGAAAATAATAAAGACATGAAAAAAAAAAATATAAATATAAATATTAATG

ACATGCAAGGAGAAAAGCAAATCATTCAAAATGAAAATACACTAAATAATAAAATGATTA

ATGTATATGGTGATAATAATATCAATAATATTAATAATATTGCTAGAAATAAAAGTGTTA

TTACTAAAAGTACATCTCTATTATTCCCTAACGATTTAACCATAAAAAATAACCTCAATG

GAAATCATAATAATGTTATAGATAATATAGAAAAGAAGAAAAAAAAAAAAAAAAAAAAAA

AAGAAAAGAAAATGGAATCGGAAAATTACTTTTTCTTTAAGAAAAATGAAGAAAACAAAG

AATTTGGTGAATATTTATACAGACTAAATGAAGGCATGTTTTATTCATGGAAGCTTCGAT

ACTTTGTTATAAAAAATAACAAATTGTGTTATTATGTTAATCATAGTAAAAATGAGCTAA

AAGGAGAAATCGATCTCCTGAATGCGCAGATACAATGGATAGGTGAATATAAAGGAAGAA

ATAGTGTTTTTGTAATAAACTCCTTATCAAAGAATGTTAATTATTTAAGTTCAGAGAATG

AAATACAAACAAAGAAATGTATGATAGATATACAAATGGCTACTTTAATGAATAGTGAAA

GTGTGAAAAAGAGAGAGATTAAACAGGAAAATGGGAAAATTGTGAGTAAGGGGGTACAAA

ATGGAGATAACAAAGAAGATGTTGATGATGATAAAGAAGATGATGATAATGATGATGATG

AAGATGAAGATACTGAGGATGATAATAATGATGATGACAATGATGATGACAATGATGATG

ATAATGATGATGATAATGATGATGATAATGATGACGATAATGATGACGATAATGATAACA

ATAATGATGATGATAATGATGACGATAATGATGATGATAATGATAACAATAATGATGATG

ATAATTATGTGGATAACAACCATTACAACAACAATGATGATAACCATATTAATTATAATA

ATAAAGACACACCGAATTATCATACACATAAAAACAAAAGAATAGTTGATAACAAAAAAT

CACAAGATATGCCAAAAAATAAAAATGATGGAACTATAAATAATTATAACTTACCATATT

CAAAAGAATCAAATAAAGAGATCGATATAAATAATAATCATTATAATAATTATATATGTG

TAAATGATAAGGACGAAATTGTTCATAGTAAGTTAATTCTTAGCGATAATGATTTATCAC

AATATCATAATGAAGAGAATAAAACAATTTCAAATAACTCACATTATATATCCACATTAC

AAAATAATACTGATCAATTTAATGGTATTTTCGTTACGAATTATGATCATATATTTAACA

AGAAGATGGAAAAAGAAACTTGTTCTTCAAATAAAAAAATGAATAAATTAAAAAAAAGGA

AAAAGAAAAAAGGATTAAAAATGAAAACACAAGACGATAAGAACAAATTTGTTGAAAGCG

CGGTGAATAATTATTGGGATAATAAAATGGACAAAAATTTTAATAAAAAAAATTACATAA

CATCTGAAAATTTTAATGACAAATATGAAGGTTACTCAGACAATAATAAAAACAATTATG

ACATGTCTTATATAAATGATGATAATAATAAAGAAAAAAAGAAAAAAAGCAAAAAAAAAA

GAGCTTGTCAAAAATCTGAACAAATGAATGAAAAAAGAAACATATTAAATTATTTTTATA

AAAAGGATGATTTATATAATACATATAATAATAATTATGTTGATAAATTATATATGAATA

TAGATCCATGTGGACTATATGATATAAAAAATATTATGAGTACCTTATTAAATGTAGAAA

CCAAAAATGAGCATGTTGATACTTTTAAAAATATAGAATCACATATTACAAAATTACCTA

TATTTCATAATAATAAAATGAATAATAAAAAAAAGAATAAACATTTATATTTATTTTATT

TTATTTTTAATATAATATCGATTATGTCATGTATGTATTCATTTATATCATTATATTTAT

ATTTCAAAATATATTTCCTGTCGTTCTTCTTGTTATTCTTTTGTATATGTATTATATATA

TATACATATCTAATCATAAACCAATTAATCATATGTATAAGTGTTCGTTAAATATACCGT

ATAATATTAATTATGTGATTAATTTCTTGACAAGCCATAATAAATATCACCCAAACGAAA

TAAATAAAAAGGTATTTAAAACCAAAAATGATAATATACAATATGTTTATAGTACATTTA

ATATTTATTGTAAAAGTATTTTGTTTAAAATATTCCCATGTCACATATTTTTTAAGCCAA

GGAAAATTGTGTGTACTCAATTTACTCAGATATGTAATAGCGATAATAATATATTAAAAG

ATAAAGAAGGCGGATTGCGTAAATATATAATTCTGCAATATACAAATAAGAATACAAAAG

CATTTCTTAAATCTTTAGAATTTAATGAAGATATTAATAAGTTAGATATGAATATATATA

AAAACGTAATAGGTTTAAAACGTAACGATAAGGAGGAGGAGTTTTTTAAGCATGGACAGA

TTAAAATTGAGGACGTAAATAATGAGAACTTTAATAATGAACAGATGAAAAATAAATATA

ATGATGAAGAAAAAATGGATATATCAAAAAGTTTAAAAAGTGATTTTTTAGAAAAGGACC

AATCGCAAGTTATTACTTCAGAAGAGGATTCTGAGATGAAATCTTCATGTAGGGAGAAAA

ATAATAATATAAGAATGGGAATCCCCAAAATGAAAAACAATAGTAATTATAATAAAAATG

ATGATAATAATAATGATGATAATAATAATGATGATAATAATGATGGTAATAATAATAATG

ATGAAAATGATGACAATAATTATGGTAATAATAATAACAACAACAATAATAATAATAGTG

TTGATAAGTGTAAGGGGGGAAATACAAATGAACCATGCCATTATGTTGATGATACAAAAA

AATGGTCTAGTGATTTAAATCAAAATAGTGAATATCATATTTATAATAATTTGAATTATA

TATCGGCATATTGTTATAAACATATTAAAAAATTATTAGACAATGTAATAAAGAAAATAA

TATGTTCAATTCTTTTGAAATATAAATATGTATATAAATATTTTAAAAAATATTATTATA

ATAAATATGTAAATGTTGATGGATGTGATATTTTTTTAATACAAGAAAAAAAGGATAATA

CATGTGAACTTATTTTTTATACTTATTATAATTATAATAGTTTCTTTTTTAATGAATCAA

TAAATTATACTCGATGTAATAATATTCGCGAAAAGTTATTAACAGTTGATATTCCTAAAT

ATTTTAAACATGAGAATATTTTTTATTGTGACAAATATTATGTAGAAGAAAAAAATGAGA

CATTATTATTTCTTGAAAAAAAAAAAATCGAAAAAGAAATATGTAAAAATATATTTTTTA

ATATTAATAGTAATGAAACATATAATAAAAATATATTATTATATATATATGAATTAACAT

ATAAAGGATATTTTAATGAAAAATTTTTATCAAGTATTCACAATATAGATGATACATTCT

TTATAAGAAATATATCTAAAGTCTTTTTATATATAATACAAAATATAAATCTATATGATG

TTAAACAGTTTAAAAATATAAGTGAATTTATAAAAAATAAAAAAAATAAAATTGATTATA

CAGATGAATATTTTATATCTAATATTATTCATATAATTAATACATTACCTTTATTACATT

ATACATTCTCTACTTCACTCATATTTCCAAAAAAAAATGAAAAAATTGAATGTTCCTATG

AACAATCCAATATTGATATTACACTAAAAAACGATAATCCAATAACTTTTTTTATAATAG

CAGAAAATTATAAAAACAAGACAAAAATTACCTCAGAACTTATTTTAAAAACACTATGTA

CATTTGATCAAATAATCATATATATGCAACATCAAATGCAAATAACAAATAAAAGTAATT

ATAATATTACCTTTAATTATCCACACATTATATTAAGAAATCTATTACATGGACATATGA

ACTTTTCTTTTGCTCATAAAATGGTTATCAAGGATACTCTGAGCAACACTGCAGAAATTA

GGGTAATACACCAGAAAAAAGGAAATTTAAATGAAAATAAAATAAATGAATATTATATAT

ATATATATATATGTATGTATGTATGATTTCATTTTGTTTCTTATTTGTTTGTATATTTAT

TTATTTATTTATTTTATTTTTCTTTTATCATTGTAGTTTATTGATAAGGATAAATCTAAT

GGAGAGTTGTTTGGGGTTATTAAAAGGAATGAACTAATTACCGATTTTTTGAGCGGAAAT

ATTTTTGATAAAATAATTTTAAATAATGATAAACAGTAAGTATATAAACAAATTTTATTG

ATTATATGGCATTATTCATTTTTTCAATATTTCTTTAAATTATAAAAAGTTCATATGTGA

ATAATATAGAATTAATATGTTAATATGTAAATATGTAATTATGTATATGTATTATGTTAT

ATATATATATATATTTTTATTTATTTATTTGTTTAATGTCTTTTTAACTTTAGATATAAT

AGTGTCAAGGATATTAAAATAAATGTAATACACAAAGAAAATACCAAAAATTTGATAAAG

TAAATTAAAAAAAAAAAAAAAATTAAAAATACATAAATAGGAATAATATATATATTAAGG

AGTTAATATAATGCATACATATCATAAACCTTTTTTATAATATTTATAGGTCCTTTTTTA

GATAAcgtaattaattcatcagatataattaataataataataaaaaaaaaaaaaaaaaa

aaaaaaagtgtaactcaatctgaagttacatataaaagaaaatctttaaacttttttaaa

acggatacatacatgtaaatatatgtatgtttatgtagttcaaaaatataagcaaattaa

aaattttgtattgtttttaaaaaaaatatgtgtacttaatattttcatatatgaatatta

atatatatatatatatatatatatatttataagtgtttgagatatgtattctttatatat

tacattctatatacatttaattatttatttatttattatttatttatttttattctattt

catttttattttttttgttgcatattaaattcgttttaataatatttaaattatttattt

tatttttttttttttgaaaacaatcaaatattttaattcttaaaaagttataatttttaa

ataaacattggaataagtttgtaaa

**Original Locus (OL) amplicon size: 1123bp**s

**> PF3D7_1351000-mNG yDHODH mNG recodonPfPITP**

ataagaatatgaaaaatatgcaagtgtttacattttatttgtaaaaaaaataacaataccttacaaaagtgtgaataaataaataaataaataaataaatatatatatatatatatatatatatatatatatgtaatgatcagtatgtgtaaggataaatatatttaagtatatttcttctttctatgttttttttaaattatacatagaataaatataaggaataatcatatatatatatatatatatatatatgttatcatttcattcatatatttattttatgaattatatatttttaataatatatagctctattttcttctttttgataattaaaagtatttataatttataaagtttatttgtgtgcatattattacgtatataaataaatatatacatatatatatacatatatatatatatctatatattgtacatatgtttgtatgtgtatcaattcatttagatatgtttacatacacgcttaatataaaATG**AAAATTGTCGAATTTCGGCTTGCTATGCCTTTAACCATAGAAGAATATAAAGTATGTCAATTATATTTTGTAGCTAAAGCTTCATTAGAAGACGCAGAAAATAATATAAATAGTAATTGTGAAAAAGGTGATGAGAATAATGATTCAAAAAAAGGAATAGTAATATTAAAAAATGAAAGTTATATAAATGAGGATGGTACATGTGGTCAGTATACATATAAGAGAATAAATTTAATAAATAAGTTACCAAAGTGGCTATTAAATTTTATAGATCCTAAATATTGTATTATAGATGAAAAGTCATGGAATGCATATCCATATTTAAAAACAGTTTATGAGTCTAGTGGTTTTCCAAAAGCGAAGATTCAAGTTGAATCCGCACATTTTAATGGTTATGACACTGAAGAGAATGCTTTGAATTTATCAGAAGAAGCGCTCTCTTTAAGAAAAGTAATATATGTTGATATAGTAAATGATAAAATATCTTATAAAGATTATAATGAAAGTGAAGATCCTTCGTTATTTTATAGTGATAAAGCTAAAAGAGGTAAACTCGAAAAAAACTGGAAAGAAAATCATTCTATTATTATGACATGTTATAAGGTATTTACTATTAATATACCTTATTTTGGTATATTCTGTTCAAAATTAGAAAACTGGATTATATCTGCGCTCAGGGATAATATATTAAAATATCATAGGAAAGCATTCTGTTGGATTGATGAATGGATAGATTTGACAATTGATGATATAAGAAATTTAGAAAGAGATGTTCAAAAGAAATTGAATAAATTTTGGAACGACTCACAAGACGAAAATGTTACAGATGATATTTTGGATGAAGAAAATAACAATAAAAACGAAAATGTAAAAGATCAATATGTTATACATCATAACAATAATGAAGAAAATAATGATACACTAAGTAATGATATAATAGATTCGTTAACTTTTAAAAATAACCAAAAAGATGATATACAAAGAAATATGTCTAATGGT**GGTTTAAACGAGCAGAAGTTAATATCAGAAGAGGATTTGGGTGAACAAAAACTCATAAGCGAAGAAGATTTAATAACTTCGTATAGCATACATTATACGAAGTTATCCGGAGGAAGTGAAGGTAGAGGTTCTTTGTTGACTTGTGGTGATGTTGAAGAAAATCCAGGTCCAGCTAGCCCTAGGATGAAGATAGTTGAGTTCAGATTAGCAATGCCACTTACAATTGAGGAGTACAAGGTTTGCCAGCTTTACTTCGTTGCAAAGGCAAGTCTTGAGGATGCTGAGAACAACATTAACTCAAACTGCGAGAAGGGAGACGAAAACAACGACAGTAAGAAGGGTATTGTTATTCTTAAGAACGAGTCATACATTAACGAAGACGGAACTTGCGGACAATACACTTACAAACGTATTAACCTTATTAACAAACTTCCTAAATGGTTGCTTAACTTCATTGACCCAAAGTACTGCATAATTGACGAGAAAAGTTGGAACGCTTACCCTTACCTTAAGACTGTATACGAAAGCTCAGGATTCCCTAAGGCAAAAATACAGGTAGAGAGTGCTCACTTCAACGGATACGATACAGAGGAAAACGCACTTAACCTTAGTGAGGAGGCATTAAGCCTTCGTAAGGTTATTTACGTAGACATTGTTAACGACAAGATTAGCTACAAGGACTACAACGAGTCAGAGGACCCAAGTCTTTTCTACTCAGACAAGGCAAAGCGTGGAAAGTTAGAGAAGAATTGGAAGGAGAACCACAGCATAATAATGACTTGCTACAAAGTTTTCACAATAAACATTCCATACTTCGGAATTTTTTGCAGTAAGCTTGAGAATTGGATAATTAGCGCATTACGTGACAACATTCTTAAGTACCACCGTAAGGCTTTTTGCTGGATAGACGAGTGGATTGACCTTACTATAGACGACATTCGTAACCTTGAGCGTGACGTACAGAAAAAGCTTAACAAGTTCTGGAATGATAGTCAGGATGAGAACGTAACTGACGACATACTTGACGAGGAGAACAATAACAAGAATGAGAACGTTAAGGACCAGTACGTAATTCACCACAATAACAACGAGGAGAACAACGACACTTTGTCAAACGACATTATTGACAGTCTTACATTCAAGAACAATCAGAAGGACGACATTCAGCGTAACATGAGCAACGGAAACAACATGATGAACAACGACAAGATTTGCAACAACAACAACAACAACAACATATACTACAACGGTAATATACACAACAATGAGTGCACAAACAAGCAGAACCTTAGCCAGGGAAACGACCAGGGTCGTAAGGAGATTAGTAAGGATTTACCAAACGGTACATACATAACTAGTAGTTTGAACATGGACAACGACAACCTTGACTTCATTTGCACAAAGTACAACGAAGCATACGACGACAACGACAAGGAGAACAACAAGGATATGAAGAAGAAGAACATTAACATTAACATAAACGATATGCAGGGTGAGAAACAGATAATACAGAACGAGAACACTTTGAACAACAAGATGATAAACGTTTACGGAGACAACAACATAAACAACATAAACAACATAGCACGTAACAAGTCAGTAATAACAAAGTCAACTAGCTTGCTTTTTCCAAATGACCTTACAATTAAGAACAATTTAAACGGTAACCACAACAACGTAATTGACAACATTGAGAAAAAAAAGAAGAAGAAGAAGAAGAAGAAGGAGAAAAAGATGGAGAGTGAGAACTATTTCTTTTTCAAAAAGAACGAGGAGAATAAGGAGTTCGGAGAGTACCTTTATCGTTTGAACGAGGGAATGTTCTACAGTTGGAAATTAAGGTATTTCGTAATTAAGAACAATAAGCTTTGCTACTACGTAAACCACTCAAAGAACGAATTGAAGGGTGAGATAGACTTATTAAACGCACAAATTCAGTGGATTGGAGAGTACAAGGGTCGTAACTCAGTATTCGTTATTAATAGTCTTAGTAAAAACGTAAACTACCTTTCAAGTGAAAACGAGATTCAGACTAAAAAGTGCATGATTGACATTCAGATGGCAACACTTATGAACTCAGAGTCAGTTAAGAAACGTGAAATAAAGCAAGAGAACGGAAAGATAGTTTCAAAAGGAGTTCAGAACGGTGACAATAAGGAGGACGTAGACGACGACAAGGAGGACGACGACAACGACGACGACGAGGACGAGGACACAGAAGACGACAACAACGACGACGATAACGACGACGATAACGACGACGACAACGACGACGACAACGACGACGACAACGACGATGACAACGACGATGACAACGACAATAACAACGACGACGACAACGACGATGACAACGACGACGACAACGACAATAACAACGACGACGACAACTACGTTGACAATAATCACTATAATAATAACGACGACAATCACATAAACTACAACAACAAGGATACTCCAAACTACCACACTCACAAGAATAAGCGTATTGTAGACAATAAGAAGAGTCAGGACATGCCTAAGAACAAGAACGACGGTACAATTAACAACTACAATCTTCCTTACAGTAAGGAGAGTAACAAGGAAATAGACATTAACAACAACCACTACAACAACTACATTTGCGTTAACGACAAAGATGAGATAGTACACTCAAAACTTATATTATCAGACAACGACCTTAGTCAGTACCACAACGAGGAAAACAAGACTATAAGTAACAATAGTCACTACATTAGTACTCTTCAGAACAACACAGACCAGTTCAACGGAATATTTGTAACAAACTACGACCACATTTTCAATAAAAAAATGGAGAAGGAGACATGCAGCAGTAACAAGAAGATGAACAAGCTTAAGAAGCGTAAGAAAAAGAAGGGTCTTAAGATGAAGACTCAGGATGACAAAAATAAGTTCGTAGAGTCAGCAGTTAACAACTACTGGGACAACAAGATGGATAAGAACTTCAACAAGAAGAACTATATTACTAGCGAGAACTTCAACGATAAGTACGAGGGATATAGTGATAACAACAAGAATAACTACGATATGAGCTACATTAACGACGACAACAACAAGGAGAAGAAAAAGAAGTCAAAGAAGAAGCGTGCATGCCAGAAGAGCGAGCAGATGAACGAGAAGCGTAATATTCTTAACTACTTCTACAAGAAAGACGACCTTTACAACACTTACAACAACAACTACGTAGACAAGCTTTACATGAACATTGACCCTTGCGGTTTGTACGACATTAAGAACATAATGTCAACACTTCTTAACGTTGAGACAAAGAACGAACACGTAGACACATTCAAGAACATTGAGAGTCACATAACTAAGCTTCCAATTTTCCACAACAACAAGATGAACAACAAGAAGAAAAACAAGCACCTTTACCTTTTCTACTTCATATTCAACATTATTAGTATAATGAGTTGCATGTACAGTTTCATTAGTCTTTACCTTTACTTTAAGATTTACTTTTTAAGTTTTTTTCTTCTTTTTTTCTGCATTTGCATAATTTACATTTATATTAGCAACCACAAGCCTATAAACCACATGTACAAATGCAGTCTTAACATTCCATACAACATAAACTACGTTATAAACTTTCTTACTTCACACAACAAGTACCATCCTAATGAGATTAACAAGAAAGTTTTCAAGACAAAGAACGACAACATTCAGTACGTATACTCAACTTTCAACATATACTGCAAGTCAATACTTTTCAAGATTTTTCCTTGCCATATTTTCTTCAAACCTCGTAAGATAGTTTGCACACAGTTCACACAAATTTGCAACTCAGACAACAACATTCTTAAGGACAAGGAGGGAGGTCTTAGAAAGTACATTATATTACAGTACACTAACAAAAACACTAAGGCTTTCTTAAAGAGCCTTGAGTTCAACGAGGACATAAACAAACTTGACATGAACATTTACAAGAATGTTATTGGACTTAAGAGAAATGACAAAGAAGAAGAATTCTTCAAACACGGTCAAATAAAGATAGAAGATGTTAACAACGAAAATTTCAACAACGAGCAAATGAAGAACAAGTACAACGACGAGGAGAAGATGGACATTAGTAAGTCACTTAAGTCAGACTTCCTTGAGAAAGATCAGAGTCAGGTAATAACAAGTGAGGAAGACAGCGAAATGAAGAGCAGTTGCCGTGAAAAGAACAACAACATTCGTATGGGTATACCAAAGATGAAGAATAACTCAAACTACAACAAGAACGACGACAACAACAACGACGACAACAACAACGACGACAACAACGACGGAAACAACAACAACGACGAGAACGACGATAACAACTACGGAAACAACAACAAcAAcAATAAtAACAACAACTCAGTAGACAAATGCAAAGGAGGTAACACTAACGAGCCTTGTCACTACGTAGACGACACTAAGAAGTGGAGCTCAGACCTTAACCAGAACTCAGAGTACCACATATACAACAACCTTAACTACATTAGTGCTTACTGCTACAAGCACATAAAGAAGCTTCTTGATAACGTTATTAAAAAGATTATTTGCAGTATATTACTTAAGTACAAGTACGTTTACAAGTACTTCAAGAAGTACTACTACAACAAGTACGTTAACGTAGACGGTTGCGACATATTCCTTATTCAGGAGAAGAAAGACAACACTTGCGAGTTAATATTCTACACATACTACAACTACAACTCATTTTTCTTCAACGAGAGTATTAACTACACAAGGTGCAACAACATAAGAGAGAAACTTCTTACTGTAGACATACCAAAGTACTTCAAGCACGAAAACATATTCTACTGCGATAAGTACTACGTTGAGGAGAAGAACGAAACTCTTCTTTTCTTAGAGAAGAAGAAGATAGAGAAGGAGATTTGCAAGAACATTTTCTTCAACATAAACTCAAACGAGACTTACAACAAGAACATTCTTCTTTACATTTACGAGCTTACTTACAAGGGTTACTTCAACGAGAAGTTCCTTAGTTCAATACATAACATTGACGACACTTTTTTCATTCGTAACATTAGCAAGGTTTTCCTTTACATTATTCAGAACATTAACTTGTACGACGTAAAGCAATTCAAGAACATTTCAGAGTTCATTAAGAACAAGAAGAACAAGATAGACTACACTGACGAGTACTTCATTAGCAACATAATACACATTATAAACACTCTTCCACTTCTTCACTACACTTTTAGCACAAGTTTAATTTTCCCTAAGAAGAACGAGAAGATAGAGTGCAGTTACGAGCAGAGTAACATAGACATAACTTTGAAGAATGACAACCCTATTACATTCTTCATTATTGCTGAGAACTACAAGAATAAAACTAAGATAACAAGTGAGTTAATACTTAAGACTTTGTGCACTTTCGACCAGATTATAATTTACATGCAGCACCAGATGCAGATTACTAACAAGTCAAACTACAACATAACATTCAACTACCCTCATATAATTCTTCGTAACTTGCTTCACGGTCACATGAATTTCAGCTTCGCACACAAGATGGTAATAAAAGACACATTATCAAATACAGCTGAGATACGTTTCATAGACAAAGACAAGAGCAACGGTGAACTTTTCGGAGTAATAAAGCGTAACGAGTTGATAACAGACTTCCTTTCAGGTAACATATTCGACAAGATTATACTTAACAACGACAAGCAGTACAACTCAGTTAAAGACATAAAGATTAACGTTATTCATAAGGAGAACACAAAGAACCTTATTAAAAGTTTCTTCCGTGCTAGAGGTGCTGCTGCTGGTGCTGGAGGTGCAGGTAGACGTACGATGGTAAGTAAGGGAGAAGAAGACAACATGGCATCATTGCCAGCCACCCACGAGTTGCATATTTTCGGTTCTATTAACGGAGTGGACTTTGATATGGTTGGTCAGGGTACAGGAAATCCTAATGATGGTTATGAGGAGCTTAATTTGAAATCTACTAAAGGTGACTTGCAGTTTTCACCTTGGATACTTGTTCCACATATAGGTTACGGTTTTCACCAGTACCTTCCATATCCAGATGGAATGTCTCCATTCCAGGCTGCTATGGTGGATGGAAGTGGTTATCAAGTACATAGGACCATGCAGTTTGAGGACGGTGCATCTTTAACCGTTAACTATAGATATACCTATGAAGGATCTCACATTAAAGGTGAAGCCCAAGTGAAAGGTACCGGATTTCCTGCTGACGGTCCAGTGATGACTAACAGTTTAACTGCCGCAGACTGGTGCCGTTCTAAAAAGACATATCCAAACGATAAGACTATAATTTCTACCTTTAAATGGTCATACACAACAGGAAATGGAAAACGTTATAGATCAACTGCAAGGACTACTTATACATTCGCTAAACCAATGGCCGCCAATTATCTTAAAAATCAACCTATGTATGTTTTCCGTAAAACTGAATTAAAGCACTCAAAGACAGAACTTAATTTTAAGGAATGGCAAAAAGCATTTACTGACGTAATGGGAATGGATGAGTTGTATAAGGAGGGTCGTGGTTCACTTCTTACTTGCGGTGACGTTGAGGAGAACCCTGGTCCTATAACTTCGTATAGCATACATTATACGAAGTTATTAGGAGAAGGAAGAGGAAGTTTATTAACATGTGGAGATGTAGAAGAAAATCCAGGACCAATGATTGAACAAGATGGATTGCACGCAGGTTCTCCGGCCGCTTGGGTGGAGAGGCTATTCGGCTATGACTGGGCACAACAGACAATCGGCTGCTCTGATGCCGCCGTGTTCCGGCTGTCAGCGCAGGGGCGCCCGGTTCTTTTTGTCAAGACCGACCTGTCCGGTGCCCTGAATGAACTGCAGGACGAGGCAGCGCGGCTATCGTGGCTGGCCACGACGGGCGTTCCTTGCGCAGCTGTGCTCGACGTTGTCACTGAAGCGGGAAGGGACTGGCTGCTATTGGGCGAAGTGCCGGGGCAGGATCTCCTGTCATCTCACCTTGCTCCTGCCGAGAAAGTATCCATCATGGCTGATGCAATGCGGCGGCTGCATACGCTTGATCCGGCTACCTGCCCATTCGACCACCAAGCGAAACATCGCATCGAGCGAGCACGTACTCGGATGGAAGCCGGTCTTGTCGATCAGGATGATCTGGACGAAGAGCATCAGGGGCTCGCGCCAGCCGAACTGTTCGCCAGGCTCAAGGCGCGCATGCCCGACGGCGAGGATCTCGTCGTGACCCATGGCGATGCCTGCTTGCCGAATATCATGGTGGAAAATGGCCGCTTTTCTGGATTCATCGACTGTGGCCGGCTGGGTGTGGCGGACCGCTATCAGGACATAGCGTTGGCTACCCGTGATATTGCTGAAGAGCTTGGCGGCGAATGGGCTGACCGCTTCCTCGTGCTaTACGGTATCGCCGCTCCCGATTCGCAGCGCATCGCCTTCTATCGCCTTCTTGACGAGTTCTTCTGActcgagggatatggcagcttaatgttcgtttttcttatttatatatttataccaattgattgtatttataactgtaaaaatgtgtatgttgtgtgcatatttttttttgtgcatgcacatgcatgtaaatagctaaaattatgaacattttattttttgttcagaaaaaaaaaactttacacacataaaatggctagtatgaatagccatattttatataaattaaatcctatgaatttatgaccatattaaaaatttagatatttatggaacataatatgtttgaaacaataagacaaaattattattattattattatttttactgttataattatgtgtctccttcaatgattcataaatagttggacttgatttttaaaatgtttataatatgattagcatagttaaataaaaaaagttgaaaaattaaaaaaaaacatataaacacaaatgatggtttttccttcaatttcgatatcaatttatagaaacaaaatatatacttgtataattttatttttttatataaatcattacatatataattatacaatattttttctaagagataattatatattaatatatataaaaaaaggtgttttttttttttttttttatttttatttttattttatggtaatattttattttccttattttataaattatattagtttatatgtgattaattttatatattatcaatttatatatttttaaatgcttacttaattatctttttttttttttttttttttttttcccctctttttatattaatttatttttgaaaaaattgatatatatatatatatataatatatatatatacatgtagtagtattaaacaatgtataatatatataaataatatatttatatatttcatttcaattttaattttttttggttttttttttttttctttttgtcatatttaaaaaaaattatattcatataagttatgcattttttataaacattattcaatatatgtataatataatatatatatatatattaatgtattattccaatgtgcatgataaaagaaaaaaataatatttataaaaaaaaagaaaaataaaacaaaaaaagaaaaaaaaaaaaaaaaaaaaaaaaatacaaaaataaataatataatttataattatatattcttgtcacaataaaaatatatatatatatatatatatttataatatgtatattttaaactagaaaaggaataactaatattttatttattatcattcaagatttatattttataataataaatacctaatagaaatatatcaggatccatgcatggttcgctaaactgcatcgtcgctgtgtcccagaacatgggcatcggcaagaacggggactacccctggccaccgctcaggaacgaatttagatatttccagagaatgaccacaacctcttcagtagaaggtaaacagaatctggtgattatgggtaagaagacctggttctccattcctgagaagaatcgacctttaaagggtagaattaatttagttctcagcagagaactcaaggaacctccacaaggagctcattttctttccagaagtctagatgatgccttaaaacttactgaacaaccagaattagcaaataaagtagacatggtctggatagttggtggcagttctgtttataaggaagccatgaatcacccaggccatcttaaactatttgtgacaaggatcatgcaagactttgaaagtgacacgttttttccagaaattgatttggagaaatataaacttctgccagaatacccaggtgttctctctgatgtccaggaggagaaaggcattaagtacaaatttgaagtatatgagaagaatgattaagcttatttaataatagattaaaaatattataaaaataaaaacataaacacagaaattacaaaaaaaatacatatgaattttttttttgtaatcttccttataaatatagaataatgaatcatataaaacatatcattattcatttatttacatttaaaattattgtttcagtatctttaatttattatgtatatataaaaataacttacaattttattaataaacaatatatgtttattaattcatgttttgtaatttatgggatagcgattttttttactgtctgtatttttcttttttaattatgttttaattgtattttatttttattattgttctttttatagtattattttaaaacaaaatgtattttctaagaacttataataataataaatataaattttaataaaaattatatttatcttttacaatatgaacataaagtacaacattaatatatagcttttaatatttttattcctaatcatgtaaatcttaaatttttctttttaaacatatgttaaatatttatttctcattatatataagaacatatttattaaatctagaattctatagtgagtcgtattacaattcactggccgtcgttttacaacgtcgtgactgggaaaaccctggcgttacccaacttaatcgccttgcagcacatccccctttcgccagctggcgtaatagcgaagaggcccgcaccgatcgcccttcccaacagttgcgcagcctgaatggcgaatggcgcctgatgcggtattttctccttacgcatctgtgcggtatttcacaccgcatatggtgcactctcagtacaatctgctctgatgccgcatagttaagccagccccgacacccgccaacacccgctgacgcgccctgacgggcttgtctgctcccggcatccgcttacagacaagctgtgaccgtctccgggagctgcatgtgtcagaggttttcaccgtcatcaccgaaacgcgcgagacgaaagggcctcgtgatacgcctatttttataggttaatgtcatgataataatggtttcttagacgtcaggtggcacttttcggggaaatgtgcgcggaacccctatttgtttatttttctaaatacattcaaatatgtatccgctcatgagacaataaccctgataaatgcttcaataatattgaaaaaggaagagtatgagtattcaacatttccgtgtcgcccttattcccttttttgcggcattttgccttcctgtttttgctcacccagaaacgctggtgaaagtaaaagatgctgaagatcagttgggtgcacgagtgggttacatcgaactggatctcaacagcggtaagatccttgagagttttcgccccgaagaacgttttccaatgatgagcacttttaaagttctgctatgtggcgcggtattatcccgtattgacgccgggcaagagcaactcggtcgccgcatacactattctcagaatgacttggttgagtactcaccagtcacagaaaagcatcttacggatggcatgacagtaagagaattatgcagtgctgccataaccatgagtgataacactgcggccaacttacttctgacaacgatcggaggaccgaaggagctaaccgcttttttgcacaacatgggggatcatgtaactcgccttgatcgttgggaaccggagctgaatgaagccataccaaacgacgagcgtgacaccacgatgcctgtagcaatgccaacaacgttgcgcaaactattaactggcgaactacttactctagcttcccggcaacaattaatagactggatggaggcggataaagttgcaggaccacttctgcgctcggcccttccggctggctggtttattgctgataaatctggagccggtgagcgtgggtctcgcggtatcattgcagcactggggccagatggtaagccctcccgtatcgtagttatctacacgacggggagtcaggcaactatggatgaacgaaatagacagatcgctgagataggtgcctcactgattaagcattggtaactgtcagaccaagtttactcatatatactttagattgatttaaaacttcatttttaatttaaaaggatctaggtgaagatcctttttgataatctcatgaccaaaatcccttaacgtgagttttcgttccactgagcgtcagaccccgtagaaaagatcaaaggatcttcttgagatcctttttttctgcgcgtaatctgctgcttgcaaacaaaaaaaccaccgctaccagcggtggtttgtttgccggatcaagagctaccaactctttttccgaaggtaactggcttcagcagagcgcagataccaaatactgtccttctagtgtagccgtagttaggccaccacttcaagaactctgtagcaccgcctacatacctcgctctgctaatcctgttaccagtggctgctgccagtggcgataagtcgtgtcttaccgggttggactcaagacgatagttaccggataaggcgcagcggtcgggctgaacggggggttcgtgcacacagcccagcttggagcgaacgacctacaccgaactgagatacctacagcgtgagctatgagaaagcgccacgcttcccgaagggagaaaggcggacaggtatccggtaagcggcagggtcggaacaggagagcgcacgagggagcttccagggggaaacgcctggtatctttatagtcctgtcgggtttcgccacctctgacttgagcgtcgatttttgtgatgctcgtcaggggggcggagcctatcgaaaaacgccagcaacgcggcctttttacggttcctggccttttgctggccttttgctcacatgttctttcctgcgttatcccctgattctgtggataaccgtattaccgcctttgagtgagctgataccgctcgccgcagccgaacgaccgagcgcagcgagtcagtgagcgaggaagcggaagagcgcccaatacgcaaaccgcctctccccgcgcgttggccgattcattaatgcagctggcacgacaggtttcccgactggaaagcgggcagtgagcgcaacgcaattaatgtgagttagctcactcattaggcaccccaggctttacactttatgcttccggctcgtatgttgtgtggaattgtgagcggataacaatttcacacaggaaacagctatgaccatgattacgccaagctatttaggtgacactatagaatactcgcggccgcTAGaaaattgtcgaatttcggcttgctatgcctttaaccatagaagaatataaagtatgtcaattatattttgtagctaaagcttcattagaagacgcagaaaataatataaatagtaattgtgaaaaaggtgatgagaataatgattcaaaaaaaggaatagtaatattaaaaaatgaaagttatataaatgaggatggtacatgtggtcagtatacatataagagaataaatttaataaataagttaccaaagtggctattaaattttatagatcctaaatattgtattatagatgaaaagtcatggaatgcatatccatatttaaaaacagtttatgagtctagtggttttccaaaagcgaagattcaagttgaatccgcacattttaatggttatgacactgaagagaatgctttgaatttatcagaagaagcgctctctttaagaaaagtaatatatgttgatatagtaaatgataaaatatcttataaagattataatgaaagtgaagatccttcgttattttatagtgataaagctaaaagaggtaaactcgaaaaaaactggaaagaaaatcattctattattatgacatgttataaggtatttactattaatataccttattttggtatattctgttcaaaattagaaaactggattatatctgcgctcagggataatatattaaaatatcataggaaagcattctgttggattgatgaatggatagatttgacaattgatgatataagaaatttagaaagagatgttcaaaagaaattgaataaattttggaacgactcacaagacgaaaatgttacagatgatattttggatgaagaaaataacaataaaaacgaaaatgtaaaagatcaatatgttatacatcataacaataatgaagaaaataatgatacactaagtaatgatataatagattcgttaacttttaaaaataaccaaaaagatgatatacaaagaaatatgtctaatggtAATAATATGATGAATAATGATAAAATATGTAATAATAATAATAATAATAATATTTATTATAATGGAAACATTCATAATAACGAATGTACTAATAAACAAAATTTATCTCAAGGTAATGATCAAGGAAGGAAAGAAATATCAAAAGACCTGCCCAATGGAACTTATATTACATCATCACTAAATATGGATAATGATAATTTAGATTTTATATGTACCAAATATAATGAGGCTTATGATGATAATGATAAAGAAAATAATAAAGACATGAAAAAAAAAAATATAAATATAAATATTAATGACATGCAAGGAGAAAAGCAAATCATTCAAAATGAAAATACACTAAATAATAAAATGATTAATGTATATGGTGATAATAATATCAATAATATTAATAATATTGCTAGAAATAAAAGTGTTATTACTAAAAGTACATCTCTATTATTCCCTAACGATTTAACCATAAAAAATAACCTCAATGGAAATCATAATAATGTTATAGATAATATAGAAAAGAAGAAAAAAAAAAAAAAAAAAAAAAAAGAAAAGAAAATGGAATCGGAAAATTACTTTTTCTTTAAGAAAAATGAAGAAAACAAAGAATTTGGTGAATATTTATACAGACTAAATGAAGGCATGTTTTATTCATGGAAGCTTCGATACTTTGTTATAAAAAATAACAAATTGTGTTATTATGTTAATCATAGTAAAAATGAGCTAAAAGGAGAAATCGATCTCCTGAATGCGCAGATACAATGGATAGGTGAATATAAAGGAAGAAATAGTGTTTTTGTAATAAACTCCTTATCAAAGAATGTTAATTATTTAAGTTCAGAGAATGAAATACAAACAAAGAAATGTATGATAGATATACAAATGGCTACTTTAATGAATAGTGAAAGTGTGAAAAAGAGAGAGATTAAACAGGAAAATGGGAAAATTGTGAGTAAGGGGGTACAAAATGGAGATAACAAAGAAGATGTTGATGATGATAAAGAAGATGATGATAATGATGATGATGAAGATGAAGATACTGAGGATGATAATAATGATGATGACAATGATGATGACAATGATGATGATAATGATGATGATAATGATGATGATAATGATGACGATAATGATGACGATAATGATAACAATAATGATGATGATAATGATGACGATAATGATGATGATAATGATAACAATAATGATGATGATAATTATGTGGATAACAACCATTACAACAACAATGATGATAACCATATTAATTATAATAATAAAGACACACCGAATTATCATACACATAAAAACAAAAGAATAGTTGATAACAAAAAATCACAAGATATGCCAAAAAATAAAAATGATGGAACTATAAATAATTATAACTTACCATATTCAAAAGAATCAAATAAAGAGATCGATATAAATAATAATCATTATAATAATTATATATGTGTAAATGATAAGGACGAAATTGTTCATAGTAAGTTAATTCTTAGCGATAATGATTTATCACAATATCATAATGAAGAGAATAAAACAATTTCAAATAACTCACATTATATATCCACATTACAAAATAATACTGATCAATTTAATGGTATTTTCGTTACGAATTATGATCATATATTTAACAAGAAGATGGAAAAAGAAACTTGTTCTTCAAATAAAAAAATGAATAAATTAAAAAAAAGGAAAAAGAAAAAAGGATTAAAAATGAAAACACAAGACGATAAGAACAAATTTGTTGAAAGCGCGGTGAATAATTATTGGGATAATAAAATGGACAAAAATTTTAATAAAAAAAATTACATAACATCTGAAAATTTTAATGACAAATATGAAGGTTACTCAGACAATAATAAAAACAATTATGACATGTCTTATATAAATGATGATAATAATAAAGAAAAAAAGAAAAAAAGCAAAAAAAAAAGAGCTTGTCAAAAATCTGAACAAATGAATGAAAAAAGAAACATATTAAATTATTTTTATAAAAAGGATGATTTATATAATACATATAATAATAATTATGTTGATAAATTATATATGAATATAGATCCATGTGGACTATATGATATAAAAAATATTATGAGTACCTTATTAAATGTAGAAACCAAAAATGAGCATGTTGATACTTTTAAAAATATAGAATCACATATTACAAAATTACCTATATTTCATAATAATAAAATGAATAATAAAAAAAAGAATAAACATTTATATTTATTTTATTTTATTTTTAATATAATATCGATTATGTCATGTATGTATTCATTTATATCATTATATTTATATTTCAAAATATATTTCCTGTCGTTCTTCTTGTTATTCTTTTGTATATGTATTATATATATATACATATCTAATCATAAACCAATTAATCATATGTATAAGTGTTCGTTAAATATACCGTATAATATTAATTATGTGATTAATTTCTTGACAAGCCATAATAAATATCACCCAAACGAAATAAATAAAAAGGTATTTAAAACCAAAAATGATAATATACAATATGTTTATAGTACATTTAATATTTATTGTAAAAGTATTTTGTTTAAAATATTCCCATGTCACATATTTTTTAAGCCAAGGAAAATTGTGTGTACTCAATTTACTCAGATATGTAATAGCGATAATAATATATTAAAAGATAAAGAAGGCGGATTGCGTAAATATATAATTCTGCAATATACAAATAAGAATACAAAAGCATTTCTTAAATCTTTAGAATTTAATGAAGATATTAATAAGTTAGATATGAATATATATAAAAACGTAATAGGTTTAAAACGTAACGATAAGGAGGAGGAGTTTTTTAAGCATGGACAGATTAAAATTGAGGACGTAAATAATGAGAACTTTAATAATGAACAGATGAAAAATAAATATAATGATGAAGAAAAAATGGATATATCAAAAAGTTTAAAAAGTGATTTTTTAGAAAAGGACCAATCGCAAGTTATTACTTCAGAAGAGGATTCTGAGATGAAATCTTCATGTAGGGAGAAAAATAATAATATAAGAATGGGAATCCCCAAAATGAAAAACAATAGTAATTATAATAAAAATGATGATAATAATAATGATGATAATAATAATGATGATAATAATGATGGTAATAATAATAATGATGAAAATGATGACAATAATTATGGTAATAATAATAACAACAACAATAATAATAATAGTGTTGATAAGTGTAAGGGGGGAAATACAAATGAACCATGCCATTATGTTGATGATACAAAAAAATGGTCTAGTGATTTAAATCAAAATAGTGAATATCATATTTATAATAATTTGAATTATATATCGGCATATTGTTATAAACATATTAAAAAATTATTAGACAATGTAATAAAGAAAATAATATGTTCAATTCTTTTGAAATATAAATATGTATATAAATATTTTAAAAAATATTATTATAATAAATATGTAAATGTTGATGGATGTGATATTTTTTTAATACAAGAAAAAAAGGATAATACATGTGAACTTATTTTTTATACTTATTATAATTATAATAGTTTCTTTTTTAATGAATCAATAAATTATACTCGATGTAATAATATTCGCGAAAAGTTATTAACAGTTGATATTCCTAAATATTTTAAACATGAGAATATTTTTTATTGTGACAAATATTATGTAGAAGAAAAAAATGAGACATTATTATTTCTTGAAAAAAAAAAAATCGAAAAAGAAATATGTAAAAATATATTTTTTAATATTAATAGTAATGAAACATATAATAAAAATATATTATTATATATATATGAATTAACATATAAAGGATATTTTAATGAAAAATTTTTATCAAGTATTCACAATATAGATGATACATTCTTTATAAGAAATATATCTAAAGTCTTTTTATATATAATACAAAATATAAATCTATATGATGTTAAACAGTTTAAAAATATAAGTGAATTTATAAAAAATAAAAAAAATAAAATTGATTATACAGATGAATATTTTATATCTAATATTATTCATATAATTAATACATTACCTTTATTACATTATACATTCTCTACTTCACTCATATTTCCAAAAAAAAATGAAAAAATTGAATGTTCCTATGAACAATCCAATATTGATATTACACTAAAAAACGATAATCCAATAACTTTTTTTATAATAGCAGAAAATTATAAAAACAAGACAAAAATTACCTCAGAACTTATTTTAAAAACACTATGTACATTTGATCAAATAATCATATATATGCAACATCAAATGCAAATAACAAATAAAAGTAATTATAATATTACCTTTAATTATCCACACATTATATTAAGAAATCTATTACATGGACATATGAACTTTTCTTTTGCTCATAAAATGGTTATCAAGGATACTCTGAGCAACACTGCAGAAATTAGGGTAATACACCAGAAAAAAGGAAATTTAAATGAAAATAAAATAAATGAATATTATATATATATATATATATGTATGTATGTATGATTTCATTTTGTTTCTTATTTGTTTGTATATTTATTTATTTATTTATTTTATTTTTCTTTTATCATTGTAGTTTATTGATAAGGATAAATCTAATGGAGAGTTGTTTGGGGTTATTAAAAGGAATGAACTAATTACCGATTTTTTGAGCGGAAATATTTTTGATAAAATAATTTTAAATAATGATAAACAGTAAGTATATAAACAAATTTTATTGATTATATGGCATTATTCATTTTTTCAATATTTCTTTAAATTATAAAAAGTTCATATGTGAATAATATAGAATTAATATGTTAATATGTAAATATGTAATTATGTATATGTATTATGTTATATATATATATATATTTTTATTTATTTATTTGTTTAATGTCTTTTTAACTTTAGATATAATAGTGTCAAGGATATTAAAATAAATGTAATACACAAAGAAAATACCAAAAATTTGATAAAGTAAATTAAAAAAAAAAAAAAAATTAAAAATACATAAATAGGAATAATATATATATTAAGGAGTTAATATAATGCATACATATCATAAACCTTTTTTATAATATTTATAGGTCCTTTTTTAGATAAcgtaattaattcatcagatataattaataataataataaaaaaaaaaaaaaaaaaaaaaaaagtgtaactcaatctgaagttacatataaaagaaaatctttaaacttttttaaaacggatacatacatgtaaatatatgtatgtttatgtagttcaaaaatataagcaaattaaaaattttgtattgtttttaaaaaaaatatgtgtacttaatattttcatatatgaatattaatatatatatatatatatatatatatttataagtgtttgagatatgtattctttatatattacattctatatacatttaattatttatttatttattatttatttatttttattctatttcatttttattttttttgttgcatattaaattcgttttaataatatttaaattatttattttatttttttttttttgaaaacaatcaaatattttaattcttaaaaagttataatttttaa

ataaacattggaataagtttgtaaa

**5’ amplicon size: 1835bps**

**3’ amplicon size: 1183bps**

### **> PF3D7_1131800 (PfOSBP)**

tatatatatatatatatatatattgaggtctattatatacaactacataatatatatatatatatatatatatatatattatatatgtgtgtttacttatttgtttttttatttttctaattatataatttgttgtttattaaaaaaagaaaaaaatcataataaataagtaaaaagaaaaagaagaattaattagcatgtgttatctttttttattttttaaaatgaaagcaaaataataaaagagtataatataattatttaacacaatatatatatatatatatatatatatatgtcgcattaaatttatgtacatttcatatatattattcatattaatatttttatttgtatcttcattttaattttatattatatatatatataatatatatataatataatataatatatataagaaaaaattgaatcatgaatacattataagtgctgaaaaaaaaaaaaaaaaaaaaaaaaaaaacgataaaaagaaaattATGATAGGTGGTAAGTATTTAAATATAAAAATTTTTGGAGAGaAAACGTAATTCATTAGGTAGCAATAAAACTTCAGATAATACGATAAATAATAATAATGTAAATTATATAAAAAGGAAAAAAAATAATAATAAGAAAGATCAAGAAAAAAAATATAATGATAATAATTATGAAGAAGAATATATAAAGAAAAGAAAATCAAAACCAGGATCATTATATAATAATATAGACAAAAGTAAAATTTATCGAGATAAAAAAATTATACATGAAGGATGGTTAAATAAATGGACTAATATTATTGGAAGTTATAGACCTAGATATTTTATTTTAGAAAATGGATTATTAAGATATTCACTTGATAAATATTCTCCTACGAAAGAGTCTTTTGTATTAACACATTGTAAAATTAAAGTATGTCCAGATGATCAATTACATTTTGAAATCGATACAAACGAACAAGGTGTGTTATATTTAAAAGCAAACTCACCAGAAGATAAACATAAATGGTATATCTCTTTTAAAAAAGCACAACTAAATTATATTAATGGTAATAATTATAAAAACAAAAATGGACATCATAATATACCTGATGCAGTACAAACATTTAATATGTCTAATAATTCTTTATTTTTAAAAAATATTATTAAAAATTCATCCAAATGTATTAAAAGGGAAGATCAACATTTGGATGATCCTATAAAGAACACTAACAATCAAACAAAAAAGGAGTTGTCTAATAATAATAAAGAAATAATAGAAAAAGAAAAAAACGTTGATTGTATTATAGTTAATAAAATTCAGGAAAATGTAAAAGGGGACGAATCTAATAATATTGTAAAAAATGTGATATACGAAAGGGATGATAAAGATGTGCCATATGAAAGAAAAGATATATGTATGTCACATGATAAAAATTATAGAAGTCCTTTATATATTGACAATGATAATATGTCATCATATAGTAAGATAAATAAAAATGCAAATATTAAATATAAATCAAATGAATATATTTTTTATAAAGATGATAATGTTGAAAAATTATCATCATCATCATCATCCACCTTGGATACATCTGAATTTATTGAAAAAATGAATAGTTATAATAAAAAATATAATCTAGAAGATATGTTTATTAGTTCAACAGATTTTGAAGATAAAAGTCCAACCTTATGTTTAATGAAGAATATAATATCTCTCAAGGAAATGACAACGGATGTTTTAAAGGGTTCTGAGTATTATCAAGCCAGATCTATATTAAATAAAAAATTAAAAAATAATATTAAAAAGGGTGTTAATAGTAATATTATTGTTGATGTTAATAATAATATTATTGTTGATGTTAATAATAACATTATTGGTGATGTTAATAATAATATTATTGTTGATGTTAATAATAACATTATTGGTGATGTTAATAACAACATTATTGGTGATGTTAATAACAACATGATTGGTGATGTTAATAATAATATTATTGTTGATGTTAATAACAACATGATTGGTGATGTTAATAACAACATGATTGGTGATGTTAATAACAACATGATTGGTGATGCTAATGATTATGTTAATAACAATTTTAGTAATGGTAATAATTTACAAATAAAAAGTGATGAATTACTTTCCTTATTATCACAACTATACTTTTCCATACAATATGCTCATATTATTATAGAGAAATATATACAATGTACCGAATTATTATTAAAGGAAGAATCCATACATTCTAAATATATGAATCAGTCTTTAAAACTTTTAGCCAAACAAAATTATTATCTTGAAAAATCCCAAGAAATTAATAATCTTACTCATATGGAAGAAGAAATAAAAGATAAATTAAAACACTTTCAACTTTATCAATGTGCTACTTTAAGTGAAGAGGACGAAGAAGAAGAAGATAAAGAAATGTATAATTCATCAAAACAAATAAAACAGAAAAAACAACAGCAAGAGGATTTATTTTATGATTGTGATGAATTTATTTATGATAAAAATATATTATCAAATGACGTTTTTGAAGACGAATCCAAAAACTCTATCAGCTTATATAGTAGTGAGGGAGATAAGAATGAAAACTCAGATAAAATAAAAAAGGTGCATAATACAAATAACATACCAGATCAAAATCGTACCTATAATAATGAACACATGTTAACAACACATAGTACTATTAATAATAAAAATGAAAAGATGAATGCTCATATAGATGAACATTATTATAATTATATTAATAATGATCAGGAAAATGAAAAAACTTTTTCACATCAAACATCAGAGCAAAGAAGTTTAAATATAAAAACAAAATCTAGTGATCATGATGATAATATAACTACCATTCAAGAGGATTGTACAACAGATGTTTTTCTGAAAACAATTTTGAATAATAAAAATATTAATGAAACAAAAAGGAATACGGTGAGTACAGAGGATGATATTGCTCTTGGCGATGTGAGGGACGGTTTAATAACATGCGATGTTTATAAAAAGGATCATTTTGTGTGTGATGACAGTGTGGAGGATAAGGATATAATACGCAAACGAATGGGGATTGAAGATGATCATGGCAATAATGAAGATGTTCATGGCAATAATGAAGATGTTCATGGCAATAATGAAGATGTTCATGGCAATAATGAAGATGTTCATGGCAATAATAAAGATGTTCATAACAATAATAAAGATGATCATAACAATAATGATAATATTTGTGTAAAGGACACGAGTAAGAATAAAACAATATATAGAGATAATCACAAATGTGATATATGCCCTTGTGATAATCATTTATATGACAAAAACAATTTAGATAATCATTTGAAAAGTATTAATTTTTCACTTATCCCTTCGTGTAATTCTAAAAATATAAAAGCATTAAATTTTAAAGATATCGAAATATATACTGATAAAAGTATTAAGAGAAGAAAAAAACTACCGAGCCCTAGAACAGAAATAAAAATTAGTATGTGGTCTCTTTTAAAAGATTGCATAGGAAAGGATTTATCGCGTATAGGTATGCCCATATATTTAAATGAGCCTTCTTCATTTCTTCAAAGATTAGCTGAAGATTTTCAATATATATATTTATTAAAATATGCGTCAAATGAAAAAGAAAGCACAAGTCGATTAGCTTTTGTAACAGCATTTACTATATCACCATACGCTTCAGTCATAGGAAGAACGTATAAACCATTTAATCCTTTACTTGGGGAAACATATGAATTGACACATAGAAAGTTTCGTTTTATATCAGAGCAGGTGGTTCATCACCCTCCTATAACAGCATATCATTGTCATAATGAATATATGGAAAATTATGCAAGTATTATGGCTAATGTTCATATATTGGGAAAATCCGTTGAGGTCACGATGCCTGGATCAAGTCATCTGATCCTTAAGTACAAGAAAACAATGATGTCTCAAAATAATTTGGAAAAAAAAATAAATAATTATATTCATAATAATATTCATAATAATATTCATAATAATAAGGTGAAGGAGGGAAATACAGAATCAAACATGATTAATAATTATAATTTTTGTGGAAAAAAGAAACATACACATGATACATATATACACACAGATGGAATGTCTAATGATATAAATTCAAAAAATGATACAAGCGAACAAATTCAAAATGAATTATTAAATGATAATATGCAAAGTGAGGAGAACGGAAATAATTGTATGCCCCCTATAAGGAAAGAATTTTATAAGGACAGAGGAACATG**TAAGAACACAAAAGATGGTGATAATAATGATGGTCATAATAATGATGATAATAATGATGATGATAATAATGATGATGATAATAATGATGATGATAATAATGATGGTCATAATAATGATGATAATAATGATGATGATAATGATGGTCATAATAATGGTCATAATAATTGTCATAATAATGGTCATAATAATGATGACCATTTATTATATGATCAAGAACACTACACATATGAAAGAGCAAATATGATTATCCACAATGTAATATTTGGAAAGCTATGGGTTGAACTCCACGGGAATATAATAATACGTAATCATAATAATGGAGACTTTTCAATCGTTAAATATATAAGAAAAGGATGGTTTGAAAAAGAAATTCATAATGTTAGAGGTGTAGTTTGTGATAGGTATAAAAACATAATTTTTTTTATTTATGGGAAATGGTCACAAGAAATATATATTGCATATGTAAAAGATATGAAGAAATTAGAATATGATAATTATTTTTTTAATGAAGATGGTAGTGAAAATATTTATCATTATAATAAAAATACATTGAATGAATTTATTAATACTATTGATTGGCAACTATATGAAAATAATATAGAAAATTTAAATTCTATATGTGTATGGAAAGCACATAAAAGACCTCACAACTCAGAACAATATTATGGTTTCAATTATATGACTGTTGAATTAAACGAAATAACAAAAGAATATGATAAAGATAAAGGAGCAGCTATCGCCTGTACAGATAGTAGATTTAGACCAGATCAAAGATACTATGAAAATGGTAATATAGAGGTTGCTATGAATGAAAAACAAAGATTAGAAAATAAACAAAGACAAAACGCAAAAAAATTTAATAATACAAATAAATATGAACCTAAATGGTTTTACAAACATAAAGATCCTATATTTAATGATAGAGATATCTATTTGTTTAATAATAAATATTGGATAACCAAAGATAAAAATCAATTCTCCGACACACCCGATATATTT**TAAaatattataaaagattaaaaatgtattaataataattatgaacatatcataagattcaagactttttcttttatgcacataataataataacaagaagaatatatatatatatatatatatatatatatatatatacatacacattttatgcattatatttgtttttttaaagaaaatattatttgttttgtttttttataatatcattataatcaaagtcaaaaaaaaaaatttttatatatcacatttgttaaaccatagtttatccaaaaaaaaaaaaaaaaaaaaagaaagatatatatatataaatatatatatatttatatataattattttattttattttattttttttttttttttatggttcttttgttcctataaaatttttaaaacgaaatttatagcatattgtgcatgcctttattttattattatttttcataatattttgtacaatgtctggaaaaagaaaaatccacatgcttgtaacatcac

**Original Locus (OL) amplicon size: 1484bp**s

**> PF3D7_1131800-HALO-SW Halo Neo-R**

tatatatatatatatatatatattgaggtctattatatacaactacataatatatatatatatatatatatatatatattatatatgtgtgtttacttatttgtttttttatttttctaattatataatttgttgtttattaaaaaaagaaaaaaatcataataaataagtaaaaagaaaaagaagaattaattagcatgtgttatctttttttattttttaaaatgaaagcaaaataataaaagagtataatataattatttaacacaatatatatatatatatatatatatatatgtcgcattaaatttatgtacatttcatatatattattcatattaatatttttatttgtatcttcattttaattttatattatatatatatataatatatatataatataatataatatatataagaaaaaattgaatcatgaatacattataagtgctgaaaaaaaaaaaaaaaaaaaaaaaaaaaacgataaaaagaaaattATGATAGGTGGTAAGTATTTAAATATAAAAATTTTTGGAGAGaAAACGTAATTCATTAGGTAGCAATAAAACTTCAGATAATACGATAAATAATAATAATGTAAATTATATAAAAAGGAAAAAAAATAATAATAAGAAAGATCAAGAAAAAAAATATAATGATAATAATTATGAAGAAGAATATATAAAGAAAAGAAAATCAAAACCAGGATCATTATATAATAATATAGACAAAAGTAAAATTTATCGAGATAAAAAAATTATACATGAAGGATGGTTAAATAAATGGACTAATATTATTGGAAGTTATAGACCTAGATATTTTATTTTAGAAAATGGATTATTAAGATATTCACTTGATAAATATTCTCCTACGAAAGAGTCTTTTGTATTAACACATTGTAAAATTAAAGTATGTCCAGATGATCAATTACATTTTGAAATCGATACAAACGAACAAGGTGTGTTATATTTAAAAGCAAACTCACCAGAAGATAAACATAAATGGTATATCTCTTTTAAAAAAGCACAACTAAATTATATTAATGGTAATAATTATAAAAACAAAAATGGACATCATAATATACCTGATGCAGTACAAACATTTAATATGTCTAATAATTCTTTATTTTTAAAAAATATTATTAAAAATTCATCCAAATGTATTAAAAGGGAAGATCAACATTTGGATGATCCTATAAAGAACACTAACAATCAAACAAAAAAGGAGTTGTCTAATAATAATAAAGAAATAATAGAAAAAGAAAAAAACGTTGATTGTATTATAGTTAATAAAATTCAGGAAAATGTAAAAGGGGACGAATCTAATAATATTGTAAAAAATGTGATATACGAAAGGGATGATAAAGATGTGCCATATGAAAGAAAAGATATATGTATGTCACATGATAAAAATTATAGAAGTCCTTTATATATTGACAATGATAATATGTCATCATATAGTAAGATAAATAAAAATGCAAATATTAAATATAAATCAAATGAATATATTTTTTATAAAGATGATAATGTTGAAAAATTATCATCATCATCATCATCCACCTTGGATACATCTGAATTTATTGAAAAAATGAATAGTTATAATAAAAAATATAATCTAGAAGATATGTTTATTAGTTCAACAGATTTTGAAGATAAAAGTCCAACCTTATGTTTAATGAAGAATATAATATCTCTCAAGGAAATGACAACGGATGTTTTAAAGGGTTCTGAGTATTATCAAGCCAGATCTATATTAAATAAAAAATTAAAAAATAATATTAAAAAGGGTGTTAATAGTAATATTATTGTTGATGTTAATAATAATATTATTGTTGATGTTAATAATAACATTATTGGTGATGTTAATAATAATATTATTGTTGATGTTAATAATAACATTATTGGTGATGTTAATAACAACATTATTGGTGATGTTAATAACAACATGATTGGTGATGTTAATAATAATATTATTGTTGATGTTAATAACAACATGATTGGTGATGTTAATAACAACATGATTGGTGATGTTAATAACAACATGATTGGTGATGCTAATGATTATGTTAATAACAATTTTAGTAATGGTAATAATTTACAAATAAAAAGTGATGAATTACTTTCCTTATTATCACAACTATACTTTTCCATACAATATGCTCATATTATTATAGAGAAATATATACAATGTACCGAATTATTATTAAAGGAAGAATCCATACATTCTAAATATATGAATCAGTCTTTAAAACTTTTAGCCAAACAAAATTATTATCTTGAAAAATCCCAAGAAATTAATAATCTTACTCATATGGAAGAAGAAATAAAAGATAAATTAAAACACTTTCAACTTTATCAATGTGCTACTTTAAGTGAAGAGGACGAAGAAGAAGAAGATAAAGAAATGTATAATTCATCAAAACAAATAAAACAGAAAAAACAACAGCAAGAGGATTTATTTTATGATTGTGATGAATTTATTTATGATAAAAATATATTATCAAATGACGTTTTTGAAGACGAATCCAAAAACTCTATCAGCTTATATAGTAGTGAGGGAGATAAGAATGAAAACTCAGATAAAATAAAAAAGGTGCATAATACAAATAACATACCAGATCAAAATCGTACCTATAATAATGAACACATGTTAACAACACATAGTACTATTAATAATAAAAATGAAAAGATGAATGCTCATATAGATGAACATTATTATAATTATATTAATAATGATCAGGAAAATGAAAAAACTTTTTCACATCAAACATCAGAGCAAAGAAGTTTAAATATAAAAACAAAATCTAGTGATCATGATGATAATATAACTACCATTCAAGAGGATTGTACAACAGATGTTTTTCTGAAAACAATTTTGAATAATAAAAATATTAATGAAACAAAAAGGAATACGGTGAGTACAGAGGATGATATTGCTCTTGGCGATGTGAGGGACGGTTTAATAACATGCGATGTTTATAAAAAGGATCATTTTGTGTGTGATGACAGTGTGGAGGATAAGGATATAATACGCAAACGAATGGGGATTGAAGATGATCATGGCAATAATGAAGATGTTCATGGCAATAATGAAGATGTTCATGGCAATAATGAAGATGTTCATGGCAATAATGAAGATGTTCATGGCAATAATAAAGATGTTCATAACAATAATAAAGATGATCATAACAATAATGATAATATTTGTGTAAAGGACACGAGTAAGAATAAAACAATATATAGAGATAATCACAAATGTGATATATGCCCTTGTGATAATCATTTATATGACAAAAACAATTTAGATAATCATTTGAAAAGTATTAATTTTTCACTTATCCCTTCGTGTAATTCTAAAAATATAAAAGCATTAAATTTTAAAGATATCGAAATATATACTGATAAAAGTATTAAGAGAAGAAAAAAACTACCGAGCCCTAGAACAGAAATAAAAATTAGTATGTGGTCTCTTTTAAAAGATTGCATAGGAAAGGATTTATCGCGTATAGGTATGCCCATATATTTAAATGAGCCTTCTTCATTTCTTCAAAGATTAGCTGAAGATTTTCAATATATATATTTATTAAAATATGCGTCAAATGAAAAAGAAAGCACAAGTCGATTAGCTTTTGTAACAGCATTTACTATATCACCATACGCTTCAGTCATAGGAAGAACGTATAAACCATTTAATCCTTTACTTGGGGAAACATATGAATTGACACATAGAAAGTTTCGTTTTATATCAGAGCAGGTGGTTCATCACCCTCCTATAACAGCATATCATTGTCATAATGAATATATGGAAAATTATGCAAGTATTATGGCTAATGTTCATATATTGGGAAAATCCGTTGAGGTCACGATGCCTGGATCAAGTCATCTGATCCTTAAGTACAAGAAAACAATGATGTCTCAAAATAATTTGGAAAAAAAAATAAATAATTATATTCATAATAATATTCATAATAATATTCATAATAATAAGGTGAAGGAGGGAAATACAGAATCAAACATGATTAATAATTATAATTTTTGTGGAAAAAAGAAACATACACATGATACATATATACACACAGATGGAATGTCTAATGATATAAATTCAAAAAATGATACAAGCGAACAAATTCAAAATGAATTATTAAATGATAATATGCAAAGTGAGGAGAACGGAAATAATTGTATGCCCCCTATAAGGAAAGAATTTTATAAGGACAGAGGAACATG**TAAGAACACAAAAGATGGTGATAATAATGATGGTCATAATAATGATGATAATAATGATGATGATAATAATGATGATGATAATAATGATGATGATAATAATGATGGTCATAATAATGATGATAATAATGATGATGATAATGATGGTCATAATAATGGTCATAATAATTGTCATAATAATGGTCATAATAATGATGACCATTTATTATATGATCAAGAACACTACACATATGAAAGAGCAAATATGATTATCCACAATGTAATATTTGGAAAGCTATGGGTTGAACTCCACGGGAATATAATAATACGTAATCATAATAATGGAGACTTTTCAATCGTTAAATATATAAGAAAAGGATGGTTTGAAAAAGAAATTCATAATGTTAGAGGTGTAGTTTGTGATAGGTATAAAAACATAATTTTTTTTATTTATGGGAAATGGTCACAAGAAATATATATTGCATATGTAAAAGATATGAAGAAATTAGAATATGATAATTATTTTTTTAATGAAGATGGTAGTGAAAATATTTATCATTATAATAAAAATACATTGAATGAATTTATTAATACTATTGATTGGCAACTATATGAAAATAATATAGAAAATTTAAATTCTATATGTGTATGGAAAGCACATAAAAGACCTCACAACTCAGAACAATATTATGGTTTCAATTATATGACTGTTGAATTAAACGAAATAACAAAAGAATATGATAAAGATAAAGGAGCAGCTATCGCCTGTACAGATAGTAGATTTAGACCAGATCAAAGATACTATGAAAATGGTAATATAGAGGTTGCTATGAATGAAAAACAAAGATTAGAAAATAAACAAAGACAAAACGCAAAAAAATTTAATAATACAAATAAATATGAACCTAAATGGTTTTACAAACATAAAGATCCTATATTTAATGATAGAGATATCTATTTGTTTAATAATAAATATTGGATAACCAAAGATAAAAATCAATTCTCCGACACACCCGATATATTT**CAAGGATCGGGTTCCGCTTCGcctaggTCAGGATTGAGATCAAGATCTGCTGCTGCTGGTGCTGGTGGTGCTGCTAGAGCTGCTctgcagAGAGGAGTACAAGTTGAAACAATATCACCAGGAGATGGTCGTACATTTCCAAAAAGAGGTCAAACTTGTGTTGTACATTATACTGGAATGCTTGAAGATGGAAAGAAATTTGATTCATCTCGTGATAGAAATAAACCATTTAAATTTATGCTAGGTAAACAAGAAGTAATACGAGGTTGGGAAGAAGGAGTTGCTCAAATGAGTGTAGGTCAAAGAGCAAAACTTACTATATCTCCAGATTATGCTTATGGTGCAACTGGACATCCAGGTATAATTCCACCTCATGCAACTCTTGTATTTGATGTGGAGCTTCTAAAACTAGAAACTAGAGGTGTTCAGGTTGAAACAATTTCACCTGGAGATGGCAGAACCTTTCCTAAAAGAGGACAGACTTGCGTAGTTCATTATACAGGCATGCTAGAGGATGGTAAGAAATTTGATTCTAGTCGAGATAGAAATAAGCCATTCAAGTTTATGCTAGGTAAACAGGAAGTAATAAGAGGTTGGGAAGAGGGTGTAGCACAGATGTCAGTTGGACAAAGAGCAAAGTTAACAATATCACCAGATTATGCATACGGTGCAACAGGCCATCCTGGCATCATCCCTCCACATGCAACTTTAGTATTCGACGTTGAATTGTTAAAGTTAGAGACAACGCGTGCTAGAGGTGCTGCTGCTGGTGCTGGAGGTGCAGGTAGACGTACGATGGCAGAAATTGGTACGGGTTTTCCATTTGATCCTCATTATGTGGAGGTGCTGGGGGAAAGGATGCATTATGTTGACGTAGGACCAAGAGATGGTACTCCAGTGTTATTTTTGCATGGAAACCCAACCTCGAGTTATGTATGGAGAAATATAATTCCACATGTAGCACCAACACATAGATGTATAGCTCCTGATTTAATTGGTATGGGAAAAAGTGATAAACCTGACTTAGGATATTTTTTTGATGATCATGTCCGTTTTATGGATGCTTTCATTGAAGCTCTTGGCCTTGAAGAAGTAGTATTAGTTATACATGATTGGGGATCCGCCTTAGGATTTCATTGGGCCAAGAGGAATCCTGAAAGAGTAAAAGGAATAGCATTCATGGAATTCATACGACCAATCCCCACATGGGATGAATGGCCAGAATTTGCACGCGAAACATTTCAAGCTTTTAGAACTACAGATGTTGGTAGAAAATTAATAATAGATCAAAATGTATTTATAGAAGGAACTTTACCTATGGGTGTTGTAAGGCCGTTAACAGAAGTTGAAATGGACCACTACCGTGAACCTTTTTTAAATCCAGTAGATAGAGAGCCCTTATGGAGATTTCCTAATGAATTACCTATTGCAGGTGAACCCGCGAATATTGTTGCTTTAGTAGAAGAATATATGGATTGGTTACATCAGTCTCCTGTTCCTAAACTTCTATTTTGGGGTACACCTGGAGTTCTAATACCACCAGCTGAAGCAGCAAGATTAGCAAAATCATTACCAAATTGTAAAGCTGTTGATATAGGTCCTGGGTTGAATTTATTACAAGAAGATAATCCAGATTTGATTGGATCTGAGATAGCTAGATGGCTAAGTACATTAGAAATTTCAGGTACCGGTGCCAGGGGAGCAGCCGCAGGAGCAGGGGGGGCAGGAAGGCGTGGTGTTCAGGTCGAGACTATTAGCCCTGGAGATGGACGCACGTTTCCTAAGCGTGGACAGACATGCGTAGTTCACTACACAGGTATGTTGGAGGACGGTAAAAAGTTCGACAGCTCACGCGACCGCAATAAACCTTTCAAGTTTATGCTTGGCAAGCAGGAGGTTATTCGTGGATGGGAGGAGGGTGTAGCACAGATGTCTGTTGGACAGCGTGCTAAGTTGACAATTTCACCTGACTATGCTTATGGCGCTACGGGCCATCCCGGGATCATTCCGCCACATGCGACTCTGGTATTCGACGTTGAATTATTAAAGTTAGAGACAGCTAGAGGGGCCGCTGCAGGTGCTGGTGGAGCTGGAAGACGTGGAGTACAAGTAGAGACTATCTCTCCAGGTGACGGTCGCACTTTCCCAAAGCGTGGCCAAACCTGTGTTGTACATTACACTGGTATGCTGGAGGATGGGAAAAAGTTCGATTCCAGTCGCGACCGTAACAAACCGTTCAAATTCATGTTGGGAAAGCAGGAAGTGATCCGCGGGTGGGAGGAAGGCGTGGCGCAAATGAGCGTCGGTCAGCGGGCTAAATTGACCATTTCCCCTGACTACGCGTATGGGGCTACTGGGCACCCAGGGATTATTCCGCCTCACGCTACACTTGTGTTTGATGTCGAACTTTTGAAACTGGAAACTGTCGACGGAGAAGGAAGAGGAAGTTTATTAACATGTGGAGATGTAGAAGAAAATCCAGGACCAATGATTGAACAAGATGGATTGCACGCAGGTTCTCCGGCCGCTTGGGTGGAGAGGCTATTCGGCTATGACTGGGCACAACAGACAATCGGCTGCTCTGATGCCGCCGTGTTCCGGCTGTCAGCGCAGGGGCGCCCGGTTCTTTTTGTCAAGACCGACCTGTCCGGTGCCCTGAATGAACTGCAGGACGAGGCAGCGCGGCTATCGTGGCTGGCCACGACGGGCGTTCCTTGCGCAGCTGTGCTCGACGTTGTCACTGAAGCGGGAAGGGACTGGCTGCTATTGGGCGAAGTGCCGGGGCAGGATCTCCTGTCATCTCACCTTGCTCCTGCCGAGAAAGTATCCATCATGGCTGATGCAATGCGGCGGCTGCATACGCTTGATCCGGCTACCTGCCCATTCGACCACCAAGCGAAACATCGCATCGAGCGAGCACGTACTCGGATGGAAGCCGGTCTTGTCGATCAGGATGATCTGGACGAAGAGCATCAGGGGCTCGCGCCAGCCGAACTGTTCGCCAGGCTCAAGGCGCGCATGCCCGACGGCGAGGATCTCGTCGTGACCCATGGCGATGCCTGCTTGCCGAATATCATGGTGGAAAATGGCCGCTTTTCTGGATTCATCGACTGTGGCCGGCTGGGTGTGGCGGACCGCTATCAGGACATAGCGTTGGCTACCCGTGATATTGCTGAAGAGCTTGGCGGCGAATGGGCTGACCGCTTCCTCGTGCTTTACGGTATCGCCGCTCCCGATTCGCAGCGCATCGCCTTCTATCGCCTTCTTGACGAGTTCTTCTAActcgagggatatggcagcttaatgttcgtttttcttatttatatatttataccaattgattgtatttataactgtaaaaatgtgtatgttgtgtgcatatttttttttgtgcatgcacatgcatgtaaatagctaaaattatgaacattttattttttgttcagaaaaaaaaaactttacacacataaaatggctagtatgaatagccatattttatataaattaaatcctatgaatttatgaccatattaaaaatttagatatttatggaacataatatgtttgaaacaataagacaaaattattattattattattatttttactgttataattatgtgtctccttcaatgattcataaatagttggacttgatttttaaaatgtttataatatgattagcatagttaaataaaaaaagttgaaaaattaaaaaaaaacatataaacacaaatgatggtttttccttcaatttcgatatcaatttatagaaacaaaatatatacttgtataattttatttttttatataaatcattacatatataattatacaatattttttctaagagataattatatattaatatatataaaaaaaggtgttttttttttttttttttatttttatttttattttatggtaatattttattttccttattttataaattatattagtttatatgtgattaattttatatattatcaatttatatatttttaaatgcttacttaattatctttttttttttttttttttttttttcccctctttttatattaatttatttttgaaaaaattgatatatatatatatatataatatatatatatacatgtagtagtattaaacaatgtataatatatataaataatatatttatatatttcatttcaattttaattttttttggttttttttttttttctttttgtcatatttaaaaaaaattatattcatataagttatgcattttttataaacattattcaatatatgtataatataatatatatatatatattaatgtattattccaatgtgcatgataaaagaaaaaaataatatttataaaaaaaaagaaaaataaaacaaaaaaagaaaaaaaaaaaaaaaaaaaaaaaaatacaaaaataaataatataatttataattatatattcttgtcacaataaaaatatatatatatatatatatatttataatatgtatattttaaactagaaaaggaataactaatattttatttattatcattcaagatttatattttataataataaatacctaatagaaatatatcaggatccatgcatggttcgctaaactgcatcgtcgctgtgtcccagaacatgggcatcggcaagaacggggactacccctggccaccgctcaggaacgaatttagatatttccagagaatgaccacaacctcttcagtagaaggtaaacagaatctggtgattatgggtaagaagacctggttctccattcctgagaagaatcgacctttaaagggtagaattaatttagttctcagcagagaactcaaggaacctccacaaggagctcattttctttccagaagtctagatgatgccttaaaacttactgaacaaccagaattagcaaataaagtagacatggtctggatagttggtggcagttctgtttataaggaagccatgaatcacccaggccatcttaaactatttgtgacaaggatcatgcaagactttgaaagtgacacgttttttccagaaattgatttggagaaatataaacttctgccagaatacccaggtgttctctctgatgtccaggaggagaaaggcattaagtacaaatttgaagtatatgagaagaatgattaagcttatttaataatagattaaaaatattataaaaataaaaacataaacacagaaattacaaaaaaaatacatatgaattttttttttgtaatcttccttataaatatagaataatgaatcatataaaacatatcattattcatttatttacatttaaaattattgtttcagtatctttaatttattatgtatatataaaaataacttacaattttattaataaacaatatatgtttattaattcatgttttgtaatttatgggatagcgattttttttactgtctgtatttttcttttttaattatgttttaattgtattttatttttattattgttctttttatagtattattttaaaacaaaatgtattttctaagaacttataataataataaatataaattttaataaaaattatatttatcttttacaatatgaacataaagtacaacattaatatatagcttttaatatttttattcctaatcatgtaaatcttaaatttttctttttaaacatatgttaaatatttatttctcattatatataagaacatatttattaaatctagaattctatagtgagtcgtattacaattcactggccgtcgttttacaacgtcgtgactgggaaaaccctggcgttacccaacttaatcgccttgcagcacatccccctttcgccagctggcgtaatagcgaagaggcccgcaccgatcgcccttcccaacagttgcgcagcctgaatggcgaatggcgcctgatgcggtattttctccttacgcatctgtgcggtatttcacaccgcatatggtgcactctcagtacaatctgctctgatgccgcatagttaagccagccccgacacccgccaacacccgctgacgcgccctgacgggcttgtctgctcccggcatccgcttacagacaagctgtgaccgtctccgggagctgcatgtgtcagaggttttcaccgtcatcaccgaaacgcgcgagacgaaagggcctcgtgatacgcctatttttataggttaatgtcatgataataatggtttcttagacgtcaggtggcacttttcggggaaatgtgcgcggaacccctatttgtttatttttctaaatacattcaaatatgtatccgctcatgagacaataaccctgataaatgcttcaataatattgaaaaaggaagagtatgagtattcaacatttccgtgtcgcccttattcccttttttgcggcattttgccttcctgtttttgctcacccagaaacgctggtgaaagtaaaagatgctgaagatcagttgggtgcacgagtgggttacatcgaactggatctcaacagcggtaagatccttgagagttttcgccccgaagaacgttttccaatgatgagcacttttaaagttctgctatgtggcgcggtattatcccgtattgacgccgggcaagagcaactcggtcgccgcatacactattctcagaatgacttggttgagtactcaccagtcacagaaaagcatcttacggatggcatgacagtaagagaattatgcagtgctgccataaccatgagtgataacactgcggccaacttacttctgacaacgatcggaggaccgaaggagctaaccgcttttttgcacaacatgggggatcatgtaactcgccttgatcgttgggaaccggagctgaatgaagccataccaaacgacgagcgtgacaccacgatgcctgtagcaatgccaacaacgttgcgcaaactattaactggcgaactacttactctagcttcccggcaacaattaatagactggatggaggcggataaagttgcaggaccacttctgcgctcggcccttccggctggctggtttattgctgataaatctggagccggtgagcgtgggtctcgcggtatcattgcagcactggggccagatggtaagccctcccgtatcgtagttatctacacgacggggagtcaggcaactatggatgaacgaaatagacagatcgctgagataggtgcctcactgattaagcattggtaactgtcagaccaagtttactcatatatactttagattgatttaaaacttcatttttaatttaaaaggatctaggtgaagatcctttttgataatctcatgaccaaaatcccttaacgtgagttttcgttccactgagcgtcagaccccgtagaaaagatcaaaggatcttcttgagatcctttttttctgcgcgtaatctgctgcttgcaaacaaaaaaaccaccgctaccagcggtggtttgtttgccggatcaagagctaccaactctttttccgaaggtaactggcttcagcagagcgcagataccaaatactgtccttctagtgtagccgtagttaggccaccacttcaagaactctgtagcaccgcctacatacctcgctctgctaatcctgttaccagtggctgctgccagtggcgataagtcgtgtcttaccgggttggactcaagacgatagttaccggataaggcgcagcggtcgggctgaacggggggttcgtgcacacagcccagcttggagcgaacgacctacaccgaactgagatacctacagcgtgagctatgagaaagcgccacgcttcccgaagggagaaaggcggacaggtatccggtaagcggcagggtcggaacaggagagcgcacgagggagcttccagggggaaacgcctggtatctttatagtcctgtcgggtttcgccacctctgacttgagcgtcgatttttgtgatgctcgtcaggggggcggagcctatcgaaaaacgccagcaacgcggcctttttacggttcctggccttttgctggccttttgctcacatgttctttcctgcgttatcccctgattctgtggataaccgtattaccgcctttgagtgagctgataccgctcgccgcagccgaacgaccgagcgcagcgagtcagtgagcgaggaagcggaagagcgcccaatacgcaaaccgcctctccccgcgcgttggccgattcattaatgcagctggcacgacaggtttcccgactggaaagcgggcagtgagcgcaacgcaattaatgtgagttagctcactcattaggcaccccaggctttacactttatgcttccggctcgtatgttgtgtggaattgtgagcggataacaatttcacacaggaaacagctatgaccatgattacgccaagctatttaggtgacactatagaagcggccgctagaagaacacaaaagatggtgataataatgatggtcataataatgatgataataatgatgatgataataatgatgatgataataatgatgatgataataatgatggtcataataatgatgataataatgatgatgataatgatggtcataataatggtcataataattgtcataataatggtcataataatgatgaccatttattatatgatcaagaacactacacatatgaaagagcaaatatgattatccacaatgtaatatttggaaagctatgggttgaactccacgggaatataataatacgtaatcataataatggagacttttcaatcgttaaatatataagaaaaggatggtttgaaaaagaaattcataatgttagaggtgtagtttgtgataggtataaaaacataattttttttatttatgggaaatggtcacaagaaatatatattgcatatgtaaaagatatgaagaaattagaatatgataattatttttttaatgaagatggtagtgaaaatatttatcattataataaaaatacattgaatgaatttattaatactattgattggcaactatatgaaaataatatagaaaatttaaattctatatgtgtatggaaagcacataaaagacctcacaactcagaacaatattatggtttcaattatatgactgttgaattaaacgaaataacaaaagaatatgataaagataaaggagcagctatcgcctgtacagatagtagatttagaccagatcaaagatactatgaaaatggtaatatagaggttgctatgaatgaaaaacaaagattagaaaataaacaaagacaaaacgcaaaaaaatttaataatacaaataaatatgaacctaaatggttttacaaacataaagatcctatatttaatgatagagatatctatttgtttaataataaatattggataaccaaagataaaaatcaattctccgacacacccgatatatttTAAaatattataaaagattaaaaatgtattaataataattatgaacatatcataagattcaagactttttcttttatgcacataataataataacaagaagaatatatatatatatatatatatatatatatatatatacatacacattttatgcattatatttgtttttttaaagaaaatattatttgttttgtttttttataatatcattataatcaaagtcaaaaaaaaaaatttttatatatcacatttgttaaaccatagtttatccaaaaaaaaaaaaaaaaaaaaagaaagatatatatatataaatatatatatatttatatataattattttattttattttattttttttttttttttatggttcttttgttcctataaaatttttaaaacgaaatttatagcatattgtgcatgcctttattttattattatttttcataatattttgtacaatgtctggaaaaagaaaaatccacatgcttgtaacatcac

**5’ amplicon size: 1210bps**

**3’ amplicon size: 1519bps**

### **> PF3D7_1346400 (PfVPS13L1)**

atattttcttatatattatgtatacataaatattattttaattttatataataatagcct

tatggcaattttaaaatgtagtaataattatattatgtaaaaaaagttcaaaattgataa

acaatgatatgtttttataattatacatttttttagcttttgctattctttttttttttt

ttttttttttttcgttaatttttctattttattccttttttttttttttttttttttgtt

tttgttttcttttttgtttgtttcaaattttttttttattttcttttttgtaattttaat

tttttcaagaattttataaaaatacatgtatcatatgtatttggtatttttatagaaaaa

ttgacaaaaaatatgtactgtatgtatatgtatgcatgtatatatatatatatatatata

tataaggacataatggaaaataaaaaataaaaaagaaagattaataaaaataaatataaa

aaagattaaaaagcataacaATGTTCGAATCACTAGTTGAAAAACTGTTGAATAAATTTT

TGGCTCCATACGTAGAAGGTATAGAACGGAATTTACATTTGGGTGTATGGTCTGGTAATA

TTGTATTAGAGAATTTAAAATTGAAACCACAAATAACTGAAATATTAGACTTGTCATTTA

AGATAATTCATGGGAATATAGGAAGAATAAATATACAGATACCATGGAGCAGTTTAGGAA

AAAACCCTGTGTGCGTTTTAATAAAAAATGTTCACATATATATAAAACCACGATGTTATA

AAAAAAGTGAAGAAGTTATTATTGAAGAATTAAGAAAAGCAAAAATGCATAGATTACAGT

TATTAGAAGAAGAAATTTCTCTTATTAAGTTACAAAAAAATAATGAGAAATCTTCTGAAA

AATCTACACTTATATTTAAATTATTAAATAAAATTATTAATAATATTCAAATAGATATTC

AAGATATACTAATTCATTTTGAGGATCCAGATAAAAACTTCTCTATTGGTTTTATTTTGA

AATCCTCTTCAGTCAAAAATTGTCAAAATAACGACGAGATGTCAACACAAGCAAATTCAA

ATGCAGAAAATAAAACATTGAATCATATAATTGAATTTAAAGGATTATGTATTTATAGTA

ATAGTGATATAAAATCAAGGACAAAGAAAAAAATTAGGAAGAAAAATAAAAAACGAAGTA

AAGATATGGTAGAAATATTGGAAGATGATAAAAATAAAGACAATGTAAAGAATTCATCAA

ATGGTAAGAACATAAGTGAAATGAAATATGAGAAAAGTGAACTTTCATTAAGTAATGAAA

AGTTTCCTAATATGAATAAGACAGATACCGAGGATATGGAAATATTTGATATTTCGTCAA

AAATAAAAGATAGTGTTTATAATAATTTAGATAGTAATGATAATAATATAAATAGTAATG

ATAATAATATAAATAATAATGATAATAATTTAAATAATAATCATAATAAATTAAATAATA

ATGATAATAATATAAATAATAATGATAATAATATAAATAATAATAATAATAATTTAAATA

ATAATGATAATAATTTAAATAATAATAATAACAATTTAAATAATAATGATAATAATTTAA

ATAATAATGATAATAATTTAAATAATAATGACCAAAATAAAGACACACATGATTACTCAA

AATTAAGTAGTTGTGATAGTAGCACCCAAGATGATTATTTAAAAACCACCTTAAATAGAA

TAAGCAGTTTTTTAAAAGAAGAAGAAAATATTATTTTACATAATTATAATTTTGAATATT

TAATAAAACCCTTCGATTTAGTATTACCAGTAGAACAATCAAGTAATAAAAAAGAATTAA

AAGCTAAATTAGAAATAAGTGATAAATGGGAAGGTATTACTCTCACGAAGACCCAAATAA

CGAAAATTATAGAAATAATGAATGAAGCTAATAAATCTAGGAATCAAACAAATAAATTAT

TATTAAAGCATGCATGGACAGTTCGATTAGATATCGAATCGTTAAGAAATGAGACGAAAA

ACGAATTTATAAATCTATATAATAAAGTATTAGGAGAAGAATATAATATAAGCAATACAG

AATTATCTGCACAAGAAATTAATAGATTACAAATTTTATATGATGTTGTGGGTGTTCGAC

ATTTGGCAAAATGGAGATTACACTGTAGAACCACATTAGAAAAATTTATTGAAGAAAAAA

ATTTAAAGAAGAAATTTTTATATGATTCAATATATAAGCAGCAATCTTGGTGGTCTTGGG

TAACAGGCAATAAAAAAGAAATTGAGAATAAAGTACAAAACATTTTAAATAGTGAACAGG

ATATAATAAATGAAAATGAGTTGTTTATGATACAAGAAGCTATGACAAATGATGATAACT

ATGATGTTGTTATGCCATCAAAATATGATTTTCAATTTAAACTAGCTAATTTTTCGATAA

ATGTATATGATGATTGTAAAAAGAGGATAATAGATAATAGAAATGTGAATAATCACCATA

ATAATCATAATAATAATAATAATAATAACAATATATATATTAGGGGGGTAAGTAACAAAA

GTGAAGTAACTAATAATAGACAACATATAAAAGAAGATAAATTAAATTTTAATGATATTA

TAAAAGATGAATTAAGTATAAATAGCAGTTGTGAAACCTTATTGACAAGTTCCATAAATT

TAAGTTCCACTGATTCCAAATTAAATAAAAGACGTGTAAGATATGAACGGGTAAATATCT

TGTCTATAAATTTTTATCAAATATATTCTTCTTTATCTCTACAATCAGTTGTTGATCATA

ATGATCATGATAATTTCCAATGGAAGTTTATTATTGAATTACAAAATTTTATAGCAAAAC

ACAAAAATAAGGTATTTATGGAATTTAGGAATAATAAAAATTCCTTTCCATATAATATGA

ATACATCAGGAAGTCTATATAAAAATTCTATTTTCTATTCATTATTATATAGTCAAACTG

TTTGCGCCTATTTAGAAATTAACCATTTAGTAACAGAAAAGGGTAATACATTATCCACTA

TATTAAGACTGAACCCTTTGGAGCTTTATTTATCTCCATTATTAATTAAAAGTATATTGT

CATTTACATTTCCTCTATTAGATATAATTAATAAAAGTCATAAGACAGCAATGAAAAAGT

ACAAGAGTAAAAATGTGAGTTTATCTGTGGAAAAGCAAAAACAAAAGGAAGAAACTTCAC

CTGGTGTTATTAAAAAGAAGGAATCAATTTCGGGCACAAGAAAAAGAAGTAATAAAAATA

AGAAGAGCCATCATGAAATTGGAAGTATAAAGAAGGATGTAAATAATGAAGAAAATGATA

CGATAAAGCAAGATGATCATAAAAATATGAATGCTGATAGTAATAAAAATAATGATAATA

ATTATGATAATAATGATGAATGTGATGATAATAATGATGAATGTGGTGAATATAATGATG

AAGATGATGAGGACAAAAATAACGATAATGAGGATAATAATAACGATGATGATGAGGAAG

ATGAGGATGATGATGACGAGGATGACGAAGAAGCATTATTACAAATAAAAAAAATGGAAC

AATCCGAATTATTAGAAGGATTAAAAGAAAGAGGAGAAAATGTTTATAATAGAGCTGTTC

AACACCTTCCAGAATTATTTGAGTTTTATATACATATATGTGGTCCCATTTTACATTTTG

ACAATTTAACGAATGGCATAGTGGAATTGCATTTAGGTAATTTGGTAGCAAAAACAGAAC

ACCCATGTACCTATAATAAATTTAATTTAATATTTGAATTTAATGAAACGCAAATAACAT

GCTTAAAAATAGGCTCAGGATATATGAAGAAGGATGAAAAATGTAAAAAGGATATAATTT

TTAATGAGAACCAAAAAGATTTGCATAAATCAACAAAAGCATATAAGAAGATGGATAAAA

GTAATATATATTTAGAAATAAATGAAGAAGGATTATTTTCAAATAATGCAATGGATAATA

CCGAAATGGATGATTTAAAAGATAGTTATTTTAATAGTATAAAGGAAAAGGTATTAACAG

GTGAAAAATTTTATATATTACAACCAATTCCAGTAAAAATCTATGTAGAATATGACTTGA

AAATTTTGAAAACAAATATTATATTAGATGGAATATTTTTTCAAATTAATCCAGATGCTA

TTAGTATTATATTAGCAGTTCCAACATCGATAACTAGATACCTGACTGGAGTTTATTCTA

AAAATAAAAATGAGAATGAATTTAATAAGAACAAGAAGAAAACTTCAGTTGAAAAAAAGG

AGGGAGATATAATGAAAGGAGATGATATTACAAGAGCAAGTATGTCAAATATCGATAGTG

GTGTACAGAAAAGAAATAGTATTACTGATATTATAAAAGGGAATAAAGATAATGATAATG

TGGTAGAAATGAATAATAAAGGTATGCCAAAGGAAGAAGAGTCGTTTTTATATGATATTG

ATTTTTTAATTAAAAATTCTTCTTTTTCAATTAAGAATAAGAAAAATTGTGAAATATTGA

AATATGAAGCATGTGGTATATTTTATAAGAATTATTTACAAAAGAAGAGGAAAATTATTA

AAATGGAAATTGAACAGTTATGGATTTGTGATCCTAGTAATAAGCAACCTATATTTTTTA

CTTTAACGAAGGATATAAATTCAAAGGATATATTTTTATTTAGATCCTTATCTACATATT

TGGAAGAGAAAAGTCATAACAATAATAGTATTGAACAAAAGGAAATTGATATGGAAAGTG

ATAATCATATAAATGTTGGTATAGACGACGTATTAGAAGAAGATCATTTTATGATGGGCA

ACAACAATAATATTAATAATAATAGCAGTCATATTAGGAAGAATAGTTTTAATATAGGGT

CAAATGCTTTTGGTAGTTTAGAAAGATTCGAATCTGCTAAAGTGATGGATGATAATGTAT

ACACAAATGGAAAACATAAAGGTTCAATCAAGAATGTAATGATTGATAATTCTATTATAA

ATAATGAACAATTAATAAACAATAGAGCAGAAGGAGGAAAATATATGAGCACAAATAGAT

TTTCTGCTAGTGGAATAAATCAGTTTAATAAATATGTTGGTATGAATATAGATAGAAAAA

ATGAAGAGACCAAATTGAATGGGAAAAAAGAAAATGATGAAAATAAAAATGTTGAGGAAG

AAGATGATGATTTTATGGATGCTATTGAAGAAAAACAATTATCTATAAGTCTTCAAATTT

TACAAGAACATAATGAAAAGAATATTGAAGAAACTCATATTAATTTTATTATTAGTGATA

TCGAATTACATTGGAAATATAAAACAATAAAACAGATTTTTAAAACAATGAAAGAATATA

AGAAAACGTTACAATATGGTATAGAAAAAGATATGACATATATAAAAAATAAATTAAAAA

ATGAAAAGGATTTAAAAAATTATAAAATGTTCATATCTGAAAATACCTTAAGAAGTGTAC

AAGAAACTTTGAAAAATGTTAAGGATTCATTAAATATTTTAGATATGGAATCTGCAAAGG

ATAAGGATGATATAATAAATAATGTACAAGAAAAAGAAATGGACAAAGATAATGTTAAAA

TGAATGCAGAAGGTATACATCTTATGAATCCTTATTATGCGAGGGACGAAGATTTAAATA

TGATAGCTGATAATAAGCTGCTTTTTAATTTAAGAAATGAAAAAAATAATGAAGAAATAA

ATATTAATAATGCTAACACTGTAAATGATAATAATGGCATTAATAATAATGATGATATTA

ATAATAACGATGATATGATTAATAATGATGATATTATTAATAATCATAATTATAACATTT

TGAATGATAATATTCTTAGTAAGGGACTTAATGATACATTACCCAAATTACACGGTCAAC

AAAATTCATATCCCAAATATTTTTTCAACTGTTTTATAAAAAGTGCTTCCTTAGCCTTCT

GGAAGAAGAAGAAAATTTTTTCTAGAATTCAAGTGAGTAACATATTTTATGAAAATAAAA

TATATATCAATTTTGATCAGAAGATGTTTTTGAATATAGAAAAGGGGATAATGTCAATGA

ATAATAAGAACATTATATCTAATAATATTAACGATTATAAATACGACCTGTTTATTTCTA

AAGATAGGAAAAGATTTGGTTCAAGTGATATCGAGAATGAGGATAGAAATATGTTAAGTC

ATTTAAAATATGGAAAAAAAGAAGGGAACTTAAATGCAATGCAGAATGATGATGATAAAG

AAAAAGAAGAATATACAGCAGAGGATACATTTTCGGAAAGACGAAATAGTCGAGATATAT

TAAATAATATAAATCTAGATAATTTATCTATAGAAATAAAGAAGAAAAAGAAAACAATGC

ATAAAGAATCCAAAGAAGATTTGTTTATTGGTAAGATAAAAGTATATAATGATAAGAGGA

ATTATAATATATGTTTATTATGTGAAATTCCAAGAATGTATTATATTTTTTACCTGAAAG

ATTTAAGATTATTTTTGGAATATTTAGATGATGGTATATTAAATGTTTTTATTAGTAAAT

CATATAAAAAGGTAGTTCAGGCTGCTCAGACCAAATACTTTTTATTTAATTTTACATTTA

TGGATCCTATAGTGATCATACCTGAAGATAAAAATATGATATATAATTATAAGGGAGAGG

TGAAAAATGTTTCTAGATATTTTAAGAGTAGCACTAGTAATAATAACACAAAGGAAAATG

AGGACACAAATGATATTGCAAAAAGAAGCTCAAAGGAATATATTCTTTCCAATGAAAAAA

AAGATACAAAGACTGAAGCGTGTAACAATTATAATAATGTGAATAATAAGAATATACATA

ACACTAATTTTGATTGTGATATGAAATCCAATTATGATAAAAATTCAATTGGTCGTAATA

CATATAATAGTAATAATATACCATACATTGAATCTTATTTACAATTCCATTTAAGCAAAT

TAAAACTTAAAAATTCTTACACTCTAAAATGTACGGAAGAAATAAGAGTAGCACAGAGAA

GACAAATGATATTAAAAAAAAAGAAATATAAAGGAAGAAGTGAAAAGGAAAAAATAAATA

AAAATAAATATGAAAATGAATCTTTTATAAGAAGTGATATTGTATCTAAGGAACAATTAA

GACCATTAGAGCGAAAAAGGAATAGTGTAAATAATGATTTGGTGAAATTAGATTTTACGT

TATATATTGATATATTAGATATAGAAAGTAAAGCTTGTGAAAATGGGGGTAGTAGTAATA

TAGAAGGGGAAATCTTGCATAAGGTAAATTTAGGTTTCTGTTTAATAAATGCTAGGAATG

GTATATTTATATATTTAAATGGAAATGACTTATGTTTAGATTTAACAGTATTTCAATTAG

CTTTTTTGTTGGATATAATAAATGAAAACTTTTGTTATAAGGGATATTTCCCAATGTGCT

TTATATGTGATAAGGATGTGAGTTTAGACAGTATGATAAATTCGGAGTGGTTTAAAAAAG

GATACATGAATTTTAACAATTTTAATAATAGGAAAGAAATGAAAGGAAAAACAAATGACA

ATAAAAAGGAAGATATAATGAACAATATAAATAATATCAATAATATCAACAACAACAACA

ATAATAATATCAATAATATCAACAACAACAATAATAATAATAATAATAATAATAATAATA

AGGTAAATAATTATGAAATTGGAGATGGTGCTATAAATAACAATGTGTATGATAGCGATC

GCACTATTGATAAGGAAATGGATCACGGTAAAAATAAATTGGATATGTCAATGAACCAAG

AAAAAAAAAGTGATTCATTAACAAATGATAAACTAATAATAAAAGATTCAAAAGAGAATG

TAACAAAAAAGACAGGCTTGAAATTATATGTATATATAAATTTTGAAAGTTTAAAAATTA

AAACAGCATTTGATACAAATACACCTGTAGCCATTATAACATTCCAATATATAAGTATGT

CATTTCGTTTAGTTCTCTTAGATTTCTATTATGTTTACTTTTTTGATTTACATGGTAATT

CATTGTATATAGATGATGCTAGGAAAAATTCAATCAATTATTACAAAAGGGTAGCCTACT

GTTGTATTGAAAATGAAAAGAAAAAGTTCAAGAGGGAAAATAAAAAAGCGCATGAGTTGT

ATAAAAATGTATATAATGAAGAAGCATATAATGATACTCTTAATAAGAGTAATGTATATG

AGTATAATAATGTGGAAAGGAATTCGTATCAAATTAATAGAGAAAATAAAATATATGATA

ATAATGGGTCACATTTTTATAATAATAAAGATGTTCAAGATGAATATGATGAAAAAGTAC

AAAAACTACATAATAATAGTAGTAATAATATTTCTGTTACGACACAATCTGATATAAAAG

ATTATAAAAATTTATTAATATATATGTTTGAAAATAATGTATTTTTAAATGAAAAGAAGA

AAAGGAAAAAACAAAAAGGAATAAAAATAAAAATTAATTCTTTCATAGAAGATTTGTTAT

TAAATATTGAACTTGATGATGCGTATATTTGCTTTTTTTTCCTTATTTTTATAGATATAT

ATAAATTTTTAACGACAGGATTTAATTTGAGCACGTTACATTTGTATCCGAAGCCATCAC

CTTATATTGTATCTTACAAAAATGAAAATAAAAAGAAAAAGATAAAGGGTCATGAATATG

TAGAGAATAGTGAAAGTAAAAGTATCCATAGAAAAAAAAAGGTTGTGTGTTCGAAGACGG

AGGAAGAAAAACATATAAAGGAACAGATTAGTATTAGATGTACTAGATCTCAATATAAAA

ATACCTCAAATGAAAATAAAAATAAAAATATTAATAATAATAATAATAATAATAATAATA

ATAGTAATAATAAGAGTAGTATCAACTGTAAGGAAAATACCAGTATAAATATCGAAGAAA

ATGTGTTTAAAAGAGAAAGAATTCATTATGATGAAGAAATTATTGAACAAAATGTTCAAC

TCATCAATGAAGAAGACAATGAAAAACATTGGGTACAAAAAAAGAAGAAAATTATTCGAC

GTAGTAGTGTTTTAGAACAATCAAAATATATAAAAGGGAAATTAAAAGAACAGAAAAATA

AAATAATAACAAATATAAACGAAGTATTAAAATTAGATAATAAACCATTTCATGTAAGTT

TTAAAGTGAATAATGGGAATTTCATTATTTTTACAGATGTCGAAAAAGTATCTCATCCAA

TAATAATGTGGTCAAATAATTTTGTTTTTTCATTTTCTTTATTTAATAAATGTATTGTAT

TTAGAAAAATATATGCAATAGATAGTAAAGTGAAGAGGATTAATTATGTATCGTCATCTC

ATGTACATTATAAGTATGATACTATTAAGAAGAGGAACAATAAAAGCAACCATAACAATA

AAAATAACAATAACAATAAAAATAACAACAACAATAAAAATAACAATAAAAATAACAATA

ACAATAAGGATAATAATAAAAACAACAGTAACGATAATAATAACATCAATATGGACAATA

ACAATAAAAATAACAATAACAAAATATTTGATCCTAAGAATATATCGAACCCACTTCATA

ATTATCATAAAAAGAAAATATTACTATGTGATAATTTGAATGTAATGGGTGAAGCTGTTT

ACGAATCAGTTGATACAAGAACTGAGGATTATAAATTAAAAAGTAGTGTAATAAGAAAAG

ATAACAATTCACCATTTATAACCGTTTTTGAATTAGATATAAATATTGGTAATTTTGATA

TACGTTTATCAAATGACGATGTTGAAATATTATTGAAAGCATCATCCACATTATTTGGTG

ATGTCCCTAGTTCATTTACAAATATAGTAGTTGGCCCAGTTATACCACCGTTAACTTTTA

TACATAAAAAAATAAAGAATAAATTTATGCAAAACAATATTTATTATATAAGTACAAATA

TCAAGGAGGAGACCGAATTGTTTTGTGGATCCGATTTGCAAATAACAAGTATAAATATAT

TAAAGAAGAGAAGGAAAAAAATTATAAGAAAGAAGGAAAAGAAAGATGCAAGAAATAGAA

AGGAAATGGAGAGAAATGATCATAATAAAATAAAGAATATAAGAAATGATGAAATTAATA

TGAATGAAAAGAAAAGTATAGTAGATTGTCATATTGAGGAAAAGAAATTAGATCAGAAAG

AATATAACGAGTTTAGTAATAAGGATGAAATATATACTGATTCTAGTAACATGATGGAAT

ATGTATATTATAAAAAAATATCTGATGTCTCTTCAAGTACTAATACTTTAAGTTATTCCA

ATACATCTTTAAGTGAATTACATGGAGAAAAAAATATATTATATAAGAGAGGTATGAAAT

ATGATGATATATCAGATATAAATAATAACAGTAGTAGTAATAATAACAACATCAATAATA

ATAATAATAATAATCAAACAAATAGTTATCGAGATATAAAAATTAATATAAAGTTACATA

ATGTAATGTGTACCTTTATAGATAATATAAGAAATTCCATTGTTCCCATTTTAAGAATGA

TATTTAGTATGAACATAAGTATTAATTTATATACTGATGAATGTTCATATAATATTAGTG

ACTTAAATTCTAAGTTAGAATATTTTAATAATTGTATAGGTGAATGGGAACCATTTCTTG

AGAAATGCAATATTTCATTAGATATACATAATATTTTTCCAAATGACGAATATGATGAAG

ATAAGGAAGCAAACAAATCTCCTATAAGTATCATTAAAATTAATAGTATCAAAGCTTTGT

GGTTTAATATTACACCTCAGTTAATTAATTTGTTATTTTTATTCGTGCCAGTCTTTTCTG

AGAAGGTATCTAATGGGTTAAGAAAAAATGCCAAAAAGGAAATAGACCATTATAATGATA

AGGATAATAAATATGTAAGTGAATGTATAAAAAATGATATATTAATGGAAGATATGAATAATAAATTAAATGTTAGGAATACATCTATTAATAAGGATATAGAAACATCTGAACAGATGAATAGTCTAAATGACTTTTCCAGATGTGATAGTGGATCCGTCGAATTGGAATACAAAAATATGAGTGTTAAGAGTAAGAGCATATTCGAAGATTGTTCTTCGGTTTTTAATTCATTTCCGTGTGATACAGATGGTAAGGAAGCAAATGATTTTCATTTGGATGACTTTTTAAAAAAAGAAGAAAATCGTAGTACATACTATAATAGGATAGATAACTCAGTAATATATTACGTTAATTTGACAAGCGAATACTTTTATGCCTTTTTAATGCCTGAAAGTAAGATTGAAGAAATTATGAAGAGAAAAAAATCTGTCACTATGAAAAATGTATACAAAAGTGTGAACGAGGCGGATGGGACAAATATGACTATAAAATCTTATAAATGGGATGAGAAAAAGAAAAAGGTAGAAAAGGGATTCATTCAAAGGGAGAATGTTACGAAGGCGAATGGAATGAAAGGAAATGATATAAAACAAAATGGCATAAAACAAAATGAAATAAAAGAAAATATAATGATGGAACATACATTAATTAGTTCAGATGATAATAGTACCCATAATGAAGGGAATGAAGATTATATATCTGAGGAAGATCTTTTCAATATGATACCTGATGTATATGCAAAAATTATAACAACAAATGAATTAATATCATTAGACAGATTGTTAATCAACGAAATAGAAAATAACTCCTTAATAGAAAAAAAATGTTTATATTTATATTTGATACCCATTCCACCAACAAATGTTGTGAATATAGTTCATGATATGTTTTCAAATATTAGTAAAAGAGATATTGTTTTAGATCTCATGATTACAAAAAATCCAAAGAAAACTATTGAAAATTTATTAATTTACAATCAACATAAGGAACATATACGTATTAGGAGAAATAGTATAAATAGCATTTGGACGGAAAATATGATGTATGATGACAATAATAAAAAGAAACATATAGAATTGTATTTGAAAAAAATGCGTATGAAAGCGGAAGAGGATGGTGGTAACAAGGTAGGAACGGATAATATACATTTATTGGAAGATCAAAATATGTATTTATTAAACAAAGAAATGTGGAATGATTATAAAAACGTTAAGGGTATTATAAAAGGGGAAGGAAAATATACTAATAGTACTATTAGTAAAACGGACCATACTTCAAGGAGTAACAATATGGACAATAATGTAAAAGGTAATGATAATATTAATGGTAATCACAGTGATAGAGAATCTGAAAAAAGACAAGACATGAATCAAAATAATGTAAATGAAAATATCTTAAATGTTATGGTAGAAGGAAACAATGAGGTATTAGATAATGAGCAACGGGTTGATAAATCATTGAATTTATATACAAAAGGTTGTGAAAATGATGATGCTAATGAGGATGTCAATGATGAAGAAGAAGGGTCTTATGATAGTTCTTCTGATAGCTGTGAGGATTATGAATCAACTGATGATACCCATTCAATTGATGAAAGGGATAATTTGAATTATGAATATTATTTTAAATATAATAAGAAAAAGAAAAATGATTCAAGTTCTATTTTTTCATCGAATAAATTTGTTGGACCCTTTTTAAGAAACACAGAATGTGTAATAATAAATTTGATTAAGAATAGTTGTACATCATTAACTACATCTTTGAAAAATAGCAACGTTATACATTTATTGAAGCCTGGTGATTTTGTGTTTGATGAAAAATATAAATATAATTTTTCTAGCAAAAAGACTAATGAAATTATGGAAAATTATAGTATTACTGATGGGATTATGGCAGACGATGACATAACTAATGGATCCATAAATAATAATAATAATAATAATATGTGTAACAAAGGTATTTATGATAATACACCAACAAAACAGATGGTCGATAAATATGTTTCTAAAAATAGTATGAGTACTAATAAGAATAATAAATCGAGTATATTAAATAAAGAAAATATTGACAAGATAAATGAATGGGAAAGAAGAAATGAAGAAATGATTAAACAAACATATATAGCATGTAAATATAGATTGAAAAATAATTATTTGCCACAAACATCGTTGGTTGAAAATGTATGTGAAATTATATCTCCTATGCCAAATTATAAAATATTATTTATGTCATCAACCGTTAGAATTATAAACAAATGTGGTATTCCCTTAGAATTTTGTTTTTTTGATGGTAGTCGAAATCCTATATTATTAACGTCATTAGAGAATAGGACTATACCTATAAATACTTTATATCCTAATCATAGTGATTCATTTAAAAATTATAAGGTATCTAATAGTTTGAATATTAATCCGAATATAAAATTAAGACAGAAAAATAATAATTTAAGTTTCACTGTAATATTAAATCATGAGTATTTATTATCAGCACCCGAGTGTGTTTTTTGTGGACCTTCTCATGTGTACATGTCTTTTAAACCTATCAATGTGATAACAGCAGAGAATGAATATTATGACATAATAGCTGGTGGTCCCACCACAAATATTGCGAACAATAATAATAATAATAATAATAATAATAGTAGTAGTAGTAGTCATAATTATTATGGTAATAAATCAAATATTAATGATTTATTAAAAAAGGACAAGGGGGGAAGCAATGATTTAATAGATCCTATGTCTATTAATATAGAAAATGGATGGTCTGATATTTTTAGTTCTGATATATCTCAAGGGACGTATGTGAAAAAGTGTAAATACAAAGATATGAATTCCTTCATGTCTAATATAAATAAAAGTAACGGTAGTGGAAATAATTTTTTTTATTTCTTGGTAAAGATTGAAAATAAGATAAGTGCTTTACCTGCTGAGAAAAATTTGAAAATTATTACTATATATCCTCATGTAAGTGTTGTAAATGCTATACCAGCCTTTGTAGACATAATAATAACATCTGACGTAGTAGAAGAAAAGCAGAATGATTATATGTATGAAGAAAGGAATAAAATAGATGCATTAAAAAATAAGAGAGCAAAGTTATTAAAACAGTGTGAGGATATAAATTCATATTTGAATAAGATGAATTCATCAAAATATGGAGGAAGTAAAATTTATAGTGATAGTTCATCTAGAGTAAAAGGTAACAAAGATAATAGTAATAGTAGTAATAGCAATAATAGTGAACATGTATACATGCAAGAGTTGAACAAATTAGAAAGAGAAATATATCATATAGATGAAGAAATAAAATATAAGAAAAAAGAATTATCTTCTCCATTAGTAAAAGGATATGTGAATAGAAGATTAAATCCATTTTCTATATTTTATATATATGAAATTAAGAAATATACTTGTTTGAATTTAAAGATGAAAATAGGGAATTCACAATGTGAATGGTCTGAAAAGTTATTTTTAGTAGAAGATGATGAAGAATCTGTAACTAGATTTTCATTACATTTTAAAAAATATGCATCAGTCGAAGTTGAAATTATTAAAAACTTTAGTGGATATTTTAATTCTTTGAGTAGTATCTTAGGAAATAAACAATTATATATTTTTTCCTTACCAAGATGTTTTATAGATAGAACAGGTTTAGGAATTAAGGCAATAAATTCGAATAAATATTATCCTGTGATTAATGGAATAACCTTATTAGGGGATCATTCCCAGATTGATTTGTTATTACCTCACAAAACTTATACTCAACCGAACAATAATGTGAACAGTGGTAATAATAATGTTGAAGATGATACGTATAATTATTATTCTAAGGGTGATATGAAGAGATCGAATAGGATGTATGATAAACGAGGAGATCATAGAAAAGATTATAACGATACAAACGAAAGTTATGGTAATGTAGATGATGTTAATAAAAATGATAGTCGTTATGTTAGTAAAAATGCAATTATAATAGATTACAATGATATATTTAATGATTTTAATTACCCTTCACCTGTAAATAATTCTGTAATATTATTCAAAGCTACATTACCTCCGATTGGTAGTTATACAGAAACCAATGTGATTTGTAAAAATTTTTTTTATACCTTTTGTTTAAATACAGAAAAAATAAAAACAACGAATATTCCATATATTATAAGTAGAATAATTACAGTGGTTCCACAATTTATTATTAGTAATAAATTGAATTTTCCCTTATTAATTAAACAATATCAGAATGACCAGATGCAAGGAGTTCGTGCAAATGATACATCTCCTTTATATTTTACCAAAAAAAGTAGCATTTTACTATTTCAATTTAAATGTTTAGATCCATGTAGAACTAATAATGAGGAGAAGAACCATATGTTATTAGATCCGAAAAATGTAATAAAGAGTAATGGAAATAAATATTATAAGTTGAATAAAAATATATATTGGTCGTCTGTTATATATCCTAGTGAAAATTTTGTTGGAACAAATTATATGGTAATTAATACAGGAAATCACGAAAAATCTGATGTATACGCGATAACGTGTATACCTGATAGAGGTACGAAAAATATTATCATAGAAAAATTAGAAAATAAGAAGAAAGGTTTTATAGCATATAATAATTCAAGTATTGCAAAATATTTAAAGATTAGAACATTCCATGATGATACGAGACATATGAAAATTCACGATGAAGAGAATTATATGGAAAAGAATATGTTTTTAAGTAATTTTTTAAAACCAACAGATATGGAACATTATTTTAATATAGAGCCAAATAAATATAGTTATTTGGGATGGGTAAATCCATTTATATATGTTACTAGAAATGTACAGATAGAAATAGTTTTGGAAGATTTAAAAATAATACCTAAATCTCCATTTGTTTTAAAATTTGCTATATATAATTATTCACAAAAAACATATTATATAAATTATTATAATATTACATTTGTTATATGTATAGAATATATAGAAGATTTGATAACCATAAAATTGAGCCATAAATTAAATATAAGCTCAAATCTATTAGATCGTTCCTCTTATTATTATATGATGAATAATAAAAGTAATATAACAGATGAAATAGCATATTCTAATAGTTCACATAATAATATTATGTTCAGCAAACATATGAATAGTTGTGACAAATCTGTTACATCTAATAATGACATTAAATTAGGATTAAATAAAAAGAAGAAATATATGGAAATGGAAAATCAGGTAAATATAGAAAAATATAATATAATAAATGGTAAAGATATTTCAAATGAAAGGAATATTATAGATAAGAATTCAGAGAATAGTTTAAGTTTATCAAAAAATAGTGTTTATGCCCAAAAACAAATGATTATAGATCATGATGAATATCATTATGTAAAATATATAGATGAGAAGGATGGGGGTACTGGTAGTTATAAAAATAATAGTATTAGTAGTAGTACTAATAATTATTATAAGAATGAATCTGTGAAGATGATAAGTAATACATATAAAAATGTGCATATAATAATAAATGTAACCCAGATTGGTGTAAGTATTATATCTAATATATTAAAAGAGGAAGTATTCTTTATTGAACTATCAAAATTGTGTGCTTTGTTTTATATGAAAAATGAAGAAGAGGTAATAGATATAAAAATAACAGATGTACAAATAGATTGTCAGTTGGAATCTTGTGAAAAGTGTGTGTTGTTAGCGAATAGGGGAATAAGTTCTTCAAATAATAATAATAGTAGTAGTAGTAGTAATAATATGAATAATATGATAAGTGGAGGGGATATGAGTGGTAGTAATAATGTATATCAGAATTATCGTAATGAAAATAAAAGTTTTACAATTAATGAGAGTGACCTCCTTCGTTTTAAGAAAACATTTGATCGTAATTCAAACGACAAAAATAATCATATGAATCCTCTATATGATAATAATAAGAATAAAGGTACTACTACTACTACTACTACTAGTGGTAGTAATAATAATAACAATTATAACAATAATAGTAGTAGTGTGTTAAATAAAAATATGAATGCAAATGAAGAAAAGATCTTTTTAAATATATATGTAGAAAGGTCTTTTATATCACATAATGATGTTATATTTAAAAAAATACAAGTTTCGTTAGATGATGTAGAAATAGAAATGGATGCTGAAACTTTGAATGGTATAAATTTATTAACAGCAGAATATATTGAAAGTATAAGTATTGTTCAAAAAAAAAATTTGTTATATGAAGAAATACAGAAATGGACGATTTTACCTGTATATGTAAATTACAAATCACCAGAAATACCATTAGCCATAAATATCCAATACATGCAAATAGATAAATTTACCTTAATAGTGTGGTGTTCATTTTTATTAGATAAAATGCACATGATGAGTGATTTGTTACGAATAGGATTACGAATATTAATGGTATCGGGAAAATTAGAATTATTAGGTGCACCTGTAACCTTAAATCAAGAAATTTTTAATAATATTCGT**GTAAGTATTAAATCCTTTTATGCATTACTAAAAGATAAGTATAGTCATTCTATTTTGGCATGTTTAGGTTTTATAGTAGGATATAGTAGTTTAATAAATATACCTAAGATACCATTAGAAATTGGTCGAAATACTATAGGTTTAGCTGTATATGCAGTAGATAATGTGAGTGTTGGAATTGGAAGCTTTTTATCAAATTTAACATTTGATTCCGAATATATTAATAGAAGACAAAAGGAGAGAACCTTTAAGACGAACACCAATATGAAAGAAGGTTTAATTAGTGCTGTTAAAAATATTGGTGAAGGAGTATTAAGTTTATCTAATATTGTAACGAAGCCAATTGAAGGGGCACAAAAAGAAGGATTTGGTGGATTTTTCAAAGGAATAGGAAAAGGTGTTGCAGGTTCTTTAGTAAAACCATTAGATAAAGTAGGACAAGCTGTATCTGATGTAACTCGTGGAATAAAAGCAGAAGTATCGAAACCTATAGGTGGTCATAAATATAAGACTAAAAGACACAGAAAACCAAGAATGCTATGGGGAGAATATGGAAAATTAAAAGAATATAATATAAACGAAGCAGAATTAAGAGAATGCTTAGGTTTGAAATTTTCAAAAAATATTATGAAATGTTTAACAGTACATAAACAAGAAAATCATCCTCCATCACATTATGCATTATTATTATATCCAAAGGTAATTATATATGCAAATCTATATGCTAATACTAATGCTCAAAAGAATGATTTCTTATTATCCGAAAAAAAAACAGATATAGTTATATGGTCTATAAAAATAGAGGATATAACAGAAATACGAGCATCTAGTCATGGACTCATAGTAAGAACGAATACAAGTACAAATACGGTTTATAAAATACCATGTAATAATGCTATATTGATAAATAAAATATATAGGGAATTGCATAACTCAAAAAATTCAATAAACTCTACTGTTATTTTGGGCGCCAATAATTCTGCCCTGTACAAATATTTA**TAAtaagaaaaaataaaatggcatatatatatatatatatatatatatatataatatatgttcatttatatgtttatttgtttatatatttatttacttatttattcatttatttatttattcatttattcatttatttattcatttatttattttatttattaatttattaatttatttattcattcatttatttattcatttatttattcatttatagttttattttcattatagcatatttccatattttttatttttatatatttaaacttgttttgattaaatatttaaaacttcttcaattctaatatatccattatttatctaattatatatagagagtgtatttttttttttttttttttttttttcttttttccagtttgccttcataaaaacaggcctatttaaataaaataaaaaatataaactatatatttattatatatacttttctatatgataaaatttgtttttaaatatatattattatgatattt

**Original Locus (OL) amplicon size: 1451bp**s

**> PF3D7_1346400-GFP-SW GFP Neo-R**

ttttttttttttgtttttgttttcttttttgtttgtttcaaattttttttttattttcttttttgtaattttaattttttcaagaattttataaaaatacatgtatcatatgtatttggtatttttatagaaaaattgacaaaaaatatgtactgtatgtatatgtatgcatgtatatatatatatatatatatatataaggacataatggaaaataaaaaataaaaaagaaagattaataaaaataaatataaaaaagattaaaaagcataacaATGTTCGAATCACTAGTTGAAAAACTGTTGAATAAATTTTTGGCTCCATACGTAGAAGGTATAGAACGGAATTTACATTTGGGTGTATGGTCTGGTAATATTGTATTAGAGAATTTAAAATTGAAACCACAAATAACTGAAATATTAGACTTGTCATTTAAGATAATTCATGGGAATATAGGAAGAATAAATATACAGATACCATGGAGCAGTTTAGGAAAAAACCCTGTGTGCGTTTTAATAAAAAATGTTCACATATATATAAAACCACGATGTTATAAAAAAAGTGAAGAAGTTATTATTGAAGAATTAAGAAAAGCAAAAATGCATAGATTACAGTTATTAGAAGAAGAAATTTCTCTTATTAAGTTACAAAAAAATAATGAGAAATCTTCTGAAAAATCTACACTTATATTTAAATTATTAAATAAAATTATTAATAATATTCAAATAGATATTCAAGATATACTAATTCATTTTGAGGATCCAGATAAAAACTTCTCTATTGGTTTTATTTTGAAATCCTCTTCAGTCAAAAATTGTCAAAATAACGACGAGATGTCAACACAAGCAAATTCAAATGCAGAAAATAAAACATTGAATCATATAATTGAATTTAAAGGATTATGTATTTATAGTAATAGTGATATAAAATCAAGGACAAAGAAAAAAATTAGGAAGAAAAATAAAAAACGAAGTAAAGATATGGTAGAAATATTGGAAGATGATAAAAATAAAGACAATGTAAAGAATTCATCAAATGGTAAGAACATAAGTGAAATGAAATATGAGAAAAGTGAACTTTCATTAAGTAATGAAAAGTTTCCTAATATGAATAAGACAGATACCGAGGATATGGAAATATTTGATATTTCGTCAAAAATAAAAGATAGTGTTTATAATAATTTAGATAGTAATGATAATAATATAAATAGTAATGATAATAATATAAATAATAATGATAATAATTTAAATAATAATCATAATAAATTAAATAATAATGATAATAATATAAATAATAATGATAATAATATAAATAATAATAATAATAATTTAAATAATAATGATAATAATTTAAATAATAATAATAACAATTTAAATAATAATGATAATAATTTAAATAATAATGATAATAATTTAAATAATAATGACCAAAATAAAGACACACATGATTACTCAAAATTAAGTAGTTGTGATAGTAGCACCCAAGATGATTATTTAAAAACCACCTTAAATAGAATAAGCAGTTTTTTAAAAGAAGAAGAAAATATTATTTTACATAATTATAATTTTGAATATTTAATAAAACCCTTCGATTTAGTATTACCAGTAGAACAATCAAGTAATAAAAAAGAATTAAAAGCTAAATTAGAAATAAGTGATAAATGGGAAGGTATTACTCTCACGAAGACCCAAATAACGAAAATTATAGAAATAATGAATGAAGCTAATAAATCTAGGAATCAAACAAATAAATTATTATTAAAGCATGCATGGACAGTTCGATTAGATATCGAATCGTTAAGAAATGAGACGAAAAACGAATTTATAAATCTATATAATAAAGTATTAGGAGAAGAATATAATATAAGCAATACAGAATTATCTGCACAAGAAATTAATAGATTACAAATTTTATATGATGTTGTGGGTGTTCGACATTTGGCAAAATGGAGATTACACTGTAGAACCACATTAGAAAAATTTATTGAAGAAAAAAATTTAAAGAAGAAATTTTTATATGATTCAATATATAAGCAGCAATCTTGGTGGTCTTGGGTAACAGGCAATAAAAAAGAAATTGAGAATAAAGTACAAAACATTTTAAATAGTGAACAGGATATAATAAATGAAAATGAGTTGTTTATGATACAAGAAGCTATGACAAATGATGATAACTATGATGTTGTTATGCCATCAAAATATGATTTTCAATTTAAACTAGCTAATTTTTCGATAAATGTATATGATGATTGTAAAAAGAGGATAATAGATAATAGAAATGTGAATAATCACCATAATAATCATAATAATAATAATAATAATAACAATATATATATTAGGGGGGTAAGTAACAAAAGTGAAGTAACTAATAATAGACAACATATAAAAGAAGATAAATTAAATTTTAATGATATTATAAAAGATGAATTAAGTATAAATAGCAGTTGTGAAACCTTATTGACAAGTTCCATAAATTTAAGTTCCACTGATTCCAAATTAAATAAAAGACGTGTAAGATATGAACGGGTAAATATCTTGTCTATAAATTTTTATCAAATATATTCTTCTTTATCTCTACAATCAGTTGTTGATCATAATGATCATGATAATTTCCAATGGAAGTTTATTATTGAATTACAAAATTTTATAGCAAAACACAAAAATAAGGTATTTATGGAATTTAGGAATAATAAAAATTCCTTTCCATATAATATGAATACATCAGGAAGTCTATATAAAAATTCTATTTTCTATTCATTATTATATAGTCAAACTGTTTGCGCCTATTTAGAAATTAACCATTTAGTAACAGAAAAGGGTAATACATTATCCACTATATTAAGACTGAACCCTTTGGAGCTTTATTTATCTCCATTATTAATTAAAAGTATATTGTCATTTACATTTCCTCTATTAGATATAATTAATAAAAGTCATAAGACAGCAATGAAAAAGTACAAGAGTAAAAATGTGAGTTTATCTGTGGAAAAGCAAAAACAAAAGGAAGAAACTTCACCTGGTGTTATTAAAAAGAAGGAATCAATTTCGGGCACAAGAAAAAGAAGTAATAAAAATAAGAAGAGCCATCATGAAATTGGAAGTATAAAGAAGGATGTAAATAATGAAGAAAATGATACGATAAAGCAAGATGATCATAAAAATATGAATGCTGATAGTAATAAAAATAATGATAATAATTATGATAATAATGATGAATGTGATGATAATAATGATGAATGTGGTGAATATAATGATGAAGATGATGAGGACAAAAATAACGATAATGAGGATAATAATAACGATGATGATGAGGAAGATGAGGATGATGATGACGAGGATGACGAAGAAGCATTATTACAAATAAAAAAAATGGAACAATCCGAATTATTAGAAGGATTAAAAGAAAGAGGAGAAAATGTTTATAATAGAGCTGTTCAACACCTTCCAGAATTATTTGAGTTTTATATACATATATGTGGTCCCATTTTACATTTTGACAATTTAACGAATGGCATAGTGGAATTGCATTTAGGTAATTTGGTAGCAAAAACAGAACACCCATGTACCTATAATAAATTTAATTTAATATTTGAATTTAATGAAACGCAAATAACATGCTTAAAAATAGGCTCAGGATATATGAAGAAGGATGAAAAATGTAAAAAGGATATAATTTTTAATGAGAACCAAAAAGATTTGCATAAATCAACAAAAGCATATAAGAAGATGGATAAAAGTAATATATATTTAGAAATAAATGAAGAAGGATTATTTTCAAATAATGCAATGGATAATACCGAAATGGATGATTTAAAAGATAGTTATTTTAATAGTATAAAGGAAAAGGTATTAACAGGTGAAAAATTTTATATATTACAACCAATTCCAGTAAAAATCTATGTAGAATATGACTTGAAAATTTTGAAAACAAATATTATATTAGATGGAATATTTTTTCAAATTAATCCAGATGCTATTAGTATTATATTAGCAGTTCCAACATCGATAACTAGATACCTGACTGGAGTTTATTCTAAAAATAAAAATGAGAATGAATTTAATAAGAACAAGAAGAAAACTTCAGTTGAAAAAAAGGAGGGAGATATAATGAAAGGAGATGATATTACAAGAGCAAGTATGTCAAATATCGATAGTGGTGTACAGAAAAGAAATAGTATTACTGATATTATAAAAGGGAATAAAGATAATGATAATGTGGTAGAAATGAATAATAAAGGTATGCCAAAGGAAGAAGAGTCGTTTTTATATGATATTGATTTTTTAATTAAAAATTCTTCTTTTTCAATTAAGAATAAGAAAAATTGTGAAATATTGAAATATGAAGCATGTGGTATATTTTATAAGAATTATTTACAAAAGAAGAGGAAAATTATTAAAATGGAAATTGAACAGTTATGGATTTGTGATCCTAGTAATAAGCAACCTATATTTTTTACTTTAACGAAGGATATAAATTCAAAGGATATATTTTTATTTAGATCCTTATCTACATATTTGGAAGAGAAAAGTCATAACAATAATAGTATTGAACAAAAGGAAATTGATATGGAAAGTGATAATCATATAAATGTTGGTATAGACGACGTATTAGAAGAAGATCATTTTATGATGGGCAACAACAATAATATTAATAATAATAGCAGTCATATTAGGAAGAATAGTTTTAATATAGGGTCAAATGCTTTTGGTAGTTTAGAAAGATTCGAATCTGCTAAAGTGATGGATGATAATGTATACACAAATGGAAAACATAAAGGTTCAATCAAGAATGTAATGATTGATAATTCTATTATAAATAATGAACAATTAATAAACAATAGAGCAGAAGGAGGAAAATATATGAGCACAAATAGATTTTCTGCTAGTGGAATAAATCAGTTTAATAAATATGTTGGTATGAATATAGATAGAAAAAATGAAGAGACCAAATTGAATGGGAAAAAAGAAAATGATGAAAATAAAAATGTTGAGGAAGAAGATGATGATTTTATGGATGCTATTGAAGAAAAACAATTATCTATAAGTCTTCAAATTTTACAAGAACATAATGAAAAGAATATTGAAGAAACTCATATTAATTTTATTATTAGTGATATCGAATTACATTGGAAATATAAAACAATAAAACAGATTTTTAAAACAATGAAAGAATATAAGAAAACGTTACAATATGGTATAGAAAAAGATATGACATATATAAAAAATAAATTAAAAAATGAAAAGGATTTAAAAAATTATAAAATGTTCATATCTGAAAATACCTTAAGAAGTGTACAAGAAACTTTGAAAAATGTTAAGGATTCATTAAATATTTTAGATATGGAATCTGCAAAGGATAAGGATGATATAATAAATAATGTACAAGAAAAAGAAATGGACAAAGATAATGTTAAAATGAATGCAGAAGGTATACATCTTATGAATCCTTATTATGCGAGGGACGAAGATTTAAATATGATAGCTGATAATAAGCTGCTTTTTAATTTAAGAAATGAAAAAAATAATGAAGAAATAAATATTAATAATGCTAACACTGTAAATGATAATAATGGCATTAATAATAATGATGATATTAATAATAACGATGATATGATTAATAATGATGATATTATTAATAATCATAATTATAACATTTTGAATGATAATATTCTTAGTAAGGGACTTAATGATACATTACCCAAATTACACGGTCAACAAAATTCATATCCCAAATATTTTTTCAACTGTTTTATAAAAAGTGCTTCCTTAGCCTTCTGGAAGAAGAAGAAAATTTTTTCTAGAATTCAAGTGAGTAACATATTTTATGAAAATAAAATATATATCAATTTTGATCAGAAGATGTTTTTGAATATAGAAAAGGGGATAATGTCAATGAATAATAAGAACATTATATCTAATAATATTAACGATTATAAATACGACCTGTTTATTTCTAAAGATAGGAAAAGATTTGGTTCAAGTGATATCGAGAATGAGGATAGAAATATGTTAAGTCATTTAAAATATGGAAAAAAAGAAGGGAACTTAAATGCAATGCAGAATGATGATGATAAAGAAAAAGAAGAATATACAGCAGAGGATACATTTTCGGAAAGACGAAATAGTCGAGATATATTAAATAATATAAATCTAGATAATTTATCTATAGAAATAAAGAAGAAAAAGAAAACAATGCATAAAGAATCCAAAGAAGATTTGTTTATTGGTAAGATAAAAGTATATAATGATAAGAGGAATTATAATATATGTTTATTATGTGAAATTCCAAGAATGTATTATATTTTTTACCTGAAAGATTTAAGATTATTTTTGGAATATTTAGATGATGGTATATTAAATGTTTTTATTAGTAAATCATATAAAAAGGTAGTTCAGGCTGCTCAGACCAAATACTTTTTATTTAATTTTACATTTATGGATCCTATAGTGATCATACCTGAAGATAAAAATATGATATATAATTATAAGGGAGAGGTGAAAAATGTTTCTAGATATTTTAAGAGTAGCACTAGTAATAATAACACAAAGGAAAATGAGGACACAAATGATATTGCAAAAAGAAGCTCAAAGGAATATATTCTTTCCAATGAAAAAAAAGATACAAAGACTGAAGCGTGTAACAATTATAATAATGTGAATAATAAGAATATACATAACACTAATTTTGATTGTGATATGAAATCCAATTATGATAAAAATTCAATTGGTCGTAATACATATAATAGTAATAATATACCATACATTGAATCTTATTTACAATTCCATTTAAGCAAATTAAAACTTAAAAATTCTTACACTCTAAAATGTACGGAAGAAATAAGAGTAGCACAGAGAAGACAAATGATATTAAAAAAAAAGAAATATAAAGGAAGAAGTGAAAAGGAAAAAATAAATAAAAATAAATATGAAAATGAATCTTTTATAAGAAGTGATATTGTATCTAAGGAACAATTAAGACCATTAGAGCGAAAAAGGAATAGTGTAAATAATGATTTGGTGAAATTAGATTTTACGTTATATATTGATATATTAGATATAGAAAGTAAAGCTTGTGAAAATGGGGGTAGTAGTAATATAGAAGGGGAAATCTTGCATAAGGTAAATTTAGGTTTCTGTTTAATAAATGCTAGGAATGGTATATTTATATATTTAAATGGAAATGACTTATGTTTAGATTTAACAGTATTTCAATTAGCTTTTTTGTTGGATATAATAAATGAAAACTTTTGTTATAAGGGATATTTCCCAATGTGCTTTATATGTGATAAGGATGTGAGTTTAGACAGTATGATAAATTCGGAGTGGTTTAAAAAAGGATACATGAATTTTAACAATTTTAATAATAGGAAAGAAATGAAAGGAAAAACAAATGACAATAAAAAGGAAGATATAATGAACAATATAAATAATATCAATAATATCAACAACAACAACAATAATAATATCAATAATATCAACAACAACAATAATAATAATAATAATAATAATAATAATAAGGTAAATAATTATGAAATTGGAGATGGTGCTATAAATAACAATGTGTATGATAGCGATCGCACTATTGATAAGGAAATGGATCACGGTAAAAATAAATTGGATATGTCAATGAACCAAGAAAAAAAAAGTGATTCATTAACAAATGATAAACTAATAATAAAAGATTCAAAAGAGAATGTAACAAAAAAGACAGGCTTGAAATTATATGTATATATAAATTTTGAAAGTTTAAAAATTAAAACAGCATTTGATACAAATACACCTGTAGCCATTATAACATTCCAATATATAAGTATGTCATTTCGTTTAGTTCTCTTAGATTTCTATTATGTTTACTTTTTTGATTTACATGGTAATTCATTGTATATAGATGATGCTAGGAAAAATTCAATCAATTATTACAAAAGGGTAGCCTACTGTTGTATTGAAAATGAAAAGAAAAAGTTCAAGAGGGAAAATAAAAAAGCGCATGAGTTGTATAAAAATGTATATAATGAAGAAGCATATAATGATACTCTTAATAAGAGTAATGTATATGAGTATAATAATGTGGAAAGGAATTCGTATCAAATTAATAGAGAAAATAAAATATATGATAATAATGGGTCACATTTTTATAATAATAAAGATGTTCAAGATGAATATGATGAAAAAGTACAAAAACTACATAATAATAGTAGTAATAATATTTCTGTTACGACACAATCTGATATAAAAGATTATAAAAATTTATTAATATATATGTTTGAAAATAATGTATTTTTAAATGAAAAGAAGAAAAGGAAAAAACAAAAAGGAATAAAAATAAAAATTAATTCTTTCATAGAAGATTTGTTATTAAATATTGAACTTGATGATGCGTATATTTGCTTTTTTTTCCTTATTTTTATAGATATATATAAATTTTTAACGACAGGATTTAATTTGAGCACGTTACATTTGTATCCGAAGCCATCACCTTATATTGTATCTTACAAAAATGAAAATAAAAAGAAAAAGATAAAGGGTCATGAATATGTAGAGAATAGTGAAAGTAAAAGTATCCATAGAAAAAAAAAGGTTGTGTGTTCGAAGACGGAGGAAGAAAAACATATAAAGGAACAGATTAGTATTAGATGTACTAGATCTCAATATAAAAATACCTCAAATGAAAATAAAAATAAAAATATTAATAATAATAATAATAATAATAATAATAATAGTAATAATAAGAGTAGTATCAACTGTAAGGAAAATACCAGTATAAATATCGAAGAAAATGTGTTTAAAAGAGAAAGAATTCATTATGATGAAGAAATTATTGAACAAAATGTTCAACTCATCAATGAAGAAGACAATGAAAAACATTGGGTACAAAAAAAGAAGAAAATTATTCGACGTAGTAGTGTTTTAGAACAATCAAAATATATAAAAGGGAAATTAAAAGAACAGAAAAATAAAATAATAACAAATATAAACGAAGTATTAAAATTAGATAATAAACCATTTCATGTAAGTTTTAAAGTGAATAATGGGAATTTCATTATTTTTACAGATGTCGAAAAAGTATCTCATCCAATAATAATGTGGTCAAATAATTTTGTTTTTTCATTTTCTTTATTTAATAAATGTATTGTATTTAGAAAAATATATGCAATAGATAGTAAAGTGAAGAGGATTAATTATGTATCGTCATCTCATGTACATTATAAGTATGATACTATTAAGAAGAGGAACAATAAAAGCAACCATAACAATAAAAATAACAATAACAATAAAAATAACAACAACAATAAAAATAACAATAAAAATAACAATAACAATAAGGATAATAATAAAAACAACAGTAACGATAATAATAACATCAATATGGACAATAACAATAAAAATAACAATAACAAAATATTTGATCCTAAGAATATATCGAACCCACTTCATAATTATCATAAAAAGAAAATATTACTATGTGATAATTTGAATGTAATGGGTGAAGCTGTTTACGAATCAGTTGATACAAGAACTGAGGATTATAAATTAAAAAGTAGTGTAATAAGAAAAGATAACAATTCACCATTTATAACCGTTTTTGAATTAGATATAAATATTGGTAATTTTGATATACGTTTATCAAATGACGATGTTGAAATATTATTGAAAGCATCATCCACATTATTTGGTGATGTCCCTAGTTCATTTACAAATATAGTAGTTGGCCCAGTTATACCACCGTTAACTTTTATACATAAAAAAATAAAGAATAAATTTATGCAAAACAATATTTATTATATAAGTACAAATATCAAGGAGGAGACCGAATTGTTTTGTGGATCCGATTTGCAAATAACAAGTATAAATATATTAAAGAAGAGAAGGAAAAAAATTATAAGAAAGAAGGAAAAGAAAGATGCAAGAAATAGAAAGGAAATGGAGAGAAATGATCATAATAAAATAAAGAATATAAGAAATGATGAAATTAATATGAATGAAAAGAAAAGTATAGTAGATTGTCATATTGAGGAAAAGAAATTAGATCAGAAAGAATATAACGAGTTTAGTAATAAGGATGAAATATATACTGATTCTAGTAACATGATGGAATATGTATATTATAAAAAAATATCTGATGTCTCTTCAAGTACTAATACTTTAAGTTATTCCAATACATCTTTAAGTGAATTACATGGAGAAAAAAATATATTATATAAGAGAGGTATGAAATATGATGATATATCAGATATAAATAATAACAGTAGTAGTAATAATAACAACATCAATAATAATAATAATAATAATCAAACAAATAGTTATCGAGATATAAAAATTAATATAAAGTTACATAATGTAATGTGTACCTTTATAGATAATATAAGAAATTCCATTGTTCCCATTTTAAGAATGATATTTAGTATGAACATAAGTATTAATTTATATACTGATGAATGTTCATATAATATTAGTGACTTAAATTCTAAGTTAGAATATTTTAATAATTGTATAGGTGAATGGGAACCATTTCTTGAGAAATGCAATATTTCATTAGATATACATAATATTTTTCCAAATGACGAATATGATGAAGATAAGGAAGCAAACAAATCTCCTATAAGTATCATTAAAATTAATAGTATCAAAGCTTTGTGGTTTAATATTACACCTCAGTTAATTAATTTGTTATTTTTATTCGTGCCAGTCTTTTCTGAGAAGGTATCTAATGGGTTAAGAAAAAATGCCAAAAAGGAAATAGACCATTATAATGATAAGGATAATAAATATGTAAGTGAATGTATAAAAAATGATATATTAATGGAAGATATGAATAATAAATTAAATGTTAGGAATACATCTATTAATAAGGATATAGAAACATCTGAACAGATGAATAGTCTAAATGACTTTTCCAGATGTGATAGTGGATCCGTCGAATTGGAATACAAAAATATGAGTGTTAAGAGTAAGAGCATATTCGAAGATTGTTCTTCGGTTTTTAATTCATTTCCGTGTGATACAGATGGTAAGGAAGCAAATGATTTTCATTTGGATGACTTTTTAAAAAAAGAAGAAAATCGTAGTACATACTATAATAGGATAGATAACTCAGTAATATATTACGTTAATTTGACAAGCGAATACTTTTATGCCTTTTTAATGCCTGAAAGTAAGATTGAAGAAATTATGAAGAGAAAAAAATCTGTCACTATGAAAAATGTATACAAAAGTGTGAACGAGGCGGATGGGACAAATATGACTATAAAATCTTATAAATGGGATGAGAAAAAGAAAAAGGTAGAAAAGGGATTCATTCAAAGGGAGAATGTTACGAAGGCGAATGGAATGAAAGGAAATGATATAAAACAAAATGGCATAAAACAAAATGAAATAAAAGAAAATATAATGATGGAACATACATTAATTAGTTCAGATGATAATAGTACCCATAATGAAGGGAATGAAGATTATATATCTGAGGAAGATCTTTTCAATATGATACCTGATGTATATGCAAAAATTATAACAACAAATGAATTAATATCATTAGACAGATTGTTAATCAACGAAATAGAAAATAACTCCTTAATAGAAAAAAAATGTTTATATTTATATTTGATACCCATTCCACCAACAAATGTTGTGAATATAGTTCATGATATGTTTTCAAATATTAGTAAAAGAGATATTGTTTTAGATCTCATGATTACAAAAAATCCAAAGAAAACTATTGAAAATTTATTAATTTACAATCAACATAAGGAACATATACGTATTAGGAGAAATAGTATAAATAGCATTTGGACGGAAAATATGATGTATGATGACAATAATAAAAAGAAACATATAGAATTGTATTTGAAAAAAATGCGTATGAAAGCGGAAGAGGATGGTGGTAACAAGGTAGGAACGGATAATATACATTTATTGGAAGATCAAAATATGTATTTATTAAACAAAGAAATGTGGAATGATTATAAAAACGTTAAGGGTATTATAAAAGGGGAAGGAAAATATACTAATAGTACTATTAGTAAAACGGACCATACTTCAAGGAGTAACAATATGGACAATAATGTAAAAGGTAATGATAATATTAATGGTAATCACAGTGATAGAGAATCTGAAAAAAGACAAGACATGAATCAAAATAATGTAAATGAAAATATCTTAAATGTTATGGTAGAAGGAAACAATGAGGTATTAGATAATGAGCAACGGGTTGATAAATCATTGAATTTATATACAAAAGGTTGTGAAAATGATGATGCTAATGAGGATGTCAATGATGAAGAAGAAGGGTCTTATGATAGTTCTTCTGATAGCTGTGAGGATTATGAATCAACTGATGATACCCATTCAATTGATGAAAGGGATAATTTGAATTATGAATATTATTTTAAATATAATAAGAAAAAGAAAAATGATTCAAGTTCTATTTTTTCATCGAATAAATTTGTTGGACCCTTTTTAAGAAACACAGAATGTGTAATAATAAATTTGATTAAGAATAGTTGTACATCATTAACTACATCTTTGAAAAATAGCAACGTTATACATTTATTGAAGCCTGGTGATTTTGTGTTTGATGAAAAATATAAATATAATTTTTCTAGCAAAAAGACTAATGAAATTATGGAAAATTATAGTATTACTGATGGGATTATGGCAGACGATGACATAACTAATGGATCCATAAATAATAATAATAATAATAATATGTGTAACAAAGGTATTTATGATAATACACCAACAAAACAGATGGTCGATAAATATGTTTCTAAAAATAGTATGAGTACTAATAAGAATAATAAATCGAGTATATTAAATAAAGAAAATATTGACAAGATAAATGAATGGGAAAGAAGAAATGAAGAAATGATTAAACAAACATATATAGCATGTAAATATAGATTGAAAAATAATTATTTGCCACAAACATCGTTGGTTGAAAATGTATGTGAAATTATATCTCCTATGCCAAATTATAAAATATTATTTATGTCATCAACCGTTAGAATTATAAACAAATGTGGTATTCCCTTAGAATTTTGTTTTTTTGATGGTAGTCGAAATCCTATATTATTAACGTCATTAGAGAATAGGACTATACCTATAAATACTTTATATCCTAATCATAGTGATTCATTTAAAAATTATAAGGTATCTAATAGTTTGAATATTAATCCGAATATAAAATTAAGACAGAAAAATAATAATTTAAGTTTCACTGTAATATTAAATCATGAGTATTTATTATCAGCACCCGAGTGTGTTTTTTGTGGACCTTCTCATGTGTACATGTCTTTTAAACCTATCAATGTGATAACAGCAGAGAATGAATATTATGACATAATAGCTGGTGGTCCCACCACAAATATTGCGAACAATAATAATAATAATAATAATAATAATAGTAGTAGTAGTAGTCATAATTATTATGGTAATAAATCAAATATTAATGATTTATTAAAAAAGGACAAGGGGGGAAGCAATGATTTAATAGATCCTATGTCTATTAATATAGAAAATGGATGGTCTGATATTTTTAGTTCTGATATATCTCAAGGGACGTATGTGAAAAAGTGTAAATACAAAGATATGAATTCCTTCATGTCTAATATAAATAAAAGTAACGGTAGTGGAAATAATTTTTTTTATTTCTTGGTAAAGATTGAAAATAAGATAAGTGCTTTACCTGCTGAGAAAAATTTGAAAATTATTACTATATATCCTCATGTAAGTGTTGTAAATGCTATACCAGCCTTTGTAGACATAATAATAACATCTGACGTAGTAGAAGAAAAGCAGAATGATTATATGTATGAAGAAAGGAATAAAATAGATGCATTAAAAAATAAGAGAGCAAAGTTATTAAAACAGTGTGAGGATATAAATTCATATTTGAATAAGATGAATTCATCAAAATATGGAGGAAGTAAAATTTATAGTGATAGTTCATCTAGAGTAAAAGGTAACAAAGATAATAGTAATAGTAGTAATAGCAATAATAGTGAACATGTATACATGCAAGAGTTGAACAAATTAGAAAGAGAAATATATCATATAGATGAAGAAATAAAATATAAGAAAAAAGAATTATCTTCTCCATTAGTAAAAGGATATGTGAATAGAAGATTAAATCCATTTTCTATATTTTATATATATGAAATTAAGAAATATACTTGTTTGAATTTAAAGATGAAAATAGGGAATTCACAATGTGAATGGTCTGAAAAGTTATTTTTAGTAGAAGATGATGAAGAATCTGTAACTAGATTTTCATTACATTTTAAAAAATATGCATCAGTCGAAGTTGAAATTATTAAAAACTTTAGTGGATATTTTAATTCTTTGAGTAGTATCTTAGGAAATAAACAATTATATATTTTTTCCTTACCAAGATGTTTTATAGATAGAACAGGTTTAGGAATTAAGGCAATAAATTCGAATAAATATTATCCTGTGATTAATGGAATAACCTTATTAGGGGATCATTCCCAGATTGATTTGTTATTACCTCACAAAACTTATACTCAACCGAACAATAATGTGAACAGTGGTAATAATAATGTTGAAGATGATACGTATAATTATTATTCTAAGGGTGATATGAAGAGATCGAATAGGATGTATGATAAACGAGGAGATCATAGAAAAGATTATAACGATACAAACGAAAGTTATGGTAATGTAGATGATGTTAATAAAAATGATAGTCGTTATGTTAGTAAAAATGCAATTATAATAGATTACAATGATATATTTAATGATTTTAATTACCCTTCACCTGTAAATAATTCTGTAATATTATTCAAAGCTACATTACCTCCGATTGGTAGTTATACAGAAACCAATGTGATTTGTAAAAATTTTTTTTATACCTTTTGTTTAAATACAGAAAAAATAAAAACAACGAATATTCCATATATTATAAGTAGAATAATTACAGTGGTTCCACAATTTATTATTAGTAATAAATTGAATTTTCCCTTATTAATTAAACAATATCAGAATGACCAGATGCAAGGAGTTCGTGCAAATGATACATCTCCTTTATATTTTACCAAAAAAAGTAGCATTTTACTATTTCAATTTAAATGTTTAGATCCATGTAGAACTAATAATGAGGAGAAGAACCATATGTTATTAGATCCGAAAAATGTAATAAAGAGTAATGGAAATAAATATTATAAGTTGAATAAAAATATATATTGGTCGTCTGTTATATATCCTAGTGAAAATTTTGTTGGAACAAATTATATGGTAATTAATACAGGAAATCACGAAAAATCTGATGTATACGCGATAACGTGTATACCTGATAGAGGTACGAAAAATATTATCATAGAAAAATTAGAAAATAAGAAGAAAGGTTTTATAGCATATAATAATTCAAGTATTGCAAAATATTTAAAGATTAGAACATTCCATGATGATACGAGACATATGAAAATTCACGATGAAGAGAATTATATGGAAAAGAATATGTTTTTAAGTAATTTTTTAAAACCAACAGATATGGAACATTATTTTAATATAGAGCCAAATAAATATAGTTATTTGGGATGGGTAAATCCATTTATATATGTTACTAGAAATGTACAGATAGAAATAGTTTTGGAAGATTTAAAAATAATACCTAAATCTCCATTTGTTTTAAAATTTGCTATATATAATTATTCACAAAAAACATATTATATAAATTATTATAATATTACATTTGTTATATGTATAGAATATATAGAAGATTTGATAACCATAAAATTGAGCCATAAATTAAATATAAGCTCAAATCTATTAGATCGTTCCTCTTATTATTATATGATGAATAATAAAAGTAATATAACAGATGAAATAGCATATTCTAATAGTTCACATAATAATATTATGTTCAGCAAACATATGAATAGTTGTGACAAATCTGTTACATCTAATAATGACATTAAATTAGGATTAAATAAAAAGAAGAAATATATGGAAATGGAAAATCAGGTAAATATAGAAAAATATAATATAATAAATGGTAAAGATATTTCAAATGAAAGGAATATTATAGATAAGAATTCAGAGAATAGTTTAAGTTTATCAAAAAATAGTGTTTATGCCCAAAAACAAATGATTATAGATCATGATGAATATCATTATGTAAAATATATAGATGAGAAGGATGGGGGTACTGGTAGTTATAAAAATAATAGTATTAGTAGTAGTACTAATAATTATTATAAGAATGAATCTGTGAAGATGATAAGTAATACATATAAAAATGTGCATATAATAATAAATGTAACCCAGATTGGTGTAAGTATTATATCTAATATATTAAAAGAGGAAGTATTCTTTATTGAACTATCAAAATTGTGTGCTTTGTTTTATATGAAAAATGAAGAAGAGGTAATAGATATAAAAATAACAGATGTACAAATAGATTGTCAGTTGGAATCTTGTGAAAAGTGTGTGTTGTTAGCGAATAGGGGAATAAGTTCTTCAAATAATAATAATAGTAGTAGTAGTAGTAATAATATGAATAATATGATAAGTGGAGGGGATATGAGTGGTAGTAATAATGTATATCAGAATTATCGTAATGAAAATAAAAGTTTTACAATTAATGAGAGTGACCTCCTTCGTTTTAAGAAAACATTTGATCGTAATTCAAACGACAAAAATAATCATATGAATCCTCTATATGATAATAATAAGAATAAAGGTACTACTACTACTACTACTACTAGTGGTAGTAATAATAATAACAATTATAACAATAATAGTAGTAGTGTGTTAAATAAAAATATGAATGCAAATGAAGAAAAGATCTTTTTAAATATATATGTAGAAAGGTCTTTTATATCACATAATGATGTTATATTTAAAAAAATACAAGTTTCGTTAGATGATGTAGAAATAGAAATGGATGCTGAAACTTTGAATGGTATAAATTTATTAACAGCAGAATATATTGAAAGTATAAGTATTGTTCAAAAAAAAAATTTGTTATATGAAGAAATACAGAAATGGACGATTTTACCTGTATATGTAAATTACAAATCACCAGAAATACCATTAGCCATAAATATCCAATACATGCAAATAGATAAATTTACCTTAATAGTGTGGTGTTCATTTTTATTAGATAAAATGCACATGATGAGTGATTTGTTACGAATAGGATTACGAATATTAATGGTATCGGGAAAATTAGAATTATTAGGTGCACCTGTAACCTTAAATCAAGAAATTTTTAATAATATTCGT**GTAAGTATTAAATCCTTTTATGCATTACTAAAAGATAAGTATAGTCATTCTATTTTGGCATGTTTAGGTTTTATAGTAGGATATAGTAGTTTAATAAATATACCTAAGATACCATTAGAAATTGGTCGAAATACTATAGGTTTAGCTGTATATGCAGTAGATAATGTGAGTGTTGGAATTGGAAGCTTTTTATCAAATTTAACATTTGATTCCGAATATATTAATAGAAGACAAAAGGAGAGAACCTTTAAGACGAACACCAATATGAAAGAAGGTTTAATTAGTGCTGTTAAAAATATTGGTGAAGGAGTATTAAGTTTATCTAATATTGTAACGAAGCCAATTGAAGGGGCACAAAAAGAAGGATTTGGTGGATTTTTCAAAGGAATAGGAAAAGGTGTTGCAGGTTCTTTAGTAAAACCATTAGATAAAGTAGGACAAGCTGTATCTGATGTAACTCGTGGAATAAAAGCAGAAGTATCGAAACCTATAGGTGGTCATAAATATAAGACTAAAAGACACAGAAAACCAAGAATGCTATGGGGAGAATATGGAAAATTAAAAGAATATAATATAAACGAAGCAGAATTAAGAGAATGCTTAGGTTTGAAATTTTCAAAAAATATTATGAAATGTTTAACAGTACATAAACAAGAAAATCATCCTCCATCACATTATGCATTATTATTATATCCAAAGGTAATTATATATGCAAATCTATATGCTAATACTAATGCTCAAAAGAATGATTTCTTATTATCCGAAAAAAAAACAGATATAGTTATATGGTCTATAAAAATAGAGGATATAACAGAAATACGAGCATCTAGTCATGGACTCATAGTAAGAACGAATACAAGTACAAATACGGTTTATAAAATACCATGTAATAATGCTATATTGATAAATAAAATATATAGGGAATTGCATAACTCAAAAAATTCAATAAACTCTACTGTTATTTTGGGCGCCAATAATTCTGCCCTGTACAAATATTTA**CCTAGGTCAGGATTGAGATCAAGATCTGCTGCTGCTGGTGCTGGTGGTGCTGCTAGAGCTGCTCTGCAGAGAGGAGTACAAGTTGAAACAATATCACCAGGAGATGGTCGTACATTTCCAAAAAGAGGTCAAACTTGTGTTGTACATTATACTGGAATGCTTGAAGATGGAAAGAAATTTGATTCATCTCGTGATAGAAATAAACCATTTAAATTTATGCTAGGTAAACAAGAAGTAATACGAGGTTGGGAAGAAGGAGTTGCTCAAATGAGTGTAGGTCAAAGAGCAAAACTTACTATATCTCCAGATTATGCTTATGGTGCAACTGGACATCCAGGTATAATTCCACCTCATGCAACTCTTGTATTTGATGTGGAGCTTCTAAAACTAGAAACTAGAGGTGTTCAGGTTGAAACAATTTCACCTGGAGATGGCAGAACCTTTCCTAAAAGAGGACAGACTTGCGTAGTTCATTATACAGGCATGCTAGAGGATGGTAAGAAATTTGATTCTAGTCGAGATAGAAATAAGCCATTCAAGTTTATGCTAGGTAAACAGGAAGTAATAAGAGGTTGGGAAGAGGGTGTAGCACAGATGTCAGTTGGACAAAGAGCAAAGTTAACAATATCACCAGATTATGCATACGGTGCAACAGGCCATCCTGGCATCATCCCTCCACATGCAACTTTAGTATTCGACGTTGAATTGTTAAAGTTAGAGACAACGCGTGCTAGAGGTGCTGCTGCTGGTGCTGGAGGTGCAGGTAGACGTACGATGAGTAAAGGAGAAGAACTTTTCACTGGAGTTGTCCCAATTCTTGTTGAATTAGATGGTGATGTTAATGGGCACAAATTTTCTGTCAGTGGAGAGGGTGAAGGTGATGCAACATACGGAAAACTTACCCTTAAATTTATTTGCACTACTGGAAAACTACCTGTTCCATGGCCAACACTTGTCACTACTTTCGCGTATGGTCTTCAATGCTTTGCGAGATACCCAGATCATATGAAACAGCATGACTTTTTCAAGAGTGCCATGCCCGAAGGTTATGTACAGGAAAGAACTATATTTTTCAAAGATGACGGGAACTACAAGACACGTGCTGAAGTCAAGTTTGAAGGTGATACCCTTGTTAATAGAATCGAGTTAAAAGGTATTGATTTTAAAGAAGATGGAAACATTCTTGGACACAAATTGGAATACAACTATAACTCACACAATGTATACATCATGGCAGACAAACAAAAGAATGGAATCAAAGTTAACTTCAAAATTAGACACAACATTGAAGATGGAAGCGTTCAACTAGCAGACCATTATCAACAAAATACTCCAATTGGCGATGGCCCTGTCCTTTTACCAGACAACCATTACCTGTCCACACAATCTGCCCTTTCGAAAGATCCCAACGAAAAGAGAGACCACATGGTCCTTCTTGAGTTTGTAACAGCTGCTGGGATTACACATGGCATGGATGAGCTCTACAAAGTCGACGCCAGGGGAGCAGCCGCAGGAGCAGGGGGGGCAGGAAGGCGTGGTGTTCAGGTCGAGACTATTAGCCCTGGAGATGGACGCACGTTTCCTAAGCGTGGACAGACATGCGTAGTTCACTACACAGGTATGTTGGAGGACGGTAAAAAGTTCGACAGCTCACGCGACCGCAATAAACCTTTCAAGTTTATGCTTGGCAAGCAGGAGGTTATTCGTGGATGGGAGGAGGGTGTAGCACAGATGTCTGTTGGACAGCGTGCTAAGTTGACAATTTCACCTGACTATGCTTATGGCGCTACGGGCCATCCCGGGATCATTCCGCCACATGCGACTCTGGTATTCGACGTTGAATTATTAAAGTTAGAGACAGCTAGAGGGGCCGCTGCAGGTGCTGGTGGAGCTGGAAGACGTGGAGTACAAGTAGAGACTATCTCTCCAGGTGACGGTCGCACTTTCCCAAAGCGTGGCCAAACCTGTGTTGTACATTACACTGGTATGCTGGAGGATGGGAAAAAGTTCGATTCCAGTCGCGACCGTAACAAACCGTTCAAATTCATGTTGGGAAAGCAGGAAGTGATCCGCGGGTGGGAGGAAGGCGTGGCGCAAATGAGCGTCGGTCAGCGGGCTAAATTGACCATTTCCCCTGACTACGCGTATGGGGCTACTGGGCACCCAGGGATTATTCCGCCTCACGCTACACTTGTGTTTGATGTCGAACTTTTGAAACTGGAAACTGTCGACGGAGAAGGAAGAGGAAGTTTATTAACATGTGGAGATGTAGAAGAAAATCCAGGACCAATGATTGAACAAGATGGATTGCACGCAGGTTCTCCGGCCGCTTGGGTGGAGAGGCTATTCGGCTATGACTGGGCACAACAGACAATCGGCTGCTCTGATGCCGCCGTGTTCCGGCTGTCAGCGCAGGGGCGCCCGGTTCTTTTTGTCAAGACCGACCTGTCCGGTGCCCTGAATGAACTGCAGGACGAGGCAGCGCGGCTATCGTGGCTGGCCACGACGGGCGTTCCTTGCGCAGCTGTGCTCGACGTTGTCACTGAAGCGGGAAGGGACTGGCTGCTATTGGGCGAAGTGCCGGGGCAGGATCTCCTGTCATCTCACCTTGCTCCTGCCGAGAAAGTATCCATCATGGCTGATGCAATGCGGCGGCTGCATACGCTTGATCCGGCTACCTGCCCATTCGACCACCAAGCGAAACATCGCATCGAGCGAGCACGTACTCGGATGGAAGCCGGTCTTGTCGATCAGGATGATCTGGACGAAGAGCATCAGGGGCTCGCGCCAGCCGAACTGTTCGCCAGGCTCAAGGCGCGCATGCCCGACGGCGAGGATCTCGTCGTGACCCATGGCGATGCCTGCTTGCCGAATATCATGGTGGAAAATGGCCGCTTTTCTGGATTCATCGACTGTGGCCGGCTGGGTGTGGCGGACCGCTATCAGGACATAGCGTTGGCTACCCGTGATATTGCTGAAGAGCTTGGCGGCGAATGGGCTGACCGCTTCCTCGTGCTTTACGGTATCGCCGCTCCCGATTCGCAGCGCATCGCCTTCTATCGCCTTCTTGACGAGTTCTTCTAActcgagggatatggcagcttaatgttcgtttttcttatttatatatttataccaattgattgtatttataactgtaaaaatgtgtatgttgtgtgcatatttttttttgtgcatgcacatgcatgtaaatagctaaaattatgaacattttattttttgttcagaaaaaaaaaactttacacacataaaatggctagtatgaatagccatattttatataaattaaatcctatgaatttatgaccatattaaaaatttagatatttatggaacataatatgtttgaaacaataagacaaaattattattattattattatttttactgttataattatgtgtctccttcaatgattcataaatagttggacttgatttttaaaatgtttataatatgattagcatagttaaataaaaaaagttgaaaaattaaaaaaaaacatataaacacaaatgatggtttttccttcaatttcgatatcaatttatagaaacaaaatatatacttgtataattttatttttttatataaatcattacatatataattatacaatattttttctaagagataattatatattaatatatataaaaaaaggtgttttttttttttttttttatttttatttttattttatggtaatattttattttccttattttataaattatattagtttatatgtgattaattttatatattatcaatttatatatttttaaatgcttacttaattatctttttttttttttttttttttttttcccctctttttatattaatttatttttgaaaaaattgatatatatatatatatataatatatatatatacatgtagtagtattaaacaatgtataatatatataaataatatatttatatatttcatttcaattttaattttttttggttttttttttttttctttttgtcatatttaaaaaaaattatattcatataagttatgcattttttataaacattattcaatatatgtataatataatatatatatatatattaatgtattattccaatgtgcatgataaaagaaaaaaataatatttataaaaaaaaagaaaaataaaacaaaaaaagaaaaaaaaaaaaaaaaaaaaaaaaatacaaaaataaataatataatttataattatatattcttgtcacaataaaaatatatatatatatatatatatttataatatgtatattttaaactagaaaaggaataactaatattttatttattatcattcaagatttatattttataataataaatacctaatagaaatatatcaggatccatgcatggttcgctaaactgcatcgtcgctgtgtcccagaacatgggcatcggcaagaacggggactacccctggccaccgctcaggaacgaatttagatatttccagagaatgaccacaacctcttcagtagaaggtaaacagaatctggtgattatgggtaagaagacctggttctccattcctgagaagaatcgacctttaaagggtagaattaatttagttctcagcagagaactcaaggaacctccacaaggagctcattttctttccagaagtctagatgatgccttaaaacttactgaacaaccagaattagcaaataaagtagacatggtctggatagttggtggcagttctgtttataaggaagccatgaatcacccaggccatcttaaactatttgtgacaaggatcatgcaagactttgaaagtgacacgttttttccagaaattgatttggagaaatataaacttctgccagaatacccaggtgttctctctgatgtccaggaggagaaaggcattaagtacaaatttgaagtatatgagaagaatgattaagcttatttaataatagattaaaaatattataaaaataaaaacataaacacagaaattacaaaaaaaatacatatgaattttttttttgtaatcttccttataaatatagaataatgaatcatataaaacatatcattattcatttatttacatttaaaattattgtttcagtatctttaatttattatgtatatataaaaataacttacaattttattaataaacaatatatgtttattaattcatgttttgtaatttatgggatagcgattttttttactgtctgtatttttcttttttaattatgttttaattgtattttatttttattattgttctttttatagtattattttaaaacaaaatgtattttctaagaacttataataataataaatataaattttaataaaaattatatttatcttttacaatatgaacataaagtacaacattaatatatagcttttaatatttttattcctaatcatgtaaatcttaaatttttctttttaaacatatgttaaatatttatttctcattatatataagaacatatttattaaatctagaattctatagtgagtcgtattacaattcactggccgtcgttttacaacgtcgtgactgggaaaaccctggcgttacccaacttaatcgccttgcagcacatccccctttcgccagctggcgtaatagcgaagaggcccgcaccgatcgcccttcccaacagttgcgcagcctgaatggcgaatggcgcctgatgcggtattttctccttacgcatctgtgcggtatttcacaccgcatatggtgcactctcagtacaatctgctctgatgccgcatagttaagccagccccgacacccgccaacacccgctgacgcgccctgacgggcttgtctgctcccggcatccgcttacagacaagctgtgaccgtctccgggagctgcatgtgtcagaggttttcaccgtcatcaccgaaacgcgcgagacgaaagggcctcgtgatacgcctatttttataggttaatgtcatgataataatggtttcttagacgtcaggtggcacttttcggggaaatgtgcgcggaacccctatttgtttatttttctaaatacattcaaatatgtatccgctcatgagacaataaccctgataaatgcttcaataatattgaaaaaggaagagtatgagtattcaacatttccgtgtcgcccttattcccttttttgcggcattttgccttcctgtttttgctcacccagaaacgctggtgaaagtaaaagatgctgaagatcagttgggtgcacgagtgggttacatcgaactggatctcaacagcggtaagatccttgagagttttcgccccgaagaacgttttccaatgatgagcacttttaaagttctgctatgtggcgcggtattatcccgtattgacgccgggcaagagcaactcggtcgccgcatacactattctcagaatgacttggttgagtactcaccagtcacagaaaagcatcttacggatggcatgacagtaagagaattatgcagtgctgccataaccatgagtgataacactgcggccaacttacttctgacaacgatcggaggaccgaaggagctaaccgcttttttgcacaacatgggggatcatgtaactcgccttgatcgttgggaaccggagctgaatgaagccataccaaacgacgagcgtgacaccacgatgcctgtagcaatgccaacaacgttgcgcaaactattaactggcgaactacttactctagcttcccggcaacaattaatagactggatggaggcggataaagttgcaggaccacttctgcgctcggcccttccggctggctggtttattgctgataaatctggagccggtgagcgtgggtctcgcggtatcattgcagcactggggccagatggtaagccctcccgtatcgtagttatctacacgacggggagtcaggcaactatggatgaacgaaatagacagatcgctgagataggtgcctcactgattaagcattggtaactgtcagaccaagtttactcatatatactttagattgatttaaaacttcatttttaatttaaaaggatctaggtgaagatcctttttgataatctcatgaccaaaatcccttaacgtgagttttcgttccactgagcgtcagaccccgtagaaaagatcaaaggatcttcttgagatcctttttttctgcgcgtaatctgctgcttgcaaacaaaaaaaccaccgctaccagcggtggtttgtttgccggatcaagagctaccaactctttttccgaaggtaactggcttcagcagagcgcagataccaaatactgtccttctagtgtagccgtagttaggccaccacttcaagaactctgtagcaccgcctacatacctcgctctgctaatcctgttaccagtggctgctgccagtggcgataagtcgtgtcttaccgggttggactcaagacgatagttaccggataaggcgcagcggtcgggctgaacggggggttcgtgcacacagcccagcttggagcgaacgacctacaccgaactgagatacctacagcgtgagctatgagaaagcgccacgcttcccgaagggagaaaggcggacaggtatccggtaagcggcagggtcggaacaggagagcgcacgagggagcttccagggggaaacgcctggtatctttatagtcctgtcgggtttcgccacctctgacttgagcgtcgatttttgtgatgctcgtcaggggggcggagcctatcgaaaaacgccagcaacgcggcctttttacggttcctggccttttgctggccttttgctcacatgttctttcctgcgttatcccctgattctgtggataaccgtattaccgcctttgagtgagctgataccgctcgccgcagccgaacgaccgagcgcagcgagtcagtgagcgaggaagcggaagagcgcccaatacgcaaaccgcctctccccgcgcgttggccgattcattaatgcagctggcacgacaggtttcccgactggaaagcgggcagtgagcgcaacgcaattaatgtgagttagctcactcattaggcaccccaggctttacactttatgcttccggctcgtatgttgtgtggaattgtgagcggataacaatttcacacaggaaacagctatgaccatgattacgccaagctatttaggtgacactatagaatactcgcggccgcTAgGTAAGTATTAAATCCTTTTATGCATTACTAAAAGATAAGTATAGTCATTCTATTTTGGCATGTTTAGGTTTTATAGTAGGATATAGTAGTTTAATAAATATACCTAAGATACCATTAGAAATTGGTCGAAATACTATAGGTTTAGCTGTATATGCAGTAGATAATGTGAGTGTTGGAATTGGAAGCTTTTTATCAAATTTAACATTTGATTCCGAATATATTAATAGAAGACAAAAGGAGAGAACCTTTAAGACGAACACCAATATGAAAGAAGGTTTAATTAGTGCTGTTAAAAATATTGGTGAAGGAGTATTAAGTTTATCTAATATTGTAACGAAGCCAATTGAAGGGGCACAAAAAGAAGGATTTGGTGGATTTTTCAAAGGAATAGGAAAAGGTGTTGCAGGTTCTTTAGTAAAACCATTAGATAAAGTAGGACAAGCTGTATCTGATGTAACTCGTGGAATAAAAGCAGAAGTATCGAAACCTATAGGTGGTCATAAATATAAGACTAAAAGACACAGAAAACCAAGAATGCTATGGGGAGAATATGGAAAATTAAAAGAATATAATATAAACGAAGCAGAATTAAGAGAATGCTTAGGTTTGAAATTTTCAAAAAATATTATGAAATGTTTAACAGTACATAAACAAGAAAATCATCCTCCATCACATTATGCATTATTATTATATCCAAAGGTAATTATATATGCAAATCTATATGCTAATACTAATGCTCAAAAGAATGATTTCTTATTATCCGAAAAAAAAACAGATATAGTTATATGGTCTATAAAAATAGAGGATATAACAGAAATACGAGCATCTAGTCATGGACTCATAGTAAGAACGAATACAAGTACAAATACGGTTTATAAAATACCATGTAATAATGCTATATTGATAAATAAAATATATAGGGAATTGCATAACTCAAAAAATTCAATAAACTCTACTGTTATTTTGGGCGCCAATAATTCTGCCCTGTACAAATATTTATAAtaagaaaaaataaaatggcatatatatatatatatatatatatatatataatatatgttcatttatatgtttatttgtttatatatttatttacttatttattcatttatttatttattcatttattcatttatttattcatttatttattttatttattaatttattaatttatttattcattcatttatttattcatttatttattcatttatagttttattttcattatagcatatttccatattttttatttttatatatttaaacttgttttgattaaatatttaaaacttcttcaattctaatatatccattatttatctaattatatatagagagtgtatttttttttttttttttttttttttcttttttccagtttgccttcataaaaacaggcctatttaaataaaataaaaaatataaactatatatttattatatatacttttctatatgataaaatttgtttttaaatatatattattatgatattt

**5’ amplicon size: 1176bps**

**3’ amplicon size: 1504bps**

**> PF3D7_1346400-Halo-SW Halo Neo-R**

atattttcttatatattatgtatacataaatattattttaattttatataataatagcct

tatggcaattttaaaatgtagtaataattatattatgtaaaaaaagttcaaaattgataa

acaatgatatgtttttataattatacatttttttagcttttgctattctttttttttttt

ttttttttttttcgttaatttttctattttattccttttttttttttttttttttttgtt

tttgttttcttttttgtttgtttcaaattttttttttattttcttttttgtaattttaat

tttttcaagaattttataaaaatacatgtatcatatgtatttggtatttttatagaaaaa

ttgacaaaaaatatgtactgtatgtatatgtatgcatgtatatatatatatatatatata

tataaggacataatggaaaataaaaaataaaaaagaaagattaataaaaataaatataaa

aaagattaaaaagcataacaATGTTCGAATCACTAGTTGAAAAACTGTTGAATAAATTTT

TGGCTCCATACGTAGAAGGTATAGAACGGAATTTACATTTGGGTGTATGGTCTGGTAATA

TTGTATTAGAGAATTTAAAATTGAAACCACAAATAACTGAAATATTAGACTTGTCATTTA

AGATAATTCATGGGAATATAGGAAGAATAAATATACAGATACCATGGAGCAGTTTAGGAA

AAAACCCTGTGTGCGTTTTAATAAAAAATGTTCACATATATATAAAACCACGATGTTATA

AAAAAAGTGAAGAAGTTATTATTGAAGAATTAAGAAAAGCAAAAATGCATAGATTACAGT

TATTAGAAGAAGAAATTTCTCTTATTAAGTTACAAAAAAATAATGAGAAATCTTCTGAAA

AATCTACACTTATATTTAAATTATTAAATAAAATTATTAATAATATTCAAATAGATATTC

AAGATATACTAATTCATTTTGAGGATCCAGATAAAAACTTCTCTATTGGTTTTATTTTGA

AATCCTCTTCAGTCAAAAATTGTCAAAATAACGACGAGATGTCAACACAAGCAAATTCAA

ATGCAGAAAATAAAACATTGAATCATATAATTGAATTTAAAGGATTATGTATTTATAGTA

ATAGTGATATAAAATCAAGGACAAAGAAAAAAATTAGGAAGAAAAATAAAAAACGAAGTA

AAGATATGGTAGAAATATTGGAAGATGATAAAAATAAAGACAATGTAAAGAATTCATCAA

ATGGTAAGAACATAAGTGAAATGAAATATGAGAAAAGTGAACTTTCATTAAGTAATGAAA

AGTTTCCTAATATGAATAAGACAGATACCGAGGATATGGAAATATTTGATATTTCGTCAA

AAATAAAAGATAGTGTTTATAATAATTTAGATAGTAATGATAATAATATAAATAGTAATG

ATAATAATATAAATAATAATGATAATAATTTAAATAATAATCATAATAAATTAAATAATA

ATGATAATAATATAAATAATAATGATAATAATATAAATAATAATAATAATAATTTAAATA

ATAATGATAATAATTTAAATAATAATAATAACAATTTAAATAATAATGATAATAATTTAA

ATAATAATGATAATAATTTAAATAATAATGACCAAAATAAAGACACACATGATTACTCAA

AATTAAGTAGTTGTGATAGTAGCACCCAAGATGATTATTTAAAAACCACCTTAAATAGAA

TAAGCAGTTTTTTAAAAGAAGAAGAAAATATTATTTTACATAATTATAATTTTGAATATT

TAATAAAACCCTTCGATTTAGTATTACCAGTAGAACAATCAAGTAATAAAAAAGAATTAA

AAGCTAAATTAGAAATAAGTGATAAATGGGAAGGTATTACTCTCACGAAGACCCAAATAA

CGAAAATTATAGAAATAATGAATGAAGCTAATAAATCTAGGAATCAAACAAATAAATTAT

TATTAAAGCATGCATGGACAGTTCGATTAGATATCGAATCGTTAAGAAATGAGACGAAAA

ACGAATTTATAAATCTATATAATAAAGTATTAGGAGAAGAATATAATATAAGCAATACAG

AATTATCTGCACAAGAAATTAATAGATTACAAATTTTATATGATGTTGTGGGTGTTCGAC

ATTTGGCAAAATGGAGATTACACTGTAGAACCACATTAGAAAAATTTATTGAAGAAAAAA

ATTTAAAGAAGAAATTTTTATATGATTCAATATATAAGCAGCAATCTTGGTGGTCTTGGG

TAACAGGCAATAAAAAAGAAATTGAGAATAAAGTACAAAACATTTTAAATAGTGAACAGG

ATATAATAAATGAAAATGAGTTGTTTATGATACAAGAAGCTATGACAAATGATGATAACT

ATGATGTTGTTATGCCATCAAAATATGATTTTCAATTTAAACTAGCTAATTTTTCGATAA

ATGTATATGATGATTGTAAAAAGAGGATAATAGATAATAGAAATGTGAATAATCACCATA

ATAATCATAATAATAATAATAATAATAACAATATATATATTAGGGGGGTAAGTAACAAAA

GTGAAGTAACTAATAATAGACAACATATAAAAGAAGATAAATTAAATTTTAATGATATTA

TAAAAGATGAATTAAGTATAAATAGCAGTTGTGAAACCTTATTGACAAGTTCCATAAATT

TAAGTTCCACTGATTCCAAATTAAATAAAAGACGTGTAAGATATGAACGGGTAAATATCT

TGTCTATAAATTTTTATCAAATATATTCTTCTTTATCTCTACAATCAGTTGTTGATCATA

ATGATCATGATAATTTCCAATGGAAGTTTATTATTGAATTACAAAATTTTATAGCAAAAC

ACAAAAATAAGGTATTTATGGAATTTAGGAATAATAAAAATTCCTTTCCATATAATATGA

ATACATCAGGAAGTCTATATAAAAATTCTATTTTCTATTCATTATTATATAGTCAAACTG

TTTGCGCCTATTTAGAAATTAACCATTTAGTAACAGAAAAGGGTAATACATTATCCACTA

TATTAAGACTGAACCCTTTGGAGCTTTATTTATCTCCATTATTAATTAAAAGTATATTGT

CATTTACATTTCCTCTATTAGATATAATTAATAAAAGTCATAAGACAGCAATGAAAAAGT

ACAAGAGTAAAAATGTGAGTTTATCTGTGGAAAAGCAAAAACAAAAGGAAGAAACTTCAC

CTGGTGTTATTAAAAAGAAGGAATCAATTTCGGGCACAAGAAAAAGAAGTAATAAAAATA

AGAAGAGCCATCATGAAATTGGAAGTATAAAGAAGGATGTAAATAATGAAGAAAATGATA

CGATAAAGCAAGATGATCATAAAAATATGAATGCTGATAGTAATAAAAATAATGATAATA

ATTATGATAATAATGATGAATGTGATGATAATAATGATGAATGTGGTGAATATAATGATG

AAGATGATGAGGACAAAAATAACGATAATGAGGATAATAATAACGATGATGATGAGGAAG

ATGAGGATGATGATGACGAGGATGACGAAGAAGCATTATTACAAATAAAAAAAATGGAAC

AATCCGAATTATTAGAAGGATTAAAAGAAAGAGGAGAAAATGTTTATAATAGAGCTGTTC

AACACCTTCCAGAATTATTTGAGTTTTATATACATATATGTGGTCCCATTTTACATTTTG

ACAATTTAACGAATGGCATAGTGGAATTGCATTTAGGTAATTTGGTAGCAAAAACAGAAC

ACCCATGTACCTATAATAAATTTAATTTAATATTTGAATTTAATGAAACGCAAATAACAT

GCTTAAAAATAGGCTCAGGATATATGAAGAAGGATGAAAAATGTAAAAAGGATATAATTT

TTAATGAGAACCAAAAAGATTTGCATAAATCAACAAAAGCATATAAGAAGATGGATAAAA

GTAATATATATTTAGAAATAAATGAAGAAGGATTATTTTCAAATAATGCAATGGATAATA

CCGAAATGGATGATTTAAAAGATAGTTATTTTAATAGTATAAAGGAAAAGGTATTAACAG

GTGAAAAATTTTATATATTACAACCAATTCCAGTAAAAATCTATGTAGAATATGACTTGA

AAATTTTGAAAACAAATATTATATTAGATGGAATATTTTTTCAAATTAATCCAGATGCTA

TTAGTATTATATTAGCAGTTCCAACATCGATAACTAGATACCTGACTGGAGTTTATTCTA

AAAATAAAAATGAGAATGAATTTAATAAGAACAAGAAGAAAACTTCAGTTGAAAAAAAGG

AGGGAGATATAATGAAAGGAGATGATATTACAAGAGCAAGTATGTCAAATATCGATAGTG

GTGTACAGAAAAGAAATAGTATTACTGATATTATAAAAGGGAATAAAGATAATGATAATG

TGGTAGAAATGAATAATAAAGGTATGCCAAAGGAAGAAGAGTCGTTTTTATATGATATTG

ATTTTTTAATTAAAAATTCTTCTTTTTCAATTAAGAATAAGAAAAATTGTGAAATATTGA

AATATGAAGCATGTGGTATATTTTATAAGAATTATTTACAAAAGAAGAGGAAAATTATTA

AAATGGAAATTGAACAGTTATGGATTTGTGATCCTAGTAATAAGCAACCTATATTTTTTA

CTTTAACGAAGGATATAAATTCAAAGGATATATTTTTATTTAGATCCTTATCTACATATT

TGGAAGAGAAAAGTCATAACAATAATAGTATTGAACAAAAGGAAATTGATATGGAAAGTG

ATAATCATATAAATGTTGGTATAGACGACGTATTAGAAGAAGATCATTTTATGATGGGCA

ACAACAATAATATTAATAATAATAGCAGTCATATTAGGAAGAATAGTTTTAATATAGGGT

CAAATGCTTTTGGTAGTTTAGAAAGATTCGAATCTGCTAAAGTGATGGATGATAATGTAT

ACACAAATGGAAAACATAAAGGTTCAATCAAGAATGTAATGATTGATAATTCTATTATAA

ATAATGAACAATTAATAAACAATAGAGCAGAAGGAGGAAAATATATGAGCACAAATAGAT

TTTCTGCTAGTGGAATAAATCAGTTTAATAAATATGTTGGTATGAATATAGATAGAAAAA

ATGAAGAGACCAAATTGAATGGGAAAAAAGAAAATGATGAAAATAAAAATGTTGAGGAAG

AAGATGATGATTTTATGGATGCTATTGAAGAAAAACAATTATCTATAAGTCTTCAAATTT

TACAAGAACATAATGAAAAGAATATTGAAGAAACTCATATTAATTTTATTATTAGTGATA

TCGAATTACATTGGAAATATAAAACAATAAAACAGATTTTTAAAACAATGAAAGAATATA

AGAAAACGTTACAATATGGTATAGAAAAAGATATGACATATATAAAAAATAAATTAAAAA

ATGAAAAGGATTTAAAAAATTATAAAATGTTCATATCTGAAAATACCTTAAGAAGTGTAC

AAGAAACTTTGAAAAATGTTAAGGATTCATTAAATATTTTAGATATGGAATCTGCAAAGG

ATAAGGATGATATAATAAATAATGTACAAGAAAAAGAAATGGACAAAGATAATGTTAAAA

TGAATGCAGAAGGTATACATCTTATGAATCCTTATTATGCGAGGGACGAAGATTTAAATA

TGATAGCTGATAATAAGCTGCTTTTTAATTTAAGAAATGAAAAAAATAATGAAGAAATAA

ATATTAATAATGCTAACACTGTAAATGATAATAATGGCATTAATAATAATGATGATATTA

ATAATAACGATGATATGATTAATAATGATGATATTATTAATAATCATAATTATAACATTT

TGAATGATAATATTCTTAGTAAGGGACTTAATGATACATTACCCAAATTACACGGTCAAC

AAAATTCATATCCCAAATATTTTTTCAACTGTTTTATAAAAAGTGCTTCCTTAGCCTTCT

GGAAGAAGAAGAAAATTTTTTCTAGAATTCAAGTGAGTAACATATTTTATGAAAATAAAA

TATATATCAATTTTGATCAGAAGATGTTTTTGAATATAGAAAAGGGGATAATGTCAATGA

ATAATAAGAACATTATATCTAATAATATTAACGATTATAAATACGACCTGTTTATTTCTA

AAGATAGGAAAAGATTTGGTTCAAGTGATATCGAGAATGAGGATAGAAATATGTTAAGTC

ATTTAAAATATGGAAAAAAAGAAGGGAACTTAAATGCAATGCAGAATGATGATGATAAAG

AAAAAGAAGAATATACAGCAGAGGATACATTTTCGGAAAGACGAAATAGTCGAGATATAT

TAAATAATATAAATCTAGATAATTTATCTATAGAAATAAAGAAGAAAAAGAAAACAATGC

ATAAAGAATCCAAAGAAGATTTGTTTATTGGTAAGATAAAAGTATATAATGATAAGAGGA

ATTATAATATATGTTTATTATGTGAAATTCCAAGAATGTATTATATTTTTTACCTGAAAG

ATTTAAGATTATTTTTGGAATATTTAGATGATGGTATATTAAATGTTTTTATTAGTAAAT

CATATAAAAAGGTAGTTCAGGCTGCTCAGACCAAATACTTTTTATTTAATTTTACATTTA

TGGATCCTATAGTGATCATACCTGAAGATAAAAATATGATATATAATTATAAGGGAGAGG

TGAAAAATGTTTCTAGATATTTTAAGAGTAGCACTAGTAATAATAACACAAAGGAAAATG

AGGACACAAATGATATTGCAAAAAGAAGCTCAAAGGAATATATTCTTTCCAATGAAAAAA

AAGATACAAAGACTGAAGCGTGTAACAATTATAATAATGTGAATAATAAGAATATACATA

ACACTAATTTTGATTGTGATATGAAATCCAATTATGATAAAAATTCAATTGGTCGTAATA

CATATAATAGTAATAATATACCATACATTGAATCTTATTTACAATTCCATTTAAGCAAAT

TAAAACTTAAAAATTCTTACACTCTAAAATGTACGGAAGAAATAAGAGTAGCACAGAGAA

GACAAATGATATTAAAAAAAAAGAAATATAAAGGAAGAAGTGAAAAGGAAAAAATAAATA

AAAATAAATATGAAAATGAATCTTTTATAAGAAGTGATATTGTATCTAAGGAACAATTAA

GACCATTAGAGCGAAAAAGGAATAGTGTAAATAATGATTTGGTGAAATTAGATTTTACGT

TATATATTGATATATTAGATATAGAAAGTAAAGCTTGTGAAAATGGGGGTAGTAGTAATA

TAGAAGGGGAAATCTTGCATAAGGTAAATTTAGGTTTCTGTTTAATAAATGCTAGGAATG

GTATATTTATATATTTAAATGGAAATGACTTATGTTTAGATTTAACAGTATTTCAATTAG

CTTTTTTGTTGGATATAATAAATGAAAACTTTTGTTATAAGGGATATTTCCCAATGTGCT

TTATATGTGATAAGGATGTGAGTTTAGACAGTATGATAAATTCGGAGTGGTTTAAAAAAG

GATACATGAATTTTAACAATTTTAATAATAGGAAAGAAATGAAAGGAAAAACAAATGACA

ATAAAAAGGAAGATATAATGAACAATATAAATAATATCAATAATATCAACAACAACAACA

ATAATAATATCAATAATATCAACAACAACAATAATAATAATAATAATAATAATAATAATA

AGGTAAATAATTATGAAATTGGAGATGGTGCTATAAATAACAATGTGTATGATAGCGATC

GCACTATTGATAAGGAAATGGATCACGGTAAAAATAAATTGGATATGTCAATGAACCAAG

AAAAAAAAAGTGATTCATTAACAAATGATAAACTAATAATAAAAGATTCAAAAGAGAATG

TAACAAAAAAGACAGGCTTGAAATTATATGTATATATAAATTTTGAAAGTTTAAAAATTA

AAACAGCATTTGATACAAATACACCTGTAGCCATTATAACATTCCAATATATAAGTATGT

CATTTCGTTTAGTTCTCTTAGATTTCTATTATGTTTACTTTTTTGATTTACATGGTAATT

CATTGTATATAGATGATGCTAGGAAAAATTCAATCAATTATTACAAAAGGGTAGCCTACT

GTTGTATTGAAAATGAAAAGAAAAAGTTCAAGAGGGAAAATAAAAAAGCGCATGAGTTGT

ATAAAAATGTATATAATGAAGAAGCATATAATGATACTCTTAATAAGAGTAATGTATATG

AGTATAATAATGTGGAAAGGAATTCGTATCAAATTAATAGAGAAAATAAAATATATGATA

ATAATGGGTCACATTTTTATAATAATAAAGATGTTCAAGATGAATATGATGAAAAAGTAC

AAAAACTACATAATAATAGTAGTAATAATATTTCTGTTACGACACAATCTGATATAAAAG

ATTATAAAAATTTATTAATATATATGTTTGAAAATAATGTATTTTTAAATGAAAAGAAGA

AAAGGAAAAAACAAAAAGGAATAAAAATAAAAATTAATTCTTTCATAGAAGATTTGTTAT

TAAATATTGAACTTGATGATGCGTATATTTGCTTTTTTTTCCTTATTTTTATAGATATAT

ATAAATTTTTAACGACAGGATTTAATTTGAGCACGTTACATTTGTATCCGAAGCCATCAC

CTTATATTGTATCTTACAAAAATGAAAATAAAAAGAAAAAGATAAAGGGTCATGAATATG

TAGAGAATAGTGAAAGTAAAAGTATCCATAGAAAAAAAAAGGTTGTGTGTTCGAAGACGG

AGGAAGAAAAACATATAAAGGAACAGATTAGTATTAGATGTACTAGATCTCAATATAAAA

ATACCTCAAATGAAAATAAAAATAAAAATATTAATAATAATAATAATAATAATAATAATA

ATAGTAATAATAAGAGTAGTATCAACTGTAAGGAAAATACCAGTATAAATATCGAAGAAA

ATGTGTTTAAAAGAGAAAGAATTCATTATGATGAAGAAATTATTGAACAAAATGTTCAAC

TCATCAATGAAGAAGACAATGAAAAACATTGGGTACAAAAAAAGAAGAAAATTATTCGAC

GTAGTAGTGTTTTAGAACAATCAAAATATATAAAAGGGAAATTAAAAGAACAGAAAAATA

AAATAATAACAAATATAAACGAAGTATTAAAATTAGATAATAAACCATTTCATGTAAGTT

TTAAAGTGAATAATGGGAATTTCATTATTTTTACAGATGTCGAAAAAGTATCTCATCCAA

TAATAATGTGGTCAAATAATTTTGTTTTTTCATTTTCTTTATTTAATAAATGTATTGTAT

TTAGAAAAATATATGCAATAGATAGTAAAGTGAAGAGGATTAATTATGTATCGTCATCTC

ATGTACATTATAAGTATGATACTATTAAGAAGAGGAACAATAAAAGCAACCATAACAATA

AAAATAACAATAACAATAAAAATAACAACAACAATAAAAATAACAATAAAAATAACAATA

ACAATAAGGATAATAATAAAAACAACAGTAACGATAATAATAACATCAATATGGACAATA

ACAATAAAAATAACAATAACAAAATATTTGATCCTAAGAATATATCGAACCCACTTCATA

ATTATCATAAAAAGAAAATATTACTATGTGATAATTTGAATGTAATGGGTGAAGCTGTTT

ACGAATCAGTTGATACAAGAACTGAGGATTATAAATTAAAAAGTAGTGTAATAAGAAAAG

ATAACAATTCACCATTTATAACCGTTTTTGAATTAGATATAAATATTGGTAATTTTGATA

TACGTTTATCAAATGACGATGTTGAAATATTATTGAAAGCATCATCCACATTATTTGGTG

ATGTCCCTAGTTCATTTACAAATATAGTAGTTGGCCCAGTTATACCACCGTTAACTTTTA

TACATAAAAAAATAAAGAATAAATTTATGCAAAACAATATTTATTATATAAGTACAAATA

TCAAGGAGGAGACCGAATTGTTTTGTGGATCCGATTTGCAAATAACAAGTATAAATATAT

TAAAGAAGAGAAGGAAAAAAATTATAAGAAAGAAGGAAAAGAAAGATGCAAGAAATAGAA

AGGAAATGGAGAGAAATGATCATAATAAAATAAAGAATATAAGAAATGATGAAATTAATA

TGAATGAAAAGAAAAGTATAGTAGATTGTCATATTGAGGAAAAGAAATTAGATCAGAAAG

AATATAACGAGTTTAGTAATAAGGATGAAATATATACTGATTCTAGTAACATGATGGAAT

ATGTATATTATAAAAAAATATCTGATGTCTCTTCAAGTACTAATACTTTAAGTTATTCCA

ATACATCTTTAAGTGAATTACATGGAGAAAAAAATATATTATATAAGAGAGGTATGAAAT

ATGATGATATATCAGATATAAATAATAACAGTAGTAGTAATAATAACAACATCAATAATA

ATAATAATAATAATCAAACAAATAGTTATCGAGATATAAAAATTAATATAAAGTTACATA

ATGTAATGTGTACCTTTATAGATAATATAAGAAATTCCATTGTTCCCATTTTAAGAATGA

TATTTAGTATGAACATAAGTATTAATTTATATACTGATGAATGTTCATATAATATTAGTG

ACTTAAATTCTAAGTTAGAATATTTTAATAATTGTATAGGTGAATGGGAACCATTTCTTG

AGAAATGCAATATTTCATTAGATATACATAATATTTTTCCAAATGACGAATATGATGAAG

ATAAGGAAGCAAACAAATCTCCTATAAGTATCATTAAAATTAATAGTATCAAAGCTTTGT

GGTTTAATATTACACCTCAGTTAATTAATTTGTTATTTTTATTCGTGCCAGTCTTTTCTG

AGAAGGTATCTAATGGGTTAAGAAAAAATGCCAAAAAGGAAATAGACCATTATAATGATA

AGGATAATAAATATGTAAGTGAATGTATAAAAAATGATATATTAATGGAAGATATGAATA

ATAAATTAAATGTTAGGAATACATCTATTAATAAGGATATAGAAACATCTGAACAGATGA

ATAGTCTAAATGACTTTTCCAGATGTGATAGTGGATCCGTCGAATTGGAATACAAAAATA

TGAGTGTTAAGAGTAAGAGCATATTCGAAGATTGTTCTTCGGTTTTTAATTCATTTCCGT

GTGATACAGATGGTAAGGAAGCAAATGATTTTCATTTGGATGACTTTTTAAAAAAAGAAG

AAAATCGTAGTACATACTATAATAGGATAGATAACTCAGTAATATATTACGTTAATTTGA

CAAGCGAATACTTTTATGCCTTTTTAATGCCTGAAAGTAAGATTGAAGAAATTATGAAGA

GAAAAAAATCTGTCACTATGAAAAATGTATACAAAAGTGTGAACGAGGCGGATGGGACAA

ATATGACTATAAAATCTTATAAATGGGATGAGAAAAAGAAAAAGGTAGAAAAGGGATTCA

TTCAAAGGGAGAATGTTACGAAGGCGAATGGAATGAAAGGAAATGATATAAAACAAAATG

GCATAAAACAAAATGAAATAAAAGAAAATATAATGATGGAACATACATTAATTAGTTCAG

ATGATAATAGTACCCATAATGAAGGGAATGAAGATTATATATCTGAGGAAGATCTTTTCA

ATATGATACCTGATGTATATGCAAAAATTATAACAACAAATGAATTAATATCATTAGACA

GATTGTTAATCAACGAAATAGAAAATAACTCCTTAATAGAAAAAAAATGTTTATATTTAT

ATTTGATACCCATTCCACCAACAAATGTTGTGAATATAGTTCATGATATGTTTTCAAATA

TTAGTAAAAGAGATATTGTTTTAGATCTCATGATTACAAAAAATCCAAAGAAAACTATTG

AAAATTTATTAATTTACAATCAACATAAGGAACATATACGTATTAGGAGAAATAGTATAA

ATAGCATTTGGACGGAAAATATGATGTATGATGACAATAATAAAAAGAAACATATAGAAT

TGTATTTGAAAAAAATGCGTATGAAAGCGGAAGAGGATGGTGGTAACAAGGTAGGAACGG

ATAATATACATTTATTGGAAGATCAAAATATGTATTTATTAAACAAAGAAATGTGGAATG

ATTATAAAAACGTTAAGGGTATTATAAAAGGGGAAGGAAAATATACTAATAGTACTATTA

GTAAAACGGACCATACTTCAAGGAGTAACAATATGGACAATAATGTAAAAGGTAATGATA

ATATTAATGGTAATCACAGTGATAGAGAATCTGAAAAAAGACAAGACATGAATCAAAATA

ATGTAAATGAAAATATCTTAAATGTTATGGTAGAAGGAAACAATGAGGTATTAGATAATG

AGCAACGGGTTGATAAATCATTGAATTTATATACAAAAGGTTGTGAAAATGATGATGCTA

ATGAGGATGTCAATGATGAAGAAGAAGGGTCTTATGATAGTTCTTCTGATAGCTGTGAGG

ATTATGAATCAACTGATGATACCCATTCAATTGATGAAAGGGATAATTTGAATTATGAAT

ATTATTTTAAATATAATAAGAAAAAGAAAAATGATTCAAGTTCTATTTTTTCATCGAATA

AATTTGTTGGACCCTTTTTAAGAAACACAGAATGTGTAATAATAAATTTGATTAAGAATA

GTTGTACATCATTAACTACATCTTTGAAAAATAGCAACGTTATACATTTATTGAAGCCTG

GTGATTTTGTGTTTGATGAAAAATATAAATATAATTTTTCTAGCAAAAAGACTAATGAAA

TTATGGAAAATTATAGTATTACTGATGGGATTATGGCAGACGATGACATAACTAATGGAT

CCATAAATAATAATAATAATAATAATATGTGTAACAAAGGTATTTATGATAATACACCAA

CAAAACAGATGGTCGATAAATATGTTTCTAAAAATAGTATGAGTACTAATAAGAATAATA

AATCGAGTATATTAAATAAAGAAAATATTGACAAGATAAATGAATGGGAAAGAAGAAATG

AAGAAATGATTAAACAAACATATATAGCATGTAAATATAGATTGAAAAATAATTATTTGC

CACAAACATCGTTGGTTGAAAATGTATGTGAAATTATATCTCCTATGCCAAATTATAAAA

TATTATTTATGTCATCAACCGTTAGAATTATAAACAAATGTGGTATTCCCTTAGAATTTT

GTTTTTTTGATGGTAGTCGAAATCCTATATTATTAACGTCATTAGAGAATAGGACTATAC

CTATAAATACTTTATATCCTAATCATAGTGATTCATTTAAAAATTATAAGGTATCTAATA

GTTTGAATATTAATCCGAATATAAAATTAAGACAGAAAAATAATAATTTAAGTTTCACTG

TAATATTAAATCATGAGTATTTATTATCAGCACCCGAGTGTGTTTTTTGTGGACCTTCTC

ATGTGTACATGTCTTTTAAACCTATCAATGTGATAACAGCAGAGAATGAATATTATGACA

TAATAGCTGGTGGTCCCACCACAAATATTGCGAACAATAATAATAATAATAATAATAATA

ATAGTAGTAGTAGTAGTCATAATTATTATGGTAATAAATCAAATATTAATGATTTATTAA

AAAAGGACAAGGGGGGAAGCAATGATTTAATAGATCCTATGTCTATTAATATAGAAAATG

GATGGTCTGATATTTTTAGTTCTGATATATCTCAAGGGACGTATGTGAAAAAGTGTAAAT

ACAAAGATATGAATTCCTTCATGTCTAATATAAATAAAAGTAACGGTAGTGGAAATAATT

TTTTTTATTTCTTGGTAAAGATTGAAAATAAGATAAGTGCTTTACCTGCTGAGAAAAATT

TGAAAATTATTACTATATATCCTCATGTAAGTGTTGTAAATGCTATACCAGCCTTTGTAG

ACATAATAATAACATCTGACGTAGTAGAAGAAAAGCAGAATGATTATATGTATGAAGAAA

GGAATAAAATAGATGCATTAAAAAATAAGAGAGCAAAGTTATTAAAACAGTGTGAGGATA

TAAATTCATATTTGAATAAGATGAATTCATCAAAATATGGAGGAAGTAAAATTTATAGTG

ATAGTTCATCTAGAGTAAAAGGTAACAAAGATAATAGTAATAGTAGTAATAGCAATAATA

GTGAACATGTATACATGCAAGAGTTGAACAAATTAGAAAGAGAAATATATCATATAGATG

AAGAAATAAAATATAAGAAAAAAGAATTATCTTCTCCATTAGTAAAAGGATATGTGAATA

GAAGATTAAATCCATTTTCTATATTTTATATATATGAAATTAAGAAATATACTTGTTTGA

ATTTAAAGATGAAAATAGGGAATTCACAATGTGAATGGTCTGAAAAGTTATTTTTAGTAG

AAGATGATGAAGAATCTGTAACTAGATTTTCATTACATTTTAAAAAATATGCATCAGTCG

AAGTTGAAATTATTAAAAACTTTAGTGGATATTTTAATTCTTTGAGTAGTATCTTAGGAA

ATAAACAATTATATATTTTTTCCTTACCAAGATGTTTTATAGATAGAACAGGTTTAGGAA

TTAAGGCAATAAATTCGAATAAATATTATCCTGTGATTAATGGAATAACCTTATTAGGGG

ATCATTCCCAGATTGATTTGTTATTACCTCACAAAACTTATACTCAACCGAACAATAATG

TGAACAGTGGTAATAATAATGTTGAAGATGATACGTATAATTATTATTCTAAGGGTGATA

TGAAGAGATCGAATAGGATGTATGATAAACGAGGAGATCATAGAAAAGATTATAACGATA

CAAACGAAAGTTATGGTAATGTAGATGATGTTAATAAAAATGATAGTCGTTATGTTAGTA

AAAATGCAATTATAATAGATTACAATGATATATTTAATGATTTTAATTACCCTTCACCTG

TAAATAATTCTGTAATATTATTCAAAGCTACATTACCTCCGATTGGTAGTTATACAGAAA

CCAATGTGATTTGTAAAAATTTTTTTTATACCTTTTGTTTAAATACAGAAAAAATAAAAA

CAACGAATATTCCATATATTATAAGTAGAATAATTACAGTGGTTCCACAATTTATTATTA

GTAATAAATTGAATTTTCCCTTATTAATTAAACAATATCAGAATGACCAGATGCAAGGAG

TTCGTGCAAATGATACATCTCCTTTATATTTTACCAAAAAAAGTAGCATTTTACTATTTC

AATTTAAATGTTTAGATCCATGTAGAACTAATAATGAGGAGAAGAACCATATGTTATTAG

ATCCGAAAAATGTAATAAAGAGTAATGGAAATAAATATTATAAGTTGAATAAAAATATAT

ATTGGTCGTCTGTTATATATCCTAGTGAAAATTTTGTTGGAACAAATTATATGGTAATTAATACAGGAAATCACGAAAAATCTGATGTATACGCGATAACGTGTATACCTGATAGAGGTACGAAAAATATTATCATAGAAAAATTAGAAAATAAGAAGAAAGGTTTTATAGCATATAATAATTCAAGTATTGCAAAATATTTAAAGATTAGAACATTCCATGATGATACGAGACATATGAAAATTCACGATGAAGAGAATTATATGGAAAAGAATATGTTTTTAAGTAATTTTTTAAAACCAACAGATATGGAACATTATTTTAATATAGAGCCAAATAAATATAGTTATTTGGGATGGGTAAATCCATTTATATATGTTACTAGAAATGTACAGATAGAAATAGTTTTGGAAGATTTAAAAATAATACCTAAATCTCCATTTGTTTTAAAATTTGCTATATATAATTATTCACAAAAAACATATTATATAAATTATTATAATATTACATTTGTTATATGTATAGAATATATAGAAGATTTGATAACCATAAAATTGAGCCATAAATTAAATATAAGCTCAAATCTATTAGATCGTTCCTCTTATTATTATATGATGAATAATAAAAGTAATATAACAGATGAAATAGCATATTCTAATAGTTCACATAATAATATTATGTTCAGCAAACATATGAATAGTTGTGACAAATCTGTTACATCTAATAATGACATTAAATTAGGATTAAATAAAAAGAAGAAATATATGGAAATGGAAAATCAGGTAAATATAGAAAAATATAATATAATAAATGGTAAAGATATTTCAAATGAAAGGAATATTATAGATAAGAATTCAGAGAATAGTTTAAGTTTATCAAAAAATAGTGTTTATGCCCAAAAACAAATGATTATAGATCATGATGAATATCATTATGTAAAATATATAGATGAGAAGGATGGGGGTACTGGTAGTTATAAAAATAATAGTATTAGTAGTAGTACTAATAATTATTATAAGAATGAATCTGTGAAGATGATAAGTAATACATATAAAAATGTGCATATAATAATAAATGTAACCCAGATTGGTGTAAGTATTATATCTAATATATTAAAAGAGGAAGTATTCTTTATTGAACTATCAAAATTGTGTGCTTTGTTTTATATGAAAAATGAAGAAGAGGTAATAGATATAAAAATAACAGATGTACAAATAGATTGTCAGTTGGAATCTTGTGAAAAGTGTGTGTTGTTAGCGAATAGGGGAATAAGTTCTTCAAATAATAATAATAGTAGTAGTAGTAGTAATAATATGAATAATATGATAAGTGGAGGGGATATGAGTGGTAGTAATAATGTATATCAGAATTATCGTAATGAAAATAAAAGTTTTACAATTAATGAGAGTGACCTCCTTCGTTTTAAGAAAACATTTGATCGTAATTCAAACGACAAAAATAATCATATGAATCCTCTATATGATAATAATAAGAATAAAGGTACTACTACTACTACTACTACTAGTGGTAGTAATAATAATAACAATTATAACAATAATAGTAGTAGTGTGTTAAATAAAAATATGAATGCAAATGAAGAAAAGATCTTTTTAAATATATATGTAGAAAGGTCTTTTATATCACATAATGATGTTATATTTAAAAAAATACAAGTTTCGTTAGATGATGTAGAAATAGAAATGGATGCTGAAACTTTGAATGGTATAAATTTATTAACAGCAGAATATATTGAAAGTATAAGTATTGTTCAAAAAAAAAATTTGTTATATGAAGAAATACAGAAATGGACGATTTTACCTGTATATGTAAATTACAAATCACCAGAAATACCATTAGCCATAAATATCCAATACATGCAAATAGATAAATTTACCTTAATAGTGTGGTGTTCATTTTTATTAGATAAAATGCACATGATGAGTGATTTGTTACGAATAGGATTACGAATATTAATGGTATCGGGAAAATTAGAATTATTAGGTGCACCTGTAACCTTAAATCAAGAAATTTTTAATAATATTCGT**GTAAGTATTAAATCCTTTTATGCATTACTAAAAGATAAGTATAGTCATTCTATTTTGGCATGTTTAGGTTTTATAGTAGGATATAGTAGTTTAATAAATATACCTAAGATACCATTAGAAATTGGTCGAAATACTATAGGTTTAGCTGTATATGCAGTAGATAATGTGAGTGTTGGAATTGGAAGCTTTTTATCAAATTTAACATTTGATTCCGAATATATTAATAGAAGACAAAAGGAGAGAACCTTTAAGACGAACACCAATATGAAAGAAGGTTTAATTAGTGCTGTTAAAAATATTGGTGAAGGAGTATTAAGTTTATCTAATATTGTAACGAAGCCAATTGAAGGGGCACAAAAAGAAGGATTTGGTGGATTTTTCAAAGGAATAGGAAAAGGTGTTGCAGGTTCTTTAGTAAAACCATTAGATAAAGTAGGACAAGCTGTATCTGATGTAACTCGTGGAATAAAAGCAGAAGTATCGAAACCTATAGGTGGTCATAAATATAAGACTAAAAGACACAGAAAACCAAGAATGCTATGGGGAGAATATGGAAAATTAAAAGAATATAATATAAACGAAGCAGAATTAAGAGAATGCTTAGGTTTGAAATTTTCAAAAAATATTATGAAATGTTTAACAGTACATAAACAAGAAAATCATCCTCCATCACATTATGCATTATTATTATATCCAAAGGTAATTATATATGCAAATCTATATGCTAATACTAATGCTCAAAAGAATGATTTCTTATTATCCGAAAAAAAAACAGATATAGTTATATGGTCTATAAAAATAGAGGATATAACAGAAATACGAGCATCTAGTCATGGACTCATAGTAAGAACGAATACAAGTACAAATACGGTTTATAAAATACCATGTAATAATGCTATATTGATAAATAAAATATATAGGGAATTGCATAACTCAAAAAATTCAATAAACTCTACTGTTATTTTGGGCGCCAATAATTCTGCCCTGTACAAATATTTA**CCTAGGTCAGGATTGAGATCAAGATCTGCTGCTGCTGGTGCTGGTGGTGCTGCTAGAGCTGCTctgcagAGAGGAGTACAAGTTGAAACAATATCACCAGGAGATGGTCGTACATTTCCAAAAAGAGGTCAAACTTGTGTTGTACATTATACTGGAATGCTTGAAGATGGAAAGAAATTTGATTCATCTCGTGATAGAAATAAACCATTTAAATTTATGCTAGGTAAACAAGAAGTAATACGAGGTTGGGAAGAAGGAGTTGCTCAAATGAGTGTAGGTCAAAGAGCAAAACTTACTATATCTCCAGATTATGCTTATGGTGCAACTGGACATCCAGGTATAATTCCACCTCATGCAACTCTTGTATTTGATGTGGAGCTTCTAAAACTAGAAACTAGAGGTGTTCAGGTTGAAACAATTTCACCTGGAGATGGCAGAACCTTTCCTAAAAGAGGACAGACTTGCGTAGTTCATTATACAGGCATGCTAGAGGATGGTAAGAAATTTGATTCTAGTCGAGATAGAAATAAGCCATTCAAGTTTATGCTAGGTAAACAGGAAGTAATAAGAGGTTGGGAAGAGGGTGTAGCACAGATGTCAGTTGGACAAAGAGCAAAGTTAACAATATCACCAGATTATGCATACGGTGCAACAGGCCATCCTGGCATCATCCCTCCACATGCAACTTTAGTATTCGACGTTGAATTGTTAAAGTTAGAGACAacgcgtGCTAGAGGTGCTGCTGCTGGTGCTGGAGGTGCAGGTAGACGTACGATGGCAGAAATTGGTACGGGTTTTCCATTTGATCCTCATTATGTGGAGGTGCTGGGGGAAAGGATGCATTATGTTGACGTAGGACCAAGAGATGGTACTCCAGTGTTATTTTTGCATGGAAACCCAACCTCGAGTTATGTATGGAGAAATATAATTCCACATGTAGCACCAACACATAGATGTATAGCTCCTGATTTAATTGGTATGGGAAAAAGTGATAAACCTGACTTAGGATATTTTTTTGATGATCATGTCCGTTTTATGGATGCTTTCATTGAAGCTCTTGGCCTTGAAGAAGTAGTATTAGTTATACATGATTGGGGATCCGCCTTAGGATTTCATTGGGCCAAGAGGAATCCTGAAAGAGTAAAAGGAATAGCATTCATGGAATTCATACGACCAATCCCCACATGGGATGAATGGCCAGAATTTGCACGCGAAACATTTCAAGCTTTTAGAACTACAGATGTTGGTAGAAAATTAATAATAGATCAAAATGTATTTATAGAAGGAACTTTACCTATGGGTGTTGTAAGGCCGTTAACAGAAGTTGAAATGGACCACTACCGTGAACCTTTTTTAAATCCAGTAGATAGAGAGCCCTTATGGAGATTTCCTAATGAATTACCTATTGCAGGTGAACCCGCGAATATTGTTGCTTTAGTAGAAGAATATATGGATTGGTTACATCAGTCTCCTGTTCCTAAACTTCTATTTTGGGGTACACCTGGAGTTCTAATACCACCAGCTGAAGCAGCAAGATTAGCAAAATCATTACCAAATTGTAAAGCTGTTGATATAGGTCCTGGGTTGAATTTATTACAAGAAGATAATCCAGATTTGATTGGATCTGAGATAGCTAGATGGCTAAGTACATTAGAAATTTCAGGTACCGGTGCCAGGGGAGCAGCCGCAGGAGCAGGGGGGGCAGGAAGGCGTGGTGTTCAGGTCGAGACTATTAGCCCTGGAGATGGACGCACGTTTCCTAAGCGTGGACAGACATGCGTAGTTCACTACACAGGTATGTTGGAGGACGGTAAAAAGTTCGACAGCTCACGCGACCGCAATAAACCTTTCAAGTTTATGCTTGGCAAGCAGGAGGTTATTCGTGGATGGGAGGAGGGTGTAGCACAGATGTCTGTTGGACAGCGTGCTAAGTTGACAATTTCACCTGACTATGCTTATGGCGCTACGGGCCATCCCGGGATCATTCCGCCACATGCGACTCTGGTATTCGACGTTGAATTATTAAAGTTAGAGACAGCTAGAGGGGCCGCTGCAGGTGCTGGTGGAGCTGGAAGACGTGGAGTACAAGTAGAGACTATCTCTCCAGGTGACGGTCGCACTTTCCCAAAGCGTGGCCAAACCTGTGTTGTACATTACACTGGTATGCTGGAGGATGGGAAAAAGTTCGATTCCAGTCGCGACCGTAACAAACCGTTCAAATTCATGTTGGGAAAGCAGGAAGTGATCCGCGGGTGGGAGGAAGGCGTGGCGCAAATGAGCGTCGGTCAGCGGGCTAAATTGACCATTTCCCCTGACTACGCGTATGGGGCTACTGGGCACCCAGGGATTATTCCGCCTCACGCTACACTTGTGTTTGATGTCGAACTTTTGAAACTGGAAACTGTCGACGGAGAAGGAAGAGGAAGTTTATTAACATGTGGAGATGTAGAAGAAAATCCAGGACCAATGATTGAACAAGATGGATTGCACGCAGGTTCTCCGGCCGCTTGGGTGGAGAGGCTATTCGGCTATGACTGGGCACAACAGACAATCGGCTGCTCTGATGCCGCCGTGTTCCGGCTGTCAGCGCAGGGGCGCCCGGTTCTTTTTGTCAAGACCGACCTGTCCGGTGCCCTGAATGAACTGCAGGACGAGGCAGCGCGGCTATCGTGGCTGGCCACGACGGGCGTTCCTTGCGCAGCTGTGCTCGACGTTGTCACTGAAGCGGGAAGGGACTGGCTGCTATTGGGCGAAGTGCCGGGGCAGGATCTCCTGTCATCTCACCTTGCTCCTGCCGAGAAAGTATCCATCATGGCTGATGCAATGCGGCGGCTGCATACGCTTGATCCGGCTACCTGCCCATTCGACCACCAAGCGAAACATCGCATCGAGCGAGCACGTACTCGGATGGAAGCCGGTCTTGTCGATCAGGATGATCTGGACGAAGAGCATCAGGGGCTCGCGCCAGCCGAACTGTTCGCCAGGCTCAAGGCGCGCATGCCCGACGGCGAGGATCTCGTCGTGACCCATGGCGATGCCTGCTTGCCGAATATCATGGTGGAAAATGGCCGCTTTTCTGGATTCATCGACTGTGGCCGGCTGGGTGTGGCGGACCGCTATCAGGACATAGCGTTGGCTACCCGTGATATTGCTGAAGAGCTTGGCGGCGAATGGGCTGACCGCTTCCTCGTGCTTTACGGTATCGCCGCTCCCGATTCGCAGCGCATCGCCTTCTATCGCCTTCTTGACGAGTTCTTCTAActcgagggatatggcagcttaatgttcgtttttcttatttatatatttataccaattgattgtatttataactgtaaaaatgtgtatgttgtgtgcatatttttttttgtgcatgcacatgcatgtaaatagctaaaattatgaacattttattttttgttcagaaaaaaaaaactttacacacataaaatggctagtatgaatagccatattttatataaattaaatcctatgaatttatgaccatattaaaaatttagatatttatggaacataatatgtttgaaacaataagacaaaattattattattattattatttttactgttataattatgtgtctccttcaatgattcataaatagttggacttgatttttaaaatgtttataatatgattagcatagttaaataaaaaaagttgaaaaattaaaaaaaaacatataaacacaaatgatggtttttccttcaatttcgatatcaatttatagaaacaaaatatatacttgtataattttatttttttatataaatcattacatatataattatacaatattttttctaagagataattatatattaatatatataaaaaaaggtgttttttttttttttttttatttttatttttattttatggtaatattttattttccttattttataaattatattagtttatatgtgattaattttatatattatcaatttatatatttttaaatgcttacttaattatctttttttttttttttttttttttttcccctctttttatattaatttatttttgaaaaaattgatatatatatatatatataatatatatatatacatgtagtagtattaaacaatgtataatatatataaataatatatttatatatttcatttcaattttaattttttttggttttttttttttttctttttgtcatatttaaaaaaaattatattcatataagttatgcattttttataaacattattcaatatatgtataatataatatatatatatatattaatgtattattccaatgtgcatgataaaagaaaaaaataatatttataaaaaaaaagaaaaataaaacaaaaaaagaaaaaaaaaaaaaaaaaaaaaaaaatacaaaaataaataatataatttataattatatattcttgtcacaataaaaatatatatatatatatatatatttataatatgtatattttaaactagaaaaggaataactaatattttatttattatcattcaagatttatattttataataataaatacctaatagaaatatatcaggatccatgcatggttcgctaaactgcatcgtcgctgtgtcccagaacatgggcatcggcaagaacggggactacccctggccaccgctcaggaacgaatttagatatttccagagaatgaccacaacctcttcagtagaaggtaaacagaatctggtgattatgggtaagaagacctggttctccattcctgagaagaatcgacctttaaagggtagaattaatttagttctcagcagagaactcaaggaacctccacaaggagctcattttctttccagaagtctagatgatgccttaaaacttactgaacaaccagaattagcaaataaagtagacatggtctggatagttggtggcagttctgtttataaggaagccatgaatcacccaggccatcttaaactatttgtgacaaggatcatgcaagactttgaaagtgacacgttttttccagaaattgatttggagaaatataaacttctgccagaatacccaggtgttctctctgatgtccaggaggagaaaggcattaagtacaaatttgaagtatatgagaagaatgattaagcttatttaataatagattaaaaatattataaaaataaaaacataaacacagaaattacaaaaaaaatacatatgaattttttttttgtaatcttccttataaatatagaataatgaatcatataaaacatatcattattcatttatttacatttaaaattattgtttcagtatctttaatttattatgtatatataaaaataacttacaattttattaataaacaatatatgtttattaattcatgttttgtaatttatgggatagcgattttttttactgtctgtatttttcttttttaattatgttttaattgtattttatttttattattgttctttttatagtattattttaaaacaaaatgtattttctaagaacttataataataataaatataaattttaataaaaattatatttatcttttacaatatgaacataaagtacaacattaatatatagcttttaatatttttattcctaatcatgtaaatcttaaatttttctttttaaacatatgttaaatatttatttctcattatatataagaacatatttattaaatctagaattctatagtgagtcgtattacaattcactggccgtcgttttacaacgtcgtgactgggaaaaccctggcgttacccaacttaatcgccttgcagcacatccccctttcgccagctggcgtaatagcgaagaggcccgcaccgatcgcccttcccaacagttgcgcagcctgaatggcgaatggcgcctgatgcggtattttctccttacgcatctgtgcggtatttcacaccgcatatggtgcactctcagtacaatctgctctgatgccgcatagttaagccagccccgacacccgccaacacccgctgacgcgccctgacgggcttgtctgctcccggcatccgcttacagacaagctgtgaccgtctccgggagctgcatgtgtcagaggttttcaccgtcatcaccgaaacgcgcgagacgaaagggcctcgtgatacgcctatttttataggttaatgtcatgataataatggtttcttagacgtcaggtggcacttttcggggaaatgtgcgcggaacccctatttgtttatttttctaaatacattcaaatatgtatccgctcatgagacaataaccctgataaatgcttcaataatattgaaaaaggaagagtatgagtattcaacatttccgtgtcgcccttattcccttttttgcggcattttgccttcctgtttttgctcacccagaaacgctggtgaaagtaaaagatgctgaagatcagttgggtgcacgagtgggttacatcgaactggatctcaacagcggtaagatccttgagagttttcgccccgaagaacgttttccaatgatgagcacttttaaagttctgctatgtggcgcggtattatcccgtattgacgccgggcaagagcaactcggtcgccgcatacactattctcagaatgacttggttgagtactcaccagtcacagaaaagcatcttacggatggcatgacagtaagagaattatgcagtgctgccataaccatgagtgataacactgcggccaacttacttctgacaacgatcggaggaccgaaggagctaaccgcttttttgcacaacatgggggatcatgtaactcgccttgatcgttgggaaccggagctgaatgaagccataccaaacgacgagcgtgacaccacgatgcctgtagcaatgccaacaacgttgcgcaaactattaactggcgaactacttactctagcttcccggcaacaattaatagactggatggaggcggataaagttgcaggaccacttctgcgctcggcccttccggctggctggtttattgctgataaatctggagccggtgagcgtgggtctcgcggtatcattgcagcactggggccagatggtaagccctcccgtatcgtagttatctacacgacggggagtcaggcaactatggatgaacgaaatagacagatcgctgagataggtgcctcactgattaagcattggtaactgtcagaccaagtttactcatatatactttagattgatttaaaacttcatttttaatttaaaaggatctaggtgaagatcctttttgataatctcatgaccaaaatcccttaacgtgagttttcgttccactgagcgtcagaccccgtagaaaagatcaaaggatcttcttgagatcctttttttctgcgcgtaatctgctgcttgcaaacaaaaaaaccaccgctaccagcggtggtttgtttgccggatcaagagctaccaactctttttccgaaggtaactggcttcagcagagcgcagataccaaatactgtccttctagtgtagccgtagttaggccaccacttcaagaactctgtagcaccgcctacatacctcgctctgctaatcctgttaccagtggctgctgccagtggcgataagtcgtgtcttaccgggttggactcaagacgatagttaccggataaggcgcagcggtcgggctgaacggggggttcgtgcacacagcccagcttggagcgaacgacctacaccgaactgagatacctacagcgtgagctatgagaaagcgccacgcttcccgaagggagaaaggcggacaggtatccggtaagcggcagggtcggaacaggagagcgcacgagggagcttccagggggaaacgcctggtatctttatagtcctgtcgggtttcgccacctctgacttgagcgtcgatttttgtgatgctcgtcaggggggcggagcctatcgaaaaacgccagcaacgcggcctttttacggttcctggccttttgctggccttttgctcacatgttctttcctgcgttatcccctgattctgtggataaccgtattaccgcctttgagtgagctgataccgctcgccgcagccgaacgaccgagcgcagcgagtcagtgagcgaggaagcggaagagcgcccaatacgcaaaccgcctctccccgcgcgttggccgattcattaatgcagctggcacgacaggtttcccgactggaaagcgggcagtgagcgcaacgcaattaatgtgagttagctcactcattaggcaccccaggctttacactttatgcttccggctcgtatgttgtgtggaattgtgagcggataacaatttcacacaggaaacagctatgaccatgattacgccaagctatttaggtgacactatagaagcggccgcTAgGTAAGTATTAAATCCTTTTATGCATTACTAAAAGATAAGTATAGTCATTCTATTTTGGCATGTTTAGGTTTTATAGTAGGATATAGTAGTTTAATAAATATACCTAAGATACCATTAGAAATTGGTCGAAATACTATAGGTTTAGCTGTATATGCAGTAGATAATGTGAGTGTTGGAATTGGAAGCTTTTTATCAAATTTAACATTTGATTCCGAATATATTAATAGAAGACAAAAGGAGAGAACCTTTAAGACGAACACCAATATGAAAGAAGGTTTAATTAGTGCTGTTAAAAATATTGGTGAAGGAGTATTAAGTTTATCTAATATTGTAACGAAGCCAATTGAAGGGGCACAAAAAGAAGGATTTGGTGGATTTTTCAAAGGAATAGGAAAAGGTGTTGCAGGTTCTTTAGTAAAACCATTAGATAAAGTAGGACAAGCTGTATCTGATGTAACTCGTGGAATAAAAGCAGAAGTATCGAAACCTATAGGTGGTCATAAATATAAGACTAAAAGACACAGAAAACCAAGAATGCTATGGGGAGAATATGGAAAATTAAAAGAATATAATATAAACGAAGCAGAATTAAGAGAATGCTTAGGTTTGAAATTTTCAAAAAATATTATGAAATGTTTAACAGTACATAAACAAGAAAATCATCCTCCATCACATTATGCATTATTATTATATCCAAAGGTAATTATATATGCAAATCTATATGCTAATACTAATGCTCAAAAGAATGATTTCTTATTATCCGAAAAAAAAACAGATATAGTTATATGGTCTATAAAAATAGAGGATATAACAGAAATACGAGCATCTAGTCATGGACTCATAGTAAGAACGAATACAAGTACAAATACGGTTTATAAAATACCATGTAATAATGCTATATTGATAAATAAAATATATAGGGAATTGCATAACTCAAAAAATTCAATAAACTCTACTGTTATTTTGGGCGCCAATAATTCTGCCCTGTACAAATATTTATAAtaagaaaaaataaaatggcatatatatatatatatatatatatatatataatatatgttcatttatatgtttatttgtttatatatttatttacttatttattcatttatttatttattcatttattcatttatttattcatttatttattttatttattaatttattaatttatttattcattcatttatttattcatttatttattcatttatagttttattttcattatagcatatttccatattttttatttttatatatttaaacttgttttgattaaatatttaaaacttcttcaattctaatatatccattatttatctaattatatatagagagtgtatttttttttttttttttttttttttcttttttccagtttgccttcataaaaacaggcctatttaaataaaataaaaaatataaactatatatttattatatatacttttctatatgataaaatttgtttttaaatatatattattatgatattt

**5’ amplicon size: 1176bps**

**3’ amplicon size: 1499bps**

### **> PF3D7_1343800 (PfVPS13L2)**

tttattatggatccaaaggaaaaaaataaaaaataaaacaaaaaaaatattacatttata

tatatatatagatttttaaatatttgtatgtatatattgtttaatttttttaatttttca

tttaaaaaattttatttttatattgctaaaaatttactaaaatatgcaaaaaaatataat

gcatataaaatttctttaaataaacatattgtctgaatgtattaatttcttcagaaaaaa

aaaataaatataaaaatataattgtcaatatatattgataaaaaaaaaatataaataagt

gaataaatatatacatatatatatatatatatatatatatatatttttatgtatgtattt

atttgttgttatatttaaaaaaatagaccatagtaatgtaatataatataatatatatat

aatatatatatatttatttatttatttgtttatttttttatatttattaatttttgaggc

aaataaagaaatatgtaagaATGTTTCAGAAATTTTTTTCAGATATATTATACAAAATAT

TGGGAAATTTTATATATGGTTTAAATGATGAAGAATTAGCATTAAGTGATTTATTAAGTG

GGAAATTGGAACTAACTAATATTCATTTAAAACAATCAGTAATAGATTTGATAGATATAC

CATGTCGCTTAAATTTTGGATGTGTTGGTTATTTTAAAATTAATTTACCCTTATTATATT

TTATGAAGAATCCAATAAATATATATATGGAAGATGTAATAATAGTTTTGTCTACTATAC

CTTCTAAATATTTTGACGATAAGTTATATAAAGAAAAATATATTGAAAATAAAAAGAATT

TATTACTTTCTTCTGAATATAGAGCTATCGTTTGTTCAGCAGAGGGGGGTGTTATATGGC

AAATGATTTTGTCATTAATAAATAATATTAATATAACCATTAAAAATGTACATATTAGAA

TAGAAGATTTTACAAGTAATCCTTCCAACTGTTTTTCTTTAGGAATTAAAATTGAAGAAT

TATTTTTGAATATGCCGAAAGAAGGTTTACCTTTTAATAGAGATGGATATTATACGAAAA

AAAAAGAACTCGTCAGCAATACTATGATTAATATGATAACCGTTAAGAGGTTAGCGTTTT

ATATGGATTCTTTGGATATGAAAAAAATTACCAACAAAAAATATTTTAACAAGTTTAGAG

ATATATTTATAAATTATTGTAATATGGATGAAAGGAATTATCCAGAAATAAATATGAAAT

CAAATTACACAAACGACAGTAAAAATATAAAAAAAGTACATATGAATTGTGAAATACAAA

ACATGAATAATTTTAAACGTAATAGAAAGAATAAAAAAAATAAATCAAAACATAATAATC

ATAAAAATAATAATAAAATTAATAATAATACCAAAAAGTTGTTTAAAAAAAAAAAAAATA

AACGTTCTAATCAACAAAACTTAGAGAGCTTTCTCAATATAATTAAAAGTATTAAGAAAA

GTTCTTATTATGTTTGCAAAAATGATAATAATATAAATAAAATGGACAAAATAAATCAAA

GGATTTTATTATTAAATGATACAATTTCAGCAAAAAAAAAAAAAAATAAAATAATAAATA

ATAATATAAAGATGAAGAACACACATAATGAACAACAAAACAAAGATAATAATTGTTCAA

TATCTATAAGTAATTTCTACAAATTTAATCCATTCTCATCAAAGAAAGAGGAGAAAGTAG

ACATGCCTATATTGATTGATGATATAAAACTTGATAATGATATAAGTTGGATAGCATGTA

TTACATTGTTTGCAGATTCTTTGGAAAATTCTTTTATCAAAGAAAAAAAAGAAGAATTAT

TAAAAAAGCAAAGAAATAAAAAGAAAAATAATACAAATGATAAAAATGCACAACCGGAGA

AGGATGATGATGGTTTCTATCAAACTTGTAAGACTATTGATGAGAATAAGAATATTAAAG

ATCACAATCATTCTTTTAATAATAAATTGAACAAAAATAATGAACAAATTAAAAGTAATA

ATTATGAGAAGGAGAAGATGGGGAGTCATACTTTTATTGATAAAGATAATAATAATAATA

ATAATAAAAATAATCCTTCCTTTAATAGTGATATAAGTAATACCAGAAAACTTACTTCCA

ATTATTATTTAAACAAAGCTGTCTCTTTTCTTTCTATGAAATCCTTCACCAAAATACTTC

AGAAGAAAAAGTTATCAAAAAATATGCACCGTCATAAAAATAAAGAAAATGATTTTAGTT

CAAATGATCAAAATATAAGTGATGACTATTTTACTACAAAAGAAAAAATGTCCGATATAG

ATTTTAACAGTGAAGATATATATTCTTCTAAATATAAAAAGAAACGAAAGAACGAAATGA

AAAAAAAAGATTTCAATTTTAAAAAGAATGGTTATTCGAAATATGAAAAAAAAGATGAAG

AGAAATATTACGACGTTAAGGATAAAAACCAAAGTGATGAACATATTTCTACATTTCATC

ATTCTAAAAATACTGACAAGAATTATTTTAGTTTTAGTGATAATGATAAGAATAATAAGA

AATATATGTCACAGAACAAAAAAAAAGGAAAAAAAAAAACATATATTGAAAAGCAAGAAG

ATTATGGAAAGAATTATAAATATAAAACGAGAAACGAATATGACAAAACAAATAATGAGA

GTTATAAAAATCAAAATTTTAATGATTCAGCTCATAATTTAATATCTAATTATGAAAAAT

GTATACAACTGAATGATAAGAGGAAAGAAATAAAAATGGAACAATATAGAATTATGAATT

ATATCAAAAAGAACGAAATAAAAGAAAAAATAAGAAATAAGAAAAGAAAAAAAATTCATT

CAGTTTATTTGATGTATGAAAAAACATACAAAATAAATAAAAGTAAATTGAATTATTTCT

ATATATTTTTGAGGCTTACAAAAAATATCAAGCATGAATATATTATTAACCCAAATTATT

TTGAAGGATATGTAGAAATATATTTAAGTTTAATTCCAACAGATCATGGTGAACGTCCAT

TTTCTAGATGTTTTTCATCTTGGGGTATACAAAGAAGAAGTCAAAATGAATTTGATGAGT

GCTTACCTAAAATAAGTGTTTTTGTTACATTAAAAAATATGAAAATTATATTTTCCGATA

AACAAATGTCGAATTTAATAAAATGGTTAAATTTAAATTTTGTAACATATTCTACTTGGA

AAGCAGGTATCTTATCTCAATTTGAGAAGCCTAAAGCAACACCTAATGAAGAAATGTGTT

ATATAAATAAATGGGTTCTAAAATTATTAGATACTCATTCTTCAAAAGAAGAAAAGGAAG

ATGCCGAAAAATATTGTGAATTATTTGAAAATACTCATTCTATACATATTATAAATTTGC

TAAGAACTAAGGCATTTTCAAAATTACAGGAATTAAGAAAGGATATGGAAACAAAAATGA

AACCAAACAATAATAATAACAACAATAATAATAATACGTATGATATGGTAAATGACACAA

ACCAATATAATGAAATAGGCAAAAAAGAAAATGAAGTGAAACAAGTTGCTAATGAAAGTA

TTAATGTGTTAAAAAATTTAATGAAAAATGAAAAAGCATTTAATAAAATAAAGACATCAT

TTAGAGATAAAAATTTTACCATAGACTTTTGTTTCGATATATGTATTATTAATAGCACAA

TTTTGTTAACTATTGAAACATTTAATAAAATAAATACTAGAAAGGTGAACTTTGTAAAAA

CATATGCTCTATTATTACAATGTATACATTATCATAATCAAATTTCTAATACTCAAGAAT

TTAAGAATTCCATGGATATCGAATTATATCCATTAACATTACTTTTAATAATTAACAATT

TAGATAACAAATATATTAAACCCGAAATTAATGTATTATATATTTTAACCAAATCAGATA

AGAAGGATAAATATAAATATAAAGATAAAAAGTTTCTATTCAATATGGAGAACTTTGATA

TATTAAAAAGATACAAATATATTAGAAATACATATCTATATCGAAATGATAATCAAAAGA

ATTCAAACGAAGACAGGCAAAAAATTGAAAAGGAACAAGAATTATATAACCATGAAAATG

AAAATGAAAATAATCACAATAAACAAAGAGATACATTAAAATTACATAATTTAAAACAAA

AATTTAAATATTTAATGGAAGAAAATTTTTTCTTTTCAAAAAGACAAAATAATATGTATT

GTAATGAATTATGTGATATTACATCTTTGTCAGATCCATCAATTTATTTACGATATGATT

CTCAATCATATACCATATTAGAAAAACCTGATTGTCATTTGTTTTTACATAATTGTGCTT

ATTCAATATTTAATATAAACGGAGAAATAGCTAATTCATTTGTTAATGATTATTTCCATT

TTGTAGATAATAGATCAGTTTTATTTAAAAGGAAAACAAATAATTTTTCTTTTAAAAAAA

AAGAATATTTCTTAGAATTAGGAGCTAAATATTGTAATGATATAATGGATAATTATACAA

AACATACTAGTTTTTTCATCGATATAGAGTTAGAAAAAATGTTATCATTTATTATAACAA

AATCAGGACAAAAAGCAGAAATTCAGGGTTATAGCCTTAATTTAAATCCAGTAAAAGTCG

TAAGTCAATTATCTAAAAGATCAGAATCGTATAATATAGATAATTCCAAAAAAAATATAT

ATGATTCTTTTTTTGTTCAATTTGAAGGAATAAATATAAAAAGAATATTAAATTGGCATG

AAGCAGCTAAACATTATAAAACATATATTGATGATAATTTTATTAAACCTTTTCATGTCC

TTTTTCATCCTATAAATAAAGATATATATACAAATAGAAACCAAACCCCAGATAAATTCT

TTTTTAATTTAAATCGTAAGAAGAATATACCTCAATTAAATGAAAATGGAAATAGTAATT

CAAATGGTAATTCAAATGGTAATTCAAATGGTAATTCAAATGGTAATTCAAATGGAAATT

CAAATGGAAATTCAAATAAAAATTCAAATAAAAATTCAAATGAAAATTCAAATGAAAATT

CAAATAAGGATAACAAAATATCATACCAAAATGATAATATTTATAATAACAATAGTAATA

ATATGCATAAGAGAAGTAGTTCTTATAAGAATTTAAATAATGAAACAATAAATTATAATA

ATGATGAAAAAAAAAAGTCGGAAATTTATAATCTTATATATAATACTAGAAAATCACTAA

GTTCATATACACATTCCGAATCTGTTTCAAAAAATCATATATACGATAATAATAAAAATA

TATATATAAAAAAAAAGAAAATGAATAATAAAAAAAAGAAAAGTAAAAACAAAAAAAACA

AAAAAACAAAATTAAATAATATTAGAAGATCTTCAAGTTGTCCACCTCCTAAAAATATAA

ATATTTATTTTGATAGTAGTAATAAAAGTGATATATTATATAATGATAGAAAAACCACTT

TATATTCATTAAATTATATGTATGAACAAAATAGACAATCTACTATAACATATAATAATT

GTGATATTGGTTGCAAAGCATGTGATGTTTCTGGAAATTGTTGTAATATATCTAATATAT

GTTGTTGTAATTTCACTAGAGATTATTATAATTCTTATTTACTTAAATCATGGGGTAACA

AATGTGAACATAAAAAAAAGAATATTACACATAGCAATAATGACTTATATAAAAAATATG

AAAAACAATGGAATGACATTACTAATTGTACTATATATGATAATAATGATATTGATTATA

ATAAAAAGAAAAATAATAATGACAATTCTAATTATGTATTATCCATGAGTGGAAATACAT

TTTTTACAATACATCTATGTCATATAAAATTTAATCCTACTCATCCTATGTTTATTGTTA

AACTGAATGAAGAAAAATTATGTATTAATATTTGTGATTATGATATGGAATTTTTTTTCT

TAATATTTTCACAAATATTTGATCATTATTCTCAACAAGAATTAGCCTGTAAAAAAAAGA

ATATCATCTTTTTAAAAAATAAAATCAATTCTATATCACATTTGAATGATAAATATTATT

ACAACACCTATCATAAAAATGAAGTCAAAAAATTGGGAATTTCTAAAGATATTACAAATG

AGTTATATACAATAGCAAAACATCCTTCTCATATGTATGATGGAGAATTTATAAACTTAA

AGAAAAAGAATAAAAAATTAAAAAAACAAAAAAGACCAAAAAAGCAAAAAATAAAAAATA

AAAAATACAAACACAAAAATGATGATTTAATAAATAATCATGAAATTAAATCAATAAATG

GTAAGAGAAAACATGACGAATTAGAACAAAATGAGGTAAATTTTTCAATTTTAAAAAAAA

CAAATAAAAGAAATTATTCACTACAAGGTCAAAGAAATAAGGAAGATATAAAAAAATACA

ATATGATCAACACACAAGACACTAATGAATTACCTATTGACATAAAGGAAAAAGAATATA

AAGAAACAAGGAAAGAAATGAAAACAAAAAGTAAAACCTTTTGTAATAAATCTAATTCAT

ACAGAAAAAATACATTTGTAAATTATTCAAGTGATAACAATACAAGAACAACAAAAGAAT

TGTCTTTGTTCAAGTTAAAAAATGATATTTTTAAATATAGGCAAAGTGCTGAAAAAAAAT

ATGAAGACAAAAATATAAACGTAGATCAATATATTAATAAAAAGGATACCAAAACAGATA

TACTATTATATAATAAAAAGGATGATGTTTCATTAAAAAGCAAAAAATTACTATTTCCTA

ATTTATCTATAGATAGAATTAAAAACAGTATAAAATTTAAGAATAGCTTGGATTCGACGT

ATGTATCCAATGGAAGTCAGTATAGTGATACTATTTATGTTCACAAAACAGAATATAAGA

ATGGTAATAAAAATAAAAATGAACATGTAAATGAACATATAAATAAAAATGAACACAAAA

GAAAAAAGAATAAACAAAAAAAAAAAAAGAATAAACTTAAAAATAAGAAGAAACATATAA

ATAAAACAAAACATAATACAAATATATTATATCAGAAATATCAAATGTTTGATGAAACAT

GCAATTCAAATAATATGCAGTATGGTTATCATAATGATGTACTCATTCCTCATGATAATT

ATAGCTTTTCATCTGACGATGACTCATGGGTTGATTTATTAGAAAACTATGATAGTGATG

AATCTTTAGATGATTATGAAAAACGAAATGTTAAAAATATGCTCATCGATATTGATAACA

TAAAGAAAAGATTCTCTAATAATAATTTTATAATTACATGCAGTTTTTCTTTTTCTATCA

AAAATGTATTTATCAGATTATGGAAAAGGAAAAGCCCTATGAAAAATCAAAATTTCAAAA

CAACTGAATGGGAAAATTTGAATAGTGTGCAACTATTAGATAATATATCAAATATAGAAT

ATAATAAAGATAATAGAAATGATAACATATATAAAAGTAAAACAAAAAATATAAAAACAG

AATATGATGAATATAAATTATTAGATAAGAATAGTTACTATAATCTAACAAGTGTAGTAA

AAGAAAATGAAATAAATAGATTTCATAAAAAAGAAAGACACAATATTATAGAAGATGATG

ATAATATTAGTTTGTGTAATAAAAATTTTATTATTTTAAATGAGCATCTACAAAAAAATA

TGTTTAATAATATACAATATGGAATGCCCTTATATACAATAATATTATCAGATATATTTT

TTGTAGCTAAAGTAGAAAAAAATAATTGTGCTCATGTTTTATACTCTATGAATAACATAG

ATATTAAAGATGAAACTAATATAACATTATTAAGTATGAATTCTATATATCATAGTGATT

TAAAATCTATGGAAAAAAAAACATTCCCAAACAGTTATAAATATAAAAATAATATACCTG

ATAATATATATATAAATAAACGAAAAAAAAATAGTTCAATGAATTCAAAAGGTGATATGG

AAAAAATCGAAACTGAAGATACCTTTTTTTCTTTTGATTATTCTATAAATAATTTAGATT

TACCAGATCTTGAAAAAACACATTTGGTGTTAATGTATTTTGGTTCTCTTTTCAAAAATA

ATCACACTTTTAAATTTATTTTGAATAAAACAAAACTAAAAATTTTCTGGGAAGTTGTAG

ATGAATTTCTTTCATGGATAATAAACACCAGTGATTTATTTCCACAATTAAATACAATAC

AATATGATATTATTAAAAAATTAGATGATAGTGAAAATAAACATTTAGATTATTTACATA

AAATACAAAAAATAAATTATGATATATTAAATGTGAAAAAATTGAGTTCATCGTTTTCTA

CTAATGTAGGTTTTAAAATACCATTCCATATATTAATCAATCGTTATAATATAAATTATG

AAATATATTCAGGTCGAAAAAATCTTGGAAACTATAATGAAAATATGTTAGATTTTTACA

ATGTTGATGTAGAACCAATTGAAGAATATTACATAATTACTAATGAACAGACTTGTAGAA

GTTACAACAACGTATGTTTTATGAATAATCCAGAAGATATGAATAATATGATAAAAGAAA

TTTTGTTTAATGAGAATTATTTTTTTAATTATACATTTAATAAACTAAATAAAAATAAGA

ACATATCTGACATATTATATAAGGGTAATAATACAAATCAAAATGTTGTAAATTTTTATA

ATGACATCTCAAATGATAATGTTATTATAAATAATAATAATAATAATAATAATATACAAA

ATTGTAATCTTAATAACAATAATATTGATAATTATAATTTGGAAAATCAAAAGAGTTATA

TAAAAATTGAATCATCTTTTTTAAAAGACAACAATGTAACAATAAAATCAAATGAAGAAT

TATTGAGTAATGAATATTTGTTTGAACCTTCAACAAATAATGCGAAAATTATTAATGAAA

ATAATATTTCTATTGATAATTTATATGAAGAAAATGAATGCTCAGACTGTTTCCATATGA

TGAACTTAAATTATACATCATATACAAAAGAATTATGTAAAACCAAAAATTATCCATATA

TTCACGTAGATATCGAAATTAATTATTTTGAATTTTGGATATCACTAGATCCTATTAGGA

TACCACAAAAGAGAAAACGCCATTGTGGTATGTATATGAATAAAAAATTATATTATCATG

AAAGAAGAAAAGAAAGACAAAAGAAAAACAGGAAAAGAGATAAACGAAAACACACCATAA

TTAATAATAATAGATTTAATAAAGAAAAATCATTTTGCACATATTATAGTACAAGTAAAG

TTCTAGATGAAGAAAAATTAAGTAATACAACAAAATTGTGTAAAAAAATAAAATATTACA

AAAATTTGAATACAGATAAACCATATGTATTAGCAACAAAAGGTTGTTTAAAATCAGTTC

TTTTTATTTTATTATATAATAGACAAGAAATAAATAATGAAGTGGATAATACATATGAAT

TCATAAACGATGATTCATCATATGATCCTTTATATGATTATTTGTTAAATAATGAAAAAG

ATAAGATAAAAGGGAATAATTATTATATTTCTAAAATGTCATCATATAATTCAGAAAATT

ATATTAAATCACAAATTAAGAATAAAGAACGTCAAAGTAATCATAATTTAAAAAACAAAT

TAATGGAGGTATTACAATATTCAAATTTAAACAATATTATAAATATAGAAGAATTTTTAA

ATAGTAAGTTCATATGTTTTCTTAATAATATGTCATCATATCTTCCATGTGTAGTGAATA

TGAATATCTTTTTATCTCAAATTACAAGTGCAATAGCTAAACCAGTTACCAAAAATTCTT

TCTTAAAAGCTTATATTGATTGGAATAATGTACCATATAATAATATATTATTACAACCAT

TTCGAGCCAATTTTCAATTTGAATTGAAATTACCTAATCCAGAAATATTATTAGAACGCC

TTTTAGGAATTAATAAAATCACAAGTTTAAAAGTTATGAAAAATAATAATATAATATATA

AAAATGAAGCTGAAGGAATCGATATTAATTGTTCTTTTGAACCTATCGTTCTCAATGTTG

ATTCTAATACATTAACCTTATTAAATCAATTATATAATATCATTTTTGATTTTTCCAATT

ATATTTTAAAATTATGTGCTCCAAAAAATAAAGACGCAAATGAACTCTTGATGATTAATTCATTTAATGACAATTTATATGACAACTTTACCGAATATATGATGAGTGATGATTTTTATTTTAATCTTAATGAGTCCTCATTATTATCAACATCATTAAGTAGGTCAACATCCTTATCACTTTCATCTTTTGAATTAAATAATAAATATAAATTAAAGAATATTCAAAGCAATAATGAATTATTATATACTTCAAAGAATACTCATAAATTCCAAATGAACAATGAAGATGATGACAATATGCTTGTTAATTTTAAGTTAAATTATAATAATCATAATCCAGTTAATAAGATAAATAATATAGATGAAAATATAATACCTATTGATGCGGAGGCACCAATGGTAGAACATATTGAACAAAGCGAATTGCAGGTGAAAAAAAATAATCAAGAAAAAGTTTATAGTGAAACAGAAAAAAATAAAGAAGGATTTGATGGAGTATCAATACATGATGAAAAGAAGAAATTAAATGTATCTGATAGAAATATACCTACCGAGGACAATACAAAAAGAAGTTTGAATAAACCATTGATACTTCATCATAAATATAATACTTTTATGAACAATACAAATGAACCAATTTTAAATACAGATTGTAATATGAAGGCCAACAATCAGACATATTTATTAGAATCCGATTTGAGTAATATACTACTATTTAATTTTGTTGAAAATGTTAAGGTTAGTGTTCATTTGAATTTTGATGTAATCCTTTTACAAATATGGGATTCTAATACAGCATATAACAGATGTTCATTCAATTTTGTTATTGAATATATTAATATCCGTTTCAATACTACCAATATATATGAAATGAATTTACAAAATGACAAGTTGGAATTATATAATATAAAAGAAAATAATACCTACGATGAAATATCAAAAAATAAAAAAGATCCTAATGACAATATAAATTATAGAAATAAAAATGAAAATTTTGATCAATCAAATAGGACATTGCTTCAAAGGGTATCTGAAACAAAAAATATACAAGATAGTAATATAGACAAGAGTCATTTAATTCATTCTGGTATAACTAATGATGCATGTTCAAACATGAATTCTGACATAAATAAAAAGAAATATACCTATGGAAAAAGTGCAGTAAATTTTCTAACAAATGAAAATCAATTATATGTTAATAATATAAAAAAAAAATATAGGTCAGTTGCTGGAGATATAAATATAAATATGAACAAAGAAACTAATGATAACATGTTAACCAAAAAAAAGGAACTCGATTATTTTGAAAGTATAGATAACAATATGATGAATTGTAATATAGAAAAGGATAAAATACTAGATTTGTCAAGAAAAGAAAAAATAGATATATATAATTCAACAAAAGAAAACACAAAAATAGTAAATAGTAATCTAGAAAATAATATCAATAATAATAGTAACTATAGTAACAATAAAAATCTGTTTAGTAAATTATTTAATAGGAACAATTTCGTAAAGAATCTGAAACAATTCAAAGAAGAAATAATGAAAATTGTTAAAAAACTTTTTTGTAGAAAAAAGTTGGATGAAATATTAGAAATTATTGAAGAAAAAGGAAATGAACTTGAAACTGAAAATGAAATGTTAGGAAGAAGAAAGAAACATGAGCATTTTAAAAATATTTTTTTAAAAAATGAATTGATGTCTAATGATTTAAAAAAAAATAATGATAAGAAAAAAAGTAAAGACATAGAATGTACAATTGAATTTTTAATATATTGTGAAAATTTTAATAAAAAAGAAAATTCTTATCAAACATTTATTGAACCTATATGTACAGAAATAATAATTATGAAAAAAGAATTAAATACACCAATTCATATATATTATTATTTTTCATGGATCAATATTAACTGGAATTTTAACTTTCTAGATAATATTCTATCTTTATGTTGTAGTATCATTTTTTCTATGATTACACAAAGAACACGATTATTATTAGGAAGACATATGAATGATATAGATAAAAGAAAAAACAGAAAAAAAATAATTCAAGGACATACTAAAAATGAAAAGATGGAAGATGGATCATTGAAAAATCCACATGTTAATACAAATACAACACATCCAACTGAATACGATGATAATAATACAGAGCGAAAAAATTTAAGTGAAATCGATGATACTCAAACATCTATGTATTTACCAAATTATTATGGAGCTCTTAATGATGAACAATATAAAGATTTAGCACTTATAACTAATGAAAATATCCTTTTCAGATATAATAAATTTGATGTATTACATACCGTAAATAAGTCATTAGATACTTACATTTTCGGTGAGTTAATATTAGAAAAACTTCTTCGTAATTATCAATATTCATGTCAATATCCAGGTGATTATTTGTCAAAATATGAAAATGAATTTAATGTGGACAAAGTAAATTCTCCTTCTCAACAATTATTACATATTAAATTATTTAATTATATAAAGAAAAAGAAAAGACTAGAAATGAAAAATATATGTAAGTTAAATAATTTGTTGGGACAACCTATAGCTGTATGCACAATAATAGAAGAAAAAAAAAACATTCAATATAAAAATATAAAAATGACTAAAAGTCAAATGAAATATGAAAATAATAAATACAATAGTAAAGAAATTTATAATAATGAAAATACAATTTATGATTATAATAAAAGAAACATATTATTAAATTCTATTAATGCAATAACTAATAGTAATAGGAAGAAATATAATTCATTTAATCCATTCGAAGATTATAAATGTTTACAAATTTCAGATATGAATCATAAAAAATTAGGAAAATGTCAAAATCATAGTGAAGAATTTGGTAATTCAGATAATAGTTCTTTAAATGATACTAAGGATATTTTAAAATATCAATGGAAAATTATTAGTAATAATGAATCGCTTGATTTACCAATTCATGAAAATGGAAAGGTGAAATTTTTTCTAATAAGATTTAGATTACTAAACTACATATATGATATACCGTCAAATATATTAAATACTGAAAAAGAAAATGAAGATGTTATAAGATTAATTATACCTGAGAGAAGATTACCTCCTAATATAAATGAGCAAGTAGAACAAGAATTATATAACCAAGAAATATTAGAAATGGAACAAGAAAATGATACTATGGATTCTAATTCTCATACAATATATCCTACTAATTATCTATTTAAAAAATATGTTACAGGAAAAAATGATACTGTATGGTGTCCTAGTAATATTCCGCAACCACGTAATCATTTATTTATTTATTTTAGAACAAGTAGTAGAATTAGTCATGAGGAAAAACAAGAATATGATTTTTTTCTATCTTCCGTAGTTGCTATTAAAAATAACACTGATTTTCCTCTTTATATATTTAAATCAGTACCAGGTAATGTAAATAAAAGTAGTATCTTTAAGGAATATTTTCACAAATATAATACCAAACCATCTCCACCACCTACGTATACATTGAAGGTTGATAAAAAAATCGAATACTTAATTGAAAACCAGAATAAAGATAAAAATGATTTCTCTAATGGAAATAATGAAAATGAACAAGATAATAATATAGAAAAGAGAAATAATATATTTTACAGCACTGATCAAATTATTATTAATAATAATAATAATAATAACAATTATAATAACAATTATAATAACAATTATAATTATTGTTTTGATCCCAATAAAGACCATAATAGTTATTCAACTAGACAACACAATGAAAAACTTTCTAAACAAAAAACAAAACAAAACAAAAATAAGAACCATAAAAAAAATAATAATAGTAATGATATTTATGAGAAGAAAGATAATAAACAAGGTAATAGTAAAAACTATTCCAATAATCACCAAAATAATAATAAAGATCAGGAAGAAGGAAAGAAAAAAAATATAAACTCATCAAATGAATCTTACTTGCATAATAGAAAAATGATTAAAAATATTTTCCCCTTAATGAAATTGTCTTCTATGGAACATTATATTTCAGGAGATATAAAATTCAGTTCATTAGCTCCATTGAAATATAAAAGAAAGAAAAAGAAAAAGAAGAAGAAAAATAAATTGAAATATGAACTGCAAAAAGAATATATTAAAAAGAAAATAAAAGAATTATATTTAAAAAGTATATGGAAAAAAACACAACGTAAAAAAATTAATAAATATAAAAATGACATCAATAAAATAAATCAAAAGATAAAAGAATATTCATATGAAGAACATGCACCTTACAGAAATATTATGTCTATCAATGATCCTTTTTTTAATTTTATAAAACAAAATTCATCTTCATCTCAAAAATTATCAAATAATTCATTAAATATTTCAGATGTCACAAATATTACATTATCATCCTTATCAAGTACTTTTTTTTCTATCCATTCAAGTCAAAGTAATAATTCAACATATGAAGATTCTTTATATGATTATGATAAAGATAATTGCTTAGAACAAGCATTATTAGATGAATATTTTAAAAAGCAAAATGATGATGATGTTTTATTGGATATTATTGAAATATCTAATAATAGTAATAAATTAATTCCAATACCATTATATTGGCTAGTTGCTGAAAATTCATTCATATGGTTATCTATACAAAATGAAGAAATTTATAAAAATAAAGATTTGTATAATTTTTTAAATGAAGAAATGAATGCAGACTATTTATGTACAAAAAAAACATTAGATAATATCTTAAATTATCAATCTCCTTTTGTAGCTGATACACTTAAATTATTTAAACCATATTTGATTAAATTAAAAAATCATTTATTGAAAGAAGATAATTTATTTTCAAAAAATAAAAAAGATGAAAATGAAACTTTTATGACCAATTATAATACAAACTATATGAATTTAAAAGAAGTGTATTTCACTTCATCAGTTGTTAGTGTCTATAATCGTAATTATGTACTAAATCGAAAAGATGCTCACAATTTTTTTCAAATAAGTATTGAACATGTTCTAATGATCAATAATGGATTACCTAAAACTATGTATATAAATTGTGAAGTCTTAAAAAGTAAGCTCAAAAAGAAAAATACTATAAAGAAAAACATAAGTCACTATAAGAGTTTTCCATCAATCGAAAAATCTAAGCAATATAAAGATTCAGAAAATAGAGAAGACTCTTATGTTATAGAAAGTTCTTGTAATAAAACAAATGAAGAATATTCATCATATTATAATGACGATAAAAATGATAATGATAGTAAATATTATGATGATGGCAAACATGATGATGACAGCAAATATGATGATAACAGCAAATATGATGATAACAGCAAATATGATAATTACAACAACAACCACCATAATAATAATAATAACAATAATGATAATAATAATTTTATGTACAATGAAGGTAGTGACATGTTCTCCCAAAATTCTCATGAAGTTAAAAGAGATAAAGGTGTTGACACCTTCTATGACATACATGACGAATTAACCTTGGAAGAAAAACAAAACATGAAGAAAAAGAAAATTACATTAGTTGTTGATAAATTTAACCAACCATTAAATGAAGATACTAATAAGGATCATATATATATCGTTGATAAATATAGTGATAATAACAGAACAAATAATAAACGAGTTTTTATAAAAGAAGCTAATATTGATAGAAATGTATTTATATATAAAAATACAAAAAAAAATGAAAATGATTACATTGATGACAACGAAAAAAGTAAGAATTATATGGATGATAATAATTTTATTGTTAATATAAAAAAAGAAGAAAATAAAGATATTTATGAAATAAAAATTAATCCTGGACAAAGTATTAAATTACCTTTCCTTCTTCATAATATTACCATATCATTTGATGGATATCATTCAAGAAATATCTTTATAAATTATCCTCATAAAAAAGGAAGGGAAAAAATCATGTTAGAAAGACAAATATCTCCAGATATTTCCTCCTTTTTCACATTTGATTATGAAGAATCCAATGAAAATAATGATGAACATAGTAGTAATAAAAGTACAATGAATTATGTAAATAATAAAAATATTAGTGAAAATACAAATAAAAATTATCATCATTTTGATAGTTCCTTATCTCTTACTGATAAAACAAAAAATCAGTTAACTTATAAAAATAGGAATCATCAAAAAAACGATGATTACCCACTATCTAAATATAGTGATTATCATAAAAGAAAAACAACAAAATATGAAGAATATCAAAAGCAATTATCTATATGGGCAATATTTAATGACTTTTATGAAACAGGCAAAAATCGATATCAAGATATATCATTAGAAAAAATTTACCATGGAACATATTCTATTTCTTGCACATTTTTTGCATCTGCATGTATCATAAATAGGTTAAGTTATCCAATTAAACTGAAACGATTAAATAATGAAAATGTGGTTGTTATTGAACCTTATATTAGAACGTTACTCCCATGTGAATTTTCCATGCGTAGAAATAAAGTAGTAATATCACATGAAACCTTAAAAGATATTAAGAATTGGAAGAATTACCCTTTAAAAAAATATATGATTAAACGAACAAAATTGATTAAAACTCGAAATTTTATTAAGAAGGATAACCAAATAATTGTGCATTCTACAAAATGTGATGATGGTAAACAGATTAAGCATAAATGGGATAATAAGCAGTTAATTCGAGGCGATAAAAATGTAAGCATAAAAAAAAACAAATCTAATGAAGAAATAATTTTAACAAAACAAAATGAAACAACAAATGAAATAAATAATGAAAAGATAAAAAGTGATATAATAAAAAATGATGGTAGTTACCAATATGATAGGAAAGAAGAAGACCATTATCATGTGGACGATAATAATGATGATGATGAAATAGAATATAATAAAAAAAATATTCCTTATAATAAGAATACTCTTTGTGGATATAAATATAAAAAGAAAAGAAGATTCAGATTTTTTTTTTATAGAAATAATATTAATAAACCCTTAGAATCAGAAGATTTTCGTATAACCGATTTATCAGTCACAAGATTAACATGTTCCAATAAAAATAAAAATATACCACAATTAAAATATAGTACTTATATGTCTATTGCTCCTATGCCTTTTTTTAGAACAACCGTTATGGAAATATTACCTTACATTGTTATAATAAATAATACAAAAATGAATATAGTTTTTCAAGAGTTTATAAAAAATTATACATATTTTAAATATAACAAACTACAACCTGGAGATTATGTAGAATTCCATCCACAGTCAAAAAAGAAAGTTATGTTAAAAATGTGCGGTATATATCAAAACACATTCGATCATTTAATTGATGATTATTTACAAAAGGAATTATATTTAGAAAACTTCGAAAAGAAAAATGAAAAATTAATAGATGATATAAAGGCAGACAAACATTATGATAAGAAAAATATGGAAAAAAAAAAAAAATACTCAACAAATAAAGAATCTAAAAAGAAAAAAAAAAATAACATTGCCAATAATTATATATGGAATATATCTAAAAGGATAACTGCATTAAAAGATATGGAAAACGTATATGATAGTAATAATAGTGCAAAATATAATGATAATGAAAAAAATTATATTAATAATAAAATGAAAAACAAAAACAAAAAAAATAAATATAAAAATAATAAGAATAAGAAGAATAATAATGAAAATAATGAAAATAATTATAGCACATCTTCAAAAAACTCATCTAACAGTACCATTGAAAATTATTCAGAAGAAGAAAATTCAAATGGACCTTTGAATTCTTCTAAACATAATAATATTAAAAAACATAAATATATGAATGAATACACATTTTTATACTGGTCTGAAGTTCTAGATTTAAATAGAGTAAGTATGATTTATTTTCGTCATCCAGTTATATCAGAAACCAATTTTAAAATAGAAGAAAAAAAATTACAAAAACTAAATTCTATATTAAGGAAATATGGTAAGAATGATGATGCTTATATAAACATTTTAAAAGAATTAATTAAAAAAAATAATCAAATAACATATAAATATAGTTGTTGTTCTATGGAAATTACAACATTTAAAGGATGTAAATTTGTACGTTTTCAAGATATTGATGTTGTACCTTATTATTTAATAAACTTAACCAAATATAATATAAAATTAAGACAAGTGGGAATTTATGAACATTCAGAAATTCTTCAAAAAATACCTCATAAAAAAAAAACATCATTTAATGAAATATTTAAAAATTATGCATTCAATTTCTCTTTTTACAATCCTTATAAAGACCCTAAATTAAAAGTTACTCTCTTATCGAATAATAAGAAAAATGATTATTATAATAATAACAGCAAATATAAAAAGGAGAAGAAAAGAACACTATTAAAAAAAATACCAAATAAATTAAAACATAATCTTAAGCATGAATTAAATTATCTAAGCTTTTTAACAAAAAATTATATAACAGCGAAAGAAGATGCAAAAATCTTGAATGACAAAATTTTCCAACTCAAAAGATTTTCACAACAAATTGAAAAATTAAATGATGAAAAAATGAATATTATTGATTTAGGAAATAATAATAATATTACCTTACTAAAATTAAATTATAAAAATTCTCTTATTTATATACTATGTACCAAAAGAACATATAATGGAAAAACCATCTATTTATTTGAAAATTATGATGATTTAATAAGTAACATCTTAAATTTTAATACTATATCATTTAATAGTAAATGTAATAGCAAATTGCTAAAATCATTCAGTTTGTTTCGTGATACAATTAAAAGCAAAAATAAATATAGTCGTAAGGATACATATAAGAAAAAAAAGAAACATATTAAATATATGAAAACGTTTATACACAATCCAAATGTTAAATATTTATTAAATTTCTTTAAAAAGAAAAGTTCGGCATCATCTATAGAAAATAAATTAAAAAAAAATATAAAACAAAAGAAAAAAAATAAAATAAACAAGGTATACAAAAATTCAAGAACAAATAAAAGGACAACACATATTATTACGGAATTATACCAAAAAAATATGTTCCACCAAGAAGAAGAAACAAAGAAAAAAAATAAAAATATCATATTAAATATGAATATATTCGCACATTTAATATTTAAAGGCATAGGAATAAGTATTAGTAATCACAATTTAGAAGAAATCCTATATTTTTCGATGGAATATATTTTAATTATTTATAAGCAAATGATAGATCGTGATTTGTTTCATATAAATATAGGATGGTTACAAATAGACAATCATACTAAAAATACTGACTATAAGAATGTGTTGTTACCAATTGTTGATTTACAAAAACAAACAAATAAAAATAACGCAAAGGAAATTAATGAAGAAAAGAAATATTTTATAAATGAGAGTAATATAGCATTATATAATAGTATTAGTAATAATAAAAGAGATGACAAATTTAATAAAATTAATAAGCATAATACCTTTCTCCATAATAGATATGATACAATTTCAGGGCATATAATTTCTGGTGATATCAATTCACGTAATTTAAAATCAAATAATTTCAAATCTTTTAATACTTTTTCTCGTAATATAAATTCTTGTAATACATTTTCTTGTAATATTAATTCTTGTAATATAAATTCTTGTAATATTAATTCTTGTAATATTAATTCTTGTAATACCTTTTCTCGTGATACCTTTCCATATAGTTTTAGATATAGCAATAATTTTTCAAAGTCTAGTACATTAAATATGATTAATTCGTTTTACTCAAATAGAAATAAATCATATAATTCATTTATAGATAATGATGTATTAATTGTTACAAATTTAAATAACCCTCTACATAATTCTTATAAAGTATTTATACCACAATTTTTTTATTCTGAATATAATTCTGTAATATATATTAGTATAATTAAATTAAAAAGAGGAAATATGAAAAATTTCTTAGAATTGTCCGATGTTCAAATAAAAGTTTCTCCTTTGTCTTTAAATATTGATTCCTATTTTATTATCGAAATGTTAAAAATATTCGACGAGTTTTTAAAAATAATCGAATTTGATTTAAGTAACTACAATTCACTTAATTTAGAAAAGAACATTGAATTGACAGATAGAGATTTCGATAGTCTAGAATATAAAAAGGATATAGGTCCTATAGAAGAATTAATTAAATATCATATATTAAAAGATTATTATTTATATGAACAAATAATTAATTTCAAGTTTAATCAAAAAGTAGATGGTATACGAAATCAAACAAATAACAAGTCACATAATTATAAAGAATTAGAAAATTATAACAAAATAAAAAAAAACAGTCAAGATAGTATATTTAATACGAATAATTTATATGACAAATCTAATTTGGATGTTATTGAATTCAAAGAATATATTAATATGCTAAATGAATATGAAAATATTAAAAAAAATAAAGCTATATCAAAAGATATAAGAAATAAGGATATGTTATCCAAAATGCATTGTACTAAAAATAATATATTAAAAAAAGGATGTACCAAAAAAAGTAGAAGAACAAAAAATATAAAATTTTACAACAAAAATGATGATGATAATAATAATAAGGTTAACGATAATAAAAAGGATTACAAAATAGATACACAAATAAATGAATATATAAAGGATAATAATAATAATAATAATAATAATAATAATAATAATACGGATAATAAACAGTACATTAATATAAAAAAAAATATAATAAACATTAACAAGTCCTGTCTAATTAATAATAAAGAGAACATGTATTATGACACAAAGAAATATTCAGATATAGATAAGAAGAAAAAATCAGATGATAATGAAGATACACAAGTTGAACTATTAGATAAAGAGAAAATTGAATATATACAAAGATGTGATAAATATTACAAGAATTCCTTAAATAATAGAAAGAAGATATTGAGAAAAATACAAAATATAAATATAAAAAATAATTTAATAAGTTTTGACAAGATATTCATATCTAAATTAGAAATTAACGAAATGAAATTAATCATAAATATAAAAAATAAAAGTCTGAGTTATGATAAAAAAAGTGAAATAATGGAATCAAATGTAATGAATGCTTTAGTTGTATTAATTATGAATATACCAAATATAAGTGATGCACAATTACATTTTGATAAAGAGGTTAAGAAAAATATCTGTGGATCTTTATATGAACTAATATTTAATGAAATTATGAATGATTATATTACACCTGGAATGAATCAAATGGTTAATGTATTAGGTGCTATAGATTTTATAGGGAATCCATTAATAATATATAAACATTGGAAAAAGGGATATGAAACATTTATACATGATTTAAAAAAAAGTCTAGATTCATGCTCTTTTCCATTATTATTTTGTATCTTATTTTTAAATACATTAGGTAAATTTGGAAAGAGCGTATTATCAGGAATTCTAGAAGGTATTAATAGATTATGTAGTAGCTGGAGTACATGTTTTGAAAGATTTTCTAGAAATTCAGATAACTATAGTATTGTTACGAATAGTAACCTCTTTCAACATGAAATATTAGATCAACCTTCTAATTGTGCAGAAGGTATATATTATGGTTGTCAATCCTTTATAAATATTCTTACTATTAGTTTTGTTAATATAATATATAAACCATTTATAGCTATACAAAAATCAAAAGAATTTAAAAAAAAAACAGAAAAAGGGAAATACAAAATATTTTTTTTATCATTGATAGTTCTATCTTATGGTATATTTAGTGGACTTAGTAGTTTCTTTTTTGGATTTTTTAATAGCGTATTTTCTTTTTTTACATTTCTATTTATTGGTACATTAAATCAAATTCAAACCTTAACAATGAAATCTGTTGTTAGACCAAAAAAAATGAGTATAATAAAAGAATATTCCAAGTTTCTAAATTATGAATATCCATTATCCTTTTCATCTCATGTAATTAATGAAAAAAAAAGGAAGAAAGAACTTTTAAAAAAAAATATTGTTGCTATTATACTTTTATATACTATTAACGATAACACTCCAAATAATATTAAAAGTTATTTATGGGTGAGTAATAACGAATTAGGATATTGTCAAAAGGATAAATTATTATGGTCTTTATATATTAATTGTATA**ATCAAAGTGGATATATTATTAATCGAAATGGATAACGGAAATTTCAAGCGATATTCAATAAGCAGCCGTAAAGAAAATAAATATTTGAATAATATGAATAATAAGGAAAAGAACGATGCTAACAAACACACCAAAAATAGCATGAATAATAATAGTAATAATAATAATAATAATAATTATAATATAAATGAAAAACTTACTTGGAATGAAAAAAATAAAGAACAAAATAATCAAATTAATAATAACAGATCCACATGGAAAATAAAAAATTTATTCACCCCACAACCTTTTAGACAACTTTCCGATTTTAAACCATCGTACTATATTAGAATATTATATAAATCATTATATAAAAATAAGAAGGAAAAAAGAAAAACAAATATTTCTAATTTATTTTTATATAAAAAAAGTGATTCAAATAATAGATTGAAACGTATAATGAGGAAATCCACTTATGAAAAATATTTTGAAAATGAATTTAAAGAAGAACAAAAAGTTCATCGTTATCATACTACTTCTAAAGATATGCCTGAAGAACCAAGTGCTAAGATTTCACATATAAATGACGAAAGAAAAAAAAGAGAAAAAAAAAACAATAAAATTAATATATATGATAACTTATATCCAGACAAAACCAAAAATATTAAACATCATAATACAAAACATTATAAAAAAGAAACAAAAAAAAAACATTTAAATAAAAATAAAAATAATATTACATCATACATTGACGATCAGAATAATCATATGAAATTATGGAAACAAAATAAAAAACATAAAACAAAAAATTATAAGTATAATAATGAACATAATACTGAATATGAACATGTGAATATAAAAAAATTATTTAAATATAACAATCATAAGAAAAAAAAAAATAAATATTTATATATCTCAAAATTGGTCAAATGTGAAAATAAGGAAGTTGCATTACATGCTTTTTCACTTCTTATGTCGTTCTTAATGCACAAATCACCATATTATTTAAAAACCACTAAT**TGAatatacacaagataataataacaatattacaaaaaaaaagaaaaatataaatataaataaatatatatatatatatatatatatatatatatatatatttatatggggtgtgtatgttccttttcattatcatattttaaatcaaaaatagagtattataaatattttaaaccttattcttttaaaaatctctttttattgtattataattatatatatatattattaatatttaattacatattatttttctcttttattaattatcggtagttatatatagatattccttgtcaattttttaaaaatttatcttttcttttttttttttaataaaaatatataaattttgtaattaattctcaatatatatttattatacatattttaatttaaaactaatacttattatataatatatagtgtctgttaattattgggcttacattcttattaatttttaatatatatttatatattcatgaatattatatttactt

**Original Locus (OL) amplicon size: 1103bp**s

**> PF3D7_1343800-Halo-SW Halo Neo-R**

tttattatggatccaaaggaaaaaaataaaaaataaaacaaaaaaaatattacatttatatatatatatagatttttaaatatttgtatgtatatattgtttaatttttttaatttttcatttaaaaaattttatttttatattgctaaaaatttactaaaatatgcaaaaaaatataatgcatataaaatttctttaaataaacatattgtctgaatgtattaatttcttcagaaaaaaaaaataaatataaaaatataattgtcaatatatattgataaaaaaaaaatataaataagtgaataaatatatacatatatatatatatatatatatatatatatttttatgtatgtatttatttgttgttatatttaaaaaaatagaccatagtaatgtaatataatataatatatatataatatatatatatttatttatttatttgtttatttttttatatttattaatttttgaggcaaataaagaaatatgtaagaATGTTTCAGAAATTTTTTTCAGATATATTATACAAAATATTGGGAAATTTTATATATGGTTTAAATGATGAAGAATTAGCATTAAGTGATTTATTAAGTG

GGAAATTGGAACTAACTAATATTCATTTAAAACAATCAGTAATAGATTTGATAGATATACCATGTCGCTTAAATTTTGGATGTGTTGGTTATTTTAAAATTAATTTACCCTTATTATATTTTATGAAGAATCCAATAAATATATATATGGAAGATGTAATAATAGTTTTGTCTACTATACCTTCTAAATATTTTGACGATAAGTTATATAAAGAAAAATATATTGAAAATAAAAAGAATTTATTACTTTCTTCTGAATATAGAGCTATCGTTTGTTCAGCAGAGGGGGGTGTTATATGGCAAATGATTTTGTCATTAATAAATAATATTAATATAACCATTAAAAATGTACATATTAGAATAGAAGATTTTACAAGTAATCCTTCCAACTGTTTTTCTTTAGGAATTAAAATTGAAGAATTATTTTTGAATATGCCGAAAGAAGGTTTACCTTTTAATAGAGATGGATATTATACGAAAAAAAAAGAACTCGTCAGCAATACTATGATTAATATGATAACCGTTAAGAGGTTAGCGTTTTATATGGATTCTTTGGATATGAAAAAAATTACCAACAAAAAATATTTTAACAAGTTTAGAGATATATTTATAAATTATTGTAATATGGATGAAAGGAATTATCCAGAAATAAATATGAAATCAAATTACACAAACGACAGTAAAAATATAAAAAAAGTACATATGAATTGTGAAATACAAAACATGAATAATTTTAAACGTAATAGAAAGAATAAAAAAAATAAATCAAAACATAATAATCATAAAAATAATAATAAAATTAATAATAATACCAAAAAGTTGTTTAAAAAAAAAAAAAATAAACGTTCTAATCAACAAAACTTAGAGAGCTTTCTCAATATAATTAAAAGTATTAAGAAAAGTTCTTATTATGTTTGCAAAAATGATAATAATATAAATAAAATGGACAAAATAAATCAAAGGATTTTATTATTAAATGATACAATTTCAGCAAAAAAAAAAAAAAATAAAATAATAAATAATAATATAAAGATGAAGAACACACATAATGAACAACAAAACAAAGATAATAATTGTTCAATATCTATAAGTAATTTCTACAAATTTAATCCATTCTCATCAAAGAAAGAGGAGAAAGTAGACATGCCTATATTGATTGATGATATAAAACTTGATAATGATATAAGTTGGATAGCATGTATTACATTGTTTGCAGATTCTTTGGAAAATTCTTTTATCAAAGAAAAAAAAGAAGAATTATTAAAAAAGCAAAGAAATAAAAAGAAAAATAATACAAATGATAAAAATGCACAACCGGAGAAGGATGATGATGGTTTCTATCAAACTTGTAAGACTATTGATGAGAATAAGAATATTAAAGATCACAATCATTCTTTTAATAATAAATTGAACAAAAATAATGAACAAATTAAAAGTAATAATTATGAGAAGGAGAAGATGGGGAGTCATACTTTTATTGATAAAGATAATAATAATAATAATAATAAAAATAATCCTTCCTTTAATAGTGATATAAGTAATACCAGAAAACTTACTTCCAATTATTATTTAAACAAAGCTGTCTCTTTTCTTTCTATGAAATCCTTCACCAAAATACTTCAGAAGAAAAAGTTATCAAAAAATATGCACCGTCATAAAAATAAAGAAAATGATTTTAGTTCAAATGATCAAAATATAAGTGATGACTATTTTACTACAAAAGAAAAAATGTCCGATATAGATTTTAACAGTGAAGATATATATTCTTCTAAATATAAAAAGAAACGAAAGAACGAAATGAAAAAAAAAGATTTCAATTTTAAAAAGAATGGTTATTCGAAATATGAAAAAAAAGATGAAGAGAAATATTACGACGTTAAGGATAAAAACCAAAGTGATGAACATATTTCTACATTTCATCATTCTAAAAATACTGACAAGAATTATTTTAGTTTTAGTGATAATGATAAGAATAATAAGAAATATATGTCACAGAACAAAAAAAAAGGAAAAAAAAAAACATATATTGAAAAGCAAGAAGATTATGGAAAGAATTATAAATATAAAACGAGAAACGAATATGACAAAACAAATAATGAGAGTTATAAAAATCAAAATTTTAATGATTCAGCTCATAATTTAATATCTAATTATGAAAAATGTATACAACTGAATGATAAGAGGAAAGAAATAAAAATGGAACAATATAGAATTATGAATTATATCAAAAAGAACGAAATAAAAGAAAAAATAAGAAATAAGAAAAGAAAAAAAATTCATTCAGTTTATTTGATGTATGAAAAAACATACAAAATAAATAAAAGTAAATTGAATTATTTCTATATATTTTTGAGGCTTACAAAAAATATCAAGCATGAATATATTATTAACCCAAATTATTTTGAAGGATATGTAGAAATATATTTAAGTTTAATTCCAACAGATCATGGTGAACGTCCATTTTCTAGATGTTTTTCATCTTGGGGTATACAAAGAAGAAGTCAAAATGAATTTGATGAGTGCTTACCTAAAATAAGTGTTTTTGTTACATTAAAAAATATGAAAATTATATTTTCCGATAAACAAATGTCGAATTTAATAAAATGGTTAAATTTAAATTTTGTAACATATTCTACTTGGAAAGCAGGTATCTTATCTCAATTTGAGAAGCCTAAAGCAACACCTAATGAAGAAATGTGTTATATAAATAAATGGGTTCTAAAATTATTAGATACTCATTCTTCAAAAGAAGAAAAGGAAGATGCCGAAAAATATTGTGAATTATTTGAAAATACTCATTCTATACATATTATAAATTTGCTAAGAACTAAGGCATTTTCAAAATTACAGGAATTAAGAAAGGATATGGAAACAAAAATGAAACCAAACAATAATAATAACAACAATAATAATAATACGTATGATATGGTAAATGACACAAACCAATATAATGAAATAGGCAAAAAAGAAAATGAAGTGAAACAAGTTGCTAATGAAAGTATTAATGTGTTAAAAAATTTAATGAAAAATGAAAAAGCATTTAATAAAATAAAGACATCATTTAGAGATAAAAATTTTACCATAGACTTTTGTTTCGATATATGTATTATTAATAGCACAATTTTGTTAACTATTGAAACATTTAATAAAATAAATACTAGAAAGGTGAACTTTGTAAAAACATATGCTCTATTATTACAATGTATACATTATCATAATCAAATTTCTAATACTCAAGAATTTAAGAATTCCATGGATATCGAATTATATCCATTAACATTACTTTTAATAATTAACAATTTAGATAACAAATATATTAAACCCGAAATTAATGTATTATATATTTTAACCAAATCAGATAAGAAGGATAAATATAAATATAAAGATAAAAAGTTTCTATTCAATATGGAGAACTTTGATATATTAAAAAGATACAAATATATTAGAAATACATATCTATATCGAAATGATAATCAAAAGAATTCAAACGAAGACAGGCAAAAAATTGAAAAGGAACAAGAATTATATAACCATGAAAATGAAAATGAAAATAATCACAATAAACAAAGAGATACATTAAAATTACATAATTTAAAACAAAAATTTAAATATTTAATGGAAGAAAATTTTTTCTTTTCAAAAAGACAAAATAATATGTATTGTAATGAATTATGTGATATTACATCTTTGTCAGATCCATCAATTTATTTACGATATGATTCTCAATCATATACCATATTAGAAAAACCTGATTGTCATTTGTTTTTACATAATTGTGCTTATTCAATATTTAATATAAACGGAGAAATAGCTAATTCATTTGTTAATGATTATTTCCATTTTGTAGATAATAGATCAGTTTTATTTAAAAGGAAAACAAATAATTTTTCTTTTAAAAAAAAAGAATATTTCTTAGAATTAGGAGCTAAATATTGTAATGATATAATGGATAATTATACAAAACATACTAGTTTTTTCATCGATATAGAGTTAGAAAAAATGTTATCATTTATTATAACAAAATCAGGACAAAAAGCAGAAATTCAGGGTTATAGCCTTAATTTAAATCCAGTAAAAGTCGTAAGTCAATTATCTAAAAGATCAGAATCGTATAATATAGATAATTCCAAAAAAAATATATATGATTCTTTTTTTGTTCAATTTGAAGGAATAAATATAAAAAGAATATTAAATTGGCATGAAGCAGCTAAACATTATAAAACATATATTGATGATAATTTTATTAAACCTTTTCATGTCCTTTTTCATCCTATAAATAAAGATATATATACAAATAGAAACCAAACCCCAGATAAATTCTTTTTTAATTTAAATCGTAAGAAGAATATACCTCAATTAAATGAAAATGGAAATAGTAATTCAAATGGTAATTCAAATGGTAATTCAAATGGTAATTCAAATGGTAATTCAAATGGAAATTCAAATGGAAATTCAAATAAAAATTCAAATAAAAATTCAAATGAAAATTCAAATGAAAATTCAAATAAGGATAACAAAATATCATACCAAAATGATAATATTTATAATAACAATAGTAATAATATGCATAAGAGAAGTAGTTCTTATAAGAATTTAAATAATGAAACAATAAATTATAATAATGATGAAAAAAAAAAGTCGGAAATTTATAATCTTATATATAATACTAGAAAATCACTAAGTTCATATACACATTCCGAATCTGTTTCAAAAAATCATATATACGATAATAATAAAAATATATATATAAAAAAAAAGAAAATGAATAATAAAAAAAAGAAAAGTAAAAACAAAAAAAACAAAAAAACAAAATTAAATAATATTAGAAGATCTTCAAGTTGTCCACCTCCTAAAAATATAAATATTTATTTTGATAGTAGTAATAAAAGTGATATATTATATAATGATAGAAAAACCACTTTATATTCATTAAATTATATGTATGAACAAAATAGACAATCTACTATAACATATAATAATTGTGATATTGGTTGCAAAGCATGTGATGTTTCTGGAAATTGTTGTAATATATCTAATATATGTTGTTGTAATTTCACTAGAGATTATTATAATTCTTATTTACTTAAATCATGGGGTAACAAATGTGAACATAAAAAAAAGAATATTACACATAGCAATAATGACTTATATAAAAAATATGAAAAACAATGGAATGACATTACTAATTGTACTATATATGATAATAATGATATTGATTATAATAAAAAGAAAAATAATAATGACAATTCTAATTATGTATTATCCATGAGTGGAAATACATTTTTTACAATACATCTATGTCATATAAAATTTAATCCTACTCATCCTATGTTTATTGTTAAACTGAATGAAGAAAAATTATGTATTAATATTTGTGATTATGATATGGAATTTTTTTTCTTAATATTTTCACAAATATTTGATCATTATTCTCAACAAGAATTAGCCTGTAAAAAAAAGAATATCATCTTTTTAAAAAATAAAATCAATTCTATATCACATTTGAATGATAAATATTATTACAACACCTATCATAAAAATGAAGTCAAAAAATTGGGAATTTCTAAAGATATTACAAATGAGTTATATACAATAGCAAAACATCCTTCTCATATGTATGATGGAGAATTTATAAACTTAAAGAAAAAGAATAAAAAATTAAAAAAACAAAAAAGACCAAAAAAGCAAAAAATAAAAAATAAAAAATACAAACACAAAAATGATGATTTAATAAATAATCATGAAATTAAATCAATAAATGGTAAGAGAAAACATGACGAATTAGAACAAAATGAGGTAAATTTTTCAATTTTAAAAAAAACAAATAAAAGAAATTATTCACTACAAGGTCAAAGAAATAAGGAAGATATAAAAAAATACAATATGATCAACACACAAGACACTAATGAATTACCTATTGACATAAAGGAAAAAGAATATAAAGAAACAAGGAAAGAAATGAAAACAAAAAGTAAAACCTTTTGTAATAAATCTAATTCATACAGAAAAAATACATTTGTAAATTATTCAAGTGATAACAATACAAGAACAACAAAAGAATTGTCTTTGTTCAAGTTAAAAAATGATATTTTTAAATATAGGCAAAGTGCTGAAAAAAAATATGAAGACAAAAATATAAACGTAGATCAATATATTAATAAAAAGGATACCAAAACAGATATACTATTATATAATAAAAAGGATGATGTTTCATTAAAAAGCAAAAAATTACTATTTCCTAATTTATCTATAGATAGAATTAAAAACAGTATAAAATTTAAGAATAGCTTGGATTCGACGTATGTATCCAATGGAAGTCAGTATAGTGATACTATTTATGTTCACAAAACAGAATATAAGAATGGTAATAAAAATAAAAATGAACATGTAAATGAACATATAAATAAAAATGAACACAAAAGAAAAAAGAATAAACAAAAAAAAAAAAAGAATAAACTTAAAAATAAGAAGAAACATATAAATAAAACAAAACATAATACAAATATATTATATCAGAAATATCAAATGTTTGATGAAACATGCAATTCAAATAATATGCAGTATGGTTATCATAATGATGTACTCATTCCTCATGATAATTATAGCTTTTCATCTGACGATGACTCATGGGTTGATTTATTAGAAAACTATGATAGTGATGAATCTTTAGATGATTATGAAAAACGAAATGTTAAAAATATGCTCATCGATATTGATAACATAAAGAAAAGATTCTCTAATAATAATTTTATAATTACATGCAGTTTTTCTTTTTCTATCAAAAATGTATTTATCAGATTATGGAAAAGGAAAAGCCCTATGAAAAATCAAAATTTCAAAACAACTGAATGGGAAAATTTGAATAGTGTGCAACTATTAGATAATATATCAAATATAGAATATAATAAAGATAATAGAAATGATAACATATATAAAAGTAAAACAAAAAATATAAAAACAGAATATGATGAATATAAATTATTAGATAAGAATAGTTACTATAATCTAACAAGTGTAGTAAAAGAAAATGAAATAAATAGATTTCATAAAAAAGAAAGACACAATATTATAGAAGATGATGATAATATTAGTTTGTGTAATAAAAATTTTATTATTTTAAATGAGCATCTACAAAAAAATATGTTTAATAATATACAATATGGAATGCCCTTATATACAATAATATTATCAGATATATTTTTTGTAGCTAAAGTAGAAAAAAATAATTGTGCTCATGTTTTATACTCTATGAATAACATAGATATTAAAGATGAAACTAATATAACATTATTAAGTATGAATTCTATATATCATAGTGATTTAAAATCTATGGAAAAAAAAACATTCCCAAACAGTTATAAATATAAAAATAATATACCTGATAATATATATATAAATAAACGAAAAAAAAATAGTTCAATGAATTCAAAAGGTGATATGGAAAAAATCGAAACTGAAGATACCTTTTTTTCTTTTGATTATTCTATAAATAATTTAGATTTACCAGATCTTGAAAAAACACATTTGGTGTTAATGTATTTTGGTTCTCTTTTCAAAAATAATCACACTTTTAAATTTATTTTGAATAAAACAAAACTAAAAATTTTCTGGGAAGTTGTAGATGAATTTCTTTCATGGATAATAAACACCAGTGATTTATTTCCACAATTAAATACAATACAATATGATATTATTAAAAAATTAGATGATAGTGAAAATAAACATTTAGATTATTTACATAAAATACAAAAAATAAATTATGATATATTAAATGTGAAAAAATTGAGTTCATCGTTTTCTACTAATGTAGGTTTTAAAATACCATTCCATATATTAATCAATCGTTATAATATAAATTATGAAATATATTCAGGTCGAAAAAATCTTGGAAACTATAATGAAAATATGTTAGATTTTTACAATGTTGATGTAGAACCAATTGAAGAATATTACATAATTACTAATGAACAGACTTGTAGAAGTTACAACAACGTATGTTTTATGAATAATCCAGAAGATATGAATAATATGATAAAAGAAATTTTGTTTAATGAGAATTATTTTTTTAATTATACATTTAATAAACTAAATAAAAATAAGAACATATCTGACATATTATATAAGGGTAATAATACAAATCAAAATGTTGTAAATTTTTATAATGACATCTCAAATGATAATGTTATTATAAATAATAATAATAATAATAATAATATACAAAATTGTAATCTTAATAACAATAATATTGATAATTATAATTTGGAAAATCAAAAGAGTTATATAAAAATTGAATCATCTTTTTTAAAAGACAACAATGTAACAATAAAATCAAATGAAGAATTATTGAGTAATGAATATTTGTTTGAACCTTCAACAAATAATGCGAAAATTATTAATGAAAATAATATTTCTATTGATAATTTATATGAAGAAAATGAATGCTCAGACTGTTTCCATATGATGAACTTAAATTATACATCATATACAAAAGAATTATGTAAAACCAAAAATTATCCATATATTCACGTAGATATCGAAATTAATTATTTTGAATTTTGGATATCACTAGATCCTATTAGGATACCACAAAAGAGAAAACGCCATTGTGGTATGTATATGAATAAAAAATTATATTATCATGAAAGAAGAAAAGAAAGACAAAAGAAAAACAGGAAAAGAGATAAACGAAAACACACCATAATTAATAATAATAGATTTAATAAAGAAAAATCATTTTGCACATATTATAGTACAAGTAAAGTTCTAGATGAAGAAAAATTAAGTAATACAACAAAATTGTGTAAAAAAATAAAATATTACAAAAATTTGAATACAGATAAACCATATGTATTAGCAACAAAAGGTTGTTTAAAATCAGTTCTTTTTATTTTATTATATAATAGACAAGAAATAAATAATGAAGTGGATAATACATATGAATTCATAAACGATGATTCATCATATGATCCTTTATATGATTATTTGTTAAATAATGAAAAAGATAAGATAAAAGGGAATAATTATTATATTTCTAAAATGTCATCATATAATTCAGAAAATTATATTAAATCACAAATTAAGAATAAAGAACGTCAAAGTAATCATAATTTAAAAAACAAATTAATGGAGGTATTACAATATTCAAATTTAAACAATATTATAAATATAGAAGAATTTTTAAATAGTAAGTTCATATGTTTTCTTAATAATATGTCATCATATCTTCCATGTGTAGTGAATATGAATATCTTTTTATCTCAAATTACAAGTGCAATAGCTAAACCAGTTACCAAAAATTCTTTCTTAAAAGCTTATATTGATTGGAATAATGTACCATATAATAATATATTATTACAACCATTTCGAGCCAATTTTCAATTTGAATTGAAATTACCTAATCCAGAAATATTATTAGAACGCCTTTTAGGAATTAATAAAATCACAAGTTTAAAAGTTATGAAAAATAATAATATAATATATAAAAATGAAGCTGAAGGAATCGATATTAATTGTTCTTTTGAACCTATCGTTCTCAATGTTGATTCTAATACATTAACCTTATTAAATCAATTATATAATATCATTTTTGATTTTTCCAATTATATTTTAAAATTATGTGCTCCAAAAAATAAAGACGCAAATGAACTCTTGATGATTAATTCATTTAATGACAATTTATATGACAACTTTACCGAATATATGATGAGTGATGATTTTTATTTTAATCTTAATGAGTCCTCATTATTATCAACATCATTAAGTAGGTCAACATCCTTATCACTTTCATCTTTTGAATTAAATAATAAATATAAATTAAAGAATATTCAAAGCAATAATGAATTATTATATACTTCAAAGAATACTCATAAATTCCAAATGAACAATGAAGATGATGACAATATGCTTGTTAATTTTAAGTTAAATTATAATAATCATAATCCAGTTAATAAGATAAATAATATAGATGAAAATATAATACCTATTGATGCGGAGGCACCAATGGTAGAACATATTGAACAAAGCGAATTGCAGGTGAAAAAAAATAATCAAGAAAAAGTTTATAGTGAAACAGAAAAAAATAAAGAAGGATTTGATGGAGTATCAATACATGATGAAAAGAAGAAATTAAATGTATCTGATAGAAATATACCTACCGAGGACAATACAAAAAGAAGTTTGAATAAACCATTGATACTTCATCATAAATATAATACTTTTATGAACAATACAAATGAACCAATTTTAAATACAGATTGTAATATGAAGGCCAACAATCAGACATATTTATTAGAATCCGATTTGAGTAATATACTACTATTTAATTTTGTTGAAAATGTTAAGGTTAGTGTTCATTTGAATTTTGATGTAATCCTTTTACAAATATGGGATTCTAATACAGCATATAACAGATGTTCATTCAATTTTGTTATTGAATATATTAATATCCGTTTCAATACTACCAATATATATGAAATGAATTTACAAAATGACAAGTTGGAATTATATAATATAAAAGAAAATAATACCTACGATGAAATATCAAAAAATAAAAAAGATCCTAATGACAATATAAATTATAGAAATAAAAATGAAAATTTTGATCAATCAAATAGGACATTGCTTCAAAGGGTATCTGAAACAAAAAATATACAAGATAGTAATATAGACAAGAGTCATTTAATTCATTCTGGTATAACTAATGATGCATGTTCAAACATGAATTCTGACATAAATAAAAAGAAATATACCTATGGAAAAAGTGCAGTAAATTTTCTAACAAATGAAAATCAATTATATGTTAATAATATAAAAAAAAAATATAGGTCAGTTGCTGGAGATATAAATATAAATATGAACAAAGAAACTAATGATAACATGTTAACCAAAAAAAAGGAACTCGATTATTTTGAAAGTATAGATAACAATATGATGAATTGTAATATAGAAAAGGATAAAATACTAGATTTGTCAAGAAAAGAAAAAATAGATATATATAATTCAACAAAAGAAAACACAAAAATAGTAAATAGTAATCTAGAAAATAATATCAATAATAATAGTAACTATAGTAACAATAAAAATCTGTTTAGTAAATTATTTAATAGGAACAATTTCGTAAAGAATCTGAAACAATTCAAAGAAGAAATAATGAAAATTGTTAAAAAACTTTTTTGTAGAAAAAAGTTGGATGAAATATTAGAAATTATTGAAGAAAAAGGAAATGAACTTGAAACTGAAAATGAAATGTTAGGAAGAAGAAAGAAACATGAGCATTTTAAAAATATTTTTTTAAAAAATGAATTGATGTCTAATGATTTAAAAAAAAATAATGATAAGAAAAAAAGTAAAGACATAGAATGTACAATTGAATTTTTAATATATTGTGAAAATTTTAATAAAAAAGAAAATTCTTATCAAACATTTATTGAACCTATATGTACAGAAATAATAATTATGAAAAAAGAATTAAATACACCAATTCATATATATTATTATTTTTCATGGATCAATATTAACTGGAATTTTAACTTTCTAGATAATATTCTATCTTTATGTTGTAGTATCATTTTTTCTATGATTACACAAAGAACACGATTATTATTAGGAAGACATATGAATGATATAGATAAAAGAAAAAACAGAAAAAAAATAATTCAAGGACATACTAAAAATGAAAAGATGGAAGATGGATCATTGAAAAATCCACATGTTAATACAAATACAACACATCCAACTGAATACGATGATAATAATACAGAGCGAAAAAATTTAAGTGAAATCGATGATACTCAAACATCTATGTATTTACCAAATTATTATGGAGCTCTTAATGATGAACAATATAAAGATTTAGCACTTATAACTAATGAAAATATCCTTTTCAGATATAATAAATTTGATGTATTACATACCGTAAATAAGTCATTAGATACTTACATTTTCGGTGAGTTAATATTAGAAAAACTTCTTCGTAATTATCAATATTCATGTCAATATCCAGGTGATTATTTGTCAAAATATGAAAATGAATTTAATGTGGACAAAGTAAATTCTCCTTCTCAACAATTATTACATATTAAATTATTTAATTATATAAAGAAAAAGAAAAGACTAGAAATGAAAAATATATGTAAGTTAAATAATTTGTTGGGACAACCTATAGCTGTATGCACAATAATAGAAGAAAAAAAAAACATTCAATATAAAAATATAAAAATGACTAAAAGTCAAATGAAATATGAAAATAATAAATACAATAGTAAAGAAATTTATAATAATGAAAATACAATTTATGATTATAATAAAAGAAACATATTATTAAATTCTATTAATGCAATAACTAATAGTAATAGGAAGAAATATAATTCATTTAATCCATTCGAAGATTATAAATGTTTACAAATTTCAGATATGAATCATAAAAAATTAGGAAAATGTCAAAATCATAGTGAAGAATTTGGTAATTCAGATAATAGTTCTTTAAATGATACTAAGGATATTTTAAAATATCAATGGAAAATTATTAGTAATAATGAATCGCTTGATTTACCAATTCATGAAAATGGAAAGGTGAAATTTTTTCTAATAAGATTTAGATTACTAAACTACATATATGATATACCGTCAAATATATTAAATACTGAAAAAGAAAATGAAGATGTTATAAGATTAATTATACCTGAGAGAAGATTACCTCCTAATATAAATGAGCAAGTAGAACAAGAATTATATAACCAAGAAATATTAGAAATGGAACAAGAAAATGATACTATGGATTCTAATTCTCATACAATATATCCTACTAATTATCTATTTAAAAAATATGTTACAGGAAAAAATGATACTGTATGGTGTCCTAGTAATATTCCGCAACCACGTAATCATTTATTTATTTATTTTAGAACAAGTAGTAGAATTAGTCATGAGGAAAAACAAGAATATGATTTTTTTCTATCTTCCGTAGTTGCTATTAAAAATAACACTGATTTTCCTCTTTATATATTTAAATCAGTACCAGGTAATGTAAATAAAAGTAGTATCTTTAAGGAATATTTTCACAAATATAATACCAAACCATCTCCACCACCTACGTATACATTGAAGGTTGATAAAAAAATCGAATACTTAATTGAAAACCAGAATAAAGATAAAAATGATTTCTCTAATGGAAATAATGAAAATGAACAAGATAATAATATAGAAAAGAGAAATAATATATTTTACAGCACTGATCAAATTATTATTAATAATAATAATAATAATAACAATTATAATAACAATTATAATAACAATTATAATTATTGTTTTGATCCCAATAAAGACCATAATAGTTATTCAACTAGACAACACAATGAAAAACTTTCTAAACAAAAAACAAAACAAAACAAAAATAAGAACCATAAAAAAAATAATAATAGTAATGATATTTATGAGAAGAAAGATAATAAACAAGGTAATAGTAAAAACTATTCCAATAATCACCAAAATAATAATAAAGATCAGGAAGAAGGAAAGAAAAAAAATATAAACTCATCAAATGAATCTTACTTGCATAATAGAAAAATGATTAAAAATATTTTCCCCTTAATGAAATTGTCTTCTATGGAACATTATATTTCAGGAGATATAAAATTCAGTTCATTAGCTCCATTGAAATATAAAAGAAAGAAAAAGAAAAAGAAGAAGAAAAATAAATTGAAATATGAACTGCAAAAAGAATATATTAAAAAGAAAATAAAAGAATTATATTTAAAAAGTATATGGAAAAAAACACAACGTAAAAAAATTAATAAATATAAAAATGACATCAATAAAATAAATCAAAAGATAAAAGAATATTCATATGAAGAACATGCACCTTACAGAAATATTATGTCTATCAATGATCCTTTTTTTAATTTTATAAAACAAAATTCATCTTCATCTCAAAAATTATCAAATAATTCATTAAATATTTCAGATGTCACAAATATTACATTATCATCCTTATCAAGTACTTTTTTTTCTATCCATTCAAGTCAAAGTAATAATTCAACATATGAAGATTCTTTATATGATTATGATAAAGATAATTGCTTAGAACAAGCATTATTAGATGAATATTTTAAAAAGCAAAATGATGATGATGTTTTATTGGATATTATTGAAATATCTAATAATAGTAATAAATTAATTCCAATACCATTATATTGGCTAGTTGCTGAAAATTCATTCATATGGTTATCTATACAAAATGAAGAAATTTATAAAAATAAAGATTTGTATAATTTTTTAAATGAAGAAATGAATGCAGACTATTTATGTACAAAAAAAACATTAGATAATATCTTAAATTATCAATCTCCTTTTGTAGCTGATACACTTAAATTATTTAAACCATATTTGATTAAATTAAAAAATCATTTATTGAAAGAAGATAATTTATTTTCAAAAAATAAAAAAGATGAAAATGAAACTTTTATGACCAATTATAATACAAACTATATGAATTTAAAAGAAGTGTATTTCACTTCATCAGTTGTTAGTGTCTATAATCGTAATTATGTACTAAATCGAAAAGATGCTCACAATTTTTTTCAAATAAGTATTGAACATGTTCTAATGATCAATAATGGATTACCTAAAACTATGTATATAAATTGTGAAGTCTTAAAAAGTAAGCTCAAAAAGAAAAATACTATAAAGAAAAACATAAGTCACTATAAGAGTTTTCCATCAATCGAAAAATCTAAGCAATATAAAGATTCAGAAAATAGAGAAGACTCTTATGTTATAGAAAGTTCTTGTAATAAAACAAATGAAGAATATTCATCATATTATAATGACGATAAAAATGATAATGATAGTAAATATTATGATGATGGCAAACATGATGATGACAGCAAATATGATGATAACAGCAAATATGATGATAACAGCAAATATGATAATTACAACAACAACCACCATAATAATAATAATAACAATAATGATAATAATAATTTTATGTACAATGAAGGTAGTGACATGTTCTCCCAAAATTCTCATGAAGTTAAAAGAGATAAAGGTGTTGACACCTTCTATGACATACATGACGAATTAACCTTGGAAGAAAAACAAAACATGAAGAAAAAGAAAATTACATTAGTTGTTGATAAATTTAACCAACCATTAAATGAAGATACTAATAAGGATCATATATATATCGTTGATAAATATAGTGATAATAACAGAACAAATAATAAACGAGTTTTTATAAAAGAAGCTAATATTGATAGAAATGTATTTATATATAAAAATACAAAAAAAAATGAAAATGATTACATTGATGACAACGAAAAAAGTAAGAATTATATGGATGATAATAATTTTATTGTTAATATAAAAAAAGAAGAAAATAAAGATATTTATGAAATAAAAATTAATCCTGGACAAAGTATTAAATTACCTTTCCTTCTTCATAATATTACCATATCATTTGATGGATATCATTCAAGAAATATCTTTATAAATTATCCTCATAAAAAAGGAAGGGAAAAAATCATGTTAGAAAGACAAATATCTCCAGATATTTCCTCCTTTTTCACATTTGATTATGAAGAATCCAATGAAAATAATGATGAACATAGTAGTAATAAAAGTACAATGAATTATGTAAATAATAAAAATATTAGTGAAAATACAAATAAAAATTATCATCATTTTGATAGTTCCTTATCTCTTACTGATAAAACAAAAAATCAGTTAACTTATAAAAATAGGAATCATCAAAAAAACGATGATTACCCACTATCTAAATATAGTGATTATCATAAAAGAAAAACAACAAAATATGAAGAATATCAAAAGCAATTATCTATATGGGCAATATTTAATGACTTTTATGAAACAGGCAAAAATCGATATCAAGATATATCATTAGAAAAAATTTACCATGGAACATATTCTATTTCTTGCACATTTTTTGCATCTGCATGTATCATAAATAGGTTAAGTTATCCAATTAAACTGAAACGATTAAATAATGAAAATGTGGTTGTTATTGAACCTTATATTAGAACGTTACTCCCATGTGAATTTTCCATGCGTAGAAATAAAGTAGTAATATCACATGAAACCTTAAAAGATATTAAGAATTGGAAGAATTACCCTTTAAAAAAATATATGATTAAACGAACAAAATTGATTAAAACTCGAAATTTTATTAAGAAGGATAACCAAATAATTGTGCATTCTACAAAATGTGATGATGGTAAACAGATTAAGCATAAATGGGATAATAAGCAGTTAATTCGAGGCGATAAAAATGTAAGCATAAAAAAAAACAAATCTAATGAAGAAATAATTTTAACAAAACAAAATGAAACAACAAATGAAATAAATAATGAAAAGATAAAAAGTGATATAATAAAAAATGATGGTAGTTACCAATATGATAGGAAAGAAGAAGACCATTATCATGTGGACGATAATAATGATGATGATGAAATAGAATATAATAAAAAAAATATTCCTTATAATAAGAATACTCTTTGTGGATATAAATATAAAAAGAAAAGAAGATTCAGATTTTTTTTTTATAGAAATAATATTAATAAACCCTTAGAATCAGAAGATTTTCGTATAACCGATTTATCAGTCACAAGATTAACATGTTCCAATAAAAATAAAAATATACCACAATTAAAATATAGTACTTATATGTCTATTGCTCCTATGCCTTTTTTTAGAACAACCGTTATGGAAATATTACCTTACATTGTTATAATAAATAATACAAAAATGAATATAGTTTTTCAAGAGTTTATAAAAAATTATACATATTTTAAATATAACAAACTACAACCTGGAGATTATGTAGAATTCCATCCACAGTCAAAAAAGAAAGTTATGTTAAAAATGTGCGGTATATATCAAAACACATTCGATCATTTAATTGATGATTATTTACAAAAGGAATTATATTTAGAAAACTTCGAAAAGAAAAATGAAAAATTAATAGATGATATAAAGGCAGACAAACATTATGATAAGAAAAATATGGAAAAAAAAAAAAAATACTCAACAAATAAAGAATCTAAAAAGAAAAAAAAAAATAACATTGCCAATAATTATATATGGAATATATCTAAAAGGATAACTGCATTAAAAGATATGGAAAACGTATATGATAGTAATAATAGTGCAAAATATAATGATAATGAAAAAAATTATATTAATAATAAAATGAAAAACAAAAACAAAAAAAATAAATATAAAAATAATAAGAATAAGAAGAATAATAATGAAAATAATGAAAATAATTATAGCACATCTTCAAAAAACTCATCTAACAGTACCATTGAAAATTATTCAGAAGAAGAAAATTCAAATGGACCTTTGAATTCTTCTAAACATAATAATATTAAAAAACATAAATATATGAATGAATACACATTTTTATACTGGTCTGAAGTTCTAGATTTAAATAGAGTAAGTATGATTTATTTTCGTCATCCAGTTATATCAGAAACCAATTTTAAAATAGAAGAAAAAAAATTACAAAAACTAAATTCTATATTAAGGAAATATGGTAAGAATGATGATGCTTATATAAACATTTTAAAAGAATTAATTAAAAAAAATAATCAAATAACATATAAATATAGTTGTTGTTCTATGGAAATTACAACATTTAAAGGATGTAAATTTGTACGTTTTCAAGATATTGATGTTGTACCTTATTATTTAATAAACTTAACCAAATATAATATAAAATTAAGACAAGTGGGAATTTATGAACATTCAGAAATTCTTCAAAAAATACCTCATAAAAAAAAAACATCATTTAATGAAATATTTAAAAATTATGCATTCAATTTCTCTTTTTACAATCCTTATAAAGACCCTAAATTAAAAGTTACTCTCTTATCGAATAATAAGAAAAATGATTATTATAATAATAACAGCAAATATAAAAAGGAGAAGAAAAGAACACTATTAAAAAAAATACCAAATAAATTAAAACATAATCTTAAGCATGAATTAAATTATCTAAGCTTTTTAACAAAAAATTATATAACAGCGAAAGAAGATGCAAAAATCTTGAATGACAAAATTTTCCAACTCAAAAGATTTTCACAACAAATTGAAAAATTAAATGATGAAAAAATGAATATTATTGATTTAGGAAATAATAATAATATTACCTTACTAAAATTAAATTATAAAAATTCTCTTATTTATATACTATGTACCAAAAGAACATATAATGGAAAAACCATCTATTTATTTGAAAATTATGATGATTTAATAAGTAACATCTTAAATTTTAATACTATATCATTTAATAGTAAATGTAATAGCAAATTGCTAAAATCATTCAGTTTGTTTCGTGATACAATTAAAAGCAAAAATAAATATAGTCGTAAGGATACATATAAGAAAAAAAAGAAACATATTAAATATATGAAAACGTTTATACACAATCCAAATGTTAAATATTTATTAAATTTCTTTAAAAAGAAAAGTTCGGCATCATCTATAGAAAATAAATTAAAAAAAAATATAAAACAAAAGAAAAAAAATAAAATAAACAAGGTATACAAAAATTCAAGAACAAATAAAAGGACAACACATATTATTACGGAATTATACCAAAAAAATATGTTCCACCAAGAAGAAGAAACAAAGAAAAAAAATAAAAATATCATATTAAATATGAATATATTCGCACATTTAATATTTAAAGGCATAGGAATAAGTATTAGTAATCACAATTTAGAAGAAATCCTATATTTTTCGATGGAATATATTTTAATTATTTATAAGCAAATGATAGATCGTGATTTGTTTCATATAAATATAGGATGGTTACAAATAGACAATCATACTAAAAATACTGACTATAAGAATGTGTTGTTACCAATTGTTGATTTACAAAAACAAACAAATAAAAATAACGCAAAGGAAATTAATGAAGAAAAGAAATATTTTATAAATGAGAGTAATATAGCATTATATAATAGTATTAGTAATAATAAAAGAGATGACAAATTTAATAAAATTAATAAGCATAATACCTTTCTCCATAATAGATATGATACAATTTCAGGGCATATAATTTCTGGTGATATCAATTCACGTAATTTAAAATCAAATAATTTCAAATCTTTTAATACTTTTTCTCGTAATATAAATTCTTGTAATACATTTTCTTGTAATATTAATTCTTGTAATATAAATTCTTGTAATATTAATTCTTGTAATATTAATTCTTGTAATACCTTTTCTCGTGATACCTTTCCATATAGTTTTAGATATAGCAATAATTTTTCAAAGTCTAGTACATTAAATATGATTAATTCGTTTTACTCAAATAGAAATAAATCATATAATTCATTTATAGATAATGATGTATTAATTGTTACAAATTTAAATAACCCTCTACATAATTCTTATAAAGTATTTATACCACAATTTTTTTATTCTGAATATAATTCTGTAATATATATTAGTATAATTAAATTAAAAAGAGGAAATATGAAAAATTTCTTAGAATTGTCCGATGTTCAAATAAAAGTTTCTCCTTTGTCTTTAAATATTGATTCCTATTTTATTATCGAAATGTTAAAAATATTCGACGAGTTTTTAAAAATAATCGAATTTGATTTAAGTAACTACAATTCACTTAATTTAGAAAAGAACATTGAATTGACAGATAGAGATTTCGATAGTCTAGAATATAAAAAGGATATAGGTCCTATAGAAGAATTAATTAAATATCATATATTAAAAGATTATTATTTATATGAACAAATAATTAATTTCAAGTTTAATCAAAAAGTAGATGGTATACGAAATCAAACAAATAACAAGTCACATAATTATAAAGAATTAGAAAATTATAACAAAATAAAAAAAAACAGTCAAGATAGTATATTTAATACGAATAATTTATATGACAAATCTAATTTGGATGTTATTGAATTCAAAGAATATATTAATATGCTAAATGAATATGAAAATATTAAAAAAAATAAAGCTATATCAAAAGATATAAGAAATAAGGATATGTTATCCAAAATGCATTGTACTAAAAATAATATATTAAAAAAAGGATGTACCAAAAAAAGTAGAAGAACAAAAAATATAAAATTTTACAACAAAAATGATGATGATAATAATAATAAGGTTAACGATAATAAAAAGGATTACAAAATAGATACACAAATAAATGAATATATAAAGGATAATAATAATAATAATAATAATAATAATAATAATAATACGGATAATAAACAGTACATTAATATAAAAAAAAATATAATAAACATTAACAAGTCCTGTCTAATTAATAATAAAGAGAACATGTATTATGACACAAAGAAATATTCAGATATAGATAAGAAGAAAAAATCAGATGATAATGAAGATACACAAGTTGAACTATTAGATAAAGAGAAAATTGAATATATACAAAGATGTGATAAATATTACAAGAATTCCTTAAATAATAGAAAGAAGATATTGAGAAAAATACAAAATATAAATATAAAAAATAATTTAATAAGTTTTGACAAGATATTCATATCTAAATTAGAAATTAACGAAATGAAATTAATCATAAATATAAAAAATAAAAGTCTGAGTTATGATAAAAAAAGTGAAATAATGGAATCAAATGTAATGAATGCTTTAGTTGTATTAATTATGAATATACCAAATATAAGTGATGCACAATTACATTTTGATAAAGAGGTTAAGAAAAATATCTGTGGATCTTTATATGAACTAATATTTAATGAAATTATGAATGATTATATTACACCTGGAATGAATCAAATGGTTAATGTATTAGGTGCTATAGATTTTATAGGGAATCCATTAATAATATATAAACATTGGAAAAAGGGATATGAAACATTTATACATGATTTAAAAAAAAGTCTAGATTCATGCTCTTTTCCATTATTATTTTGTATCTTATTTTTAAATACATTAGGTAAATTTGGAAAGAGCGTATTATCAGGAATTCTAGAAGGTATTAATAGATTATGTAGTAGCTGGAGTACATGTTTTGAAAGATTTTCTAGAAATTCAGATAACTATAGTATTGTTACGAATAGTAACCTCTTTCAACATGAAATATTAGATCAACCTTCTAATTGTGCAGAAGGTATATATTATGGTTGTCAATCCTTTATAAATATTCTTACTATTAGTTTTGTTAATATAATATATAAACCATTTATAGCTATACAAAAATCAAAAGAATTTAAAAAAAAAACAGAAAAAGGGAAATACAAAATATTTTTTTTATCATTGATAGTTCTATCTTATGGTATATTTAGTGGACTTAGTAGTTTCTTTTTTGGATTTTTTAATAGCGTATTTTCTTTTTTTACATTTCTATTTATTGGTACATTAAATCAAATTCAAACCTTAACAATGAAATCTGTTGTTAGACCAAAAAAAATGAGTATAATAAAAGAATATTCCAAGTTTCTAAATTATGAATATCCATTATCCTTTTCATCTCATGTAATTAATGAAAAAAAAAGGAAGAAAGAACTTTTAAAAAAAAATATTGTTGCTATTATACTTTTATATACTATTAACGATAACACTCCAAATAATATTAAAAGTTATTTATGGGTGAGTAATAACGAATTAGGATATTGTCAAAAGGATAAATTATTATGGTCTTTATATATTAATTGTATA**ATCAAAGTGGATATATTATTAATCGAAATGGATAACGGAAATTTCAAGCGATATTCAATAAGCAGCCGTAAAGAAAATAAATATTTGAATAATATGAATAATAAGGAAAAGAACGATGCTAACAAACACACCAAAAATAGCATGAATAATAATAGTAATAATAATAATAATAATAATTATAATATAAATGAAAAACTTACTTGGAATGAAAAAAATAAAGAACAAAATAATCAAATTAATAATAACAGATCCACATGGAAAATAAAAAATTTATTCACCCCACAACCTTTTAGACAACTTTCCGATTTTAAACCATCGTACTATATTAGAATATTATATAAATCATTATATAAAAATAAGAAGGAAAAAAGAAAAACAAATATTTCTAATTTATTTTTATATAAAAAAAGTGATTCAAATAATAGATTGAAACGTATAATGAGGAAATCCACTTATGAAAAATATTTTGAAAATGAATTTAAAGAAGAACAAAAAGTTCATCGTTATCATACTACTTCTAAAGATATGCCTGAAGAACCAAGTGCTAAGATTTCACATATAAATGACGAAAGAAAAAAAAGAGAAAAAAAAAACAATAAAATTAATATATATGATAACTTATATCCAGACAAAACCAAAAATATTAAACATCATAATACAAAACATTATAAAAAAGAAACAAAAAAAAAACATTTAAATAAAAATAAAAATAATATTACATCATACATTGACGATCAGAATAATCATATGAAATTATGGAAACAAAATAAAAAACATAAAACAAAAAATTATAAGTATAATAATGAACATAATACTGAATATGAACATGTGAATATAAAAAAATTATTTAAATATAACAATCATAAGAAAAAAAAAAATAAATATTTATATATCTCAAAATTGGTCAAATGTGAAAATAAGGAAGTTGCATTACATGCTTTTTCACTTCTTATGTCGTTCTTAATGCACAAATCACCATATTATTTAAAAACCACTAAT**CCTAGGTCAGGATTGAGATCAAGATCTGCTGCTGCTGGTGCTGGTGGTGCTGCTAGAGCTGCTCTGCAGAGAGGAGTACAAGTTGAAACAATATCACCAGGAGATGGTCGTACATTTCCAAAAAGAGGTCAAACTTGTGTTGTACATTATACTGGAATGCTTGAAGATGGAAAGAAATTTGATTCATCTCGTGATAGAAATAAACCATTTAAATTTATGCTAGGTAAACAAGAAGTAATACGAGGTTGGGAAGAAGGAGTTGCTCAAATGAGTGTAGGTCAAAGAGCAAAACTTACTATATCTCCAGATTATGCTTATGGTGCAACTGGACATCCAGGTATAATTCCACCTCATGCAACTCTTGTATTTGATGTGGAGCTTCTAAAACTAGAAACTAGAGGTGTTCAGGTTGAAACAATTTCACCTGGAGATGGCAGAACCTTTCCTAAAAGAGGACAGACTTGCGTAGTTCATTATACAGGCATGCTAGAGGATGGTAAGAAATTTGATTCTAGTCGAGATAGAAATAAGCCATTCAAGTTTATGCTAGGTAAACAGGAAGTAATAAGAGGTTGGGAAGAGGGTGTAGCACAGATGTCAGTTGGACAAAGAGCAAAGTTAACAATATCACCAGATTATGCATACGGTGCAACAGGCCATCCTGGCATCATCCCTCCACATGCAACTTTAGTATTCGACGTTGAATTGTTAAAGTTAGAGACAACGCGTGCTAGAGGTGCTGCTGCTGGTGCTGGAGGTGCAGGTAGACGTACGATGGCAGAAATTGGTACGGGTTTTCCATTTGATCCTCATTATGTGGAGGTGCTGGGGGAAAGGATGCATTATGTTGACGTAGGACCAAGAGATGGTACTCCAGTGTTATTTTTGCATGGAAACCCAACCTCGAGTTATGTATGGAGAAATATAATTCCACATGTAGCACCAACACATAGATGTATAGCTCCTGATTTAATTGGTATGGGAAAAAGTGATAAACCTGACTTAGGATATTTTTTTGATGATCATGTCCGTTTTATGGATGCTTTCATTGAAGCTCTTGGCCTTGAAGAAGTAGTATTAGTTATACATGATTGGGGATCCGCCTTAGGATTTCATTGGGCCAAGAGGAATCCTGAAAGAGTAAAAGGAATAGCATTCATGGAATTCATACGACCAATCCCCACATGGGATGAATGGCCAGAATTTGCACGCGAAACATTTCAAGCTTTTAGAACTACAGATGTTGGTAGAAAATTAATAATAGATCAAAATGTATTTATAGAAGGAACTTTACCTATGGGTGTTGTAAGGCCGTTAACAGAAGTTGAAATGGACCACTACCGTGAACCTTTTTTAAATCCAGTAGATAGAGAGCCCTTATGGAGATTTCCTAATGAATTACCTATTGCAGGTGAACCCGCGAATATTGTTGCTTTAGTAGAAGAATATATGGATTGGTTACATCAGTCTCCTGTTCCTAAACTTCTATTTTGGGGTACACCTGGAGTTCTAATACCACCAGCTGAAGCAGCAAGATTAGCAAAATCATTACCAAATTGTAAAGCTGTTGATATAGGTCCTGGGTTGAATTTATTACAAGAAGATAATCCAGATTTGATTGGATCTGAGATAGCTAGATGGCTAAGTACATTAGAAATTTCAGGTACCGGTGCCAGGGGAGCAGCCGCAGGAGCAGGGGGGGCAGGAAGGCGTGGTGTTCAGGTCGAGACTATTAGCCCTGGAGATGGACGCACGTTTCCTAAGCGTGGACAGACATGCGTAGTTCACTACACAGGTATGTTGGAGGACGGTAAAAAGTTCGACAGCTCACGCGACCGCAATAAACCTTTCAAGTTTATGCTTGGCAAGCAGGAGGTTATTCGTGGATGGGAGGAGGGTGTAGCACAGATGTCTGTTGGACAGCGTGCTAAGTTGACAATTTCACCTGACTATGCTTATGGCGCTACGGGCCATCCCGGGATCATTCCGCCACATGCGACTCTGGTATTCGACGTTGAATTATTAAAGTTAGAGACAGCTAGAGGGGCCGCTGCAGGTGCTGGTGGAGCTGGAAGACGTGGAGTACAAGTAGAGACTATCTCTCCAGGTGACGGTCGCACTTTCCCAAAGCGTGGCCAAACCTGTGTTGTACATTACACTGGTATGCTGGAGGATGGGAAAAAGTTCGATTCCAGTCGCGACCGTAACAAACCGTTCAAATTCATGTTGGGAAAGCAGGAAGTGATCCGCGGGTGGGAGGAAGGCGTGGCGCAAATGAGCGTCGGTCAGCGGGCTAAATTGACCATTTCCCCTGACTACGCGTATGGGGCTACTGGGCACCCAGGGATTATTCCGCCTCACGCTACACTTGTGTTTGATGTCGAACTTTTGAAACTGGAAACTGTCGACGGAGAAGGAAGAGGAAGTTTATTAACATGTGGAGATGTAGAAGAAAATCCAGGACCAATGATTGAACAAGATGGATTGCACGCAGGTTCTCCGGCCGCTTGGGTGGAGAGGCTATTCGGCTATGACTGGGCACAACAGACAATCGGCTGCTCTGATGCCGCCGTGTTCCGGCTGTCAGCGCAGGGGCGCCCGGTTCTTTTTGTCAAGACCGACCTGTCCGGTGCCCTGAATGAACTGCAGGACGAGGCAGCGCGGCTATCGTGGCTGGCCACGACGGGCGTTCCTTGCGCAGCTGTGCTCGACGTTGTCACTGAAGCGGGAAGGGACTGGCTGCTATTGGGCGAAGTGCCGGGGCAGGATCTCCTGTCATCTCACCTTGCTCCTGCCGAGAAAGTATCCATCATGGCTGATGCAATGCGGCGGCTGCATACGCTTGATCCGGCTACCTGCCCATTCGACCACCAAGCGAAACATCGCATCGAGCGAGCACGTACTCGGATGGAAGCCGGTCTTGTCGATCAGGATGATCTGGACGAAGAGCATCAGGGGCTCGCGCCAGCCGAACTGTTCGCCAGGCTCAAGGCGCGCATGCCCGACGGCGAGGATCTCGTCGTGACCCATGGCGATGCCTGCTTGCCGAATATCATGGTGGAAAATGGCCGCTTTTCTGGATTCATCGACTGTGGCCGGCTGGGTGTGGCGGACCGCTATCAGGACATAGCGTTGGCTACCCGTGATATTGCTGAAGAGCTTGGCGGCGAATGGGCTGACCGCTTCCTCGTGCTTTACGGTATCGCCGCTCCCGATTCGCAGCGCATCGCCTTCTATCGCCTTCTTGACGAGTTCTTCTAActcgagggatatggcagcttaatgttcgtttttcttatttatatatttataccaattgattgtatttataactgtaaaaatgtgtatgttgtgtgcatatttttttttgtgcatgcacatgcatgtaaatagctaaaattatgaacattttattttttgttcagaaaaaaaaaactttacacacataaaatggctagtatgaatagccatattttatataaattaaatcctatgaatttatgaccatattaaaaatttagatatttatggaacataatatgtttgaaacaataagacaaaattattattattattattatttttactgttataattatgtgtctccttcaatgattcataaatagttggacttgatttttaaaatgtttataatatgattagcatagttaaataaaaaaagttgaaaaattaaaaaaaaacatataaacacaaatgatggtttttccttcaatttcgatatcaatttatagaaacaaaatatatacttgtataattttatttttttatataaatcattacatatataattatacaatattttttctaagagataattatatattaatatatataaaaaaaggtgttttttttttttttttttatttttatttttattttatggtaatattttattttccttattttataaattatattagtttatatgtgattaattttatatattatcaatttatatatttttaaatgcttacttaattatctttttttttttttttttttttttttcccctctttttatattaatttatttttgaaaaaattgatatatatatatatatataatatatatatatacatgtagtagtattaaacaatgtataatatatataaataatatatttatatatttcatttcaattttaattttttttggttttttttttttttctttttgtcatatttaaaaaaaattatattcatataagttatgcattttttataaacattattcaatatatgtataatataatatatatatatatattaatgtattattccaatgtgcatgataaaagaaaaaaataatatttataaaaaaaaagaaaaataaaacaaaaaaagaaaaaaaaaaaaaaaaaaaaaaaaatacaaaaataaataatataatttataattatatattcttgtcacaataaaaatatatatatatatatatatatttataatatgtatattttaaactagaaaaggaataactaatattttatttattatcattcaagatttatattttataataataaatacctaatagaaatatatcaggatccatgcatggttcgctaaactgcatcgtcgctgtgtcccagaacatgggcatcggcaagaacggggactacccctggccaccgctcaggaacgaatttagatatttccagagaatgaccacaacctcttcagtagaaggtaaacagaatctggtgattatgggtaagaagacctggttctccattcctgagaagaatcgacctttaaagggtagaattaatttagttctcagcagagaactcaaggaacctccacaaggagctcattttctttccagaagtctagatgatgccttaaaacttactgaacaaccagaattagcaaataaagtagacatggtctggatagttggtggcagttctgtttataaggaagccatgaatcacccaggccatcttaaactatttgtgacaaggatcatgcaagactttgaaagtgacacgttttttccagaaattgatttggagaaatataaacttctgccagaatacccaggtgttctctctgatgtccaggaggagaaaggcattaagtacaaatttgaagtatatgagaagaatgattaagcttatttaataatagattaaaaatattataaaaataaaaacataaacacagaaattacaaaaaaaatacatatgaattttttttttgtaatcttccttataaatatagaataatgaatcatataaaacatatcattattcatttatttacatttaaaattattgtttcagtatctttaatttattatgtatatataaaaataacttacaattttattaataaacaatatatgtttattaattcatgttttgtaatttatgggatagcgattttttttactgtctgtatttttcttttttaattatgttttaattgtattttatttttattattgttctttttatagtattattttaaaacaaaatgtattttctaagaacttataataataataaatataaattttaataaaaattatatttatcttttacaatatgaacataaagtacaacattaatatatagcttttaatatttttattcctaatcatgtaaatcttaaatttttctttttaaacatatgttaaatatttatttctcattatatataagaacatatttattaaatctagaattctatagtgagtcgtattacaattcactggccgtcgttttacaacgtcgtgactgggaaaaccctggcgttacccaacttaatcgccttgcagcacatccccctttcgccagctggcgtaatagcgaagaggcccgcaccgatcgcccttcccaacagttgcgcagcctgaatggcgaatggcgcctgatgcggtattttctccttacgcatctgtgcggtatttcacaccgcatatggtgcactctcagtacaatctgctctgatgccgcatagttaagccagccccgacacccgccaacacccgctgacgcgccctgacgggcttgtctgctcccggcatccgcttacagacaagctgtgaccgtctccgggagctgcatgtgtcagaggttttcaccgtcatcaccgaaacgcgcgagacgaaagggcctcgtgatacgcctatttttataggttaatgtcatgataataatggtttcttagacgtcaggtggcacttttcggggaaatgtgcgcggaacccctatttgtttatttttctaaatacattcaaatatgtatccgctcatgagacaataaccctgataaatgcttcaataatattgaaaaaggaagagtatgagtattcaacatttccgtgtcgcccttattcccttttttgcggcattttgccttcctgtttttgctcacccagaaacgctggtgaaagtaaaagatgctgaagatcagttgggtgcacgagtgggttacatcgaactggatctcaacagcggtaagatccttgagagttttcgccccgaagaacgttttccaatgatgagcacttttaaagttctgctatgtggcgcggtattatcccgtattgacgccgggcaagagcaactcggtcgccgcatacactattctcagaatgacttggttgagtactcaccagtcacagaaaagcatcttacggatggcatgacagtaagagaattatgcagtgctgccataaccatgagtgataacactgcggccaacttacttctgacaacgatcggaggaccgaaggagctaaccgcttttttgcacaacatgggggatcatgtaactcgccttgatcgttgggaaccggagctgaatgaagccataccaaacgacgagcgtgacaccacgatgcctgtagcaatgccaacaacgttgcgcaaactattaactggcgaactacttactctagcttcccggcaacaattaatagactggatggaggcggataaagttgcaggaccacttctgcgctcggcccttccggctggctggtttattgctgataaatctggagccggtgagcgtgggtctcgcggtatcattgcagcactggggccagatggtaagccctcccgtatcgtagttatctacacgacggggagtcaggcaactatggatgaacgaaatagacagatcgctgagataggtgcctcactgattaagcattggtaactgtcagaccaagtttactcatatatactttagattgatttaaaacttcatttttaatttaaaaggatctaggtgaagatcctttttgataatctcatgaccaaaatcccttaacgtgagttttcgttccactgagcgtcagaccccgtagaaaagatcaaaggatcttcttgagatcctttttttctgcgcgtaatctgctgcttgcaaacaaaaaaaccaccgctaccagcggtggtttgtttgccggatcaagagctaccaactctttttccgaaggtaactggcttcagcagagcgcagataccaaatactgtccttctagtgtagccgtagttaggccaccacttcaagaactctgtagcaccgcctacatacctcgctctgctaatcctgttaccagtggctgctgccagtggcgataagtcgtgtcttaccgggttggactcaagacgatagttaccggataaggcgcagcggtcgggctgaacggggggttcgtgcacacagcccagcttggagcgaacgacctacaccgaactgagatacctacagcgtgagctatgagaaagcgccacgcttcccgaagggagaaaggcggacaggtatccggtaagcggcagggtcggaacaggagagcgcacgagggagcttccagggggaaacgcctggtatctttatagtcctgtcgggtttcgccacctctgacttgagcgtcgatttttgtgatgctcgtcaggggggcggagcctatcgaaaaacgccagcaacgcggcctttttacggttcctggccttttgctggccttttgctcacatgttctttcctgcgttatcccctgattctgtggataaccgtattaccgcctttgagtgagctgataccgctcgccgcagccgaacgaccgagcgcagcgagtcagtgagcgaggaagcggaagagcgcccaatacgcaaaccgcctctccccgcgcgttggccgattcattaatgcagctggcacgacaggtttcccgactggaaagcgggcagtgagcgcaacgcaattaatgtgagttagctcactcattaggcaccccaggctttacactttatgcttccggctcgtatgttgtgtggaattgtgagcggataacaatttcacacaggaaacagctatgaccatgattacgccaagctatttaggtgacactatagaagcggccgctagATCAAAGTGGATATATTATTAATCGAAATGGATAACGGAAATTTCAAGCGATATTCAATAAGCAGCCGTAAAGAAAATAAATATTTGAATAATATGAATAATAAGGAAAAGAACGATGCTAACAAACACACCAAAAATAGCATGAATAATAATAGTAATAATAATAATAATAATAATTATAATATAAATGAAAAACTTACTTGGAATGAAAAAAATAAAGAACAAAATAATCAAATTAATAATAACAGATCCACATGGAAAATAAAAAATTTATTCACCCCACAACCTTTTAGACAACTTTCCGATTTTAAACCATCGTACTATATTAGAATATTATATAAATCATTATATAAAAATAAGAAGGAAAAAAGAAAAACAAATATTTCTAATTTATTTTTATATAAAAAAAGTGATTCAAATAATAGATTGAAACGTATAATGAGGAAATCCACTTATGAAAAATATTTTGAAAATGAATTTAAAGAAGAACAAAAAGTTCATCGTTATCATACTACTTCTAAAGATATGCCTGAAGAACCAAGTGCTAAGATTTCACATATAAATGACGAAAGAAAAAAAAGAGAAAAAAAAAACAATAAAATTAATATATATGATAACTTATATCCAGACAAAACCAAAAATATTAAACATCATAATACAAAACATTATAAAAAAGAAACAAAAAAAAAACATTTAAATAAAAATAAAAATAATATTACATCATACATTGACGATCAGAATAATCATATGAAATTATGGAAACAAAATAAAAAACATAAAACAAAAAATTATAAGTATAATAATGAACATAATACTGAATATGAACATGTGAATATAAAAAAATTATTTAAATATAACAATCATAAGAAAAAAAAAAATAAATATTTATATATCTCAAAATTGGTCAAATGTGAAAATAAGGAAGTTGCATTACATGCTTTTTCACTTCTTATGTCGTTCTTAATGCACAAATCACCATATTATTTAAAAACCACTAATTGAatatacacaagataataataacaatattacaaaaaaaaagaaaaatataaatataaataaatatatatatatatatatatatatatatatatatatatttatatggggtgtgtatgttccttttcattatcatattttaaatcaaaaatagagtattataaatattttaaaccttattcttttaaaaatctctttttattgtattataattatatatatatattattaatatttaattacatattatttttctcttttattaattatcggtagttatatatagatattccttgtcaattttttaaaaatttatcttttcttttttttttttaataaaaatatataaattttgtaattaattctcaatatatatttattatacatattttaatttaaaactaatacttattatataatatatagtgtctgttaattattgggcttacattcttattaatttttaatatatatttatatattcatgaatattatatttactt

**5’ amplicon size: 1202bps**

**3’ amplicon size: 1125bps**

### **> PF3D7_1324300 (PfVPS13L6)**

tttttttaaaacaaaggtgtatatatttgttttgttaattttgatacgtttctaatattggtaaattatatttactcatatacaaaatttatgaaataaaattataactcttttgaagattttttaatcataatataaaataattaaattatatacagtaattttttttattattttattattttatttttttatcatttttttttttttttttctatcatctcaatttataatgatgtgagaaattatttatttatttgtttttcttttttttttttctttgttaaaatatactgtgtatgaatatttatacaaatacatactatattgaatgataatatatatatataacaaaaagaaatttataatttgtttgatttaattttattcttttgttaaatgtggccatccgtgttgtatcctcgtgtgggtgcctcattttattatttattctgaaccttcgaaagATGAACTTATTTAATATATTAAGGAACACCTCATTTGAGATAGATGCAATAAAAATAAATTTTCTCTTCTTTAAGAGTTTGATAATTTATAATGTACCTTTAAAAAACATTGAAATAAGAAAGATTCTTTTTCATTTTTATTTTATAAAAAGGTTCCTTTCGTTAATCTTTAACAAGCACGCAAATGAGGATGACAATAATAAAACTGAAATAAAAAAAAAAAATAATAAAATAAAAAAAAATAAAATAAAAAATAATAATAATAATAATAATAATAATAATAATAATAATAATAATAGTAGTAGTAGCAGTAGTTGTTGTAGCAATAGGAATAATGACATATATGATAATTATAATCATCATAATGATAATAAAATAGATAATTATAAAAATTTGAAAAAAGATTATATTGATAATAGACATAATAATAATCATTCCAATAAATATAATAACGAAGAAAATATAAATGTCTCGAAAAAAAACAAAGAGGAGATATTTATAAAATCTATAAAGGTATCCATAAATTGGTCCGAGGTTGAACAATATAATTCTTATTATAATATTTATTTTAAATTAGCTAATCAAAATATTATATGTAGAATATTATATTACATATTTTTTAAGTTATTATTATATTGGTTTATTAAAATATTTTTTAAATTTTATTATATTAAAATTGAACAAGTACATGTAAATATATATTATAACTATAAACATAGGCACATAAAATATTGCGATAATAAAAAAAATATTTCTAAAAGTATAAATGTATATAATATAAAATTTTATGATCTATCTGTTTTTTATAATACACTCAAAGATACTATAACAAAAATTGAACAAAATGATTCTCAAAAGAAAAAACAAATTATATCAAGGAAAGATATCCAAAAATTAAGTGCCCAAGATAATCAAACAAATAACAATAATAATATGGAGAATCACACATATGATAATGTTCAACATAGTACATATCAAAAAATATATAATAATTTATGTTGTTACAACACGTTTAATATATTTTGTAAAAAAATTATTCTTTTTAAAGAAAATAATCACTACAATAATTTAATACACGTCAAACATCCAACCATTATATTGGATAATCATAAGAACAAGTTTTTTTTTCCTTTTTTTCATATAAACCTTAAGACATGCGAGTTAAAAGAAATTTGTTATTTATTAAATTATATAAACAAAGACCATTTTATAAACGATTTAATATATATTATTGAAAGGAATGAACTTGCCAGTTATGTAAATTATAAAGAGAAAAAGAATTCAAATTATGAACACACAACAACAAGACAAAATGAAAATAAGAATTATTGTAAGTACTTTTTAAATTGTTCATTTTTTTTTAATTACTTATTTATTCATTTATATTTAAAAAATAAATTACTATTAAAAAATGAAATTCATAAATTACATATAATAAAAAAAAATTGTGGAAATAATAAGCATTGTAATAATAATGTATGTTATTATAATAATAAAAGTGAAAAAAAATTACCTGACCGTTTCGAGCTATTTTTCAATTATCTCCTGACTTATTCAGAAAAAAATGTAATTCGCATTGGTCAGAAAAAAAAAAAAAAAAAAAAATATAACAATAATAATGATAATAATAATAATAATAATAATAATAATATAATAAACGAAATATATTTAAAAGAGGAGAAGATATGTACTTCATACGATAATCCAAACAATGAGAACAAAATATTAAATAATAAATATAATTATAATGATCTAAATATTGATCCGAAAAGTATTTATGATATGAGTGAGCATCATGAACGTATGATTAAACATGAAGAGGGAGAAAACAAAATGAATTATTATGATGAGGAAATCTTTATGAAAGAAGATGAAAAAGAAAGGAACTTCAATTTAAAAATATATCAAAATTGTAAAAGTTCTTTGTCTTTAATAAAACTTTTTGTAATTTTTGAAATTAAAGAGATGTTGGAATTATACGAAATAGTACAAAAGGAATTATTACTTCCTCATTTGTATATAAATAAAAAAGAGTATGATACAACTACTACAAATAATGATAATAATAAGGATAATTTTTTGGTGAGTTCACTTCTTTCTACCGATATATATATACAAGAATTTAATCTGTTCATAAATAGAAATATAAATAAAAATATAATATTAAATCAAAACACCATTTTCATAAAAAATAGAAATGAAGAGTTGTGTTGTGATAAATCACAAATGGTTGATAAGGAAGAAATTATAAAAATAGAAATTGAAGAGATAAAAAATGAAAATAAGAAGAATGATATATACATAGGAATAAGAGATCAAAGAAGTGTTAATCAATACAATGTCGAGAATACTTATAAGGATATTCCTAAAGAAAATATAGAAAGAGAAAGAAACGTACATTTAAATAATATATTATGCACTAGTGTTTATTATGAAGAGGATGTCTTATATGAATACTCATTATGTGAAAAAATAAATATTATATTAAAAATAGAAAGAGAAGATTCATTTAATAAAAAAATTAATAATGATAAAAACATTATTGTAACGATATATGCTCATAATTTTCTTGGCTATATGTTACCTACACATGTAAATTTTATTTATTATTTATATTCTATATATTATTTAAGAAAAAAAAAAAGTACACATATTTTTAATTGTTGTATGAGAAATGTATTACGAGAAAAATGTCGAGTGAATAAAAAAAAATTATATGAATATAAAAATACTGAGTATAAAGAAAAAGATGAAAAAGATGAAAAAAAAAGTTCCCTTTTTATTATTAAAATAATTAGCAAAAATGTTAAATTACAATTTTTTTGTATTTCAAACAAATATAATGTTCTTAATAATGATTGTAATGTTGTATGTACAAAGAAAGGAAATAATAAAAATGTGATAAATAATAATATAATCTCACCTTTGTGTATTACATTTAGCTTATATTATTTTTTAAAAATGAATTTAAATTATAAACATTTGTCTTATAAAACATATTGTTTGAATAATCATTATTTTTATTTTAGTAATATAAAAGTTCAGTTGTTATCATGTTCTAATATTTTGAGAGATATTTGTATTCTTGAAAAGAAAAAAAAAAATTATACATCATGTAGAAAAAGAAGTAAAGAACTTTCAAATGATGGGAATACATATAAGAACCTTCCTTGTGATATGTATTTAAAAATTAGTGACAAAAAAATTGATATAAATATAATTAATAATGTTAAAATAATTTTTCTTAATTTTATTATATATCAAATATATATAGCTATACGAAGAATTGTTTATTTAGATAATATAAAATTAAATTATAAAAGTACGAAAAAAAAAAAATTCATAAATGAAGAAAAGAGAAAAAGGAAAGTGAAACAAAATATATATCATCATATGTTATTAAAAAAAAATTATTTCTTAAGGAAAACGAAGAATGTGCCATTTGAAAAAAATAAAGAAAAAAAATATATATATGATGATAATAATGATGATAATAATGGAGATAATAATGGAGATAATTATAATATTAAAGATTTATTCCTTCTAAATAAGAATGCGATTTATAATAGCTATTATTATCCATATGATCAACAAGAAGAACATAATAATAATAATAATAATAAGAAGAAGAAGAAGAAGAAGAAGAAAAAAAAAAAAACCGAATCAGATGTAACAATACCTATAAATTACAAAAGAAAAGATCCAAATAAGATATGCACATATTACAATTTTTATAATATGAAAAAAAAAAAAAAGACATCACTATTTGACTTTTCTCACCTTTTAAAATATATTATCCATATAAAATTAAAAAGTCATATTACATTTATATCATTTGATAATGTTTATAATAGTTTAGAAAAATTAATAAATATAAAAAAAAATGAGGATTTATTAAAAAATCAATATGAAATATATTTATGTAGTAATCTATGTATATATTATTTCGATTTGTTACATAGATATAATATAAAAAAGAATAATTTTATTAAGTTGTTCATGGATTTTTATGTATTCAAAATATATAAAAGATTATATAAGAATAAAATATTTAATATGTGTAGTTTTAAAAGATATTATAATAATCATTTGGAAAAAAATGAAAATAACATAAATACAAAGCAACATAAAATAAATAATTATAATACTATTCAAAATAATGAAGAGTCACCTTATTTTCGTTTGATAAGTACAATAAGGAATATAGATTTCTCATATGAAGTATTAAAAAGTTATCATAAGAATCAACATATTCATGATATTAATAAAAAATTTGATGACAAGGTTTATAATATGAAAAAGGGTATTATATATGATGAGTTTATAAGAAACATAAAAAAGTATAATACAAATACAAAAATATATTGTAACCATATTTCTGTTATATATAAGAGCAATGATTTTTTTAATTTTGTCAATAATTATATTTTAAGGGATTTTTTAAATATATTTTCATTAAAGCATCCATATGAAATAAAACAAAAAGAAATGAAATATATAAAAGAAATGAAATATATAAAAGAAATGAAATATATAAAAGAAATGAAATATATAAAAGAAATGAAATATATAAAAGAAATGAAATATATAAAAGAAATGAAATATATAAAAGAAATGAAAGATATAAAAGAAATGAAAGATATGAAAGATATAAAAGAAATGAAAGATATAAAAGAAATGAAAGATATGAAAGATATAAAAGAAATGAAAGATATGAAAGATATAAAAGAAATGAAAGATATGAAAGATATAAAAGATATAAAAGATATAAAAGAAATATTAAATTATAACAATATGATGATAAATACATATGGTAGTAATAAACACTTATGTAATAATCATGAGTATTATAATTTTGTGCATGATGAAGTGTGTAATATGTTCCGTTTACGTAATTTTATTATTTTTAAAAATATGATTAAAAAGGAACATGTGGAACATACTAAAGGATATGAATCGAAGGGTTTTTCTTCATATAATAATATGTATAGAATAGAAAAGAAAAATATGAAAAATATGATTATAAGTTTGAATAAATATAAATTTGATTTTAATAATGTACTTTTATATATTCCATCTATATATAATAATAATAATAAGTTTTTAATTTTTAAAAGTAGAAATATACATATAAAAAATAAATATATATTTAAAAAAGAAGATAATAAAATAAAATTATATGATAAGATATTAGTATATTTTTTAAATAATTATTGTTTTTCATATCCAACAAAAAATAAGGTTATGGATAATTTCCATATTCTTATTCAATATTATAGAAATGTTTTATATAAAAAGAAGAATCCTCTTTTTTTTCTGAAAATATTTTTATGTGGAGTATTACATATGTGTCCAGAAGATTTTAATTTATTACAACAAATATGTGTGGACAATTTGTTAAATAATAATGTGGAATATTTCTATGATAAGTTATATTATAATTTTGTACATAAACAGGAAAATCAAAAAAATAATAAAAAAAATAATAATAATAATAAAAAAAAAAATAATAATAATAATAATAAAAAAAATAATAATAATAATAATAAAAAAGAAATACTTAAATCAATATTACATTGTAATGATAATAAAAATTTTGAGGGTAACCAAGATAAGAATGATCATAGAACTGTGTATAATATAAAGAAAGGGAAATATAGAAAAGATAAAAGAAAGACTTATAAAGATATGCATATAATATTACACTTTTTAAATTTTCAGATATTTATAAAAAATAATTATTTAAAATGTAATATGCTTTATATATATAGTACATATATTAATGTAGATATTATAAAATATATAAATAATAAAGATACTATAATAACACATATAAAAGGATTCATACCTATAGTATTAAGTTATTTATATATGTATCCAAATAATATAATAATGTATCATAAAAAATTGAATAAGAAAGAATATATAAATATTATATCCAGAAAATATAATAGATATCTATATTTTGATATTTCTCAATTTAATATGTTTATAAAAAATGAAGTCAAAAAAAAAAATATATCTATATTCCATGAAAAAAATAATTTATTAAAACAAAATGAGTGTTATGAAATATTTTATAAGAGAACTAAAGATAATTTTATTGGTAAGGAAATATGCAATGATATTAACATAAAAATTAAAAGTCTTTATATAAATTCGGACATTTTTGAATTATATAGTTTATATATTTTTTATTTTGTATGTGATAAAAATCTATTTATATATGAAATAAACAAATTGGTACAGTTTGAAAGGTATATAAAAAGGTTAAATAGGAATATCTTATCATCATTAGATATGAAAAAAAAAAAAAAAAATAATAATAAAAAAAAAAATAATAATATATATAAAAATAAAAATAATAATATATATATAAATAAAAATAATAATATATATAAAAATAAAAATTTATTAATAATAAGGGATAAAGAAAGTTATCATGATAACAATTATTTTGATATGCTAATATATTTATATAAAAAAAAAAAAATGAAAAATATAAAAGATACCGATTTTAATAATTATAGTAGTTATATAAAATATAATAATATTTATAAAATAAATGTAAACATTAAACATTTATCTTTTATATTTAATTTTATGTTTTATGAACATATCATACAACCTACCAAATTATTTTATAAACAAATAGTAAAAAAAAAAAAGAAAAAAGATGTATATAATACTAATAATAAAAGACAATACAAAAATGAATTATGTATTCTGTATAAAGGATATTTCTATGTTAATTTTATATTTAATACATGGAAAAATATGGTTCATTATTTTGTTCATAATAAAAAGTATATGTGTATCAATGGTTATATTTTTGATGTTAATAACATCGTAGAGAATAAAAAAAAAAACGTAAGTAATGTAAATCGTATATGTGAACAAAATAATATGTCATTTTATAAGGATCCTATTTTTGATAATGATATTAATAAAAAAGAAACAAATAAAAATCGACATTATTTTCAAAATAAAGAATATTCCGATATTTTAAGTGTGTTACTACATTTATATGAAAATGTGAATAATATACATAATGAAAATTTAAATATACATAATGGAAATTTAAATATACATAATGAAAATATTAATATATATTTTACGATAAAAAAAAGTAATAAGTTATATTTACTAAGCCCTAACATTTCATATGTTTTAAGTTCAAATATGAATATACAATTAAATTGTCAGTATAATAAGGATATATATTTTGTGGATAATATAAAAAAAATAAAATTCAATTATTTTAATTTTAACGATATATATACAGACCATCTTGGTTATAGAGGGATATCCCCTGCCCATAAAAATAATGACCCTAAAGATGAATATGAAGATATAAATATAAATACAAATGAAAAAATAAATACAAATGAAAAAATAAATACAAATGAAAAAATAAATACAAATGAAAAAATAAATACAAATGAAAAAATAAATACAAATGAAAAAATAAATACAAATGAAAAAATAAATACAAATATATATACAAATATATGTAAAAATAATCCATTTACAAATGTTATTTATGATATCAAATTATTATACCACACGTCCGACGTAAAAACATGTAATAAAAATGAATATCCGTTATTAAATAATGTACATAAAGAATATAATAAATTGGATACATTGAAAACAAATAAGGATTGTTCGAATCCATTTTTTATAAATATAAAAATAAAAAATTTAAATATAAATATCACATTAAAAGATTTATTATATTTAATAAATAAGTTCAAAGAAATAAATTATTTTATAACATATTATGAAGATATACATGTATTATTGTTTTATAAGTTATCTAGAGAAATGTTATGTAACCAATATATATGTAGAATAAAAAAAGAAAAAAGAAATACATCTTCTCATACAATTTTTTTTTTAAATGTAGACATGTACAAATGTTCTTTCAATATATATAAACCTATATGTTGGTATAAACAAAAAGAAATGTATCATAAAAAAAAAATTGTAAAAATAATTTCTAATGGATCTTTTTCTTATACATCTTGTATTAACAAAAAAAAGGATATAGATAATACTATCTTTAAAAAAATGAATATTTCTTTACACACGGATATTTTTTTTTCAAATAATAATATAAATGAAGAATTTGTAAGAAACATATATGTTACATTTATTTATGTAAAACGTAAAGAACAAAATAATCTTTTTGTTATCCTTCAAAATGATAGTATTAATGTATTTATTACAAAAAATGTCTTCAGGAAAATTTTTTATATTTATCAATGTATAACACATGTGAATAATAATATATGTAATGTTAAAAATAAAAAAAAATATAATTGTAATTATAATAAAAAAATATATAATTGTAATTATAATAAAAAAAAATATAATTGTAATTATAATAAAAAAATATATAATTGTAATTATAATAAAAAAAAATATAATTGTAATTATAATAAAAAAAAATATAATTGTAATTATAATAAAAAAAAATATAATTGTAATTATAATAAAAAAATATATAGTTGTAATTATAATAAAAAAATATATAGTTGTAATTATAATAATAGGGACGTGAAAAATTATGACCTTACAAATATCCATATTAATGAAAAGAATAACAAATATGATACATTACAAATTATACAAAATAAAAAAGACCTTTTTCAATATAATAAAGATCATATATATTATAATTCTAGAGACATATACAGTTTATTAAATAATTATAAAGAGAAAATATTTTTTAATCAAAGTGAAGATATATATAATTTGAAAAATGATAAGGAAACCTGCTTACTAATGTTTTATATAAATAAAAAGAAGAAAAAAACAAAGAGTAAAGAATATCCACAAAGTCTAATAAATATAAATGTAGATAAGAACCAAAAAATAGATACAAAACCATTATATATCTTTCCAGTATTAAGAGAAAATTATATAAAGAAAGAGGATATAATTATGTATAGAAAAAATAATATATTTCATTTTTTTACAAATGTATGTATATATAATAATACAAATTATGATATTATATTCTTATATAAAAACTACAAAACTTTATTATATTCTAAAAAAAATAATTATATAAGCAATTTTCAACATGTTGAAGAACCATTCTGTATTTTACTTTTTTATAAAAAGAAATTGTTTGTTTCGAAAAAAATTAATATATATCATAATTTCCAAAAAGGAAGTTATCATAATAGGTTTGATACGAACCACCTTCATATAAAAGATAGAAGACAACATAAGGAAATAAAAAAAACATTTAATTTGTGTGAAAACATAAATAATAACAAAATGAACAGAAATATTAAGGATTGTAATTATTCATCTGGTGGGATATATCATATTCCTTGTAACCAAGGTGTTTGTTATAAAAACCTTGTGTCCTTAAAAGAAGATGAGAAATGGAAAGAATATTTTTCATTAATTCGATGTCACGGGAAAGAACAAGATAAAAAATGTAATAATGATATAAAATATACATGTCACATGCCTAATAATACACATAGAATGTTTGAATTAATTAAAAGAAAATATATTAAATCATTTTTAAACAATACTCATATTAGTTATGATAAAAATGGTATAGAATTTATCATCGACATGACGGCCATACAAAATAATAATTGGCTATATATACATATAGATGCACATGTAAAATTATTAAATTTGCTACCTTTACAAGTATTTTTTAAATGTCGGGATAAGATTCAAACTATTAAATCATTTCAAAAGAAAGATATATTTTTTCAGGACATTACATTAAATGTATCATCAAATGAATTCAAGGATGTAAATAAATTAAAAATAATATATCAAAAAGAAGATGAAACAAGAAAGAAAGGTGATACAATAATAATAAATCATGATAATAAAAACAAAAACAATAATAATAATGATGATGATGATAATAATATAATTCCTTGTGTATATAGAAAACAAAATTTTCATTTAAATTATTTTCGATTTAATAAAGAAATCATTATAACATGTCACACTGTTATTTTTTTATATGATGTACATATGAAAAATATACAAATTAATGATGACAAATTATTTTTTGTGACAAAAGATAAATATAGTAATAGTAATAATATAGCAAATGAAATGAAGGAACCATATATATATCCTCTTAAAAAAATTAAAAAGTTTGAATCATTTTACAACGAACCTTGTAAATATATTATTATGGAGAATACGAAAATTGAGAAAATATATTCTAATACAAATATAGAGATAAAGATAAAAGGAAATAATCAAAGTATTTCAAACTATGTTCATACATATATGATCATATGTTGGAAAAGACAAGATCAATTATCTAATTATATAAATGATAAGCTTCATATGTCTTCCATTTTTCCGTATTATGATACAAATAAGGAAACTAATGATATAACACAAGATAGAAAATGTTATAATAATAATAATAATGATATTAATAATGATAATAATGGTAACAATACTAATTATCTATTTAATAATAACATGTTAAGTCCATACAACAATTTTATTAATCAAATTATATTGATACACCCCTATTTTATTCTTATAAATAATACAAATTTTAATTTGGTCATGTGTACATCTTGTAATTTTTATAATAATAATAATAATAAAAATTATAAAACTAGTGAACGTGAAATAAATAAAAACAAAAGGAATTATTATTTCCCTAGACACTCTAGTCATGTCATCAATTTAATATCTTTAGATGTAGATAGAAATTTTTTTTTTTATATTAGTAATTCTAATAATATTAATAATTATCCTTTTATATGTGGAAAGGTAGATATAACTAAAGAATCGACTATCATCTTATGTTTTCTGAATAATAAAAATATAAAACATAAGGAGAACACAACAAAACACAATCATAACAATAATTATAATATGAACAATAATAATGACATGAAATATATAAACACAGAAAAACAAACATACCACGTAACGTTTATTCATTTAAAAATTAGCATATGCAAAGGATTTAAGTTACAAGGATATTATAAAAAAAATGTATTCCCAAATAGTATGTATATTATTATATCTGAATATGTAAATAATGTGTGTAAATTTGAAAAGTATATATCATATATAAATAATTATAATAGTAATAAAATCTATGTTAATCATAATATACAAAAAAATAATGTTATGAATATATATAAATATATAAAAAATGAATCCCAGGGTGGTCATATAGGTGTGCGAAAATGTAGTAAAAAAAAATGTATACAACATGAAGAATATAATAGAAATGGTGACGAGTATGATGGTGACGAGTATGATGGTGACGAGTATGATGGTGATGAGTGTGATGGTGATGAGTATGATGGTGATGAGTATGATGGTGATGAGTATGATGGTGATGAGTATGATGGTGATGAGCATAATGGTGATGAGTATAATGGCGATGAGTATAATGGCGATGAGTATAATGGCGATGAGTATAATGGCGATGAGTATAATGGCGATGAGTATAATGGCGATGAGTATAATGGTGATGAGTATAATGGTGATGAAGAAAGTGTAAAAAAAATATTTATCTATAATGATACCGAAAGAGAAATCCTTTTCGATATATTTGATTTAAGCGAAATGATTTACATAATCAAAAAGACAATGGACGAGGCAAAAAACAAAAACGAACATAATAATGATAATAATAATTATTGTATTCGTAATGTTCCTTTTGATGAGAAAGAAACCGTTCCTTCCATCAAAGAAAAAGAAAAAACACACGATTTGATTAAAACAAAAAAAAAAATGAAAGGAAATTTTTTCATAAACATTTTTAATTTGTATAGTAGCGATGAAAATACTTCTGGAGAGGAAAAAAAAAATAGTTCTCAAGAATTTATAGAATATGATATACAAAATAAAACAAATGTATTATCTTCTAAAAAATATATTGATTTTATATACAAAAATGATAATTCGTTTAAGAAATATATAAAGGATAATGAACATCTTTTTCGGCATGTTAAGAAAAAAAGCATTGCCTTCTTTCCCATGAGGAAAAATAATGGTATTAAAGATTGTTTTTTTATAATTAATACAAATTATGAAGAAGATGATGAGAAATACTATCCTAAAGATTATCATGATGAAAAAAATGATGTTCGCTACAATAACAATAATTGTGATTATGATAACTATCGATCTTATAAACTTTTTCATTTTTCCTTAAAAAATAAAAAGAATCATAACAAAAGATATGTGATTAAATACACCAGAAAAGAAGAAAAGAAAACGAATTACATACAAATATTACTAATACAAAAAAAAAAAAAATTATTCATTAATATTTCAAACATGGTAGATAATCTCATACAAGTCAAAGGAAAGATAAACATGAACATTCTACATAAACAAAATCAAGATGTTATAAAAACAGAAGAAGATATATCAACATGTTGTAATATGAATAAAACGTTTAAGTGTAATGTGATCTCTCCAAAAAATGTGTGTAACGAATCTCATATAGTACAAAATTTAGTATATAACAAATTTATTCAAATAATAAAAAGATACAGAAGGAAAAAAAAAATTGGAATCATAGGAAGAAATAATAAGGGGTACTTATTTTTAAATAATATATATAGAAATATTATTAATAATTGTAAGGATGATAAATATAAACATCTTTTTAAATTTTTTAAAAAAAATTGTTTTCTAAATCAAGGTATGACATATATTGTTTGGAAAGTATTGACTTATGTTCATAAGGAACGTATAAAAGTTCAAGCGAACATAAAAGATGATACAACAAAACAAAATAATAATAATAATAATAATAATAATCGTTATCATAATAATAATCGTTATCATAATAATAATCGTTATCATAATAATAATCGTTATCATAATAATAATCATCATCATCATAATAATAACCGGTGTAATGATTTTCATATGATAAAATCACATCTTTTTGGATATCATACACATAGCAGGAATATACACCTATATGAAAAAAATAATAAAAATATATTTTATATTATAAAGAATAAGCAAGAAAAAGATAATAAGGAGGCCCTGCTACCCATAGATGAATATAATTATTATGTACATATAAAAAATGATATACGGTTACATTTATTCGCTTCTGTTATAATAGAAGATAATTATATTCATCGAAACAAATATGAATTAGGACATTCGAATGATGTATATAATTTTTCTAAGATAAATTATTATCATAATAAAAATGGAAGTTCCTTAAATATTAGTTCATCACATGATACTTCTTATGAACATTCGTATAATATACATGATAATATAATAAACAGTATTGAGAGTATGGCATTTTTAAAAAAGAAAGATAATATTGATCCATATGATGATTCATATGATAAACATATAAATAATTTTATGAATAACAAAAATATTGAGATAAAGAAAGAGAATACTAATTTCTCATCAATTTTATATAATCAAAGAGAAAATAAAAAAATAAAAAAGAATGATAAAGAAGATATAAATATATATAAAAAAAAAAAAAAAAATGCACATATTTTTACTATAATATTAAATAACATAAATTTGGTGATAAATAAAAAATGTAATTATGTTGAAAAGTATCATATTTTTCTCACCATTAAAAATATCCTTTTACTATATATGTTATCTCACGAAATAAATAAAATATTTTTTAAAAAGGAGGTAGATAAAAATAAAAATGAGGATCATACAAATGATTTGTTTGTAAAAAAGAAAAAAAAAAACTATCCTCAAAAAGACAAATGTGTTAATACATATGATGATTGTAATCATATACATTCTTTAATAAAAATAGAAATACAATACTTTTATGATAATAATAAATGTTGTTTTTATATAAAAAATTTTTCTTTTTTCTTATCACCTATTTATATACGCTTATCATCAACACATGTAGAAGAGATATTTTTTTTTTATGATTACGTTTGTGAATTTATTAAAAGAAAAAGATGGGAAAAAGAAAAAAAAAAGAAAAACAAAAAAGAAAATGTGTACGAAGATAACTTTATCTTTTTTACAATAGAACAAAAGAAATCAAACGAAGAAACAAAAAAAAATATAAGCGAAAATATGAAATATCAAAATGGAATATGTAAAAACATTTGTGTGGGGAAAAAAAATTGTGTGGGGAAAAAAAATTGTGTGGGGAAAAAAAATTGTGTGTGTAAAAACATTTGTGTGGGGAAAAACATTTGTGTGGGGAAAAAAAATTGTGTGGGGAAAAAAAATTGTGTGTGTAAAAACATTTGTGTGGGGAAAAACATATGTGTGTGTAACAAAGAAGTAATATATTTAAACCTGGAAAAAGATACAAATATAACATACCCTTGGTATGTTAAAGAATCTTCATTTTGTTTTAATATAAAAAAGAAGAAGAAAATAAAAATTAATAATATATATATTAGTACATTAAATATTAATTTATCATGTAATAAAATGAAGAGAAATGTAAATATATATATAAAGAAAAGAGACCCCCCAAAAGGAAAAGATAATATATATAATATTTTAAATTATTTTTACGATGTAATAAAAGATGCAGATTTTGTATTAACATCATTAAATTTAAAAAATATTAAAGAAAATGATATGTTGAACTTTTTTAATTACATATTCAATTTTTATTTAAAAGAAATATATAAAAATTTTTTCTTTTATTTAACAAGAATGAATTTAATAAATAAATATCTACATGATTATTATAATTTTTTTCATGTTATTATAGATAATAGTAAAATTTTTAATATCGATATATCAAATAGTATTAAAACATTCAAACAAGAATTACAAAAAAGAGGAAATCAGTTTCTTATTCCACAATACATTAAAGGAGTATATGAAGAAAAAGAATATTCAATACAAGAGAATACTTATAAATATAAAGATATGGATCCTATAGATGATAAACAAAATCAACCTTTTGATATTACACAACAAAAATATATTAATAAAGAAACAGAAATAACAAATGAAATGTTCTTATCCAGAAAATACAAAAGTTTATTATTATTACAAAATAAAAGGAACATCAGTGAACATGAACAAATCTACACATCTGCAGATGCTCTAGAGACTTATAGTGATTCTAATGTATATATAAAATCAAAAAGAAAAAAAAAAATACAAAATGTTAACAAAGAACTTTTTTATATACATAATGGTATAAAAAGATATATATTTAATTACCCAACAGTTAATTCTAATTATTATAAGTATGAAAATAATGATGATCATAATATAATTAATCATATTATTATGGATAAAAATATAAAACAAAATCAACAAATGAAAAACATAAAAGAAAACGAAAAGGAAAAAGAAAAATATAATTATTCAAATTTTTATTTATTTAATAATACTCTAAAATCATTTGGAAAGGATATTATATCAAGTATTAATTATTTTATATATAATAATGATGATCATATGATAAAAAATTATGAAAAAGATACTTTATATTTTGATCTTGATAAGAAGCATATTATAAAGTATTTTACTTCAACAATGAAAAAAATAAAATGTATAAAAATGAAG*gtaacctcacataagcataaaaaaaaaaaaaaaaatgaacatatatattgtatatatatatatatatatatatatatatatatatatattatatgtatatatgtatttatttatttttttattttcttttaattttatatcttcag***GAGTGTTTACATTCATATGAAAAAAAAGATCAATTTATTACCTACTTCCCAATTTATAAAAAAAATACAATTTTTTTTAATCCGTCTTCTATCAAAAATTGGATAATTTTTCAAAACCTTCCACT*gtaaggatattatttataagtctatctgtttatataatctcatggatatataacaacattgtataatatatatatatatttatttattcatttattcatttatttatttatttatttatgtttatatttaatttttttctatgtgattttgtcccttctcag*ATACAACTACATAATCCTCAATTATCTGTATGACAAAAAAGAAGACATGCTAATATTATTTTGTGAAGATACCATTATTCATATAAAGAGTAAGAAAAAACATTGTGTCATAAGAAAAGAAAATATTATAAAAATTGAAATGTGCTGTATTTATTTTAATATAACATTTGAGTGTGATAAAAATATTATCTTTAACATTAAAAATATTATAAAAAAGAATAAAAATAATTTCCTTGGTCTTTATTTTGACATACTTTTTAAAAACAAAAACAAAAAAAGGTTAATAAAAAAAATACAGAAAAACCTCGTAAAAATTTTGTACTTCCTTTATTATGTATTTCATTTTATATATAATTATAATAAAAAAATAGAAAACATGAACAAATTAAAAAATACTAATTTATATCATAATATTAATCATCAACACACAAATAAAAAAAAAAATACTACTTCTTACAAATCTAAAGAACACCCATTTATAACAATATTTGTACAAAATCAAAAAACGATATTTTCGTTTTCATCCATAACATATTTTTATCACTTCTTTATTAACATAAGAGATCAATTATTTAATACACCCAAT**TAAtagaaagatatacaatatgtatgatcatatataaataaaatatatatatatatatataatatatatatatatgtaatgtacataaattttataccttttcaaaatttcttaattacattattaattaaacgtttgtttttttgtgtgatttataattacctatatatgtatatatatccttttatttttaattaaaaaatatttatattatattattatttttttttttttttacgtttatacaatgatatatataatattatgttttcctaaaaaatatatacattactacttgtatatatatatattatataataatttattacttttttatatattctcgctatattttttaatatgtatgaaattaaataattatatatatatatatatatatatattttattattattattttttttttttttttgttgtttttatatcataattaaaaatttaaaactttttttttttttttaatcttttttaaacgaacaaataataatatacttaaatataaaataatgtaggctctagtttatcaataattttaatgtgatatattatatatatatatatatatacaatattattaatatatatattaataatatcaatatatatttttgttattttttttttttcaagatatgattattaaataaaatattcaaatggtaaaataatatttcatatatataatattcttc

**Original Locus (OL) amplicon size: 1131bp**s

**> PF3D7_1324300-Halo-SW Halo Neo-R**

tttttttaaaacaaaggtgtatatatttgttttgttaattttgatacgtttctaatattggtaaattatatttactcatatacaaaatttatgaaataaaattataactcttttgaagattttttaatcataatataaaataattaaattatatacagtaattttttttattattttattattttatttttttatcatttttttttttttttttctatcatctcaatttataatgatgtgagaaattatttatttatttgtttttcttttttttttttctttgttaaaatatactgtgtatgaatatttatacaaatacatactatattgaatgataatatatatatataacaaaaagaaatttataatttgtttgatttaattttattcttttgttaaatgtggccatccgtgttgtatcctcgtgtgggtgcctcattttattatttattctgaaccttcgaaagATGAACTTATTTAATATATTAAGGAACACCTCATTTGAGATAGATGCAATAAAAATAAATTTTCTCTTCTTTAAGAGTTTGATAATTTATAATGTACCTTTAAAAAACATTGAAATAAGAAAGATTCTTTTTCATTTTTATTTTATAAAAAGGTTCCTTTCGTTAATCTTTAACAAGCACGCAAATGAGGATGACAATAATAAAACTGAAATAAAAAAAAAAAATAATAAAATAAAAAAAAATAAAATAAAAAATAATAATAATAATAATAATAATAATAATAATAATAATAATAATAGTAGTAGTAGCAGTAGTTGTTGTAGCAATAGGAATAATGACATATATGATAATTATAATCATCATAATGATAATAAAATAGATAATTATAAAAATTTGAAAAAAGATTATATTGATAATAGACATAATAATAATCATTCCAATAAATATAATAACGAAGAAAATATAAATGTCTCGAAAAAAAACAAAGAGGAGATATTTATAAAATCTATAAAGGTATCCATAAATTGGTCCGAGGTTGAACAATATAATTCTTATTATAATATTTATTTTAAATTAGCTAATCAAAATATTATATGTAGAATATTATATTACATATTTTTTAAGTTATTATTATATTGGTTTATTAAAATATTTTTTAAATTTTATTATATTAAAATTGAACAAGTACATGTAAATATATATTATAACTATAAACATAGGCACATAAAATATTGCGATAATAAAAAAAATATTTCTAAAAGTATAAATGTATATAATATAAAATTTTATGATCTATCTGTTTTTTATAATACACTCAAAGATACTATAACAAAAATTGAACAAAATGATTCTCAAAAGAAAAAACAAATTATATCAAGGAAAGATATCCAAAAATTAAGTGCCCAAGATAATCAAACAAATAACAATAATAATATGGAGAATCACACATATGATAATGTTCAACATAGTACATATCAAAAAATATATAATAATTTATGTTGTTACAACACGTTTAATATATTTTGTAAAAAAATTATTCTTTTTAAAGAAAATAATCACTACAATAATTTAATACACGTCAAACATCCAACCATTATATTGGATAATCATAAGAACAAGTTTTTTTTTCCTTTTTTTCATATAAACCTTAAGACATGCGAGTTAAAAGAAATTTGTTATTTATTAAATTATATAAACAAAGACCATTTTATAAACGATTTAATATATATTATTGAAAGGAATGAACTTGCCAGTTATGTAAATTATAAAGAGAAAAAGAATTCAAATTATGAACACACAACAACAAGACAAAATGAAAATAAGAATTATTGTAAGTACTTTTTAAATTGTTCATTTTTTTTTAATTACTTATTTATTCATTTATATTTAAAAAATAAATTACTATTAAAAAATGAAATTCATAAATTACATATAATAAAAAAAAATTGTGGAAATAATAAGCATTGTAATAATAATGTATGTTATTATAATAATAAAAGTGAAAAAAAATTACCTGACCGTTTCGAGCTATTTTTCAATTATCTCCTGACTTATTCAGAAAAAAATGTAATTCGCATTGGTCAGAAAAAAAAAAAAAAAAAAAAATATAACAATAATAATGATAATAATAATAATAATAATAATAATAATATAATAAACGAAATATATTTAAAAGAGGAGAAGATATGTACTTCATACGATAATCCAAACAATGAGAACAAAATATTAAATAATAAATATAATTATAATGATCTAAATATTGATCCGAAAAGTATTTATGATATGAGTGAGCATCATGAACGTATGATTAAACATGAAGAGGGAGAAAACAAAATGAATTATTATGATGAGGAAATCTTTATGAAAGAAGATGAAAAAGAAAGGAACTTCAATTTAAAAATATATCAAAATTGTAAAAGTTCTTTGTCTTTAATAAAACTTTTTGTAATTTTTGAAATTAAAGAGATGTTGGAATTATACGAAATAGTACAAAAGGAATTATTACTTCCTCATTTGTATATAAATAAAAAAGAGTATGATACAACTACTACAAATAATGATAATAATAAGGATAATTTTTTGGTGAGTTCACTTCTTTCTACCGATATATATATACAAGAATTTAATCTGTTCATAAATAGAAATATAAATAAAAATATAATATTAAATCAAAACACCATTTTCATAAAAAATAGAAATGAAGAGTTGTGTTGTGATAAATCACAAATGGTTGATAAGGAAGAAATTATAAAAATAGAAATTGAAGAGATAAAAAATGAAAATAAGAAGAATGATATATACATAGGAATAAGAGATCAAAGAAGTGTTAATCAATACAATGTCGAGAATACTTATAAGGATATTCCTAAAGAAAATATAGAAAGAGAAAGAAACGTACATTTAAATAATATATTATGCACTAGTGTTTATTATGAAGAGGATGTCTTATATGAATACTCATTATGTGAAAAAATAAATATTATATTAAAAATAGAAAGAGAAGATTCATTTAATAAAAAAATTAATAATGATAAAAACATTATTGTAACGATATATGCTCATAATTTTCTTGGCTATATGTTACCTACACATGTAAATTTTATTTATTATTTATATTCTATATATTATTTAAGAAAAAAAAAAAGTACACATATTTTTAATTGTTGTATGAGAAATGTATTACGAGAAAAATGTCGAGTGAATAAAAAAAAATTATATGAATATAAAAATACTGAGTATAAAGAAAAAGATGAAAAAGATGAAAAAAAAAGTTCCCTTTTTATTATTAAAATAATTAGCAAAAATGTTAAATTACAATTTTTTTGTATTTCAAACAAATATAATGTTCTTAATAATGATTGTAATGTTGTATGTACAAAGAAAGGAAATAATAAAAATGTGATAAATAATAATATAATCTCACCTTTGTGTATTACATTTAGCTTATATTATTTTTTAAAAATGAATTTAAATTATAAACATTTGTCTTATAAAACATATTGTTTGAATAATCATTATTTTTATTTTAGTAATATAAAAGTTCAGTTGTTATCATGTTCTAATATTTTGAGAGATATTTGTATTCTTGAAAAGAAAAAAAAAAATTATACATCATGTAGAAAAAGAAGTAAAGAACTTTCAAATGATGGGAATACATATAAGAACCTTCCTTGTGATATGTATTTAAAAATTAGTGACAAAAAAATTGATATAAATATAATTAATAATGTTAAAATAATTTTTCTTAATTTTATTATATATCAAATATATATAGCTATACGAAGAATTGTTTATTTAGATAATATAAAATTAAATTATAAAAGTACGAAAAAAAAAAAATTCATAAATGAAGAAAAGAGAAAAAGGAAAGTGAAACAAAATATATATCATCATATGTTATTAAAAAAAAATTATTTCTTAAGGAAAACGAAGAATGTGCCATTTGAAAAAAATAAAGAAAAAAAATATATATATGATGATAATAATGATGATAATAATGGAGATAATAATGGAGATAATTATAATATTAAAGATTTATTCCTTCTAAATAAGAATGCGATTTATAATAGCTATTATTATCCATATGATCAACAAGAAGAACATAATAATAATAATAATAATAAGAAGAAGAAGAAGAAGAAGAAGAAAAAAAAAAAAACCGAATCAGATGTAACAATACCTATAAATTACAAAAGAAAAGATCCAAATAAGATATGCACATATTACAATTTTTATAATATGAAAAAAAAAAAAAAGACATCACTATTTGACTTTTCTCACCTTTTAAAATATATTATCCATATAAAATTAAAAAGTCATATTACATTTATATCATTTGATAATGTTTATAATAGTTTAGAAAAATTAATAAATATAAAAAAAAATGAGGATTTATTAAAAAATCAATATGAAATATATTTATGTAGTAATCTATGTATATATTATTTCGATTTGTTACATAGATATAATATAAAAAAGAATAATTTTATTAAGTTGTTCATGGATTTTTATGTATTCAAAATATATAAAAGATTATATAAGAATAAAATATTTAATATGTGTAGTTTTAAAAGATATTATAATAATCATTTGGAAAAAAATGAAAATAACATAAATACAAAGCAACATAAAATAAATAATTATAATACTATTCAAAATAATGAAGAGTCACCTTATTTTCGTTTGATAAGTACAATAAGGAATATAGATTTCTCATATGAAGTATTAAAAAGTTATCATAAGAATCAACATATTCATGATATTAATAAAAAATTTGATGACAAGGTTTATAATATGAAAAAGGGTATTATATATGATGAGTTTATAAGAAACATAAAAAAGTATAATACAAATACAAAAATATATTGTAACCATATTTCTGTTATATATAAGAGCAATGATTTTTTTAATTTTGTCAATAATTATATTTTAAGGGATTTTTTAAATATATTTTCATTAAAGCATCCATATGAAATAAAACAAAAAGAAATGAAATATATAAAAGAAATGAAATATATAAAAGAAATGAAATATATAAAAGAAATGAAATATATAAAAGAAATGAAATATATAAAAGAAATGAAATATATAAAAGAAATGAAATATATAAAAGAAATGAAAGATATAAAAGAAATGAAAGATATGAAAGATATAAAAGAAATGAAAGATATAAAAGAAATGAAAGATATGAAAGATATAAAAGAAATGAAAGATATGAAAGATATAAAAGAAATGAAAGATATGAAAGATATAAAAGATATAAAAGATATAAAAGAAATATTAAATTATAACAATATGATGATAAATACATATGGTAGTAATAAACACTTATGTAATAATCATGAGTATTATAATTTTGTGCATGATGAAGTGTGTAATATGTTCCGTTTACGTAATTTTATTATTTTTAAAAATATGATTAAAAAGGAACATGTGGAACATACTAAAGGATATGAATCGAAGGGTTTTTCTTCATATAATAATATGTATAGAATAGAAAAGAAAAATATGAAAAATATGATTATAAGTTTGAATAAATATAAATTTGATTTTAATAATGTACTTTTATATATTCCATCTATATATAATAATAATAATAAGTTTTTAATTTTTAAAAGTAGAAATATACATATAAAAAATAAATATATATTTAAAAAAGAAGATAATAAAATAAAATTATATGATAAGATATTAGTATATTTTTTAAATAATTATTGTTTTTCATATCCAACAAAAAATAAGGTTATGGATAATTTCCATATTCTTATTCAATATTATAGAAATGTTTTATATAAAAAGAAGAATCCTCTTTTTTTTCTGAAAATATTTTTATGTGGAGTATTACATATGTGTCCAGAAGATTTTAATTTATTACAACAAATATGTGTGGACAATTTGTTAAATAATAATGTGGAATATTTCTATGATAAGTTATATTATAATTTTGTACATAAACAGGAAAATCAAAAAAATAATAAAAAAAATAATAATAATAATAAAAAAAAAAATAATAATAATAATAATAAAAAAAATAATAATAATAATAATAAAAAAGAAATACTTAAATCAATATTACATTGTAATGATAATAAAAATTTTGAGGGTAACCAAGATAAGAATGATCATAGAACTGTGTATAATATAAAGAAAGGGAAATATAGAAAAGATAAAAGAAAGACTTATAAAGATATGCATATAATATTACACTTTTTAAATTTTCAGATATTTATAAAAAATAATTATTTAAAATGTAATATGCTTTATATATATAGTACATATATTAATGTAGATATTATAAAATATATAAATAATAAAGATACTATAATAACACATATAAAAGGATTCATACCTATAGTATTAAGTTATTTATATATGTATCCAAATAATATAATAATGTATCATAAAAAATTGAATAAGAAAGAATATATAAATATTATATCCAGAAAATATAATAGATATCTATATTTTGATATTTCTCAATTTAATATGTTTATAAAAAATGAAGTCAAAAAAAAAAATATATCTATATTCCATGAAAAAAATAATTTATTAAAACAAAATGAGTGTTATGAAATATTTTATAAGAGAACTAAAGATAATTTTATTGGTAAGGAAATATGCAATGATATTAACATAAAAATTAAAAGTCTTTATATAAATTCGGACATTTTTGAATTATATAGTTTATATATTTTTTATTTTGTATGTGATAAAAATCTATTTATATATGAAATAAACAAATTGGTACAGTTTGAAAGGTATATAAAAAGGTTAAATAGGAATATCTTATCATCATTAGATATGAAAAAAAAAAAAAAAAATAATAATAAAAAAAAAAATAATAATATATATAAAAATAAAAATAATAATATATATATAAATAAAAATAATAATATATATAAAAATAAAAATTTATTAATAATAAGGGATAAAGAAAGTTATCATGATAACAATTATTTTGATATGCTAATATATTTATATAAAAAAAAAAAAATGAAAAATATAAAAGATACCGATTTTAATAATTATAGTAGTTATATAAAATATAATAATATTTATAAAATAAATGTAAACATTAAACATTTATCTTTTATATTTAATTTTATGTTTTATGAACATATCATACAACCTACCAAATTATTTTATAAACAAATAGTAAAAAAAAAAAAGAAAAAAGATGTATATAATACTAATAATAAAAGACAATACAAAAATGAATTATGTATTCTGTATAAAGGATATTTCTATGTTAATTTTATATTTAATACATGGAAAAATATGGTTCATTATTTTGTTCATAATAAAAAGTATATGTGTATCAATGGTTATATTTTTGATGTTAATAACATCGTAGAGAATAAAAAAAAAAACGTAAGTAATGTAAATCGTATATGTGAACAAAATAATATGTCATTTTATAAGGATCCTATTTTTGATAATGATATTAATAAAAAAGAAACAAATAAAAATCGACATTATTTTCAAAATAAAGAATATTCCGATATTTTAAGTGTGTTACTACATTTATATGAAAATGTGAATAATATACATAATGAAAATTTAAATATACATAATGGAAATTTAAATATACATAATGAAAATATTAATATATATTTTACGATAAAAAAAAGTAATAAGTTATATTTACTAAGCCCTAACATTTCATATGTTTTAAGTTCAAATATGAATATACAATTAAATTGTCAGTATAATAAGGATATATATTTTGTGGATAATATAAAAAAAATAAAATTCAATTATTTTAATTTTAACGATATATATACAGACCATCTTGGTTATAGAGGGATATCCCCTGCCCATAAAAATAATGACCCTAAAGATGAATATGAAGATATAAATATAAATACAAATGAAAAAATAAATACAAATGAAAAAATAAATACAAATGAAAAAATAAATACAAATGAAAAAATAAATACAAATGAAAAAATAAATACAAATGAAAAAATAAATACAAATGAAAAAATAAATACAAATATATATACAAATATATGTAAAAATAATCCATTTACAAATGTTATTTATGATATCAAATTATTATACCACACGTCCGACGTAAAAACATGTAATAAAAATGAATATCCGTTATTAAATAATGTACATAAAGAATATAATAAATTGGATACATTGAAAACAAATAAGGATTGTTCGAATCCATTTTTTATAAATATAAAAATAAAAAATTTAAATATAAATATCACATTAAAAGATTTATTATATTTAATAAATAAGTTCAAAGAAATAAATTATTTTATAACATATTATGAAGATATACATGTATTATTGTTTTATAAGTTATCTAGAGAAATGTTATGTAACCAATATATATGTAGAATAAAAAAAGAAAAAAGAAATACATCTTCTCATACAATTTTTTTTTTAAATGTAGACATGTACAAATGTTCTTTCAATATATATAAACCTATATGTTGGTATAAACAAAAAGAAATGTATCATAAAAAAAAAATTGTAAAAATAATTTCTAATGGATCTTTTTCTTATACATCTTGTATTAACAAAAAAAAGGATATAGATAATACTATCTTTAAAAAAATGAATATTTCTTTACACACGGATATTTTTTTTTCAAATAATAATATAAATGAAGAATTTGTAAGAAACATATATGTTACATTTATTTATGTAAAACGTAAAGAACAAAATAATCTTTTTGTTATCCTTCAAAATGATAGTATTAATGTATTTATTACAAAAAATGTCTTCAGGAAAATTTTTTATATTTATCAATGTATAACACATGTGAATAATAATATATGTAATGTTAAAAATAAAAAAAAATATAATTGTAATTATAATAAAAAAATATATAATTGTAATTATAATAAAAAAAAATATAATTGTAATTATAATAAAAAAATATATAATTGTAATTATAATAAAAAAAAATATAATTGTAATTATAATAAAAAAAAATATAATTGTAATTATAATAAAAAAAAATATAATTGTAATTATAATAAAAAAATATATAGTTGTAATTATAATAAAAAAATATATAGTTGTAATTATAATAATAGGGACGTGAAAAATTATGACCTTACAAATATCCATATTAATGAAAAGAATAACAAATATGATACATTACAAATTATACAAAATAAAAAAGACCTTTTTCAATATAATAAAGATCATATATATTATAATTCTAGAGACATATACAGTTTATTAAATAATTATAAAGAGAAAATATTTTTTAATCAAAGTGAAGATATATATAATTTGAAAAATGATAAGGAAACCTGCTTACTAATGTTTTATATAAATAAAAAGAAGAAAAAAACAAAGAGTAAAGAATATCCACAAAGTCTAATAAATATAAATGTAGATAAGAACCAAAAAATAGATACAAAACCATTATATATCTTTCCAGTATTAAGAGAAAATTATATAAAGAAAGAGGATATAATTATGTATAGAAAAAATAATATATTTCATTTTTTTACAAATGTATGTATATATAATAATACAAATTATGATATTATATTCTTATATAAAAACTACAAAACTTTATTATATTCTAAAAAAAATAATTATATAAGCAATTTTCAACATGTTGAAGAACCATTCTGTATTTTACTTTTTTATAAAAAGAAATTGTTTGTTTCGAAAAAAATTAATATATATCATAATTTCCAAAAAGGAAGTTATCATAATAGGTTTGATACGAACCACCTTCATATAAAAGATAGAAGACAACATAAGGAAATAAAAAAAACATTTAATTTGTGTGAAAACATAAATAATAACAAAATGAACAGAAATATTAAGGATTGTAATTATTCATCTGGTGGGATATATCATATTCCTTGTAACCAAGGTGTTTGTTATAAAAACCTTGTGTCCTTAAAAGAAGATGAGAAATGGAAAGAATATTTTTCATTAATTCGATGTCACGGGAAAGAACAAGATAAAAAATGTAATAATGATATAAAATATACATGTCACATGCCTAATAATACACATAGAATGTTTGAATTAATTAAAAGAAAATATATTAAATCATTTTTAAACAATACTCATATTAGTTATGATAAAAATGGTATAGAATTTATCATCGACATGACGGCCATACAAAATAATAATTGGCTATATATACATATAGATGCACATGTAAAATTATTAAATTTGCTACCTTTACAAGTATTTTTTAAATGTCGGGATAAGATTCAAACTATTAAATCATTTCAAAAGAAAGATATATTTTTTCAGGACATTACATTAAATGTATCATCAAATGAATTCAAGGATGTAAATAAATTAAAAATAATATATCAAAAAGAAGATGAAACAAGAAAGAAAGGTGATACAATAATAATAAATCATGATAATAAAAACAAAAACAATAATAATAATGATGATGATGATAATAATATAATTCCTTGTGTATATAGAAAACAAAATTTTCATTTAAATTATTTTCGATTTAATAAAGAAATCATTATAACATGTCACACTGTTATTTTTTTATATGATGTACATATGAAAAATATACAAATTAATGATGACAAATTATTTTTTGTGACAAAAGATAAATATAGTAATAGTAATAATATAGCAAATGAAATGAAGGAACCATATATATATCCTCTTAAAAAAATTAAAAAGTTTGAATCATTTTACAACGAACCTTGTAAATATATTATTATGGAGAATACGAAAATTGAGAAAATATATTCTAATACAAATATAGAGATAAAGATAAAAGGAAATAATCAAAGTATTTCAAACTATGTTCATACATATATGATCATATGTTGGAAAAGACAAGATCAATTATCTAATTATATAAATGATAAGCTTCATATGTCTTCCATTTTTCCGTATTATGATACAAATAAGGAAACTAATGATATAACACAAGATAGAAAATGTTATAATAATAATAATAATGATATTAATAATGATAATAATGGTAACAATACTAATTATCTATTTAATAATAACATGTTAAGTCCATACAACAATTTTATTAATCAAATTATATTGATACACCCCTATTTTATTCTTATAAATAATACAAATTTTAATTTGGTCATGTGTACATCTTGTAATTTTTATAATAATAATAATAATAAAAATTATAAAACTAGTGAACGTGAAATAAATAAAAACAAAAGGAATTATTATTTCCCTAGACACTCTAGTCATGTCATCAATTTAATATCTTTAGATGTAGATAGAAATTTTTTTTTTTATATTAGTAATTCTAATAATATTAATAATTATCCTTTTATATGTGGAAAGGTAGATATAACTAAAGAATCGACTATCATCTTATGTTTTCTGAATAATAAAAATATAAAACATAAGGAGAACACAACAAAACACAATCATAACAATAATTATAATATGAACAATAATAATGACATGAAATATATAAACACAGAAAAACAAACATACCACGTAACGTTTATTCATTTAAAAATTAGCATATGCAAAGGATTTAAGTTACAAGGATATTATAAAAAAAATGTATTCCCAAATAGTATGTATATTATTATATCTGAATATGTAAATAATGTGTGTAAATTTGAAAAGTATATATCATATATAAATAATTATAATAGTAATAAAATCTATGTTAATCATAATATACAAAAAAATAATGTTATGAATATATATAAATATATAAAAAATGAATCCCAGGGTGGTCATATAGGTGTGCGAAAATGTAGTAAAAAAAAATGTATACAACATGAAGAATATAATAGAAATGGTGACGAGTATGATGGTGACGAGTATGATGGTGACGAGTATGATGGTGATGAGTGTGATGGTGATGAGTATGATGGTGATGAGTATGATGGTGATGAGTATGATGGTGATGAGTATGATGGTGATGAGCATAATGGTGATGAGTATAATGGCGATGAGTATAATGGCGATGAGTATAATGGCGATGAGTATAATGGCGATGAGTATAATGGCGATGAGTATAATGGCGATGAGTATAATGGTGATGAGTATAATGGTGATGAAGAAAGTGTAAAAAAAATATTTATCTATAATGATACCGAAAGAGAAATCCTTTTCGATATATTTGATTTAAGCGAAATGATTTACATAATCAAAAAGACAATGGACGAGGCAAAAAACAAAAACGAACATAATAATGATAATAATAATTATTGTATTCGTAATGTTCCTTTTGATGAGAAAGAAACCGTTCCTTCCATCAAAGAAAAAGAAAAAACACACGATTTGATTAAAACAAAAAAAAAAATGAAAGGAAATTTTTTCATAAACATTTTTAATTTGTATAGTAGCGATGAAAATACTTCTGGAGAGGAAAAAAAAAATAGTTCTCAAGAATTTATAGAATATGATATACAAAATAAAACAAATGTATTATCTTCTAAAAAATATATTGATTTTATATACAAAAATGATAATTCGTTTAAGAAATATATAAAGGATAATGAACATCTTTTTCGGCATGTTAAGAAAAAAAGCATTGCCTTCTTTCCCATGAGGAAAAATAATGGTATTAAAGATTGTTTTTTTATAATTAATACAAATTATGAAGAAGATGATGAGAAATACTATCCTAAAGATTATCATGATGAAAAAAATGATGTTCGCTACAATAACAATAATTGTGATTATGATAACTATCGATCTTATAAACTTTTTCATTTTTCCTTAAAAAATAAAAAGAATCATAACAAAAGATATGTGATTAAATACACCAGAAAAGAAGAAAAGAAAACGAATTACATACAAATATTACTAATACAAAAAAAAAAAAAATTATTCATTAATATTTCAAACATGGTAGATAATCTCATACAAGTCAAAGGAAAGATAAACATGAACATTCTACATAAACAAAATCAAGATGTTATAAAAACAGAAGAAGATATATCAACATGTTGTAATATGAATAAAACGTTTAAGTGTAATGTGATCTCTCCAAAAAATGTGTGTAACGAATCTCATATAGTACAAAATTTAGTATATAACAAATTTATTCAAATAATAAAAAGATACAGAAGGAAAAAAAAAATTGGAATCATAGGAAGAAATAATAAGGGGTACTTATTTTTAAATAATATATATAGAAATATTATTAATAATTGTAAGGATGATAAATATAAACATCTTTTTAAATTTTTTAAAAAAAATTGTTTTCTAAATCAAGGTATGACATATATTGTTTGGAAAGTATTGACTTATGTTCATAAGGAACGTATAAAAGTTCAAGCGAACATAAAAGATGATACAACAAAACAAAATAATAATAATAATAATAATAATAATCGTTATCATAATAATAATCGTTATCATAATAATAATCGTTATCATAATAATAATCGTTATCATAATAATAATCATCATCATCATAATAATAACCGGTGTAATGATTTTCATATGATAAAATCACATCTTTTTGGATATCATACACATAGCAGGAATATACACCTATATGAAAAAAATAATAAAAATATATTTTATATTATAAAGAATAAGCAAGAAAAAGATAATAAGGAGGCCCTGCTACCCATAGATGAATATAATTATTATGTACATATAAAAAATGATATACGGTTACATTTATTCGCTTCTGTTATAATAGAAGATAATTATATTCATCGAAACAAATATGAATTAGGACATTCGAATGATGTATATAATTTTTCTAAGATAAATTATTATCATAATAAAAATGGAAGTTCCTTAAATATTAGTTCATCACATGATACTTCTTATGAACATTCGTATAATATACATGATAATATAATAAACAGTATTGAGAGTATGGCATTTTTAAAAAAGAAAGATAATATTGATCCATATGATGATTCATATGATAAACATATAAATAATTTTATGAATAACAAAAATATTGAGATAAAGAAAGAGAATACTAATTTCTCATCAATTTTATATAATCAAAGAGAAAATAAAAAAATAAAAAAGAATGATAAAGAAGATATAAATATATATAAAAAAAAAAAAAAAAATGCACATATTTTTACTATAATATTAAATAACATAAATTTGGTGATAAATAAAAAATGTAATTATGTTGAAAAGTATCATATTTTTCTCACCATTAAAAATATCCTTTTACTATATATGTTATCTCACGAAATAAATAAAATATTTTTTAAAAAGGAGGTAGATAAAAATAAAAATGAGGATCATACAAATGATTTGTTTGTAAAAAAGAAAAAAAAAAACTATCCTCAAAAAGACAAATGTGTTAATACATATGATGATTGTAATCATATACATTCTTTAATAAAAATAGAAATACAATACTTTTATGATAATAATAAATGTTGTTTTTATATAAAAAATTTTTCTTTTTTCTTATCACCTATTTATATACGCTTATCATCAACACATGTAGAAGAGATATTTTTTTTTTATGATTACGTTTGTGAATTTATTAAAAGAAAAAGATGGGAAAAAGAAAAAAAAAAGAAAAACAAAAAAGAAAATGTGTACGAAGATAACTTTATCTTTTTTACAATAGAACAAAAGAAATCAAACGAAGAAACAAAAAAAAATATAAGCGAAAATATGAAATATCAAAATGGAATATGTAAAAACATTTGTGTGGGGAAAAAAAATTGTGTGGGGAAAAAAAATTGTGTGGGGAAAAAAAATTGTGTGTGTAAAAACATTTGTGTGGGGAAAAACATTTGTGTGGGGAAAAAAAATTGTGTGGGGAAAAAAAATTGTGTGTGTAAAAACATTTGTGTGGGGAAAAACATATGTGTGTGTAACAAAGAAGTAATATATTTAAACCTGGAAAAAGATACAAATATAACATACCCTTGGTATGTTAAAGAATCTTCATTTTGTTTTAATATAAAAAAGAAGAAGAAAATAAAAATTAATAATATATATATTAGTACATTAAATATTAATTTATCATGTAATAAAATGAAGAGAAATGTAAATATATATATAAAGAAAAGAGACCCCCCAAAAGGAAAAGATAATATATATAATATTTTAAATTATTTTTACGATGTAATAAAAGATGCAGATTTTGTATTAACATCATTAAATTTAAAAAATATTAAAGAAAATGATATGTTGAACTTTTTTAATTACATATTCAATTTTTATTTAAAAGAAATATATAAAAATTTTTTCTTTTATTTAACAAGAATGAATTTAATAAATAAATATCTACATGATTATTATAATTTTTTTCATGTTATTATAGATAATAGTAAAATTTTTAATATCGATATATCAAATAGTATTAAAACATTCAAACAAGAATTACAAAAAAGAGGAAATCAGTTTCTTATTCCACAATACATTAAAGGAGTATATGAAGAAAAAGAATATTCAATACAAGAGAATACTTATAAATATAAAGATATGGATCCTATAGATGATAAACAAAATCAACCTTTTGATATTACACAACAAAAATATATTAATAAAGAAACAGAAATAACAAATGAAATGTTCTTATCCAGAAAATACAAAAGTTTATTATTATTACAAAATAAAAGGAACATCAGTGAACATGAACAAATCTACACATCTGCAGATGCTCTAGAGACTTATAGTGATTCTAATGTATATATAAAATCAAAAAGAAAAAAAAAAATACAAAATGTTAACAAAGAACTTTTTTATATACATAATGGTATAAAAAGATATATATTTAATTACCCAACAGTTAATTCTAATTATTATAAGTATGAAAATAATGATGATCATAATATAATTAATCATATTATTATGGATAAAAATATAAAACAAAATCAACAAATGAAAAACATAAAAGAAAACGAAAAGGAAAAAGAAAAATATAATTATTCAAATTTTTATTTATTTAATAATACTCTAAAATCATTTGGAAAGGATATTATATCAAGTATTAATTATTTTATATATAATAATGATGATCATATGATAAAAAATTATGAAAAAGATACTTTATATTTTGATCTTGATAAGAAGCATATTATAAAGTATTTTACTTCAACAATGAAAAAAATAAAATGTATAAAAATGAAG*gtaacctcacataagcataaaaaaaaaaaaaaaaatgaacatatatattgtatatatatatatatatatatatatatatatatatatattatatgtatatatgtatttatttatttttttattttcttttaattttatatcttcag***GAGTGTTTACATTCATATGAAAAAAAAGATCAATTTATTACCTACTTCCCAATTTATAAAAAAAATACAATTTTTTTTAATCCGTCTTCTATCAAAAATTGGATAATTTTTCAAAACCTTCCACT*gtaaggatattatttataagtctatctgtttatataatctcatggatatataacaacattgtataatatatatatatatttatttattcatttattcatttatttatttatttatttatgtttatatttaatttttttctatgtgattttgtcccttctcag*ATACAACTACATAATCCTCAATTATCTGTATGACAAAAAAGAAGACATGCTAATATTATTTTGTGAAGATACCATTATTCATATAAAGAGTAAGAAAAAACATTGTGTCATAAGAAAAGAAAATATTATAAAAATTGAAATGTGCTGTATTTATTTTAATATAACATTTGAGTGTGATAAAAATATTATCTTTAACATTAAAAATATTATAAAAAAGAATAAAAATAATTTCCTTGGTCTTTATTTTGACATACTTTTTAAAAACAAAAACAAAAAAAGGTTAATAAAAAAAATACAGAAAAACCTCGTAAAAATTTTGTACTTCCTTTATTATGTATTTCATTTTATATATAATTATAATAAAAAAATAGAAAACATGAACAAATTAAAAAATACTAATTTATATCATAATATTAATCATCAACACACAAATAAAAAAAAAAATACTACTTCTTACAAATCTAAAGAACACCCATTTATAACAATATTTGTACAAAATCAAAAAACGATATTTTCGTTTTCATCCATAACATATTTTTATCACTTCTTTATTAACATAAGAGATCAATTATTTAATACACCCAAT**cctaggTCAGGATTGAGATCAAGATCTGCTGCTGCTGGTGCTGGTGGTGCTGCTAGAGCTGCTctgcagAGAGGAGTACAAGTTGAAACAATATCACCAGGAGATGGTCGTACATTTCCAAAAAGAGGTCAAACTTGTGTTGTACATTATACTGGAATGCTTGAAGATGGAAAGAAATTTGATTCATCTCGTGATAGAAATAAACCATTTAAATTTATGCTAGGTAAACAAGAAGTAATACGAGGTTGGGAAGAAGGAGTTGCTCAAATGAGTGTAGGTCAAAGAGCAAAACTTACTATATCTCCAGATTATGCTTATGGTGCAACTGGACATCCAGGTATAATTCCACCTCATGCAACTCTTGTATTTGATGTGGAGCTTCTAAAACTAGAAACTAGAGGTGTTCAGGTTGAAACAATTTCACCTGGAGATGGCAGAACCTTTCCTAAAAGAGGACAGACTTGCGTAGTTCATTATACAGGCATGCTAGAGGATGGTAAGAAATTTGATTCTAGTCGAGATAGAAATAAGCCATTCAAGTTTATGCTAGGTAAACAGGAAGTAATAAGAGGTTGGGAAGAGGGTGTAGCACAGATGTCAGTTGGACAAAGAGCAAAGTTAACAATATCACCAGATTATGCATACGGTGCAACAGGCCATCCTGGCATCATCCCTCCACATGCAACTTTAGTATTCGACGTTGAATTGTTAAAGTTAGAGACAACGCGTGCTAGAGGTGCTGCTGCTGGTGCTGGAGGTGCAGGTAGACGTACGATGGCAGAAATTGGTACGGGTTTTCCATTTGATCCTCATTATGTGGAGGTGCTGGGGGAAAGGATGCATTATGTTGACGTAGGACCAAGAGATGGTACTCCAGTGTTATTTTTGCATGGAAACCCAACCTCGAGTTATGTATGGAGAAATATAATTCCACATGTAGCACCAACACATAGATGTATAGCTCCTGATTTAATTGGTATGGGAAAAAGTGATAAACCTGACTTAGGATATTTTTTTGATGATCATGTCCGTTTTATGGATGCTTTCATTGAAGCTCTTGGCCTTGAAGAAGTAGTATTAGTTATACATGATTGGGGATCCGCCTTAGGATTTCATTGGGCCAAGAGGAATCCTGAAAGAGTAAAAGGAATAGCATTCATGGAATTCATACGACCAATCCCCACATGGGATGAATGGCCAGAATTTGCACGCGAAACATTTCAAGCTTTTAGAACTACAGATGTTGGTAGAAAATTAATAATAGATCAAAATGTATTTATAGAAGGAACTTTACCTATGGGTGTTGTAAGGCCGTTAACAGAAGTTGAAATGGACCACTACCGTGAACCTTTTTTAAATCCAGTAGATAGAGAGCCCTTATGGAGATTTCCTAATGAATTACCTATTGCAGGTGAACCCGCGAATATTGTTGCTTTAGTAGAAGAATATATGGATTGGTTACATCAGTCTCCTGTTCCTAAACTTCTATTTTGGGGTACACCTGGAGTTCTAATACCACCAGCTGAAGCAGCAAGATTAGCAAAATCATTACCAAATTGTAAAGCTGTTGATATAGGTCCTGGGTTGAATTTATTACAAGAAGATAATCCAGATTTGATTGGATCTGAGATAGCTAGATGGCTAAGTACATTAGAAATTTCAGGTACCGGTGCCAGGGGAGCAGCCGCAGGAGCAGGGGGGGCAGGAAGGCGTGGTGTTCAGGTCGAGACTATTAGCCCTGGAGATGGACGCACGTTTCCTAAGCGTGGACAGACATGCGTAGTTCACTACACAGGTATGTTGGAGGACGGTAAAAAGTTCGACAGCTCACGCGACCGCAATAAACCTTTCAAGTTTATGCTTGGCAAGCAGGAGGTTATTCGTGGATGGGAGGAGGGTGTAGCACAGATGTCTGTTGGACAGCGTGCTAAGTTGACAATTTCACCTGACTATGCTTATGGCGCTACGGGCCATCCCGGGATCATTCCGCCACATGCGACTCTGGTATTCGACGTTGAATTATTAAAGTTAGAGACAGCTAGAGGGGCCGCTGCAGGTGCTGGTGGAGCTGGAAGACGTGGAGTACAAGTAGAGACTATCTCTCCAGGTGACGGTCGCACTTTCCCAAAGCGTGGCCAAACCTGTGTTGTACATTACACTGGTATGCTGGAGGATGGGAAAAAGTTCGATTCCAGTCGCGACCGTAACAAACCGTTCAAATTCATGTTGGGAAAGCAGGAAGTGATCCGCGGGTGGGAGGAAGGCGTGGCGCAAATGAGCGTCGGTCAGCGGGCTAAATTGACCATTTCCCCTGACTACGCGTATGGGGCTACTGGGCACCCAGGGATTATTCCGCCTCACGCTACACTTGTGTTTGATGTCGAACTTTTGAAACTGGAAACTGTCGACGGAGAAGGAAGAGGAAGTTTATTAACATGTGGAGATGTAGAAGAAAATCCAGGACCAATGATTGAACAAGATGGATTGCACGCAGGTTCTCCGGCCGCTTGGGTGGAGAGGCTATTCGGCTATGACTGGGCACAACAGACAATCGGCTGCTCTGATGCCGCCGTGTTCCGGCTGTCAGCGCAGGGGCGCCCGGTTCTTTTTGTCAAGACCGACCTGTCCGGTGCCCTGAATGAACTGCAGGACGAGGCAGCGCGGCTATCGTGGCTGGCCACGACGGGCGTTCCTTGCGCAGCTGTGCTCGACGTTGTCACTGAAGCGGGAAGGGACTGGCTGCTATTGGGCGAAGTGCCGGGGCAGGATCTCCTGTCATCTCACCTTGCTCCTGCCGAGAAAGTATCCATCATGGCTGATGCAATGCGGCGGCTGCATACGCTTGATCCGGCTACCTGCCCATTCGACCACCAAGCGAAACATCGCATCGAGCGAGCACGTACTCGGATGGAAGCCGGTCTTGTCGATCAGGATGATCTGGACGAAGAGCATCAGGGGCTCGCGCCAGCCGAACTGTTCGCCAGGCTCAAGGCGCGCATGCCCGACGGCGAGGATCTCGTCGTGACCCATGGCGATGCCTGCTTGCCGAATATCATGGTGGAAAATGGCCGCTTTTCTGGATTCATCGACTGTGGCCGGCTGGGTGTGGCGGACCGCTATCAGGACATAGCGTTGGCTACCCGTGATATTGCTGAAGAGCTTGGCGGCGAATGGGCTGACCGCTTCCTCGTGCTTTACGGTATCGCCGCTCCCGATTCGCAGCGCATCGCCTTCTATCGCCTTCTTGACGAGTTCTTCTAActcgagggatatggcagcttaatgttcgtttttcttatttatatatttataccaattgattgtatttataactgtaaaaatgtgtatgttgtgtgcatatttttttttgtgcatgcacatgcatgtaaatagctaaaattatgaacattttattttttgttcagaaaaaaaaaactttacacacataaaatggctagtatgaatagccatattttatataaattaaatcctatgaatttatgaccatattaaaaatttagatatttatggaacataatatgtttgaaacaataagacaaaattattattattattattatttttactgttataattatgtgtctccttcaatgattcataaatagttggacttgatttttaaaatgtttataatatgattagcatagttaaataaaaaaagttgaaaaattaaaaaaaaacatataaacacaaatgatggtttttccttcaatttcgatatcaatttatagaaacaaaatatatacttgtataattttatttttttatataaatcattacatatataattatacaatattttttctaagagataattatatattaatatatataaaaaaaggtgttttttttttttttttttatttttatttttattttatggtaatattttattttccttattttataaattatattagtttatatgtgattaattttatatattatcaatttatatatttttaaatgcttacttaattatctttttttttttttttttttttttttcccctctttttatattaatttatttttgaaaaaattgatatatatatatatatataatatatatatatacatgtagtagtattaaacaatgtataatatatataaataatatatttatatatttcatttcaattttaattttttttggttttttttttttttctttttgtcatatttaaaaaaaattatattcatataagttatgcattttttataaacattattcaatatatgtataatataatatatatatatatattaatgtattattccaatgtgcatgataaaagaaaaaaataatatttataaaaaaaaagaaaaataaaacaaaaaaagaaaaaaaaaaaaaaaaaaaaaaaaatacaaaaataaataatataatttataattatatattcttgtcacaataaaaatatatatatatatatatatatttataatatgtatattttaaactagaaaaggaataactaatattttatttattatcattcaagatttatattttataataataaatacctaatagaaatatatcaggatccatgcatggttcgctaaactgcatcgtcgctgtgtcccagaacatgggcatcggcaagaacggggactacccctggccaccgctcaggaacgaatttagatatttccagagaatgaccacaacctcttcagtagaaggtaaacagaatctggtgattatgggtaagaagacctggttctccattcctgagaagaatcgacctttaaagggtagaattaatttagttctcagcagagaactcaaggaacctccacaaggagctcattttctttccagaagtctagatgatgccttaaaacttactgaacaaccagaattagcaaataaagtagacatggtctggatagttggtggcagttctgtttataaggaagccatgaatcacccaggccatcttaaactatttgtgacaaggatcatgcaagactttgaaagtgacacgttttttccagaaattgatttggagaaatataaacttctgccagaatacccaggtgttctctctgatgtccaggaggagaaaggcattaagtacaaatttgaagtatatgagaagaatgattaagcttatttaataatagattaaaaatattataaaaataaaaacataaacacagaaattacaaaaaaaatacatatgaattttttttttgtaatcttccttataaatatagaataatgaatcatataaaacatatcattattcatttatttacatttaaaattattgtttcagtatctttaatttattatgtatatataaaaataacttacaattttattaataaacaatatatgtttattaattcatgttttgtaatttatgggatagcgattttttttactgtctgtatttttcttttttaattatgttttaattgtattttatttttattattgttctttttatagtattattttaaaacaaaatgtattttctaagaacttataataataataaatataaattttaataaaaattatatttatcttttacaatatgaacataaagtacaacattaatatatagcttttaatatttttattcctaatcatgtaaatcttaaatttttctttttaaacatatgttaaatatttatttctcattatatataagaacatatttattaaatctagaattctatagtgagtcgtattacaattcactggccgtcgttttacaacgtcgtgactgggaaaaccctggcgttacccaacttaatcgccttgcagcacatccccctttcgccagctggcgtaatagcgaagaggcccgcaccgatcgcccttcccaacagttgcgcagcctgaatggcgaatggcgcctgatgcggtattttctccttacgcatctgtgcggtatttcacaccgcatatggtgcactctcagtacaatctgctctgatgccgcatagttaagccagccccgacacccgccaacacccgctgacgcgccctgacgggcttgtctgctcccggcatccgcttacagacaagctgtgaccgtctccgggagctgcatgtgtcagaggttttcaccgtcatcaccgaaacgcgcgagacgaaagggcctcgtgatacgcctatttttataggttaatgtcatgataataatggtttcttagacgtcaggtggcacttttcggggaaatgtgcgcggaacccctatttgtttatttttctaaatacattcaaatatgtatccgctcatgagacaataaccctgataaatgcttcaataatattgaaaaaggaagagtatgagtattcaacatttccgtgtcgcccttattcccttttttgcggcattttgccttcctgtttttgctcacccagaaacgctggtgaaagtaaaagatgctgaagatcagttgggtgcacgagtgggttacatcgaactggatctcaacagcggtaagatccttgagagttttcgccccgaagaacgttttccaatgatgagcacttttaaagttctgctatgtggcgcggtattatcccgtattgacgccgggcaagagcaactcggtcgccgcatacactattctcagaatgacttggttgagtactcaccagtcacagaaaagcatcttacggatggcatgacagtaagagaattatgcagtgctgccataaccatgagtgataacactgcggccaacttacttctgacaacgatcggaggaccgaaggagctaaccgcttttttgcacaacatgggggatcatgtaactcgccttgatcgttgggaaccggagctgaatgaagccataccaaacgacgagcgtgacaccacgatgcctgtagcaatgccaacaacgttgcgcaaactattaactggcgaactacttactctagcttcccggcaacaattaatagactggatggaggcggataaagttgcaggaccacttctgcgctcggcccttccggctggctggtttattgctgataaatctggagccggtgagcgtgggtctcgcggtatcattgcagcactggggccagatggtaagccctcccgtatcgtagttatctacacgacggggagtcaggcaactatggatgaacgaaatagacagatcgctgagataggtgcctcactgattaagcattggtaactgtcagaccaagtttactcatatatactttagattgatttaaaacttcatttttaatttaaaaggatctaggtgaagatcctttttgataatctcatgaccaaaatcccttaacgtgagttttcgttccactgagcgtcagaccccgtagaaaagatcaaaggatcttcttgagatcctttttttctgcgcgtaatctgctgcttgcaaacaaaaaaaccaccgctaccagcggtggtttgtttgccggatcaagagctaccaactctttttccgaaggtaactggcttcagcagagcgcagataccaaatactgtccttctagtgtagccgtagttaggccaccacttcaagaactctgtagcaccgcctacatacctcgctctgctaatcctgttaccagtggctgctgccagtggcgataagtcgtgtcttaccgggttggactcaagacgatagttaccggataaggcgcagcggtcgggctgaacggggggttcgtgcacacagcccagcttggagcgaacgacctacaccgaactgagatacctacagcgtgagctatgagaaagcgccacgcttcccgaagggagaaaggcggacaggtatccggtaagcggcagggtcggaacaggagagcgcacgagggagcttccagggggaaacgcctggtatctttatagtcctgtcgggtttcgccacctctgacttgagcgtcgatttttgtgatgctcgtcaggggggcggagcctatcgaaaaacgccagcaacgcggcctttttacggttcctggccttttgctggccttttgctcacatgttctttcctgcgttatcccctgattctgtggataaccgtattaccgcctttgagtgagctgataccgctcgccgcagccgaacgaccgagcgcagcgagtcagtgagcgaggaagcggaagagcgcccaatacgcaaaccgcctctccccgcgcgttggccgattcattaatgcagctggcacgacaggtttcccgactggaaagcgggcagtgagcgcaacgcaattaatgtgagttagctcactcattaggcaccccaggctttacactttatgcttccggctcgtatgttgtgtggaattgtgagcggataacaatttcacacaggaaacagctatgaccatgattacgccaagctatttaggtgacactatagaagcggccgcTAgGAGTGTTTACATTCATATGAAAAAAAAGATCAATTTATTACCTACTTCCCAATTTATAAAAAAAATACAATTTTTTTTAATCCGTCTTCTATCAAAAATTGGATAATTTTTCAAAACCTTCCACTgtaaggatattatttataagtctatctgtttatataatctcatggatatataacaacattgtataatatatatatatatttatttattcatttattcatttatttatttatttatttatgtttatatttaatttttttctatgtgattttgtcccttctcagATACAACTACATAATCCTCAATTATCTGTATGACAAAAAAGAAGACATGCTAATATTATTTTGTGAAGATACCATTATTCATATAAAGAGTAAGAAAAAACATTGTGTCATAAGAAAAGAAAATATTATAAAAATTGAAATGTGCTGTATTTATTTTAATATAACATTTGAGTGTGATAAAAATATTATCTTTAACATTAAAAATATTATAAAAAAGAATAAAAATAATTTCCTTGGTCTTTATTTTGACATACTTTTTAAAAACAAAAACAAAAAAAGGTTAATAAAAAAAATACAGAAAAACCTCGTAAAAATTTTGTACTTCCTTTATTATGTATTTCATTTTATATATAATTATAATAAAAAAATAGAAAACATGAACAAATTAAAAAATACTAATTTATATCATAATATTAATCATCAACACACAAATAAAAAAAAAAATACTACTTCTTACAAATCTAAAGAACACCCATTTATAACAATATTTGTACAAAATCAAAAAACGATATTTTCGTTTTCATCCATAACATATTTTTATCACTTCTTTATTAACATAAGAGATCAATTATTTAATACACCCAATTAAtagaaagatatacaatatgtatgatcatatataaataaaatatatatatatatatataatatatatatatatgtaatgtacataaattttataccttttcaaaatttcttaattacattattaattaaacgtttgtttttttgtgtgatttataattacctatatatgtatatatatccttttatttttaattaaaaaatatttatattatattattatttttttttttttttacgtttatacaatgatatatataatattatgttttcctaaaaaatatatacattactacttgtatatatatatattatataataatttattacttttttatatattctcgctatattttttaatatgtatgaaattaaataattatatatatatatatatatatatattttattattattattttttttttttttttgttgtttttatatcataattaaaaatttaaaactttttttttttttttaatcttttttaaacgaacaaataataatatacttaaatataaaataatgtaggctctagtttatcaataattttaatgtgatatattatatatatatatatatatacaatattattaatatatatattaataatatcaatatatatttttgttattttttttttttcaagatatgattattaaataaaatattcaaatggtaaaataatatttcatatatataatattcttc

**5’ amplicon size: 1234bps**

**3’ amplicon size: 994bps**

## **> PF3D7_1364000**

ataaaaaaatatttgaatgaataatttctccttattttatatttcgaaaatggtaatatgaaaaaacaaaatataccatatcttaaatattattatatttgttcatttgataaaagaaatatatatatatatatatatatattatatatatattttattattttatgccgatatgtgtaattaaatatatataaaacataatacagacaaaaaaatgtgaaggcatgcaaatatgtaaaataatatatcgttatataaaatatgaacaaatataaatgtacactaaataaataaataaatatatatatatatatatatatatatatatgtatattttatttaattttttttattttatcctaacatatatatatatatatatatatatatgtgtatatgtttatattttgattatattactttgcaagatatataattagtagtataataccaatgatataataatgcattagatgatgaggtaattatgtataaaaagaATGGAAAATCGAGAATGCGAAATAAGAATATTTAACGGGACTAGTTGTTGTTTTTATAGATATATGCATTCAGTAAAACATGGAGTATGGTTAATTAATCCTCCTGAGTGTTTAAAAAGTAAAGGGGAAGAAAAATGTATAGCTAAATTTACGAAAACTATGGGTAATTTTTATGGTATATTACAATATTGTGTAGTTATAAATAGTGATGAATATATTTTGAATGCTCGTTTTGATATACCATTATTAGGTGATAATACAGCTACATCGATTATTGGTTTGAATATAAAGAATGAAGATAAACAAATTTTATCGCAACAGAATTATTTTTTAGTAAAGTATAGTTTTGATAATACTTATAACAGTAAGTTTTATATTAACATTGTTGAAACTGAAGAAGGACATAGATTTATTGAATACCATAAGAAGAAATTAAAGAAAGGTCTGCTTCGTGTATTACACATACGAACATTAGAAGAAAAGGAAAAATTACGTAAAGAAAATATGAATAATAATTTAAGAATAAACAATTGCTCGAATATTATTAGTAATAATAATAATGATAATAATAATAATAATAATTATTATTATAATAATGAAGACAATTTAACAAATGATAAATTAATACATAATTATCATCGTGATAATGATATATTAGATATATTACCATTAAGTTTTATTATTAATATAAAAAATTATGAATGGAAAAAGAGACTTAGAAAAGCACATAAATCTTTATATATAATAATTATAAATTTTACTAGACAAACATTAAATTTAACAGATTGTAGAGGTATGAAATCTGTTGAATTGAATGAAGGAAATTGGATTGAATTACCTAATGAAAAAATATCTTCTTTACGTTCCTGTGAATTTGGATGTAATAGTGATGGTATGTTTTCTAGTATCAGTGGTTTTTGTAAATATAATTTTATGAACGAATCATGTTATTTAAATATATTATGGGATATTAATAGTGTTAATTCAAAAATTAAATGTAATATTAAAAATACTAGTAAATATACAATTATAAAAAATGTAGAATTATATAATGAATGTACTGCTGTTTTTCATATCCTAGAATATTATTATTATCCTCCAATAAAAATATTAGAATGCAGAGCTTTAACAAATAATGTTATTAATAATTATAATATAAATAAAGATAATATTGATAAAAATATTAATTCTATATACCAATGTAGTATTAATGTTATTAGACAATTTTGTCAATATCTTCTTAATAATAATACATCTCAAGAAAATAATTCAAATGATGAAGACTTCATGAATTCATCAAATTTACAAAAAAAATTTGTATATAATGATGACGACGATGATTTAAAATTTATAAATAATAATATGTCTTCTTTTCCAAAGATTAATAAAAATGCAATTTCCCTTTATACGAACAAAGGCAATGAAATAATTTCTAGTAAAAAAAAAAAAAAAAATCAGAAGAATCCTTATTATAATAGCAAAGTGCCTATT**GCTTCAGAAGATATTTATAAAAAAGGGGAAAAAAATATATATAACAGCGAATATGAAAATGATAATGATAATAATAAAGATTCTTGTCATTTTAATGAGGAGGAGAATGAAGAATTAGAAGAAGATGGAGATTTTTTAAATATCGATCATTTAAAAAGTAGTACTATACAAGAAAAGAATTTAGAAATTTTAAAAAAAAGAAAAGCATGTTCAAACAATATGAATATTAATAGAAAGAATATATTTACGAATATTAATATAACTTGTGGGCGTGATTTTTTTTGTTACATGCATGGGAATAATTATTCTAATAACATACCTATAAATTCATATTTATATATATCTTGGAATATAGGAAATGTGATATATAGAAATTTATATAATTCAAGTGAAAATATAATAATAAATGCTCATAGTTTGTTAAGTAGTCATAATATTAATGATGATGTAATATTAAAATTGAATATAGATGAAAATAATAGAAGGCTTTATCCATCAAATCTAATACATAGTTTATTAATGAATACAGAAAGTAATATATATATAAATAATGAATATGTTATATTAGATATCATTTTAAGTATATTCCATATTATATCAACTAATTTATCACCTATTATTGAAAAAATATTAGATGCAGAATATGGTACTAGGTGGATTATGGATAGCAAAATACCAAAAAGTAATATATGGGAAAAATCAAAAGGAAAAAATGAATTAGACATTGAAGGAATTATACATATCATTACAACCTTCTGGATAGATATATTTGAAAAGCGTATTAAAAATATAGAAATTATTCAAAATTTACAAAAAGCATCGATTTTTTGGTCAAATCAAGAAATAGATCAATTTGATCAAGACTTTGTCAAAAAATTAATCGAAAGCTCTTCCTTACTTTTAAAGCTCTTCGGAGATTATAACACAGCAAAGAGTATTGAAAAGTTGTATTACAATAAATATTCCATTTTTAATGACTTTTGCGTGGAC**TAAgattatatatatgtatatatgtatatatttatatatgtgtatatatatttcacatttttttttttttttttttttttttaatgggttatccataaatacatgctcatggggatatttaacaaaattttcttatgatacatatttggatatttaaccatttatttttatgttgatctataaaaaaagcattcatattgtgatatataatttattctgttaaaaaatttatgtcattatttattatttctttatttattttatttttttttttttttttgtttaaatttttaaaagaatatattccattttttgccattattacatgaattttttttttttttattccttttatatttattcatttttttttattccttttatatttatttattttttttttttttgttattccgcatctcatttgtgatgtactatattatccgtttttttttttttttttttttttttttttttttataatatatcctttttttttgttt

**Original Locus (OL) amplicon size: 1194bp**s

**> PF3D7_1364000-mNG-SW mNG Neo-R**

ataaaaaaatatttgaatgaataatttctccttattttatatttcgaaaatggtaatatgaaaaaacaaaatataccatatcttaaatattattatatttgttcatttgataaaagaaatatatatatatatatatatatattatatatatattttattattttatgccgatatgtgtaattaaatatatataaaacataatacagacaaaaaaatgtgaaggcatgcaaatatgtaaaataatatatcgttatataaaatatgaacaaatataaatgtacactaaataaataaataaatatatatatatatatatatatatatatatgtatattttatttaattttttttattttatcctaacatatatatatatatatatatatatatgtgtatatgtttatattttgattatattactttgcaagatatataattagtagtataataccaatgatataataatgcattagatgatgaggtaattatgtataaaaagaATGGAAAATCGAGAATGCGAAATAAGAATATTTAACGGGACTAGTTGTTGTTTTTATAGATATATGCATTCAGTAAAACATGGAGTATGGTTAATTAATCCTCCTGAGTGTTTAAAAAGTAAAGGGGAAGAAAAATGTATAGCTAAATTTACGAAAACTATGGGTAATTTTTATGGTATATTACAATATTGTGTAGTTATAAATAGTGATGAATATATTTTGAATGCTCGTTTTGATATACCATTATTAGGTGATAATACAGCTACATCGATTATTGGTTTGAATATAAAGAATGAAGATAAACAAATTTTATCGCAACAGAATTATTTTTTAGTAAAGTATAGTTTTGATAATACTTATAACAGTAAGTTTTATATTAACATTGTTGAAACTGAAGAAGGACATAGATTTATTGAATACCATAAGAAGAAATTAAAGAAAGGTCTGCTTCGTGTATTACACATACGAACATTAGAAGAAAAGGAAAAATTACGTAAAGAAAATATGAATAATAATTTAAGAATAAACAATTGCTCGAATATTATTAGTAATAATAATAATGATAATAATAATAATAATAATTATTATTATAATAATGAAGACAATTTAACAAATGATAAATTAATACATAATTATCATCGTGATAATGATATATTAGATATATTACCATTAAGTTTTATTATTAATATAAAAAATTATGAATGGAAAAAGAGACTTAGAAAAGCACATAAATCTTTATATATAATAATTATAAATTTTACTAGACAAACATTAAATTTAACAGATTGTAGAGGTATGAAATCTGTTGAATTGAATGAAGGAAATTGGATTGAATTACCTAATGAAAAAATATCTTCTTTACGTTCCTGTGAATTTGGATGTAATAGTGATGGTATGTTTTCTAGTATCAGTGGTTTTTGTAAATATAATTTTATGAACGAATCATGTTATTTAAATATATTATGGGATATTAATAGTGTTAATTCAAAAATTAAATGTAATATTAAAAATACTAGTAAATATACAATTATAAAAAATGTAGAATTATATAATGAATGTACTGCTGTTTTTCATATCCTAGAATATTATTATTATCCTCCAATAAAAATATTAGAATGCAGAGCTTTAACAAATAATGTTATTAATAATTATAATATAAATAAAGATAATATTGATAAAAATATTAATTCTATATACCAATGTAGTATTAATGTTATTAGACAATTTTGTCAATATCTTCTTAATAATAATACATCTCAAGAAAATAATTCAAATGATGAAGACTTCATGAATTCATCAAATTTACAAAAAAAATTTGTATATAATGATGACGACGATGATTTAAAATTTATAAATAATAATATGTCTTCTTTTCCAAAGATTAATAAAAATGCAATTTCCCTTTATACGAACAAAGGCAATGAAATAATTTCTAGTAAAAAAAAAAAAAAAAATCAGAAGAATCCTTATTATAATAGCAAAGTGCCTATT**GCTTCAGAAGATATTTATAAAAAAGGGGAAAAAAATATATATAACAGCGAATATGAAAATGATAATGATAATAATAAAGATTCTTGTCATTTTAATGAGGAGGAGAATGAAGAATTAGAAGAAGATGGAGATTTTTTAAATATCGATCATTTAAAAAGTAGTACTATACAAGAAAAGAATTTAGAAATTTTAAAAAAAAGAAAAGCATGTTCAAACAATATGAATATTAATAGAAAGAATATATTTACGAATATTAATATAACTTGTGGGCGTGATTTTTTTTGTTACATGCATGGGAATAATTATTCTAATAACATACCTATAAATTCATATTTATATATATCTTGGAATATAGGAAATGTGATATATAGAAATTTATATAATTCAAGTGAAAATATAATAATAAATGCTCATAGTTTGTTAAGTAGTCATAATATTAATGATGATGTAATATTAAAATTGAATATAGATGAAAATAATAGAAGGCTTTATCCATCAAATCTAATACATAGTTTATTAATGAATACAGAAAGTAATATATATATAAATAATGAATATGTTATATTAGATATCATTTTAAGTATATTCCATATTATATCAACTAATTTATCACCTATTATTGAAAAAATATTAGATGCAGAATATGGTACTAGGTGGATTATGGATAGCAAAATACCAAAAAGTAATATATGGGAAAAATCAAAAGGAAAAAATGAATTAGACATTGAAGGAATTATACATATCATTACAACCTTCTGGATAGATATATTTGAAAAGCGTATTAAAAATATAGAAATTATTCAAAATTTACAAAAAGCATCGATTTTTTGGTCAAATCAAGAAATAGATCAATTTGATCAAGACTTTGTCAAAAAATTAATCGAAAGCTCTTCCTTACTTTTAAAGCTCTTCGGAGATTATAACACAGCAAAGAGTATTGAAAAGTTGTATTACAATAAATATTCCATTTTTAATGACTTTTGCGTGGAC**cctaggTCAGGATTGAGATCAAGATCTGCTGCTGCTGGTGCTGGTGGTGCTGCTAGAGCTGCTctgcagAGAGGAGTACAAGTTGAAACAATATCACCAGGAGATGGTCGTACATTTCCAAAAAGAGGTCAAACTTGTGTTGTACATTATACTGGAATGCTTGAAGATGGAAAGAAATTTGATTCATCTCGTGATAGAAATAAACCATTTAAATTTATGCTAGGTAAACAAGAAGTAATACGAGGTTGGGAAGAAGGAGTTGCTCAAATGAGTGTAGGTCAAAGAGCAAAACTTACTATATCTCCAGATTATGCTTATGGTGCAACTGGACATCCAGGTATAATTCCACCTCATGCAACTCTTGTATTTGATGTGGAGCTTCTAAAACTAGAAACTAGAGGTGTTCAGGTTGAAACAATTTCACCTGGAGATGGCAGAACCTTTCCTAAAAGAGGACAGACTTGCGTAGTTCATTATACAGGCATGCTAGAGGATGGTAAGAAATTTGATTCTAGTCGAGATAGAAATAAGCCATTCAAGTTTATGCTAGGTAAACAGGAAGTAATAAGAGGTTGGGAAGAGGGTGTAGCACAGATGTCAGTTGGACAAAGAGCAAAGTTAACAATATCACCAGATTATGCATACGGTGCAACAGGCCATCCTGGCATCATCCCTCCACATGCAACTTTAGTATTCGACGTTGAATTGTTAAAGTTAGAGACAacgcgtGCTAGAGGTGCTGCTGCTGGTGCTGGAGGTGCAGGTAGAcgtacgATGGTAAGTAAGGGAGAAGAAGACAACATGGCATCATTGCCAGCCACCCACGAGTTGCATATTTTCGGTTCTATTAACGGAGTGGACTTTGATATGGTTGGTCAGGGTACAGGAAATCCTAATGATGGTTATGAGGAGCTTAATTTGAAATCTACTAAAGGTGACTTGCAGTTTTCACCTTGGATACTTGTTCCACATATAGGTTACGGTTTTCACCAGTACCTTCCATATCCAGATGGAATGTCTCCATTCCAGGCTGCTATGGTGGATGGAAGTGGTTATCAAGTACATAGGACCATGCAGTTTGAGGACGGTGCATCTTTAACCGTTAACTATAGATATACCTATGAAGGATCTCACATTAAAGGTGAAGCCCAAGTGAAAGGTACCGGATTTCCTGCTGACGGTCCAGTGATGACTAACAGTTTAACTGCCGCAGACTGGTGCCGTTCTAAAAAGACATATCCAAACGATAAGACTATAATTTCTACCTTTAAATGGTCATACACAACAGGAAATGGAAAACGTTATAGATCAACTGCAAGGACTACTTATACATTCGCTAAACCAATGGCCGCCAATTATCTTAAAAATCAACCTATGTATGTTTTCCGTAAAACTGAATTAAAGCACTCAAAGACAGAACTTAATTTTAAGGAATGGCAAAAAGCATTTACTGACGTAATGGGAATGGATGAGTTGTATAAGaccGGTgccaggggagcagccgcaggagcagggggggcaggaaggcgtggtgttcaggtcgagactattagccctggagatggacgcacgtttcctaagcgtggacagacatgcgtagttcactacacaggtatgttggaggacggtaaaaagttcgacagctcacgcgaccgcaataaacctttcaagtttatgcttggcaagcaggaggttattcgtggatgggaggagggtgtagcacagatgtctgttggacagcgtgctaagttgacaatttcacctgactatgcttatggcgctacgggccatcccgggatcattccgccacatgcgactctggtattcgacgttgaattattaaagttagagacagctagaggggccgctgcaggtgctggtggagctggaagacgtggagtacaagtagagactatctctccaggtgacggtcgcactttcccaaagcgtggccaaacctgtgttgtacattacactggtatgctggaggatgggaaaaagttcgattccagtcgcgaccgtaacaaaccgttcaaattcatgttgggaaagcaggaagtgatccgcgggtgggaggaaggcgtggcgcaaatgagcgtcggtcagcgggctaaattgaccatttcccctgactacgcgtatggggctactgggcacccagggattattccgcctcacgctacacttgtgtttgatgtcgaacttttgaaactggaaactgtcgacGGAGAAGGAAGAGGAAGTTTATTAACATGTGGAGATGTAGAAGAAAATCCAGGACCAATGATTGAACAAGATGGATTGCACGCAGGTTCTCCGGCCGCTTGGGTGGAGAGGCTATTCGGCTATGACTGGGCACAACAGACAATCGGCTGCTCTGATGCCGCCGTGTTCCGGCTGTCAGCGCAGGGGCGCCCGGTTCTTTTTGTCAAGACCGACCTGTCCGGTGCCCTGAATGAACTGCAGGACGAGGCAGCGCGGCTATCGTGGCTGGCCACGACGGGCGTTCCTTGCGCAGCTGTGCTCGACGTTGTCACTGAAGCGGGAAGGGACTGGCTGCTATTGGGCGAAGTGCCGGGGCAGGATCTCCTGTCATCTCACCTTGCTCCTGCCGAGAAAGTATCCATCATGGCTGATGCAATGCGGCGGCTGCATACGCTTGATCCGGCTACCTGCCCATTCGACCACCAAGCGAAACATCGCATCGAGCGAGCACGTACTCGGATGGAAGCCGGTCTTGTCGATCAGGATGATCTGGACGAAGAGCATCAGGGGCTCGCGCCAGCCGAACTGTTCGCCAGGCTCAAGGCGCGCATGCCCGACGGCGAGGATCTCGTCGTGACCCATGGCGATGCCTGCTTGCCGAATATCATGGTGGAAAATGGCCGCTTTTCTGGATTCATCGACTGTGGCCGGCTGGGTGTGGCGGACCGCTATCAGGACATAGCGTTGGCTACCCGTGATATTGCTGAAGAGCTTGGCGGCGAATGGGCTGACCGCTTCCTCGTGCTTTACGGTATCGCCGCTCCCGATTCGCAGCGCATCGCCTTCTATCGCCTTCTTGACGAGTTCTTCTAActcgagggatatggcagcttaatgttcgtttttcttatttatatatttataccaattgattgtatttataactgtaaaaatgtgtatgttgtgtgcatatttttttttgtgcatgcacatgcatgtaaatagctaaaattatgaacattttattttttgttcagaaaaaaaaaactttacacacataaaatggctagtatgaatagccatattttatataaattaaatcctatgaatttatgaccatattaaaaatttagatatttatggaacataatatgtttgaaacaataagacaaaattattattattattattatttttactgttataattatgtgtctccttcaatgattcataaatagttggacttgatttttaaaatgtttataatatgattagcatagttaaataaaaaaagttgaaaaattaaaaaaaaacatataaacacaaatgatggtttttccttcaatttcgatatcaatttatagaaacaaaatatatacttgtataattttatttttttatataaatcattacatatataattatacaatattttttctaagagataattatatattaatatatataaaaaaaggtgttttttttttttttttttatttttatttttattttatggtaatattttattttccttattttataaattatattagtttatatgtgattaattttatatattatcaatttatatatttttaaatgcttacttaattatctttttttttttttttttttttttttcccctctttttatattaatttatttttgaaaaaattgatatatatatatatatataatatatatatatacatgtagtagtattaaacaatgtataatatatataaataatatatttatatatttcatttcaattttaattttttttggttttttttttttttctttttgtcatatttaaaaaaaattatattcatataagttatgcattttttataaacattattcaatatatgtataatataatatatatatatatattaatgtattattccaatgtgcatgataaaagaaaaaaataatatttataaaaaaaaagaaaaataaaacaaaaaaagaaaaaaaaaaaaaaaaaaaaaaaaatacaaaaataaataatataatttataattatatattcttgtcacaataaaaatatatatatatatatatatatttataatatgtatattttaaactagaaaaggaataactaatattttatttattatcattcaagatttatattttataataataaatacctaatagaaatatatcaggatccatgcatggttcgctaaactgcatcgtcgctgtgtcccagaacatgggcatcggcaagaacggggactacccctggccaccgctcaggaacgaatttagatatttccagagaatgaccacaacctcttcagtagaaggtaaacagaatctggtgattatgggtaagaagacctggttctccattcctgagaagaatcgacctttaaagggtagaattaatttagttctcagcagagaactcaaggaacctccacaaggagctcattttctttccagaagtctagatgatgccttaaaacttactgaacaaccagaattagcaaataaagtagacatggtctggatagttggtggcagttctgtttataaggaagccatgaatcacccaggccatcttaaactatttgtgacaaggatcatgcaagactttgaaagtgacacgttttttccagaaattgatttggagaaatataaacttctgccagaatacccaggtgttctctctgatgtccaggaggagaaaggcattaagtacaaatttgaagtatatgagaagaatgattaagcttatttaataatagattaaaaatattataaaaataaaaacataaacacagaaattacaaaaaaaatacatatgaattttttttttgtaatcttccttataaatatagaataatgaatcatataaaacatatcattattcatttatttacatttaaaattattgtttcagtatctttaatttattatgtatatataaaaataacttacaattttattaataaacaatatatgtttattaattcatgttttgtaatttatgggatagcgattttttttactgtctgtatttttcttttttaattatgttttaattgtattttatttttattattgttctttttatagtattattttaaaacaaaatgtattttctaagaacttataataataataaatataaattttaataaaaattatatttatcttttacaatatgaacataaagtacaacattaatatatagcttttaatatttttattcctaatcatgtaaatcttaaatttttctttttaaacatatgttaaatatttatttctcattatatataagaacatatttattaaatctagaattctatagtgagtcgtattacaattcactggccgtcgttttacaacgtcgtgactgggaaaaccctggcgttacccaacttaatcgccttgcagcacatccccctttcgccagctggcgtaatagcgaagaggcccgcaccgatcgcccttcccaacagttgcgcagcctgaatggcgaatggcgcctgatgcggtattttctccttacgcatctgtgcggtatttcacaccgcatatggtgcactctcagtacaatctgctctgatgccgcatagttaagccagccccgacacccgccaacacccgctgacgcgccctgacgggcttgtctgctcccggcatccgcttacagacaagctgtgaccgtctccgggagctgcatgtgtcagaggttttcaccgtcatcaccgaaacgcgcgagacgaaagggcctcgtgatacgcctatttttataggttaatgtcatgataataatggtttcttagacgtcaggtggcacttttcggggaaatgtgcgcggaacccctatttgtttatttttctaaatacattcaaatatgtatccgctcatgagacaataaccctgataaatgcttcaataatattgaaaaaggaagagtatgagtattcaacatttccgtgtcgcccttattcccttttttgcggcattttgccttcctgtttttgctcacccagaaacgctggtgaaagtaaaagatgctgaagatcagttgggtgcacgagtgggttacatcgaactggatctcaacagcggtaagatccttgagagttttcgccccgaagaacgttttccaatgatgagcacttttaaagttctgctatgtggcgcggtattatcccgtattgacgccgggcaagagcaactcggtcgccgcatacactattctcagaatgacttggttgagtactcaccagtcacagaaaagcatcttacggatggcatgacagtaagagaattatgcagtgctgccataaccatgagtgataacactgcggccaacttacttctgacaacgatcggaggaccgaaggagctaaccgcttttttgcacaacatgggggatcatgtaactcgccttgatcgttgggaaccggagctgaatgaagccataccaaacgacgagcgtgacaccacgatgcctgtagcaatgccaacaacgttgcgcaaactattaactggcgaactacttactctagcttcccggcaacaattaatagactggatggaggcggataaagttgcaggaccacttctgcgctcggcccttccggctggctggtttattgctgataaatctggagccggtgagcgtgggtctcgcggtatcattgcagcactggggccagatggtaagccctcccgtatcgtagttatctacacgacggggagtcaggcaactatggatgaacgaaatagacagatcgctgagataggtgcctcactgattaagcattggtaactgtcagaccaagtttactcatatatactttagattgatttaaaacttcatttttaatttaaaaggatctaggtgaagatcctttttgataatctcatgaccaaaatcccttaacgtgagttttcgttccactgagcgtcagaccccgtagaaaagatcaaaggatcttcttgagatcctttttttctgcgcgtaatctgctgcttgcaaacaaaaaaaccaccgctaccagcggtggtttgtttgccggatcaagagctaccaactctttttccgaaggtaactggcttcagcagagcgcagataccaaatactgtccttctagtgtagccgtagttaggccaccacttcaagaactctgtagcaccgcctacatacctcgctctgctaatcctgttaccagtggctgctgccagtggcgataagtcgtgtcttaccgggttggactcaagacgatagttaccggataaggcgcagcggtcgggctgaacggggggttcgtgcacacagcccagcttggagcgaacgacctacaccgaactgagatacctacagcgtgagctatgagaaagcgccacgcttcccgaagggagaaaggcggacaggtatccggtaagcggcagggtcggaacaggagagcgcacgagggagcttccagggggaaacgcctggtatctttatagtcctgtcgggtttcgccacctctgacttgagcgtcgatttttgtgatgctcgtcaggggggcggagcctatcgaaaaacgccagcaacgcggcctttttacggttcctggccttttgctggccttttgctcacatgttctttcctgcgttatcccctgattctgtggataaccgtattaccgcctttgagtgagctgataccgctcgccgcagccgaacgaccgagcgcagcgagtcagtgagcgaggaagcggaagagcgcccaatacgcaaaccgcctctccccgcgcgttggccgattcattaatgcagctggcacgacaggtttcccgactggaaagcgggcagtgagcgcaacgcaattaatgtgagttagctcactcattaggcaccccaggctttacactttatgcttccggctcgtatgttgtgtggaattgtgagcggataacaatttcacacaggaaacagctatgaccatgattacgccaagctatttaggtgacactatagaagcggccgcTAAGCTTCAGAAGATATTTATAAAAAAGGGGAAAAAAATATATATAACAGCGAATATGAAAATGATAATGATAATAATAAAGATTCTTGTCATTTTAATGAGGAGGAGAATGAAGAATTAGAAGAAGATGGAGATTTTTTAAATATCGATCATTTAAAAAGTAGTACTATACAAGAAAAGAATTTAGAAATTTTAAAAAAAAGAAAAGCATGTTCAAACAATATGAATATTAATAGAAAGAATATATTTACGAATATTAATATAACTTGTGGGCGTGATTTTTTTTGTTACATGCATGGGAATAATTATTCTAATAACATACCTATAAATTCATATTTATATATATCTTGGAATATAGGAAATGTGATATATAGAAATTTATATAATTCAAGTGAAAATATAATAATAAATGCTCATAGTTTGTTAAGTAGTCATAATATTAATGATGATGTAATATTAAAATTGAATATAGATGAAAATAATAGAAGGCTTTATCCATCAAATCTAATACATAGTTTATTAATGAATACAGAAAGTAATATATATATAAATAATGAATATGTTATATTAGATATCATTTTAAGTATATTCCATATTATATCAACTAATTTATCACCTATTATTGAAAAAATATTAGATGCAGAATATGGTACTAGGTGGATTATGGATAGCAAAATACCAAAAAGTAATATATGGGAAAAATCAAAAGGAAAAAATGAATTAGACATTGAAGGAATTATACATATCATTACAACCTTCTGGATAGATATATTTGAAAAGCGTATTAAAAATATAGAAATTATTCAAAATTTACAAAAAGCATCGATTTTTTGGTCAAATCAAGAAATAGATCAATTTGATCAAGACTTTGTCAAAAAATTAATCGAAAGCTCTTCCTTACTTTTAAAGCTCTTCGGAGATTATAACACAGCAAAGAGTATTGAAAAGTTGTATTACAATAAATATTCCATTTTTAATGACTTTTGCGTGGACTAAgattatatatatgtatatatgtatatatttatatatgtgtatatatatttcacatttttttttttttttttttttttttaatgggttatccataaatacatgctcatggggatatttaacaaaattttcttatgatacatatttggatatttaaccatttatttttatgttgatctataaaaaaagcattcatattgtgatatataatttattctgttaaaaaatttatgtcattatttattatttctttatttattttatttttttttttttttttgtttaaatttttaaaagaatatattccattttttgccattattacatgaattttttttttttttattccttttatatttattcatttttttttattccttttatatttatttattttttttttttttgttattccgcatctcatttgtgatgtactatattatccgtttttttttttttttttttttttttttttttttataatatatcctttttttttgttt

**5’ amplicon size: 1214bps**

**3’ amplicon size: 1195bps**
